# Long Noncoding RNA RROL Provides Chromatin Scaffold for MYC-WDR82 Interaction to Impact Lipid Metabolism and Tumor Cell Growth in Multiple Myeloma

**DOI:** 10.1101/2021.12.08.471297

**Authors:** Eugenio Morelli, Mariateresa Fulciniti, Mehmet K. Samur, Caroline F. Ribeiro, Leon Wert-Lamas, Jon E. Henninger, Annamaria Gullà, Anil Aktas-Samur, Katia Todoerti, Srikanth Talluri, Woojun D. Park, Cinzia Federico, Francesca Scionti, Nicola Amodio, Giada Bianchi, Megan Johnstone, Na Liu, Doriana Gramegna, Nicola A. Russo, Charles Lin, Yu-Tzu Tai, Antonino Neri, Dharminder Chauhan, Teru Hideshima, Masood A. Shammas, Pierfrancesco Tassone, Sergei Gryaznov, Richard A. Young, Kenneth C. Anderson, Carl D. Novina, Massimo Loda, Nikhil C. Munshi

## Abstract

Long noncoding RNAs (lncRNA) can drive the tumorigenesis and be susceptible to therapeutic intervention. To define the landscape of therapeutically actionable lncRNA dependencies in multiple myeloma (MM), we coupled our extensive lncRNA transcriptomic profile with lncRNA targeted CRISPR *interference* viability screen and identified RNA Regulator of Lipogenesis (RROL) as a leading lncRNA dependency in MM. RROL shares its origin with the microRNA locus MIR17HG, however supports the proliferation and survival of MM cells in a microRNA- and DROSHA- independent manner. We found that RROL provides a chromatin scaffold for the functional interaction between c-MYC and WDR82 to promote the regulation of the lipogenic pathways via the transcriptional control of the rate-limiting enzyme ACC1 in MM cells. Inhibition of RROL with clinically applicable antisense molecules disrupts its transcriptional and functional activities causing potent anti-tumor effects both *in vitro* and *in vivo* in two pre-clinical animal models. This study establishes lncRNA RROL as a therapeutically actionable dependency with a unique mechanism of action in support of myeloma cell growth.

## INTRODUCTION

In the human genome, gene loci harboring lncRNAs outnumber protein-coding genes and are susceptible to the same oncogenic pathogenetic events (Hon et al., 2017; Wang et al., 2018). These RNA molecules are defined simply by having a length greater than 200nt and a lack of protein-coding potential, therefore representing a diverse array of functional entities (Ulitsky and Bartel, 2013). LncRNAs are often classified in four separate subgroups based on their location relative to protein-coding genes: *exonic*, *intronic*, *overlapping* and *intergenic*; alternatively, they can be functionally classified into cis- and trans-acting lncRNAs (Ulitsky and Bartel, 2013). Trans-acting lncRNAs are of special interest for their diverse mechanisms of action, such as their role as precursor molecules for the biogenesis of mature microRNAs (miRNAs) (Lu et al., 2017) or through their direct interactions with proteins and nucleic acids to regulate protein function and/or stability (Tseng et al., 2014). With the plethora of biological functions that lncRNAs modulate to control cellular processes at multiple levels, it is not surprising that their aberrant expression and function has been implicated in the progressive gain of a malignant phenotype by tumor cells (Gutschner and Diederichs, 2012).

Multiple myeloma (MM) is a genetically complex malignancy of plasma cells that accounts for about the 10% of hematologic cancers (Gulla and Anderson, 2020). Despite recent advancements, MM remains largely incurable (Gulla and Anderson, 2020). A growing body of evidence points to a key role played by the noncoding RNA (ncRNA) networks in this disease context (Morelli et al., 2020), indicating that MM cells can become significantly addicted and therapeutically susceptible to the modulation of oncogenic ncRNAs (Amodio et al., 2018; Hu et al., 2018; Leone et al., 2013; Morelli et al., 2018; Morelli et al., 2015; Pichiorri et al., 2016). We have consistently observed a profound alteration of the *intergenic* lncRNA (hereafter just lncRNA) landscape in newly diagnosed MM patients and reported their role as an independent risk predictor for the clinical outcome (Samur et al., 2018); laying the foundations to functionally and therapeutically explore the dependency to lncRNAs in MM.

Here, through a large-scale CRISPR *interference* (CRISPRi) viability screen (Liu et al., 2017), we identify RNA Regulator of Lipogenesis (RROL) as a leading lncRNA dependency in MM; and describe its oncogenic function and therapeutic targeting in MM.

## RESULTS

### A genome-wide CRISPRi viability screen identifies MIR17HG as a leading dependency in MM

We analyzed RNA-seq data from 360 newly diagnosed MM patients and identified 913 lncRNA transcripts expressed in primary MM cells (**Fig. 1A, I.**) and in a panel of 70 MM cell lines (data not shown). To systematically interrogate the role of these lncRNAs in MM cell growth, we transduced 3 MM cell lines (H929, KMS-11 and KMS-12-BM) engineered to express a dCAS9-KRAB fusion protein with a pooled library consisting of 7 sgRNAs against each of the 913 transcription start sites (TSS) and 576 negative control sgRNAs (**Fig. 1A, II.** and **Supplementary table 1**). Relative representation of sgRNAs was assessed by deep sequencing after 3 weeks and analyzed using the Model-based Analysis of Genome-wide CRISPR-Cas9 Knockout (MAGeCK) robust rank aggregation (RRA) algorithm (Li et al., 2014a). The most enriched or depleted sgRNAs were further tested in secondary screens using a pooled library targeting 224 lncRNA TSS, the TSS of known protein coding oncogenes (MYC, IRF4) (Chesi et al., 2008; Shaffer et al., 2008) or tumor suppressors (TP53) (Jovanovic et al., 2018) as positive controls, and 2245 non-targeting sgRNAs as negative control (**Fig. 1A; III.** and **Supplementary table 2)**. In the secondary screens 4 MM cell lines (H929, KMS11, KMS12BM and AMO1) were used to detect and rank significantly depleted or enriched sgRNAs. As expected, sgRNAs targeting IRF4 and MYC were significantly depleted in three (MYC) or all (IRF4) cell lines, while sgRNAs targeting TP53 were significantly enriched in both TP53 wild-type cell lines (AMO1 and H929) (Tessoulin et al., 2018). Focusing on depleted sgRNAs, we identified lncRNA dependencies in MM cells that were either cell type specific (54%) or shared by two or more cell lines (46%) (**Supplementary Fig. 1A).** A ranked analysis of sgRNA depletion identified MIR17HG as the leading lncRNA dependency in the screen, with RRA scores equal or superior to those obtained by targeting MYC or IRF4 in all cell lines tested (**Fig.1B**). To validate this data further, we next transduced MM cell lines expressing dCAS9-KRAB fusion protein with the top four sgRNAs targeting MIR17HG under the regulation of a tetracycline-inducible promoter and observed reduced cell growth compared to cells infected with non-targeting sgRNAs after continued’ exposure to doxycycline (**Fig. 1C** and **Supplementary Fig. 1B**). Moreover, we used 2 different locked nucleic acid (LNA) gapmeR ASOs (simply referred as ASO), targeting the MIR17HG nascent RNA (pre-RNA) for RNase H-mediated degradation (Lai et al., 2020; Lee and Mendell, 2020), to transfect 11 MM cell lines including those resistant to conventional anti-MM agents (AMO1-ABZB resistant to bortezomib; AMO1-ACFZ resistant to carfilzomib; MM.1R resistant to dexamethasone); and confirmed significant impact on MM cell viability independent of the genetic and molecular background (**Fig. 1D** and **Supplementary Fig. 1C**).

**Figure 1.**
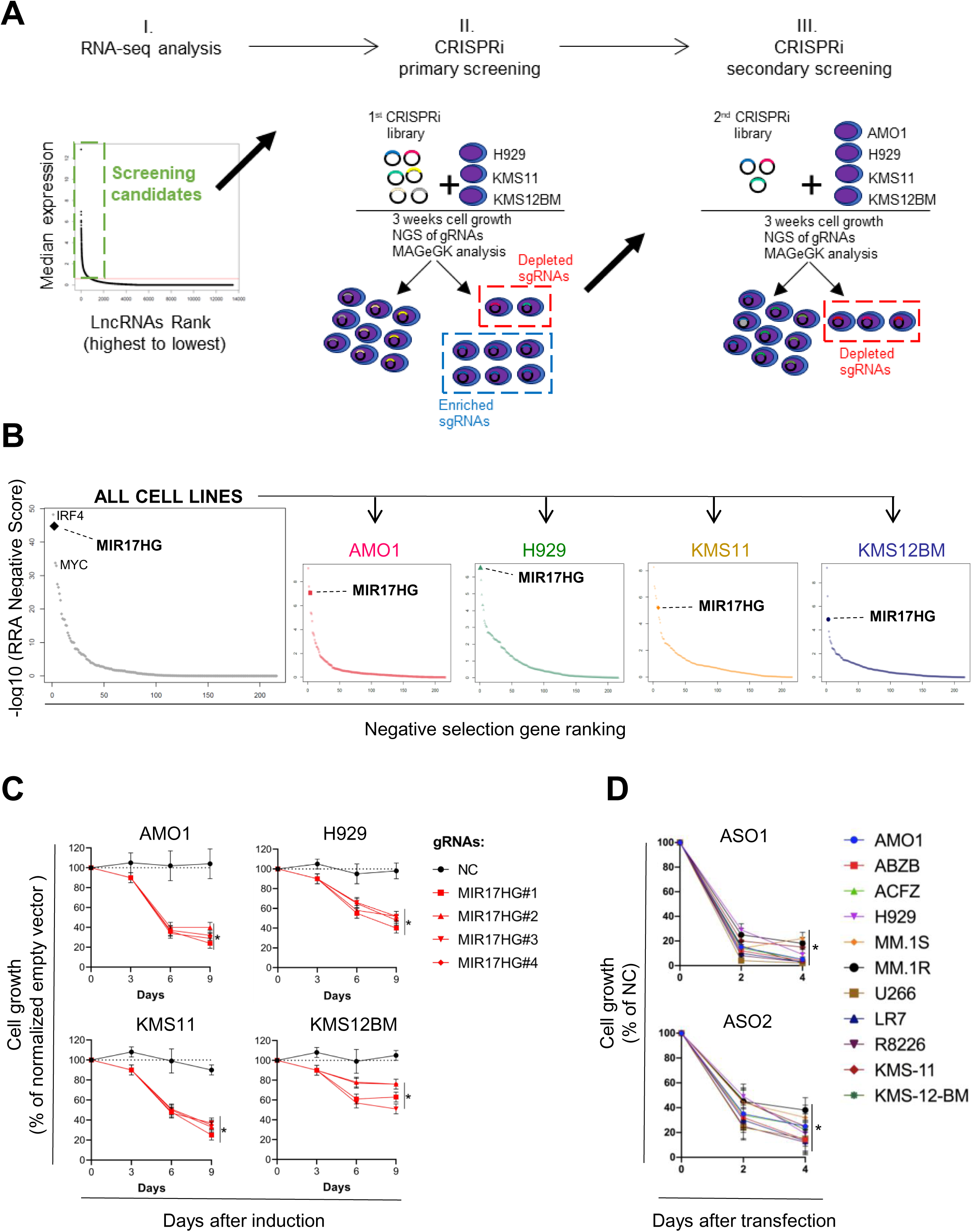
A genome-wide CRISPRi viability screen identifies MIR17HG as a leading dependency in MM. **A**) Schematic of CRISPRi viability screens. **B**) RRA-based ranked analysis of lncRNA dependencies in the secondary screen, considering 4 MM cell lines either together or individually. The top lncRNA dependency MIR17HG is highlighted, along with protein coding genes IRF4 and MYC used as positive controls. **C**) CCK-8 proliferation assay of MM cell lines (AMO1, H929, KMS11 and KMS12BM) stably expressing KRAB-dCAS9 fusion protein and transduced with lentivectors to conditionally express anti-MIR17HG sgRNAs. CCK-8 assay was performed at indicated time points after exposure to doxycycline (0.5μ g/mL). Cell proliferation is calculated compared to parental cells infected with the empty sgRNA vector and exposed to doxycycline under some conditions. **D**) CCK-8 proliferation assay of MM cell lines (n=11) transfected with 2 different ASOs targeting the MIR17HG pre-RNA or a non-targeting ASO (NC). ASOs were used at concentration of 25nM. Cell viability was measured 2 and 4 days after electroporation, and it is represented as % of viability compared to cells transfected with NC-ASO. Data from 1 out of 3 independent experiments is shown in panel D and E. Data present mean ± s.d.in D and E. \**p*<0.05 by Student’s *t* test.

These data establish a broad dependency to lncRNAs in MM cells and provide rationale to further explore the molecular and functional roles of MIR17HG in the MM setting.

### MIR17HG^RROL^ (or RROL) mediates dependency in microRNA-independent manner

Beside providing precursor for the microRNA cluster miR-17-92 (MIR17HG^miR-17-92^: miR-17/-18a/-19a/-20a/-19b/-92a1), MIR17HG also produces as yet poorly characterized lncRNA transcript (He et al., 2005; Ota et al., 2004) (**Fig. 2A**); named here RNA Regulator of Lipogenesis (RROL or MIR17HG^RROL^) based on the functional description provided in this study. We observed that RROL expression was higher during disease progression in 2 independent datasets from MM patients analyzed at diagnosis and/or relapse (**Supplementary Fig. 2A-B**); and that higher expression of RROL was associated with shorter event-free (EFS) and overall (OS) survival in 3 large cohorts of newly diagnosed MM patients (**Fig. 2B**). Expression of RROL did not significantly correlate with miR-17-92s in CD138+ MM cells from 140 patients (average Spearman *r* = 0.16), suggesting that RROL and miR-17-92 are subject to an independent regulatory control and may function in distinct molecular pathways (**Supplementary Fig. 2C**).

**Figure 2.**
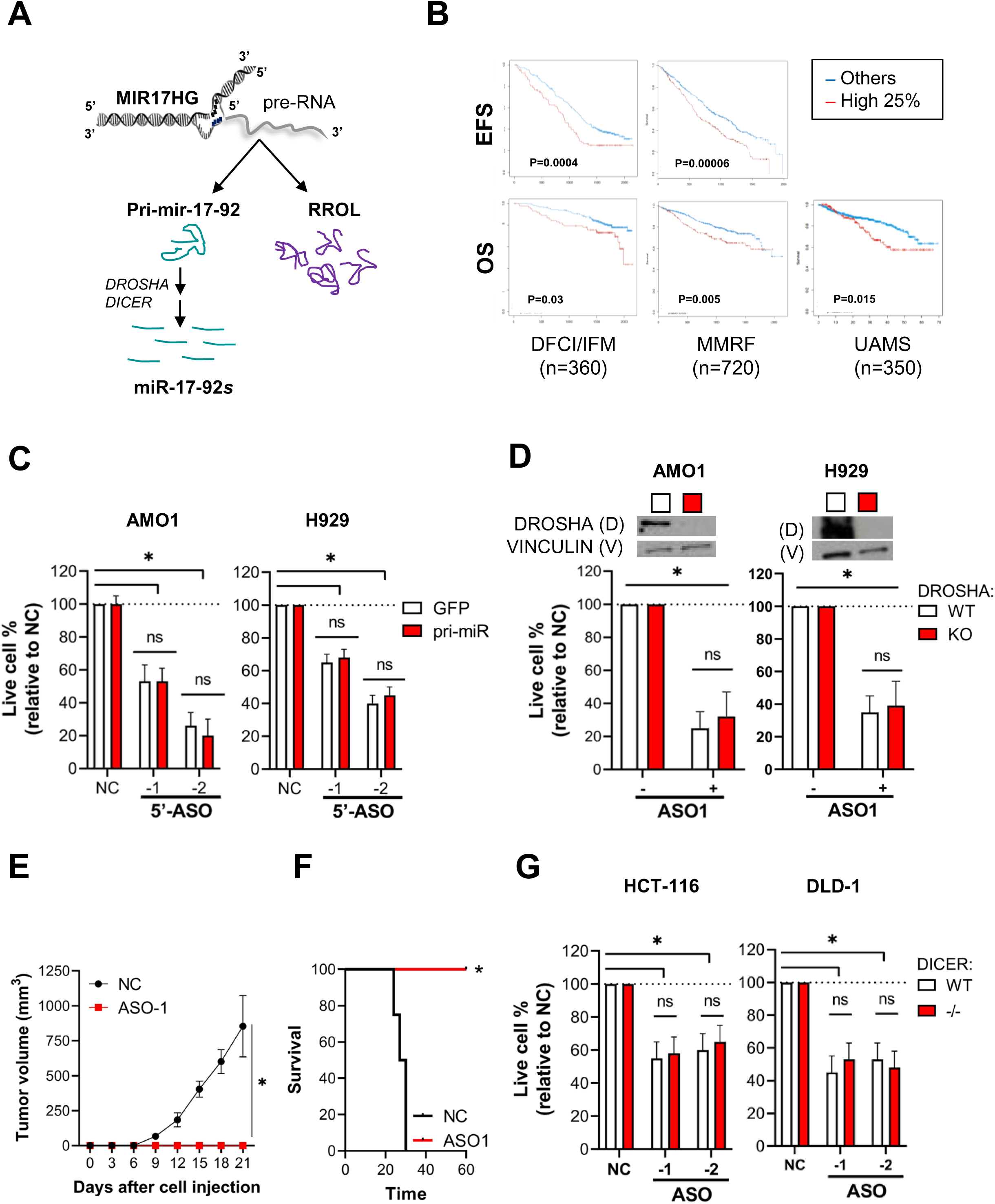
MIR17HG^RROL^ (or RROL) mediates dependency in microRNA-independent manner. **A**) Overview of MIR17HG locus, including both lncRNA (RROL) and miRNA (miR-17-92) derived transcripts. **B**) Prognostic significance (PFS and OS) of high RROL expression (top quartile) in 3 large cohorts of MM patients. **C**) CCK-8 proliferation assay in AMO1 and H929 cells stably transduced with either a lentivector carrying pri-mir-17-92 (pri-miR) or a lentiviral vector carrying GFP as control; and transfected with 2 different ASOs targeting the 5’end (5’-ASO) of MIR17HG pre-RNA or a scrambled control (NC). Effects on cell proliferation were assessed 48h after transfection. **D**) CCK-8 proliferation assay of DROSHA WT or KO AMO1 and H929 exposed to ASO1 (1µM for AMO1 and 2,5µM for H929) for 6 days. It is shown a western blot analysis of DROSHA expression in WT and KO cells. Vinculin was used as protein loading control **E**) Effects of RROL depletion in a matrigel-based AMO1^DR-KO^ xenograft in NOD SCID mice. It is shown tumor growth of AMO1^DR-KO^ with (ASO-1) or without (NC) RROL depletion. **F**) Survival analysis of tumor injected mice. **G**) CCK-8 proliferation assay in HCT-116 and DLD-1 colorectal cancer cell lines expressing either a WT or mutated (-/-) Dicer; and transfected with either a scramble control (NC) or 2 different ASO (-1 and -2) to obtain RROL depletion. Live cells %, compared to NC, was analyzed 48h after transfection. *indicates p<0.05, *ns* indicates p>0.05 after Student t test.

Supporting a miRNA-independent function of RROL, first, we observed an intact anti-proliferative activity of anti-MIR17HG ASOs in presence or absence of ectopic expression of pri-mir-17-92 in two MM cell lines (**Fig. 2C** and **Supplementary Fig. 2D**). To solely deplete RROL, ASOs were designed to target MIR17HG at the 5’-end (5’-ASOs), a sequence not included in the ectopic pri-mir-17-92. Second, we established two DROSHA knockout (DR-KO) MM cell lines (AMO1^DR-KO^ and H929^DR-KO^) which are unable to produce miR-17-92s (**Supplementary Fig. 2E**) (Bartel, 2004). ASO-mediated targeting of MIR17HG pre-RNA solely affects RROL in these cells. We observed a strong anti-proliferative activity in both DR-WT and DR-KO cell systems after RROL depletion, with no significant rescue in DR-KO cells, as assessed after gymnotic treatment with ASO1 (**Fig. 2D)** or after transfection with 3 different ASOs (-1/-2/-3) (**Supplementary Fig. 2F**). Importantly, exposure to ASO1 abrogated the ability of AMO1^DR-KO^ to establish tumors into NOD SCID mice (**Fig. 2E**) and prolonged animal survival (**Fig. 2F**). We also observed anti-proliferative activity of anti-MIR17HG ASOs in colorectal cancer cell lines HCT-116 and DLD-1, despite these cell lines carrying (-/-) or not (wt) a mutant Dicer conferring hypomorphic phenotype (Cummins et al., 2006) (**Fig. 2G**).

These results indicate that RROL is the main mediator of MIR17HG cancer dependency without relationship to miR-17-92.

### RROL interacts with chromatin to regulate gene expression

The functional role of lncRNAs depends on their subcellular localization (Ulitsky and Bartel, 2013). qRT-PCR analysis of nuclear and cytosolic compartments, with MALAT1 and GAPDH mRNA as positive controls, indicated a nuclear enrichment of RROL (**Supplementary Fig. 3A**). This finding was confirmed by RNA FISH (**Supplementary Fig. 3B**). On this basis, we next explored the transcriptional network regulated by RROL in MM. To this end, we performed an ASO-based loss-of function (LOF) study followed by an integrated gene expression analysis in both DR-WT (AMO1 and H929) and DR-KO (AMO1^DR-KO^) MM cell lines. Through a kinetic analysis of differentially expressed genes (DEGs), performed after early exposure to gymnotic ASO1 to avoid modulation of miR-17-92 in DROSHA WT cells (**Supplementary Fig. 3C**), we identified a set of genes rapidly downregulated after RROL depletion in all the cell lines tested (**Fig. 3A**). We validated these findings in CD138+ cells from 3 MM patients treated *ex-vivo* with ASO1 (**Fig. 3B**); as well as in other cellular models, including lymphoma cell lines Raji and Daudi (**Supplementary Fig. 3D**) and murine MM cell line 5TGM1 depleted of human or murine RROL (**Supplementary Fig. 3E**). Conversely, the expression of these genes was not affected by modulation of individual members of miR-17-92 by synthetic mimics or inhibitors (**Supplementary Fig. 3F-G**). Moreover, we observed significant positive correlation (Spearman *r* > 0.3; p<0.001) between RROL and its target genes in at least 1 out of 2 large RNA-seq MM patient datasets (IFM/DFCI and MMRF/CoMMpass) (**Fig. 3C**); supporting clinical relevance of these regulatory axis.

**Figure 3.**
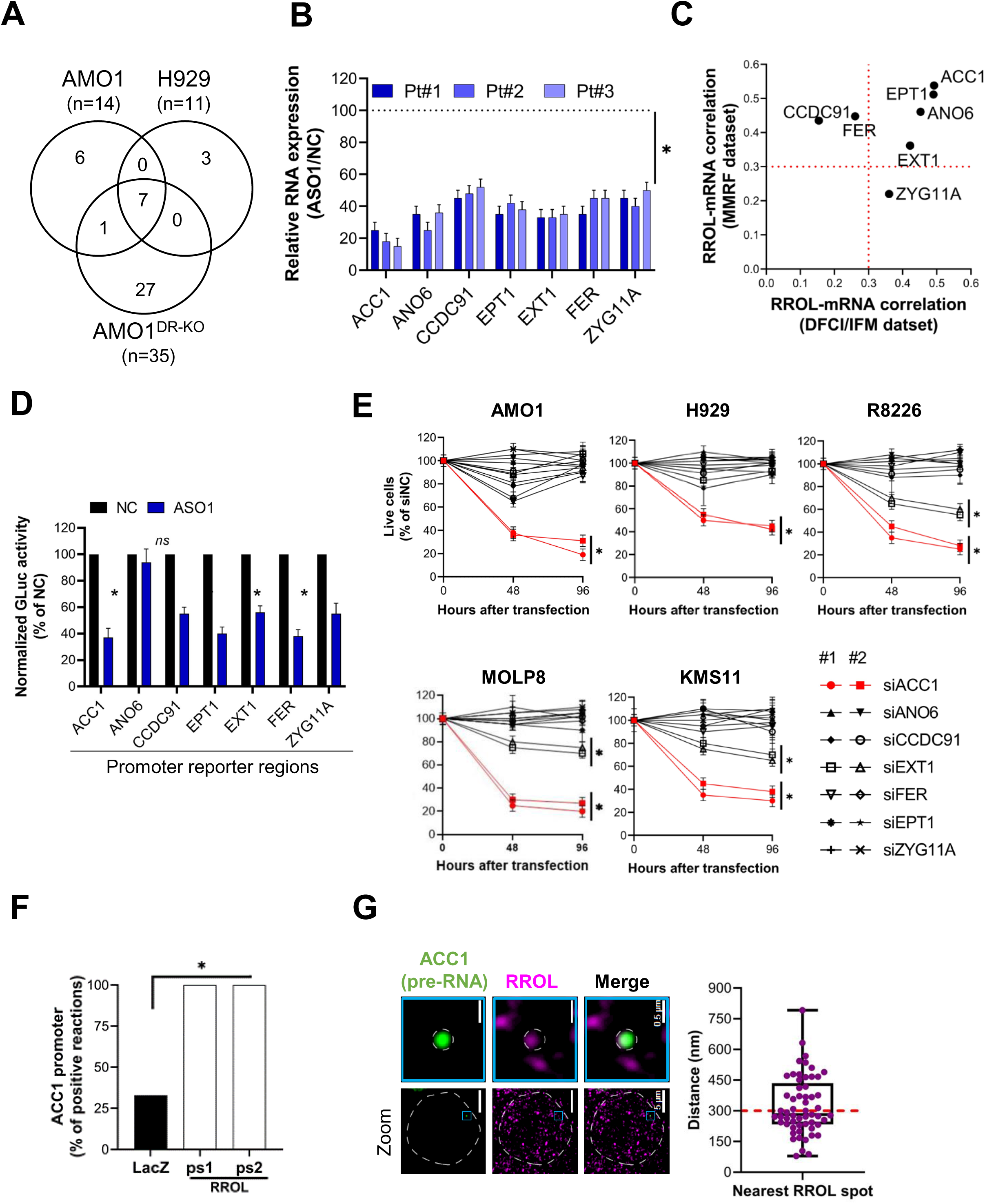
RROL interacts with chromatin to regulate gene expression. **A**) Transcriptomic analysis after RROL depletion in MM cell lines, either DROSHA WT (AMO1, H929) or KO (AMO1^DR-KO^). It is shown a venn diagram of commonly downregulated genes (adj p<0.05; lfc<-1). Cells were exposed to ASO1 for 24h. **B**) qRT-PCR analysis of RROL targets in CD138+ cells from 3 MM patients exposed to ASO1 for 24h. The results shown are average mRNA expression levels after normalization with GAPDH and ΔΔCt calculations. RNA level in cells exposed to NC (vehicle) were set as an internal reference. **C**) Correlation analysis between RROL targets (mRNA) and RROL in CD138+ MM patient cells from 2 large RNA-seq cohorts (DFCI/IFM, n=360; MMRF/CoMMpass, n=720). It is reported the Spearman *r* obtained in DFCI/IFM (*x* axis) and MMRF/CoMMpass (*y* axis) dataset. Dotted red lines indicates *r*=0.3. **D**) GLuc/ SEAP dual reporter assay showing reduced activity of ACC1, ANO6, CCDC91, EPT1, EXT1, FER and KIAA1109 promoter activity after RROL knockdown using ASO1. The reporter vectors were co-transfected into 293T cells with either ASO1 or control ASO. Cells were harvested for luciferase activity assay 48h after transfection. Results are shown as % of normalized Gluc activity in ASO1 transfected cells compared to control. **E**) CCK-8 proliferation assay in 5 MM cells lines after transfection with siRNAs against RROL targets. 2 siRNAs were used for each target, plus a scramble siRNA (NC) as control. Cell viability was measured at the indicated time points and it is represented as % of NC transfected cells. **F**) ChIRP-qPCR analysis showing effective amplification of ACC1 promoter in chromatin purified using 2 RROL antisense probe sets (ps1 and ps2), compared to chromatin purified using LacZ antisense probes (negative control). *indicates p<0.05 after Fisher Exact Test. **G**) (left) Snapshot obtained by dual RNA-FISH analysis of ACC1 pre-mRNA (green) and RROL (purple) in a representative AMO1 cell; (right) box plot showing the distance (nm) of ACC1 pre-RNA spots to the nearest RROL spots (n=57) or to the nearest random spots (160). 300nm is used as cut-off determining proximity. *indicates p<0.1 after Fisher Exact Test. In all panels, except **F** and **G**, *indicates p<0.05 after Student t test.

Using a luciferase reporter assay, performed in 293T^DR-KO^ cells in presence or absence of RROL depletion, we demonstrated that the regulatory control of RROL over these genes, except ANO6, occurs at the promoter level (**Fig. 3D**). RNAi-based LOF screen in 5 MM cell lines identified ACC1 as the RROL target gene with most significant impact on MM cell proliferation and survival (**Fig. 3E**); providing the rationale to further explore the role of RROL-ACC1 axis in MM pathobiology. Consistently, we confirmed RROL interaction at the promoter region of the top target ACC1 by chromatin isolation by RNA precipitation (ChIRP) assay followed by qRT-PCR analysis (**Fig. 3F** and **Supplementary Fig. 3H-I**). Moreover, we showed frequent proximal localization of RROL at the ACC1 gene locus by single molecule dual RNA FISH analysis of RROL and ACC1 pre-mRNA (<300nm to nearest RROL spot in ∼50% of ACC1 pre-RNA spots analyzed (n=60) (**Fig. 3G**).

Altogether, these data indicate RROL as a chromatin-interacting lncRNA with transcriptional regulatory functions.

### RROL promotes MYC occupancy at the ACC1 promoter

Through an upstream regulator analysis of RROL-related gene expression changes, we observed a strong inhibition of the MYC related network upon RROL depletion in both cell lines tested (**Fig. 4A**). Based on this analysis we evaluated if MYC and RROL cooperate to promote ACC1 expression in MM cells. RROL depletion in MM cells indeed abrogated MYC occupancy at the ACC1 promoter, while not affecting its expression (**Fig. 4B** and **Supplementary Fig. 4A**); and reduced the expression of ACC1 in the conditional MYC Tet-Off cell line P493-6 (Schuhmacher et al., 1999) only in presence of high MYC levels (**Fig. 4C**). Moreover, by coupling RNA FISH analysis of RROL and ACC1 pre-RNA with immunofluorescence analysis of MYC protein (FISH/IF), we captured the co-localization of RROL and MYC at the ACC1 gene locus (**Fig. 4D**).

**Figure 4.**
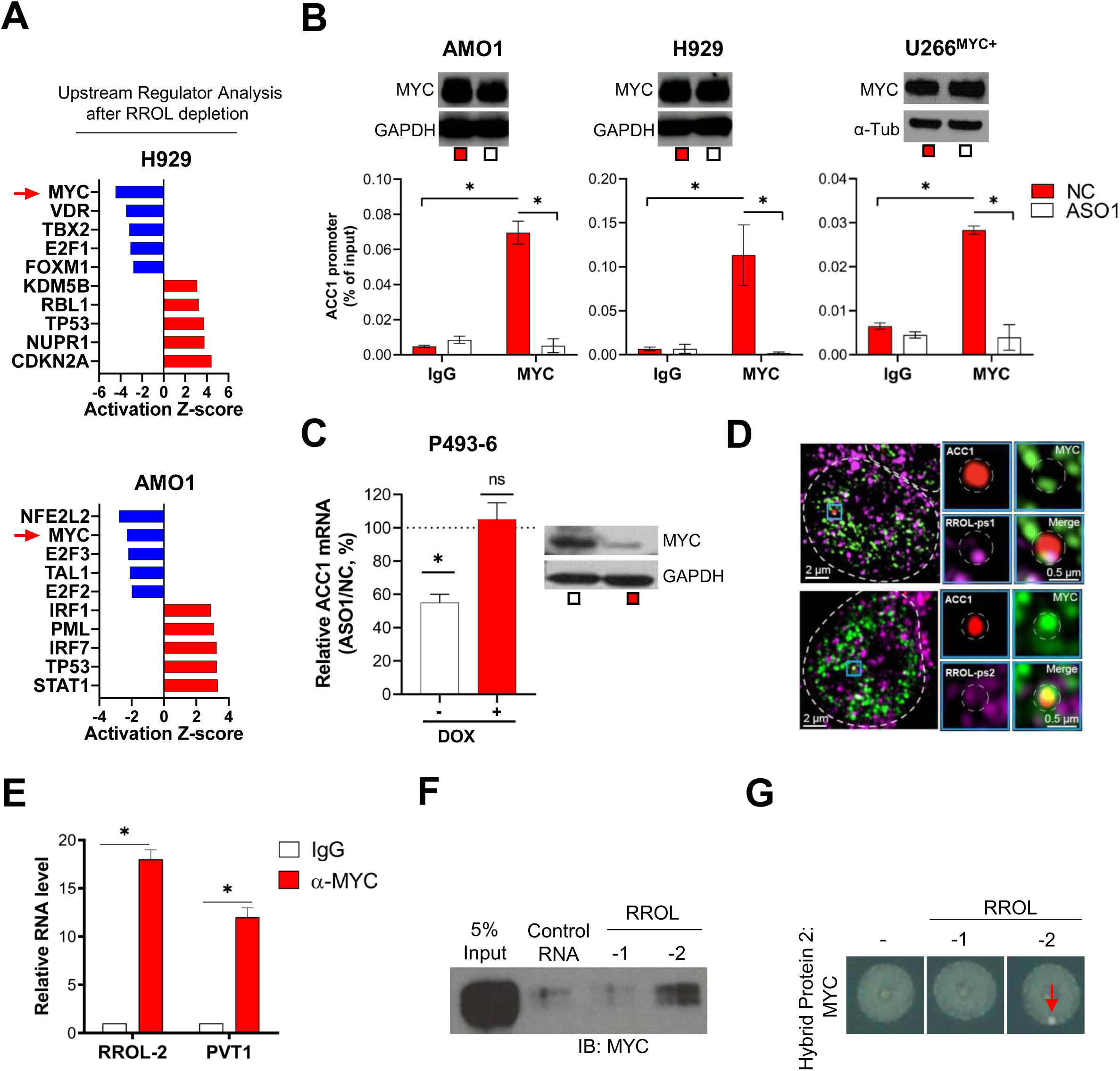
RROL promotes MYC occupancy at the ACC1 promoter. **A**) Upstream regulator analysis of transcriptional changes after 3 days of gymnotic exposure to ASO1 in H929 and AMO1 cells. MYC is highlighted by a red arrow. **B**) ChIP-qPCR analysis of MYC occupancy at ACC1 promoter in AMO1, H929 and U266^MYC+^ exposed for 24h to ASO1 or NC (vehicle). MYC occupancy at ACC1 promoter is calculated as % of input chromatin. Below each histogram plot, it is shown western blot analysis of MYC from paired samples. GAPDH or α-tubulin were used as protein loading controls. **C**) qRT-PCR analysis of ACC1 mRNA in P493-6 cells exposed for 2 days to either doxycycline or DMSO to knockdown MYC; and then, for additional 2 days exposed to either ASO1 or vehicle (NC) to deplete RROL. ACC1 expression levels in cells exposed to NC were set as an internal reference. **D**) Co-IF/FISH analysis of ACC1 pre-RNA (red), MYC protein (green) and RROL (purple). RROL was detected using 2 different probes (RROL-ps1 and RROL-ps2). A representative image is shown with each RROL probe. **E**) qRT-PCR analysis of RROL (detecting isoform 2) in RIP material precipitated using an anti-MYC antibody (α-MYC) or IgG control. LncRNA PVT1 is used as positive control for the known role as MYC interactor. **F**) Western blot analysis of MYC in RPPD material precipitated with control RNA or RROL transcripts -1 and -2. 5% input is used as reference. **G**) RNA Y3H using MYC as hybrid protein 2 and, as hybrid RNAs, either a negative control RNA (-) or RROL transcripts -1 and -2. *p < 0.05, Student t test.

To evaluate the existence of a RROL-MYC complex, we then performed RNA immunoprecipitation (RIP) assay with MYC antibody which showed a specific enrichment of RROL isoform 2 (RROL-2) in the MYC-bound RNA (**Fig. 4E** and **Supplementary Fig. 4B-C**). RNA-Protein pull-down (RPPD) experiments confirmed MYC forming a complex with RROL-2 (**Fig. 4F** and **Supplementary Fig. 4D**). Furthermore, we adapted the RNA yeast-3-hybrid (Y3H) assay to confirm the RROL-MYC interaction in an *in vivo* cellular model (Hook et al., 2005). In this assay, a direct RROL-MYC interaction activates a reporter gene allowing for yeast colony growth; and, as shown in **Fig. 4G**, we detected yeast colony growth in the presence of RROL-2 as hybrid RNA.

These data demonstrate that RROL forms an RNA-protein complex with the transcription factor MYC to promote its chromatin occupancy and transcriptional activity at the ACC1 promoter.

### RROL mediates the assembly of a MYC-WDR82 transcriptional complex, leading to transcriptional and epigenetic activation of ACC1

MYC activity is modulated through the interaction with transcriptional and epigenetic co-regulators (Gouw et al., 2019). To determine if RROL affects these protein-protein interactions, we integrated the results of proximity-dependent biotin identification (BioID) analysis with co-immunoprecipitation assay followed by mass-spectrometry analysis (Co-IP/MS) in 3 MM cell lines (AMO1, H929 and U266^MYC+^), in the presence and in the absence of RROL depletion. This integrated analysis highlighted WDR82 as a very high-confidence RROL-dependent MYC interactor (**Fig. 5A** and **Supplementary Fig. 5A; Supplementary tables 3-6**); A direct RNA-protein interaction between RROL and WDR82 was further confirmed by both RPPD (**Fig. 5B**) and RNA Y3H (**Fig. 5C**) assays. WDR82 is a regulatory component of the SET1 methyltransferase complex catalyzing the histone H3 ‘Lys-4’ trimethylation (H3K4me3) at the transcription start sites of active loci (Lee and Skalnik, 2008), a *sine qua non* condition for MYC binding to chromatin and transactivation (Amente et al., 2011). Consistently, depletion of WDR82 resulted in a global reduction of H3K4me3 (**Fig. 5D**) and a reduced occupancy of MYC at ACC1 promoter (**Fig. 5E**), resulting in decreased ACC1 expression (**Fig. 5F**) in MM cells. Furthermore, using MM cells expressing an ectopic WDR82-GFP fusion protein (**Supplementary Fig. 5B**), we demonstrated that RROL expression is essential for WDR82 occupancy at the ACC1 promoter (**Fig. 5H**). Additionally, RROL depletion resulted in reduced levels of H3K4me3 at the ACC1 promoter site, without detectable impact on the total H3K4me3 (**Fig. 5I**).

**Figure 5.**
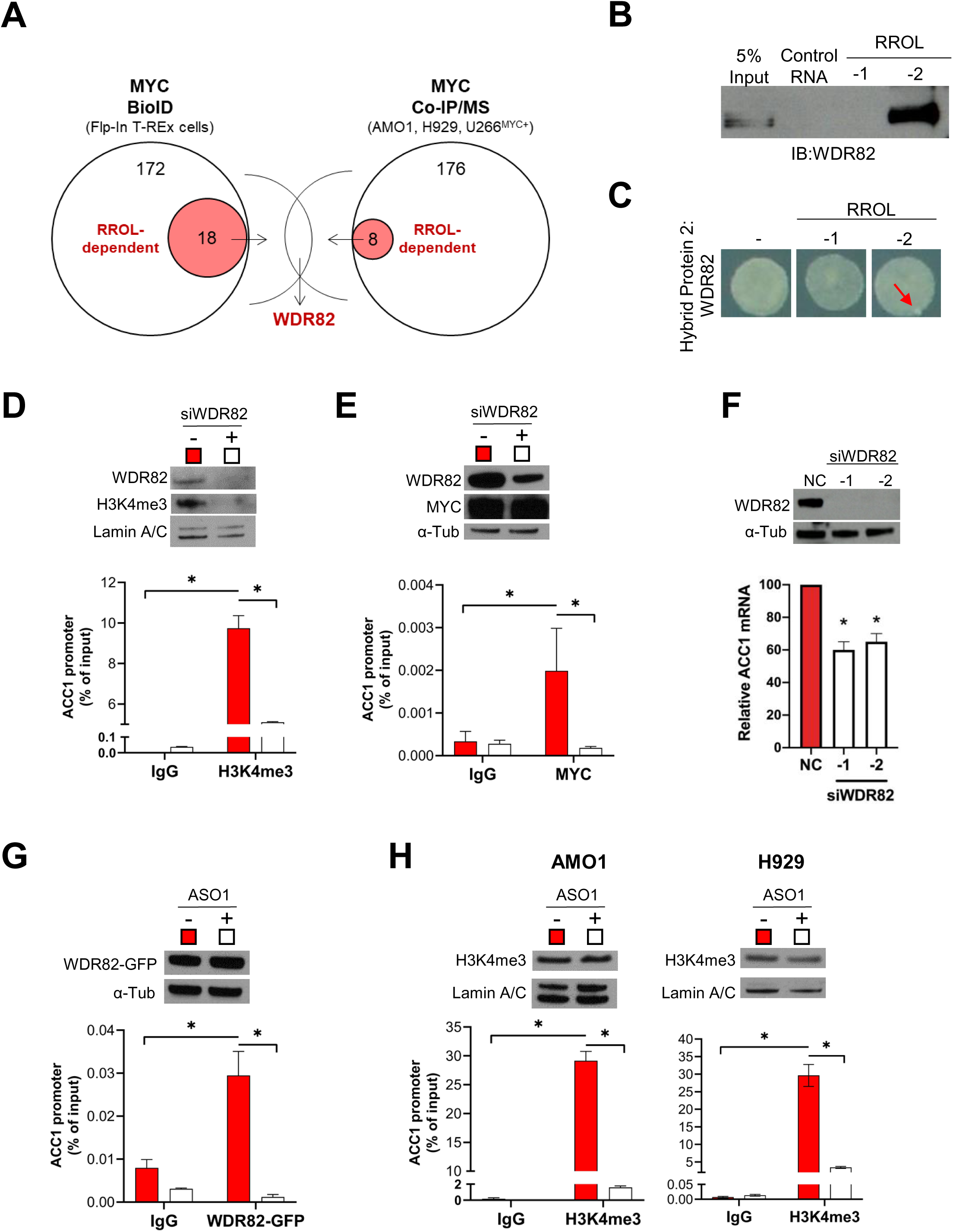
RROL mediates the assembly of a MYC-WDR82 transcriptional complex, leading to transcriptional and epigenetic activation of ACC1. **A)** Schematic of integrated BioID and Co-IP/MS assays to explore the MYC-protein interacting network in presence or absence of RROL depletion. **B)** Western blot analysis of WDR82 in RPPD material precipitated with RROL -1 and RROL-2 or with control RNA. 5% input is used as reference. **C)** RNA Y3H using WDR82 as hybrid protein 2 and, as hybrid RNAs, either a negative control RNA (-) or RROL transcripts -1 and -2. Red arrows indicate yeast colony growth. **D)** ChIP-qPCR analysis of H3K4me3 occupancy at ACC1 promoter in AMO1 after silencing of WDR82 (24h time point). Data are represented as % of input chromatin. It is also shown western blot analysis of WDR82 and H3K4me3. Lamin A/C was used as protein loading controls (nuclear lysates). **E)** ChIP-qPCR analysis of MYC occupancy at ACC1 promoter in AMO1 after silencing of WDR82 (24h time point). Data are represented as % of input chromatin. It is also shown a western blot analysis of WDR82 and MYC from paired samples. α-tubulin was used as protein loading controls. **F)** qRT-PCR analysis of ACC1 mRNA after transfection of siRNA targeting WDR82 (siWDR82-1 or -2) in AMO1. Raw Ct values were normalized to GAPDH mRNA and expressed as ΔΔCt values calculated using the comparative cross threshold method. ACC1 expression levels in cells transfected with NC were set as an internal reference. **G)** ChIP-qPCR analysis of WDR82-GFP occupancy at ACC1 promoter in AMO1 exposed for 24h to gymnotic ASO1. Data are represented as % of input chromatin. It is also shown western blot analysis of WDR82-GFP from paired samples. α-tubulin was used as protein loading controls. **H)** ChIP-qPCR analysis of H3K4me3 occupancy at ACC1 promoter in AMO1 and H929 exposed for 24h to gymnotic ASO1. Data are represented as % of input chromatin. It is also shown western blot analysis of H3K4me3 from paired samples. Lamin A/C was used as protein loading controls (nuclear lysates). *p < 0.05, Student t test.

These findings suggest the role of RROL as a chromatin scaffold mediating the assembly of MYC-WDR82 multiprotein transcriptional complexes, to control the expression of ACC1.

### The RROL/MYC-ACC1 axis regulates *de novo* lipogenesis

ACC1 catalyzes the carboxylation of acetyl-CoA into malonyl-CoA, the rate limiting step during *de novo* lipogenesis (DNL) (Beloribi-Djefaflia et al., 2016), a metabolic pathway aberrantly activated in cancer cells (Rohrig and Schulze, 2016). Here we observed that RROL depletion, similarly to either MYC or ACC1 inhibition, significantly reduced the incorporation of C^14^-radiolabeled glucose into the lipid pool – indicative of a reduced DNL (Zadra et al., 2019) – both in MM cell lines and CD138+ MM patient cells (**Fig. 6A)**. This inhibition was however not observed after transfection of MM cells with synthetic inhibitors of miR-17-92s (**Supplementary Fig. 6A)**. Liquid Chromatography - Mass Spectrometry (LC-MS) based lipid profiling after RROL inhibition in MM cells confirmed depletion of several saturated (SFA) and monounsaturated (MUFA) phospholipid species (**Fig. 6B**) – produced *via* DNL(Rysman et al., 2010; Zaidi et al., 2013). Moreover, addition of the main downstream product of ACC1 activity – palmitate (PA) – significantly rescued the anti-proliferative (**Fig. 6C**) and pro-apoptotic effects of RROL depletion in MM cells (**Fig. 6D**), confirming role of DNL in the tumor promoting activity of RROL in MM and justifying the name of RNA Regulator of Lipogenesis (**Fig. 6E**).

**Figure 6.**
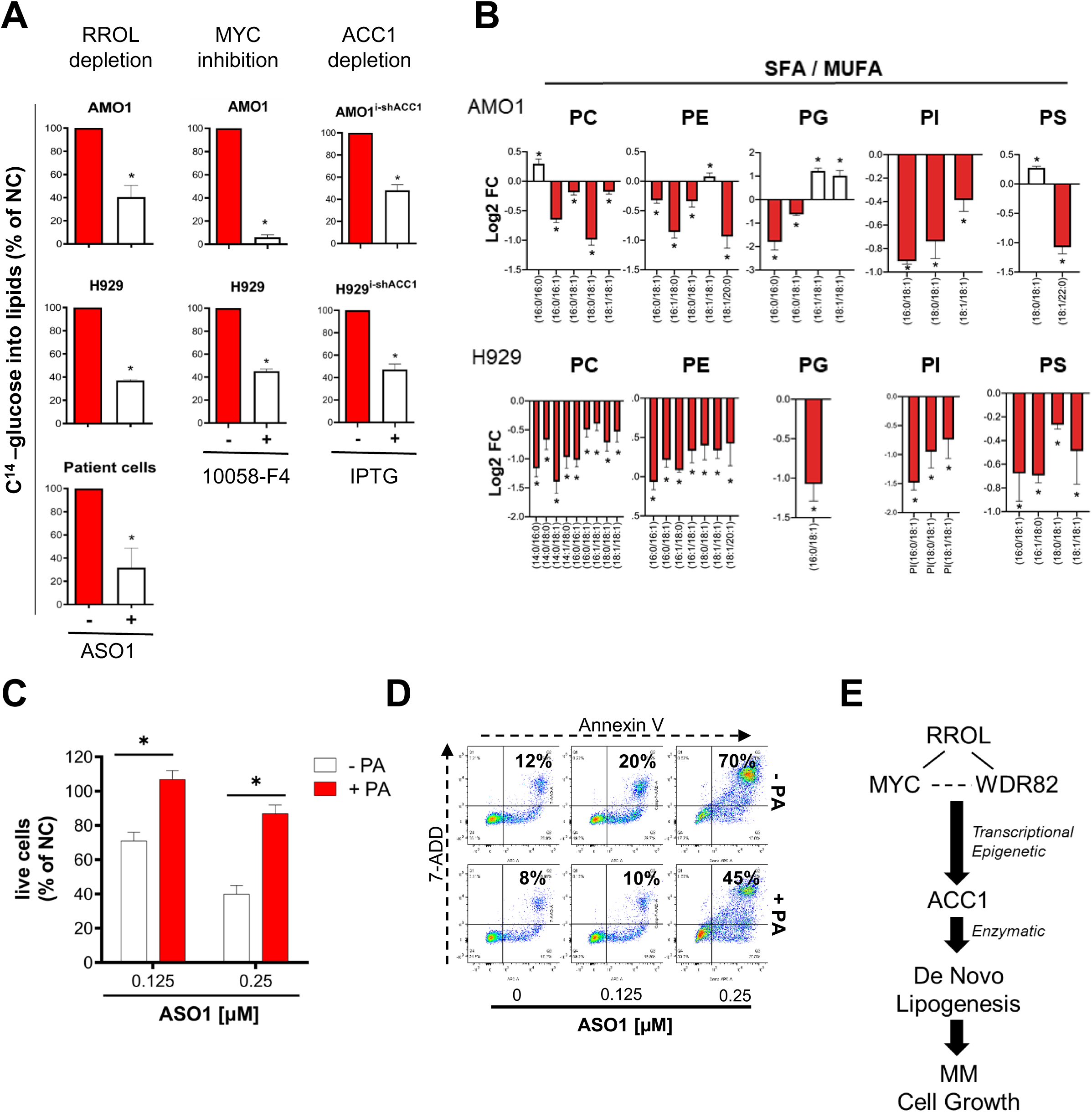
The RROL/MYC-ACC1 axis regulates *de novo* lipogenesis. **A**) Incorporation of ^14^C-glucose into lipids following either RROL or MYC or ACC1 depletion / inhibition in MM cell lines or CD138+ MM patient cells. RROL was depleted using ASO1; MYC was inhibited using small molecule 10058-F4; ACC1 was depleted using IPTG-inducble shRNAs. Results are expressed as percentage of negative controls (NC ASO, DMSO and uninduced, respectively). **B**) Lipid profiling analysis showing modulation of major membrane phospholipid classes [phosphatidylglycerol (PG), phosphatidylethanolamine (PE), phosphatidylserine (PS), phosphatidylcholin (PC) and phosphatidylinositol (PI)], with saturated (SFA) or monounsaturated (MUFA) acyl chains, after treatment of AMO1 and H929 with ASO1. **C**) CCK-8 proliferation assay and **D**) Annexin V / 7-AAD flow cytometry assay in AMO1 exposed for 6 days to ASO1 at indicated concentrations, with supplementation of either BSA (control) or BSA-PA (20μM). **E**) Proposed model of RROL lipogenic pathway. * p<0.05, Student t test.

### Therapeutic inhibitors of RROL exert potent anti-tumor activity *in vitro* and *in vivo* in animal models of human MM

To explore the therapeutic potential of RROL as a target and develop inhibitors, we screened >80 fully phosphorothioated (PS), 2’-O-methoxyethyl (2’-MOE)-modified, lipid-conjugated ASOs that could either trigger RNase H-mediated degradation of RROL (gapmeRs) or exert function *via* an RNase H-independent mechanism (steric blockers) (Puttaraju et al., 2021) (**Fig. 7A** and **Supplementary Fig. 7A**). This procedure identified an 18-mer tocopherol (T)-conjugated gapmeR G2-15b-T (“G”) and an 18-mer tocopherol (T)-conjugated steric blocker SB9-19-T (“SB”) as leading compounds, both producing a strong anti-proliferative activity (cell growth inhibition, CGI > 50%) in a large panel of MM cell lines as well as CD138+ primary MM cells; while sparing (CGI < 50%) non-malignant cell lines (THLE-2, HK-2, HS-5 and 293T) and PBMCs from 3 healthy donors (**Fig. 7B**).

**Figure 7.**
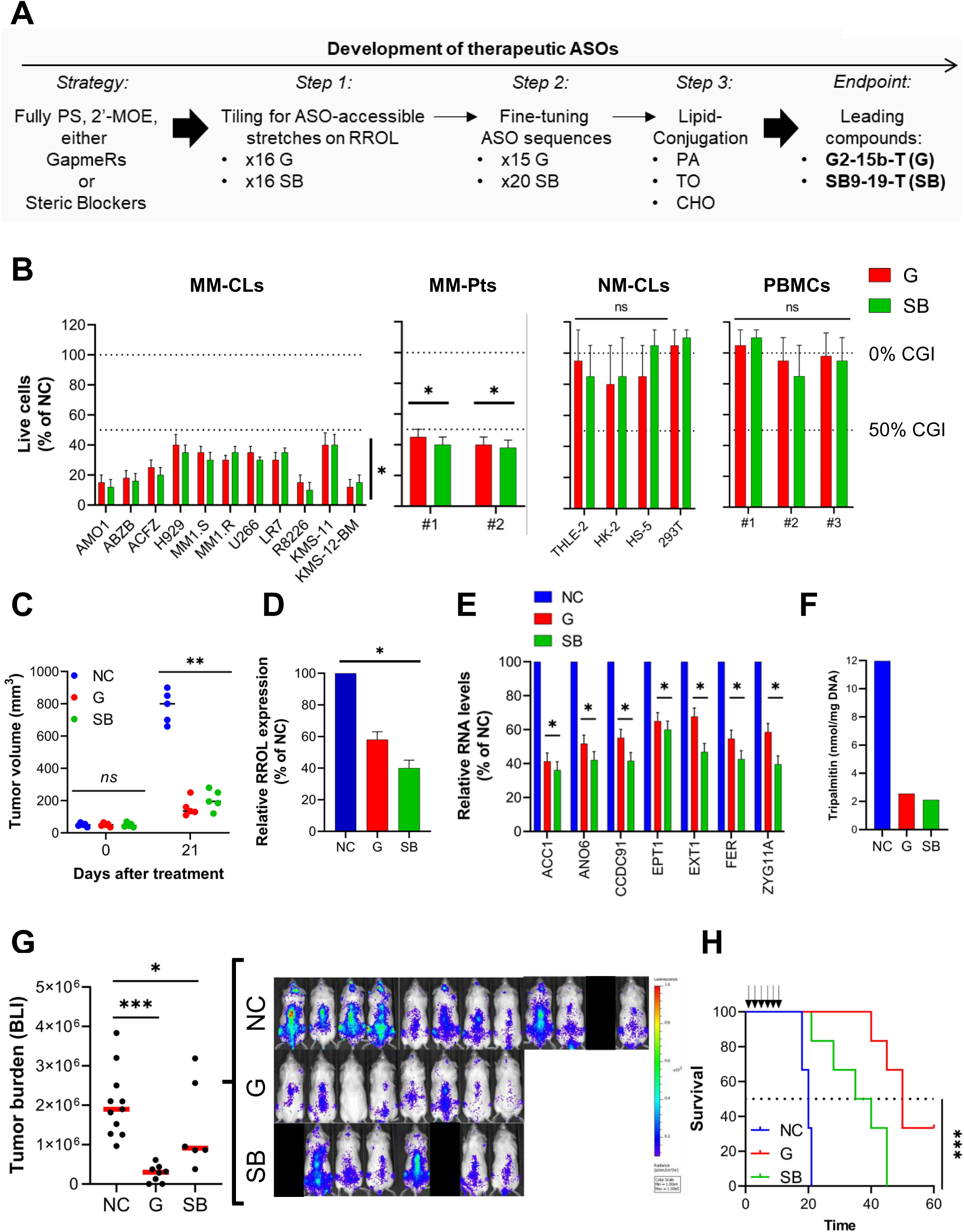
Therapeutic inhibitors of RROL exert potent anti-tumor activity *in vitro* and *in vivo* in animal models of human MM. **A**) Schematic of the multi-step screen to develop RROL therapeutic ASOs. **B**) CCK-8 proliferation assay in a panel of 11 MM cell lines (MM-CLs: AMO1, ABZB, ACFZ, H929, MM.1S, MM1.R, U266, LR7, R8226, KMS-11, KMS-12-BM), CD138+ cells from 2 MM patients (MM-Pt), 4 non-malignant cell lines (THLE-2, HK-2, HS-5 and 293T) and PMBCs from 3 healthy donors. Cell viability was measured 48h (or 24h for MM-pt) after transfection with either G2-15b*-TO (G) or SB9-19-TO (SB) or vehicle (-) as control. Cell proliferation is calculated compared to cells transfected with vehicle. **C**) Subcutaneous *in vivo* tumor growth of AMO1 cells in NOD SCID mice, 21 days after treatment with G2-15b*-TO (G; n=5) or SB9-19-TO (SB; n=5) or vehicle (NC; n=5). **D-E**) qRT-PCR analysis of RROL (D) and RROL targets (E) in AMO1 xenografts, retrieved from animals treated with G2-15b*-TO (G; n=1) or SB9-19-TO (SB; n=1) or vehicle (NC; n=1) as control. Raw Ct values were normalized to ACTB mRNA and expressed as ΔΔCt values calculated using the comparative cross threshold method. Expression levels in NC were set as an internal reference. **F**) Lipid profiling analysis showing modulation of tripalmitin in tumors retrieved from animals treated with G2-15b*-TO (G; n=1) or SB9-19-TO (SB; n=1) or vehicle (NC; n=1) as control. **G**) BLI-based measurement of *in vivo* tumor growth of MOLP8-luc+ in NSG mice, after treatment with G2-15b*-TO (G; n=8) or SB9-19-TO (SB; n=6) or vehicle (NC; n=11). On the left, a scatter plot showing the analysis of bioluminescence intensity. Red bars indicate median value. Bioluminescence was measured at the end of treatment cycle (day 15). On the right, it is shown image acquisition. Mice removed from the study due to failed I.V. injection of tumor cells are covered by a black rectangle. **F**) Survival analysis from experiment in panel E. Black arrows indicate treatments. *means p<0.05; **means p<0.01; ***means p<0.001.

To assess the *in vivo* anti-tumor activity of both compounds, we first used a subcutaneous AMO1 xenograft model in immunocompromised NOD SCID mice. Here, we observed a significant reduction of tumor growth after a treatment cycle with either G2-15b-T (tumor growth inhibition, TGI=76%) or SB9-19-T (TGI=69%) (**Fig. 7C**). Analysis of tumors retrieved from mice following this treatment confirmed reduced expression of RROL (**Fig. 7D**) and its targets (**Fig. 7E**), as well as reduced levels of tripalmitin (**Fig. 7F**), a surrogate for the DNL product palmitate (Falchook et al., 2021); demonstrating an efficient uptake of G2-15b-T and SB9-19-T by tumor cells *in vivo*. Moreover, no overt toxicity was observed in the mice.

We next confirmed a significant anti-MM activity of G2-15b-T and SB9-19-T in an aggressive model of diffused myeloma, in which tumor growth of MOLP8-luc+ MM cells is assessed by bioluminescence imaging (BLI) measurement. In this model, tumor growth was significantly antagonized after a treatment cycle with either G2-15b-T (TGI=84%) or SB9-19-T (TGI=52%). Treatment with G2-15b-T resulted in a tumor clearance in 2 out of 8 mice (25%) (**Fig. 7G**). Importantly, both inhibitors significantly prolonged the animal survival (**Fig. 7H**).

## DISCUSSION

We have identified lncRNA RROL as a leading dependency in MM and explored its functional and therapeutic roles.

RROL host gene, MIR17HG, is often amplified and/or overexpressed in human cancer with driver role (He et al., 2005; Li et al., 2014b; Mogilyansky and Rigoutsos, 2013). Its impact in tumorigenesis seems particularly relevant when co-expressed with MYC (He et al., 2005; Li et al., 2014b), with a well-documented interplay between MIR17HG^miR-17-92^ and MYC transcriptional targets to maintain cancer cell homeostasis (Izreig et al., 2016; Li et al., 2014b; Morelli et al., 2018). Our description of microRNA-, DROSHA- and DICER-independent function of MIR17HG, *via* RROL, establishes this gene as having both *short* (miR-17-92) and *long* (RROL) noncoding RNA activities; with the latter mediating the tumor promoting activity in MM and likely other cancer contexts (e.g., colorectal cancer).

We described RROL as a regulator of gene expression *via* chromatin occupancy and interaction with transcription factors and epigenetic modulators, such as MYC and WDR82. Our data lend support to the emerging paradigm whereby chromatin occupancy by transcription factors like MYC may be determined through the interaction with specific lncRNAs (Zhang et al., 2018), in addition to protein partners as previously established (Kalkat et al., 2018). In a broader perspective, our observations on RROL suggest lncRNAs as key mediators of the epigenetic and transcriptional reprogramming of MM cells. In this molecular scenario, while proteins such as transcription factors and epigenetic modulators act as catalytic effectors, the intrinsic structural flexibility of lncRNAs make them modular scaffolds able to mediate both protein-protein and protein-DNA interactions at specific chromatin regions.

We further showed that the RROL-MYC-WDR82 complex impacts tumor cell metabolism by activating the DNL pathway *via* the rate-limiting enzyme ACC1. This anabolic pathway is primarily restricted to liver and adipose tissue in normal adults but is reactivated in cancer cells via mechanisms yet to be fully described (Beloribi-Djefaflia et al., 2016; Rohrig and Schulze, 2016). Noteworthy, MYC has been implicated in the reprogramming of tumor cell metabolism by activating DNL *via* ACC1 and other genes (Stine et al., 2015). In turn, DNL has emerged as an essential pathway for the onset and progression of MYC-driven cancers, that are susceptible to pharmacologic inhibition of ACC1 (Gouw et al., 2019). The roles of ACC1 and DNL in tumorigenesis seem particularly relevant in MM, where tumor cells need to adapt their metabolic pathways to meet the high bioenergetic and biosynthetic demand posed by the malignant cell growth coupled with unceasing production of monoclonal immunoglobulin (El Arfani et al., 2018; Masarwi et al., 2019). Nevertheless, we acknowledge that the oncogenic roles of RROL – along with MYC and WDR82 – are unlikely limited to the transcriptional axis with ACC1; and will require further investigations to be comprehensively elucidated.

Finally, for translational purposes, we have developed 2 therapeutic ASOs that target RROL via different mechanisms of action (i.e., RNase H-dependent or - independent). With the recent advances in RNA medicine (Crooke et al., 2018; Damase et al., 2021; Sullenger and Nair, 2016) the use of ASOs to therapeutically antagonize disease-driver genes is becoming increasing possible (Dhuri et al., 2020; Puttaraju et al., 2021), including in MM therapy (Mondala et al., 2021; Morelli et al., 2018). Our optimization of design and chemistry has helped to overcome the major obstacles to the clinical use of ASOs, such as poor bioavailability (Dhuri et al., 2020), while limiting off-target toxicity. The inhibitors described here bear state-of-the-art chemical modifications (2’MOE, PS-backbone, lipid conjugation) and have sufficient nucleotide length (18mer) to assure high specificity for RROL; inhibitors of this kind have already been tested within clinical trials and few are already FDA approved for the use in different human diseases (Dhuri et al., 2020). Our optimized ASOs targeting RROL with demonstrated activity in 2 different murine models of human MM provide therefore the rationale to now consider clinical application in MM.

In conclusion, we here establish RROL as a novel lncRNA that facilitates MYC-WDR82 protein complex formation and its chromatin binding impacting lipid metabolism and ultimately tumor cell growth in MM.

## ACKNOWLEDGEMENTS

This work is supported by NIH/NCI grants SPORE-P50CA100707, R01-CA050947, R01CA207237, P01CA155258, R01-CA178264 (N.C.M., K.C.A.); by VA Healthcare System grant No. 5I01BX001584 (N.C.M.); by the Paula and Roger Riney Foundation grant (N.C.M., K.C.A.) by NIH/NCI grants RO1CA131945, R01CA187918, P50 CA211024 and Department of Defense grants DoD PC160357 and DoD PC180582 (M.L.); by the Sheldon and Miriam Medical Research Foundation (K.C.A.); by the Italian Association for Cancer Research (AIRC) with “Special Program for Molecular Clinical Oncology–5 per mille”, 2010/15 and its Extension Program” No. 9980, 2016/18 (P.T.). E.M. is supported by the Brian D. Novis Junior Grant from the International Myeloma Foundation (IMF) and by the Dana Farber/Harvard Cancer Center SPORE in Multiple Myeloma Career Enhancement Award (SPORE-P50CA100707). A.G. is a Fellow of The Leukemia & Lymphoma Society (LLS) and a Scholar of the American Society of Hematology (ASH). JEH is supported by a NIH F32 fellowship (NCI 1F32CA254216-01). A.N. is supported by a grant from the Italian Association for Cancer Research (AIRC, IG24365). K.C.A. is an American Cancer Society Clinical Research Professor. We gratefully acknowledge the members of our laboratories for technical advice and critical discussions. We thank Dr. Dirk Eick (Helmholtz-Zentrum München, Molecular Epigenetics), Dr. Christoph Driessen (Cantonal Hospital St Gallen, St Gallen, Switzerland) and Dr. Linda Penn (Princess Margaret Cancer Centre, University Health Network, Toronto, Ontario, Canada) for providing us relevant cellular models. We thank Dr. Pierosandro Tagliaferri (Magna Graecia University, Catanzaro, Italy) for insightful discussions on the dual long and short nature of MIR17HG. We thank Dr. Benjamin Izar and Dr. Johannes Melms (Columbia University, New York, NY, USA.) for insightful discussions on CRISPR viability screen methodology. We thank the “8^th^ Annual Miracles for Myeloma 5K Virtual Run/Walk” organizers and attendees for the fundraising in support to this research project.

## AUTHOR CONTRIBUTIONS

E.M. and N.C.M. conceived and designed the research studies. E.M., M.F. and N.C.M. wrote the manuscript. M.K.S., A.A.S. and K.T. performed in silico analysis of transcriptomic data. C.F.R. performed lipidomic studies. L.W.L. performed yeast-3-hybrid experiments. J.E.H. performed RNA FISH, dual RNA FISH and Co-IF/dual RNA FISH. S.T. provided MM cells expressing CAS9. W.D.P. analyzed ChIP-seq data. S.G. designed t-ASOs. C.F., N.R. and F.S. provided support for the identification of RROL isoforms. M.F., A.G., N.A., M.J., G.B., C.L., Y.T.T., A.N., D.C., T.H., M.A.S., P.T., R.A.Y., K.C.A., C.D.N. and M.L. contributed to the design and interpretation of key experiments. M.L. supervised lipidomic studies. C.D.N. supervised Y3H experiments.

## STAR METHODS

### Cell lines

Cell lines (CLs) were grown at 37°C, 5% CO_2_. a) <u>MM-CLs:</u> AMO1, NCI-H929, SK-MM-1, U266, JJN3 and KMS-12-BM were purchased from DSMZ (Braunschweig, Germany). MM.1S, MM.1R and RPMI-8226 were purchased from ATCC (Manassas, VA, USA). AMO1 bortezomib-resistant (ABZB) and AMO1 carfilzomib-resistant (ACFZ) were kindly provided by Dr. Christoph Driessen (Eberhand Karls University, Tübingen, Germany)(Morelli et al., 2018). U266 melphalan-resistant (LR7) were kindly provided by Dr. Atanasio Pandiella (Universidad de Salamanca, Salamanca, Spain)(Morelli et al., 2018). KMS-11 were kindly provided by Dr. K.C. Anderson (Dana-Farber Cancer Institute, Harvard Medical School, Boston, MA, USA). These cells were cultured in RPMI-1640 medium (Gibco® Life Technologies, Carlsbad, CA, USA) supplemented with 10% fetal bovine serum (Lonza Group Ltd., Basel, Switzerland) and 1% penicillin/streptomycin (Gibco®, Life Technologies). b) B-cell lymphoma cell lines (BCL0CL): Maver-1, Jeko-1 (mantle cell lymphoma), Sultan, P3HR1, Daudi and Raji (Burkitt lymphoma) (purchased from ATCC) were cultured in RPMI-1640 medium (Gibco® Life Technologies) supplemented with 10% fetal bovine serum (Lonza Group Ltd.) and 1% penicillin/streptomycin (Gibco®, Life Technologies). c) <u>Non-malignant cell lines:</u> HK-2 (human kidney cells, cortex/proximal tubule) were purchased from ATCC and cultured in K-SFM (Keratinocyte Serum Free Medium) (Thermo Fisher Scientific, Waltham, MA, USA). supplemented in accordance with ATCC guide lines; THLE-2 (human liver cells) were purchased from ATCC and cultured in BEGM (Bronchial epithelial cell growth medium) (Lonza Group Ltd.) supplemented in accordance with ATCC guide lines; d) <u>Lenti-X™ 293T</u> (human embryonic kidney, purchased from Takara (cat. no. 632180)) and Flp-In T-REx cells (kind gift of Dr. Linda Penn, University of Toronto) were cultured in DMEM (Dulbecco’s modified Eagle’s medium) (Gibco®, Life Technologies) supplemented with 10% fetal bovine serum (Lonza Group Ltd.) and 1% penicillin/streptomycin (Gibco®, Life Technologies). e) <u>P493-6</u> were kindly provided by Dr. Dirk Eick (Max Planck Institute of Biochemistry, Helmholtz-Zentrum München, Germany) and cultured in RPMI-1640 medium (Gibco® Life Technologies) supplemented with 10% fetal bovine serum (Lonza Group Ltd.) and 1% penicillin/streptomycin (Gibco®, Life Technologies). f) <u>5TGM1 murine MM cells</u> were kindly provided by Dr. Irene Ghobrial (Dana-Farber Cancer Institute, Boston, MA) and cultured in IMDM (Iscove modified Dulbecco medium) (Gibco®, Life Technologies) supplemented with 10% fetal bovine serum (Lonza Group Ltd.) and 1% penicillin/streptomycin (Gibco®, Life Technologies). g) <u>Colorectal cancer cell lines</u> HCT116 and DLD-1, parental and DICER mutants, were purchased from Horizon Discovery and cultured in ATCC-formulated McCoy’s 5a Medium Modified (Catalog No. 30-2007) supplemented with 10% fetal bovine serum (Lonza Group Ltd.) and 1% penicillin/streptomycin (Gibco®, Life Technologies). Cells were periodically tested to exclude mycoplasma contamination. Cells were STR (short tandem repeats) authenticated.

### Primary patient cells

Following informed consent approved by our Institutional Review Board of the Dana-Farber Cancer Institute, CD138+ cells were isolated from the BM aspirates of MM patients by Ficoll-Hypaque (Lonza Group, Basel, Switzerland) density gradient sedimentation; followed by antibody-mediated positive selection using anti-CD138 magnetic activated cell separation microbeads (Miltenyi Biotech, Gladbach, Germany). Purity of immunoselected cells was assessed by flow-cytometry analysis using a phycoerythrin-conjugated CD138 monoclonal antibody by standard procedures. For long-term culture (6 days), CD138+ cells were cultured physically separated from HS-5 cells by means of Falcon Cell Culture Inserts (Corning, New York, NY, USA), according to manufacturer’s instructions, as previously described(Morelli et al., 2018).

### Peripheral blood mononuclear cells

Peripheral blood mononuclear cells (PBMCs) were isolated from healthy adult donors, after informed consent approved by our Institutional Review Board of the Dana-Farber Cancer Institute. Cells were separated using Ficoll-hypaque method (Lonza Group Ltd.). PBMCs were cultured in RPMI-1640 medium (Gibco®, Life Technologies) supplemented with 10% fetal bovine serum (Lonza Group Ltd.) and 1% penicillin/streptomycin (Gibco®, Life Technologies).

### RNA-seq, microarray-based gene expression analysis and microRNA profiling of MM patients

<u>RNA-seq:</u> as primary dataset, we used previously published RNAseq from CD138+ MM cells from 360 MM patients from IFM/DFCI 20019 clinical trial (NCT01191060) (Samur et al., 2018). We used this dataset to assess expression of lncRNAs in newly diagnosed MM patients. Unstranded paired-end RNA sequencing were quantified using quasi-mapping with Salmon. Reference transcripts for GRCh38 transcripts were downloadedfrom Gencode v24. After QC controls TPM values for genes generated from isoform level TPMs with tximport. All figures were created with R and ggpubr. De-novo assembly for the RNAseq data on IFM cohort was done using TopHat. Gencode v24 GTF file were used as the reference and new isoform annotated by TopHat were identified from the output files. We also used newly diagnosis (ND) and relapse (R) samples from the continuation of the DFCI/IFM study to compare ND-MM and R-MM. Similar to diagnosis only samples these samples were sequenced with pared end sequencing and expression was quantified using the same pipeline explained above. As a secondary dataset, TPM level filtered MMRF CoMMpass data were downloaded from MMRF Research portal. Only samples those were collected from CD138+ selected BM samples at diagnosis were used for analysis.

<u>Microarray-based gene expression analysis:</u> RROL expression level was evaluated in a publicly available dataset (GSE66293)(Lionetti et al., 2015) including 129 newly diagnosed and 12 relapsed MM cases that were profiled by GeneChip Human Gene 1.0 ST array (Affymetrix, Santa Clara, CA, USA)(Todoerti et al., 2013). Normalized and re-annotated expression levels were obtained as described(Todoerti et al., 2013), using Chip Definition Files from BrainArray libraries version 20.0.0(Dai et al., 2005). Differential expression between the two groups was assessed by Wilcoxon runk sum test with continuity correction in R environment (version 4.0.4).

<u>miRNA profiling:</u> miRNA expression data for IFM cohort were generated using Affymetrix GeneChip® miRNA Array 4.0 platform. We used Affy and oligo packages from Bioconductor to normalize the miRNA expression data.

<u>Correlation analysis:</u> Spearman correlation was used to evaluate correlation between lncRNA, mRNAs and miRNAs.

<u>Survival analysis:</u> survival analysis was performed using survival package in R, and log rank test was used to compare groups.

### Generation of dCAS9-KRAB cell lines

Cell lines expressing the dCas9-KRAB fusion protein were generated as previously described (Morelli et al., 2021). Briefly, cells were infected with a lentivirus expressing the dCas9-BFP-KRAB transgene (Addgene, Plasmid #46911) and sorted for clones stably expressing high BFP. Infection was performed at low MOI (<0.4). Validation of transcriptional repression in MM cell lines expressing the dCas9-KRAB fusion protein was assessed by infecting lentivirus expressing a sgRNA for ENO1 (gRNA_ENO1: CCGGCGAGATCTCCGTGCTC) or a non-targeting negative control (gRNA_NC: GATGTGGTCATTCGTCATGA). sgRNAs were cloned into pU6-sgRNA EF1Alpha-puro-T2A-BFP (Plasmid #60955). This procedure followed protocols established by Weissman Lab and available online https://weissmanlab.ucsf.edu/CRISPR/CRISPR.html. Downregulation of ENO1 was assessed by qRT-PCR analysis following procedure described below [Reverse transcription (RT) and quantitative real-time amplification (qRT-PCR)].

### CRISPRi viability screens

<u>Library design:</u> gRNAs to target lncRNA TSSs were used designed using the Broad Institute web portal (now called CRISPick: https://portals.broadinstitute.org/gppx/crispick/public). For primary screen, target lncRNAs were selected based on median TPM>0.5 in the IFM/DFCI cohort. For secondary screen, target lncRNAs were selected based on primary screen results (i.e. targeted by significantly depleted or enriched gRNAs, FDR<0.25); plus additional lncRNAs identified through a de-novo assembly of RNA-seq data and manually selected lncRNAs selected based on their impact on the clinical outcome of MM patients enrolled in the IFM/DFCI clinical study.

<u>gRNA pool library production:</u> Primary CRISPRi library consisting of 7,500 gRNAs or secondary CRISPRi library consisting of 3,750 gRNAs were co-transfected with packaging plasmids (psPAX2, Addgene #12260; pMD2.G, Addgene #12259) into HEK293T cells using Lipofectamine 2000 transfection reagent (Thermo Fisher Scientific) following the manufacture’s protocol. Library DNA (4µg), psPAX2 DNA (4µg) and VSV-G DNA (2µg) was mixed and transfected into HEK293T cells in a T75 flask (×10). Six hours after transfection, media was removed and replaced with 10ml of virus production media (DMEM media supplemented with 10% of FBS). Forty-eight hours after transfection, lentiviral media was harvested, concentrated using Lenti-X™ Concentrator (Takara, cat. no. 631232) and stored at -80◦C.

<u>Virus titer determination</u>: 1×10^6^ cells (each cell line) were plated per well of a 6-well plate. Cells were infected with different amounts of lentivirus overnight in the presence of 8µg/ml of polybrene. The titering of lentiviral particles was performed by flow-cytometry following protocol from Cellecta, section 5.3 and 5.4: https://manuals.cellecta.com/crispr-pooled-lentiviral-sgrna-libraries/.

<u>Primary screening:</u> 4×10^7^ MM cells expressing dCAS-KRAB fusion protein were infected, using Spinoculation, with library lentiviral particles at MOI ranging from 0.1 to 0.3. Infection was performed in triplicate. Virus-containing media was removed after 1h of Spinoculation, cells were washed 2x with PBS and cultured in complete media. After 4 days, cells were selected with puromycin for 3 additional days. At day 7, cellular debris were removed by Ficoll-Hypaque (Lonza Group, Basel, Switzerland) density gradient sedimentation. Cells were cultured for additional 2 weeks ensuring a 1000x representation of library. Genomic DNA was isolated using Blood & Cell Culture DNA Maxi/Midi Kit (Qiagen #13362,13343) following the manufacturer’s protocol. Cellecta (Mountain View, CA) performed PCR amplification of the gRNA cassette for Illumina sequencing of gRNA representation. Protocols for PCR and Illumina sequencing are available online.

<u>Screening data analysis:</u> For candidate gene discovery, the normalized gRNA count table was loaded into MaGeCK (Model-based Analysis of Genome-wide CRISPR-Cas9 Knockout) (40) by comparing the experimental and control (plasmid library) conditions. Top genes were determined based on mean log2 fold change (LFC) for all gRNAs and false discovery rate (FDR).

<u>In vitro validation of MIR17HG:</u> Top scoring (n=4, see below) sgRNAs targeting MIR17HG were cloned into a pRSGT16-u6Tet-sg-CMV-TetRep-2A-TagRFP-2A-Puro (Cellecta, cat. #SVCRU6T16-L) vector and confirmed by sequencing. gRNA constructs were co-transfected with packaging plasmids (psPAX2, Addgene #12260; pMD2.G, Addgene #12259) into HEK293T cells using Lipofectamine 2000 transfection reagent (Thermo Fisher Scientific) following the manufacture’s protocol. Virus was harvested 48 hours later, concentrated and stored at -80◦C. MM cell lines stably expressing dCas9-KRAB fusion protein were infected with a lentivirus driving expression of individual sgRNAs. Infected cells were selected using puromycin. Expression of sgRNAs was obtained by doxycycline (0.5μg/mL, every other day).

*- MIR17HG* sgRNAs#1: AGTGGCGCGAAGGCGCAGGT
- *MIR17HG* sgRNAs#2: GTGGCGCGAAGGCGCAGGTC
- *MIR17HG* sgRNAs#3: CCTCGCCCGAGGGCGCGAAG
- *MIR17HG* sgRNAs#4: GAGGGCGCGAAGTGGCGCGA

### Antisense oligonucleotides, synthetic miRNA mimics and inhibitors, siRNAs

The following Long Non-Coding LNA gapmerRs were customly-designed and purchased from Exiqon (Vedbaek, Denmark):

ASO-NC (NC): GCTCCCTTCAATCCAA
ASO1: TACTTGCTTGGCTT
ASO2: CACCGTCCAAATCTAT
ASO3: AGCACTCAACATCAGC
5’-ASO1: CACCGTCCAAATCTAT
5’-ASO2: GTATGACTGGAATAGG
Murine aso1: TACAGTGGAAATCGGC
Murine aso2: GCGAGCAAACACGAAA
Murine aso3: ACTTGGATTGGATGAG

Synthetic mimics and inhibitors for miR-17a, miR-18a, miR-19a, miR-20a, miR-19b-1 and miR-92a1 were purchased from Ambion (Applied Biosystems, CA, US). Silencer selected siRNAs were purchased from Ambion (Applied Biosystems, CA, US).

Design of t-ASOs is described in **Supplementary Table 7**.

### Gymnosis

Gymonotic experiments were performed as previously described (Taiana et al., 2021). Briefly: cells were seeded at plating density to reach confluence on the final day of the experiments. Cell number at plating ranged from 0,5 to 2,5 x 10^3^ in 96-well plates, from 2,5 to 10 x 10^4^ in 12-well plates, from 1 to 3 x 10^5^ in 6-well plates. For ChIP and Co-IP experiments, cell number at plating was 1 x 10^6^ in T75 flask (10mL final volume).

### Transient transfection of cells

Adherent cell lines: cells were transfected by Lipofectamine 2000 according to manufacturer instructions with 25 nM of LNA gapmeRs (Exiqon).

<u>Suspension cell lines:</u> cells were transfected (electroporation) by Neon Transfection System (Invitrogen, CA, US), (2 pulses at 1150, 30ms). LNA gapmeRs, miRNA inhibitors/mimics and siRNAs were used at 25nM. The transfection efficiency evaluated by flow-cytometric analysis relative to a FAM dye–labeled anti-miR–negative control reached 85% to 90%.

### Stable expression using lentiviral plasmids

To generate cells stably over-expressing miR-17-92 cluster, AMO1 were transduced with PMIRH17-92PA-1 lenti-vector (System Biosciences, Palo Alto, CA, USA). To generate cells stably expressing c-MYC, U266 were transduced with Lenti ORF clone of Human v-myc myelocytomatosis viral oncogene homolog (avian) (MYC), Myc-DDK-tagged (RC201611L3) (Origene Technologies, Rockville, Maryland, MD). To generate cells stably expressing WDR82, AMO1 were transduced with Lenti ORF clone of Human WD repeat domain 82 (WDR82), mGFP tagged (RC216325L4) (Origene Technologies, Rockville, Maryland, MD). To generate cells stably expressing Cas9, AMO1 and H929 were transduced with pLX_311-Cas9 (Addgene #96924). Cells expressing the transgene were selected by antibiotic-selection for 3 to 5 days.

### CRISPR/CAS9 gene knockout

To generate DROSHA KO cells, AMO1 and H929 stably expressing Cas9 were transduced with transEDIT CRISPR single gRNA lentiviral expression vectors targeting DROSHA (CMV promoter, ZsGreen, TEVH-1203933) (transOMIC technologies Inc., Huntsville, AL, USA). ZsGreen+ cells were sorted (BD FACSARIA III; BD Biosciences, Qume Drive San Jose, CA, USA) 5 days after infection and cultured.

### Cell viability assay

Cell viability was evaluated by Cell Counting Kit-8 (CCK-8) assay (Dojindo Molecular Technologies) and 7-AminoactinoMYCin (7-AAD) flow cytometry assays (BD biosciences), according to manufacturer’s instructions. Flow cytometry analysis was performed either by FACS CANTO II (BD biosciences) or by Attune NxT Flow cytometer (Thermo Fisher Scientific).

### Detection of apoptosis

Apoptosis was investigated by Annexin V/7-AAD flow cytometry assay (BD biosciences) and by electronic microscopy. Flow cytometry analysis was performed either by FACS CANTO II (BD biosciences) or by Attune NxT Flow cytometer (Thermo Fisher Scientific).

### Reverse transcription (RT) and quantitative real-time amplification (qRT-PCR)

RNA extraction, reverse transcription (RT) and quantitative real-time amplification (qRT-PCR) were performed as previously described(Morelli et al., 2018). Briefly, total RNA was extracted from cells with TRIzol® Reagent (Thermo Fisher Scientific), according to manufacturer’s instructions. Nuclear and cytosolic subcellular RNA purification was performed using RNA Subcellular Isolation Kit (cat. no. 25501) (Active Motif, Carlsbad, CA), according to manufacturer’s instructions. The integrity of total RNA was verified by nanodrop (Celbio Nanodrop Spectrophotometer nd-1000). For RROL (MIR17HG) and mRNA dosage studies, oligo-dT-primed cDNA was obtained through the High Capacity cDNA Reverse Transcription Kit (Thermo Fisher Scientific) and then used as a template to quantify: a) *human* RROL (Hs03295901), ACACA or ACC1 (Hs01046047_m1), ANO6 (Hs03805835_m1), EXT1 (Hs00609156_m1), FER (Hs00245497_m1), MALAT1 (Hs00273907_s1) AND PVT1 (Hs00413039_m1). Normalization was performed with human GAPDH (Hs03929097_g1) or ACTB (Hs03023943_g1) or 18S (Hs03003631_g1). b) murine rrol (Mm01230322_s1), acaca (Mm01304258_m1) and fer (Mm00484303_m1). Single-tube TaqMan miRNA assay (Thermo Fisher Scientific) was used to detect and quantify miR-17 (002308), miR-18a (002422), miR-19a (000395), miR-20a (000580), miR-19b (000396) and miR-92a-1 (000431), according to the manufacturer’s instructions, by the use of ViiA7 RT reader (Thermo Fisher Scientific). Mature miRNAs expression was normalized on RNU44 (Thermo Fisher Scientific, assay Id: Hs03929097_g1). RROL isoforms were also detected by SYBR Green qRT-PCR using the following primers: RROL-1 (Fw, 5’-CCTGCAACTTCCTGGAGAAC; Rev, 5’-GTCTCAAGTGGGCATGATGA), RROL-2 (Fw, 5’-GACCCTCTTTTAAGTTGGGTG; Rev, 5’-TGGCAAAACATTTTCCTCCT). Comparative real-time polymerase chain-reaction (RT-PCR) was performed in triplicate, including no-template controls. Relative expression was calculated using the comparative cross threshold (Ct) method.

### Western blot analysis

Protein extraction and western blot analysis were performed as previously described. Briefly, cells were lysed in 1x RIPA buffer (Cell Signaling Technology) supplemented with Halt Protease Inhibitor Single-Use cocktail (100X, Thermo Scientific). Whole cells lysates (∼20 μg per lane) were separated using 4-12% Novex Bis-Tris SDS-acrylamide gels (Invitrogen), electro-transferred on Nitrocellulose membranes (Bio-Rad). Extraction of nuclear proteins was performed using the NE-PER™ Nuclear and Cytoplasmic Extraction Reagents (Thermo Fisher, #78833), according to manufacturer’s instructions. After electrophoresis the nitrocellulose membranes were blocked and probed over-night with primary antibodies at 4 ^0^C, then the membranes were washed 3 times in PBS-Tween and then incubated with a secondary antibody conjugated with horseradish peroxidase for 2 hours at room temperature. Chemiluminescence was detected using Western Blotting Luminol Reagent (sc-2048, Santa Cruz, Dallas, TX, USA).

Primary antibodies: anti-MYC [D84C1] (#5605), anti-WDR82 [D2I3B] (#99715), anti-H3K4me3 [C42D8] (#9751) and anti-Lamin A/C (#2032) antibodies were purchased from Cell Signaling Biotechnology (Danvers, MA). Anti-Drosha antibody [EPR12794] (ab183732) was purchased from Abcam (Cambridge, UK). Anti-MYC [9E10] (sc-40), GAPDH (sc-25778) and β-actin (ab96682) antibodies were purchased from Santa Cruz Biotechnology (Dallas, TX, USA). Monoclonal ANTI-FLAG® M2 antibody (F3165) was purchased from Millipore Sigma (Bedford, MA).

Secondary antibodies: Anti-rabbit IgG, HRP-linked Antibody (#7074) and Anti-mouse IgG, HRP-linked Antibody (#7076) were purchased from Cell Signaling Biotechnology (Danvers, MA).

### RNA FISH

RNA-FISH experiments were conducted according to established protocols (Raj et al., 2008; Shaffer et al., 2013). Cells were plated on coverslips coated with poly-L-lysine and allowed to attach for at least 1 hour. The media was then removed, the cells were washed once with 1X PBS, and then fixed and permeabilized in ice cold 95% methanol/5% acetic acid at 4°C for 10 minutes. After removing the fixative, cells were washed with Wash Buffer A (20% Stellaris RNA FISH Wash Buffer A, Biosearch Technologies, Inc., SMF-WA1-60; 10% Deionized Formamide, EMD Millipore, S4117; in RNAse-free water, Life Technologies, AM9932) for 5 minutes at room temperature. Cells were then incubated with RNA FISH probes (Stellaris) at a working concentration of 125 nM in Hybridization buffer (90% Stellaris RNA FISH Hybridization Buffer, Biosearch Technologies, SMF-HB1- 10; 10% Deionized Formamide) at 37°C in a humidified chamber in the dark overnight. The next day, cells were washed 3 times for 30 minutes each at 37°C in the dark with Wash Buffer A. The cells were then incubated for 15 minutes with Wash Buffer A plus 1:1000 Hoescht 33342 (Invitrogen, stock 10 mg/mL) at 37°C, followed by a wash with Wash Buffer B (Biosearch Technologies, SMF-WB1-20) for 5 minutes at room temperature. Coverslips were mounted on slides with Vectashield (VWR 101098-042), and coverslips were sealed with clear nail polish. Z-stack images were acquired on an LSM 880 with Airyscan with an oil-immersion 63X objective and a 2-3X zoom (W.M. Keck Microscopy Facility, MIT), and Airyscan processing was performed using the “Auto” strength feature. Representative images were generated using ImageJ.

### Co-immunofluorescence with RNA FISH (Co-IF/FISH)

Co-IF/FISH experiments were conducted in a similar fashion to the dual RNA-FISH experiment with the following modifications. After adhering the cells to coverslips, the cells were fixed with 4% PFA (VWR, BT140770) in RNase-free PBS for 10 minutes at room temperature. After washing the cells 3X for 5 minutes with PBS, the cells were permeabilized with ice cold 95% methanol/5% acetic acid at 4°C for 10 minutes. Cells were then blocked with 4% IgG-free Bovine Serum Albumin (VWR, 102643-516) in PBS for 30 minutes and a primary antibody mixture (1:500 Rabbit anti-c-MYC D84C12 in PBS) was then added to the cells and incubated overnight in a humidified chamber at room temperature. The next day, cells were washed 3X with PBS for 5 minutes at room temperature, and a secondary antibody mixture (1:500 Alexa Fluor 488 Goat anti-rabbit IgG, ThermoFisher A11008 in PBS) was added and incubated for 1 hour at room temperature in the dark. Cells were washed 3X with PBS for 5 minutes, and prior to RNA FISH, cells with antibody staining were re-fixed with 4% PFA in PBS for 10 minutes at room temperature, followed by 3X washes with PBS. After the antibody staining and fixation, the RNA FISH protocol was conducted as described above, starting with the wash with Wash Buffer A.

### Microarray-based gene expression profiling after RROL depletion

Microarray-based analysis of gene expression changes after treatment with ASO1 was performed as previously described (Morelli et al., 2018).

### RNA-seq analysis of AMO1^DR-KO^ after RROL depletion

Total RNA was extracted as described above and submitted to NovaSeq RNAseq analysis followed by VIPER NGS Analysis pipeline (Cornwell et al., 2018). List of differentially expressed genes (DEGs) were applied to the GSEA or IPA software to reveal biological pathways modulated by RROL.

### Luciferase reporter assay

Promoter reporter clones for human ACC1 (NM_198834), ANO6 (NM_001025356), CCDC91 (NM_018318), EPT1 (NM_033505), EXT1 (NM_000127), FER (NM_001308028) and ZYG11A (NM_001004339) were cloned into GLuc-ON™ Promoter Reporter Vector (GeneCopoeia, Rockville, MD). Luciferase reporter assay was performed according to manufacturer’s instructions.

### ChIRP

RROL and LacZ antisense DNA probes were designed using the online probe designer at singlemoleculefi sh.com. Oligonucleotides were biotinylated at the 3′ end with an 18-carbon spacer arm. AMO1 cells were collected and subjected to ChIRP using the EZ-Magna ChIRP RNA Interactome Kit (Millipore Sigma, Bedford, MA), according to manufacturer’s instructions and established protocols (Chu et al., 2012).

### De novo lipogenesis assay

Cells were seeded at 5×10^5^ cells per well in 6-well plates and incubated for 3 days in presence of treatments (ASO1 / 10058-F4 / IPTG or respective controls). Twenty-four hours before the end of treatment, 1 µCi of ^14^C-labeled glucose (ARC-0122D) was added to each well. Cells were harvested, washed with cold PBS and collected in glass tubes. Purified lipid extract is obtained by chloroform-methanol based extraction (Bligh and Dyer, 1959). Glucose incorporation in cellular lipids was quantitated by photon emission through scintillation counting and normalized to total protein content.

### Lipid profiling

Lipids were extracted from MM cells, dried and stored under argon until analysis. Lipid species were analyzed by liquid chromatography electrospray ionization tandem mass spectrometry (LC-ESI/MS/MS) on a Nexera X2 UHPLC system (Shimadzu) coupled with hybrid triple quadrupole/linear ion trap mass spectrometer (6500+ QTRAP system; AB SCIEX) by Lipometrix, at KU Leuven, Belgium.

<u>Lipid extraction:</u> lipid extraction was performed with 1 N HCl:CH_3_OH 1:8 (v/v), 900 μl CHCl_3_ and 200 μg/ml of the antioxidant 2,6-di-tert-butyl-4-methylphenol (BHT; Sigma Aldrich). A mixture of deuterium labeled lipids SPLASH® LIPIDOMIX® Mass Spec Standard (#330707, Avanti Polar Lipids) was spiked into the extract mix. The organic fraction was evaporated using a Savant Speedvac spd111v (Thermo Fisher Scientific) at room temperature and the remaining lipid pellet was stored at - 20°C under argon.

<u>Mass spectrometry:</u> just before mass spectrometry analysis, lipid pellets were reconstituted in 100% ethanol. Lipid species were analyzed by liquid chromatography electrospray ionization tandem mass spectrometry (LC-ESI/MS/MS) on a Nexera X2 UHPLC system (Shimadzu) coupled with hybrid triple quadrupole/linear ion trap mass spectrometer (6500+ QTRAP system; AB SCIEX). Chromatographic separation was performed on a XBridge amide column (150 mm × 4.6 mm, 3.5 μm; Waters) maintained at 35°C using mobile phase A [1 mM ammonium acetate in water-acetonitrile 5:95 (v/v)] and mobile phase B [1 mM ammonium acetate in water-acetonitrile 50:50 (v/v)] in the following gradient: (0-6 min: 0% B > 6% B; 6-10 min: 6% B > 25% B; 10-11 min: 25% B > 98% B; 11-13 min: 98% B > 100% B; 13-19 min: 100% B; 19-24 min: 0% B) at a flow rate of 0.7 mL/min which was increased to 1.5 mL/min from 13 minutes onwards. Sphingomyelins, ceramides, dihydroceramides, hexosylceramides and lactosylceramides were measured in positive ion mode with a precursor scan of 184.1, 264.4, 266.4, 264.4 and 264.4 respectively. Triacylglycerides and diacylglycerides were measured in positive ion mode with a neutral loss scan for one of the fatty acyl moieties. Phosphatidylcholine, lysophosphatidylcholine, phosphatidylethanolamine, lysophosphatidylethanolamine, Phosphatidylglycerol, phosphatidylinositol and phosphatidylserine were measured in negative ion mode by fatty acyl fragment ions. Lipid quantification was performed by scheduled multiple reactions monitoring (MRM), the transitions being based on the neutral losses or the typical product ions as described above. The instrument parameters were as follows: Curtain Gas = 35 psi; Collision Gas = 8 a.u. (medium); IonSpray Voltage = 5500 V and −4,500 V; Temperature = 550°C; Ion Source Gas 1 = 50 psi; Ion Source Gas 2 = 60 psi; Declustering Potential = 60 V and −80 V; Entrance Potential = 10 V and −10 V; Collision Cell Exit Potential = 15 V and −15 V. The following fatty acyl moieties were taken into account for the lipidomic analysis: 14:0, 14:1, 16:0, 16:1, 16:2, 18:0, 18:1, 18:2, 18:3, 20:0, 20:1, 20:2, 20:3, 20:4, 20:5, 22:0, 22:1, 22:2, 22:4, 22:5 and 22:6 except for TGs which considered: 16:0, 16:1, 18:0, 18:1, 18:2, 18:3, 20:3, 20:4, 20:5, 22:2, 22:3, 22:4, 22:5, 22:6.

<u>Data Analysis:</u> peak integration was performed with the MultiQuantTM software version 3.0.3. Lipid species signals were corrected for isotopic contributions (calculated with Python Molmass 2019.1.1) and were quantified based on internal standard signals and adheres to the guidelines of the Lipidomics Standards Initiative (LSI) (level 2 type quantification as defined by the LSI).

### ChIP-qPCR

ChIP-qPCR was performed as previously described (Fulciniti et al., 2018). Briefly, 1×10^7^ cells (AMO1, H929 and U266^MYC+^, with corresponding treatments) were cross-linked with 1% formaldehyde for 10 minutes at 37°C. The cross-linked chromatin was then extracted, diluted with lysis buffer, and sheared by sonication. The chromatin was divided into equal samples for immunoprecipitation with specific antibodies. The immunoprecipitates were pelleted by centrifugation and incubated at 68°C to reverse the protein-DNA cross-linking. The DNA was extracted from the elute by the Qiaquick PCR purification kit (QIAGEN). Antibodies used were as follows: endogenous MYC (Cell Signaling Technology, #13987), MYC-DDK (Santa Cruz Biotechnology, 9E10-x), GFP (Abcam, #ab290), H3K4me3 (#ab8580), Normal Rabbit IgG (Cell Signaling Technology, #2729), Normal Mouse IgG (Santa Cruz Biotechnology, sc-2025). A parallel sample of input DNA from the same cells was used as control. ChIP and input DNA were analyzed using SYBR Green real-time PCR analysis (Applied Biosystems).

Primers for ChIP-qPCR:

ACC1 Fw: TTTCTCTCTTGCAGAGTGAGGTGTGG
ACC1 Rv: TACAAAGGCACGGAGAGAGCAAGT

### RNA-Protein Pull-Down

RROL transcripts were cloned into a pBlueScript vector (kindly provided by Dr. Giada Bianchi, Dana-Farber Cancer Institute, Boston, MA) and sequence verified. In vitro transcription and biotynilation was performed using AmpliScribe™ T7-Flash™ Biotin-RNA Transcription Kit (Lucigen, cat. no. #ASB71110), according to manufacturer’s instructions. Cell nuclear lysates (from 1×10^7^ AMO1 cells) were incubated with biotinylated RNA and streptavidin beads for RNA pull-down incubation, using Pierce™ Magnetic RNA-Protein Pull-Down Kit (Thermo Fisher Scientific, cat. no. #20164), according to manufacturer’s instructions. RNA-associated proteins were eluted and analyzed by western blotting.

### RNA Yeast 3 Hybrid

Saccharomyces cerevisiae strain YLW3 was transformed with RNA plasmids, using standard protocols. They were tested for viability by spotting on SC plates depleted of uracil (SC-U). Protein plasmids were transformed into the yeast strain Y8800 and grown in SC-plates depleted of tryptophan (SC-W). Yeast strains YLW3 containing the examined RNA plasmid were mated with the Y8800 yeast strains containing the protein plasmid. Mating was performed according to the manufacturer’s protocol in YPD media. Diploids carrying both plasmids were selected in SC media depleted of tryptophan and uracil (SD-WU), and dimerization was tested by growth in (SC-WUH) media, also depleted of histidine. The following day the diploids in the SC-WUH media were transferred to solid agar plates containing different levels of 3AT, a competitive inhibitor of the HIS3 gene product, to increase the stringency of the selection. Only the diploids with significant interaction should be able to produce enough histidine for survival. After 1-3 days the growth of the different colonies in the different conditions was examined to seek out the diploids with the strongest interactions.

### RIP-qPCR

RNA immunoprecipitation (RIP) experiments were performed using the Magna RIP RNA-binding Protein Immunoprecipitation Kit (Millipore Sigma, cat. no. 17-701), according to manufacturer’s instructions. The anti-MYC antibody [Y69] used for RIP was purchased from Abcam (ab32072). Normal Rabbit IgG was purchased from Cell Signaling Technology (cat. no. #2729). The primers used for detecting RROL are listed above.

### Co-immunoprecipitation (Co-IP)

Protein lysates were obtained from 1×10^7^ cells (AMO1, H929 and U266^MYC+^, with corresponding treatments). Coimmunoprecipitation was performed using Pierce™ Co-Immunoprecipitation Kit (Thermo Fisher Scientific, cat. no. 26149), according to manufacturer’s instructions. IP antibodies used were as follows: anti-MYC antibody [Y69] was purchased from Abcam (ab32072), Anti-FLAG® M2 antibody was purchased from Millipore Sigma (F3165), Normal Rabbit IgG was purchased from Cell Signaling Technology (2729).

### Proximity-dependent biotin identification (BioID)

BioID was performed as described by Kalkat et al.(Kalkat et al., 2018). FBA and FBA-MYC cells were kindly provided by Dr. Linda Penn (University of Toronto). Briefly, FBA-MYC cells were grown to 60% confluence into T75 flasks prior to transfection with ASO1 (50nM, using Lipofectamine 2000 as described above) and treatment with 1 mg/mL doxycycline (Millipore Sigma), 1 μM MG132 (Millipore Sigma) and 50 mM biotin (Bio Basic) for 24 hours. Experiments with FBA-MYC cells, exposed to doxycycline, included 16 biological replicates (8 with RROL depletion and 8 without RROL depletion). Negative controls used for the analysis included 6 biological replicates of FBA-MYC cells not exposed to doxycycline. Cells were harvested by scraping and washed three times with 50mL of PBS prior to flash freezing. Cell pellets were lysed in 1 mL of modified RIPA buffer (1% NP-40, 50 mM Tris–HCl pH 7.5, 150 mM NaCl, 1 mM EDTA, 1 mM EGTA, 0.1% SDS, 1:100 protease inhibitor cocktail (Thermo Fisher Scientific), 0.5% sodium deoxycholate), with 250U of benzonase (Millipore). Lysate was rotated for 1 h at 4°C, sonicated 3x30 s, then centrifuged at 27000 g for 30 min at 4°C. Biotinylated proteins were isolated by affinity purification with 30 mg of washed streptavidin-Sepharose beads (GE) with rotation for 2 h at 4°C. Beads were then washed 7x1 mL 50 mM ammonium bicarbonate (pH 8.0) prior to tryptic digestion.

### Mass Spectrometry

Mass Spectrometry analysis of Co-IP and BioID samples was performed at the Taplin Mass Spectrometry Facility (Harvard Medical School, Boston, MA), according to established protocols.

### Animal study

6-week old female immunodeficient NOD.CB17-Prkdcscid/NCrCrl (NOD/SCID) mice (Charles River) or NSG mice (Jackson Laboratory) were housed in our animal facility at Dana-Farber Cancer Institute (DFCI). All experiments were performed after approval by the Animal Ethics Committee of the DFCI and performed using institutional guidelines. AMO1^DR-KO^ xenograft model: AMO^DR-KO^ were gymnotically exposed to ASO1 (2,5uM) or ASO-NC (2,5uM) for 2 days before subcutaneous injection into SCID NOD mice. The day of injection (day 0), cell viability was assessed by Annexin V / 7-AAD flow cytometry assay, confirming no detectable pro-apoptotic activity of ASO-1 at this time point (not shown). For tumor cells injection, cells were resuspended in PBS1X supplemented with ASO1 (5uM) or ASO-NC (5uM); and then mixed with equivalent volume of Matrigel (Corning, #354230) reaching a final oligo concentration of 2,5uM. 5×10^6^ cells were subcutaneously injected per mice (5 mice per group). Tumor sizes were measured by electronic caliper.

<u>AMO1 xenograft model:</u> 5*10^6^ AMO1 cells were subcutaneously injected in NOD SCID mice. As tumor became palpable (∼50mm), mice were randomized to receive G2-15b*- TO or SB9-19-TO or vehicle (-) as control (3 groups, 5 mice/group). Treatments were administered via I.P. injection, every other day per 2 weeks, at 10mg/kg. Tumor sizes were measured by electronic caliper. In an independent experiment used for qRT-PCR analysis of RROL and ACC1, mice were enrolled to receive treatment after tumors reached the volume of ∼200mm and treated at day 1-3-5. Tumors were then collected at day 6.

<u>MOLP8-luc+ xenograft model:</u> 1*10^6^ MOLP8-luc+ cells were injected via tail vein in 28 NSG mice. 3 mice (marked by a X) were then excluded for failed injection. The day after, 11 mice were assigned to the control group, 8 mice for treatment with G2-19b*-TO and 6 mice for treatment with SB9-19-TO. Treatments were administered via I.P. injection, every other day per 2 weeks, at 10mg/kg. At the end of the treatment cycle (day 15), BLI was measured as indication of tumor growth.

Tumor growth inhibition (%TGI) was determined, as previously described (Buck et al., 2008), by the formula: %TGI = (1- [Tt/T0 / Ct/C0] 1-[C0/Ct]) X 100 where Tt = median tumor volume of treated at time t, T0 = median tumor volume of treated at time 0, Ct = median tumor volume of control at time t and C0 = median tumor volume of control at time 0.

### Statistical Analysis

All in vitro experiments were repeated at least three times and performed in triplicate; a representative experiment was showed in figures. Statistical significances of differences were determined using Student’s t test (unless otherwise specified), with minimal level of significance specified as p<0.05. Kaplan-Meier survival curves were compared by log-rank test. Statistical analyses were determined using GraphPad software (http://www.graphpad.com). Graphs were obtained using GraphPad software (unless otherwise specified).

### Data availability

The authors declare that all data supporting the findings of this study are available within the article and its Supplementary Information. Files or reagents are available from the corresponding authors on request.

## REFERENCES

Amente, S., Lania, L., and Majello, B. (2011). Epigenetic reprogramming of Myc target genes. Am J Cancer Res 1, 413–418.

Amodio, N., Stamato, M. A., Juli, G., Morelli, E., Fulciniti, M., Manzoni, M., Taiana, E., Agnelli, L., Cantafio, M. E. G., Romeo, E., et al. (2018). Drugging the lncRNA MALAT1 via LNA gapmeR ASO inhibits gene expression of proteasome subunits and triggers anti-multiple myeloma activity. Leukemia 32, 1948–1957.

Bartel, D. P. (2004). MicroRNAs: genomics, biogenesis, mechanism, and function. Cell 116, 281–297.

Beloribi-Djefaflia, S., Vasseur, S., and Guillaumond, F. (2016). Lipid metabolic reprogramming in cancer cells. Oncogenesis 5, e189.

Bligh, E. G., and Dyer, W. J. (1959). A rapid method of total lipid extraction and purification. Can J Biochem Physiol 37, 911–917.

Buck, E., Eyzaguirre, A., Rosenfeld-Franklin, M., Thomson, S., Mulvihill, M., Barr, S., Brown, E., O’Connor, M., Yao, Y., Pachter, J., et al. (2008). Feedback mechanisms promote cooperativity for small molecule inhibitors of epidermal and insulin-like growth factor receptors. Cancer Res 68, 8322–8332.

Chesi, M., Robbiani, D. F., Sebag, M., Chng, W. J., Affer, M., Tiedemann, R., Valdez, R., Palmer, S. E., Haas, S. S., Stewart, A. K., et al. (2008). AID-dependent activation of a MYC transgene induces multiple myeloma in a conditional mouse model of post-germinal center malignancies. Cancer Cell 13, 167–180.

Chu, C., Quinn, J., and Chang, H. Y. (2012). Chromatin isolation by RNA purification (ChIRP). J Vis Exp.

Cornwell, M., Vangala, M., Taing, L., Herbert, Z., Koster, J., Li, B., Sun, H., Li, T., Zhang, J., Qiu, X., et al. (2018). VIPER: Visualization Pipeline for RNA-seq, a Snakemake workflow for efficient and complete RNA-seq analysis. BMC Bioinformatics 19, 135.

Crooke, S. T., Witztum, J. L., Bennett, C. F., and Baker, B. F. (2018). RNA-Targeted Therapeutics. Cell Metab 27, 714–739.

Cummins, J. M., He, Y., Leary, R. J., Pagliarini, R., Diaz, L. A., Jr., Sjoblom, T., Barad, O., Bentwich, Z., Szafranska, A. E., Labourier, E., et al. (2006). The colorectal microRNAome. Proc Natl Acad Sci U S A 103, 3687–3692.

Dai, M., Wang, P., Boyd, A. D., Kostov, G., Athey, B., Jones, E. G., Bunney, W. E., Myers, R. M., Speed, T. P., Akil, H., et al. (2005). Evolving gene/transcript definitions significantly alter the interpretation of GeneChip data. Nucleic Acids Res 33, e175.

Damase, T. R., Sukhovershin, R., Boada, C., Taraballi, F., Pettigrew, R. I., and Cooke, J. P. (2021). The Limitless Future of RNA Therapeutics. Front Bioeng Biotechnol 9, 628137.

Dhuri, K., Bechtold, C., Quijano, E., Pham, H., Gupta, A., Vikram, A., and Bahal, R. (2020). Antisense Oligonucleotides: An Emerging Area in Drug Discovery and Development. J Clin Med 9.

El Arfani, C., De Veirman, K., Maes, K., De Bruyne, E., and Menu, E. (2018). Metabolic Features of Multiple Myeloma. Int J Mol Sci 19.

Falchook, G., Infante, J., Arkenau, H. T., Patel, M. R., Dean, E., Borazanci, E., Brenner, A., Cook, N., Lopez, J., Pant, S., et al. (2021). First-in-human study of the safety, pharmacokinetics, and pharmacodynamics of first-in-class fatty acid synthase inhibitor TVB-2640 alone and with a taxane in advanced tumors. EClinicalMedicine 34, 100797.

Fulciniti, M., Lin, C. Y., Samur, M. K., Lopez, M. A., Singh, I., Lawlor, M. A., Szalat, R. E., Ott, C. J., Avet-Loiseau, H., Anderson, K. C., et al. (2018). Non-overlapping Control of Transcriptome by Promoter- and Super-Enhancer-Associated Dependencies in Multiple Myeloma. Cell Rep 25, 3693–3705 e3696.

Gouw, A. M., Margulis, K., Liu, N. S., Raman, S. J., Mancuso, A., Toal, G. G., Tong, L., Mosley, A., Hsieh, A. L., Sullivan, D. K., et al. (2019). The MYC Oncogene Cooperates with Sterol-Regulated Element-Binding Protein to Regulate Lipogenesis Essential for Neoplastic Growth. Cell Metab 30, 556–572 e555.

Gulla, A., and Anderson, K. C. (2020). Multiple myeloma: the (r)evolution of current therapy and a glance into future. Haematologica.

Gutschner, T., and Diederichs, S. (2012). The hallmarks of cancer: a long non-coding RNA point of view. RNA Biol 9, 703–719.

He, L., Thomson, J. M., Hemann, M. T., Hernando-Monge, E., Mu, D., Goodson, S., Powers, S., Cordon-Cardo, C., Lowe, S. W., Hannon, G. J., and Hammond, S. M. (2005). A microRNA polycistron as a potential human oncogene. Nature 435, 828–833.

Hon, C. C., Ramilowski, J. A., Harshbarger, J., Bertin, N., Rackham, O. J., Gough, J., Denisenko, E., Schmeier, S., Poulsen, T. M., Severin, J., et al. (2017). An atlas of human long non-coding RNAs with accurate 5’ ends. Nature 543, 199–204.

Hook, B., Bernstein, D., Zhang, B., and Wickens, M. (2005). RNA-protein interactions in the yeast three-hybrid system: affinity, sensitivity, and enhanced library screening. RNA 11, 227–233.

Hu, Y., Lin, J., Fang, H., Fang, J., Li, C., Chen, W., Liu, S., Ondrejka, S., Gong, Z., Reu, F., et al. (2018). Targeting the MALAT1/PARP1/LIG3 complex induces DNA damage and apoptosis in multiple myeloma. Leukemia 32, 2250–2262.

Izreig, S., Samborska, B., Johnson, R. M., Sergushichev, A., Ma, E. H., Lussier, C., Loginicheva, E., Donayo, A. O., Poffenberger, M. C., Sagan, S. M., et al. (2016). The miR-17 approximately 92 microRNA Cluster Is a Global Regulator of Tumor Metabolism. Cell Rep 16, 1915–1928.

Jovanovic, K. K., Escure, G., Demonchy, J., Willaume, A., Van de Wyngaert, Z., Farhat, M., Chauvet, P., Facon, T., Quesnel, B., and Manier, S. (2018). Deregulation and Targeting of TP53 Pathway in Multiple Myeloma. Front Oncol 8, 665.

Kalkat, M., Resetca, D., Lourenco, C., Chan, P. K., Wei, Y., Shiah, Y. J., Vitkin, N., Tong, Y., Sunnerhagen, M., Done, S. J., et al. (2018). MYC Protein Interactome Profiling Reveals Functionally Distinct Regions that Cooperate to Drive Tumorigenesis. Mol Cell 72, 836–848 e837.

Lai, F., Damle, S. S., Ling, K. K., and Rigo, F. (2020). Directed RNase H Cleavage of Nascent Transcripts Causes Transcription Termination. Mol Cell 77, 1032–1043 e1034.

Lee, J. H., and Skalnik, D. G. (2008). Wdr82 is a C-terminal domain-binding protein that recruits the Setd1A Histone H3-Lys4 methyltransferase complex to transcription start sites of transcribed human genes. Mol Cell Biol 28, 609–618.

Lee, J. S., and Mendell, J. T. (2020). Antisense-Mediated Transcript Knockdown Triggers Premature Transcription Termination. Mol Cell 77, 1044–1054 e1043.

Leone, E., Morelli, E., Di Martino, M. T., Amodio, N., Foresta, U., Gulla, A., Rossi, M., Neri, A., Giordano, A., Munshi, N. C., et al. (2013). Targeting miR-21 inhibits in vitro and in vivo multiple myeloma cell growth. Clin Cancer Res 19, 2096–2106.

Li, W., Xu, H., Xiao, T., Cong, L., Love, M. I., Zhang, F., Irizarry, R. A., Liu, J. S., Brown, M., and Liu, X. S. (2014a). MAGeCK enables robust identification of essential genes from genome-scale CRISPR/Cas9 knockout screens. Genome Biol 15, 554.

Li, Y., Choi, P. S., Casey, S. C., Dill, D. L., and Felsher, D. W. (2014b). MYC through miR-17-92 suppresses specific target genes to maintain survival, autonomous proliferation, and a neoplastic state. Cancer Cell 26, 262–272.

Lionetti, M., Barbieri, M., Todoerti, K., Agnelli, L., Marzorati, S., Fabris, S., Ciceri, G., Galletti, S., Milesi, G., Manzoni, M., et al. (2015). Molecular spectrum of BRAF, NRAS and KRAS gene mutations in plasma cell dyscrasias: implication for MEK-ERK pathway activation. Oncotarget 6, 24205–24217.

Liu, S. J., Horlbeck, M. A., Cho, S. W., Birk, H. S., Malatesta, M., He, D., Attenello, F. J., Villalta, J. E., Cho, M. Y., Chen, Y., et al. (2017). CRISPRi-based genome-scale identification of functional long noncoding RNA loci in human cells. Science 355.

Lu, Y., Zhao, X., Liu, Q., Li, C., Graves-Deal, R., Cao, Z., Singh, B., Franklin, J. L., Wang, J., Hu, H., et al. (2017). lncRNA MIR100HG-derived miR-100 and miR-125b mediate cetuximab resistance via Wnt/beta-catenin signaling. Nat Med 23, 1331–1341.

Masarwi, M., DeSchiffart, A., Ham, J., and Reagan, M. R. (2019). Multiple Myeloma and Fatty Acid Metabolism. JBMR Plus 3, e10173.

Mogilyansky, E., and Rigoutsos, I. (2013). The miR-17/92 cluster: a comprehensive update on its genomics, genetics, functions and increasingly important and numerous roles in health and disease. Cell Death Differ 20, 1603–1614.

Mondala, P. K., Vora, A. A., Zhou, T., Lazzari, E., Ladel, L., Luo, X., Kim, Y., Costello, C., MacLeod, A. R., Jamieson, C. H. M., and Crews, L. A. (2021). Selective antisense oligonucleotide inhibition of human IRF4 prevents malignant myeloma regeneration via cell cycle disruption. Cell Stem Cell 28, 623–636 e629.

Morelli, E., Biamonte, L., Federico, C., Amodio, N., Di Martino, M. T., Gallo Cantafio, M. E., Manzoni, M., Scionti, F., Samur, M. K., Gulla, A., et al. (2018). Therapeutic vulnerability of multiple myeloma to MIR17PTi, a first-in-class inhibitor of pri-miR-17-92. Blood 132, 1050–1063.

Morelli, E., Gulla, A., Amodio, N., Taiana, E., Neri, A., Fulciniti, M., and Munshi, N. C. (2021). CRISPR Interference (CRISPRi) and CRISPR Activation (CRISPRa) to Explore the Oncogenic lncRNA Network. Methods Mol Biol 2348, 189–204.

Morelli, E., Gulla, A., Rocca, R., Federico, C., Raimondi, L., Malvestiti, S., Agosti, V., Rossi, M., Costa, G., Giavaresi, G., et al. (2020). The Non-Coding RNA Landscape of Plasma Cell Dyscrasias. Cancers (Basel) 12.

Morelli, E., Leone, E., Cantafio, M. E., Di Martino, M. T., Amodio, N., Biamonte, L., Gulla, A., Foresta, U., Pitari, M. R., Botta, C., et al. (2015). Selective targeting of IRF4 by synthetic microRNA-125b-5p mimics induces anti-multiple myeloma activity in vitro and in vivo. Leukemia 29, 2173–2183.

Ota, A., Tagawa, H., Karnan, S., Tsuzuki, S., Karpas, A., Kira, S., Yoshida, Y., and Seto, M. (2004). Identification and characterization of a novel gene, C13orf25, as a target for 13q31-q32 amplification in malignant lymphoma. Cancer Res 64, 3087–3095.

Pichiorri, F., Suh, S. S., Rocci, A., De Luca, L., Taccioli, C., Santhanam, R., Zhou, W., Benson, D. M., Jr., Hofmainster, C., Alder, H., et al. (2016). Downregulation of p53-inducible microRNAs 192, 194, and 215 Impairs the p53/MDM2 Autoregulatory Loop in Multiple Myeloma Development. Cancer Cell 30, 349–351.

Puttaraju, M., Jackson, M., Klein, S., Shilo, A., Bennett, C. F., Gordon, L., Rigo, F., and Misteli, T. (2021). Systematic screening identifies therapeutic antisense oligonucleotides for Hutchinson-Gilford progeria syndrome. Nat Med.

Raj, A., van den Bogaard, P., Rifkin, S. A., van Oudenaarden, A., and Tyagi, S. (2008). Imaging individual mRNA molecules using multiple singly labeled probes. Nat Methods 5, 877–879.

Rohrig, F., and Schulze, A. (2016). The multifaceted roles of fatty acid synthesis in cancer. Nat Rev Cancer 16, 732–749.

Rysman, E., Brusselmans, K., Scheys, K., Timmermans, L., Derua, R., Munck, S., Van Veldhoven, P. P., Waltregny, D., Daniels, V. W., Machiels, J., et al. (2010). De novo lipogenesis protects cancer cells from free radicals and chemotherapeutics by promoting membrane lipid saturation. Cancer Res 70, 8117–8126.

Samur, M. K., Minvielle, S., Gulla, A., Fulciniti, M., Cleynen, A., Aktas Samur, A., Szalat, R., Shammas, M., Magrangeas, F., Tai, Y. T., et al. (2018). Long intergenic non-coding RNAs have an independent impact on survival in multiple myeloma. Leukemia 32, 2626–2635.

Schuhmacher, M., Staege, M. S., Pajic, A., Polack, A., Weidle, U. H., Bornkamm, G. W., Eick, D., and Kohlhuber, F. (1999). Control of cell growth by c-Myc in the absence of cell division. Curr Biol 9, 1255–1258.

Shaffer, A. L., Emre, N. C., Lamy, L., Ngo, V. N., Wright, G., Xiao, W., Powell, J., Dave, S., Yu, X., Zhao, H., et al. (2008). IRF4 addiction in multiple myeloma. Nature 454, 226–231.

Shaffer, S. M., Wu, M. T., Levesque, M. J., and Raj, A. (2013). Turbo FISH: a method for rapid single molecule RNA FISH. PLoS One 8, e75120.

Stine, Z. E., Walton, Z. E., Altman, B. J., Hsieh, A. L., and Dang, C. V. (2015). MYC, Metabolism, and Cancer. Cancer Discov 5, 1024–1039.

Sullenger, B. A., and Nair, S. (2016). From the RNA world to the clinic. Science 352, 1417–1420.

Taiana, E., Favasuli, V., Ronchetti, D., Morelli, E., Tassone, P., Viglietto, G., Munshi, N. C., Neri, A., and Amodio, N. (2021). In Vitro Silencing of lncRNAs Using LNA GapmeRs. Methods Mol Biol 2348, 157–166.

Tessoulin, B., Moreau-Aubry, A., Descamps, G., Gomez-Bougie, P., Maiga, S., Gaignard, A., Chiron, D., Menoret, E., Le Gouill, S., Moreau, P., et al. (2018). Whole-exon sequencing of human myeloma cell lines shows mutations related to myeloma patients at relapse with major hits in the DNA regulation and repair pathways. J Hematol Oncol 11, 137.

Todoerti, K., Agnelli, L., Fabris, S., Lionetti, M., Tuana, G., Mosca, L., Lombardi, L., Grieco, V., Bianchino, G., D’Auria, F., et al. (2013). Transcriptional characterization of a prospective series of primary plasma cell leukemia revealed signatures associated with tumor progression and poorer outcome. Clin Cancer Res 19, 3247–3258.

Tseng, Y. Y., Moriarity, B. S., Gong, W., Akiyama, R., Tiwari, A., Kawakami, H., Ronning, P., Reuland, B., Guenther, K., Beadnell, T. C., et al. (2014). PVT1 dependence in cancer with MYC copy-number increase. Nature 512, 82–86.

Ulitsky, I., and Bartel, D. P. (2013). lincRNAs: genomics, evolution, and mechanisms. Cell 154, 26–46.

Wang, Z., Yang, B., Zhang, M., Guo, W., Wu, Z., Wang, Y., Jia, L., Li, S., Cancer Genome Atlas Research, N., Xie, W., and Yang, D. (2018). lncRNA Epigenetic Landscape Analysis Identifies EPIC1 as an Oncogenic lncRNA that Interacts with MYC and Promotes Cell-Cycle Progression in Cancer. Cancer Cell 33, 706–720 e709.

Zadra, G., Ribeiro, C. F., Chetta, P., Ho, Y., Cacciatore, S., Gao, X., Syamala, S., Bango, C., Photopoulos, C., Huang, Y., et al. (2019). Inhibition of de novo lipogenesis targets androgen receptor signaling in castration-resistant prostate cancer. Proc Natl Acad Sci U S A 116, 631–640.

Zaidi, N., Lupien, L., Kuemmerle, N. B., Kinlaw, W. B., Swinnen, J. V., and Smans, K. (2013). Lipogenesis and lipolysis: the pathways exploited by the cancer cells to acquire fatty acids. Prog Lipid Res 52, 585–589.

Zhang, B., Lu, H. Y., Xia, Y. H., Jiang, A. G., and Lv, Y. X. (2018). Long non-coding RNA EPIC1 promotes human lung cancer cell growth. Biochem Biophys Res Commun 503, 1342–1348.

