## Supplementary Figures for "Long Noncoding RNA RROL Provides Chromatin Scaffold for MYC-WDR82 Interaction to Impact Lipid Metabolism and Tumor Cell Growth in Multiple Myeloma"

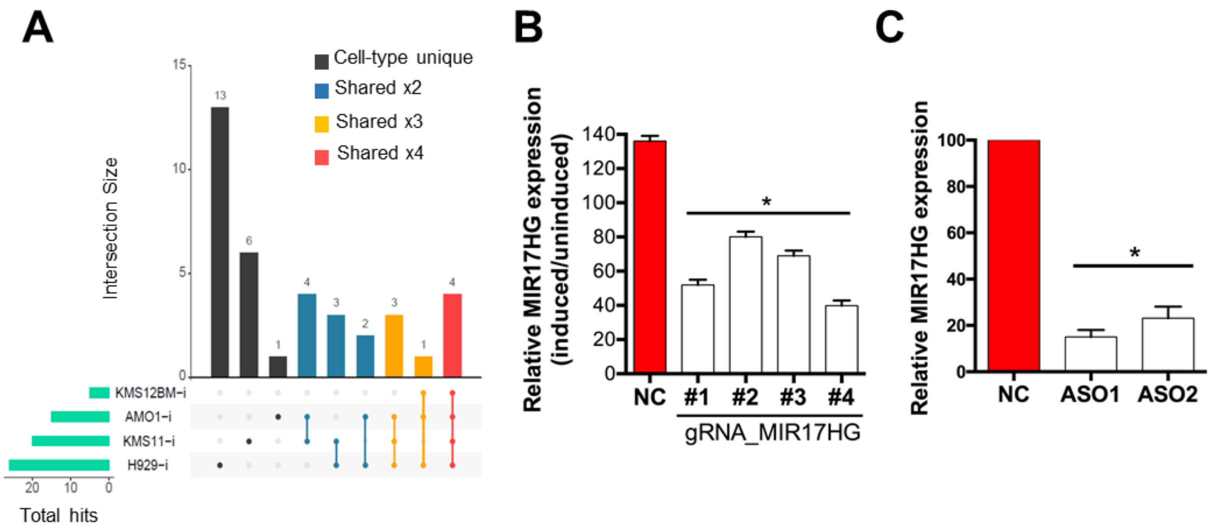


**Supplementary Fig. 1. A**) Analysis of screening data, with the upset plot showing the identification of cell-type unique and shared lncRNA dependencies in MM cells. **B**) Knockdown of MIR17HG obtained in AMO1 engineered to express a dCas9-KRAB fusion protein and anti-MIR17HG gRNAs under the regulation of a conditional promoter. MIR17HG expression was investigated by qRT-PCR five days after induction of gRNAs with doxycycline. The results shown are average RNA expression levels after normalization with ACTB and ΔΔCt calculations. **C**) Knockdown of MIR17HG obtained in AMO1 transfected with two different gapmeR ASOs (25nM) targeting the MIR17HG nascent RNA (ASO1 and -2), or a control ASO. MIR17HG expression was investigated by qRT-PCR 24h after transfection. The results shown are average RNA expression levels after normalization with ACTB and ΔΔCt calculations. 1 of three independent experiments is shown. *Indicates $p<0.05$

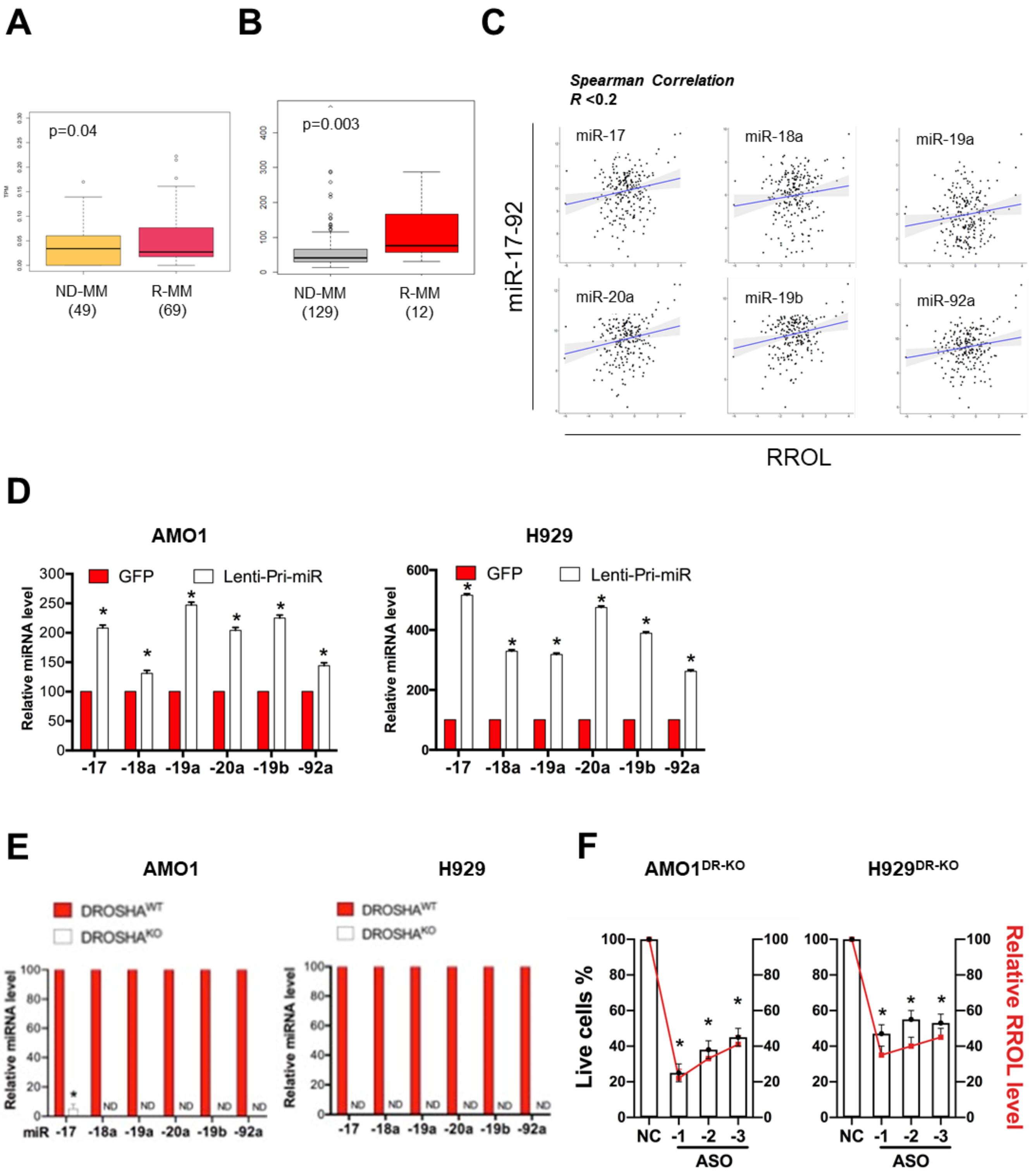


**Supplementary Fig. 2. A**) RROL expression in newly diagnosed (ND) vs relapsed (R) MM patients enrolled in the IFM/DFCI clinical trial (NCT01191060). **B**) RROL expression in ND vs R MM patients from GSE66293. **C**) Spearman's correlation between RROL and miR-17-92 in

CD138+ cells from 140 MM patients analyzed by RNA-seq and miRNA profiling. **D**) qRT-PCR analysis of miR-17-92 microRNAs in AMO1 and H929 either infected with a lentiviral vector carrying the expression of pri-mir-17-92 (Lenti-pri-mir) or with lentiviral vector carrying the expression of GFP. The results shown are average miRNA expression levels after normalization with RNU44 and ΔΔCt calculations. **E**) qRT-PCR analysis of miR-17-92 microRNAs in AMO1 and H929 either WT or KO for DROSHA. The results shown are average miRNA expression levels after normalization with RNU44 and ΔΔCt calculations. **F**) Knockdown of MIR17HG using 3 different ASOs, or a scramble control (NC), in $AMO1^{DR\text{-}KO}$ and $H929^{DR\text{-}KO}$. Live cells %, compared to NC, was analyzed 48h after transfection by CCK-8 assay. RROL expression was analyzed 48h after transfection by qRT-PCR. 1 of three independent experiments is shown D-E-F. *Indicates $p<0.05$ after Student t test calculation.

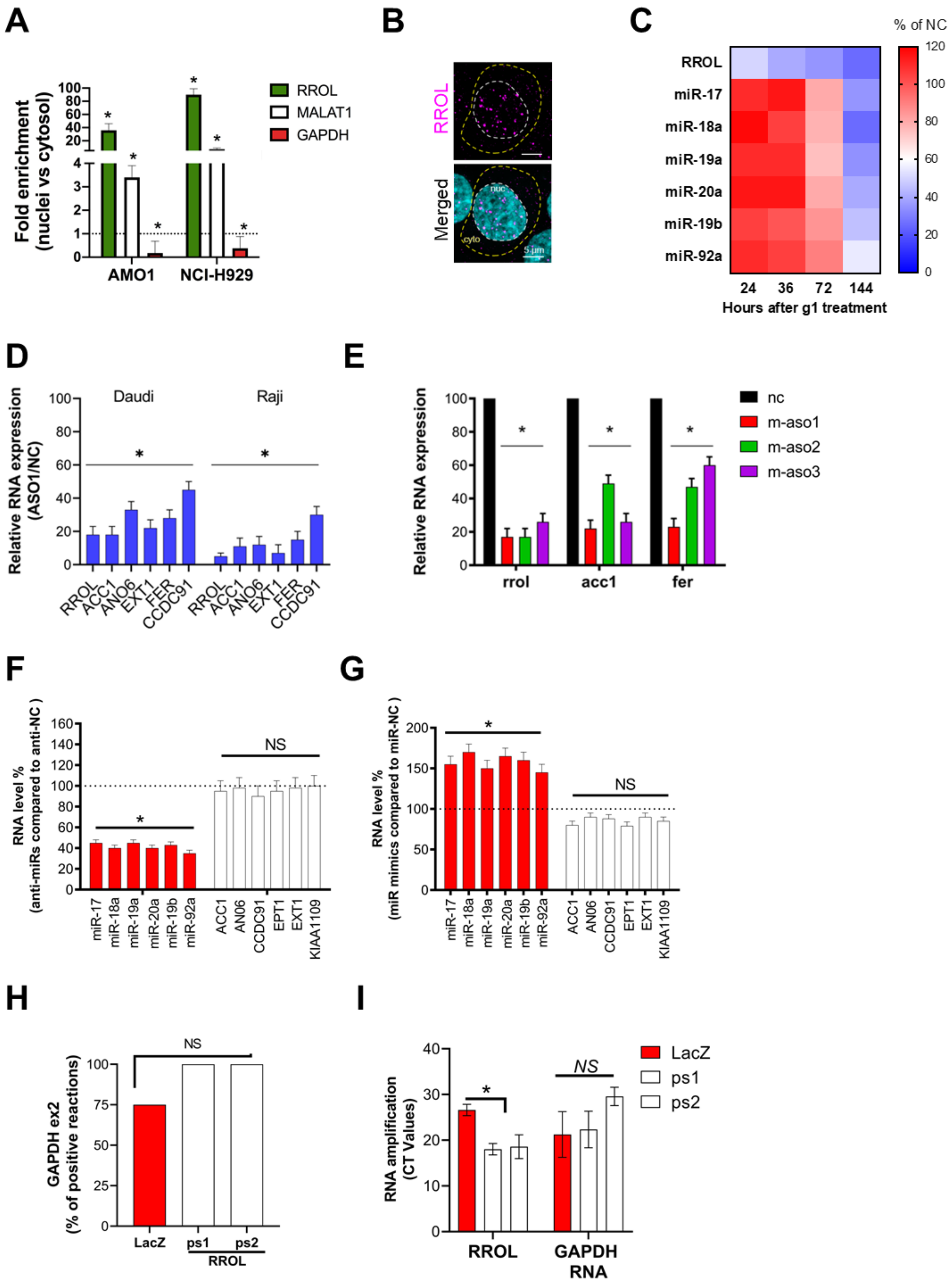

A
Fold enrichment (nuclei vs cytosol)
100
80
60
40
20
4
3
2
1
0
AMO1
NCI-H929
RROL
MALAT1
GAPDH
B
RROL
Merged
nuc
cyto
5 μm
C
% of NC
RROL
miR-17
miR-18a
miR-19a
miR-20a
miR-19b
miR-92a
24
36
72
144
Hours after g1 treatment
120
100
80
60
40
20
0
D
Relative RNA expression (ASO1/NC)
Daudi
Raji
RROL
ACC1
ANO6
EXT1
FER
CCDC91
E
Relative RNA expression
nc
m-aso1
m-aso2
m-aso3
rrol
acc1
fer
F
RNA level % (anti-miRs compared to anti-NC )
160
140
120
100
80
60
40
20
0
NS
miR-17
miR-18a
miR-19a
miR-20a
miR-19b
miR-92a
ACC1
AN06
CCDC91
EPT1
EXT1
KIAA1109
G
RNA level % (miR mimics compared to miR-NC )
200
150
100
50
0
NS
miR-17
miR-18a
miR-19a
miR-20a
miR-19b
miR-92a
ACC1
AN06
CCDC91
EPT1
EXT1
KIAA1109
H
GAPDH ex2 (% of positive reactions)
NS
100
75
50
25
0
LacZ
ps1
ps2
RROL
I
RNA amplification (CT Values)
40
30
20
10
0
NS
LacZ
ps1
ps2
RROL
GAPDH RNA

**Supplementary Fig. 3. A**) Sub-cellular qRT-PCR analysis of RROL in AMO1 and H929. MALAT1 and GAPDH were used as internal controls for nuclear- and cytosolic- enriched RNAs, respectively. The results are average fold enrichment of nuclear vs cytosolic fraction after 2-ΔCt calculations. **B**) RNA-FISH analysis of subcellular localization of RROL in AMO1. Cell nuclei are stained by DAPI. **C**) qRT-PCR analysis of RROL and miR-17-92 miRNAs in AMO1 gymnotically exposed to ASO1 or ASO-NC for the indicated time. The heatmap shows average RROL or miRNA expression levels after normalization with GAPDH or RNU44, respectively, and ΔΔCt calculations. **D**) qRT-PCR analysis of RROL transcriptional targets in Daudi and Raji cells gymnotically exposed to ASO1 for 24h. The results shown are average mRNA expression levels after normalization with GAPDH and ΔΔCt calculations. **E**) qRT-PCR analysis of murine *rrol*, *acc1* and *fer* in 5TGM1 murine MM cells transfected with 3 different ASOs targeting the murine *mir17hg* nascent RNA. The results shown are average mRNA expression levels after normalization with murine *gapdh* and ΔΔCt calculations. **F-G**) qRT-PCR analysis of miR-17-92 miRNAs (red bars) and RROL transcriptional targets (white bars) in AMO1 transfected with pooled miR-17-92 miRNA inhibitors (F) or mimics (G). The results shown are average miRNA or mRNA expression levels after normalization with RNU44 or GAPDH, respectively, and ΔΔCt calculations. **H**) ChIRP-qPCR analysis showing the % of positive reactions (amplification) of GAPDH exon2 in chromatin purified using 2 RROL antisense probe sets (ps1 and ps2) or using LacZ antisense probes (negative control). **I**) ChIRP-qPCR analysis of RROL or GAPDH RNA in the RNA purified using 2 RROL antisense probe sets (ps1 and ps2) or using LacZ antisense probes (negative control). Results are shown as average CT values. *indicates $p<0.05$ after Student t test in a) c) d) f) h). *NS* indicates $p>0.05$ after Student t test d) and h); or after Fisher Exact Test in g).

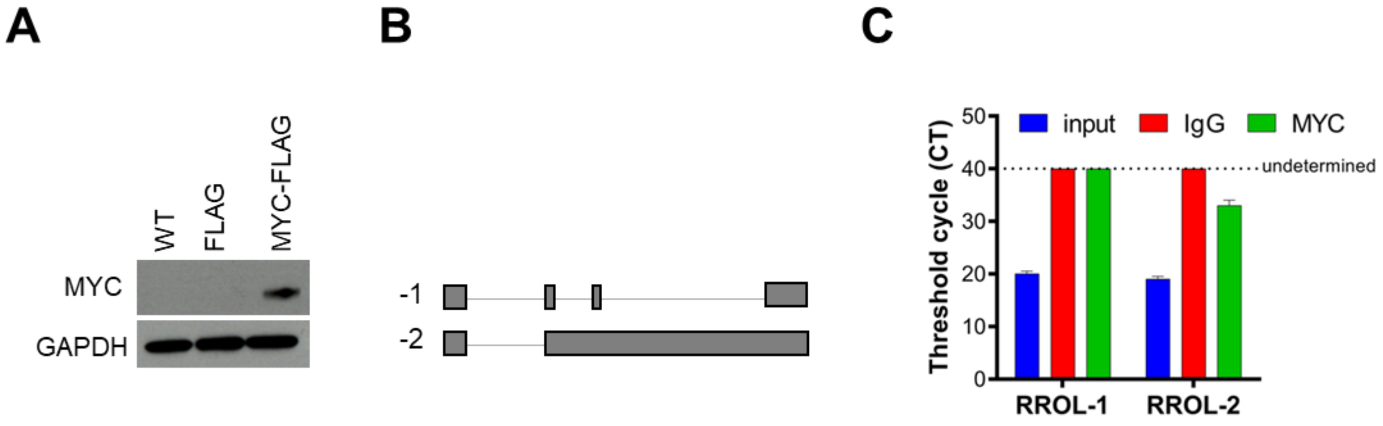


**Supplementary Fig. 4. A**) Western blot analysis of MYC in U266 cells WT or infected with a vector carrying the expression of FLAG or MYC-FLAG. GAPDH was used as protein loading control. **B**) Schema of RROL isoforms considered for this study. *Indicates $p<0.05$ after Student t test calculation. **C**) qRT-PCR analysis of RROL isoforms -1 and 2 in RIP material precipitated using an anti-MYC antibody (α-MYC) or IgG control; or in the 15% input. CT values are shown in each condition.

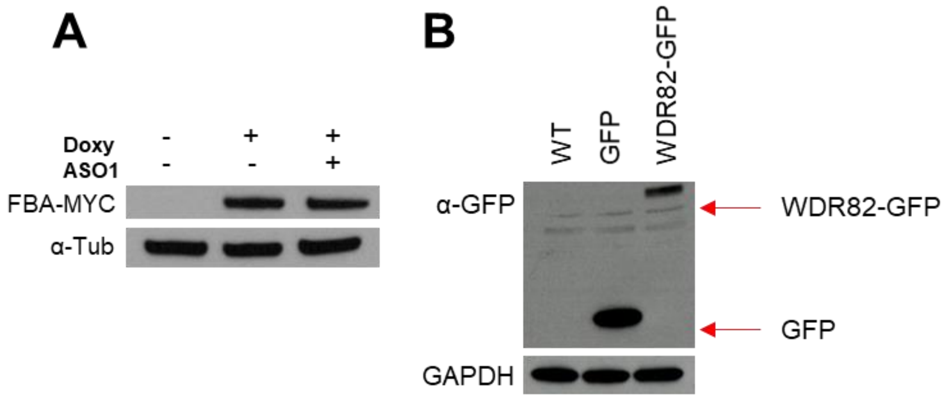


**Supplementary Fig. 5. A**) Western blot analysis of MYC in FBA-MYC cells with or without doxycycline to induce the FBA-MYC fusion protein; and transfected with either NC or ASO1 to knockdown RROL. α-tubulin used as protein loading controls. **B**) Western blot analysis of GFP in AMO1 cells WT or infected with a vector carrying the expression of GFP or WDR82-GFP. GAPDH was used as protein loading control. *Indicates $p<0.05$ after Student t test calculation. Cell proliferation is calculated compared to cells transfected with vehicle (0 nM).

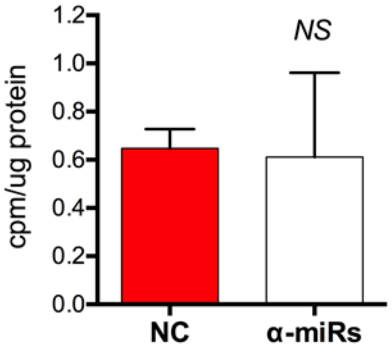


**Supplementary Fig. 6. A**) Incorporation of $C^{14}$-glucose into lipids, 48h after transfection of AMO1 with miR-17-92 anti-miRs (25nM each). Results are expressed as percentage of uninduced cells. *NS* indicates $p > 0.05$ after Student t test.

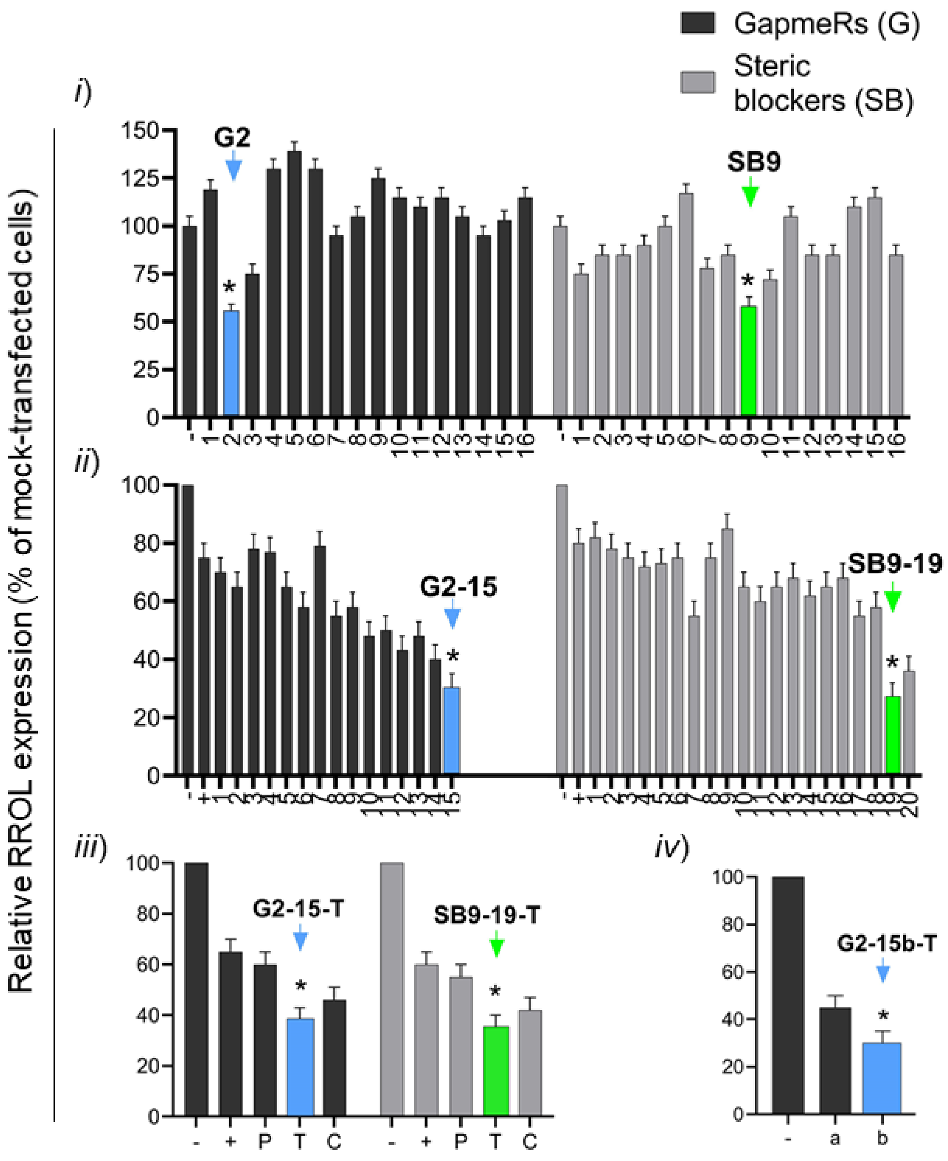


**Supplementary Fig. 7. A**) Multi-step screen to develop therapeutic ASOs targeting RROL. *In step1*, to identify ASO-accessible stretches on RROL, we tested 16 sequences (>20-mer) either in "G" or "SB" configuration for a total of 32 ASOs; based on KD activity in AMO1 cells assessed by qRT-PCR, we selected for further investigation the sequences G2 (21-mer) and SB9 (22-mer). *In step 2*, to optimize the G2 and SB9 designs, we fine-tuned these sequences to obtain 20-mer (n=8) and 18-mer (n=5) derivative ASOs for G2, and 22-mer (7), 20-mer (n=8) and 18-mer (n=5) derivative ASOs for SB9; based on KD activity in AMO1 cells assessed by qRT-PCR, we selected for further investigation the 18-mer derivative sequences G2-15 and SB9-19. *In step 3*, we tested these 2 molecules conjugated with palmitic acid (P) or cholesterol (C) or tocopherol (T). Based on KD activity in AMO1 cells assessed by qRT-PCR, we selected the TO-conjugated molecule as the leading compounds. The G2-15-T ASO was further optimized by replacing the 10-mer "core" DNA gap with a 8-mer "core" DNA gap (G2-15b-T).
