## Supplementary Tables 1-7 for "Long Noncoding RNA RROL Provides Chromatin Scaffold for MYC-WDR82 Interaction to Impact Lipid Metabolism and Tumor Cell Growth in Multiple Myeloma"

| Seq# | Gene Symbol | sgRNA |
|---|---|---|
| 1 | Non_Targeting_Human | CTGCATGGGGCGCGAATCA |
| 2 | Non_Targeting_Human | TGCACCCGGCTAGGACCGG |
| 3 | Non_Targeting_Human | TGTGCAACCTCCGCCGTTG |
| 4 | Non_Targeting_Human | CCCGAGCAGTGGCTCGCTA |
| 5 | Non_Targeting_Human | GTCAGGTAGAGGGATTGAG |
| 6 | Non_Targeting_Human | GAGGACGATCGTACTCCAG |
| 7 | Non_Targeting_Human | TATGGAGGGCTGGATCTGC |
| 8 | Non_Targeting_Human | CCTTGGCTAAACCGCTCCC |
| 9 | Non_Targeting_Human | GAGTCGGGTAAATAGACAA |
| 10 | Non_Targeting_Human | AGTGGACGACCCTGTCAGG |
| 11 | Non_Targeting_Human | TATCGGTCTGTACGGGTAG |
| 12 | Non_Targeting_Human | TGGTACGGGGCCGGCCTGT |
| 13 | Non_Targeting_Human | CAGTCGCCGCGCATCGGGC |
| 14 | Non_Targeting_Human | AGAGGTCGGATACGCTTCC |
| 15 | Non_Targeting_Human | GCCCTACATCGCTAGGCTA |
| 16 | Non_Targeting_Human | CGCCGGTTCCGTCGACTAG |
| 17 | Non_Targeting_Human | AGCAAGGTTCGGCAAAGCG |
| 18 | Non_Targeting_Human | CCAGTACTGACAGCCTTAT |
| 19 | Non_Targeting_Human | GTCGCCGTAGGGTCCGCCA |
| 20 | Non_Targeting_Human | CTAGAATGGTTGCGCGAAC |
| 21 | Non_Targeting_Human | AGCCTAACGATCGGACCGA |
| 22 | Non_Targeting_Human | CCGCTCCATCAGACTTTCT |
| 23 | Non_Targeting_Human | CTACGCCCGGGGGAAAAGA |
| 24 | Non_Targeting_Human | CAGGAGCTAAGGGTCCCGT |
| 25 | Non_Targeting_Human | TACTCCCCGAATGCAGCGG |
| 26 | Non_Targeting_Human | AGGCCGCAAACCGCGTGCG |
| 27 | Non_Targeting_Human | AGCCTCTTGGTTCCATGGA |
| 28 | Non_Targeting_Human | CTCCCCCGCTTCACTTGGG |
| 29 | Non_Targeting_Human | GTGGCCTTTGCAATTGGCG |
| 30 | Non_Targeting_Human | CGCTAGGAAACCAACAAGG |
| 31 | Non_Targeting_Human | GGTGGAGTGCAGCCACAAA |
| 32 | Non_Targeting_Human | TTCCGATGGTTAAGCACGA |
| 33 | Non_Targeting_Human | TGCGTACCACACGACACAT |
| 34 | Non_Targeting_Human | AGAACTCCCGGCAATTAGG |
| 35 | Non_Targeting_Human | TCATTCACGTTGGGTTTTC |
| 36 | Non_Targeting_Human | ATCTCCGGGGCGTTAGGCC |
| 37 | Non_Targeting_Human | CAGCGAGCAGCTTTTACGA |
| 38 | Non_Targeting_Human | ACGTGTGTTACGGGGGGAA |
| 39 | Non_Targeting_Human | GAGAGGAAGTCGTCGGAAA |
| 40 | Non_Targeting_Human | CGTACGACAATACGCGCGA |
| 41 | Non_Targeting_Human | GACTGGAGAGTGCCCGGCC |
| 42 | Non_Targeting_Human | CAATTCCCCGCAATCCAGC |

| | | |
|---|---|---|
| 43 | Non_Targeting_Human | TCAAAAGCAACGTACGCAA |
| 44 | Non_Targeting_Human | TACCAGCACGGAGCGGCAG |
| 45 | Non_Targeting_Human | CAGGCAGCCGCTGAAGCAT |
| 46 | Non_Targeting_Human | AGGGGTCGTCGGGCAACCC |
| 47 | Non_Targeting_Human | TCACGGGCGTGCTCGAGTC |
| 48 | Non_Targeting_Human | TACATTGCTCTGGGGCGTA |
| 49 | Non_Targeting_Human | GTGGTGACAAGTCCTGAGG |
| 50 | Non_Targeting_Human | ACACAGGCCGTTAAAAAGA |
| 51 | Non_Targeting_Human | GTGATCAAATGCCTATGAT |
| 52 | Non_Targeting_Human | GAGCTAGCGGAATGCATGG |
| 53 | Non_Targeting_Human | GTGTGACCTCGTCTGGCAG |
| 54 | Non_Targeting_Human | CGTGATGGGCACGAGTAAG |
| 55 | Non_Targeting_Human | GGCCTGGACGAGCCTAAAA |
| 56 | Non_Targeting_Human | GATCGCGTCTAGTGCAGAT |
| 57 | Non_Targeting_Human | AACCTGAGACGCTAAGTGA |
| 58 | Non_Targeting_Human | CCCCCTACGACAGGTACAT |
| 59 | Non_Targeting_Human | CGGCGAGGGAGAATGCCCT |
| 60 | Non_Targeting_Human | CCACAGGGAAGCGAATACG |
| 61 | Non_Targeting_Human | GGGAAGCGTCGCGCCGGAA |
| 62 | Non_Targeting_Human | ATGTCGTAGGAGAAGGACG |
| 63 | Non_Targeting_Human | CCCAAAGGGCGGCAAGCAG |
| 64 | Non_Targeting_Human | TGAGACACCGTACACGATT |
| 65 | Non_Targeting_Human | CGCCCTGGGCCCCACCAGC |
| 66 | Non_Targeting_Human | GCGGCGACGCACGCTACCT |
| 67 | Non_Targeting_Human | GCAGAGGCAAGGGCATAAA |
| 68 | Non_Targeting_Human | GCTCATGCGCGCGAACGCC |
| 69 | Non_Targeting_Human | CCTCCCCCATGCTGAGTAA |
| 70 | Non_Targeting_Human | TGTTTAGCCGCGGGCGGAC |
| 71 | Non_Targeting_Human | GCCTCGGGCGTACATTAAG |
| 72 | Non_Targeting_Human | TCAGCGGGCCGCATCTTGT |
| 73 | Non_Targeting_Human | GGCGCTTTGAGACGATGGG |
| 74 | Non_Targeting_Human | CGGCTCCATGGGCGGCCGT |
| 75 | Non_Targeting_Human | GGTGGGTCACGCGGAAGGG |
| 76 | Non_Targeting_Human | GATCGCTTCAATCACCGAG |
| 77 | Non_Targeting_Human | GAAGGTGCTGAATGTGCCG |
| 78 | Non_Targeting_Human | GAAACGACGGAGTATTCCG |
| 79 | Non_Targeting_Human | AAGCACCCATCGCCGGCTG |
| 80 | Non_Targeting_Human | GTCTAGTATCTTGAGCGTT |
| 81 | Non_Targeting_Human | TGCCACTCACGAGTTGACT |
| 82 | Non_Targeting_Human | TCCCGTTGAGAGAGGACAC |
| 83 | Non_Targeting_Human | ACGTTGAGGGACTGTCCTA |
| 84 | Non_Targeting_Human | ATCCTGACGGGGACTACAT |
| 85 | Non_Targeting_Human | GGTTGCGCGGGGTGGCGTA |

| | | |
|---|---|---|
| 86 | Non_Targeting_Human | CATCACCCGATGTACCGAG |
| 87 | Non_Targeting_Human | TACCGATTCCCGGCCAAGA |
| 88 | Non_Targeting_Human | ATAACCACAAGGCGCTCGA |
| 89 | Non_Targeting_Human | AGGACCCAGTAGACCGGCT |
| 90 | Non_Targeting_Human | GGGTGAGGGTCCAATTCGG |
| 91 | Non_Targeting_Human | TAGCTATGCGGTAGGACGG |
| 92 | Non_Targeting_Human | TAGACGTGCAAAAGCCGCA |
| 93 | Non_Targeting_Human | GATTAACCCAGGGCTTTGA |
| 94 | Non_Targeting_Human | TGCTAATGGGGAAAGCCGG |
| 95 | Non_Targeting_Human | CGATGGGACAGACGCCGGA |
| 96 | Non_Targeting_Human | CCGGGGCCTTGGAAAACAC |
| 97 | Non_Targeting_Human | TGACAGGATGCGTTCACGG |
| 98 | Non_Targeting_Human | CGACATCGGTTACATGTGG |
| 99 | Non_Targeting_Human | GTCGATGTGGGGACGGCAC |
| 100 | Non_Targeting_Human | TGCGCAAACCGGATCGGGG |
| 101 | Non_Targeting_Human | GCAAGATAGACGCTACACA |
| 102 | Non_Targeting_Human | CGACAATTGCAGCTGGACC |
| 103 | Non_Targeting_Human | ATCCGTCACTGACCCTTCC |
| 104 | Non_Targeting_Human | CGCAGCGGTTGTCAAACGA |
| 105 | Non_Targeting_Human | AGACTCTAGGTCCAGGAGT |
| 106 | Non_Targeting_Human | GGGAAAGAGAATCGCCCTA |
| 107 | Non_Targeting_Human | AAACTCCGCAGGCCAAGTA |
| 108 | Non_Targeting_Human | GGCAGGGCTACGATGCAGG |
| 109 | Non_Targeting_Human | GCCGGGTATTCCTGAGGGG |
| 110 | Non_Targeting_Human | TCGGTTGGACAATTTCCCG |
| 111 | Non_Targeting_Human | GCGCGTGGTTGCATTAGAC |
| 112 | Non_Targeting_Human | TAGACAGCAGGGGCGTTTG |
| 113 | Non_Targeting_Human | GTAAGGAAAAACGGCGGTG |
| 114 | Non_Targeting_Human | TTCAGGGTGGCCGACAGGC |
| 115 | Non_Targeting_Human | ATGAGTTCGCGAGGCCGTA |
| 116 | Non_Targeting_Human | CTTCTAGGAGGCACTACGG |
| 117 | Non_Targeting_Human | CAGGAGCTCAACGACGGTG |
| 118 | Non_Targeting_Human | AGCTCTTTGACGGACTGTG |
| 119 | Non_Targeting_Human | GTATCGGGTGGCGTTAATT |
| 120 | Non_Targeting_Human | GCGCCACGAGGATCAAAGC |
| 121 | Non_Targeting_Human | CGATCTGGTTGCACAACTA |
| 122 | Non_Targeting_Human | AGGGCCCGCTCGTCCTCAG |
| 123 | Non_Targeting_Human | GAGATCAGGATGATCGTAC |
| 124 | Non_Targeting_Human | GGGGGGACCCCGATCTCCG |
| 125 | Non_Targeting_Human | GCTAGAGTGGCGCAGGGAG |
| 126 | Non_Targeting_Human | GGCGGGTCGGACATCGCCT |
| 127 | Non_Targeting_Human | TTCTACACATTGGAAATAA |
| 128 | Non_Targeting_Human | GGCTAATCCCGGTGAAAAA |

| | | |
|---|---|---|
| 129 | Non_Targeting_Human | CGCCAGACCGGCCAGCGAG |
| 130 | Non_Targeting_Human | TCGGTTCGAGCGAAACGTG |
| 131 | Non_Targeting_Human | GAGAGTCGGGAGGGATCGT |
| 132 | Non_Targeting_Human | CGCGTCGTTACAGGGGCGG |
| 133 | Non_Targeting_Human | CACGCCACGAGACGTTGGA |
| 134 | Non_Targeting_Human | CGAGGAGCAGTTCAACACC |
| 135 | Non_Targeting_Human | TGACGTTGTGAGGACGGTC |
| 136 | Non_Targeting_Human | CCCACCCGAACGGAGGCGG |
| 137 | Non_Targeting_Human | GCCGGGCGCATATCCTCAC |
| 138 | Non_Targeting_Human | GCCATGCGGAAGATGTGGA |
| 139 | Non_Targeting_Human | ACTGTTTGCGCGTGTAGAG |
| 140 | Non_Targeting_Human | CACACAGGCCTAGCGATCT |
| 141 | Non_Targeting_Human | CCCTGGCAAACGAAGGGGG |
| 142 | Non_Targeting_Human | AGGCGTCAGTGAGCGGTGT |
| 143 | Non_Targeting_Human | TGTAACTGCCACTCCTAGG |
| 144 | Non_Targeting_Human | CCCGGGGGGGGTCGTAGGT |
| 145 | Non_Targeting_Human | TGTCCAGTTCCGTAGGATG |
| 146 | Non_Targeting_Human | CCTTGCCGCGACGAAAACT |
| 147 | Non_Targeting_Human | GACCTGAGTCGTGACACAG |
| 148 | Non_Targeting_Human | AAGCACTATACGGACGGAG |
| 149 | Non_Targeting_Human | GTATGGAGGATCTTCGTGG |
| 150 | Non_Targeting_Human | CGCGCTGACTCGTTACGAG |
| 151 | Non_Targeting_Human | CGGGCGCTCTATCTTGCAG |
| 152 | Non_Targeting_Human | CGATTTGTCCCGGCAAGGG |
| 153 | Non_Targeting_Human | CGTATACCGGGTCCAAGAG |
| 154 | Non_Targeting_Human | CCTCATATGCTACTTTATG |
| 155 | Non_Targeting_Human | CCCCTCGTTGAAGCCGGCT |
| 156 | Non_Targeting_Human | GACTACTCGTGGCGCTTGC |
| 157 | Non_Targeting_Human | GCGGCGGAGGCCCCGCCAA |
| 158 | Non_Targeting_Human | GGCCTCATATAGAACACTG |
| 159 | Non_Targeting_Human | CTCGTAAGGAGGATATGCG |
| 160 | Non_Targeting_Human | ATGTTGTATCAGCCAGTAC |
| 161 | Non_Targeting_Human | TGGACGAGACGGCGGACGG |
| 162 | Non_Targeting_Human | CGGGCGTTAACGCGTGCAA |
| 163 | Non_Targeting_Human | TGGGCGGCTCGGAGAACTG |
| 164 | Non_Targeting_Human | CGATGCGCCGACCCGTCAA |
| 165 | Non_Targeting_Human | AGTCACGCGGCTCCCATCG |
| 166 | Non_Targeting_Human | GACGGGGACCCGGTTGGCA |
| 167 | Non_Targeting_Human | AGCCCCGCCTGGGTACGCG |
| 168 | Non_Targeting_Human | CCCGATGCCGCAGTCCATA |
| 169 | Non_Targeting_Human | AAAGCGGGATAACTTGAGG |
| 170 | Non_Targeting_Human | GCGCCGGTTAACAATAACT |
| 171 | Non_Targeting_Human | GATGGTTTCGCGTACATAG |

| | | |
|---|---|---|
| 172 | Non_Targeting_Human | CCTGCGCGACAAGCGGACG |
| 173 | Non_Targeting_Human | CAGGGGCCTGCGCCAGCGG |
| 174 | Non_Targeting_Human | GCAGGTACAACGAGCGTCA |
| 175 | Non_Targeting_Human | GCGCCCCAGTCGGCCCCTA |
| 176 | Non_Targeting_Human | ACAATACTGTCGATGGCAT |
| 177 | Non_Targeting_Human | GCGTGTCCTGAGCCGTGAA |
| 178 | Non_Targeting_Human | CTACGCCGACACAAAGATG |
| 179 | Non_Targeting_Human | GACCCTGACATCAAGACAG |
| 180 | Non_Targeting_Human | CCATGCCTCGTACAGGAGA |
| 181 | Non_Targeting_Human | CTATTAAGGTTATGGCCGA |
| 182 | Non_Targeting_Human | TATGGCTAAGGGCGGGCGT |
| 183 | Non_Targeting_Human | TACCGCACGAAATGCTCCG |
| 184 | Non_Targeting_Human | GGGGAATCTGCAAGACCAT |
| 185 | Non_Targeting_Human | TCGGAGTAATGACCACAGC |
| 186 | Non_Targeting_Human | GGAGCTCCGGATTGCCGAA |
| 187 | Non_Targeting_Human | CCACCCGGTGGGTGACGGG |
| 188 | Non_Targeting_Human | AGACGTGGCACGTCTCGAC |
| 189 | Non_Targeting_Human | AAAAATGAGCCATGGGCGG |
| 190 | Non_Targeting_Human | ACCGGTCGGCGGACGTGGG |
| 191 | Non_Targeting_Human | TGCGCGCCGCCCAATTCTT |
| 192 | Non_Targeting_Human | TGTGGCTCGGCCAGCGATA |
| 193 | Non_Targeting_Human | TGTACGGCACCCGTGGCAA |
| 194 | Non_Targeting_Human | GGGCGGGGGCCGGGCGGAT |
| 195 | Non_Targeting_Human | TGCCAGCTTGTGGTGTCGT |
| 196 | Non_Targeting_Human | GCCGTAACGGTGTCACACT |
| 197 | Non_Targeting_Human | GAGATGTCCAGGCGTACAG |
| 198 | Non_Targeting_Human | CGTGGGCTTACTAAGCAGC |
| 199 | Non_Targeting_Human | TGCACACGGCAGTAGAAAA |
| 200 | Non_Targeting_Human | GAGTTGCAGCCGTCACGAG |
| 201 | Non_Targeting_Human | ACCGCCGCGCGGCCTAACG |
| 202 | Non_Targeting_Human | TCAGCGGAGATGTGCACTG |
| 203 | Non_Targeting_Human | CTACCCCTGATTGCCAGCC |
| 204 | Non_Targeting_Human | GAGGCCTGTACGCGGAGGC |
| 205 | Non_Targeting_Human | GTCGGGCGCATCACGCGGG |
| 206 | Non_Targeting_Human | TTCGCACCGGCCACCACAC |
| 207 | Non_Targeting_Human | ATGGCATACACTTCGCGAG |
| 208 | Non_Targeting_Human | GCCTGGCCCTCGGCAACTG |
| 209 | Non_Targeting_Human | GGAGCCAAGCCGATGCACA |
| 210 | Non_Targeting_Human | CGCCCGGAAACGTTAACCT |
| 211 | Non_Targeting_Human | GTTCGCTTACACGCCGTCC |
| 212 | Non_Targeting_Human | ACACTGAGGGCCGCAGCCG |
| 213 | Non_Targeting_Human | AACAGTTTAAGGAGCACCG |
| 214 | Non_Targeting_Human | CAACAAGCGCGCGTAGGTA |

| | | |
|---|---|---|
| 215 | Non_Targeting_Human | AGCGCCGACAAGCTGACGC |
| 216 | Non_Targeting_Human | TCCGCCGAGTCTAAACTTC |
| 217 | Non_Targeting_Human | GCGTTGGCTTAAGCTGTAG |
| 218 | Non_Targeting_Human | TGAACTCAAAAATCCCGAC |
| 219 | Non_Targeting_Human | GCCTGGCGTGAAACGGCCG |
| 220 | Non_Targeting_Human | TGACAGCCCCGTAACTGAG |
| 221 | Non_Targeting_Human | TCAAACGCGTGCACCTTGA |
| 222 | Non_Targeting_Human | ATTGACCGGACGGTGTCGC |
| 223 | Non_Targeting_Human | GCGGAACCGACGGTCACTG |
| 224 | Non_Targeting_Human | AGCAGAACGGATGCGCGTC |
| 225 | Non_Targeting_Human | ACGAAGGGCAAAATTCCAG |
| 226 | Non_Targeting_Human | AAGCATAATGGACGGAAGT |
| 227 | Non_Targeting_Human | GAGGGGCGAGCCGTCGCCG |
| 228 | Non_Targeting_Human | CGTGACATGCCGATCCCTG |
| 229 | Non_Targeting_Human | GGAGTGCCAGGCCGAGAGG |
| 230 | Non_Targeting_Human | GTCCAGCTAATATTGCAGG |
| 231 | Non_Targeting_Human | CCCAGCGAAGCGGGGACCT |
| 232 | Non_Targeting_Human | TCGTCTGGAGGTGCGACGG |
| 233 | Non_Targeting_Human | AACGGTATAGCGGCATGCG |
| 234 | Non_Targeting_Human | GGGCCCCGGAGGCAGAACG |
| 235 | Non_Targeting_Human | GAGCCCCGACGCGCGCGTC |
| 236 | Non_Targeting_Human | TTACCTCAGATATGCGGGG |
| 237 | Non_Targeting_Human | ACTTGCGACCCCGAAGAAT |
| 238 | Non_Targeting_Human | ACGCCCCATGCTGAAACCG |
| 239 | Non_Targeting_Human | TGCGGGCCCGCTTCTGCAG |
| 240 | Non_Targeting_Human | ATTGTCACTTTAGATCTGT |
| 241 | Non_Targeting_Human | GCAGCGACCGGCATGCCGC |
| 242 | Non_Targeting_Human | CGAGCGGTTGCCGCCCTGG |
| 243 | Non_Targeting_Human | TTGAATTACAGTTCGACGG |
| 244 | Non_Targeting_Human | TATGGGCGCAAGGGTACAG |
| 245 | Non_Targeting_Human | GCGCTTGGTGCGCGACACG |
| 246 | Non_Targeting_Human | CGGGCAGCGATAGTGGCTG |
| 247 | Non_Targeting_Human | CGGCATTGACATAGGGTCA |
| 248 | Non_Targeting_Human | CGGTGGGCTCCCGGACCGC |
| 249 | Non_Targeting_Human | AACAAGTAACCCAGGCGCC |
| 250 | Non_Targeting_Human | CGGGTGGACCATGACTACG |
| 251 | Non_Targeting_Human | CGAGCCTAGAACCCCGCTC |
| 252 | Non_Targeting_Human | ACCACGCCGGCCCCGTTGA |
| 253 | Non_Targeting_Human | CGACACCGGTAAAGGCACG |
| 254 | Non_Targeting_Human | CCCCCCAGGAGGGCATTTT |
| 255 | Non_Targeting_Human | TACTTGGGGGAGTCGGTGA |
| 256 | Non_Targeting_Human | TCGGGTAACAAGAGGAATC |
| 257 | Non_Targeting_Human | TTTGACGCGCGTCATCCAG |

| | | |
|---|---|---|
| 258 | Non_Targeting_Human | GTAGCAGTTACAGGCTCGC |
| 259 | Non_Targeting_Human | CGTAGCGCCCTCACAGTGG |
| 260 | Non_Targeting_Human | AAGGATGTGTCCCGAATTG |
| 261 | Non_Targeting_Human | TGGGATCGTGCGCGCCCCA |
| 262 | Non_Targeting_Human | GCTATACCACGTTCAGTCA |
| 263 | Non_Targeting_Human | TGCCCGCACGAAAGTGGGT |
| 264 | Non_Targeting_Human | TATGGCGGGGATGCACGAA |
| 265 | Non_Targeting_Human | TCGGTAAGCGACCCCATAG |
| 266 | Non_Targeting_Human | TTAGCGCGCTAGTATATAG |
| 267 | Non_Targeting_Human | CGGACGAACCAGGCTAAAG |
| 268 | Non_Targeting_Human | AGCATCATTGATCTTCCGG |
| 269 | Non_Targeting_Human | CTAGGAGTCGGATCTACTG |
| 270 | Non_Targeting_Human | CACACTCCGCGGGGAAGTG |
| 271 | Non_Targeting_Human | ATACGCTACGTAGCGAAAA |
| 272 | Non_Targeting_Human | GCGCGATCATGTGATAAGT |
| 273 | Non_Targeting_Human | AACGTCGCGGGCGCCCGGA |
| 274 | Non_Targeting_Human | AATCCCCTACTGGGATCAC |
| 275 | Non_Targeting_Human | CCCACGGAGTAGCGCACCG |
| 276 | Non_Targeting_Human | CAAAACTCAGCTACCAGGT |
| 277 | Non_Targeting_Human | GCCGGCCCGGACCCCTAGT |
| 278 | Non_Targeting_Human | GGTGCCCACAGCGCCGTCA |
| 279 | Non_Targeting_Human | GTACTGCGAGGCCGTTATG |
| 280 | Non_Targeting_Human | TAATCGAATCAGGGCTCAC |
| 281 | Non_Targeting_Human | TGTTAACACCCCAGCCACT |
| 282 | Non_Targeting_Human | GATCGTAGACAATAAGCGG |
| 283 | Non_Targeting_Human | GGCCGACGGATAGGAGGGA |
| 284 | Non_Targeting_Human | CACGGACGCGATGACACGC |
| 285 | Non_Targeting_Human | TCAGGAAGAGCGCGGGAGT |
| 286 | Non_Targeting_Human | TTTTGCCCGCGGCGTAACG |
| 287 | Non_Targeting_Human | GTGCCTCACTTGTTCGACG |
| 288 | Non_Targeting_Human | GTACAAGTGCCCACAAGTG |
| 289 | Non_Targeting_Human | CCCCTGACCGGCACCCTCG |
| 290 | Non_Targeting_Human | CCGCGCCCTGGGGTCGAGA |
| 291 | Non_Targeting_Human | CGCGCTGAGGTACATGGGG |
| 292 | Non_Targeting_Human | GCGAACGTCGACCAGCCCT |
| 293 | Non_Targeting_Human | AGCTTCCCGGGTCGACCGC |
| 294 | Non_Targeting_Human | TTAGCCGGACAAGAACGAT |
| 295 | Non_Targeting_Human | TCAGAGAGATAAGCTATCG |
| 296 | Non_Targeting_Human | CGTGGGCGTGCGGACTCAG |
| 297 | Non_Targeting_Human | GTCGCCGTGCGCCTACCGA |
| 298 | Non_Targeting_Human | CGGACTAGGCGTAGCGGGT |
| 299 | Non_Targeting_Human | CGAGTATGTCGCCTTGCAG |
| 300 | Non_Targeting_Human | GACGGAACGCGGACAGCAA |

| | | |
|---|---|---|
| 301 | Non_Targeting_Human | AAGTAGCTGAGTCATCACG |
| 302 | Non_Targeting_Human | CTCCGACCCCCGAGATGAG |
| 303 | Non_Targeting_Human | GGGCGAGACGACCGAGCAA |
| 304 | Non_Targeting_Human | GACCGATCGAGTATACCGT |
| 305 | Non_Targeting_Human | ATTGTTCAGGCGAGGGTGT |
| 306 | Non_Targeting_Human | ACGGCGCGTGCGCTTAGTG |
| 307 | Non_Targeting_Human | TATCCTCTACCTGCTACGT |
| 308 | Non_Targeting_Human | CGACAGCGGCTCGCCTGCG |
| 309 | Non_Targeting_Human | CTCCCCGCAGGGTTAGTCG |
| 310 | Non_Targeting_Human | CAGCATTTGAACACGGAAG |
| 311 | Non_Targeting_Human | CAAAAACTTGTGCAGCTAG |
| 312 | Non_Targeting_Human | CTTACCACCGACGACCAAT |
| 313 | Non_Targeting_Human | TGCGGTGCCGAGCCCTATG |
| 314 | Non_Targeting_Human | AACGGGGGGGCCCTTCGG |
| 315 | Non_Targeting_Human | CTTGTAACCTGGGACGGCT |
| 316 | Non_Targeting_Human | ATCGACAGTCTAGCGGCTG |
| 317 | Non_Targeting_Human | GGTCGACCACCGCCCTCTG |
| 318 | Non_Targeting_Human | GCGCAGACGCCGACCAGAG |
| 319 | Non_Targeting_Human | TGAGGCAGTCTGAGTGCGA |
| 320 | Non_Targeting_Human | GAGGATACCCGGGGAACCT |
| 321 | Non_Targeting_Human | GGCACGTTCGCGAAAACGA |
| 322 | Non_Targeting_Human | ACAAGTCTAGGGCCTCCCC |
| 323 | Non_Targeting_Human | AGAGGGCATGGCGCATGCA |
| 324 | Non_Targeting_Human | CTGGGCCTGGCAAATTGAA |
| 325 | Non_Targeting_Human | GTCAGGGGTTTGGTCGGCA |
| 326 | Non_Targeting_Human | CCGCGTTTTACGCGACCGA |
| 327 | Non_Targeting_Human | GTTCTCGCGGCTGTGCGGT |
| 328 | Non_Targeting_Human | GCAGGGCCCAGTCGTGCAG |
| 329 | Non_Targeting_Human | TCGGTACGGCGGTTGATCG |
| 330 | Non_Targeting_Human | ACTGGCCACCGGCACCTGG |
| 331 | Non_Targeting_Human | GCCCCGGACGTCCTGACCC |
| 332 | Non_Targeting_Human | AAAGGATGGTGATATCTCG |
| 333 | Non_Targeting_Human | AGGGAAAATTGACGTTCGG |
| 334 | Non_Targeting_Human | GGAACGACTATGACCGCCA |
| 335 | Non_Targeting_Human | CCTGGCGAACCCGACGTCT |
| 336 | Non_Targeting_Human | ACAGAGGAAGGTGCCGTAT |
| 337 | Non_Targeting_Human | TATTCGTACCGGGCAGCAG |
| 338 | Non_Targeting_Human | GCCAGATCCCGCCCAGCAG |
| 339 | Non_Targeting_Human | GTGATGGGGCACGACATCG |
| 340 | Non_Targeting_Human | GGAGCCGGAAGGCCGATGG |
| 341 | Non_Targeting_Human | CCCGCGCTGCGGGACGCAG |
| 342 | Non_Targeting_Human | GCTAAAAAACCCCTTCCCC |
| 343 | Non_Targeting_Human | CGAATGAGGGGGACGGACT |

| | | |
|---|---|---|
| 344 | Non_Targeting_Human | CAGCAGGCGCGATGACCCT |
| 345 | Non_Targeting_Human | ACAGGATCAGTGAGCTAAA |
| 346 | Non_Targeting_Human | ATCGCGCGGCACGCGTCAG |
| 347 | Non_Targeting_Human | TTGGTCGGTCACCACTGGG |
| 348 | Non_Targeting_Human | GCTTCTCCAGTCAACGGTT |
| 349 | Non_Targeting_Human | GGCGCATGACAGCCAGGCG |
| 350 | Non_Targeting_Human | AGTCGGTGAACGAGGCCGG |
| 351 | Non_Targeting_Human | TCGGGAGCATACCAAGTGT |
| 352 | Non_Targeting_Human | GCGAGGCGGGAATTCACCA |
| 353 | Non_Targeting_Human | GGCAGACTGATGAAGACGT |
| 354 | Non_Targeting_Human | GGTAGGCCCCGAGTCGCCC |
| 355 | Non_Targeting_Human | GTACTCGCCGGAACGCGAC |
| 356 | Non_Targeting_Human | TATGAGAAATACGCGCGGC |
| 357 | Non_Targeting_Human | GGATCAGGAGGGACGTCGC |
| 358 | Non_Targeting_Human | TCGAGGACGCACCCCATAA |
| 359 | Non_Targeting_Human | ACGGTCGTTGAATCGGCTA |
| 360 | Non_Targeting_Human | CCTACCTCGATGAAGCGTG |
| 361 | Non_Targeting_Human | GTACCAGCGGGGTTTTAGA |
| 362 | Non_Targeting_Human | ACCGCGGGCGTAAGCCCCC |
| 363 | Non_Targeting_Human | GGAAGAGCGAGATCGGGTT |
| 364 | Non_Targeting_Human | CTTCGGCCTCCTAGTGCAG |
| 365 | Non_Targeting_Human | TGAATGAGTGCCCACCCGG |
| 366 | Non_Targeting_Human | CTTAGACCCTCCTGGACTC |
| 367 | Non_Targeting_Human | GGCGTAACGCGGTGGCACA |
| 368 | Non_Targeting_Human | GCGAAGTGCAACCGATGGG |
| 369 | Non_Targeting_Human | GCGGTGCGGGGGATCGAAG |
| 370 | Non_Targeting_Human | CGCTGGTAAAGGCGGAAAG |
| 371 | Non_Targeting_Human | CGAACGTTCAGGATTCCAC |
| 372 | Non_Targeting_Human | TCGAAGTACGGTGTCCCTA |
| 373 | Non_Targeting_Human | TAAACCACGGGCAACGTCT |
| 374 | Non_Targeting_Human | GTAGTGCGCTTCCCCGAAG |
| 375 | Non_Targeting_Human | GGGCAGACCAGGCTCGAGC |
| 376 | Non_Targeting_Human | TCCCCGCGACGGGAATGAA |
| 377 | Non_Targeting_Human | GAATCCATTCGGGCTGCGA |
| 378 | Non_Targeting_Human | AACGACTTGCGTTCATGCC |
| 379 | Non_Targeting_Human | AATATCAAAGGTCTTACGG |
| 380 | Non_Targeting_Human | GGGGTGGGCGCTCTCTCCA |
| 381 | Non_Targeting_Human | CGCGATCAGTAGGACAGTT |
| 382 | Non_Targeting_Human | CGGTGGGGCCGCTCACTAG |
| 383 | Non_Targeting_Human | AGCTGGCACCTCTTAGCGA |
| 384 | Non_Targeting_Human | GTCCCGCATGGGACGGTGG |
| 385 | Non_Targeting_Human | CGACACACGCGACTTCAGA |
| 386 | Non_Targeting_Human | CCCGCATTCAGCACCAGGC |

| | | |
|---|---|---|
| 387 | Non_Targeting_Human | GAATATCGCCTGTCGACCA |
| 388 | Non_Targeting_Human | GACTAGGGAGGCGACACGG |
| 389 | Non_Targeting_Human | GCTTGTTATACACGTCAGT |
| 390 | Non_Targeting_Human | GGCGAGCTGCCATCGTCGA |
| 391 | Non_Targeting_Human | CGCACTATACTAAAGGCGT |
| 392 | Non_Targeting_Human | CCGAGAAGCAGGCTCCAAC |
| 393 | Non_Targeting_Human | AGGGGTGGCTAAGTCCGGC |
| 394 | Non_Targeting_Human | CCCCTCGATTCCCGAACAG |
| 395 | Non_Targeting_Human | CGACCCAACTCTGTGCGTC |
| 396 | Non_Targeting_Human | TTACGTCGATCGATAAAAT |
| 397 | Non_Targeting_Human | GTGAGGCGCGGTACATGGA |
| 398 | Non_Targeting_Human | TGAACGCATGCGGGAGCGT |
| 399 | Non_Targeting_Human | GAGCCGACGTGAGAACGAT |
| 400 | Non_Targeting_Human | GCCATGAGACGTTATGCAG |
| 401 | Non_Targeting_Human | CACATGAAAGTCCAAAAAT |
| 402 | Non_Targeting_Human | GCGCCGGACTGGACCTCGA |
| 403 | Non_Targeting_Human | GAGACAACAGCCGGCTCCC |
| 404 | Non_Targeting_Human | ATGTGATCGAGGGGCACTC |
| 405 | Non_Targeting_Human | TTGCGCAGTGGCGAAATGG |
| 406 | Non_Targeting_Human | TAGGGTTGCGAGGGGTGCC |
| 407 | Non_Targeting_Human | GCGGGCAGCAGGCGCGAGA |
| 408 | Non_Targeting_Human | GGCGGCGCACCGAGTCGAC |
| 409 | Non_Targeting_Human | GCTTGACCGGCTTCCCAGG |
| 410 | Non_Targeting_Human | AGTAAGATTGCACGATGCG |
| 411 | Non_Targeting_Human | GGGGATCGGAGTCCCCCGG |
| 412 | Non_Targeting_Human | CCACGACGTCGGTACTCGA |
| 413 | Non_Targeting_Human | GTGATCATCGCACTGGCAG |
| 414 | Non_Targeting_Human | CAGCGAACACGGACGTCTG |
| 415 | Non_Targeting_Human | TCGCCTCGGAGCAAACCGT |
| 416 | Non_Targeting_Human | ACCCCCGCCGAGAGTTGGT |
| 417 | Non_Targeting_Human | CAGTATTTCCCTTCAAGCG |
| 418 | Non_Targeting_Human | AACCTAAAGGGTATAGTTT |
| 419 | Non_Targeting_Human | CCCGTTTAGGTCATAACCG |
| 420 | Non_Targeting_Human | AACTGCTGGCGTCTGACGC |
| 421 | Non_Targeting_Human | TCCGTGCCCCCGGAGGAAT |
| 422 | Non_Targeting_Human | ACTTAGCCTATTTCTTGCG |
| 423 | Non_Targeting_Human | CAGCAACTGCACTCGCATG |
| 424 | Non_Targeting_Human | CAGCGCCCGCAATGGGCTA |
| 425 | Non_Targeting_Human | TCTCGTTGCATGCTTCGCG |
| 426 | Non_Targeting_Human | GACCGGCCAGCGAAACCAT |
| 427 | Non_Targeting_Human | CACGGAGCGGGGTAAAGAG |
| 428 | Non_Targeting_Human | TTGGCTCCTGCTTCCCCAT |
| 429 | Non_Targeting_Human | CCCGGAACGCTGGTACCCC |

| | | |
|---|---|---|
| 430 | Non_Targeting_Human | TGGGGCGAATCAGGTGTAG |
| 431 | Non_Targeting_Human | TTCGGGAAGCGGGGACGCT |
| 432 | Non_Targeting_Human | ATCATCCGTTTAAACAAAT |
| 433 | Non_Targeting_Human | TCGCAAGTGCATTCCCTGG |
| 434 | Non_Targeting_Human | AGGGTACGTGCTCACCCCC |
| 435 | Non_Targeting_Human | GCTAATTCCGAGGTCGTAT |
| 436 | Non_Targeting_Human | GTTATGCCGTTACTATTAA |
| 437 | Non_Targeting_Human | TGTCCTCACTGGCCGAACG |
| 438 | Non_Targeting_Human | GTGCCGTTGCTGAACTAAG |
| 439 | Non_Targeting_Human | CTGAACTTCGGTAAAGGCG |
| 440 | Non_Targeting_Human | GAGCCAATCTATGGTCGAC |
| 441 | Non_Targeting_Human | GAAGGACCCGGCGCCACTT |
| 442 | Non_Targeting_Human | GCCGTGACAATCACAACTA |
| 443 | Non_Targeting_Human | CGGGCCCGACTCAGTAGCC |
| 444 | Non_Targeting_Human | CCGGACGTCGCTCTTGGTG |
| 445 | Non_Targeting_Human | GTTGGCGGTCAGCGGTGAT |
| 446 | Non_Targeting_Human | CACGGCTCATCGAACTTGG |
| 447 | Non_Targeting_Human | GGCGAAAAGCGAGCCTAGG |
| 448 | Non_Targeting_Human | AAGAGTCGCCTAATTCCGT |
| 449 | Non_Targeting_Human | GGATGGGCGCGCGTGGGGA |
| 450 | Non_Targeting_Human | AGGTCGCGCGCCGGCGTAG |
| 451 | Non_Targeting_Human | CACGCCATGTTGCGAGAAC |
| 452 | Non_Targeting_Human | TCTGGGGCCACGAAACGTT |
| 453 | Non_Targeting_Human | AGGCAGAGCCATGTGGCGG |
| 454 | Non_Targeting_Human | CTCACCGTAGAGACGGCAC |
| 455 | Non_Targeting_Human | CCGCATTGGGAGATGAAAG |
| 456 | Non_Targeting_Human | TAGGACTGAAAACGGATCG |
| 457 | Non_Targeting_Human | CGTTTGACTCACCGAATTG |
| 458 | Non_Targeting_Human | TACGGCGTGCGCCGGCACG |
| 459 | Non_Targeting_Human | TAACATATCAGACGGTCCA |
| 460 | Non_Targeting_Human | ACGGCTCCGTGTGTACCGC |
| 461 | Non_Targeting_Human | TCTGCGGACGCCGTACTAG |
| 462 | Non_Targeting_Human | TGGAGACGCGACTACTACA |
| 463 | Non_Targeting_Human | TCTGTGTATAACTCGGAGG |
| 464 | Non_Targeting_Human | ACATTATTAAGGCCAGCGG |
| 465 | Non_Targeting_Human | GGAAGTTCGGCCCGCACTA |
| 466 | Non_Targeting_Human | AGCCCTGACCCCCACCGGG |
| 467 | Non_Targeting_Human | TGCAGTCAAAGGGCATAAG |
| 468 | Non_Targeting_Human | TCCGACCGCGCTGAAGCAA |
| 469 | Non_Targeting_Human | ACGTCAGTATAGACTGCAA |
| 470 | Non_Targeting_Human | CACGCGCTTAAGTTAATAT |
| 471 | Non_Targeting_Human | GGCCCACGGATTTACGGTG |
| 472 | Non_Targeting_Human | AGGGGCGCACATCGATGTG |

| | | |
|---|---|---|
| 473 | Non_Targeting_Human | GGCCCGGGCCTAACCCCCG |
| 474 | Non_Targeting_Human | GACACGGCGTCGTCATTGA |
| 475 | Non_Targeting_Human | GGTCTAGCTATGAAATTAT |
| 476 | Non_Targeting_Human | GATGGTGCCGGGAACCCGT |
| 477 | Non_Targeting_Human | CGATGACGGGTCCAGACGC |
| 478 | Non_Targeting_Human | CGCGTCAACACGCCATTTA |
| 479 | Non_Targeting_Human | CTTGGCCCCGGGTGACTGG |
| 480 | Non_Targeting_Human | AGGATGGCACCAGATGGGG |
| 481 | Non_Targeting_Human | ACGTGTCTGGCCAGCTAGG |
| 482 | Non_Targeting_Human | GCGACTCCTCACCTGTACC |
| 483 | Non_Targeting_Human | TAGCGCGTTGCGTTCTGAC |
| 484 | Non_Targeting_Human | GGCTATCCAAGGTGCGCAC |
| 485 | Non_Targeting_Human | TGGCGCCCGTAAGAACAAG |
| 486 | Non_Targeting_Human | CGCAGACTCGATTGCCCAC |
| 487 | Non_Targeting_Human | GGTTGCGGGGGGCTCTTTT |
| 488 | Non_Targeting_Human | CCCGACGTAGCACCAATGG |
| 489 | Non_Targeting_Human | AGTGCACCCATACCGAACG |
| 490 | Non_Targeting_Human | GTGCTTCCGGTGAGCCGAC |
| 491 | Non_Targeting_Human | GGGTCGTGGACACACGGTG |
| 492 | Non_Targeting_Human | CCCAAAGTGAGGACGCATT |
| 493 | Non_Targeting_Human | CTTGCGACACACTCTCCAG |
| 494 | Non_Targeting_Human | CTACCGGCGGTCCGCCTCG |
| 495 | Non_Targeting_Human | AGGTGGCGGACGCCCGCAT |
| 496 | Non_Targeting_Human | CGCTGACCCCTAGAACGAG |
| 497 | Non_Targeting_Human | AATCTCCCCGAGAAGAGTC |
| 498 | Non_Targeting_Human | GAATCCCGGGCGCGTGAAA |
| 499 | Non_Targeting_Human | AGAGGGCCCGACTAGGCCG |
| 500 | Non_Targeting_Human | CCGCCAACGAGACATACCT |
| 501 | Non_Targeting_Human | CTACAGTGTCGGTACTCAG |
| 502 | Non_Targeting_Human | GTCGGTGGGGAACACACCG |
| 503 | Non_Targeting_Human | TAAGCTGGACGCAAATGCC |
| 504 | Non_Targeting_Human | CGGCCCGCGAAGCGGGGGT |
| 505 | Non_Targeting_Human | ACAGCACGCAGACACCTAC |
| 506 | Non_Targeting_Human | GGCACTAATGTGGGCGATT |
| 507 | Non_Targeting_Human | TAAGGACGACCCGTCACGG |
| 508 | Non_Targeting_Human | GGGCATAGGCTTGACCGAG |
| 509 | Non_Targeting_Human | GACGGCCCAGAGTACAATA |
| 510 | Non_Targeting_Human | AGGGTACGCGCAAGAGTAG |
| 511 | Non_Targeting_Human | TCCTAGAGGCTCTATAGCC |
| 512 | Non_Targeting_Human | ATTGCGCAAGCCAGGTTAC |
| 513 | Non_Targeting_Human | CGCGTCTTGATGAATGTTG |
| 514 | Non_Targeting_Human | TTTCGATTCGCATGGGGTC |
| 515 | Non_Targeting_Human | TCACGCGAGTTCAACAACG |

| | | |
|---|---|---|
| 516 | Non_Targeting_Human | TCGAACCGCGGCCGTACCG |
| 517 | Non_Targeting_Human | TCAATTCAAACCCACTCGG |
| 518 | Non_Targeting_Human | CAAGAGCTATTCGCTTCGG |
| 519 | Non_Targeting_Human | CTTATCGGGGGCAAAAACG |
| 520 | Non_Targeting_Human | CCGCGTTACGCCGCTACGG |
| 521 | Non_Targeting_Human | TGGGAGCAGATCAAGACTC |
| 522 | Non_Targeting_Human | CATGGGAGGACACCCTCGC |
| 523 | Non_Targeting_Human | ACATCCGCAGAACCGGACT |
| 524 | Non_Targeting_Human | GGCCAGGCAAGCGTTTCCG |
| 525 | Non_Targeting_Human | TTACTACAGGAGCCGAAGG |
| 526 | Non_Targeting_Human | AGGCGGCGCTAACCTCGGG |
| 527 | Non_Targeting_Human | CCAGAAGATCCAAAGGCCC |
| 528 | Non_Targeting_Human | CGGGTGAGGTCCCACCAGA |
| 529 | Non_Targeting_Human | GCTAGTGCACAACATTTGC |
| 530 | Non_Targeting_Human | ACGGGAACCAAGAAGCGGC |
| 531 | Non_Targeting_Human | TGCCCCGCGCATACGACAT |
| 532 | Non_Targeting_Human | AGTCAGAGATGTGTTCCTG |
| 533 | Non_Targeting_Human | TGGAACCGCGACAGGCCCG |
| 534 | Non_Targeting_Human | CGTGAGGAATCTTAAGAGT |
| 535 | Non_Targeting_Human | GTCCCAGGAGCCGCCGTTA |
| 536 | Non_Targeting_Human | TAGATCTCAGCGCGAGTCA |
| 537 | Non_Targeting_Human | CCCGGACAGGAAAATCGCA |
| 538 | Non_Targeting_Human | CAGACCAGTGCGTGCCGAG |
| 539 | Non_Targeting_Human | GGACTAAGGGCGCCTAAGA |
| 540 | Non_Targeting_Human | CGCGAGCCGGCCCTCGTCT |
| 541 | Non_Targeting_Human | AGAGGCGAGTCGTCCTGAT |
| 542 | Non_Targeting_Human | CGGATGGGCCCGATAGCTG |
| 543 | Non_Targeting_Human | CCCCGGCGGACAAAACCGG |
| 544 | Non_Targeting_Human | GACGTAGCACCTCGGCGGT |
| 545 | Non_Targeting_Human | GCAGCCGCGAGTCGTCCCA |
| 546 | Non_Targeting_Human | GGGCACGCGCGCGTCTGAT |
| 547 | Non_Targeting_Human | TTAATCAATCATCGCGTAT |
| 548 | Non_Targeting_Human | GTGGCCGAGGACTAGAAAG |
| 549 | Non_Targeting_Human | CCATGGAGACAGCGCCGGT |
| 550 | Non_Targeting_Human | GGTCGACGAAGATCCGATG |
| 551 | Non_Targeting_Human | CGAGGTGGCGGGGGACGGT |
| 552 | Non_Targeting_Human | GCCCAACGCTTTTTAGATT |
| 553 | Non_Targeting_Human | TCCTTCCCGGCGGCCGTCT |
| 554 | Non_Targeting_Human | GCCGATCTAGGGTGTGTAG |
| 555 | Non_Targeting_Human | GTTGGACGCGCCCCCCTCG |
| 556 | Non_Targeting_Human | CGTGCGCGCGGATGGGCCG |
| 557 | Non_Targeting_Human | CCGATGGTCTTGTACTACA |
| 558 | Non_Targeting_Human | CTCCGGCGACGACAGGCGC |

| | | |
|---|---|---|
| 559 | Non_Targeting_Human | CCGGCAGTTGGGCAGTTTG |
| 560 | Non_Targeting_Human | CCTGGTGCAGTAGACAGCG |
| 561 | Non_Targeting_Human | CGTCAGAGATCGGGGGGAG |
| 562 | Non_Targeting_Human | GGCACATGAGCCTACCATG |
| 563 | Non_Targeting_Human | GTACCGGTGGCGATAAGGA |
| 564 | Non_Targeting_Human | ACGCCGCATGGCCGATTGT |
| 565 | Non_Targeting_Human | ACCGCTGCCCTAACCACTG |
| 566 | Non_Targeting_Human | GTATCGTGCTGGCCGTGGA |
| 567 | Non_Targeting_Human | TAGTCCCGCGGGTCCGAAC |
| 568 | Non_Targeting_Human | GTCGGGCGCTCCATCACCG |
| 569 | Non_Targeting_Human | GAGGGGCGGACGACGTCCG |
| 570 | Non_Targeting_Human | TCGGGCGCCAGTCAGCGAG |
| 571 | Non_Targeting_Human | TGCGGGGCAGTGGGCACGC |
| 572 | Non_Targeting_Human | TGTGGTCGCTTGCAAGTCC |
| 573 | Non_Targeting_Human | GATGAAAGCGGGGGTCGGT |
| 574 | Non_Targeting_Human | GCGCACCCTCGGAAGCGAG |
| 575 | Non_Targeting_Human | ATAGCCAGCCGCCCAAAAT |
| 576 | Non_Targeting_Human | CCGGCCGCTGCCGTCGTCT |
| 577 | AC004257.1-001 | AAAAAAAAGTCTAGGCACAG |
| 578 | AC004257.1-001 | CAAATATTAGCCAGGCGTGG |
| 579 | AC004257.1-001 | ATACAAATATTAGCCAGGCG |
| 580 | AC004257.1-001 | CCATCTCAAAAAAAAAGTCT |
| 581 | AC004257.1-001 | TATTAGCCAGGCGTGGTGGT |
| 582 | AC004257.1-001 | TAAAAATACAAATATTAGCC |
| 583 | AC004257.1-001 | ACTTCGTGTTCCTCCCGCCT |
| 584 | AC004702.2-001 | TCTGTCTTGTCATCCTCCTG |
| 585 | AC004702.2-001 | CCCTCTTCTACCAGCTGAAG |
| 586 | AC004702.2-001 | TACCAGCTGAAGCGGTTCTA |
| 587 | AC004702.2-001 | AGCCCTAGAACCGCTTCAGC |
| 588 | AC004702.2-001 | AATGCAATCTGCTCCACCTC |
| 589 | AC004702.2-001 | GCAATCTGCTCCACCTCAGG |
| 590 | AC004702.2-001 | AAGTGCTCCTCTGACTGCTG |
| 591 | AC004951.6-001 | CACCTCCTCCAACTCCTCTG |
| 592 | AC004951.6-001 | CAGGGCTTGGGTTTCCCCAG |
| 593 | AC004951.6-001 | ACGTTGCCCAGGCTTGAAGC |
| 594 | AC004951.6-001 | AAACGCCCTTTGAGAAGACG |
| 595 | AC004951.6-001 | TTCCCCAGAGGAGTTGGAGG |
| 596 | AC004951.6-001 | GAGGTGGGGAGGAATTGGGT |
| 597 | AC004951.6-001 | GGAGTTGGAGGAGGTGAGTG |
| 598 | AC005154.6-025 | GGGTTACTGATGATTACACT |
| 599 | AC005154.6-025 | AATTGAAACCCATAATGAAG |
| 600 | AC005154.6-025 | ATGCTGAGGCAGTGTCACGC |
| 601 | AC005154.6-025 | TTACCTGTGGATAGAACACA |

| | | |
|---|---|---|
| 602 | AC005154.6-025 | TTGAGTCAATGGAACTGTTC |
| 603 | AC005154.6-025 | TGTTCAGGCATTCTGTAAGG |
| 604 | AC005154.6-025 | TTAAAATATATCAGCTGCTA |
| 605 | AC005616.2-001 | CACTGCTGATGCGTAAGACG |
| 606 | AC005616.2-001 | CTGATGCGTAAGACGAGGAG |
| 607 | AC005616.2-001 | TGCAGGCAGTCAACCAGCTG |
| 608 | AC005616.2-001 | ACTGTTAAAAATGGTCACAG |
| 609 | AC005616.2-001 | GTTTGTTGCTATGCCTCAGG |
| 610 | AC005616.2-001 | GGTGTTTGTTGCTATGCCTC |
| 611 | AC005616.2-001 | GCTATGCCTCAGGAGGTAGG |
| 612 | AC006129.1-001 | CTGGTTCCAAGGACTTGTCG |
| 613 | AC006129.1-001 | ACTTCTGAGAGTTCCAGTAC |
| 614 | AC006129.1-001 | TCTTACCTTCTTACTGGACA |
| 615 | AC006129.1-001 | GAAGAGATCTGCTGGTTCCA |
| 616 | AC006129.1-001 | TCAGTTTCCTCAGAGTCTCC |
| 617 | AC006129.1-001 | AATTACGGAAGAAAGTATCT |
| 618 | AC006129.1-001 | CCGTGAACGAAGAGATCTGC |
| 619 | AC006129.1-002 | CACACTCACGACGTTCTGCA |
| 620 | AC006129.1-002 | CCAGCAGATCTCTTCGTTCA |
| 621 | AC006129.1-002 | CCAGAAGGACTCTTCACACT |
| 622 | AC006129.1-002 | TTCTCTGATTATGATTCTGA |
| 623 | AC006129.1-002 | GAAGATTCTTACCTTCTTAC |
| 624 | AC006129.1-002 | GATTCTGAAGGGAAAAGCGA |
| 625 | AC006129.1-002 | TGAAGGGAAAAGCGATGGAA |
| 626 | AC006129.2-001 | TAAAAACTGAGGAAACTCAG |
| 627 | AC006129.2-001 | ATGTCTCACCTCCCGCCACA |
| 628 | AC006129.2-001 | CTATGCCTGGACGTGCACTC |
| 629 | AC006129.2-001 | ACCAGCACTCAGCACACCAA |
| 630 | AC006129.2-001 | GACCAGCACTCAGCACACCA |
| 631 | AC006129.2-001 | TCTGGCCTGAGTGCACGTCC |
| 632 | AC006129.2-001 | AGCACACCAAGGGCCTGCAC |
| 633 | AC006129.2-003 | TGAGCCCAGGAATAAGTTGG |
| 634 | AC006129.2-003 | ACCCCTTAAAAAGTCACTTG |
| 635 | AC006129.2-003 | ACAGCCTCCAACTTATTCCT |
| 636 | AC006129.2-003 | GACTGAGTCCTGCAGGACAC |
| 637 | AC006129.2-003 | CACCCCTTAAAAAGTCACTT |
| 638 | AC006129.2-003 | TTAAGGGGTGGCTCACCAGG |
| 639 | AC006129.2-003 | GCTTGAGCCCAGGAATAAGT |
| 640 | AC006262.5-006 | TGCCGAAGACCTGAGACAGG |
| 641 | AC006262.5-006 | TGAGGACCCGAGGTCATGGG |
| 642 | AC006262.5-006 | ATGAGATCCACCCATGACCT |
| 643 | AC006262.5-006 | GGTGAGAAGTTCCTTCGGCT |
| 644 | AC006262.5-006 | CGGGTCCTCAGCCCAGCCGA |

| | | |
|---|---|---|
| 645 | AC006262.5-006 | GGGTGGATCTCATGGAGAGA |
| 646 | AC006262.5-006 | TGGGTGGATCTCATGGAGAG |
| 647 | AC006273.4-001 | AAAAAATTAGCTGGGCGTGG |
| 648 | AC006273.4-001 | TGTGGTCCCAGCTACTCAGG |
| 649 | AC006273.4-001 | GAGAATGGCCTGAACCCGAG |
| 650 | AC006273.4-001 | AATGGCCTGAACCCGAGAGG |
| 651 | AC006273.4-001 | TACAAAAAAAAAAATTAGCT |
| 652 | AC006273.4-001 | AAAAAAAAATTAGCTGGGCG |
| 653 | AC006273.4-001 | GGCGTGGTGGCAGGCGCCTG |
| 654 | AC007228.5-001 | AAAGTGTGGGTTGCAAACCC |
| 655 | AC007228.5-001 | GCTGCGGAATCTAAAAAGGT |
| 656 | AC007228.5-001 | ATCTAAAAAGGTAGGTGCCA |
| 657 | AC007228.5-001 | TAAGTCCCTACTGCAGGCTG |
| 658 | AC007228.5-001 | CCAGTCTATGGCAAAGAAGG |
| 659 | AC007228.5-001 | ACAGGCTAAGTCCCTACTGC |
| 660 | AC007228.5-001 | ATTCCAGTCTATGGCAAAGA |
| 661 | AC007383.3-001 | AGGGGAAACTCCTACATCTG |
| 662 | AC007383.3-001 | GGAAGATGTAGGGAGCAGCG |
| 663 | AC007383.3-001 | CTGGAGCAGTCCCCAGATGT |
| 664 | AC007383.3-001 | GAGGGGAAACTCCTACATCT |
| 665 | AC007383.3-001 | GCGAGGAGACTGTCAGTGAG |
| 666 | AC007383.3-001 | CCGGCGGAGAGGAAGATGTA |
| 667 | AC007383.3-001 | GAGGAGACTGTCAGTGAGAG |
| 668 | AC007383.3-002 | CCTCGGGCCGGTAGGCCCCA |
| 669 | AC007383.3-002 | CCGTGGGGCCTACCGGCCCG |
| 670 | AC007383.3-002 | CCTACCGGCCCGAGGCCTAC |
| 671 | AC007383.3-002 | GGCGCCGGTAGGCCTCGGGC |
| 672 | AC007383.3-002 | GCGGCTGCCGTGGGGCCTAC |
| 673 | AC007383.3-002 | CCGGTAGGCCTCGGGCCGGT |
| 674 | AC007383.3-002 | GGCAGGGCGCGGCTGCCGTG |
| 675 | AC007389.3-001 | TCTGACTAATTATATAGCAG |
| 676 | AC007389.3-001 | CCTGCTATATAATTAGTCAG |
| 677 | AC007389.3-001 | TATATAATTAGTCAGAGGGG |
| 678 | AC007389.3-001 | AGGGGCAGTGATTTACCAAA |
| 679 | AC007389.3-001 | TGCTATATAATTAGTCAGAG |
| 680 | AC007389.3-001 | AAAAGGTAAGTCACAAATGG |
| 681 | AC007389.3-001 | GGCAGTGATTTACCAAAAGG |
| 682 | AC008592.8-001 | ATGGATGGGTAGGTAAGTAG |
| 683 | AC008592.8-001 | TAGATAAAAATTAAAAGTTG |
| 684 | AC008592.8-001 | TGGATGGGTAGGTAAGTAGC |
| 685 | AC008592.8-001 | TGGATGGATGGATGAATGGA |
| 686 | AC008592.8-001 | AAAGTTGAGGTCAGGAGCAC |
| 687 | AC008592.8-001 | TAATAAACAAAAAAAAGGAT |

| | | |
|---|---|---|
| 688 | AC008592.8-001 | AGAGCTAATAAACAAAAAAA |
| 689 | AC009120.10-001 | ATTACAGGCATGAGCCACCA |
| 690 | AC009120.10-001 | TGAGGTGGGTGGATCACCTG |
| 691 | AC009120.10-001 | ACAGGCATGCACACCACGCC |
| 692 | AC009120.10-001 | TGTAATCCCAGCACTTTGGG |
| 693 | AC009120.10-001 | CCTCAGCCTCCCAAAGTGCT |
| 694 | AC009120.10-001 | TGCCTGTAATCCCAGCACTT |
| 695 | AC009120.10-001 | AATACACAATTAGCCAGGCG |
| 696 | AC009158.1-007 | ACAAACTAGGTAGTGAACAC |
| 697 | AC009158.1-007 | GAATCAGACAGGATGAGAGG |
| 698 | AC009158.1-007 | CATCAACACATGAATCAGAC |
| 699 | AC009158.1-007 | ACCTTGGTGACAAGGAATGG |
| 700 | AC009158.1-007 | CTGAGTTAGGCAATTGCTTG |
| 701 | AC009158.1-007 | GAGGATAAGGCTAAAGACAG |
| 702 | AC009158.1-007 | GCTACCTTGGTGACAAGGAA |
| 703 | AC009229.6-001 | ACAAGTGGTCAAGAGAGCAT |
| 704 | AC009229.6-001 | GGATAGGCTGGTCTGCATGG |
| 705 | AC009229.6-001 | ACAGGTCTGCTGGAGACAAG |
| 706 | AC009229.6-001 | AAAGGATAGGCTGGTCTGCA |
| 707 | AC009229.6-001 | CGTTTGTTAAAAGGATAGGC |
| 708 | AC009229.6-001 | AAACTTGGGTGAACCCCTGG |
| 709 | AC009229.6-001 | TCCTTCAACCTGCCTGACCC |
| 710 | AC009299.3-002 | CGGAAAGCGAAACTGAGATG |
| 711 | AC009299.3-002 | GGAAAGCGAAACTGAGATGC |
| 712 | AC009299.3-002 | TTCTAGCCTCCGCCTCCAGG |
| 713 | AC009299.3-002 | GGTGAGAGGGAGGAATTCTC |
| 714 | AC009299.3-002 | TCGCTTTCCGGCAGGGTCAG |
| 715 | AC009299.3-002 | TCACCTCCTCACCCGCCTGG |
| 716 | AC009299.3-002 | GAAACTGAGATGCGGGCATT |
| 717 | AC009404.2-002 | CCCTGCACTTTCAACGCCGT |
| 718 | AC009404.2-002 | ACAGTCTTACCCGGACGCCA |
| 719 | AC009404.2-002 | CCGGGTAAGACTGTCCTTCC |
| 720 | AC009404.2-002 | TACGGCGTTGAAAGTGCAGG |
| 721 | AC009404.2-002 | CGGGTAAGACTGTCCTTCCA |
| 722 | AC009404.2-002 | CTACGGCGTTGAAAGTGCAG |
| 723 | AC009404.2-002 | ACTGTCCTTCCAGGGCTGCC |
| 724 | AC011288.2-002 | GCAGGATGCTAAATGATAGC |
| 725 | AC011288.2-002 | GAAGAATAACATGTTTCCTG |
| 726 | AC011288.2-002 | TAAAATCTCAAAGTCTTGTG |
| 727 | AC011288.2-002 | CACTACTCACCTATTCCCAG |
| 728 | AC011288.2-002 | GGTAATTGGAATTCAAGGGC |
| 729 | AC011288.2-002 | ATCATTTAGCATCCTGCCTC |
| 730 | AC011288.2-002 | ATCAGAATAGATGCTAAGCC |

| | | |
|---|---|---|
| 731 | AC013448.1-001 | AAGGTGTGCCCAGTACAGGA |
| 732 | AC013448.1-001 | CATGGCCCAGTTAGAAGGTG |
| 733 | AC013448.1-001 | TGTGGAAATGACAGTGCGTG |
| 734 | AC013448.1-001 | CTGGCATACCCTCCTGTACT |
| 735 | AC013448.1-001 | GGGCCATGTTGCTGTCCCTG |
| 736 | AC013448.1-001 | TTTCCTCAGGGACAGCAACA |
| 737 | AC013448.1-001 | GAGGGTATGCCAGCATCTCC |
| 738 | AC015849.16-002 | AATATTCACATATTTAACTG |
| 739 | AC015849.16-002 | ATTCAGCAGCTAAATACCCC |
| 740 | AC015849.16-002 | CGGAAGTTCATCATTTCCAG |
| 741 | AC015849.16-002 | AAACGAACTTACAATGTCTG |
| 742 | AC015849.16-002 | TGGGCTCTTGATAGATTTGG |
| 743 | AC015849.16-002 | CTGAGTTGGAGATGGAGTCA |
| 744 | AC015849.16-002 | GCGGAAGTTCATCATTTCCA |
| 745 | AC016722.3-001 | TTCGCCCAGTACAATGTTGG |
| 746 | AC016722.3-001 | TTGGGAGCCAAATATCAGGT |
| 747 | AC016722.3-001 | TCCCAAGATCCTGTTCCTGT |
| 748 | AC016722.3-001 | TTACCTTTGATCCTACCCCA |
| 749 | AC016722.3-001 | TTTCATTCTGGCAGGCCTTG |
| 750 | AC016722.3-001 | AGGCCTTGGGGTAGGATCAA |
| 751 | AC016722.3-001 | CAACAAAGAAAACAGCCGAC |
| 752 | AC016747.3-001 | GCACCATCCCTGCTATGAAG |
| 753 | AC016747.3-001 | GGCATTCCCTCTTCATAGCA |
| 754 | AC016747.3-001 | TGCTGTCGGAAAATGTCCAG |
| 755 | AC016747.3-001 | TTCCGACAGCAGGGATCGTT |
| 756 | AC016747.3-001 | ACAGCAGGGATCGTTGGGTG |
| 757 | AC016747.3-001 | CACCATCCCTGCTATGAAGA |
| 758 | AC016747.3-001 | CACCCAACGATCCCTGCTGT |
| 759 | AC016747.3-002 | TCGCTTTAGACACCGTTTGG |
| 760 | AC016747.3-002 | CGCTTTAGACACCGTTTGGA |
| 761 | AC016747.3-002 | GCTTTAGACACCGTTTGGAG |
| 762 | AC016747.3-002 | GCTCATAAACCTGCTCTCCT |
| 763 | AC016747.3-002 | AATCACTGCATTACATCAGC |
| 764 | AC016747.3-002 | ATCAGCAGGTATTCAGCACC |
| 765 | AC016747.3-002 | AAGGCTGCTTCCGCTAACCC |
| 766 | AC016831.7-001 | CCTTCCAGCCCATAAATCAA |
| 767 | AC016831.7-001 | ACTTAACAATATGTGCAGGC |
| 768 | AC016831.7-001 | TTGTTAAGTGTGATCAACTC |
| 769 | AC016831.7-001 | TCACACTTAACAATATGTGC |
| 770 | AC016831.7-001 | GGCAGGACCGAAAAAAGGTG |
| 771 | AC016831.7-001 | TTCTTGAAGAAGGCACACCA |
| 772 | AC016831.7-001 | CCTTTGATTTATGGGCTGGA |
| 773 | AC017002.2-001 | CCTGTAGTGTGACTGTCAGG |

| | | |
|---|---|---|
| 774 | AC017002.2-001 | ACACTACAGGACAGTAACCA |
| 775 | AC017002.2-001 | CACTACAGGACAGTAACCAT |
| 776 | AC017002.2-001 | ACTACAGGACAGTAACCATG |
| 777 | AC017002.2-001 | GCTTCACTTTGGCTTCTCAG |
| 778 | AC017002.2-001 | TTGGAAGAGACTGCAGTGCT |
| 779 | AC017002.2-001 | CAGGACAGTAACCATGGGGC |
| 780 | AC017104.2-001 | GGTGGCAGCATAAGTCCACA |
| 781 | AC017104.2-001 | AGGGCCAGAAAGAGACATTG |
| 782 | AC017104.2-001 | TGCTGGAAACATGACAAATG |
| 783 | AC017104.2-001 | GACAGGTACTAGGGAAGACT |
| 784 | AC017104.2-001 | TGTAAGGTGGGTTACCCTGC |
| 785 | AC017104.2-001 | GAAACTAGAGACAGGTACTA |
| 786 | AC017104.2-001 | GTAAGGTGGGTTACCCTGCA |
| 787 | AC021224.1-201 | CACCATATCTACCACCAGCA |
| 788 | AC021224.1-201 | CGTGCTGGTGGTAGATATGG |
| 789 | AC021224.1-201 | TGGTAGATATGGTGGCAGTG |
| 790 | AC021224.1-201 | TTTGGTGGCAGCCGTGCTGG |
| 791 | AC021224.1-201 | AGCCGTGCTGGTGGTAGATA |
| 792 | AC021224.1-201 | AGAAACTCTGGCCCCTATGG |
| 793 | AC021224.1-201 | AGCAGCAGTAGCTGTGTCAG |
| 794 | AC022007.5-006 | CTGGACAAAGACACAGAGCG |
| 795 | AC022007.5-006 | CAAAGACACAGAGCGCGGCC |
| 796 | AC022007.5-006 | ACACAGGCGGGCACTCTCTG |
| 797 | AC022007.5-006 | GGGAGGCGGAGCTCTGCGGG |
| 798 | AC022007.5-006 | GCCTGGGAGGCGGAGCTCTG |
| 799 | AC022007.5-006 | TTGGGGCAGCAGCGCCTGGA |
| 800 | AC022007.5-006 | GGGCAGCAGCGCCTGGAAGG |
| 801 | AC022182.1-003 | TCCCGCCACACCGCCCACCA |
| 802 | AC022182.1-003 | AGGGAATGCGAAGGGCGCCG |
| 803 | AC022182.1-003 | GGGATGTGCCCGCCTGCCCC |
| 804 | AC022182.1-003 | CGGGATGTGCCCGCCTGCCC |
| 805 | AC022182.1-003 | CCGCGGGCCCTGCGCGCCCG |
| 806 | AC022182.1-003 | CCGCCTGCCCCGGGCGCGCA |
| 807 | AC022182.1-003 | CCCTGCGCGCCCGGGGCAGG |
| 808 | AC022182.3-001 | GCACCCACACCCTGAGCAGT |
| 809 | AC022182.3-001 | GAAAGCAAAAGGGATGCCGG |
| 810 | AC022182.3-001 | GAGGAAGCATCTGCCACCAC |
| 811 | AC022182.3-001 | AGCACCCACACCCTGAGCAG |
| 812 | AC022182.3-001 | GGATTTGGGATTTGGCCAGG |
| 813 | AC022182.3-001 | TGGTGGCAGATGCTTCCTCC |
| 814 | AC022182.3-001 | CCTCTCCTGCCCACTGCTCA |
| 815 | AC034187.2-001 | ATGACATGGAAATTCAGACC |
| 816 | AC034187.2-001 | AGAGAAAAGAGATTCTACTG |

| | | |
|---|---|---|
| 817 | AC034187.2-001 | ATACTATTTATAGAATGACA |
| 818 | AC034187.2-001 | AAAAATAGAGAAATATGGCT |
| 819 | AC034187.2-001 | GAGATTCTACTGTGGCCACC |
| 820 | AC034187.2-001 | CAAAAATTAGCTGGGTGTGG |
| 821 | AC034187.2-001 | ATATGCAAATATATTCTCAC |
| 822 | AC058791.1-001 | AGTCTCCTACTGAACTGTCA |
| 823 | AC058791.1-001 | GCAGAAGAATCGCTGTTTGG |
| 824 | AC058791.1-001 | AAAACCTGCCATTATAAAGG |
| 825 | AC058791.1-001 | GTCTGCCGTGACAGTTCAGT |
| 826 | AC058791.1-001 | AAACCTGCCATTATAAAGGA |
| 827 | AC058791.1-001 | ACGTTCATTCCAAAAAACTG |
| 828 | AC058791.1-001 | TGCAGAAGAATCGCTGTTTG |
| 829 | AC058791.1-002 | GACTCAAGACGACCAACACT |
| 830 | AC058791.1-002 | GCTGGTGATTTCCTAAGCAG |
| 831 | AC058791.1-002 | GATTATTGGTAACCCAGTGT |
| 832 | AC058791.1-002 | AAATGTTACACTCGGTGAGT |
| 833 | AC058791.1-002 | ACTACAAGAAATGTTACACT |
| 834 | AC058791.1-002 | TGTTACACTCGGTGAGTAGG |
| 835 | AC058791.1-002 | GGAAGTATGGATTCATTGCC |
| 836 | AC068286.1-001 | CCGAGTGCCTCCCTCCACCG |
| 837 | AC068286.1-001 | TCATTGACTTCTCACAACAG |
| 838 | AC068286.1-001 | AGTGGAGTGATGTACCAAAG |
| 839 | AC068286.1-001 | TCCAAGTTTCCTTCTAGGCA |
| 840 | AC068286.1-001 | ATCAGATGCTGTGTTCTCTC |
| 841 | AC068286.1-001 | ACTGTTGTGAGAAGTCAATG |
| 842 | AC068286.1-001 | ACAGCATCTGATTTCCTCTT |
| 843 | AC069277.2-004 | CTAAAACAAAAAAAGCAGCC |
| 844 | AC069277.2-004 | CCCAGCACTTTGAGAGGCCG |
| 845 | AC069277.2-004 | ATAAAGCTGTGATATTTAGA |
| 846 | AC069277.2-004 | AACTTTCTTCAGGCCAGGCG |
| 847 | AC069277.2-004 | TTTCTTCAGGCCAGGCGCGG |
| 848 | AC069277.2-004 | CTCCAGACTTTATTTGGTAA |
| 849 | AC069277.2-004 | CCCCTTTACCAAATAAAGTC |
| 850 | AC073046.25-001 | CCCCCACTCCGGAGCCGGAA |
| 851 | AC073046.25-001 | CCTTTCCGGCTCCGGAGTGG |
| 852 | AC073046.25-001 | AGTCGGTGCCTTTCCGGCTC |
| 853 | AC073046.25-001 | GGCGGGCAGGTCGGTGCAGT |
| 854 | AC073046.25-001 | GCCTTTCCGGCTCCGGAGTG |
| 855 | AC073046.25-001 | TGCCTTTCCGGCTCCGGAGT |
| 856 | AC073046.25-001 | CCCCTCCCCCACTCCGGAGC |
| 857 | AC073257.1-001 | GGGTAGGTGTGTCCAACACA |
| 858 | AC073257.1-001 | AGTGGGAGAGAACACAGCAG |
| 859 | AC073257.1-001 | AGGAAGTGATGAGATCAGCA |

| | | |
|---|---|---|
| 860 | AC073257.1-001 | AGGGAGAGCAAGAGTCAATG |
| 861 | AC073257.1-001 | AGGGTAGGTGTGTCCAACAC |
| 862 | AC073257.1-001 | GGGGAAGAAAAGCTTCCGTA |
| 863 | AC073257.1-001 | CTGATCTCATCACTTCCTTA |
| 864 | AC079610.2-001 | AAAGGTGAGTTCTCAACACC |
| 865 | AC079610.2-001 | TCTTCATTGCTAAGCATCTG |
| 866 | AC079610.2-001 | CCTAGTTAACGCGAGAGCTC |
| 867 | AC079610.2-001 | TAATAGAAAGACTGACCCAG |
| 868 | AC079610.2-001 | TAGATCCCCAAGTGTTCCCA |
| 869 | AC079610.2-001 | ATGGGCAAGAATCAAGAGAA |
| 870 | AC079610.2-001 | GAAAACCAAGCTGCTTGGGT |
| 871 | AC090505.4-001 | TCCATGGATAGGGACCAGCA |
| 872 | AC090505.4-001 | GAACTGGGTATCCTAAGATG |
| 873 | AC090505.4-001 | AGGGCCATTAACTCCCCTGC |
| 874 | AC090505.4-001 | CCATGGATAGGGACCAGCAG |
| 875 | AC090505.4-001 | TATTACTGATAATTTGCAGC |
| 876 | AC090505.4-001 | CCCCTGCTGGTCCCTATCCA |
| 877 | AC090505.4-001 | TTAATGGCCCTGGAGACAGA |
| 878 | AC090505.6-002 | AGCAGTTATTAAGTGTCAGG |
| 879 | AC090505.6-002 | TTAATTCTCACGATGCCATG |
| 880 | AC090505.6-002 | ATTCCAACCTCGTGTGAGAG |
| 881 | AC090505.6-002 | TCAGGTGAAGCATTTAGAAG |
| 882 | AC090505.6-002 | GCCAACCTATATTAGTGCTC |
| 883 | AC090505.6-002 | AAAGAGCACAGTAAACTGCA |
| 884 | AC090505.6-002 | TTTGGTTGTATCTCTAATCC |
| 885 | AC091814.3-001 | CCGTCTGGGACGGAGATGGG |
| 886 | AC091814.3-001 | CCCCATCTCCGTCCCAGACG |
| 887 | AC091814.3-001 | CATCTCCGTCCCAGACGGGG |
| 888 | AC091814.3-001 | CCCCCCATCTCCGTCCCAGA |
| 889 | AC091814.3-001 | CCCGTCTGGGACGGAGATGG |
| 890 | AC091814.3-001 | CCTCACTTCCCAGTAGGGGG |
| 891 | AC091814.3-001 | CCCCGTCTGGGACGGAGATG |
| 892 | AC092415.1-002 | CAGGAACAAATAGGACTGGG |
| 893 | AC092415.1-002 | ACAATGACACCATGTTTGCT |
| 894 | AC092415.1-002 | CTTACAAAGGAATGAGCAAG |
| 895 | AC092415.1-002 | AAATCAGGAACAAATAGGAC |
| 896 | AC092415.1-002 | TTTATCTTCTGGGAATCTGG |
| 897 | AC092415.1-002 | CTAATTACAGTGAAGAGATG |
| 898 | AC092415.1-002 | AATTTCAACTCTGCTTACCT |
| 899 | AC092669.3-001 | TAAATTGTTACCATCCACGG |
| 900 | AC092669.3-001 | CATCAGTAGACCAAACACTG |
| 901 | AC092669.3-001 | GACATCAGTAGACCAAACAC |
| 902 | AC092669.3-001 | TAACTACTTACCCCAGTGTT |

| | | |
|---|---|---|
| 903 | AC092669.3-001 | ACATCAGTAGACCAAACACT |
| 904 | AC092669.3-001 | AAATTGCTAATTATCCAGAG |
| 905 | AC092669.3-001 | AGAAAAGCTATGAAAATAGT |
| 906 | AC093323.3-002 | CCTGTTGCTAGTCTGGTCGT |
| 907 | AC093323.3-002 | TCTGGTCGTTGGTGACAGCG |
| 908 | AC093323.3-002 | TGCCCCAACTTTCCGCAGCG |
| 909 | AC093323.3-002 | CCAACGACCAGACTAGCAAC |
| 910 | AC093323.3-002 | TTTGGATACCGTCCTCGCTG |
| 911 | AC093323.3-002 | GTCCTCGCTGCGGAAAGTTG |
| 912 | AC093323.3-002 | CCAACTTTCCGCAGCGAGGA |
| 913 | AC093642.3-002 | TTTCAACGTAAGGATTTGGG |
| 914 | AC093642.3-002 | ATTTCAACGTAAGGATTTGG |
| 915 | AC093642.3-002 | GATTTCAACGTAAGGATTTG |
| 916 | AC093642.3-002 | GGGTTTAGGATTTCAACGTA |
| 917 | AC093642.3-002 | AGGGCACTGCTGTGGGCTGA |
| 918 | AC093642.3-002 | TGACAACATTCACACCCACA |
| 919 | AC093642.3-002 | GGAGGAAGTCTAGGAGAGGA |
| 920 | AC096772.6-001 | GAGAGTTCTGACATGCATCA |
| 921 | AC096772.6-001 | AGGAGTAAAGGGAATCATGT |
| 922 | AC096772.6-001 | TTTCCTGGTAAAAGTGCTAA |
| 923 | AC096772.6-001 | TCCTGAGGTTAAGGCACACC |
| 924 | AC096772.6-001 | ACATGCATCATGGAATGCAC |
| 925 | AC096772.6-001 | ACTCTTCATATAGATGCTCT |
| 926 | AC096772.6-001 | TATATGAAGAGTAGCCAGCA |
| 927 | AC114730.2-001 | GATACTCACAAAAAGAGACA |
| 928 | AC114730.2-001 | ACTCACAAAAAGAGACAAGG |
| 929 | AC114730.2-001 | CTCACAAAAAGAGACAAGGC |
| 930 | AC114730.2-001 | ACAAAAAGAGACAAGGCGGG |
| 931 | AC114730.2-001 | AGGAAGCGGTCGCTGTCCCC |
| 932 | AC114730.2-001 | GCGAAGTGCTGACTGCGCCG |
| 933 | AC114730.2-001 | CAACACACCGCGGACGCCCG |
| 934 | AC131056.5-001 | AAATATTCTAAAATTGATTG |
| 935 | AC131056.5-001 | TATTCTAAAATTGATTGTGG |
| 936 | AC131056.5-001 | ATTGTGGTGGCCAGGCACAG |
| 937 | AC131056.5-001 | TCCAGTCTGGCGAGAGAGTG |
| 938 | AC131056.5-001 | CGAGACTAGCCTGACCAACA |
| 939 | AC131056.5-001 | CGAGGCGGGCGGATCACCTG |
| 940 | AC131056.5-001 | GCACTTTGAGAGGCCGAGGC |
| 941 | AC133644.2-002 | ATCTGCTTCAAAGAACACAT |
| 942 | AC133644.2-002 | AAAGAACACATCGGCGACAG |
| 943 | AC133644.2-002 | CAAAGAACACATCGGCGACA |
| 944 | AC133644.2-002 | TGTATTCAGCCCTAACCGTA |
| 945 | AC133644.2-002 | GTTGACTCACTCAAGGCGGT |

| | | |
|---|---|---|
| 946 | AC133644.2-002 | CGGTTAGGGCTGAATACAAT |
| 947 | AC133644.2-002 | AAAGTTGCTACCATACGGTT |
| 948 | AC133644.2-003 | GTTTGTTGACTCACTCAAGG |
| 949 | AC133644.2-003 | GGTCTGAAGCCAACCGTCCT |
| 950 | AC133644.2-003 | CCCTTGCACTCACTCAGGAC |
| 951 | AC133644.2-003 | ACAGGCCAGGAAACCGAGGA |
| 952 | AC133644.2-003 | GCCAGGAAACCGAGGACGGT |
| 953 | AC133644.2-003 | TCCAACCAGACTGATGCCAC |
| 954 | AC133644.2-003 | GGAAAAGCAGCAGATCTGCA |
| 955 | AC137934.1-006 | AGGAAGAAGTTGAACACCCT |
| 956 | AC137934.1-006 | TGCGGTGTTTGAAGTGACAA |
| 957 | AC137934.1-006 | CTGGAAGTAGTCACAGCCCA |
| 958 | AC137934.1-006 | GCTGCTGTGACAAAAAAGCA |
| 959 | AC137934.1-006 | GAGGAAGAAGTTGAACACCC |
| 960 | AC137934.1-006 | TGTGACAAAAAAGCAGGGAA |
| 961 | AC137934.1-006 | CCTGGAAGTAGTCACAGCCC |
| 962 | AC138035.2-001 | CCATAAACACCCAAACAGCA |
| 963 | AC138035.2-001 | GCCAGTAATAGAAAATAAGG |
| 964 | AC138035.2-001 | TTGATAAAATCACTGGCCAT |
| 965 | AC138035.2-001 | GGCATGATTGATAAAATCAC |
| 966 | AC138035.2-001 | AAACACAAGGAGGTCCTGGG |
| 967 | AC138035.2-001 | GAGATTGACTTAACTACCAA |
| 968 | AC138035.2-001 | CTGTGTTGCTAAACACAAGG |
| 969 | AC138035.2-003 | CCTCCAGCTCTGTGGCCCTG |
| 970 | AC138035.2-003 | GCTGCTCACCTCCAGCTCTG |
| 971 | AC138035.2-003 | GTCAGATCAGCAGCAGCATT |
| 972 | AC138035.2-003 | CTCTGAAGAATGGCGTGGGT |
| 973 | AC138035.2-003 | CAGCAGCATTAGGTTGTCAT |
| 974 | AC159540.1-001 | TAATTGATATGTGTATGTTG |
| 975 | AC159540.1-001 | TTTGCATTGATGTTCATCGA |
| 976 | AC159540.1-001 | ACTATTTGAAAATATTGAGG |
| 977 | AC159540.1-001 | GGAACTATTTGAAAATATTG |
| 978 | AC159540.1-001 | TGGCCTCATAGAATGAGTTG |
| 979 | AC159540.1-001 | ATTTGAAAATATTGAGGAGG |
| 980 | AC159540.1-001 | GTTGGCCTCATAGAATGAGT |
| 981 | AC159540.1-004 | GAGAGGTAGGACAAAATGAG |
| 982 | AC159540.1-004 | AAAGCAGTGCTTCACTACAG |
| 983 | AC159540.1-004 | TTCTAAGAGACTCTTCTCGC |
| 984 | AC159540.1-004 | AGAAGCTGAAGTTAAAGTTG |
| 985 | AC159540.1-004 | TTTCACGGTCACAGAGAGGT |
| 986 | AC159540.1-004 | CGTCTTTCACGGTCACAGAG |
| 987 | AC159540.1-004 | CTCTCTGTGACCGTGAAAGA |
| 988 | AC159540.1-014 | ATGATCCAATATTGACAATG |

| | | |
|---|---|---|
| 989 | AC159540.1-014 | AGATGATCCAATATTGACAA |
| 990 | AC159540.1-014 | AGGCAGAAAATTGACCAGAA |
| 991 | AC159540.1-014 | TTGTCAATATTGGATCATCT |
| 992 | AC159540.1-014 | AGGCTACCACACAACAATAG |
| 993 | AC159540.1-014 | GATGATCCAATATTGACAAT |
| 994 | AC159540.1-014 | TATAGTCTGTTGTTTCGTGT |
| 995 | AC159540.1-016 | ATGTTTATGTGTCTTGAGGA |
| 996 | AC159540.1-016 | CCTCAAGACACATAAACATC |
| 997 | AC159540.1-016 | CCTGATGTTTATGTGTCTTG |
| 998 | AC159540.1-016 | GATGGCCTGCCCTTTCTCTC |
| 999 | AC159540.1-016 | CTGAAATCCTACAAGCCAGA |
| 1000 | AC159540.1-016 | GAGATCTGCTGTTAGCCTAA |
| 1001 | AC159540.1-016 | GGCAGCCAGAGAGAAAGGGC |
| 1002 | AC159540.1-017 | ACATACTTAAACACACAGAC |
| 1003 | AC159540.1-017 | AAACTAAGCTTTCTAAATGA |
| 1004 | AC159540.1-017 | GATCTGCTGTTAGCCTAATG |
| 1005 | AC159540.1-017 | TTACAGACAAGCAAATTCTG |
| 1006 | AC159540.1-017 | TACAAGCCAGAAGGGATTGG |
| 1007 | AC159540.1-017 | AAGGGCAGGCCATCTACAAA |
| 1008 | AC159540.1-017 | AATCTCTTGTAAGGCAGATG |
| 1009 | AC159540.1-020 | ATGTGGCATATGATGATGTG |
| 1010 | AC159540.1-020 | GTATGTTTCACAGTAATAGT |
| 1011 | AC159540.1-020 | AATATATATTCATCCAATGC |
| 1012 | AC159540.1-020 | CTAATACTGTCATTGGAGTG |
| 1013 | AC159540.1-020 | CAGTATTAGACAGATGATTG |
| 1014 | AC159540.1-020 | AGTATGTTTCACAGTAATAG |
| 1015 | AC159540.1-020 | TTGAGTGCAGAGTTCTGTAA |
| 1016 | AC159540.2-001 | CATTTCCAAAGCCAAGCACG |
| 1017 | AC159540.2-001 | TCCAAAGCCAAGCACGTGGT |
| 1018 | AC159540.2-001 | GACAACAAATGCTTGCTCCT |
| 1019 | AC159540.2-001 | CTTCATCAGTCAGACAACAT |
| 1020 | AC159540.2-001 | AGAGATGTACAAAGTGCCAT |
| 1021 | AC159540.2-001 | AAAATGTAGGGCTTCAATGT |
| 1022 | AC159540.2-001 | GCCATGGGATCACCAAGCAG |
| 1023 | AF011889.2-001 | GGGTCCTTCAGCTCCTCGGG |
| 1024 | AF011889.2-001 | TGTGTCCTCCCTACCACCCG |
| 1025 | AF011889.2-001 | CCTACCACCCGAGGAGCTGA |
| 1026 | AF011889.2-001 | TCGGGTGGTAGGGAGGACAC |
| 1027 | AF011889.2-001 | TGGTAGGGAGGACACAGGCT |
| 1028 | AF011889.2-001 | GGTAGGGAGGACACAGGCTG |
| 1029 | AF011889.2-001 | AGGGAGGACACAGGCTGGGG |
| 1030 | AF127936.3-002 | TGTGAAGAGGCATTCTGTCT |
| 1031 | AF127936.3-002 | TAGCATAATTTGAATGTCTG |

| | | |
|---|---|---|
| 1032 | AF127936.3-002 | AGCATAATTTGAATGTCTGG |
| 1033 | AF127936.3-002 | AGCAATTCATGTTTCTGTGT |
| 1034 | AF127936.3-002 | CTAGCATAATTTGAATGTCT |
| 1035 | AF127936.3-002 | ACTGAGCTGTGCTAACTTGG |
| 1036 | AF127936.3-002 | GAGCAATTCATGTTTCTGTG |
| 1037 | AF127936.5-001 | GAACATGCAGATAATACAAG |
| 1038 | AF127936.5-001 | GAGGTGCGGTTTGTGACTGA |
| 1039 | AF127936.5-001 | ACACAAAGCCCCTCTCCTTG |
| 1040 | AF127936.5-001 | GCCAGAATATTCTAAGCCTG |
| 1041 | AF127936.5-001 | TGCCAGAATATTCTAAGCCT |
| 1042 | AF127936.5-001 | AATAGAATGCAAAGTCCACA |
| 1043 | AF127936.5-001 | TGCAAAGTCCACAAGGAGAG |
| 1044 | AF127936.7-006 | ATGCCCGATAGTTCAACACA |
| 1045 | AF127936.7-006 | TGCCCGATAGTTCAACACAA |
| 1046 | AF127936.7-006 | GTGACCGAGCCACCAAAGAC |
| 1047 | AF127936.7-006 | AGTCCACATTTCCCCAGTAT |
| 1048 | AF127936.7-006 | TAGCCCTTGTGTTGAACTAT |
| 1049 | AF127936.7-006 | CTTTAACTGAAGGGCAGCCA |
| 1050 | AF127936.7-006 | AAGGGCTAAGCAATATTTCA |
| 1051 | AL122127.25-001 | CAGCCTTAGATGCCAGGTCA |
| 1052 | AL122127.25-001 | CCTTAGATGCCAGGTCAGGG |
| 1053 | AL122127.25-001 | GCTGCAGGGATCTGGGCATG |
| 1054 | AL122127.25-001 | CAGCTGTGTCTAGTGCTGCA |
| 1055 | AL122127.25-001 | GCAGCACTAGACACAGCTGG |
| 1056 | AL122127.25-001 | CTGGAGCAGCCTTAGATGCC |
| 1057 | AL122127.25-001 | CCTGCAGCACTAGACACAGC |
| 1058 | AL122127.25-002 | TCTTCCCCGCCTGCCATCGA |
| 1059 | AL122127.25-002 | TCAGACCCACCTGCCCTCGA |
| 1060 | AL122127.25-002 | GCCTGCCATCGAGGGCAGGT |
| 1061 | AL122127.25-002 | ATCTTCCCCGCCTGCCATCG |
| 1062 | AL122127.25-002 | ACCCACCTGCCCTCGATGGC |
| 1063 | AL122127.25-002 | TGGGTCTGACTCACCCTCCC |
| 1064 | AL122127.25-002 | TCACCTCAGCCCCCTCCTGG |
| 1065 | AL928768.3-001 | TGTGGTGCTGGACTTGCCGC |
| 1066 | AL928768.3-001 | TCCAGCACCACATAGGGAGG |
| 1067 | AL928768.3-001 | AAGTCCAGCACCACATAGGG |
| 1068 | AL928768.3-001 | GCAGATGCCGCCTCCCTATG |
| 1069 | AL928768.3-001 | GGCAAGTCCAGCACCACATA |
| 1070 | AL928768.3-001 | AGGTGATGGTGGTGGGCCAC |
| 1071 | AL928768.3-001 | CGGCAAGTCCAGCACCACAT |
| 1072 | AP000442.4-001 | CCCTTCTCTCACTGTCAGAG |
| 1073 | AP000442.4-001 | CCCCTCTGACAGTGAGAGAA |
| 1074 | AP000442.4-001 | CGGCAAGCCCGCGACCAAGC |

| | | |
|---|---|---|
| 1075 | AP000442.4-001 | GACCCTTCTCTCACTGTCAG |
| 1076 | AP000442.4-001 | GCCCCTCTGACAGTGAGAGA |
| 1077 | AP000442.4-001 | ACCCTTCTCTCACTGTCAGA |
| 1078 | AP000442.4-001 | TTCTCTCACTGTCAGAGGGG |
| 1079 | AP000476.1-001 | TAGATGTAGGAAGTGACTGT |
| 1080 | AP000476.1-001 | CTTAACCTCCTAAATTGGGA |
| 1081 | AP000476.1-001 | CCCCAGAGAAAGACGCCGGA |
| 1082 | AP000476.1-001 | CCTTCCGGCGTCTTTCTCTG |
| 1083 | AP000476.1-001 | GGTCTGTTGGTTGAGCTCTG |
| 1084 | AP000476.1-001 | TATGTCCTTCCCAATTTAGG |
| 1085 | AP000476.1-001 | TGACTGTAGGTTTAGATGAA |
| 1086 | AP000477.3-001 | TCAGATACTGAGTTTATGAC |
| 1087 | AP000477.3-001 | TTTGTTGAATTCGTTGACAA |
| 1088 | AP000477.3-001 | TGTAGTTGAATGCAAAAATG |
| 1089 | AP000477.3-001 | TTTAGAGGTTCAAGGTACCT |
| 1090 | AP000477.3-001 | GCTGTCTTAATTAGTATTGA |
| 1091 | AP000477.3-001 | AAAGCTGTAAGCAAAGCCAA |
| 1092 | AP000477.3-001 | CTTAAAACAACTTGAAAATG |
| 1093 | AP000705.7-001 | AGAAACCAATAGGTCGGGCG |
| 1094 | AP000705.7-001 | AGAATTGCTTGAACCCAAGA |
| 1095 | AP000705.7-001 | TCACTGCACTCCACCCTGGG |
| 1096 | AP000705.7-001 | AAAAAAGAAACCAATAGGTC |
| 1097 | AP000705.7-001 | GAGAATTGCTTGAACCCAAG |
| 1098 | AP000705.7-001 | GAGTCTCACTCTGCCACCCA |
| 1099 | AP000705.7-001 | TGTCAAAAAAAGAAACCAAT |
| 1100 | AP001046.5-001 | AGGGTCGTGAGAAAATACGC |
| 1101 | AP001046.5-001 | ATTCCGCCTGAATTAAGCAG |
| 1102 | AP001046.5-001 | CCGCCACTGCTTAATTCAGG |
| 1103 | AP001046.5-001 | CCAAGGGAAACATGTCCACG |
| 1104 | AP001046.5-001 | AGGCGGAATCACGGCTCAGA |
| 1105 | AP001046.5-001 | CAGGCGGAATCACGGCTCAG |
| 1106 | AP001046.5-001 | ACGCAGGAAGAGATACCCCG |
| 1107 | AP001055.6-001 | TTACCCGATTGTTGCAGGAG |
| 1108 | AP001055.6-001 | CTTACCCGATTGTTGCAGGA |
| 1109 | AP001055.6-001 | CCTTACCCGATTGTTGCAGG |
| 1110 | AP001055.6-001 | TGGCCCCTCCTGCAACAATC |
| 1111 | AP001055.6-001 | TTGCAGGAGGGGCCAGCACA |
| 1112 | AP001055.6-001 | CTGGCCCCTCCTGCAACAAT |
| 1113 | AP001055.6-001 | TCCCCTTACCCGATTGTTGC |
| 1114 | AP001055.6-002 | TCACGAGTCACCAGAGAGAG |
| 1115 | AP001055.6-002 | TATCACGAGTCACCAGAGAG |
| 1116 | AP001055.6-002 | AGTCACCAGAGAGAGGGGAA |
| 1117 | AP001055.6-002 | ATCACGAGTCACCAGAGAGA |

| | | |
|---|---|---|
| 1118 | AP001055.6-002 | AGAGAGAGGGGAACGGAGAG |
| 1119 | AP001055.6-002 | CAGAGCTGAAGGGGAACTGG |
| 1120 | AP001055.6-002 | TCCTGGGCACAGAGCTGAAG |
| 1121 | AP001056.1-001 | CACAGAGACTCTGTCCCAGG |
| 1122 | AP001056.1-001 | GAGACTCTGTCCCAGGAGGT |
| 1123 | AP001056.1-001 | GCGGGCATGTGCATGGTTCA |
| 1124 | AP001056.1-001 | AGAAAAAATGCCTACCTCCT |
| 1125 | AP001056.1-001 | AGGCCTGGCTGTCTGCAGGG |
| 1126 | AP001056.1-001 | TCCTGAGTACCTGGGACTAG |
| 1127 | AP001056.1-001 | TGGTGCGTACCCCTAGTCCC |
| 1128 | AP001058.3-001 | GGCCCGGGGAGCATGGGTCA |
| 1129 | AP001058.3-001 | CCATGCTCCCCGGGCCCCTG |
| 1130 | AP001058.3-001 | TCGCCCTGACCCATGCTCCC |
| 1131 | AP001058.3-001 | CTCAGGGGCCCGGGGAGCAT |
| 1132 | AP001058.3-001 | CAGGGCGAGTGGCTGCTCCC |
| 1133 | AP001058.3-001 | GGTCCTCCTGGGTGCCTCAG |
| 1134 | AP001058.3-001 | CGCCCTGACCCATGCTCCCC |
| 1135 | AP001059.7-001 | TCAGCTGAACCACCCTCCAA |
| 1136 | AP001059.7-001 | GGACATGGGCATCCTTTGGA |
| 1137 | AP001059.7-001 | AGGGTGGTTCAGCTGAGCAT |
| 1138 | AP001059.7-001 | GGATCCGGGAATGGGGACAT |
| 1139 | AP001059.7-001 | AGGTAACATTCACAGGATCC |
| 1140 | AP001059.7-001 | AATGTTACCTCATGTGGCAA |
| 1141 | AP001059.7-001 | ATTCACAGGATCCGGGAATG |
| 1142 | AP001462.6-001 | AGCCGGAACGAGATTTACGG |
| 1143 | AP001462.6-001 | CCGGGATTAAGCGAGAGTCG |
| 1144 | AP001462.6-001 | CGGAACGAGATTTACGGCGG |
| 1145 | AP001462.6-001 | GCCGGAACGAGATTTACGGC |
| 1146 | AP001462.6-001 | CCGTAAATCTCGTTCCGGCT |
| 1147 | AP001462.6-001 | GGATTAAGCGAGAGTCGCGG |
| 1148 | AP001462.6-001 | CCGGAACGAGATTTACGGCG |
| 1149 | AP001505.10-001 | TTCACAAGAAACCCAGACGG |
| 1150 | AP001505.10-001 | TCACAAGAAACCCAGACGGT |
| 1151 | AP001505.10-001 | GCATCCCAATCCCACCGTCT |
| 1152 | AP001505.10-001 | CGGTGGGATTGGGATGCTGG |
| 1153 | AP001505.10-001 | GCGCTGTGCTCTGTGCATCC |
| 1154 | AP001505.10-001 | ATGCACAGAGCACAGCGCGG |
| 1155 | AP001505.10-001 | GCACAGAGCACAGCGCGGTG |
| 1156 | AP006222.2-004 | CACTATCTGGTATGTCACGT |
| 1157 | AP006222.2-004 | GTCACGTAGGTAATTTACAA |
| 1158 | AP006222.2-004 | TATCTGACATGAAATGACAT |
| 1159 | AP006222.2-004 | TGTCACGTAGGTAATTTACA |
| 1160 | AP006222.2-004 | TCCAGGCACTATTCTGACTG |

| | | |
|---|---|---|
| 1161 | AP006222.2-004 | TGTGGCAGGAGTGAACAAAC |
| 1162 | AP006222.2-004 | TTTCAGAACAGAGTACTAAA |
| 1163 | AP006621.5-002 | GCCAATAGGTAGGGTCGCGA |
| 1164 | AP006621.5-002 | GGCCAATAGGTAGGGTCGCG |
| 1165 | AP006621.5-002 | CTGGCAGGGGCCAATAGGTA |
| 1166 | AP006621.5-002 | CCTGGCAGGGGCCAATAGGT |
| 1167 | AP006621.5-002 | GCGGGTTCGGCATCCGAGCC |
| 1168 | AP006621.5-002 | TTCGGCATCCGAGCCTGGCA |
| 1169 | AP006621.5-002 | GCTCGGATGCCGAACCCGCC |
| 1170 | AP006621.5-003 | CGTTCGTACCCGGTCCGGGT |
| 1171 | AP006621.5-003 | CTCGGATGCCGAACCCGCCG |
| 1172 | AP006621.5-003 | TGCGCGTTCGTACCCGGTCC |
| 1173 | AP006621.5-003 | CTGCGCGTTCGTACCCGGTC |
| 1174 | AP006621.5-003 | AGGCGCGACCTGCGACCGCC |
| 1175 | AP006621.5-003 | GGCGCGACCTGCGACCGCCC |
| 1176 | AP006621.5-003 | GAACCCGCCGGGGCCAAAGC |
| 1177 | APH1A | GGAACCGAAACCCCAAATGA |
| 1178 | APH1A | GTGGCGCGGACCTTCATTTG |
| 1179 | APH1A | GCAATGTCACCCCCAGACCC |
| 1180 | APH1A | AATGTCACCCCCAGACCCCG |
| 1181 | APH1A | CAATGTCACCCCCAGACCCC |
| 1182 | APH1A | AGTGGCGCGGACCTTCATTT |
| 1183 | APH1A | CCCCCCCTTCCCCTTCCCCG |
| 1184 | APTR-003 | GCCACAGGGACCCTCTTCGG |
| 1185 | APTR-003 | TTGCTGAACGCGCAGATGCA |
| 1186 | APTR-003 | GCACCCCAGGAAAAATGGAA |
| 1187 | APTR-003 | CCGAGCTCTGACCTCCGAAG |
| 1188 | APTR-003 | GGCACCCCAGGAAAAATGGA |
| 1189 | APTR-003 | CGAGCTCTGACCTCCGAAGA |
| 1190 | APTR-003 | CTTGCTGAACGCGCAGATGC |
| 1191 | APTR-004 | CGAGTTCACGCCAGCGACAG |
| 1192 | APTR-004 | CGTGAACTCGCGTGCCCGGT |
| 1193 | APTR-004 | GCGTGAACTCGCGTGCCCGG |
| 1194 | APTR-004 | GTTCTTCCCTGATACCCACC |
| 1195 | APTR-004 | TCGGAGGTCAGAGCTCGGCG |
| 1196 | APTR-004 | CTTCGGAGGTCAGAGCTCGG |
| 1197 | APTR-004 | GGTTCTTCCCTGATACCCAC |
| 1198 | ATAD2 | GTCTGGAGCACATCGGCCGG |
| 1199 | ATAD2 | CCACCATCTTCTCTCCCTAC |
| 1200 | ATAD2 | CGCGCACGCCGAGGCCAGTA |
| 1201 | ATAD2 | TCGCGCACGCCGAGGCCAGT |
| 1202 | ATAD2 | CCGGAGAGAGATCCAGCTCC |
| 1203 | ATAD2 | TGTGCAGCTCCAAGCTGCTG |

| | | |
|---|---|---|
| 1204 | ATAD2 | GGTGGTTCTCCGCAGCAGCT |
| 1205 | ATAD5 | TCGGACCACAGAGCTTCGGA |
| 1206 | ATAD5 | ATCGGACCACAGAGCTTCGG |
| 1207 | ATAD5 | GAATTCCCTCCGAAGCTCTG |
| 1208 | ATAD5 | AAGCTCTGTGGTCCGATCTG |
| 1209 | ATAD5 | CAGATCGGACCACAGAGCTT |
| 1210 | ATAD5 | AAAGCAAGCGGACCGCAGAT |
| 1211 | ATAD5 | GGAGGGAATTCTTGTCAATT |
| 1212 | AURKA | CCCACGGTCTACCCACCCGT |
| 1213 | AURKA | CCGACGGGTGGGTAGACCGT |
| 1214 | AURKA | GGTCGCAGGTGGGAGCCGAC |
| 1215 | AURKA | CGACGGGTGGGTAGACCGTG |
| 1216 | AURKA | GACGGGTGGGTAGACCGTGG |
| 1217 | AURKA | CGGCTCCCACCTGCGACCCA |
| 1218 | AURKA | GCTGAGCTCTTGGAAGACTT |
| 1219 | AURKB | CAGATTCAGTTGTTTGCGGG |
| 1220 | AURKB | GACAGGAGGCCAGCTCACCT |
| 1221 | AURKB | GTGGCAGATTCAGTTGTTTG |
| 1222 | AURKB | CCAAGGCACTGCTACTCTCC |
| 1223 | AURKB | CCGGGAGAGTAGCAGTGCCT |
| 1224 | AURKB | GGCGCAAGGCCTGCGACAGG |
| 1225 | AURKB | GTGAGCTGGCCTCCTGTCGC |
| 1226 | BARD1 | ATGAAAGGCTCCTCGCAGAG |
| 1227 | BARD1 | GGAAGCCTCGGGAAACCACA |
| 1228 | BARD1 | GCGGCCCGCCGTCCCAGACG |
| 1229 | BARD1 | GCTGGCCTGCAGCTTCCCTG |
| 1230 | BARD1 | AGGAAGCCTCGGGAAACCAC |
| 1231 | BARD1 | GCAGGCCAGCGACTCGAAAC |
| 1232 | BARD1 | GCGAGGAGCCTTTCATCCGA |
| 1233 | BISPR-002 | AGCTGGGTGCCAGCCAACTG |
| 1234 | BISPR-002 | CAGTTTCCCAGAAAGGAGGT |
| 1235 | BISPR-002 | CCCAGCTGGGAAGGGCCCAA |
| 1236 | BISPR-002 | TCAGTTTCAGTTTCCCAGAA |
| 1237 | BISPR-002 | GTGTCCCAGGCAGCCACAGT |
| 1238 | BISPR-002 | GTTTCAGTTTCCCAGAAAGG |
| 1239 | BISPR-002 | CCCAGGCAGCCACAGTTGGC |
| 1240 | BISPR-007 | TAACACACAGCAGAGGCACA |
| 1241 | BISPR-007 | ACAGTTGCGTGGTGGAGTGA |
| 1242 | BISPR-007 | ACTGAGGTTGCACAGTTGCG |
| 1243 | BISPR-007 | GAGGTTGCACAGTTGCGTGG |
| 1244 | BISPR-007 | GAGGCAGAGGCCATCACTGT |
| 1245 | BISPR-007 | AGAGGCCATCACTGTTGGTC |
| 1246 | BISPR-007 | GCAACCTCAGTTTCCCCAGG |

| | | |
|---|---|---|
| 1247 | BISPR-009 | TAGTACAGATGAATACAGGC |
| 1248 | BISPR-009 | CAGATGAATACAGGCTGGCA |
| 1249 | BISPR-009 | CACTCAGTAACACACAGCAG |
| 1250 | BISPR-009 | AATACAGGCTGGCACGGCCT |
| 1251 | BISPR-009 | GAACACAAAAACCAGTCCCT |
| 1252 | BISPR-009 | TGAACACAAAAACCAGTCCC |
| 1253 | BISPR-009 | GTCATAGTACAGATGAATAC |
| 1254 | BISPR-012 | GGGACTATGGGAGACATGGA |
| 1255 | BISPR-012 | ACATTGTTTGTTGGCTGGGT |
| 1256 | BISPR-012 | GACCGACATTGTTTGTTGGC |
| 1257 | BISPR-012 | ACCGACATTGTTTGTTGGCT |
| 1258 | BISPR-012 | CTCCCATAGTCCCCTCGGTG |
| 1259 | BISPR-012 | GTCCCCACCGAGGGGACTAT |
| 1260 | BISPR-012 | GGGGACTATGGGAGACATGG |
| 1261 | bP-21264C1.2-001 | AAGCCGGCCCGGCTACAGGA |
| 1262 | bP-21264C1.2-001 | CCAGAGGCCGTCCTGTAGCC |
| 1263 | bP-21264C1.2-001 | CGCCCTTGCACCGACCCGCA |
| 1264 | bP-21264C1.2-001 | CCTGTAGCCGGGCCGGCTTG |
| 1265 | bP-21264C1.2-001 | AGGCCGTCCTGTAGCCGGGC |
| 1266 | bP-21264C1.2-001 | CCGGCTACAGGACGGCCTCT |
| 1267 | bP-21264C1.2-001 | TCCGGAAGTGCGGCACCCAG |
| 1268 | bP-21264C1.2-002 | CTAGAACCCGTTTCACTGCC |
| 1269 | bP-21264C1.2-002 | AGAGGAACCCGTGGGCCTCA |
| 1270 | bP-21264C1.2-002 | CTAGAGGCGCAGGAGCAGGT |
| 1271 | bP-21264C1.2-002 | CAGACGCGTGGGCACATTGC |
| 1272 | bP-21264C1.2-002 | GAAACGGGTTCTAGAGGCGC |
| 1273 | bP-21264C1.2-002 | GAGCAGGTAGGAGAGGCCGG |
| 1274 | bP-21264C1.2-002 | CGGGATCCCCTGAGGCCCAC |
| 1275 | bP-21264C1.2-003 | GCAGACGCGTGGGCACATTG |
| 1276 | bP-21264C1.2-003 | GGCAGTGAAACGGGTTCTAG |
| 1277 | bP-21264C1.2-003 | GGGTCGGTGCAAGGGCGAAG |
| 1278 | bP-21264C1.2-003 | CGGCCCCCCGGGATCCCCTG |
| 1279 | bP-21264C1.2-003 | GAGGAACCCGTGGGCCTCAG |
| 1280 | bP-21264C1.2-003 | TCTAGAACCCGTTTCACTGC |
| 1281 | bP-21264C1.2-003 | ACGCGTGGGCACATTGCGGG |
| 1282 | bP-21264C1.2-006 | CCATTAAGCCATCTCCCTCG |
| 1283 | bP-21264C1.2-006 | AGTTTGCCCATCTGTCCCAG |
| 1284 | bP-21264C1.2-006 | GAGAACTGTGTGTTACCACT |
| 1285 | bP-21264C1.2-006 | TTCTGTAAATGCGACCCAGG |
| 1286 | bP-21264C1.2-006 | GTGTTACCACTGGGACAGAT |
| 1287 | bP-21264C1.2-006 | TGTGTTACCACTGGGACAGA |
| 1288 | bP-21264C1.2-006 | AAACTGAGCGTCATGCAGGT |
| 1289 | BRCA1 | CGCTGAGACTTCCTGGACGG |

| | | |
|---|---|---|
| 1290 | BRCA1 | ACTACCTTTACCCAGAGCAG |
| 1291 | BRCA1 | TGAAGGCCTCCTGAGCGCAG |
| 1292 | BRCA1 | TTACCCAGAGCAGAGGGTGA |
| 1293 | BRCA1 | TCGCTGAGACTTCCTGGACG |
| 1294 | BRCA1 | CTCGCTGAGACTTCCTGGAC |
| 1295 | BRCA1 | GTGAAGGCCTCCTGAGCGCA |
| 1296 | BRIP1 | GGGTCGAGGAAAGGTAACGG |
| 1297 | BRIP1 | CAGTCCTGCACACAAGGCCG |
| 1298 | BRIP1 | CCAGTCCTGCACACAAGGCC |
| 1299 | BRIP1 | GGGGAAGTAGCAGCACCCCC |
| 1300 | BRIP1 | TTCCTCCTCCCTCTTCCTGG |
| 1301 | BRIP1 | CGGCCCCAGTCCTGCACACA |
| 1302 | BRIP1 | TGGGGGTGCTGCTACTTCCC |
| 1303 | BUB1 | GCGCGGGGTATTCGAATCGG |
| 1304 | BUB1 | ACCAGACGGACACTTACTGA |
| 1305 | BUB1 | ACATTCCAAACCCAGGAAGG |
| 1306 | BUB1 | CGGGGTATTCGAATCGGCGG |
| 1307 | BUB1 | CGGCGGCGGCTTCTAGTTTG |
| 1308 | BUB1 | GGTTCAGGTTTGGCCGCTGC |
| 1309 | BUB1 | TCTAGTTTGCGGTTCAGGTT |
| 1310 | BUB1B | GGTCGCTTCTGTAGCTCCGA |
| 1311 | BUB1B | CGGAGCTACAGAAGCGACCA |
| 1312 | BUB1B | TACACGACTCCCTAACAAAC |
| 1313 | BUB1B | AGGGAGTCGTGTACGTGCCT |
| 1314 | BUB1B | AAGAAAGCCCAGGCGGTCTG |
| 1315 | BUB1B | TGGTCGCTTCTGTAGCTCCG |
| 1316 | BUB1B | GTTGCGGAAGAAAGCCCAGG |
| 1317 | C11orf71-002 | TTAGTCATCGGGTCCGACAC |
| 1318 | C11orf71-002 | GCCGCCTTGCCTTTAACGAG |
| 1319 | C11orf71-002 | ACCCCTCGTTAAAGGCAAGG |
| 1320 | C11orf71-002 | GATACCCCTCGTTAAAGGCA |
| 1321 | C11orf71-002 | AGCCGCCTTGCCTTTAACGA |
| 1322 | C11orf71-002 | CAGATCAGTCCAGCCTGTGT |
| 1323 | C11orf71-002 | GGAACCCATAACTGAGCCTG |
| 1324 | C11orf72-001 | ATGCGAGGTTTGTTTACTTG |
| 1325 | C11orf72-001 | CCAGCAGAAGGCGACATGCG |
| 1326 | C11orf72-001 | TATAGAAAGTGGAACACACC |
| 1327 | C11orf72-001 | TACTTGTGGCATCAAAGGCT |
| 1328 | C11orf72-001 | GGGAGCCGGCGAGCTAAGGA |
| 1329 | C11orf72-001 | TGGGAGCCGGCGAGCTAAGG |
| 1330 | C11orf72-001 | ACACACCAGGACCCAGCAGA |
| 1331 | C1orf132-001 | TGCGCAAACAGGCTCTCCCC |
| 1332 | C1orf132-001 | CCCGAGATGCGCAGCCGCTG |

| | | |
|---|---|---|
| 1333 | C1orf132-001 | CGCATCTCGGGCCTGCAGCG |
| 1334 | C1orf132-001 | CCGCAGCGGCTGCGCATCTC |
| 1335 | C1orf132-001 | TGCGCATCTCGGGCCTGCAG |
| 1336 | C1orf132-001 | CCCGCCAAGCGCCCCGCTGC |
| 1337 | C1orf132-001 | GCCTGCAGCGGGGCGCTTGG |
| 1338 | C8orf33 | CAGTCGCCACCGGAACTACG |
| 1339 | C8orf33 | GAGAAAGGGGGGTAGCACGA |
| 1340 | C8orf33 | AGTCGCCACCGGAACTACGC |
| 1341 | C8orf33 | CGAGAGCCGCCAACAATGCA |
| 1342 | C8orf33 | GTCGCCACCGGAACTACGCG |
| 1343 | C8orf33 | CGTAGTTCCGGTGGCGACTG |
| 1344 | C8orf33 | GCGCATGGCGGTGAGCGGTG |
| 1345 | CAHM-001 | TTGGCCCGTCGCCCACGGTG |
| 1346 | CAHM-001 | GGCCAGACCCAGCCCCTAGT |
| 1347 | CAHM-001 | AGCCGACTAGGGGCTGGGTC |
| 1348 | CAHM-001 | GTGGGCGACGGGCCAAGACC |
| 1349 | CAHM-001 | TGGGTCTGGCCGTTTAGGGC |
| 1350 | CAHM-001 | CGGGGGCGGCAGCCGACTAG |
| 1351 | CAHM-001 | GCGGCAGCCGACTAGGGGCT |
| 1352 | CCNA2 | CAGGCAGGCACTGCCCAGCG |
| 1353 | CCNA2 | GCGTCTTTGGCTCGCCACGC |
| 1354 | CCNA2 | AGGGCCGAGGAGGTTGCGAA |
| 1355 | CCNA2 | AGGTTGCGAAAGGCGCAGGC |
| 1356 | CCNA2 | GCAACCTCCTCGGCCCTGCG |
| 1357 | CCNA2 | CGTCTTTGGCTCGCCACGCT |
| 1358 | CCNA2 | GAGGAGGTTGCGAAAGGCGC |
| 1359 | CCNB1 | GGGGAGAAGCAGAACACCGG |
| 1360 | CCNB1 | TTCACCAGGCAGCAGCTCAG |
| 1361 | CCNB1 | TGTTGGTTTCTGCTGGGTGT |
| 1362 | CCNB1 | GCTGGGTGTAGGTCCTTGGC |
| 1363 | CCNB1 | AACGGCTGTTGGTTTCTGCT |
| 1364 | CCNB1 | TGCTGCGGCGGAACGGCTGT |
| 1365 | CCNB1 | CACCAGGCAGCAGCTCAGCG |
| 1366 | CCNB2 | AGGGAGGACACTAGCGTACG |
| 1367 | CCNB2 | GGCTCGGAGAGCAGTCCTAA |
| 1368 | CCNB2 | GGGACGCGGACTGAAAAGGG |
| 1369 | CCNB2 | TCGAAGATCCCCAGCGCTGC |
| 1370 | CCNB2 | CTAGCGTACGAGGCGCCGTT |
| 1371 | CCNB2 | GGAGGGACGCGGACTGAAAA |
| 1372 | CCNB2 | GGGAGGGACGCGGACTGAAA |
| 1373 | CDC37L1-AS1-002 | CTAACCGCTAATGGCGCTGT |
| 1374 | CDC37L1-AS1-002 | CGAAGAGCTGCAAGAAAACG |
| 1375 | CDC37L1-AS1-002 | ACTGCAAACCTAACCGCTAA |

| | | |
|---|---|---|
| 1376 | CDC37L1-AS1-002 | TGGGTGGAGAAACGTTTGTT |
| 1377 | CDC37L1-AS1-002 | GTGATTCTAAGGCAGAGTTG |
| 1378 | CDC37L1-AS1-002 | CAGTGATTCTAAGGCAGAGT |
| 1379 | CDC37L1-AS1-002 | CCTAACCGCTAATGGCGCTG |
| 1380 | CDC7 | GCAGGCGCATCCGATCGACT |
| 1381 | CDC7 | GCGCATCCGATCGACTCGGT |
| 1382 | CDC7 | CGCGCCATGGAACAAACCGT |
| 1383 | CDC7 | TCCGATCGACTCGGTAGGTG |
| 1384 | CDC7 | TCCCCACCTACCGAGTCGAT |
| 1385 | CDC7 | AACAAACCGTCGGAAACTCA |
| 1386 | CDC7 | CATCCGATCGACTCGGTAGG |
| 1387 | CDCA5 | GAGCCTAAGTTATGTCTGGG |
| 1388 | CDCA5 | TCGCCTCCCAGACATAACTT |
| 1389 | CDCA5 | ACGGAGCCTAAGTTATGTCT |
| 1390 | CDCA5 | TCTGGGAGGCGAACGCGGTC |
| 1391 | CDCA5 | CCGCTCACCGGAGCGCTGAG |
| 1392 | CDCA5 | TTATGTCTGGGAGGCGAACG |
| 1393 | CDCA5 | GACGGAGCCTAAGTTATGTC |
| 1394 | CECR7-001 | CAGTCCGCAGGACCCCCTCG |
| 1395 | CECR7-001 | CTGCGTCAGCATCCGCGAGG |
| 1396 | CECR7-001 | GCATCCGCGAGGGGGTCCTG |
| 1397 | CECR7-001 | GCTGACGCAGAATGGAGTTG |
| 1398 | CECR7-001 | ATTCTGCGTCAGCATCCGCG |
| 1399 | CECR7-001 | ACGCAGAATGGAGTTGAGGT |
| 1400 | CECR7-001 | TTCTGCGTCAGCATCCGCGA |
| 1401 | CECR7-002 | CGCAGTCTCGGCTTCCGTCG |
| 1402 | CECR7-002 | GAGTCTCGGAGAAGATGTGG |
| 1403 | CECR7-002 | CCGAGACTGCGCTTCGTGCG |
| 1404 | CECR7-002 | AGCGAGTCTCGGAGAAGATG |
| 1405 | CECR7-002 | GGCGTGGTCAGAGCGAGTCT |
| 1406 | CECR7-002 | AGCGCAGTCTCGGCTTCCGT |
| 1407 | CECR7-002 | CCTCGCACGAAGCGCAGTCT |
| 1408 | CECR7-003 | TCCCCCTCCACTTACCGCCG |
| 1409 | CECR7-003 | AGGCCTCGGCGGTAAGTGGA |
| 1410 | CECR7-003 | TGGTGGCTTCCGTTTGTTGG |
| 1411 | CECR7-003 | GAGGAGGTGGCAGGCCTCGG |
| 1412 | CECR7-003 | TGGCTTCCGTTTGTTGGTGG |
| 1413 | CECR7-003 | CCACCTCCTCCACCAACAAA |
| 1414 | CECR7-003 | ATGTGGTGGCTTCCGTTTGT |
| 1415 | CENPE | CACTCGCACGCAGACGGCCA |
| 1416 | CENPE | CGGCCGCACTCGCACGCAGA |
| 1417 | CENPE | AGTTCAGCCTGATAGGATGG |
| 1418 | CENPE | TGGGACCAGTTCAGCCTGAT |

| | | |
|---|---|---|
| 1419 | CENPE | CCCCCCTACCTGCTGTTCAG |
| 1420 | CENPE | CCGCTGAACAGCAGGTAGGG |
| 1421 | CENPE | GAGTGCGGCCGCTGAACAGC |
| 1422 | CENPF | CGGAGAGTACAGTTCTGGCG |
| 1423 | CENPF | GCGGGCGAATTGGGCACCGG |
| 1424 | CENPF | CCAGAACTGTACTCTCCGAG |
| 1425 | CENPF | GGGACGGGAAAACGACCTCT |
| 1426 | CENPF | GTACAGTTCTGGCGCGGCTT |
| 1427 | CENPF | GGGCAGTTTGAATTAGACTC |
| 1428 | CENPF | GGCAGTTTGAATTAGACTCT |
| 1429 | CENPK | TCACAAACTCCAGGTCGCCT |
| 1430 | CENPK | TGGAGTTTGTGACGCTGTGA |
| 1431 | CENPK | ATCACAGCGTCACAAACTCC |
| 1432 | CENPK | GATATAACGGTTGAATCTTG |
| 1433 | CENPK | ACGCTGTGATGGTAAGACCT |
| 1434 | CENPK | TTGTGGAGCTCCAGTCCTGA |
| 1435 | CENPK | AGGTCGCCTAGGCGCTGCGC |
| 1436 | CENPL | CCAAACACAGATCGCCGCTG |
| 1437 | CENPL | CCCAAACACAGATCGCCGCT |
| 1438 | CENPL | AGTGCCCAAGATTACCGGGA |
| 1439 | CENPL | CGGGAGCCTCTCGAGAAGCG |
| 1440 | CENPL | GCGGCCAGAACCGGAAAACA |
| 1441 | CENPL | ACCCAAACACAGATCGCCGC |
| 1442 | CENPL | CTCTTTCCACGCTTCTCGAG |
| 1443 | CENPM | CAAGCTGCCCGGCCTGAACA |
| 1444 | CENPM | CGCACCGCAGGAGCAACGGT |
| 1445 | CENPM | AGGCCGGGCAGCTTGTCCAG |
| 1446 | CENPM | CGGCTGTGATGTCGGTGTTG |
| 1447 | CENPM | GGCGCGCACCGCAGGAGCAA |
| 1448 | CENPM | AGGAGCAACGGTTGGTCCTG |
| 1449 | CENPM | GCCTGCACCTACCAAGATGG |
| 1450 | CENPO | AAAACCAAACCAACCGGTCC |
| 1451 | CENPO | GGCAGGATCGCAAAGCACGC |
| 1452 | CENPO | CAAAGCACGCCGGGACCGGT |
| 1453 | CENPO | GCAGGATCGCAAAGCACGCC |
| 1454 | CENPO | GTCTTCAAAACCAAACCAAC |
| 1455 | CENPO | TGGGTGAGCTAGAAGGGAGA |
| 1456 | CENPO | ATGGCGGGAATTCTCGCTTC |
| 1457 | CEP55 | CCACACCTGATGGTGTGACT |
| 1458 | CEP55 | GCTGCTAGCTGGCCCAAGGG |
| 1459 | CEP55 | GAAGCGGCATCCACACCTGA |
| 1460 | CEP55 | TCAGCTGCTAGCTGGCCCAA |
| 1461 | CEP55 | GTCGGCCGAGTCACACCATC |

| | | |
|---|---|---|
| 1462 | CEP55 | TGGCCCAAGGGAGGCGACCG |
| 1463 | CEP55 | GCGAAGCGCGGCGCTCGCGT |
| 1464 | CH17-189H20.1-001 | TGGGTGAGAGGAGGAGAAGG |
| 1465 | CH17-189H20.1-001 | TCCTGGGTGAGAGGAGGAGA |
| 1466 | CH17-189H20.1-001 | CGGAAGTGATCCTGGGTGAG |
| 1467 | CH17-189H20.1-001 | AAGTGATCCTGGGTGAGAGG |
| 1468 | CH17-189H20.1-001 | AGAACTGGGAAGGGGGCGCG |
| 1469 | CH17-189H20.1-001 | TCCTTCTCCTCCTCTCACCC |
| 1470 | CH17-189H20.1-001 | AAAGCTGAAAGCGTGCGAGG |
| 1471 | CH17-340M24.3-001 | ACCGACTAGGACGAGACAAG |
| 1472 | CH17-340M24.3-001 | AAACGGACGGGCGAACGCGT |
| 1473 | CH17-340M24.3-001 | GACGGGCGAACGCGTCGGGT |
| 1474 | CH17-340M24.3-001 | ACGAGACAAGAGGGACAGGG |
| 1475 | CH17-340M24.3-001 | CCGACTAGGACGAGACAAGA |
| 1476 | CH17-340M24.3-001 | GGGGCCCAGCTAGACCGACT |
| 1477 | CH17-340M24.3-001 | AAAATAAAAATAAAAACGGA |
| 1478 | CH17-373J23.1-001 | AAAGCTCATACTTACCTGGA |
| 1479 | CH17-373J23.1-001 | TCAGCACATCCGGAACGCAA |
| 1480 | CH17-373J23.1-001 | CCTGCGATTTCCCCAAGTGT |
| 1481 | CH17-373J23.1-001 | CACACTTGGGGAAATCGCAG |
| 1482 | CH17-373J23.1-001 | CCCCAAACATGGCCTCACAA |
| 1483 | CH17-373J23.1-001 | CCACACTTGGGGAAATCGCA |
| 1484 | CH17-373J23.1-001 | CCCTGCGATTTCCCCAAGTG |
| 1485 | CH507-513H4.1-001 | CGACACCTAGTGAATGCGGG |
| 1486 | CH507-513H4.1-001 | GGAGAGGAAGAAACAACGCA |
| 1487 | CH507-513H4.1-001 | TGGAGGGCAAGGCATCATCA |
| 1488 | CH507-513H4.1-001 | AAAGCGACACCTAGTGAATG |
| 1489 | CH507-513H4.1-001 | AAGCGACACCTAGTGAATGC |
| 1490 | CH507-513H4.1-001 | GGAGTCCTGGAAAGTCGAGG |
| 1491 | CH507-513H4.1-001 | TTCTGAAACACTCAGACTGG |
| 1492 | CH507-513H4.3-001 | CGACGGGCGGTGTGTACAAA |
| 1493 | CH507-513H4.3-001 | GACCTGTTATTGCTCAATCT |
| 1494 | CH507-513H4.3-001 | TTGTCCCTCTAAGAAGTTGG |
| 1495 | CH507-513H4.3-001 | CATCACGAATGGGGTTCAAC |
| 1496 | CH507-513H4.3-001 | TGGGGGACGCCGACCGCTCG |
| 1497 | CH507-513H4.3-001 | GTGTAGCGCGCGTGCAGCCC |
| 1498 | CH507-513H4.3-001 | GCTTATGACCCGCACTTACT |
| 1499 | CH507-513H4.4-001 | TACCCGCGCCTGCCGGCGTA |
| 1500 | CH507-513H4.4-001 | TACCCTACGCCGGCAGGCGC |
| 1501 | CH507-513H4.4-001 | AACGGGTTACCCGCGCCTGC |
| 1502 | CH507-513H4.4-001 | TGTGCCTACCCTACGCCGGC |
| 1503 | CH507-513H4.4-001 | CCCCGATCCCCATCACGAAT |
| 1504 | CH507-513H4.4-001 | CCATCACGAATGGGGTTCAA |

| | | |
|---|---|---|
| 1505 | CH507-513H4.4-001 | GCGACGGGCGGTGTGTACAA |
| 1506 | CH507-513H4.5-001 | GTGTGCCTACCCTACGCCGG |
| 1507 | CH507-513H4.5-001 | TGTCCCTCTAAGAAGTTGGG |
| 1508 | CH507-513H4.5-001 | CAACGGGTTACCCCCACCGC |
| 1509 | CH507-513H4.5-001 | CGTCCCCCCAACTTCTTAGA |
| 1510 | CH507-513H4.5-001 | CCTACCCTACGCCGGCGGTG |
| 1511 | CH507-513H4.5-001 | GCGTCCCCCCAACTTCTTAG |
| 1512 | CH507-513H4.5-001 | CCCCACCGCCGGCGTAGGGT |
| 1513 | CH507-513H4.6-001 | CAATCGGTAGTAGCGACGGG |
| 1514 | CH507-513H4.6-001 | AGCGTGTGCCTACCCTACGC |
| 1515 | CH507-513H4.6-001 | CATCCAATCGGTAGTAGCGA |
| 1516 | CH507-513H4.6-001 | CCCATTCGTGATGGGGATCG |
| 1517 | CH507-513H4.6-001 | CGCCCGTCGCTACTACCGAT |
| 1518 | CH507-513H4.6-001 | CGTCGCTACTACCGATTGGA |
| 1519 | CH507-513H4.6-001 | CGCGCCTGCCGGCGTAGGGT |
| 1520 | CH507-528H12.1-001 | GAGATAGGAAGATCACCTGA |
| 1521 | CH507-528H12.1-001 | ACCTTCTGAAACACTCAGAC |
| 1522 | CH507-528H12.1-001 | TCCTATCTCAGCCTTCCAAG |
| 1523 | CH507-528H12.1-001 | CAACTTCAGTGTGTAGAAAG |
| 1524 | CH507-528H12.1-001 | TGAGATAGGAAGATCACCTG |
| 1525 | CH507-528H12.1-001 | TACTGTGGAAACAGTTGGAG |
| 1526 | CH507-528H12.1-001 | TGTAGTCTCAGCCACTTGGA |
| 1527 | CHEK1 | GTTTCTAAAGAAGGAGTTCG |
| 1528 | CHEK1 | GGGCAAAGGACAGTCCGGTG |
| 1529 | CHEK1 | ATTTGGATTCCTGCAGTGGT |
| 1530 | CHEK1 | GTGGTGGGCAAAGGACAGTC |
| 1531 | CHEK1 | AGAAACCTTCCCTACTCTAC |
| 1532 | CHEK1 | GACAGTCCGGTGAGGAAGGG |
| 1533 | CHEK1 | GAAGGGCGGCCGGTAGAGTA |
| 1534 | CITF22-1A6.3-001 | GTGTGCACATGTGCACTGTG |
| 1535 | CITF22-1A6.3-001 | GTGTGCACGTGTGCACTGTG |
| 1536 | CITF22-1A6.3-001 | ACATGTGCACTGTGTATGCA |
| 1537 | CITF22-1A6.3-001 | GTGCACTGTGTGTGGATGCA |
| 1538 | CITF22-1A6.3-001 | GTGCACGTGTGCACGGTGTG |
| 1539 | CITF22-1A6.3-001 | GTGCACGTGTGCACTGTGTG |
| 1540 | CITF22-1A6.3-001 | ATGGTGTGTGCACGTGTGCA |
| 1541 | CITF22-49E9.3-001 | CGCACAGACTCAGTGACCAG |
| 1542 | CITF22-49E9.3-001 | CACAGACTCAGTGACCAGAG |
| 1543 | CITF22-49E9.3-001 | GGGATGGGGTCACCTATTGG |
| 1544 | CITF22-49E9.3-001 | GCACAGACTCAGTGACCAGA |
| 1545 | CITF22-49E9.3-001 | GGCAGGTCCTGGGTAATCCC |
| 1546 | CITF22-49E9.3-001 | AGCTCTGGAAAAACAACAGA |
| 1547 | CITF22-49E9.3-001 | AAAAACAACAGAAGGCTGCA |

| | | |
|---|---|---|
| 1548 | CMB9-22P13.1-001 | AGGGTGGCTTCATTCTGTAA |
| 1549 | CMB9-22P13.1-001 | GGCTGGGGTAAGGATGATCA |
| 1550 | CMB9-22P13.1-001 | GGCCCATGTAGCATCCCTGG |
| 1551 | CMB9-22P13.1-001 | CTGCACACTCTTGAAGTGAC |
| 1552 | CMB9-22P13.1-001 | GCGCGCACGTAGGCGACTTG |
| 1553 | CMB9-22P13.1-001 | TTCAAGAGTGTGCAGCTCCC |
| 1554 | CMB9-22P13.1-001 | CATTATCTTTACCTTGCTCT |
| 1555 | CREB3L4 | GTAGGGGACTAAGAGTCTGA |
| 1556 | CREB3L4 | CCTTGCCAGGAGCCACGGTA |
| 1557 | CREB3L4 | TCCCCTACTCCCAGATACAG |
| 1558 | CREB3L4 | GGCAGGGTTGTAAACAAAGG |
| 1559 | CREB3L4 | CATGGCAGGGTTGTAAACAA |
| 1560 | CREB3L4 | ATCTCCCTCTTGCAAAAGCG |
| 1561 | CREB3L4 | AGAGTCTGAGGGTCCACAAA |
| 1562 | CRNDE-003 | GACACGACTAGAGGGGCGGA |
| 1563 | CRNDE-003 | ACGAGGGGACACGACTAGAG |
| 1564 | CRNDE-003 | AGGGGACACGACTAGAGGGG |
| 1565 | CRNDE-003 | GGAGAGGTGAGCCCCCGCCC |
| 1566 | CRNDE-003 | GACTAGAGGGGCGGACGGCG |
| 1567 | CRNDE-003 | GGGCGGACGGCGCGGCCAGA |
| 1568 | CRNDE-003 | GGGGCGGACGGCGCGGCCAG |
| 1569 | CRNDE-004 | GTTACCGACCGTTAGCGCCA |
| 1570 | CRNDE-004 | TTACCGACCGTTAGCGCCAG |
| 1571 | CRNDE-004 | CTGGCGCTAACGGTCGGTAA |
| 1572 | CRNDE-004 | GGGGCTCGCGCCAGCCACGA |
| 1573 | CRNDE-004 | CGCGGACGCTCCCTCGTGGC |
| 1574 | CRNDE-004 | TGGCTGGCGCGAGCCCCCGG |
| 1575 | CRNDE-004 | TGGCGCGAGCCCCCGGCGGG |
| 1576 | CRNDE-007 | CGCCCGACCAGACGTCCCGT |
| 1577 | CRNDE-007 | TAGCCCACGGGACGTCTGGT |
| 1578 | CRNDE-007 | AGCCCACGGGACGTCTGGTC |
| 1579 | CRNDE-007 | GCGCCCGACCAGACGTCCCG |
| 1580 | CRNDE-007 | CGTGGGCCGAACGGACGCGG |
| 1581 | CRNDE-007 | GGGACGTCTGGTCGGGCGCG |
| 1582 | CRNDE-007 | GCCCTAGCCCACGGGACGTC |
| 1583 | CRNDE-009 | ATAAAGATTCAGGATTACAG |
| 1584 | CRNDE-009 | CTGTACTGGCTATTGGAAGG |
| 1585 | CRNDE-009 | TCATAATGGAGCTACTGTAC |
| 1586 | CRNDE-009 | AAGAAAATCAAATGGAGTAT |
| 1587 | CRNDE-009 | AATTGTAATTCCATGAAGGA |
| 1588 | CRNDE-009 | TTATGGTTGATTAACTGAGA |
| 1589 | CRNDE-009 | GGAGCTACTGTACTGGCTAT |
| 1590 | CTA-14H9.5-001 | CAATCTCAGTCCCATAAGAT |

| | | |
|---|---|---|
| 1591 | CTA-14H9.5-001 | TGTTGTGCCTACACACACAC |
| 1592 | CTA-14H9.5-001 | GGCACAACATCCCATCTTAT |
| 1593 | CTA-14H9.5-001 | TCTAGGTCCTGTGTGTGTGT |
| 1594 | CTA-14H9.5-001 | AATGTAGTCCTACCTTTCTT |
| 1595 | CTA-14H9.5-001 | AGGCACAACATCCCATCTTA |
| 1596 | CTA-14H9.5-001 | ACAATCTCAGTCCCATAAGA |
| 1597 | CTA-212A2.2-001 | GGGAGTGACTGCCTCTCCTG |
| 1598 | CTA-212A2.2-001 | GAATTGGCTAACCCCAGGAG |
| 1599 | CTA-212A2.2-001 | CAGTCACTCCCCAGCCAGGA |
| 1600 | CTA-212A2.2-001 | AAAAAGAATTGGCTAACCCC |
| 1601 | CTA-212A2.2-001 | ATCCAGAAACCTTCCTGGCT |
| 1602 | CTA-212A2.2-001 | TCCAGAAACCTTCCTGGCTG |
| 1603 | CTA-212A2.2-001 | AAAATAAAAAATATAGGGCT |
| 1604 | CTA-228A9.3-001 | CCAGCACTTTGGGAGTCCGA |
| 1605 | CTA-228A9.3-001 | GAGGGCAGGCAGATCACCTG |
| 1606 | CTA-228A9.3-001 | CCTCGGACTCCCAAAGTGCT |
| 1607 | CTA-228A9.3-001 | TTACTTACCCAGTAACACTA |
| 1608 | CTA-228A9.3-001 | CCCAGCACTTTGGGAGTCCG |
| 1609 | CTA-228A9.3-001 | CAAAAATTAGCTGGGCATGG |
| 1610 | CTA-228A9.3-001 | TCAGGTGATCTGCCTGCCCT |
| 1611 | CTA-384D8.34-001 | AAGGAGAACCAAGAGATGGT |
| 1612 | CTA-384D8.34-001 | ATGTGCCCGGGGCCTACTGT |
| 1613 | CTA-384D8.34-001 | TGGTTCTCCTTCTACCTCAC |
| 1614 | CTA-384D8.34-001 | GGCCTGGCTAGGCCCACAGT |
| 1615 | CTA-384D8.34-001 | GAAGGAGAACCAAGAGATGG |
| 1616 | CTA-384D8.34-001 | GTAGAAGGAGAACCAAGAGA |
| 1617 | CTA-384D8.34-001 | AATGTGCCCGGGGCCTACTG |
| 1618 | CTA-384D8.35-001 | GGCGTTAGGCTGATTTCACG |
| 1619 | CTA-384D8.35-001 | CGTCATTGAGAAACGGCGTT |
| 1620 | CTA-384D8.35-001 | ATGACATGGAAGTGCTGCTG |
| 1621 | CTA-384D8.35-001 | GAGGAAGACGCTGAGCTCCG |
| 1622 | CTA-384D8.35-001 | ATTTCACGGGGCAGATGACA |
| 1623 | CTA-384D8.35-001 | CGGCGTTAGGCTGATTTCAC |
| 1624 | CTA-384D8.35-001 | CTCCGAGGTGAAAAGGGATG |
| 1625 | CTA-398F10.1-001 | GATTATGCAAAGAGGTACCG |
| 1626 | CTA-398F10.1-001 | CGTGCCCTGATTATGCAAAG |
| 1627 | CTA-398F10.1-001 | TCTTTGCATAATCAGGGCAC |
| 1628 | CTA-398F10.1-001 | CTGTACCAGTGTCACTCTGG |
| 1629 | CTA-398F10.1-001 | CTAACACAAGGTAAGGCCCT |
| 1630 | CTA-398F10.1-001 | AAGATCCCCCAGAGTGACAC |
| 1631 | CTA-398F10.1-001 | GGTACCTCTTTGCATAATCA |
| 1632 | CTA-398F10.2-001 | CCCAGAGAAGTGTTCACTTG |
| 1633 | CTA-398F10.2-001 | TGCATGAAAACCCCCCGATT |

| | | |
|---|---|---|
| 1634 | CTA-398F10.2-001 | CTTGATTACCATGCTCTGGA |
| 1635 | CTA-398F10.2-001 | CGATTTGGCCATCCAGAGCA |
| 1636 | CTA-398F10.2-001 | GTTTCTTGATTACCATGCTC |
| 1637 | CTA-398F10.2-001 | GCTCTGGATGGCCAAATCGG |
| 1638 | CTA-398F10.2-001 | CTCTGGATGGCCAAATCGGG |
| 1639 | CTA-722E9.1-001 | GAATTGGCTCCTTCTCATGG |
| 1640 | CTA-722E9.1-001 | ATGGTGATAATAGTGATGAT |
| 1641 | CTA-722E9.1-001 | CTGAGAAGTCTCACAGTATG |
| 1642 | CTA-722E9.1-001 | GATAGTGATAGTGATAGTGA |
| 1643 | CTA-722E9.1-001 | ACAGTATGTGGTCTGCAAGT |
| 1644 | CTA-722E9.1-001 | AGTGATAATGAGAGTGATGA |
| 1645 | CTA-722E9.1-001 | ATAGTGATGATGGGTGATGA |
| 1646 | CTB-113I20.2-001 | CCGGCCACATCAACTCCCGG |
| 1647 | CTB-113I20.2-001 | TCCCGGGGGATCTAGCAGCG |
| 1648 | CTB-113I20.2-001 | TCCGCGCTGCTAGATCCCCC |
| 1649 | CTB-113I20.2-001 | CCACTTTGTCCACCCAGGCG |
| 1650 | CTB-113I20.2-001 | GCCGGCCACATCAACTCCCG |
| 1651 | CTB-113I20.2-001 | GCTGGACCCAAGCCTCGCCT |
| 1652 | CTB-113I20.2-001 | AGATCCCCCGGGAGTTGATG |
| 1653 | CTB-193M12.5-001 | GAAAACTCGAACGCACCTCT |
| 1654 | CTB-193M12.5-001 | CTTCTCATTGAGATTCCTAG |
| 1655 | CTB-193M12.5-001 | CTTGCCCCAAAGACCCCTCA |
| 1656 | CTB-193M12.5-001 | TCTTAGTGGCTGCATTACCA |
| 1657 | CTB-193M12.5-001 | TTCTGCCTCAGGACCCTGAG |
| 1658 | CTB-193M12.5-001 | AATTCTGCCTCAGGACCCTG |
| 1659 | CTB-193M12.5-001 | GAAGCAGGCTACAGTCTCAG |
| 1660 | CTB-31O20.2-001 | GCGGCCGCCACTTACCACCG |
| 1661 | CTB-31O20.2-001 | GCGTCAGGCTGTGGACGAAG |
| 1662 | CTB-31O20.2-001 | GGTACCTGGCTGGGTCCCAG |
| 1663 | CTB-31O20.2-001 | GACCCAGCCAGGTACCGGTC |
| 1664 | CTB-31O20.2-001 | GCGGCCTCGGTGGTAAGTGG |
| 1665 | CTB-31O20.2-001 | AGCCAGGTACCGGTCCGGCT |
| 1666 | CTB-31O20.2-001 | GAGGCGGCCTCGGTGGTAAG |
| 1667 | CTB-58E17.1-001 | CCGCCGCCCTATGCGTGCCA |
| 1668 | CTB-58E17.1-001 | CTGCCCCAGTCCCTTCGCCG |
| 1669 | CTB-58E17.1-001 | GTCCCGCGGCGAAGGGACTG |
| 1670 | CTB-58E17.1-001 | CGCGCGGCCCTGGCACGCAT |
| 1671 | CTB-58E17.1-001 | CTCGCGCGTCCCGCGGCGAA |
| 1672 | CTB-58E17.1-001 | TCGCCGCGGGACGCGCGAGA |
| 1673 | CTB-58E17.1-001 | TCCGCCGCCCTATGCGTGCC |
| 1674 | CTB-61M7.2-001 | AAATAGGGTAATCCTCCTGT |
| 1675 | CTB-61M7.2-001 | AATTTGTAGATGAACACAGC |
| 1676 | CTB-61M7.2-001 | GAAAGTCATAATCCTACAGG |

| | | |
|---|---|---|
| 1677 | CTB-61M7.2-001 | AGTGAAAGTCATAATCCTAC |
| 1678 | CTB-61M7.2-001 | TTGTAGATGAACACAGCAGG |
| 1679 | CTB-61M7.2-001 | CTTCCCCAAATATAAATGGT |
| 1680 | CTB-61M7.2-001 | TTACCAACCATTTATATTTG |
| 1681 | CTC-242N15.1-002 | AATATACTTACCTCCCCCGA |
| 1682 | CTC-242N15.1-002 | TTCAGATTGTCCTCCTTCGG |
| 1683 | CTC-242N15.1-002 | CTTCAGATTGTCCTCCTTCG |
| 1684 | CTC-242N15.1-002 | GCTGAATGCACTAACTAGAA |
| 1685 | CTC-242N15.1-002 | ACAGCTGAATTGGAGTCACA |
| 1686 | CTC-242N15.1-002 | TGAATTGGAGTCACATGGAC |
| 1687 | CTC-242N15.1-002 | ATGCATGCACCCTCACATGT |
| 1688 | CTC-384G19.1-001 | TATAGTCCCAGCTACTCGGG |
| 1689 | CTC-384G19.1-001 | TGTTGAGGTTAATATGACAC |
| 1690 | CTC-384G19.1-001 | AATTATAACTTTCAAAGGCC |
| 1691 | CTC-384G19.1-001 | CTTCTAACTTCCAAATGTTG |
| 1692 | CTC-384G19.1-001 | CTTTCAAAGGCCGGGTGCAT |
| 1693 | CTC-384G19.1-001 | CAAAAGAAATTAGCCAGGCG |
| 1694 | CTC-384G19.1-001 | AAATACAAAAGAAATTAGCC |
| 1695 | CTC-429P9.2-001 | CATAGAAATTATAAGGATGG |
| 1696 | CTC-429P9.2-001 | GAGATGGGGTTTCCCCATGT |
| 1697 | CTC-429P9.2-001 | TCCCAGCTACCAGGAGGCTG |
| 1698 | CTC-429P9.2-001 | TACCATAGAAATTATAAGGA |
| 1699 | CTC-429P9.2-001 | AAATAAAATACAGGGTGTTA |
| 1700 | CTC-429P9.2-001 | TTGTAATCCCAGCTACCAGG |
| 1701 | CTC-429P9.2-001 | GTGCTGGGATTACAGGTTAC |
| 1702 | CTC-436P18.3-002 | GCTGGACATCATTCACCACT |
| 1703 | CTC-436P18.3-002 | CTTGAACTCTAAGGGCCATG |
| 1704 | CTC-436P18.3-002 | TAACTTATTAATGGTAAGGC |
| 1705 | CTC-436P18.3-002 | TTTATAGAGGTGAGTCCTCA |
| 1706 | CTC-436P18.3-002 | ATCTCAGTCCCTTAAAAACA |
| 1707 | CTC-436P18.3-002 | TCATATAAATAGCATCTGGG |
| 1708 | CTC-436P18.3-002 | GCTTCATATAAATAGCATCT |
| 1709 | CTC-444N24.11-001 | AAAGCCTAATTAGGTAGACA |
| 1710 | CTC-444N24.11-001 | CCTAATTAGGTAGACAAGGA |
| 1711 | CTC-444N24.11-001 | ATTAGGTAGACAAGGATGGG |
| 1712 | CTC-444N24.11-001 | CTAATTAGGTAGACAAGGAT |
| 1713 | CTC-444N24.11-001 | CAAGGATGGGAGGCTGAGGT |
| 1714 | CTC-444N24.11-001 | GAGTTACACAAAGCCTAATT |
| 1715 | CTC-444N24.11-001 | AGAAACCAAATACAGATAAA |
| 1716 | CTC-444N24.8-001 | GCTTTGTGGTAAACGCGCTG |
| 1717 | CTC-444N24.8-001 | CTTTGTGGTAAACGCGCTGA |
| 1718 | CTC-444N24.8-001 | AGAAAAGACGGAGAATGTCT |
| 1719 | CTC-444N24.8-001 | GTAAATGGCCATACACAGCA |

| | | |
|---|---|---|
| 1720 | CTC-444N24.8-001 | AGTAACATGACAGAAATTAG |
| 1721 | CTC-444N24.8-001 | TACTCAAACCAGACACATTG |
| 1722 | CTC-444N24.8-001 | GTAACATGACAGAAATTAGG |
| 1723 | CTC-542B22.2-001 | ATGAGAATTCTGTCACAATG |
| 1724 | CTC-542B22.2-001 | AGTTAGACCCACAGCTACCC |
| 1725 | CTC-542B22.2-001 | GACATGACCTGGGTAGCTGT |
| 1726 | CTC-542B22.2-001 | TGTCACAATGAGGACTGTTT |
| 1727 | CTC-542B22.2-001 | TAGACAGCAGAGACATGACC |
| 1728 | CTC-542B22.2-001 | GAGTGACTTGGTCAGAAGGC |
| 1729 | CTC-542B22.2-001 | TCTGGAGTGCCATTCCTGCT |
| 1730 | CTC-543D15.8-001 | CCCACGCTGAGCCCACCCCA |
| 1731 | CTC-543D15.8-001 | TTCCATCTGCTTGGCTGCGG |
| 1732 | CTC-543D15.8-001 | GGCTGCGGAGGACCTTGGGG |
| 1733 | CTC-543D15.8-001 | CCTTGGGGTGGGCTCAGCGT |
| 1734 | CTC-543D15.8-001 | GCTGCGGAGGACCTTGGGGT |
| 1735 | CTC-543D15.8-001 | CTTGGCTGCGGAGGACCTTG |
| 1736 | CTC-543D15.8-001 | TGGGGGCTGCAGGCAGAAAT |
| 1737 | CTD-2007H13.3-001 | AGGCCGCGACCAGAGTCCAG |
| 1738 | CTD-2007H13.3-001 | CACTTCCCGGAATTCAACGC |
| 1739 | CTD-2007H13.3-001 | GCTTGGGTGCCCGCCCTTAG |
| 1740 | CTD-2007H13.3-001 | AAGCGAGGATCAGCGGACTA |
| 1741 | CTD-2007H13.3-001 | GCTGTCGGACTCTGACTGCG |
| 1742 | CTD-2007H13.3-001 | ATTCAACGCCGGGCCGCTAA |
| 1743 | CTD-2007H13.3-001 | GGCACCCAAGCGAGGATCAG |
| 1744 | CTD-2012K14.6-002 | GATCGAGTTGCACAATCCTG |
| 1745 | CTD-2012K14.6-002 | ACTCGATCTAGAGCGCCAGT |
| 1746 | CTD-2012K14.6-002 | TAAAGCGGGAGGGAAACTCA |
| 1747 | CTD-2012K14.6-002 | CTCGATCTAGAGCGCCAGTA |
| 1748 | CTD-2012K14.6-002 | ATAAACTCATTCTGCCCTAC |
| 1749 | CTD-2012K14.6-002 | GCCCATGCGATGAAATATGA |
| 1750 | CTD-2012K14.6-002 | TGCCCATGCGATGAAATATG |
| 1751 | CTD-2013M15.1-001 | ATCAGTGCTCAATGTTGCCA |
| 1752 | CTD-2013M15.1-001 | TTACTTCCTAGATGGGATGA |
| 1753 | CTD-2013M15.1-001 | CGAGGTGCCAGGATTGCAGA |
| 1754 | CTD-2013M15.1-001 | TGCCCATCATCTGGGATATG |
| 1755 | CTD-2013M15.1-001 | TCCTAGATGGGATGACGGCC |
| 1756 | CTD-2013M15.1-001 | GCTGCCCAGTCTGGGAAGTG |
| 1757 | CTD-2013M15.1-001 | CTGGCAGATCACTCGCAGTT |
| 1758 | CTD-2013N17.7-001 | TCTATGTAACAAAATATCTG |
| 1759 | CTD-2013N17.7-001 | TTTAAAAATAAATCTACCTG |
| 1760 | CTD-2013N17.7-001 | TTCATGAGATTGAAAATACA |
| 1761 | CTD-2013N17.7-001 | TGAGATTGAAAATACAGGGC |
| 1762 | CTD-2013N17.7-001 | AAATAAATCTACCTGAGGCC |

| | | |
|---|---|---|
| 1763 | CTD-2013N17.7-001 | TGAAAATACAGGGCCGGGCG |
| 1764 | CTD-2013N17.7-001 | ATTCATGAGATTGAAAATAC |
| 1765 | CTD-2023M8.1-001 | ATCTTCTCGCTAACAACTGG |
| 1766 | CTD-2023M8.1-001 | AGCTGTAACACTTGCTGCTG |
| 1767 | CTD-2023M8.1-001 | TTTCTCAATCCAAGGGTTCC |
| 1768 | CTD-2023M8.1-001 | AGCTCGATTAGAGAAATGCA |
| 1769 | CTD-2023M8.1-001 | TCTATCTTCTCGCTAACAAC |
| 1770 | CTD-2023M8.1-001 | TCGTGCGGCTGCACACGCTT |
| 1771 | CTD-2023M8.1-001 | GGCTTCAAGGAATGGAGCCA |
| 1772 | CTD-2050E21.1-001 | TATGTGCATGCATATGCGTG |
| 1773 | CTD-2050E21.1-001 | ATACCAACGACAGTCTGGAA |
| 1774 | CTD-2050E21.1-001 | GGGAAAAAGACAGCACACAA |
| 1775 | CTD-2050E21.1-001 | AATACCAACGACAGTCTGGA |
| 1776 | CTD-2050E21.1-001 | AAAAATACAAAAAAGTAGCC |
| 1777 | CTD-2050E21.1-001 | TTTCCCTTCCAGACTGTCGT |
| 1778 | CTD-2050E21.1-001 | AAAAAAGTAGCCGGGTGTGG |
| 1779 | CTD-2054N24.2-001 | AAAACACGCGAGCATCTCAG |
| 1780 | CTD-2054N24.2-001 | CGCCTGGGCTCAAACAACGC |
| 1781 | CTD-2054N24.2-001 | GTTTGAGCCCAGGCGCGGAA |
| 1782 | CTD-2054N24.2-001 | GCGTTGTTTGAGCCCAGGCG |
| 1783 | CTD-2054N24.2-001 | TGTTTGAGCCCAGGCGCGGA |
| 1784 | CTD-2054N24.2-001 | CATGGGTCCCAAGAGGGCTC |
| 1785 | CTD-2054N24.2-001 | CTCCAGCGTTGTTTGAGCCC |
| 1786 | CTD-2117L12.1-001 | TTAAGAACAAAAGAAAAGGG |
| 1787 | CTD-2117L12.1-001 | ACTAGGGAGTGTCAGACAGT |
| 1788 | CTD-2117L12.1-001 | CACTAGGGAGTGTCAGACAG |
| 1789 | CTD-2117L12.1-001 | CATTAAGAACAAAAGAAAAG |
| 1790 | CTD-2117L12.1-001 | AAAGGGGTAACGGACGCACC |
| 1791 | CTD-2117L12.1-001 | GAAATCATCCGTCTTCTGCG |
| 1792 | CTD-2117L12.1-001 | AGAACAAAAGAAAAGGGGGG |
| 1793 | CTD-2134A5.4-001 | ATTGTCTAAATCCGGCACCC |
| 1794 | CTD-2134A5.4-001 | CGGCACCCAGGACGACACAC |
| 1795 | CTD-2134A5.4-001 | GTTCCTGCCCAAAGCAGGAG |
| 1796 | CTD-2134A5.4-001 | ACCTGCTGTGTGGCAGACAA |
| 1797 | CTD-2134A5.4-001 | AATATTTAGCTTTCCCTTGG |
| 1798 | CTD-2134A5.4-001 | TGCCCAAAGCAGGAGGGGAA |
| 1799 | CTD-2134A5.4-001 | GACAATATTTAGCTTTCCCT |
| 1800 | CTD-2184D3.6-001 | ACAAAGGTCACACCAGTCCG |
| 1801 | CTD-2184D3.6-001 | AGGTCACACCAGTCCGAGGT |
| 1802 | CTD-2184D3.6-001 | AAAATTACAACCTTGCACAT |
| 1803 | CTD-2184D3.6-001 | GTTGCAGCAGCCTATGTGCA |
| 1804 | CTD-2184D3.6-001 | AACCATGACAAATAGAGCTG |
| 1805 | CTD-2184D3.6-001 | CATGACAAATAGAGCTGGGG |

| | | |
|---|---|---|
| 1806 | CTD-2184D3.6-001 | TAAATGTCTATCATTTGTGT |
| 1807 | CTD-2186M15.3-001 | TGGGTAGTGAAGGGGAAACG |
| 1808 | CTD-2186M15.3-001 | TCCAGCCCTGGGTAGTGAAG |
| 1809 | CTD-2186M15.3-001 | CGTTTCCCCTTCACTACCCA |
| 1810 | CTD-2186M15.3-001 | CTCCAGCCCTGGGTAGTGAA |
| 1811 | CTD-2186M15.3-001 | GAAGGAGGGAGGGCACGTCA |
| 1812 | CTD-2186M15.3-001 | AGTGAAGGGGAAACGGGGAA |
| 1813 | CTD-2186M15.3-001 | AAGGAGGGAGGGCACGTCAC |
| 1814 | CTD-2192J16.26-001 | CAGGCATGAGCCACTGCGCC |
| 1815 | CTD-2192J16.26-001 | CAAGACCAGCCTGACCAACA |
| 1816 | CTD-2192J16.26-001 | AAAAAATTAGCCGAGCATGG |
| 1817 | CTD-2192J16.26-001 | GAGATGAGGTTTCACCATGT |
| 1818 | CTD-2192J16.26-001 | CCAGGTGTGGCCGGGCGCAG |
| 1819 | CTD-2192J16.26-001 | CTTTGGGAGGCTGAGGCAAG |
| 1820 | CTD-2192J16.26-001 | TGTAATCCCAGCTACTCGAG |
| 1821 | CTD-2227E11.1-001 | ACTGACCCACTATAGAGAGC |
| 1822 | CTD-2227E11.1-001 | GACTGCTCTCTCATCCCTAG |
| 1823 | CTD-2227E11.1-001 | AAGCTCATAAAACCAAAACA |
| 1824 | CTD-2227E11.1-001 | CAAATTGTAGTCCTGTTCCA |
| 1825 | CTD-2227E11.1-001 | CAGGGCAATGTCCATGGAAC |
| 1826 | CTD-2227E11.1-001 | CATGGAACAGGACTACAATT |
| 1827 | CTD-2227E11.1-001 | AGTGGAATGGAGGTTTGGGT |
| 1828 | CTD-2231E14.5-001 | AGACTAGAGGAGGCCAGGTG |
| 1829 | CTD-2231E14.5-001 | CTAGAGGAGGCCAGGTGCGG |
| 1830 | CTD-2231E14.5-001 | TGAGCTCAGGAGTTCAAGAC |
| 1831 | CTD-2231E14.5-001 | CAAGACTGGCCTGGGCAACA |
| 1832 | CTD-2231E14.5-001 | CAGGTGTGAGCCACCGCACC |
| 1833 | CTD-2231E14.5-001 | CTTTGGGAGGCTGAGGCAGG |
| 1834 | CTD-2231E14.5-001 | TAAAAATACAAAAATTAGCT |
| 1835 | CTD-2232E5.2-001 | ACCTCGTGATCCACCCACCT |
| 1836 | CTD-2232E5.2-001 | CAGGTGTGAGTCACCAAGCC |
| 1837 | CTD-2232E5.2-001 | GACCATCCTGGCTAACACAG |
| 1838 | CTD-2232E5.2-001 | TACATTCTTAGGGCCAGGCT |
| 1839 | CTD-2232E5.2-001 | GCCGAGGTGGGTGGATCACG |
| 1840 | CTD-2232E5.2-001 | TCAGGAGTCCAAGACCATCC |
| 1841 | CTD-2232E5.2-001 | TATGAAAAATTAGCCAGGTG |
| 1842 | CTD-2245F17.9-001 | CCACCGTCCTAGCAGCAAGA |
| 1843 | CTD-2245F17.9-001 | TGCCACAGCCAGGATCATGT |
| 1844 | CTD-2245F17.9-001 | TTGCCACAGCCAGGATCATG |
| 1845 | CTD-2245F17.9-001 | CCCAGCTCCATCTTGCTGCT |
| 1846 | CTD-2245F17.9-001 | CTAGCAGCAAGATGGAGCTG |
| 1847 | CTD-2245F17.9-001 | AGTTTCTGTTTCAGTCTGTA |
| 1848 | CTD-2245F17.9-001 | GCTCCATCTTGCTGCTAGGA |

| | | |
|---|---|---|
| 1849 | CTD-2251F13.1-002 | TGGGAAACTTACGATCAAGT |
| 1850 | CTD-2251F13.1-002 | GATACTTAGTGAGTATTCCA |
| 1851 | CTD-2251F13.1-002 | ACGATCAAGTCGGAAAGTGA |
| 1852 | CTD-2251F13.1-002 | GATCAAGTCGGAAAGTGAAG |
| 1853 | CTD-2251F13.1-002 | CGATCAAGTCGGAAAGTGAA |
| 1854 | CTD-2251F13.1-002 | TAAAGCAGTTAGTGCTGCCA |
| 1855 | CTD-2251F13.1-002 | GCAAGCACATCTTATCATGG |
| 1856 | CTD-2270L9.4-001 | GAGGTGAACCACGCCGTGGA |
| 1857 | CTD-2270L9.4-001 | AGAGGTGAACCACGCCGTGG |
| 1858 | CTD-2270L9.4-001 | CGTGGAGGGTGAGCCGCTCG |
| 1859 | CTD-2270L9.4-001 | GCCCAGCCGAGAGCAGGGGA |
| 1860 | CTD-2270L9.4-001 | CTCTGCCCAGCCGAGAGCAG |
| 1861 | CTD-2270L9.4-001 | CCTCTGCCCAGCCGAGAGCA |
| 1862 | CTD-2270L9.4-001 | GCGGGACCCTCCCCTGCTCT |
| 1863 | CTD-2270L9.4-003 | ACTTAAGAGTTGGCTCCGGG |
| 1864 | CTD-2270L9.4-003 | GCGAGCCTGCGAGAGCACCG |
| 1865 | CTD-2270L9.4-003 | AGCACCGAGGCTAGCCTCCG |
| 1866 | CTD-2270L9.4-003 | GGCGTCCAGAACTTAAGAGT |
| 1867 | CTD-2270L9.4-003 | CAGAACTTAAGAGTTGGCTC |
| 1868 | CTD-2270L9.4-003 | GTTGGCTCCGGGCGGCAACG |
| 1869 | CTD-2270L9.4-003 | GTTTCTGGGGTTCGCCTCGG |
| 1870 | CTD-2270P14.1-001 | AAGCTGAACTCCAGCAACGG |
| 1871 | CTD-2270P14.1-001 | GACTCTGAGGAGGCCCCGAG |
| 1872 | CTD-2270P14.1-001 | CCTAAGCTGAACTCCAGCAA |
| 1873 | CTD-2270P14.1-001 | CGTTGCTGGAGTTCAGCTTA |
| 1874 | CTD-2270P14.1-001 | GCTGGAGTTCAGCTTAGGGA |
| 1875 | CTD-2270P14.1-001 | CCGAAGCGCTCCGCCGCTCG |
| 1876 | CTD-2270P14.1-001 | CCGTTGCTGGAGTTCAGCTT |
| 1877 | CTD-2296D1.5-001 | AGAGAAGCTAAGACTCACAG |
| 1878 | CTD-2296D1.5-001 | CTTTAAAGCTCCAACATGTG |
| 1879 | CTD-2296D1.5-001 | TAAAGCTCCAACATGTGAGG |
| 1880 | CTD-2296D1.5-001 | CAGCTGTATCAGGCCAGGCG |
| 1881 | CTD-2296D1.5-001 | TGAGGAGGTTCAGCCTTGAG |
| 1882 | CTD-2296D1.5-001 | AGATACAGCTGTATCAGGCC |
| 1883 | CTD-2296D1.5-001 | TGTGAGGAGGTTCAGCCTTG |
| 1884 | CTD-2298J14.2-001 | ACCCTTGACTTCTTACAGCT |
| 1885 | CTD-2298J14.2-001 | CAATGAAAGTCTAGGCGATG |
| 1886 | CTD-2298J14.2-001 | AATCACGTTTCCTTAATGCT |
| 1887 | CTD-2298J14.2-001 | ACCTAGCTGTAAGAAGTCAA |
| 1888 | CTD-2298J14.2-001 | CCCGAAGACCTTGCATCGCT |
| 1889 | CTD-2298J14.2-001 | ATAGGAACCAATGAAAGTCT |
| 1890 | CTD-2298J14.2-001 | ACCCGAAGACCTTGCATCGC |
| 1891 | CTD-2306M5.1-003 | GCCTTTACAGTGAGGTATGG |

| | | |
|---|---|---|
| 1892 | CTD-2306M5.1-003 | TATTGAATGTATTAGCCCTG |
| 1893 | CTD-2306M5.1-003 | GTATTGAATGTATTAGCCCT |
| 1894 | CTD-2306M5.1-003 | CATGCCTTTACAGTGAGGTA |
| 1895 | CTD-2306M5.1-003 | TGAAACATGCCTTTACAGTG |
| 1896 | CTD-2306M5.1-003 | TTTACAAAAATACCTAAGTA |
| 1897 | CTD-2306M5.1-003 | CAGCGAAGGGAGATAGGATG |
| 1898 | CTD-2313J17.6-001 | AGGATTACCTAAGATAACGC |
| 1899 | CTD-2313J17.6-001 | CTGAACGTTTGTCTTGAGCA |
| 1900 | CTD-2313J17.6-001 | CTTGCTCAAGACAAACGTTC |
| 1901 | CTD-2313J17.6-001 | GAAAGACAAGAGCAGCTCCA |
| 1902 | CTD-2313J17.6-001 | CCCAGCGCCACCCTGCACTG |
| 1903 | CTD-2313J17.6-001 | AGCAAGGCCTGCGTTATCTT |
| 1904 | CTD-2313J17.6-001 | CCCGGGTCCGCAGTGCAGGG |
| 1905 | CTD-2315E11.1-001 | GACCCACCGGGACCCGCGAG |
| 1906 | CTD-2315E11.1-001 | GGGACCCGCGAGCGGGATCA |
| 1907 | CTD-2315E11.1-001 | CCCAGCCCTGATCCCGCTCG |
| 1908 | CTD-2315E11.1-001 | CTGATCCCGCTCGCGGGTCC |
| 1909 | CTD-2315E11.1-001 | TCCCGCTCGCGGGTCCCGGT |
| 1910 | CTD-2315E11.1-001 | ATCCCGCTCGCGGGTCCCGG |
| 1911 | CTD-2315E11.1-001 | GCGAGCGGGATCAGGGCTGG |
| 1912 | CTD-2319I12.2-001 | GAGTTGGAGAGGGTATGCTG |
| 1913 | CTD-2319I12.2-001 | CCGAATCTGAAACTCTAGTA |
| 1914 | CTD-2319I12.2-001 | GTCCAGGAACAAACTCTGGG |
| 1915 | CTD-2319I12.2-001 | TGGAGAGGGTATGCTGTGGA |
| 1916 | CTD-2319I12.2-001 | CTAGAGTTTCAGATTCGGGC |
| 1917 | CTD-2319I12.2-001 | AGTCAGGGCTCAAGAAGGAG |
| 1918 | CTD-2319I12.2-001 | ATTCGGGCTGGAGTTGGAGA |
| 1919 | CTD-2325M2.1-001 | TGGGGGTTGCTGTTTGAAAG |
| 1920 | CTD-2325M2.1-001 | CAAGTGATTTCCATGTCAAG |
| 1921 | CTD-2325M2.1-001 | AGCCTGTTCACCACTTGACA |
| 1922 | CTD-2325M2.1-001 | CTAAGAAGAAAAGTAAATTG |
| 1923 | CTD-2325M2.1-001 | GTAAATTGGGGAAAGAGCAT |
| 1924 | CTD-2325M2.1-001 | GGTTGCTGTTTGAAAGAGGA |
| 1925 | CTD-2325M2.1-001 | CACTTGACATGGAAATCACT |
| 1926 | CTD-2340E1.2-001 | GCACGCTCTATCTCTCCAGA |
| 1927 | CTD-2340E1.2-001 | GGTAAAATAGAGTTACATAG |
| 1928 | CTD-2340E1.2-001 | TTAACAGGTCAGGCCAGGTG |
| 1929 | CTD-2340E1.2-001 | GGAAAGCTGAAGTAAGCTGC |
| 1930 | CTD-2340E1.2-001 | TGATATTAACAGGTCAGGCC |
| 1931 | CTD-2340E1.2-001 | TGGCTTGATATTAACAGGTC |
| 1932 | CTD-2340E1.2-001 | ACATCGTTTCTTAGCAGTAA |
| 1933 | CTD-2358C21.5-001 | AATCCCATGAAACTCAATGG |
| 1934 | CTD-2358C21.5-001 | AAATCCCATGAAACTCAATG |

| | | |
|---|---|---|
| 1935 | CTD-2358C21.5-001 | AGGCTCACTACACCAGCTGA |
| 1936 | CTD-2358C21.5-001 | GTGAAGCACTCAGGTTCCCA |
| 1937 | CTD-2358C21.5-001 | TCCCAGGGCCTGACTACTTG |
| 1938 | CTD-2358C21.5-001 | TCCACAAGTAGTCAGGCCCT |
| 1939 | CTD-2358C21.5-001 | TCCGTAGACTCAAGTTCTCT |
| 1940 | CTD-2383I20.1-001 | GGAGTATTACCTGTGACGCA |
| 1941 | CTD-2383I20.1-001 | TATGTCCAAGACCCCAGTAT |
| 1942 | CTD-2383I20.1-001 | TGTCCAAGACCCCAGTATGG |
| 1943 | CTD-2383I20.1-001 | ACAGGTAATACTCCTGCAGA |
| 1944 | CTD-2383I20.1-001 | ATTACCTGTGACGCAGGGTA |
| 1945 | CTD-2383I20.1-001 | ATGTCCAAGACCCCAGTATG |
| 1946 | CTD-2383I20.1-001 | CTATGTCCAAGACCCCAGTA |
| 1947 | CTD-2410N18.3-001 | AGCTGAGTTCATCACCAGAG |
| 1948 | CTD-2410N18.3-001 | AGCCTGTAACTGGTTCCCCT |
| 1949 | CTD-2410N18.3-001 | TGTAACTGGTTCCCCTGGGA |
| 1950 | CTD-2410N18.3-001 | CACCAGAGCGGCCTTCCCAG |
| 1951 | CTD-2410N18.3-001 | GTTTCCGTGATCGGACTGCC |
| 1952 | CTD-2410N18.3-001 | TGAACTCAGCTCCTCACCTG |
| 1953 | CTD-2410N18.3-001 | TCACCAGAGCGGCCTTCCCA |
| 1954 | CTD-2538C1.2-001 | GGACCGCAGTATTCCCACGC |
| 1955 | CTD-2538C1.2-001 | CTTCCTGCGTGGGAATACTG |
| 1956 | CTD-2538C1.2-001 | ATTCACAGCTGCCTGAAAGG |
| 1957 | CTD-2538C1.2-001 | TGTGAATGACAAACGGTCGC |
| 1958 | CTD-2538C1.2-001 | AGGGCACTCTCGCGGGACAA |
| 1959 | CTD-2538C1.2-001 | TCGCGGGACAATGGGTGTCA |
| 1960 | CTD-2538C1.2-001 | TTTGGTCCAAGCTTCCTGCG |
| 1961 | CTD-2553L13.10-001 | ATGTTTATCTCCCGGGACGT |
| 1962 | CTD-2553L13.10-001 | ACCTCAAAGACCATACAGGG |
| 1963 | CTD-2553L13.10-001 | GGGTGGACAGCGTAGTAGCA |
| 1964 | CTD-2553L13.10-001 | AAGACCTCAAAGACCATACA |
| 1965 | CTD-2553L13.10-001 | AATGTTTATCTCCCGGGACG |
| 1966 | CTD-2553L13.10-001 | AGGGTGGACAGCGTAGTAGC |
| 1967 | CTD-2553L13.10-001 | ACTACGCTGTCCACCCTGTA |
| 1968 | CTD-2561J22.5-001 | ACCCTTGGAAAGAAGACACA |
| 1969 | CTD-2561J22.5-001 | TTGGGACGCGCCAGAAATGG |
| 1970 | CTD-2561J22.5-001 | ATCAGTTCACCGCGTGATGG |
| 1971 | CTD-2561J22.5-001 | GACCCTTGGAAAGAAGACAC |
| 1972 | CTD-2561J22.5-001 | GAAGAGCTAACCTCGAGCTC |
| 1973 | CTD-2561J22.5-001 | CTTGCTCCAGCCAGAGCTCG |
| 1974 | CTD-2561J22.5-001 | CTGGCTGGAGCAAGAGACAA |
| 1975 | CTD-2571L23.8-001 | GTTAAGGCTTAAAAAAACAG |
| 1976 | CTD-2571L23.8-001 | CATGTGTTCTAGTATCTTGG |
| 1977 | CTD-2571L23.8-001 | AGCTTGAACCCAGGGGACGG |

| | | |
|---|---|---|
| 1978 | CTD-2571L23.8-001 | CGAGAATAGCTTGAACCCAG |
| 1979 | CTD-2571L23.8-001 | GGCTTAAAAAAACAGAGGCC |
| 1980 | CTD-2571L23.8-001 | AATAGCTTGAACCCAGGGGA |
| 1981 | CTD-2571L23.8-001 | CCGAGAATAGCTTGAACCCA |
| 1982 | CTD-2619J13.14-001 | TGAGAACAGCCCACTCCTGG |
| 1983 | CTD-2619J13.14-001 | AAAACTGGAATTGAGCTGAG |
| 1984 | CTD-2619J13.14-001 | AGGATTATTCTCGCTTGCTG |
| 1985 | CTD-2619J13.14-001 | TGGAGGCACCAAGACCTGCG |
| 1986 | CTD-2619J13.14-001 | GGAGGCACCAAGACCTGCGA |
| 1987 | CTD-2619J13.14-001 | ACCAAGACCTGCGAGGGCAC |
| 1988 | CTD-2619J13.14-001 | GTGAGAACAGCCCACTCCTG |
| 1989 | CTD-2619J13.14-002 | TGGTCCGCAGAAACGAACGA |
| 1990 | CTD-2619J13.14-002 | CGTTCGTTTCTGCGGACCAG |
| 1991 | CTD-2619J13.14-002 | GCGGCAGTGGGATCATTGGG |
| 1992 | CTD-2619J13.14-002 | CGCGCCTTCGTTCGTTTCTG |
| 1993 | CTD-2619J13.14-002 | TGGGATCATTGGGCGGTGGG |
| 1994 | CTD-2619J13.14-002 | CAGTGGGATCATTGGGCGGT |
| 1995 | CTD-2619J13.14-002 | TTTGCGGCAGTGGGATCATT |
| 1996 | CTD-2619J13.14-201 | CCCTCATCTCCCGGACAGGG |
| 1997 | CTD-2619J13.14-201 | CACCTCCCTTCCGGACAGGG |
| 1998 | CTD-2619J13.14-201 | CGCCCTGTCCGGGAGATGAG |
| 1999 | CTD-2619J13.14-201 | AGCCGCCCTGTCCGGAAGGG |
| 2000 | CTD-2619J13.14-201 | CGCCCCTCATCTCCCGGACA |
| 2001 | CTD-2619J13.14-201 | CACCTCCCTCCCAGATGGGG |
| 2002 | CTD-2619J13.14-201 | CCCCACCTCCCTTCCGGACA |
| 2003 | CTD-3075F15.1-001 | CAGCCACAAGGCCCTTCGCT |
| 2004 | CTD-3075F15.1-001 | GGGCCCAGCGAAGGGCCTTG |
| 2005 | CTD-3075F15.1-001 | CTCGCGCCTAGGCGGCCGGT |
| 2006 | CTD-3075F15.1-001 | CCCAATAGGAAACAGCCACA |
| 2007 | CTD-3075F15.1-001 | TCCCCGCCCTCGCGCCTAGG |
| 2008 | CTD-3075F15.1-001 | CCTTCGCTGGGCCCGCCCAC |
| 2009 | CTD-3075F15.1-001 | ACAGCCACAAGGCCCTTCGC |
| 2010 | CTD-3131K8.2-001 | AAGAAGAGGACGTCCTATCG |
| 2011 | CTD-3131K8.2-001 | GACGTCCTATCGAGGTGGCA |
| 2012 | CTD-3131K8.2-001 | TCAACCTTGCTGTATCAGGG |
| 2013 | CTD-3131K8.2-001 | CAACCTTGCTGTATCAGGGA |
| 2014 | CTD-3131K8.2-001 | CGGCCCTCCCTGATACAGCA |
| 2015 | CTD-3131K8.2-001 | TGCTCAACCTTGCTGTATCA |
| 2016 | CTD-3131K8.2-001 | GGAGAGGGGAGCTCAAGAAG |
| 2017 | CTD-3220F14.1-002 | AGGTCCGCCTGTGTGCCCGT |
| 2018 | CTD-3220F14.1-002 | GGCCAGGGTAGTTACCAGGA |
| 2019 | CTD-3220F14.1-002 | GGCAACGGTGGCTTTCACTG |
| 2020 | CTD-3220F14.1-002 | GCAGCCCACGGGCACACAGG |

| | | |
|---|---|---|
| 2021 | CTD-3220F14.1-002 | GACCGTCCTGGTAACTACCC |
| 2022 | CTD-3220F14.1-002 | GGCTTTCACTGTGGCCCAAG |
| 2023 | CTD-3220F14.1-002 | CTCAGACGCACGATCACACA |
| 2024 | CTD-3220F14.1-003 | CAGCAGGAAGAAACCCCAAG |
| 2025 | CTD-3220F14.1-003 | TTTCACTTTGACCCAAGCGG |
| 2026 | CTD-3220F14.1-003 | TCCCTGACACTTGGCGAAGT |
| 2027 | CTD-3220F14.1-003 | GGCTTTCACTTTGACCCAAG |
| 2028 | CTD-3220F14.1-003 | GGCCAGGGCAGTTACCAGGA |
| 2029 | CTD-3220F14.1-003 | AGAGCAGCCCACGGGCACAC |
| 2030 | CTD-3220F14.1-003 | CCTCAGACAGATCGCCGGCG |
| 2031 | CTD-3222D19.12-001 | TTCCTTTCGCGGGCACAGTG |
| 2032 | CTD-3222D19.12-001 | TTGCTACCTCCGGGCGACCA |
| 2033 | CTD-3222D19.12-001 | TGCTACCTCCGGGCGACCAG |
| 2034 | CTD-3222D19.12-001 | CTCCTCACTGTGCCCGCGAA |
| 2035 | CTD-3222D19.12-001 | GGCCCAGGCACGAAGTCCCC |
| 2036 | CTD-3222D19.12-001 | GGGACTTCGTGCCTGGGCCA |
| 2037 | CTD-3222D19.12-001 | TCCGGGCGACCAGGGGCCTG |
| 2038 | CTD-3252C9.4-001 | CACATCCCTCCTCCAGACAC |
| 2039 | CTD-3252C9.4-001 | AGAATCAGACCTGTGTCTGG |
| 2040 | CTD-3252C9.4-001 | TCAGACCTGTGTCTGGAGGA |
| 2041 | CTD-3252C9.4-001 | TCCAAGCATCAGCCCACCCA |
| 2042 | CTD-3252C9.4-001 | TGATGCTTGGAGCAGAGATG |
| 2043 | CTD-3252C9.4-001 | CTCCAAGCATCAGCCCACCC |
| 2044 | CTD-3252C9.4-001 | AGCATCAGCCCACCCAGGGA |
| 2045 | CYP4A22-AS1-001 | AGTTGCTGGCATGACCTGAG |
| 2046 | CYP4A22-AS1-001 | CGTCCGGTCCCAGTGTGACG |
| 2047 | CYP4A22-AS1-001 | CGTGGTCCCAGGAACCTCTC |
| 2048 | CYP4A22-AS1-001 | TGGGACCGGACGCATTTCCA |
| 2049 | CYP4A22-AS1-001 | CCGGACGCATTTCCATGGCG |
| 2050 | CYP4A22-AS1-001 | CCACGCCATGGAAATGCGTC |
| 2051 | CYP4A22-AS1-001 | CTTCCCCGTCACACTGGGAC |
| 2052 | CYP4F35P-001 | AAACATGTCACGTTCTGGGA |
| 2053 | CYP4F35P-001 | GAAACATGTCACGTTCTGGG |
| 2054 | CYP4F35P-001 | AGTGCTTCTGAAAGGAAAGG |
| 2055 | CYP4F35P-001 | CAAAAGGCTCACTGTCCCAA |
| 2056 | CYP4F35P-001 | CCTTTACGGTCTGTAGGGCA |
| 2057 | CYP4F35P-001 | TGAGAAACATGTCACGTTCT |
| 2058 | CYP4F35P-001 | GGGCTGGAGAATCCGAGGGC |
| 2059 | CYP4F35P-002 | GTTGTTAGATTAAAAAACAG |
| 2060 | CYP4F35P-002 | ATGAGATGTGTGAGAAGCCA |
| 2061 | CYP4F35P-002 | TTGCTTCAAACCTGGACCAG |
| 2062 | CYP4F35P-002 | GCTTCTCACACATCTCATCG |
| 2063 | CYP4F35P-002 | GATGAGATGTGTGAGAAGCC |

| | | |
|---|---|---|
| 2064 | CYP4F35P-002 | GTTGCCAAAATTCACACCGC |
| 2065 | CYP4F35P-002 | TGTGAGAAGCCAGGGTGCCA |
| 2066 | DANCR-001 | GACGGGCGCACAAACCAGAG |
| 2067 | DANCR-001 | CGAGGCCCGCAGCCAATCCC |
| 2068 | DANCR-001 | CCCTTGCCCAGAGTCTTCCC |
| 2069 | DANCR-001 | GGGCGCACAAACCAGAGAGG |
| 2070 | DANCR-001 | GCCAATCCCGGGAAGACTCT |
| 2071 | DANCR-001 | GGGCGGGGAGGGAAGCAGGA |
| 2072 | DANCR-001 | AGGGAAGCAGGAGGGTCGCG |
| 2073 | DBET-001 | GGTGAAAAAATAAACTGCAG |
| 2074 | DBET-001 | ATACCACAGACAGCCAACTG |
| 2075 | DBET-001 | AAAAATAAACTGCAGTGGAC |
| 2076 | DBET-001 | TTTCACCCAGAACAGTAACT |
| 2077 | DBET-001 | AGATTCTTTCTCTCCTGATG |
| 2078 | DBET-001 | GTCCTTCAGCACTCCACATC |
| 2079 | DBET-001 | AATACCACAGACAGCCAACT |
| 2080 | DEDD | CCGAGCCTGAGCCGAGAGTG |
| 2081 | DEDD | TCGACTCCCTCTCCCCACTC |
| 2082 | DEDD | CCGCCCCTCCGCCTCACTCT |
| 2083 | DEDD | CCTCACTCTCGGCTCAGGCT |
| 2084 | DEDD | TGAGCCGAGAGTGAGGCGGA |
| 2085 | DEDD | GAGCCGAGAGTGAGGCGGAG |
| 2086 | DEDD | CTCCGCCTCACTCTCGGCTC |
| 2087 | DKFZP434I0714-001 | CTCCAAGGGAAGAGGAAGTG |
| 2088 | DKFZP434I0714-001 | GCTCCAAGGGAAGAGGAAGT |
| 2089 | DKFZP434I0714-001 | GCCGCGTCAGAACTGAAGAG |
| 2090 | DKFZP434I0714-001 | AGGGAAGAGGAAGTGGGGAT |
| 2091 | DKFZP434I0714-001 | GGTGGCGGCTGCGAGTCCCG |
| 2092 | DKFZP434I0714-001 | CAGTTCTGACGCGGCCGTAG |
| 2093 | DKFZP434I0714-001 | TCCTCTCTTCAGTTCTGACG |
| 2094 | DNMT1 | ATGGCCGGGACGGCCAGTGT |
| 2095 | DNMT1 | TCGTCGGGCAGCGAGATGGC |
| 2096 | DNMT1 | GATGGCCGGGACGGCCAGTG |
| 2097 | DNMT1 | GGGCAGCGAGATGGCCGGGA |
| 2098 | DNMT1 | CCCAGCCCGGGTGCCCACAC |
| 2099 | DNMT1 | GGTGCCCACACTGGCCGTCC |
| 2100 | DNMT1 | GACGGCCAGTGTGGGCACCC |
| 2101 | E2F1 | GCGGGATCGAGCCCTCGCCG |
| 2102 | E2F1 | CGGGACTTTGCAGGCAGCGG |
| 2103 | E2F1 | AAAAGTGGCCGGGACTTTGC |
| 2104 | E2F1 | CGCCGCTGCCTGCAAAGTCC |
| 2105 | E2F1 | GGCCGGGACTTTGCAGGCAG |
| 2106 | E2F1 | ATGGCGGCAGGCCTCGGCGA |

| | | |
|---|---|---|
| 2107 | E2F1 | CGCCTGTCACCCGGGCCGCG |
| 2108 | EBLN3-001 | GACACAGAAGGCGACGCCTA |
| 2109 | EBLN3-001 | GCATACCAGGCGCGACAGGT |
| 2110 | EBLN3-001 | CGCATACCAGGCGCGACAGG |
| 2111 | EBLN3-001 | ATGCGCATACCAGGCGCGAC |
| 2112 | EBLN3-001 | TTCCGGGACAACACAGCCCT |
| 2113 | EBLN3-001 | GGAGGAGAGAAGGCACTTCC |
| 2114 | EBLN3-001 | GGACACAGAAGGCGACGCCT |
| 2115 | EBLN3-002 | CCACAGCTCGGCTGGCGAGA |
| 2116 | EBLN3-002 | CGCTAGTTCCACAGCTCGGC |
| 2117 | EBLN3-002 | CCCCGAGTCACCCGCGGCCT |
| 2118 | EBLN3-002 | GGCCGGGCTGCGGGTCTAGG |
| 2119 | EBLN3-002 | AGGCAGGGGTCCCAGGCCGC |
| 2120 | EBLN3-002 | ACCCGCAGCCCGGCCGCGTG |
| 2121 | EBLN3-002 | ACCCCGAGTCACCCGCGGCC |
| 2122 | EBLN3-003 | CTAAGGTCGTGATTCTAGGG |
| 2123 | EBLN3-003 | CAGACACGCCACCCCCACCG |
| 2124 | EBLN3-003 | AGAATCACGACCTTAGCCCT |
| 2125 | EBLN3-003 | CTTGGAAAGCCAGGGAGATG |
| 2126 | EBLN3-003 | CCCTGGCTTTCCAAGGGCTA |
| 2127 | EBLN3-003 | GTCGTGATTCTAGGGCGGCT |
| 2128 | EBLN3-003 | GGGCTAAGGTCGTGATTCTA |
| 2129 | EBLN3-004 | CACGCGCTAGTTCCACAGCT |
| 2130 | EBLN3-004 | CTGCGCCCCACCACAAGTGA |
| 2131 | EBLN3-004 | TGCGCCCCACCACAAGTGAG |
| 2132 | EBLN3-004 | GCCCCACCACAAGTGAGGGG |
| 2133 | EBLN3-004 | GGCAGTGAGTATGTATGTGA |
| 2134 | EBLN3-004 | TCCGCCCCTCACTTGTGGTG |
| 2135 | EBLN3-004 | GCTGCGCCCCACCACAAGTG |
| 2136 | EBLN3-005 | CAAACAGAAAGCACCAAACA |
| 2137 | EBLN3-005 | CAGGGACCAAAACGCGTAGA |
| 2138 | EBLN3-005 | GCAAACAGAAAGCACCAAAC |
| 2139 | EBLN3-005 | ACAGGGACCAAAACGCGTAG |
| 2140 | EBLN3-005 | GGGAGGGGCGAGGACAAATG |
| 2141 | EBLN3-005 | CGAGGACAAATGAGGTAAGG |
| 2142 | EBLN3-005 | AAACGCGTAGAGGGACGGGG |
| 2143 | EBLN3-006 | GGAGGGTAGGCAGGGGTCCC |
| 2144 | EBLN3-006 | CGGCGCGCAGCTGAGGGACC |
| 2145 | EBLN3-006 | GACCCGCAGCCCGGCCGCGT |
| 2146 | EBLN3-006 | CGCAGCCCGGCCGCGTGGGG |
| 2147 | EBLN3-006 | GCGGCCGGGCTGCGGGTCTA |
| 2148 | EBLN3-006 | TAGGCAGGGGTCCCAGGCCG |
| 2149 | EBLN3-006 | CGGCCGGGCTGCGGGTCTAG |

| | | |
|---|---|---|
| 2150 | EIF3J-AS1-001 | CAGACCAGTACTGGAGTTCG |
| 2151 | EIF3J-AS1-001 | AGCACCACGAACTCCAGTAC |
| 2152 | EIF3J-AS1-001 | CAGTACTGGTCTGCGTTGTA |
| 2153 | EIF3J-AS1-001 | AGTACTGGAGTTCGTGGTGC |
| 2154 | EIF3J-AS1-001 | ACCCGGGGCCCAGAGTCGGG |
| 2155 | EIF3J-AS1-001 | CCTACAACGCAGACCAGTAC |
| 2156 | EIF3J-AS1-001 | CCAGTACTGGTCTGCGTTGT |
| 2157 | EIF3J-AS1-005 | GTGATAAAAAGGTAGATACG |
| 2158 | EIF3J-AS1-005 | TGATAAAAAGGTAGATACGT |
| 2159 | EIF3J-AS1-005 | TAGCTCTTTAAAATCCTGCG |
| 2160 | EIF3J-AS1-005 | AAAAAGGTAGATACGTGGGC |
| 2161 | EIF3J-AS1-005 | GCTAGCTCTTTAAAATCCTG |
| 2162 | EIF3J-AS1-005 | AGCTCTTTAAAATCCTGCGG |
| 2163 | EIF3J-AS1-005 | GTAGATACGTGGGCAGGACG |
| 2164 | ENO1 | GCGGCGCGGACAGTATCTGT |
| 2165 | ENO1 | CGCCGGCTTTACGTTCACCT |
| 2166 | ENO1 | CACCGAGGTGAACGTAAAGC |
| 2167 | ENO1 | CCGGAGCACGGAGATCTCGC |
| 2168 | ENO1 | CCAATCATTACCTAGCCACT |
| 2169 | ENO1 | GGAGGGTGCTGCAGACACCG |
| 2170 | ENO1 | ATCTGTGGGTACCCGGAGCA |
| 2171 | EPB41L4A-AS1-001 | CTGTCGGGTCCCAAAATACG |
| 2172 | EPB41L4A-AS1-001 | CAAGAGCATATGATCAGTGC |
| 2173 | EPB41L4A-AS1-001 | CTGGACTCATGAATTCTGTC |
| 2174 | EPB41L4A-AS1-001 | GACATCCCCCCACCCGACCA |
| 2175 | EPB41L4A-AS1-001 | CATATGCTCTTGACCCTGGT |
| 2176 | EPB41L4A-AS1-001 | TGATCATATGCTCTTGACCC |
| 2177 | EPB41L4A-AS1-001 | GCTGGACTCATGAATTCTGT |
| 2178 | EPB41L4A-AS1-002 | TTCTCCCTCCGGCCACACGG |
| 2179 | EPB41L4A-AS1-002 | ACCGAGGGGCGTGTCACCAA |
| 2180 | EPB41L4A-AS1-002 | CCGAGGGGCGTGTCACCAAA |
| 2181 | EPB41L4A-AS1-002 | AGTGCCAGTGAGGGACCGAG |
| 2182 | EPB41L4A-AS1-002 | CCCTTTGGTGACACGCCCCT |
| 2183 | EPB41L4A-AS1-002 | AAGTGCCAGTGAGGGACCGA |
| 2184 | EPB41L4A-AS1-002 | CTCACTGGCACTTCTCCCTC |
| 2185 | EPB41L4A-AS1-003 | CAGTCGACTCACCAGACTCA |
| 2186 | EPB41L4A-AS1-003 | GGGTCCTCTGGGATGGCGCC |
| 2187 | EPB41L4A-AS1-003 | TCTGGGATGGCGCCTGGCTG |
| 2188 | EPB41L4A-AS1-003 | CTCACGCAGGGGCGCCGGGC |
| 2189 | EPB41L4A-AS1-003 | CACGCAGGGGCGCCGGGCGG |
| 2190 | EPB41L4A-AS1-003 | GGCGCCTGGCTGTGGCCGCG |
| 2191 | EPB41L4A-AS1-003 | GGTGGCCGCGTTCCCCCGCC |
| 2192 | EPB41L4A-AS1-004 | ATAAAGTCAGTTATACGGTA |

| | | |
|---|---|---|
| 2193 | EPB41L4A-AS1-004 | TGAGTCTGGTGAGTCGACTG |
| 2194 | EPB41L4A-AS1-004 | GCAGTCGACTCACCAGACTC |
| 2195 | EPB41L4A-AS1-004 | TCACTGACGAAAATAAAACC |
| 2196 | EPB41L4A-AS1-004 | TTACATCATCGACTATGCCA |
| 2197 | EPB41L4A-AS1-004 | TCGACTCACCAGACTCAGGG |
| 2198 | EPB41L4A-AS1-004 | TTTACATCATCGACTATGCC |
| 2199 | EPB41L4A-AS2-201 | GCCGAGCGCTTCATACTCGG |
| 2200 | EPB41L4A-AS2-201 | TCCGCCGAGTATGAAGCGCT |
| 2201 | EPB41L4A-AS2-201 | AAAGCCGAGCGCTTCATACT |
| 2202 | EPB41L4A-AS2-201 | CGCCCATGGTCGGGGCGTCG |
| 2203 | EPB41L4A-AS2-201 | ATACTCGGCGGATCTGACCC |
| 2204 | EPB41L4A-AS2-201 | CGCCCCGACCATGGGCGCAG |
| 2205 | EPB41L4A-AS2-201 | ACCCACGACGCCCCGACCAT |
| 2206 | ERVH-1-001 | TGATGGCTTAGTGACCCCTG |
| 2207 | ERVH-1-001 | ATCAGCTGATCAAGAAAGGT |
| 2208 | ERVH-1-001 | ACATAGGTATACGTGTGCAG |
| 2209 | ERVH-1-001 | TAGGTATACGTGTGCAGTGG |
| 2210 | ERVH-1-001 | ACACCAAGGAAAGCTCGGTG |
| 2211 | ERVH-1-001 | AGCCATCAGCTGATCAAGAA |
| 2212 | ERVH-1-001 | TTTCCTTGGTGTCAGCCACA |
| 2213 | EXO1 | CCGACGCCCCTAACTCACTG |
| 2214 | EXO1 | CGGCCGGCAACGCAGAACAC |
| 2215 | EXO1 | TTGCCTACACAGCGCGCGAA |
| 2216 | EXO1 | CAGCCTTTCGCGCGCTGTGT |
| 2217 | EXO1 | CCACAGTGAGTTAGGGGCGT |
| 2218 | EXO1 | GTGAGTTAGGGGCGTCGGAG |
| 2219 | EXO1 | TGTTCTGCGTTGCCGGCCGT |
| 2220 | EXOSC2 | GAAACATCTAGTGGTGCCGG |
| 2221 | EXOSC2 | CGACACTAAGAAACATCTAG |
| 2222 | EXOSC2 | AGAAACATCTAGTGGTGCCG |
| 2223 | EXOSC2 | CGGCCCAGTCTCTCGCTAAG |
| 2224 | EXOSC2 | AAGCCTCTTAGCGAGAGACT |
| 2225 | EXOSC2 | GCAACTCATTGGCGCCAAGA |
| 2226 | EXOSC2 | GATGGAGATGAGGCTTCCAG |
| 2227 | FAM157C-004 | TTAAACTAGAATGAGCAAGA |
| 2228 | FAM157C-004 | TGTAATCCCAGAACTCTGGG |
| 2229 | FAM157C-004 | GAACTCTGGGAGGCCAAGGT |
| 2230 | FAM157C-004 | AAAAAACACAAAAATCAGCT |
| 2231 | FAM157C-004 | AGAACTCTGGGAGGCCAAGG |
| 2232 | FAM157C-004 | CAGGAGTTTGAGAACAGTCT |
| 2233 | FAM157C-004 | CAAAAATCAGCTGGGTGTGG |
| 2234 | FAM157C-013 | GCATCCCATATCCCATTGCA |
| 2235 | FAM157C-013 | CATATGTCCTGCAAGCTTAG |

| | | |
|---|---|---|
| 2236 | FAM157C-013 | AGGAGATAGACGCACACAGA |
| 2237 | FAM157C-013 | AGTGCTGAGCTCCTTGCAAT |
| 2238 | FAM157C-013 | CAGTGCTGAGCTCCTTGCAA |
| 2239 | FAM157C-013 | GGAGATAGACGCACACAGAT |
| 2240 | FAM157C-013 | TTGATGCTAGGAAGGGTACC |
| 2241 | FAM215B-001 | TGGGCAACAGGGCAAAACGC |
| 2242 | FAM215B-001 | TCCCAAGTCGCTAAGACCAT |
| 2243 | FAM215B-001 | CACTGCAGTGATTCCATATG |
| 2244 | FAM215B-001 | CCCATATGGAATCACTGCAG |
| 2245 | FAM215B-001 | TCTGCTCTTATGAGCTCACA |
| 2246 | FAM215B-001 | CTCTGCTCTTATGAGCTCAC |
| 2247 | FAM215B-001 | ACATTAAAGGAAAAACTCCT |
| 2248 | FAM27C-001 | ACACCCAAGCACTGCCACAG |
| 2249 | FAM27C-001 | AGCGGTTTCTCCCGTCACAG |
| 2250 | FAM27C-001 | GGCAAAGCAGGAGCCCTCTG |
| 2251 | FAM27C-001 | GTCTTAGAAACCAGTCCCCG |
| 2252 | FAM27C-001 | CGATGAGGAAACCCCAGGCT |
| 2253 | FAM27C-001 | CACCCAAGCACTGCCACAGA |
| 2254 | FAM27C-001 | AACCATGGGAACCACTGTGA |
| 2255 | FAM95B1-001 | CCCTTCCTAATCATACCAGA |
| 2256 | FAM95B1-001 | ATGAGATGTTTCATATGGCA |
| 2257 | FAM95B1-001 | CCTTCTGGTATGATTAGGAA |
| 2258 | FAM95B1-001 | ATGTTTCATATGGCAGGGGT |
| 2259 | FAM95B1-001 | CATAGTGGAAGTTTCTCAAG |
| 2260 | FAM95B1-001 | AGGAAGGGCCTGGTCAGTGT |
| 2261 | FAM95B1-001 | TGAGATGTTTCATATGGCAG |
| 2262 | FLJ20021-001 | GCAAAAGTGACACCTGAGCA |
| 2263 | FLJ20021-001 | GAGCAGGGACTGGTCCGTAG |
| 2264 | FLJ20021-001 | CAGGGACTGGTCCGTAGAGG |
| 2265 | FLJ20021-001 | TCTTCCAGCTGCCGGACCAG |
| 2266 | FLJ20021-001 | AGCAAAAGTGACACCTGAGC |
| 2267 | FLJ20021-001 | CCACTCTCCTGCCCTAATCT |
| 2268 | FLJ20021-001 | TGGCAGGCCGAGATTAGGGC |
| 2269 | FLJ20021-002 | GCGGCTTAACCCAGCCGACA |
| 2270 | FLJ20021-002 | ACAAGGCACGCTTGCCAAAG |
| 2271 | FLJ20021-002 | CAAGTGCAGAGGGTGTCTCC |
| 2272 | FLJ20021-002 | GGAGAGTGGGCAAGGCGTCG |
| 2273 | FLJ20021-002 | GTGTGAGTAGCCAAGTGCAG |
| 2274 | FLJ20021-002 | TGTGAGTAGCCAAGTGCAGA |
| 2275 | FLJ20021-002 | GCACGCTTGCCAAAGAGGCG |
| 2276 | FLJ22447-002 | CAGACTAGTGACTTCCAGTG |
| 2277 | FLJ22447-002 | GAAGGTGCAGAAACCATGGG |
| 2278 | FLJ22447-002 | AGAAACCATGGGAGGTAAGG |

| | | |
|---|---|---|
| 2279 | FLJ22447-002 | TTCTGCACCTTCATTTCCAA |
| 2280 | FLJ22447-002 | GAAACCATGGGAGGTAAGGA |
| 2281 | FLJ22447-002 | AAATGAAGGTGCAGAAACCA |
| 2282 | FLJ22447-002 | AACTGAGTCCAGCAGCACTG |
| 2283 | FLJ31662-001 | CAGAAAGAAAACCAGTCCAG |
| 2284 | FLJ31662-001 | GTAGCTCCATGTAAGTGACT |
| 2285 | FLJ31662-001 | GCTAATCCAAAAATACAAGC |
| 2286 | FLJ31662-001 | GGAAATATCTTTGAATTGCT |
| 2287 | FLJ31662-001 | ATTGTTCCAAGTCACTTACA |
| 2288 | FLJ31662-001 | CCTCTCAGTAAAAGCAGAGA |
| 2289 | FLJ31662-001 | ATGACTCTTCTGCTTCTGGG |
| 2290 | GAPLINC-004 | GGTGGTAGGTCAGATAGCAT |
| 2291 | GAPLINC-004 | TTACTTCCAGCATTAGACGC |
| 2292 | GAPLINC-004 | GGTTTGGGTGATAGTAACTG |
| 2293 | GAPLINC-004 | GTTTACAGTGATGTGTATGC |
| 2294 | GAPLINC-004 | TTAATGCCTGCGTCTAATGC |
| 2295 | GAPLINC-004 | CTATCAAATCTCTCTGCAAA |
| 2296 | GAPLINC-004 | TGACACCTGGATAACAGGTT |
| 2297 | GAS5-001 | CAGGGCCTACAAGTTGAGGA |
| 2298 | GAS5-001 | GAAAAGACAGTATGGTGCCT |
| 2299 | GAS5-001 | TCAACTTGTAGGCCCTGCAA |
| 2300 | GAS5-001 | GCCCTGCAAAGGCGCGCGAC |
| 2301 | GAS5-001 | CTACAAGTTGAGGATGGTGG |
| 2302 | GAS5-001 | TAAGGTCACCCTGAATTGAC |
| 2303 | GAS5-001 | TCAGATGTCCCTACCAACAC |
| 2304 | GAS5-003 | TATTTAACGTGACTGCTGTG |
| 2305 | GAS5-003 | AGAATTGGGGAAAGTATGGT |
| 2306 | GAS5-003 | CTGGATAGCACCTTATGGAC |
| 2307 | GAS5-003 | GTACCTGATCTCTTTAGGTA |
| 2308 | GAS5-003 | TTATGGACAGGTAAGAATTG |
| 2309 | GAS5-003 | ATCTCTTTAGGTATGGTGCT |
| 2310 | GAS5-003 | GATCTCTTTAGGTATGGTGC |
| 2311 | GAS5-008 | AAGGTCACCCTGAATTGACT |
| 2312 | GAS5-008 | AGGGTGACCTTAGCAGACAA |
| 2313 | GAS5-008 | AAAGATAAAACATACCTCAC |
| 2314 | GAS5-008 | TAGAAGTCCCAGTCAATTCA |
| 2315 | GAS5-008 | GGTAGGAGTCGACTCCTGTG |
| 2316 | GAS5-008 | ACTATGTTAGAGTAGAACTG |
| 2317 | GAS5-008 | CTCTAACATAGTCGCAGCTT |
| 2318 | GAS5-014 | AATGCAGGCACTTAAGCACG |
| 2319 | GAS5-014 | TGGAAATATTGGTGACACTG |
| 2320 | GAS5-014 | CCGACAGCCTTTCATGAGAG |
| 2321 | GAS5-014 | CAGCCTTTCATGAGAGCGGA |

| | | |
|---|---|---|
| 2322 | GAS5-014 | TGAGAGCGGACGGCTGATGG |
| 2323 | GAS5-014 | CAGCCGTCCGCTCTCATGAA |
| 2324 | GAS5-014 | CCGCTCTCATGAAAGGCTGT |
| 2325 | GAS5-028 | TTAATGCATTCAGCACTAGG |
| 2326 | GAS5-028 | TGAATGCATTAAATAAATGG |
| 2327 | GAS5-028 | GAAACTGTATCCAAGTAAGC |
| 2328 | GAS5-028 | TAAAAAGCCATTTGTGCCGT |
| 2329 | GAS5-028 | TGCCCAGTTACCTGCTTACT |
| 2330 | GAS5-028 | ACAGGCATTAGACAGAAAGC |
| 2331 | GAS5-028 | CAAACTTCCTACGGCACAAA |
| 2332 | GMDS-AS1-002 | CTCCTTCCTCTTCCCTGAGG |
| 2333 | GMDS-AS1-002 | CACTGCGTCCAGGCCCCCCA |
| 2334 | GMDS-AS1-002 | CACCGCCTCAGGGAAGAGGA |
| 2335 | GMDS-AS1-002 | ACGGCCAGCCAGGCGAGAGG |
| 2336 | GMDS-AS1-002 | CCGCCACCGCCTCAGGGAAG |
| 2337 | GMDS-AS1-002 | GGCCTCCTTCCTCTTCCCTG |
| 2338 | GMDS-AS1-002 | AGGGAAGAGGAAGGAGGCCG |
| 2339 | GMDS-AS1-007 | TTTATCAAAGTCCATGACCC |
| 2340 | GMDS-AS1-007 | AGGGTCCTGGGAGATTCGGG |
| 2341 | GMDS-AS1-007 | AGTGGCCTCCCGAATCTCCC |
| 2342 | GMDS-AS1-007 | TATTAGGGTCCTGGGAGATT |
| 2343 | GMDS-AS1-007 | CCTCTTCCCTGAGGCGGTGG |
| 2344 | GMDS-AS1-007 | CTCGCCTGGCTGGCCGTGGG |
| 2345 | GMDS-AS1-007 | CTTCCTCTTCCCTGAGGCGG |
| 2346 | GS1-124K5.11-001 | CAGCTACTAGAGAGGCTGAG |
| 2347 | GS1-124K5.11-001 | CTGTAGTCCCAGCTCTCAGG |
| 2348 | GS1-124K5.11-001 | GCTACTAGAGAGGCTGAGTG |
| 2349 | GS1-124K5.11-001 | CAAAAAACAGGAGCCAGGCT |
| 2350 | GS1-124K5.11-001 | AAGGCTGCAGTAAGTCATGA |
| 2351 | GS1-124K5.11-001 | TGTCTCAAAAAACAGGAGCC |
| 2352 | GS1-124K5.11-001 | ACTAGAGAGGCTGAGTGGGG |
| 2353 | GS1-124K5.3-001 | GGTTACCAGGCTCCACTCCG |
| 2354 | GS1-124K5.3-001 | TGTTGCTGGCTGGAGAAGGG |
| 2355 | GS1-124K5.3-001 | GCAACACACAGAACATGGCG |
| 2356 | GS1-124K5.3-001 | CGGAAGGATCCGTTCAGGCC |
| 2357 | GS1-124K5.3-001 | AGAACATGGCGGGGAAGTCG |
| 2358 | GS1-124K5.3-001 | GGAAAAACCAGCGCAGCTCC |
| 2359 | GS1-124K5.3-001 | AGCCAGCAACACACAGAACA |
| 2360 | GS1-124K5.3-002 | CTGAACGGATCCTTCCGCCT |
| 2361 | GS1-124K5.3-002 | GGCCTTCGGCGACGTCACAG |
| 2362 | GS1-124K5.3-002 | GGCCGGCGCTATTACGACGC |
| 2363 | GS1-124K5.3-002 | CGCGCCTGTGACGTCAGCTG |
| 2364 | GS1-124K5.3-002 | TTCCGGCGTCGTAATAGCGC |

| | | |
|---|---|---|
| 2365 | GS1-124K5.3-002 | GGCGCTATTACGACGCCGGA |
| 2366 | GS1-124K5.3-002 | ACTATGTCCAGGCTCGGCCG |
| 2367 | GS1-124K5.4-001 | GACGCATGGCCACAGACTGC |
| 2368 | GS1-124K5.4-001 | CACCGCCTGCTGACACTGAC |
| 2369 | GS1-124K5.4-001 | TGCCGGTCAGTGTCAGCAGG |
| 2370 | GS1-124K5.4-001 | GGTCAGTGTCAGCAGGCGGT |
| 2371 | GS1-124K5.4-001 | TTGTAGCTTATTGGTGTCGG |
| 2372 | GS1-124K5.4-001 | AGGGTCCGCAGGCCCAGACC |
| 2373 | GS1-124K5.4-001 | GACTGCCGGTCAGTGTCAGC |
| 2374 | GS1-124K5.4-002 | GACACCAATAAGCTACAAGG |
| 2375 | GS1-124K5.4-002 | CTGCGCTACTTCCCAGCCTG |
| 2376 | GS1-124K5.4-002 | GGCAGTAGTAGTACAGGCAG |
| 2377 | GS1-124K5.4-002 | GCCGACACCAATAAGCTACA |
| 2378 | GS1-124K5.4-002 | AGCGCAGGCAGTAGTAGTAC |
| 2379 | GS1-124K5.4-002 | TGCGGCGTAGGGCGCAGCAA |
| 2380 | GS1-124K5.4-002 | ACTTCCCAGCCTGCGGCGTA |
| 2381 | H2AFZ | AGACAGCGCAGTTTGAATCG |
| 2382 | H2AFZ | TGGCGACGGTTGGGAACGCG |
| 2383 | H2AFZ | AGGTGGTGGGATCTCACCGT |
| 2384 | H2AFZ | TAGGTGGTGGGATCTCACCG |
| 2385 | H2AFZ | ACCCGTTACACAGAGGGATG |
| 2386 | H2AFZ | AAACCCGTTACACAGAGGGA |
| 2387 | H2AFZ | GAAAAGGCTAATCGGACCCA |
| 2388 | HCG11-001 | CAGTAATATACTGCTCACAA |
| 2389 | HCG11-001 | ACTGCTGTGTACATACATAG |
| 2390 | HCG11-001 | CTTGCTCAATGGTTTAGAGG |
| 2391 | HCG11-001 | TACTGTCTATACTTGCTCAA |
| 2392 | HCG11-001 | CCTCTACATTTAATATACAT |
| 2393 | HCG11-001 | CTCTACATTTAATATACATC |
| 2394 | HCG11-001 | CCGATGTATATTAAATGTAG |
| 2395 | HELLS | GGTTGTGAGGAGTTAGCTCG |
| 2396 | HELLS | TAACTCCTCACAACCGCTGC |
| 2397 | HELLS | TGGACACTCACCCGGGAACC |
| 2398 | HELLS | GCGGCATTGCAGGCTCTGAG |
| 2399 | HELLS | GCATTGCAGGCTCTGAGAGG |
| 2400 | HELLS | CTGGGCCGGCAGCGGTTGTG |
| 2401 | HELLS | GGGGATTTGGCTAGAAGGCT |
| 2402 | HNRNPR | GGCCCGTGAGAATCAGCGCG |
| 2403 | HNRNPR | GGCCGGCGCGAGACTCACCA |
| 2404 | HNRNPR | TGCCCTGGTGAGTCTCGCGC |
| 2405 | HNRNPR | CGAGACTCACCAGGGCAGAG |
| 2406 | HNRNPR | CGCGCTGATTCTCACGGGCC |
| 2407 | HNRNPR | CGCCTCGCGCTGATTCTCAC |

| | | |
|---|---|---|
| 2408 | HNRNPR | GCGCCTCGCGCTGATTCTCA |
| 2409 | HRAT17-002 | ATTGCTAAATGCCCCTTGAG |
| 2410 | HRAT17-002 | ACCCAGTCTAGACAAAGGTT |
| 2411 | HRAT17-002 | TCCCAACCTTTGTCTAGACT |
| 2412 | HRAT17-002 | TAGAAAATGGAGTAGATACC |
| 2413 | HRAT17-002 | TAGTTTCCTAATCAGACTCC |
| 2414 | HRAT17-002 | TAGATACCAGGAGTCTGATT |
| 2415 | HRAT17-002 | GAATCACTCTGGATGCTGTG |
| 2416 | ILF2 | CCTTAAAACACGAACAATGG |
| 2417 | ILF2 | TAAGGCGCCATGAGGTAAGT |
| 2418 | ILF2 | AGTGGGTCGAAGCGGTCGAA |
| 2419 | ILF2 | GCTTCGACCCACTTACCTCA |
| 2420 | ILF2 | GCGCCTTAAAACACGAACAA |
| 2421 | ILF2 | TTCAGTTGTCTGCTACTCAG |
| 2422 | ILF2 | TTAAGGCGCCATGAGGTAAG |
| 2423 | ILF3-AS1-001 | ACACATGCGTCAAAGCCTCT |
| 2424 | ILF3-AS1-001 | CCCGCAGACCTGCCTTGGTG |
| 2425 | ILF3-AS1-001 | CCAAGAACCCGGGCGCGCAT |
| 2426 | ILF3-AS1-001 | CACTGGAGCCCGAAAATCCC |
| 2427 | ILF3-AS1-001 | ACGAAAGAACCCAAGAACCC |
| 2428 | ILF3-AS1-001 | CCACACCAAGGCAGGTCTGC |
| 2429 | ILF3-AS1-001 | GGCCCAGGGACGCCACACCA |
| 2430 | IQCH-AS1-002 | CCTGCCCGAGTTACTAATGA |
| 2431 | IQCH-AS1-002 | CGTCATTAGTAACTCGGGCA |
| 2432 | IQCH-AS1-002 | GAGTTACTAATGACGGCGTG |
| 2433 | IQCH-AS1-002 | CACGCCGTCATTAGTAACTC |
| 2434 | IQCH-AS1-002 | GTCATTAGTAACTCGGGCAG |
| 2435 | IQCH-AS1-002 | CAAAACCAGCTCCTTGACGC |
| 2436 | IQCH-AS1-002 | TCACGCCGTCATTAGTAACT |
| 2437 | IQCH-AS1-005 | GGTTTGAGTTCTGCACCTTG |
| 2438 | IQCH-AS1-005 | CTGGAGCCCAGGGCGTGGGA |
| 2439 | IQCH-AS1-005 | TTGAGTTCTGCACCTTGCGG |
| 2440 | IQCH-AS1-005 | AATAACTAACCACCTCCGCA |
| 2441 | IQCH-AS1-005 | GGGCCTGAGGAAAATTCAGG |
| 2442 | IQCH-AS1-005 | AAGTCTCCCTCCCACGCCCT |
| 2443 | IQCH-AS1-005 | TTTGGGCCTGAGGAAAATTC |
| 2444 | IRF4 | CCGAGTCCAGGGCGAGGTAA |
| 2445 | IRF4 | TCGGAGCTGAGGGCAGCGGT |
| 2446 | IRF4 | TGCCCTCAGCTCCGAGTCCA |
| 2447 | IRF4 | GCGAGGTAAGGGCTGGAGTC |
| 2448 | IRF4 | TCAGCTCCGAGTCCAGGGCG |
| 2449 | IRF4 | CCCTTACCTCGCCCTGGACT |
| 2450 | IRF4 | TCCGAGTCCAGGGCGAGGTA |

| | | |
|---|---|---|
| 2451 | JPX-003 | CGAAGCTTGGAAGTTGCAGA |
| 2452 | JPX-003 | AAAGTGTGTAGCCTCGCTAA |
| 2453 | JPX-003 | TAGAGAAGTGGCCGTTAGCG |
| 2454 | JPX-003 | TAACTTCTACTGACGAAGCT |
| 2455 | JPX-003 | GCCGGTTACTCCCCCGACTC |
| 2456 | JPX-003 | CTCGCCAGTTAATAGTATTG |
| 2457 | JPX-003 | GAGAGAGAAAAATGGAATAC |
| 2458 | JPX-007 | CAATCACTGCGTCCTTACGG |
| 2459 | JPX-007 | TTTCATCCGCCTATGCCCTA |
| 2460 | JPX-007 | TAAGTCTTCCACTAGCCCTA |
| 2461 | JPX-007 | TAGTGGAAGACTTAAGATGG |
| 2462 | JPX-007 | GCAATCACTGCGTCCTTACG |
| 2463 | JPX-007 | TTCCACTAGCCCTAGGGCAT |
| 2464 | JPX-007 | CACTAGCCCTAGGGCATAGG |
| 2465 | JPX-012 | AAAATAAAGTTAGGCCCAAG |
| 2466 | JPX-012 | ACGACCCATATATCCCTCCT |
| 2467 | JPX-012 | TGCAGAAGGTAGCTCCTAGG |
| 2468 | JPX-012 | GCAGAAGGTAGCTCCTAGGA |
| 2469 | JPX-012 | CTAGGAACAGCCTAACCTCT |
| 2470 | JPX-012 | CTAATAAGTCTGGACATGAA |
| 2471 | JPX-012 | CTGTTCCTAGGAAAGCTCTG |
| 2472 | KANTR-001 | CCCGCTGACCTAGGGTTGAG |
| 2473 | KANTR-001 | TGGGGCGGCCTCTCAACCCT |
| 2474 | KANTR-001 | CTCCGCAACCCGCTGACCTA |
| 2475 | KANTR-001 | GCCTCTCAACCCTAGGTCAG |
| 2476 | KANTR-001 | GCCCCAGTAAAAGCTCGCGC |
| 2477 | KANTR-001 | CCTCTCAACCCTAGGTCAGC |
| 2478 | KANTR-001 | GGCCGGCCGAAGCGCGCGAG |
| 2479 | KANTR-005 | GCTGGACGGAACCCTTCCAC |
| 2480 | KANTR-005 | AGGCCTACAGCAAGATCTGA |
| 2481 | KANTR-005 | TATCCTTCAGATCTTGCTGT |
| 2482 | KANTR-005 | GATCTGAAGGATACGCTGGA |
| 2483 | KANTR-005 | GCAAGATCTGAAGGATACGC |
| 2484 | KANTR-005 | CGCTGGACGGAACCCTTCCA |
| 2485 | KANTR-005 | CAGATCTTGCTGTAGGCCTC |
| 2486 | KB-1460A1.1-001 | CATACCTGCCTGAGGTACGG |
| 2487 | KB-1460A1.1-001 | CCATACCTGCCTGAGGTACG |
| 2488 | KB-1460A1.1-001 | CCCAGTCCCATACCTGCCTG |
| 2489 | KB-1460A1.1-001 | TACGGGGGTTCAGGCTGAGT |
| 2490 | KB-1460A1.1-001 | AGCCTGAACCCCCGTACCTC |
| 2491 | KB-1460A1.1-001 | CCCATACCTGCCTGAGGTAC |
| 2492 | KB-1460A1.1-001 | AAGACAGACCTGTAGCAGCA |
| 2493 | KB-1980E6.3-001 | GTGGGGGACACACTGAAGCG |

| | | |
|---|---|---|
| 2494 | KB-1980E6.3-001 | GTCTTGTCTGTAATGCCAGG |
| 2495 | KB-1980E6.3-001 | GCGAGGAAATCCTAGAGGGA |
| 2496 | KB-1980E6.3-001 | AGGTGTAGCACGAGATGGGA |
| 2497 | KB-1980E6.3-001 | GTAGCACGAGATGGGAAGGT |
| 2498 | KB-1980E6.3-001 | ATCCTCCCACAGCCACCAGG |
| 2499 | KB-1980E6.3-001 | ATCGTTCTCACTCCTCCTGG |
| 2500 | KIAA0125-003 | CCTTTGACCAGCTCTGCTGT |
| 2501 | KIAA0125-003 | GGCTCCCCTCCTGGGTGAAG |
| 2502 | KIAA0125-003 | CAGCTCTGCTGTGGGGATCT |
| 2503 | KIAA0125-003 | CTTTGACCAGCTCTGCTGTG |
| 2504 | KIAA0125-003 | CCAGATCCCCACAGCAGAGC |
| 2505 | KIAA0125-003 | ACTCACGCGGCTCCCCTCCT |
| 2506 | KIAA0125-003 | ACCTTTGACCAGCTCTGCTG |
| 2507 | KIAA0125-201 | AGACACACGGACATCGGGAA |
| 2508 | KIAA0125-201 | ACACGGACATCGGGAACGGA |
| 2509 | KIAA0125-201 | TCCTGGCCACAGGAGACACA |
| 2510 | KIAA0125-201 | CGATGTCCGTGTGTCTCCTG |
| 2511 | KIAA0125-201 | CGGACATCGGGAACGGACGG |
| 2512 | KIAA0125-201 | ACAGGAGACACACGGACATC |
| 2513 | KIAA0125-201 | CATCGGGAACGGACGGAGGA |
| 2514 | KIF11 | CCAGAGTACCGGGTAGAGAG |
| 2515 | KIF11 | AGAGTACCGGGTAGAGAGCG |
| 2516 | KIF11 | GGCGTTTGTCCCGCGCCAGT |
| 2517 | KIF11 | CTGGCGCGGGACAAACGCCA |
| 2518 | KIF11 | GAGTCGTTGCTTAGTTTCTG |
| 2519 | KIF11 | AACGCCACGGCCAGAGTACC |
| 2520 | KIF11 | TCTACCCGGTACTCTGGCCG |
| 2521 | KIF14 | GCGTGGGGACTGTGTCGAGG |
| 2522 | KIF14 | GGCGTGGGGACTGTGTCGAG |
| 2523 | KIF14 | ATTCAAATTGCGGCCTTCTG |
| 2524 | KIF14 | CGGGGCGCCCATTCAAATTG |
| 2525 | KIF14 | TGGAGGTGGTGAGTGGCGTG |
| 2526 | KIF14 | CCACTCACCACCTCCAGCGC |
| 2527 | KIF14 | TGGCGTGGGGACTGTGTCGA |
| 2528 | KIF15 | GGGCGGAATTCAGTCGCGCG |
| 2529 | KIF15 | ATTGTCTTTGAGCGACAGAG |
| 2530 | KIF15 | TTGTCTTTGAGCGACAGAGC |
| 2531 | KIF15 | TGCAGGAGCTGAGTGACCGG |
| 2532 | KIF15 | CATGCCTAGCAAGAACCTTG |
| 2533 | KIF15 | GATCCCGAAAACAATGCAGC |
| 2534 | KIF15 | TCAGTCGCGCGCGGTGCAGT |
| 2535 | KIF18A | ACATTAAAGTGAAGTGGTTG |
| 2536 | KIF18A | AGTGAGTGAGAGGCACATGA |

| | | |
|---|---|---|
| 2537 | KIF18A | GTGGTTGAGTCTGAAGCGCT |
| 2538 | KIF18A | AGGCGGACATTAAAGTGAAG |
| 2539 | KIF18A | GTGAAGTGGTTGCGGTAACC |
| 2540 | KIF18A | GTTGAGTCTGAAGCGCTGGG |
| 2541 | KIF18A | TGGTTGCGGTAACCTGGCCT |
| 2542 | KIF20B | GTGACCCGCGGGAATGTGTG |
| 2543 | KIF20B | CCTTCCCCACACATTCCCGC |
| 2544 | KIF20B | GGTGACCCGCGGGAATGTGT |
| 2545 | KIF20B | TCCTTCCCCACACATTCCCG |
| 2546 | KIF20B | GCAGGTGAGGAGGTGACCCG |
| 2547 | KIF20B | GGCGAAGTTTGCAGGTGAGG |
| 2548 | KIF20B | AGGTGACCCGCGGGAATGTG |
| 2549 | KIF21B | CGGAGCTAGCTGACAGCCGG |
| 2550 | KIF21B | ACCCGCCGCTCCCTACGGCG |
| 2551 | KIF21B | GCGCGGAGCTAGCTGACAGC |
| 2552 | KIF21B | CGCCGCCGCCCGATGGCGCG |
| 2553 | KIF21B | TGCGCGTGTGCCGCGCCGTA |
| 2554 | KIF21B | TGCCGCGCCGTAGGGAGCGG |
| 2555 | KIF21B | CGCCTCCTCGCGCCATCGGG |
| 2556 | KIF23 | AGACTGCGGGACGTTAGGAC |
| 2557 | KIF23 | AGAACTTCGCGGCTAAGGCG |
| 2558 | KIF23 | CAGCAAGAACTAGAACTTCG |
| 2559 | KIF23 | GTCCCGCATGCGCGTTTGGG |
| 2560 | KIF23 | CGCCGCCCAAACGCGCATGC |
| 2561 | KIF23 | GAACTAGAACTTCGCGGCTA |
| 2562 | KIF23 | ACGCCGCCCAAACGCGCATG |
| 2563 | KIF2C | ACGCGGCGTTAAGACTTCGT |
| 2564 | KIF2C | AGGTCAAGGGGACTCGTGAG |
| 2565 | KIF2C | CGCGGCGTTAAGACTTCGTA |
| 2566 | KIF2C | GGCCTGAAGCGACGAGTCCA |
| 2567 | KIF2C | TCGTAGGGTTAGCGAAATTG |
| 2568 | KIF2C | CCGTCCCCGGACACACAGAT |
| 2569 | KIF2C | GGCCATTCGGAGAGTCAGCA |
| 2570 | KIF4A | ATTTAGTTTGGTGAGTTACG |
| 2571 | KIF4A | TTTAGTTTGGTGAGTTACGG |
| 2572 | KIF4A | TAAAAACAGGTTCTGAGCTA |
| 2573 | KIF4A | AACCAAAGCCCTAGGGTACT |
| 2574 | KIF4A | ACAGGGCGGCGGGAGGCCCA |
| 2575 | KIF4A | ACATTTAGTTTGGTGAGTTA |
| 2576 | KIF4A | AAGCTCCGGCTGGGACAGGG |
| 2577 | KLHL7-AS1-001 | GAGGCAAAATTCACTCAGCG |
| 2578 | KLHL7-AS1-001 | AAGGCTTCATGGGACAGGGT |
| 2579 | KLHL7-AS1-001 | AGGCAAAATTCACTCAGCGC |

| | | |
|---|---|---|
| 2580 | KLHL7-AS1-001 | TGCAGTCTCATTGCTCGGAA |
| 2581 | KLHL7-AS1-001 | CGGGGCCTGACACGTCACAG |
| 2582 | KLHL7-AS1-001 | GTGCCTGCAGTCTCATTGCT |
| 2583 | KLHL7-AS1-001 | GCGACCTCGGACCAGCTCCG |
| 2584 | L3MBTL4-AS1-001 | CCACTAAGGATTCTGCTCCA |
| 2585 | L3MBTL4-AS1-001 | TAAGACAGGAACACCCGCCG |
| 2586 | L3MBTL4-AS1-001 | ACTTAAGAGTGGGTGCCCAG |
| 2587 | L3MBTL4-AS1-001 | GCAGATCGGGAGACTAAGAC |
| 2588 | L3MBTL4-AS1-001 | GGATTCTGCTCCAAGGACAG |
| 2589 | L3MBTL4-AS1-001 | ACCCACTCTTAAGTGCAGAT |
| 2590 | L3MBTL4-AS1-001 | AAGACAGGAACACCCGCCGT |
| 2591 | LA16c-380H5.6-001 | TGCAGTGAACTATGATCCTG |
| 2592 | LA16c-380H5.6-001 | TATGATCCTGCGGCCAATCG |
| 2593 | LA16c-380H5.6-001 | GACTAATCCGAAGCTCAGAA |
| 2594 | LA16c-380H5.6-001 | TCAGCCAGGAGAGCGCTGAA |
| 2595 | LA16c-380H5.6-001 | ATTCCCAAGGTCGTTCAGCC |
| 2596 | LA16c-380H5.6-001 | GCCTCGAGCAGCCACTTCCG |
| 2597 | LA16c-380H5.6-001 | TTCAGCCAGGAGAGCGCTGA |
| 2598 | LA16c-60D12.2-001 | AATAGGAACACTTACACTGT |
| 2599 | LA16c-60D12.2-001 | CACCACACTGACTTCCACAA |
| 2600 | LA16c-60D12.2-001 | CCATTCTAACTGGTGTGAGA |
| 2601 | LA16c-60D12.2-001 | GGAAAAAACAGGTGCTGGAG |
| 2602 | LA16c-60D12.2-001 | GGAACACTTACACTGTTGGT |
| 2603 | LA16c-60D12.2-001 | AACCATTGTGGAAGTCAGTG |
| 2604 | LA16c-60D12.2-001 | GTAAACTAGTTCAACCATTG |
| 2605 | LA16c-OS12.2-001 | GGGAGGAATACGGACACGGA |
| 2606 | LA16c-OS12.2-001 | GCAGGTAAGGAATACGGACA |
| 2607 | LA16c-OS12.2-001 | GTAAGGAATACGGACACGGG |
| 2608 | LA16c-OS12.2-001 | ACACGGGAGGAATACGGACA |
| 2609 | LA16c-OS12.2-001 | CGGGAGGAATACGGACACGG |
| 2610 | LA16c-OS12.2-001 | CAGGTAAGGAATACGGACAC |
| 2611 | LA16c-OS12.2-001 | ACACGGAGGGAACAGCGACA |
| 2612 | LINC00106-001 | TCAGGCTTCATCGGACGGCC |
| 2613 | LINC00106-001 | GGTGAGGTTGGCTTTGTGCG |
| 2614 | LINC00106-001 | TTCAGGCTTCATCGGACGGC |
| 2615 | LINC00106-001 | GGGGTGAGGTTGGCTTTGTG |
| 2616 | LINC00106-001 | CCGGGAAGGAATCAGAGCAG |
| 2617 | LINC00106-001 | CCATCCATTCCAGAAGCATT |
| 2618 | LINC00106-001 | GCATTTGGCAAGCACATCCT |
| 2619 | LINC00106-002 | ACAGTTGCTATGCCTGACTA |
| 2620 | LINC00106-002 | TGCCTGACTAGGGCATGACC |
| 2621 | LINC00106-002 | CACAGTTGCTATGCCTGACT |
| 2622 | LINC00106-002 | CCTTTCCCAAGGAGGCTGTG |

| | | |
|---|---|---|
| 2623 | LINC00106-002 | CTGATCCTCCTTTCCCAAGG |
| 2624 | LINC00106-002 | GCCTGACTAGGGCATGACCT |
| 2625 | LINC00106-002 | CATGACCTGGGAGGCCTTTG |
| 2626 | LINC00115-001 | GCCACGCAGTCCCCTCCCTG |
| 2627 | LINC00115-001 | AAACGGATTATCCGGAGCCA |
| 2628 | LINC00115-001 | CGGGTGTGGAGGACGCCGCA |
| 2629 | LINC00115-001 | TCCGGAGCCAAGGTCCGCTC |
| 2630 | LINC00115-001 | GCCGCAGGGAGGGGACTGCG |
| 2631 | LINC00115-001 | GGTTTGGCCACAAAAAGCGG |
| 2632 | LINC00115-001 | GTTTGGCCACAAAAAGCGGA |
| 2633 | LINC00152-003 | GAAACATACTCACCAAGGAG |
| 2634 | LINC00152-003 | ACTCACCAAGGAGAGGCTGG |
| 2635 | LINC00152-003 | CATACTCACCAAGGAGAGGC |
| 2636 | LINC00152-003 | TGGAGCCACCAGCCTCTCCT |
| 2637 | LINC00152-003 | CAGCACAGTTCCTGGGAAGG |
| 2638 | LINC00152-003 | TTCCCACAGCCCAGACTCCA |
| 2639 | LINC00152-003 | GTTGCTTCTGCAGTTTGCCT |
| 2640 | LINC00152-004 | TATCAAGATAGCTTTCGACG |
| 2641 | LINC00152-004 | GGTATCCTCAACCAGGAACA |
| 2642 | LINC00152-004 | TCCTGGTTGAGGATACCTGA |
| 2643 | LINC00152-004 | AAAGAACATTCTGACAAGAC |
| 2644 | LINC00152-004 | AAACTACCGAACCATGTTCC |
| 2645 | LINC00152-004 | CCGAACCATGTTCCTGGTTG |
| 2646 | LINC00152-004 | ACCTTCAGGTATCCTCAACC |
| 2647 | LINC00152-005 | CACAAGTATGAAGAGAATGT |
| 2648 | LINC00152-005 | AATGTCGGGAGAGGAAGTGG |
| 2649 | LINC00152-005 | ACAAGTATGAAGAGAATGTC |
| 2650 | LINC00152-005 | TATGAAGAGAATGTCGGGAG |
| 2651 | LINC00152-005 | AAATTGACATTCCAGACAAG |
| 2652 | LINC00152-005 | TCACAGCACAGTTCCTGGGA |
| 2653 | LINC00152-005 | TGAAGATCTGAAGACAGGCA |
| 2654 | LINC00152-014 | GTCCTGACAGAGGTTGAAGG |
| 2655 | LINC00152-014 | TCCAGCGTGCGTCCTGACAG |
| 2656 | LINC00152-014 | CGTCCTGACAGAGGTTGAAG |
| 2657 | LINC00152-014 | GGAGTACACCATGTGAAGAC |
| 2658 | LINC00152-014 | GTAGCTGAACCACACCAAGC |
| 2659 | LINC00152-014 | TGCGTCCTGACAGAGGTTGA |
| 2660 | LINC00152-014 | ACCTCTGTCAGGACGCACGC |
| 2661 | LINC00174-001 | AGAGATCAGGTGGTAATGTG |
| 2662 | LINC00174-001 | TTGTGATGAAATGCCAGTGG |
| 2663 | LINC00174-001 | AACAGGAGGTGAGTAAGGGT |
| 2664 | LINC00174-001 | TGATTGTGATGAAATGCCAG |
| 2665 | LINC00174-001 | ACACAACAGGAGGTGAGTAA |

| | | |
|---|---|---|
| 2666 | LINC00174-001 | GCCTGTGACCCAGGGATCAG |
| 2667 | LINC00174-001 | GACAGAAGGCAGAGATCAGG |
| 2668 | LINC00174-004 | GCCGCCCCTACTGGAAAGTG |
| 2669 | LINC00174-004 | TCCTCACTTTCCAGTAGGGG |
| 2670 | LINC00174-004 | CCCTCACCTCCTGGATAGGG |
| 2671 | LINC00174-004 | CGCCCTATCCAGGAGGTGAG |
| 2672 | LINC00174-004 | GGCCAGCCGCCCTATCCAGG |
| 2673 | LINC00174-004 | CCCTACCTCCCTCCCGGACG |
| 2674 | LINC00174-004 | GAAGTGAGGACCCCTCTGCC |
| 2675 | LINC00265-001 | CACTGGCCTTGAAGTCCGGA |
| 2676 | LINC00265-001 | AAGAAAAACAGATGCCCAAG |
| 2677 | LINC00265-001 | CATCCGGACTTCAAGGCCAG |
| 2678 | LINC00265-001 | TTGAAGTCCGGATGGAAGCT |
| 2679 | LINC00265-001 | AGAAAACCAGACCCAGCCAC |
| 2680 | LINC00265-001 | TCCTAGTCTCCTTCCCGCTT |
| 2681 | LINC00265-001 | GCAAGCGCCAAGCTTCCATC |
| 2682 | LINC00273-001 | AGCCCGGCGTCCCAGTCGCG |
| 2683 | LINC00273-001 | GGCGTCCCAGTCGCGAGGAT |
| 2684 | LINC00273-001 | CCGCGCCCATCCTCGCGACT |
| 2685 | LINC00273-001 | TCGCGAGGATGGGCGCGGCA |
| 2686 | LINC00273-001 | CCCAGTCGCGAGGATGGGCG |
| 2687 | LINC00273-001 | CGGCGTCCCAGTCGCGAGGA |
| 2688 | LINC00273-001 | ACGCCGGGCTCATCCTCGCG |
| 2689 | LINC00294-001 | TTGTTTGATCTCACTGATGG |
| 2690 | LINC00294-001 | CTGATAACTTTGTGGACCGG |
| 2691 | LINC00294-001 | CAGTCCCAAAAAGCCATTGG |
| 2692 | LINC00294-001 | AGTCCCAAAAAGCCATTGGG |
| 2693 | LINC00294-001 | CCAGTCCCAAAAAGCCATTG |
| 2694 | LINC00294-001 | AGGGTCTTCTGGCCCCCCAA |
| 2695 | LINC00294-001 | ACCCAATCAGTGAAGAAGCC |
| 2696 | LINC00339-001 | GCTGCCACCTCGGAGGACGG |
| 2697 | LINC00339-001 | CGCTCCCCCGTCCTCCGAGG |
| 2698 | LINC00339-001 | CTACGCAGCTGCCACCTCGG |
| 2699 | LINC00339-001 | GCAGCTGCCACCTCGGAGGA |
| 2700 | LINC00339-001 | CGCGCGGCGCTCCAAAGCTC |
| 2701 | LINC00339-001 | CCCCGCTCCCCCGTCCTCCG |
| 2702 | LINC00339-001 | GCGGGGGACGAGGAAAGATC |
| 2703 | LINC00339-008 | GCGCGGCGCTCCAAAGCTCG |
| 2704 | LINC00339-008 | CATGACTTCCGGTTCCCGCG |
| 2705 | LINC00339-008 | CCGAGCTTTGGAGCGCCGCG |
| 2706 | LINC00339-008 | CTCCAAAGCTCGGGGAGTGG |
| 2707 | LINC00339-008 | TAACTCCCACAAATGACTCC |
| 2708 | LINC00339-008 | CCACAAATGACTCCAGGAAA |

| | | |
|---|---|---|
| 2709 | LINC00339-008 | GCGCTCCAAAGCTCGGGGAG |
| 2710 | LINC00342-002 | TGAGGCAACCTTGCATCCTG |
| 2711 | LINC00342-002 | ATTAATTGACGTGTATGTTG |
| 2712 | LINC00342-002 | TCTAAAAAATCTGGGTATTG |
| 2713 | LINC00342-002 | CATATATTGAGATAATCACA |
| 2714 | LINC00342-002 | AAGGCTTTCAATAAAATTCA |
| 2715 | LINC00342-002 | TTAAAAACTCTAAAAAATCT |
| 2716 | LINC00342-002 | TGCTTCATCCCCAGGATGCA |
| 2717 | LINC00352-001 | GAAATCTTGGAGCATACGGG |
| 2718 | LINC00352-001 | TGTATATGTGCGCGTATGTG |
| 2719 | LINC00352-001 | TGGAAATCTTGGAGCATACG |
| 2720 | LINC00352-001 | TCACAAAACATAAGCACTGT |
| 2721 | LINC00352-001 | GGAAATCTTGGAGCATACGG |
| 2722 | LINC00352-001 | TATGTATATGTGCGCGTATG |
| 2723 | LINC00352-001 | GTGCGCGTATGTGGGGATGT |
| 2724 | LINC00426-004 | CAGGCAGCGTAATCTGACAA |
| 2725 | LINC00426-004 | TACCTTAAACTGCCACCGCA |
| 2726 | LINC00426-004 | AAGTGACCGTCTGCACATAG |
| 2727 | LINC00426-004 | CTGCGGTGGCAGTTTAAGGT |
| 2728 | LINC00426-004 | TGGGAAGGCTCTAGACGTGT |
| 2729 | LINC00426-004 | TGTCAGATTACGCTGCCTGA |
| 2730 | LINC00426-004 | AATTGGTATAAGGCTGAAAG |
| 2731 | LINC00426-005 | AAATTCATGTTGGAAGACGT |
| 2732 | LINC00426-005 | GAAGACGTGGGCTCTAGCTG |
| 2733 | LINC00426-005 | TATGCTGTAGAGGCCGGGCG |
| 2734 | LINC00426-005 | AAAATTCATGTTGGAAGACG |
| 2735 | LINC00426-005 | GTCAAGGCAGGCAGATCACG |
| 2736 | LINC00426-005 | TAAAATATGCTGTAGAGGCC |
| 2737 | LINC00426-005 | CCTTGACCTCCCAAAGTGCT |
| 2738 | LINC00430-002 | GAGTTGTCTGATTCTCATCG |
| 2739 | LINC00430-002 | AATTCGTCCTTAAAGGTCAA |
| 2740 | LINC00430-002 | ATGAGAATCAGACAACTCTG |
| 2741 | LINC00430-002 | TGTACTCTGAAATGGTGAGC |
| 2742 | LINC00430-002 | TGAAGTTAGTAAGCCTGACA |
| 2743 | LINC00430-002 | GGTCAAGGGCCCACAAAGGA |
| 2744 | LINC00430-002 | TAAAGGTCAAGGGCCCACAA |
| 2745 | LINC00467-001 | GCGCTGTGACGTTCCCTACG |
| 2746 | LINC00467-001 | GACAACCTCAACCGGACCGA |
| 2747 | LINC00467-001 | AAGGTGCTGAACAACCAGTG |
| 2748 | LINC00467-001 | GTTGTCAAGTCGGACCAAAC |
| 2749 | LINC00467-001 | TGTGACGTTCCCTACGCGGT |
| 2750 | LINC00467-001 | GTGACGTTCCCTACGCGGTC |
| 2751 | LINC00467-001 | GTTGTTCAGCACCTTCGGTC |

| | | |
|---|---|---|
| 2752 | LINC00467-003 | GTCCCCTCGGAGTGACCGAA |
| 2753 | LINC00467-003 | GAGTGACCGAAGGGAGATTG |
| 2754 | LINC00467-003 | GGGAGATTGAGGAAAGAGTG |
| 2755 | LINC00467-003 | GGGGCTACCAACAGTCCCCT |
| 2756 | LINC00467-003 | CTCCCTTCGGTCACTCCGAG |
| 2757 | LINC00467-003 | AGTCCCCTCGGAGTGACCGA |
| 2758 | LINC00467-003 | GGTCACTCCGAGGGGACTGT |
| 2759 | LINC00467-004 | AGATTACTCACCACTCGTCG |
| 2760 | LINC00467-004 | GTGTTCAACTCCCCGACGAG |
| 2761 | LINC00467-004 | ACGTGGGCGCCAGATATCGG |
| 2762 | LINC00467-004 | ACAGATTACTCACCACTCGT |
| 2763 | LINC00467-004 | CGGTGAAACATCAATCGGCT |
| 2764 | LINC00467-004 | GAGTTGAACACGCTTATCTG |
| 2765 | LINC00467-004 | GTAAGGTCTGGGTTAACTAT |
| 2766 | LINC00494-001 | ACCTGCAACTGCATGATGTG |
| 2767 | LINC00494-001 | GCAACTGCATGATGTGCGGG |
| 2768 | LINC00494-001 | GCTGTTCTAAAGGACGATAG |
| 2769 | LINC00494-001 | CTAAAGGACGATAGAGGTGA |
| 2770 | LINC00494-001 | TAAAGGACGATAGAGGTGAA |
| 2771 | LINC00494-001 | AGGCCTCCTGACCCCTGACA |
| 2772 | LINC00494-001 | CCTGCAACTGCATGATGTGC |
| 2773 | LINC00494-004 | GCTTGCCCACCTCCTCAGAG |
| 2774 | LINC00494-004 | GGGAAGAGCACCCCAAGCAG |
| 2775 | LINC00494-004 | GAAGGGCAAGGCCGTGTCAG |
| 2776 | LINC00494-004 | TGAGGAGGTGGGCAAGCCCT |
| 2777 | LINC00494-004 | GGAGGCCTCTCTGAGGAGGT |
| 2778 | LINC00494-004 | TGCTTGGGGTGCTCTTCCCA |
| 2779 | LINC00494-004 | ACCTGCTGTGCCTCTGCTTG |
| 2780 | LINC00504-001 | TGTAGGCCTCAGAGTCATGG |
| 2781 | LINC00504-001 | AAATATATGTCTTGTTTACT |
| 2782 | LINC00504-001 | GGCGAAAGGCACTTCTAACA |
| 2783 | LINC00504-001 | ATAAACACATACCTGAAACT |
| 2784 | LINC00504-001 | GAGTCATGGTGGGAGGCGAA |
| 2785 | LINC00504-001 | GGAACGAAAAGAGGTTTAAG |
| 2786 | LINC00504-001 | TATGCAATCATTTGGCTTAA |
| 2787 | LINC00504-003 | GCAGAGGCAAGAGACAAATG |
| 2788 | LINC00504-003 | AGTGGACTTGCAGTTCGACG |
| 2789 | LINC00504-003 | GCAGTTCGACGTGGCTGTGT |
| 2790 | LINC00504-003 | GGCCTCAGAGTCATGGTGGG |
| 2791 | LINC00504-003 | AGGCACTTCTAACATGGCAG |
| 2792 | LINC00504-003 | GTAGGCCTCAGAGTCATGGT |
| 2793 | LINC00504-003 | CATGAAAATAAGGTCGCTTC |
| 2794 | LINC00506-001 | TGCGAAAGGAGGGAACGTGT |

| | | |
|---|---|---|
| 2795 | LINC00506-001 | GGGAACGTGTGGGTCTGCCA |
| 2796 | LINC00506-001 | AACCGACTTGTGCGAAAGGA |
| 2797 | LINC00506-001 | CAACCGACTTGTGCGAAAGG |
| 2798 | LINC00506-001 | GTGCGAAAGGAGGGAACGTG |
| 2799 | LINC00506-001 | TTCCCTCCTTTCGCACAAGT |
| 2800 | LINC00506-001 | CAGCAACCGACTTGTGCGAA |
| 2801 | LINC00511-002 | GCGCTGGTATCCAGAGGACG |
| 2802 | LINC00511-002 | TGCCTAACAGAGGGTCCTTG |
| 2803 | LINC00511-002 | AGAACGACAAATGCCAGAGG |
| 2804 | LINC00511-002 | GGTGCGGCGCTGGTATCCAG |
| 2805 | LINC00511-002 | CGCAGAACGACAAATGCCAG |
| 2806 | LINC00511-002 | TCGTTCTGCGCTTCTCCGCA |
| 2807 | LINC00511-002 | GAGGCGGTGACCGCGTCCTC |
| 2808 | LINC00528-001 | GGCCTCGTTCTGTTGCAAAG |
| 2809 | LINC00528-001 | TTCAAGTCAGCCCACAACAG |
| 2810 | LINC00528-001 | TACCCCTTTGCAACAGAACG |
| 2811 | LINC00528-001 | TGGACACAGGGACTTTCCAG |
| 2812 | LINC00528-001 | CCTTTGCAACAGAACGAGGC |
| 2813 | LINC00528-001 | ACACCAGTCAGATAGGAGGA |
| 2814 | LINC00528-001 | CTGTTGCAAAGGGGTAACTG |
| 2815 | LINC00571-001 | TCTAAAGTACGTACATTTGG |
| 2816 | LINC00571-001 | AATAAGCATCCTTCCATGCA |
| 2817 | LINC00571-001 | ATAAGCATCCTTCCATGCAA |
| 2818 | LINC00571-001 | AAATTTAAGAACCCCTTGCA |
| 2819 | LINC00571-001 | TAAGCATCCTTCCATGCAAG |
| 2820 | LINC00571-001 | ACCATAGCGCTGAACCTTGA |
| 2821 | LINC00571-001 | AAGTACGTACATTTGGGGGA |
| 2822 | LINC00623-001 | AGCTGTTGCCAAATGAACCC |
| 2823 | LINC00623-001 | TCGCGGTCAGAGAACTGGGA |
| 2824 | LINC00623-001 | TTCTCTGACCGCGACGTGTG |
| 2825 | LINC00623-001 | CGGTCAGAGAACTGGGAAGG |
| 2826 | LINC00623-001 | GCCAAATGAACCCCGGAATG |
| 2827 | LINC00623-001 | ACCCCCAGCCACACACGTCG |
| 2828 | LINC00623-001 | GCGGTCAGAGAACTGGGAAG |
| 2829 | LINC00641-001 | AACCCAAGATAAATGCAGAT |
| 2830 | LINC00641-001 | GGGGTGAATGCAAATGACAA |
| 2831 | LINC00641-001 | AGATAAATGCAGATAGGAAG |
| 2832 | LINC00641-001 | AATAAAAGTTTGGAGAAACA |
| 2833 | LINC00641-001 | TGACAAAGGGATTAGTAGGA |
| 2834 | LINC00641-001 | TTATTACATTACCAAATTTG |
| 2835 | LINC00641-001 | AATGTGCATTAAAATTTCCT |
| 2836 | LINC00657-001 | CTGCCAAGTCAGTTCCGGTC |
| 2837 | LINC00657-001 | GAACGCAGCCCGCTCCTCCA |

| | | |
|---|---|---|
| 2838 | LINC00657-001 | AGAACGCAGCCCGCTCCTCC |
| 2839 | LINC00657-001 | CCGGGGCCGGAGGGCCTGGA |
| 2840 | LINC00657-001 | GAGGGCCTGGAGGGCCCTGG |
| 2841 | LINC00657-001 | ACCCGCCGGCCCGGGGCCGG |
| 2842 | LINC00657-001 | CCAGGGCCCTCCAGGCCCTC |
| 2843 | LINC00662-003 | CCTGGGTTAGACCTCGCACA |
| 2844 | LINC00662-003 | CACCAGTTTCAGAAGCGTGT |
| 2845 | LINC00662-003 | CCCTGTGCGAGGTCTAACCC |
| 2846 | LINC00662-003 | AACAACCTCAGTTCTGTGAA |
| 2847 | LINC00662-003 | GAAGGAGTGTGGCAAAGGGT |
| 2848 | LINC00662-003 | AACCTCAGTTCTGTGAAAGG |
| 2849 | LINC00662-003 | GCGAGGTCTAACCCAGGTGT |
| 2850 | LINC00662-004 | TCGGCCCTGCCCATAGAAAG |
| 2851 | LINC00662-004 | TGCGCATCTCTGAAGGAGTG |
| 2852 | LINC00662-004 | CTCTGAAGGAGTGTGGCAAA |
| 2853 | LINC00662-004 | GCAGCACTGCGCATCTCTGA |
| 2854 | LINC00662-004 | TGAAGGAGTGTGGCAAAGGG |
| 2855 | LINC00662-004 | GCTGCCCCTTTCTATGGGCA |
| 2856 | LINC00662-004 | GCAGTGCTGCCCCTTTCTAT |
| 2857 | LINC00667-001 | TCCTGACCTAGAACCATGCC |
| 2858 | LINC00667-001 | CGCGTTGGCTGTCATGATGT |
| 2859 | LINC00667-001 | CTAGAACCATGCCCGGCGCA |
| 2860 | LINC00667-001 | GGGCATGGTTCTAGGTCAGG |
| 2861 | LINC00667-001 | CGCAGGCCTTGCTCCCCGCT |
| 2862 | LINC00667-001 | AGCGGGGAGCAAGGCCTGCG |
| 2863 | LINC00667-001 | GCCGGGCATGGTTCTAGGTC |
| 2864 | LINC00667-004 | GGCGGAGGAATCTGACTGCA |
| 2865 | LINC00667-004 | GAAGAAAAGCTAGAGAGCGA |
| 2866 | LINC00667-004 | TCAGATTCCTCCGCCGACCG |
| 2867 | LINC00667-004 | GAGAGCGAAGGCAAAGCCCG |
| 2868 | LINC00667-004 | AAGGCAAAGCCCGAGGAGAG |
| 2869 | LINC00667-004 | GCTAGGCCTGGCCTTGTCCG |
| 2870 | LINC00667-004 | AGGGGCGGACTCCGCGGACA |
| 2871 | LINC00667-005 | GCGATCAAGCTTCCCAGCGA |
| 2872 | LINC00667-005 | TAGGAGTGAAGCGGCTCGCG |
| 2873 | LINC00667-005 | CCGCTTCACTCCTACAAGCC |
| 2874 | LINC00667-005 | GGCGATCAAGCTTCCCAGCG |
| 2875 | LINC00667-005 | CCGGGCTTGTAGGAGTGAAG |
| 2876 | LINC00667-005 | AGGGTCAAAGTCCAGCAGCT |
| 2877 | LINC00667-005 | GCAGCTTGGAACTAGTGAGC |
| 2878 | LINC00667-006 | AGAACAACGAGCTTACGGAC |
| 2879 | LINC00667-006 | GAACAACGAGCTTACGGACC |
| 2880 | LINC00667-006 | CAGGGAGAACAACGAGCTTA |

| | | |
|---|---|---|
| 2881 | LINC00667-006 | AGCTTACGGACCGGGTGCGG |
| 2882 | LINC00667-006 | ACGAGCTTACGGACCGGGTG |
| 2883 | LINC00667-006 | GAGCTTACGGACCGGGTGCG |
| 2884 | LINC00667-006 | CGAGCTTACGGACCGGGTGC |
| 2885 | LINC00847-002 | ACCGGCCTATGCGCCAAGCG |
| 2886 | LINC00847-002 | GTGTTCTTGTAAATTAGAAC |
| 2887 | LINC00847-002 | AAAAAAACATGGCCGAAACC |
| 2888 | LINC00847-002 | GACACAGGCCGGGAGAAGAG |
| 2889 | LINC00847-002 | GAAACCAGGAAGCTGCGCGA |
| 2890 | LINC00847-002 | CCCGCGCTTGGCGCATAGGC |
| 2891 | LINC00847-002 | CCGGCCTATGCGCCAAGCGC |
| 2892 | LINC00847-004 | GCTGCATCACCCTAGAAACG |
| 2893 | LINC00847-004 | GCCCCAGTAAGTTCCTGGCG |
| 2894 | LINC00847-004 | CAGCCCCAGTAAGTTCCTGG |
| 2895 | LINC00847-004 | TCCCCGCCAGGAACTTACTG |
| 2896 | LINC00847-004 | GCGAGGAGAGACTTCCACAG |
| 2897 | LINC00847-004 | CTGTGGAAACGCTGCCTCTG |
| 2898 | LINC00847-004 | AGCCCAAAGCCACGTTTCTA |
| 2899 | LINC00863-001 | GGAATAAAGAGAGACATCGG |
| 2900 | LINC00863-001 | TGGAATAAAGAGAGACATCG |
| 2901 | LINC00863-001 | GCTGGAATAAAGAGAGACAT |
| 2902 | LINC00863-001 | CTGGAATAAAGAGAGACATC |
| 2903 | LINC00863-001 | GCGCCGGCCAGTTTGACAGC |
| 2904 | LINC00863-001 | CAGGGACGCTGGCCTTCTCG |
| 2905 | LINC00863-001 | CGCCGGCCAGTTTGACAGCC |
| 2906 | LINC00869-001 | TTTCGTGGCGAAGCAGCAGG |
| 2907 | LINC00869-001 | AGGCGTTGTCGGTGACATCA |
| 2908 | LINC00869-001 | TTGTCGGTGACATCACGGAG |
| 2909 | LINC00869-001 | ATGGCTCCGGGATGCGCGTG |
| 2910 | LINC00869-001 | TGTCGGTGACATCACGGAGA |
| 2911 | LINC00869-001 | AGAGCAGACGGGGATCCGGA |
| 2912 | LINC00869-001 | CACTTGCCTCACGCGCATCC |
| 2913 | LINC00869-003 | CGGTCGGAGAACTGAGAAAG |
| 2914 | LINC00869-003 | GGATCACTTCCAAAACCACT |
| 2915 | LINC00869-003 | AGAACTGAGAAAGGGGCGCT |
| 2916 | LINC00869-003 | GCGGTCGGAGAACTGAGAAA |
| 2917 | LINC00869-003 | AGAAAGGGGCGCTGGGTCCG |
| 2918 | LINC00869-003 | ACGAAAGCTGAAAGCGTGCG |
| 2919 | LINC00869-003 | GAGAACTGAGAAAGGGGCGC |
| 2920 | LINC00869-017 | CTCCTTTGACAGTCAGCCCA |
| 2921 | LINC00869-017 | CCTGGATTGTAGCTTTCCCT |
| 2922 | LINC00869-017 | TGTTGCCACATCAGATAACA |
| 2923 | LINC00869-017 | GATCTGGAAGAAAAAGACTC |

| | | |
|---|---|---|
| 2924 | LINC00869-017 | CCTCCTTTGACAGTCAGCCC |
| 2925 | LINC00869-017 | TCCATCCTTTAGGAAGACCC |
| 2926 | LINC00869-017 | TTCCCTGGGCTGACTGTCAA |
| 2927 | LINC00869-018 | ATCACCAATCCATCGCCAGA |
| 2928 | LINC00869-018 | TCACCAATCCATCGCCAGAA |
| 2929 | LINC00869-018 | CAATCCATCGCCAGAAGGGG |
| 2930 | LINC00869-018 | AGCGGCGAGCAGCAGCTGGA |
| 2931 | LINC00869-018 | CACCAATCCATCGCCAGAAG |
| 2932 | LINC00869-018 | TCCTCCTCCTCCTCTCACCC |
| 2933 | LINC00869-018 | CGCCCGGACCAACCGCCGTG |
| 2934 | LINC00877-004 | GCACGGAGTTTCCCTTCGCG |
| 2935 | LINC00877-004 | CTGCACGGAGTTTCCCTTCG |
| 2936 | LINC00877-004 | TCCCCAGCGGGCCCCGCGAA |
| 2937 | LINC00877-004 | AGCCGCGGATGCTTCCCCAG |
| 2938 | LINC00877-004 | TTCCCCAGCGGGCCCCGCGA |
| 2939 | LINC00877-004 | TGCACGGAGTTTCCCTTCGC |
| 2940 | LINC00877-004 | GAAGCATCCGCGGCTTGTGG |
| 2941 | LINC00891-001 | ACCCGCGATAGTTTACGCAT |
| 2942 | LINC00891-001 | GACCCATGCGTAAACTATCG |
| 2943 | LINC00891-001 | ATGCGTAAACTATCGCGGGT |
| 2944 | LINC00891-001 | AACCCGCGATAGTTTACGCA |
| 2945 | LINC00891-001 | ACCCATGCGTAAACTATCGC |
| 2946 | LINC00891-001 | CTCAGACAAATCCAGCTGAG |
| 2947 | LINC00891-001 | CTTGAAGTTACGCGTGAAGG |
| 2948 | LINC00908-005 | CTACTAAGGTAAATATGGAA |
| 2949 | LINC00908-005 | GAGAGAAGAAAGGGAAGACG |
| 2950 | LINC00908-005 | CGGGGAGCCAGGCAGGTGCG |
| 2951 | LINC00908-005 | GGGGAGCCAGGCAGGTGCGT |
| 2952 | LINC00908-005 | TACTAAGGTAAATATGGAAT |
| 2953 | LINC00908-005 | AGCCAGGCAGGTGCGTGGGA |
| 2954 | LINC00908-005 | TTCTTGTTTGGGGATTGGAG |
| 2955 | LINC00909-001 | CTTCCAAGGCGACTCCGTAG |
| 2956 | LINC00909-001 | AAAAAACAAGGAACTCGACC |
| 2957 | LINC00909-001 | GTCCGGCGCTGCATCCACTA |
| 2958 | LINC00909-001 | TCCCACCCTTCCCTTAGAGA |
| 2959 | LINC00909-001 | TCCTTCTCTAAGGGAAGGGT |
| 2960 | LINC00909-001 | GAACTGGTTTAGTACAATCC |
| 2961 | LINC00909-001 | GAAGGAAAACGGAGAGCCGA |
| 2962 | LINC00909-002 | GAGCTCTTGTGGACACCTCG |
| 2963 | LINC00909-002 | GTGCCCGGGTTGCTCTCAAG |
| 2964 | LINC00909-002 | TTTCCACTTGAGAGCAACCC |
| 2965 | LINC00909-002 | TTAGCGTCTGGTTTCCAGCA |
| 2966 | LINC00909-002 | GCAACCCGGGCACGCGAGAA |

| | | |
|---|---|---|
| 2967 | LINC00909-002 | GCTTGCAGCTGTGCAGCTAG |
| 2968 | LINC00909-002 | TAGCGTCTGGTTTCCAGCAG |
| 2969 | LINC00923-001 | CCATCACGTCAGCAGTCACA |
| 2970 | LINC00923-001 | ATTTAACAGTTCATTCAGAG |
| 2971 | LINC00923-001 | TTCATTCAGAGAGGGAACCA |
| 2972 | LINC00923-001 | ATACAGCAACAGGCCCAGTG |
| 2973 | LINC00923-001 | TCCATCACGTCAGCAGTCAC |
| 2974 | LINC00923-001 | TACAGCAACAGGCCCAGTGA |
| 2975 | LINC00923-001 | TGACGTGGAAGACGCCGAAG |
| 2976 | LINC00926-003 | TGATACTCCCTAAATCACAA |
| 2977 | LINC00926-003 | ATGATACTCCCTAAATCACA |
| 2978 | LINC00926-003 | GAAATGGCTGCACGTGACAG |
| 2979 | LINC00926-003 | GTTGGAGAACAGGAAGCAAG |
| 2980 | LINC00926-003 | AGCACTTTGGAAGGCCAAGG |
| 2981 | LINC00926-003 | TGTAATCCCAGCACTTTGGA |
| 2982 | LINC00926-003 | CTCCCTAAATCACAAGGGTG |
| 2983 | LINC00936-001 | GCTCCGCAGAGCGGCACACA |
| 2984 | LINC00936-001 | GGCTCCGCAGAGCGGCACAC |
| 2985 | LINC00936-001 | GCGCCCTGTGTGCCGCTCTG |
| 2986 | LINC00936-001 | AGGTGGCTGCACCCCGCCCG |
| 2987 | LINC00936-001 | GCGGCTGCTGTTGCGGACGG |
| 2988 | LINC00936-001 | CTGTTGCGGACGGCGGCGTG |
| 2989 | LINC00936-001 | CTGCCCGTGCCGAGCGAGCG |
| 2990 | LINC00937-001 | GGGCGCCAGAGGACCACGAG |
| 2991 | LINC00937-001 | GGACCACGAGCGGCCCGCGG |
| 2992 | LINC00937-001 | AGAGGACCACGAGCGGCCCG |
| 2993 | LINC00937-001 | GCTCCTCCGCGGGCCGCTCG |
| 2994 | LINC00937-001 | CCTTCTTCTTCCGCAAGTGT |
| 2995 | LINC00937-001 | CTGCGCAAGCGGCCTCTCGG |
| 2996 | LINC00937-001 | TGCCAGTGGCTTCGGAGCTG |
| 2997 | LINC00937-003 | GCCAGGGCCCTACCTGTCCA |
| 2998 | LINC00937-003 | GACAGCGACTGCTTGGTCCG |
| 2999 | LINC00937-003 | GCGCGCACCGTGGACAGGTA |
| 3000 | LINC00937-003 | GGCAGGAGACAGCGACTGCT |
| 3001 | LINC00937-003 | CAGTCGCTGTCTCCTGCCTG |
| 3002 | LINC00937-003 | CTGTCCACGGTGCGCGCGAG |
| 3003 | LINC00937-003 | CGCGCGCACCGTGGACAGGT |
| 3004 | LINC00937-005 | TCAACCCAGGGCAGGCCACG |
| 3005 | LINC00937-005 | AGGACGGGCAGGACGCGCCG |
| 3006 | LINC00937-005 | CGCACAGCCCCAGGACTGCG |
| 3007 | LINC00937-005 | CCAGGACTGCGAGGACGGGC |
| 3008 | LINC00937-005 | TGCCCGTCCTCGCAGTCCTG |
| 3009 | LINC00937-005 | CTGCGGCTCTTGCTGGAGGG |

| | | |
|---|---|---|
| 3010 | LINC00937-005 | AGCCCCAGGACTGCGAGGAC |
| 3011 | LINC00957-001 | GGACTTCATCACCAAGCTCG |
| 3012 | LINC00957-001 | CACCAAGCTCGTGGACATCG |
| 3013 | LINC00957-001 | CGCCACGCCCTGGCGCACGG |
| 3014 | LINC00957-001 | CTGCGCCACGCCCTGGCGCA |
| 3015 | LINC00957-001 | TGGCGCACGGTGGGCTGCAG |
| 3016 | LINC00957-001 | CTCCTCGATGTCCACGAGCT |
| 3017 | LINC00957-001 | CTGCAGCCCACCGTGCGCCA |
| 3018 | LINC00957-002 | GGACACGCGTCAGTAGCAGG |
| 3019 | LINC00957-002 | GGGGCTATGGAGCTCAAGGA |
| 3020 | LINC00957-002 | CAGGTGGACTGAGGAAGGCA |
| 3021 | LINC00957-002 | GCGGGGTGACAGGTGGACTG |
| 3022 | LINC00957-002 | GGTGACAGGTGGACTGAGGA |
| 3023 | LINC00957-002 | GCGGGACACGCGTCAGTAGC |
| 3024 | LINC00957-002 | CACGCGTCAGTAGCAGGTGG |
| 3025 | LINC00960-001 | CAGGCGTCATAACCAACCCG |
| 3026 | LINC00960-001 | CAGGCGCGCGTATCTGGACC |
| 3027 | LINC00960-001 | CCAGGCGTCATAACCAACCC |
| 3028 | LINC00960-001 | ATAACCAACCCGGGGCCGCG |
| 3029 | LINC00960-001 | TCCAGGCGTCATAACCAACC |
| 3030 | LINC00960-001 | TGGTTATGACGCCTGGAATC |
| 3031 | LINC00960-001 | CCCGGGTTGGTTATGACGCC |
| 3032 | LINC00963-002 | ACATCAGCCGGCCGCGAGGG |
| 3033 | LINC00963-002 | TCCGGGAAACGGCGGCCTGG |
| 3034 | LINC00963-002 | GGAAACGGCGGCCTGGCGGG |
| 3035 | LINC00963-002 | GGGCCGGCTCCGGGAAACGG |
| 3036 | LINC00963-002 | CATCAGCCGGCCGCGAGGGC |
| 3037 | LINC00963-002 | AGGCCGCCGTTTCCCGGAGC |
| 3038 | LINC00963-002 | GGCTCCGGGAAACGGCGGCC |
| 3039 | LINC00963-003 | AAGCTCTCTTAGGTAACACG |
| 3040 | LINC00963-003 | ATTACCGTATGCAGTCCCCC |
| 3041 | LINC00963-003 | CCTTTGACCCTGTATCCTGG |
| 3042 | LINC00963-003 | CATCGCAGTTGGAAGCCCAT |
| 3043 | LINC00963-003 | AGTCACTGTTGTCACTACTG |
| 3044 | LINC00963-003 | CAGTGACTCACCTGCCCCAT |
| 3045 | LINC00963-003 | GAGAGCACAACCATCGCAGT |
| 3046 | LINC00963-007 | GTCCTACTGCAGATGATGTG |
| 3047 | LINC00963-007 | CCATGGGCTTCCAACTGCGA |
| 3048 | LINC00963-007 | CATCATCTGCAGTAGGACGG |
| 3049 | LINC00963-007 | TGCCCCACATCATCTGCAGT |
| 3050 | LINC00963-007 | ACATCATCTGCAGTAGGACG |
| 3051 | LINC00963-007 | CGTCCTACTGCAGATGATGT |
| 3052 | LINC00963-007 | AACTGTGGTCAGTAGCAGAG |

| | | |
|---|---|---|
| 3053 | LINC00963-011 | CTCAGGGGCTCCCGAGTGTG |
| 3054 | LINC00963-011 | GAGGTCCGGGAAGGGGAATG |
| 3055 | LINC00963-011 | GAGCCGCCCTCCCCACACTC |
| 3056 | LINC00963-011 | TGAGCCGCCCTCCCCACACT |
| 3057 | LINC00963-011 | TTCTCAGGGGCTCCCGAGTG |
| 3058 | LINC00963-011 | AGGGGCTCCCGAGTGTGGGG |
| 3059 | LINC00963-011 | AACGGGAAAGAGAGCAGGAG |
| 3060 | LINC00969-001 | ACACAAAAATTAGCCAGGTG |
| 3061 | LINC00969-001 | TAACACGTGGTGATATCTCC |
| 3062 | LINC00969-001 | GGGAAACGCTTCTTTAAACC |
| 3063 | LINC00969-001 | GATACAAAGCAACATTTCAC |
| 3064 | LINC00969-001 | AGAATCAACAAAATAAGGCC |
| 3065 | LINC00969-001 | TGAAAACACAAAAATTAGCC |
| 3066 | LINC00969-001 | TCAGGAATTCAAGACCAACC |
| 3067 | LINC00969-035 | CCCAGTGCCGACGTCCACAT |
| 3068 | LINC00969-035 | GCCGACGTCCACATAGGACA |
| 3069 | LINC00969-035 | AGCCCTTTGAGGTGCACTGG |
| 3070 | LINC00969-035 | TACTCATCAATCTGCACCTG |
| 3071 | LINC00969-035 | TGCCGACGTCCACATAGGAC |
| 3072 | LINC00969-035 | GATCACTCCAAGCCCATCCA |
| 3073 | LINC00969-035 | AGAAGCCCTTTGAGGTGCAC |
| 3074 | LINC00969-038 | AAGGCGGTGAGTCCCTGCCG |
| 3075 | LINC00969-038 | AGGCCCCCGAGTACACTCCG |
| 3076 | LINC00969-038 | GAGCCGCGGAGTGTACTCGG |
| 3077 | LINC00969-038 | GCACCACGGCCGGGTCAAGG |
| 3078 | LINC00969-038 | GAGCTCGGTCCTTAGCAGGT |
| 3079 | LINC00969-038 | GGAGTGTACTCGGGGGCCTG |
| 3080 | LINC00969-038 | ACTAGCGCCCCGCCTTGACC |
| 3081 | LINC00969-039 | GGCGACTACAGACGTGTCGG |
| 3082 | LINC00969-039 | CGGCGACTACAGACGTGTCG |
| 3083 | LINC00969-039 | GGCGGCGACTACAGACGTGT |
| 3084 | LINC00969-039 | GGCGCTGGCGCTGGCCAAGG |
| 3085 | LINC00969-039 | TGTCGCGGTTGCCAAGCGCT |
| 3086 | LINC00969-039 | TTGGCAACCGCGACAGGCCC |
| 3087 | LINC00969-039 | GCGGCGACTACAGACGTGTC |
| 3088 | LINC00969-040 | GCGGCAGGGACTCACCGCCT |
| 3089 | LINC00969-040 | CTTCCCGATCCCCGGGCAGG |
| 3090 | LINC00969-040 | GGTGCCCGCGCCCCCTGCCC |
| 3091 | LINC00969-040 | CCAGCCCCTTCCCGATCCCC |
| 3092 | LINC00969-040 | CGGTCCCCAGCGGGTCCGAG |
| 3093 | LINC00969-040 | GTGCCCGCGCCCCCTGCCCG |
| 3094 | LINC00969-040 | GGGTGCCCGCGCCCCCTGCC |
| 3095 | LINC00969-041 | TCCGCGGCTCCCCCTCGCAC |

| | | |
|---|---|---|
| 3096 | LINC00969-041 | AGCCCTGGGCCGGTGCGAGG |
| 3097 | LINC00969-041 | CCCTGCCCGGGGATCGGGAA |
| 3098 | LINC00969-041 | CCTTCCCGATCCCCGGGCAG |
| 3099 | LINC00969-041 | CCGGGGATCGGGAAGGGGCT |
| 3100 | LINC00969-041 | CCCCTGCCCGGGGATCGGGA |
| 3101 | LINC00969-041 | CGCGCCCCCTGCCCGGGGAT |
| 3102 | LINC00969-044 | GCATTCCCACCAGCTACGAG |
| 3103 | LINC00969-044 | ACCGTGCATTATAACATGGA |
| 3104 | LINC00969-044 | CGGCATTCCCACCAGCTACG |
| 3105 | LINC00969-044 | CCATGTTATAATGCACGGTG |
| 3106 | LINC00969-044 | GTCCATGTTATAATGCACGG |
| 3107 | LINC00969-044 | GGCATTCCCACCAGCTACGA |
| 3108 | LINC00969-044 | ACCTGCCCCTCGTAGCTGGT |
| 3109 | LINC00969-051 | GAAAAACGTTTCCTAAGACA |
| 3110 | LINC00969-051 | CAGTTGTGCTCATGTGATGG |
| 3111 | LINC00969-051 | AACGTTTCCTAAGACAAGGC |
| 3112 | LINC00969-051 | ACTAGGAAGGAGTGGACTGT |
| 3113 | LINC00969-051 | TCCACTCCTTCCTAGTGCTG |
| 3114 | LINC00969-051 | ACAGTTGTGCTCATGTGATG |
| 3115 | LINC00969-051 | AACAGTTGTGCTCATGTGAT |
| 3116 | LINC00969-066 | AAAAACAAAAAATTAACAGA |
| 3117 | LINC00969-066 | CCTGTCTCTAAAAATTAGCC |
| 3118 | LINC00969-066 | TGGGACCAGCCTGGGCAAGA |
| 3119 | LINC00969-066 | TAAAAATTAGCCAGGCATGG |
| 3120 | LINC00969-066 | CTCTAAAAATTAGCCAGGCA |
| 3121 | LINC00969-066 | TGTAGTCTCAGCTCTTAGGG |
| 3122 | LINC00969-066 | TTGGGACCAGCCTGGGCAAG |
| 3123 | LINC00969-067 | TGAGACAAAAATTTGGAAGG |
| 3124 | LINC00969-067 | CCCTGAGACAAAAATTTGGA |
| 3125 | LINC00969-067 | AAAATCTAAAGTGGTTGCTG |
| 3126 | LINC00969-067 | CGCCTGTAGAATGCAATCCA |
| 3127 | LINC00969-067 | GTGGTTGCTGTGGAGGACAG |
| 3128 | LINC00969-067 | CTGTGGAGGACAGGGGTCAG |
| 3129 | LINC00969-067 | TGGAGGACAGGGGTCAGTGG |
| 3130 | LINC00969-072 | CTGTGATGAATTCTGCCATG |
| 3131 | LINC00969-072 | GGCCTGTATGTGAGTTACCA |
| 3132 | LINC00969-072 | CAGGGTTGCAGAGGACACTG |
| 3133 | LINC00969-072 | GGTGGCAGTCTGGTGCCACA |
| 3134 | LINC00969-072 | GGGCCTGTATGTGAGTTACC |
| 3135 | LINC00969-072 | GACGGATCACCGCTGCAAGA |
| 3136 | LINC00969-072 | ACGGATCACCGCTGCAAGAG |
| 3137 | LINC00969-126 | TTCATATGAGTATGCATGCA |
| 3138 | LINC00969-126 | AGACATACGGCTGGGCGCAG |

| | | |
|---|---|---|
| 3139 | LINC00969-126 | CCAAGGCAGGTGGATCACTG |
| 3140 | LINC00969-126 | GCAGGTGGATCACTGAGGTC |
| 3141 | LINC00969-126 | ATACAAAAATTAGCCAAGTG |
| 3142 | LINC00969-126 | CGACACCAGCCTGGCCAACA |
| 3143 | LINC00969-126 | CAAAAATTAGCCAAGTGTGG |
| 3144 | LINC00969-129 | CAGCTTTGTGTGGATGTACT |
| 3145 | LINC00969-129 | TGGAAGAAGCATAAGAACAT |
| 3146 | LINC00969-129 | TCCAATTAAACCAAATGCTG |
| 3147 | LINC00969-129 | CTCCAATTAAACCAAATGCT |
| 3148 | LINC00969-129 | GATGCAGGTGCAGCTTTGTG |
| 3149 | LINC00969-129 | CTTTGTGTGGATGTACTGGG |
| 3150 | LINC00969-129 | GACAAATTGAGATTTGCTGA |
| 3151 | LINC00969-130 | GGTTAGAAGTGCAGCCAGCG |
| 3152 | LINC00969-130 | GTTTCAAGCCTGCCCTATGG |
| 3153 | LINC00969-130 | AAACTGGCATTTCCTCCATA |
| 3154 | LINC00969-130 | GTGGATGCCGACCCAAAGCA |
| 3155 | LINC00969-130 | GCTGGCTGCACTTCTAACCT |
| 3156 | LINC00969-130 | CTTCTAACAGCAAAGCACAT |
| 3157 | LINC00969-130 | GCATCCACTGATGCCAGCAG |
| 3158 | LINC00969-132 | CAAGTGTTGAAGTGTTGCTG |
| 3159 | LINC00969-132 | AAGTGTTGAAGTGTTGCTGA |
| 3160 | LINC00969-132 | CTAACTACAGGCTCAGAAGT |
| 3161 | LINC00969-132 | CCACTTTGAAAATCAATTTG |
| 3162 | LINC00969-132 | AAAATCAATTTGGGGAGATG |
| 3163 | LINC00969-132 | TTAGCCATTCATCAGTTGAA |
| 3164 | LINC00969-132 | CTACTTCTGAGCCTGTAGTT |
| 3165 | LINC00976-002 | AAGAAGCGCACAGCAAAACG |
| 3166 | LINC00976-002 | CTGAGTCCTCTCCAGCACAG |
| 3167 | LINC00976-002 | ACAGTTCCACTGTGCTGGAG |
| 3168 | LINC00976-002 | ACACGTCTTTCTTCACATGT |
| 3169 | LINC00976-002 | ATCTGTTCAAAGTGCTTATG |
| 3170 | LINC00976-002 | GAACTTCAGTTAAGGAGTTG |
| 3171 | LINC00976-002 | GACTCACAGTTCCACTGTGC |
| 3172 | LINC00996-001 | AGTGACTACCCCGACCTACC |
| 3173 | LINC00996-001 | CACAACAGCCTGGTAGGTCG |
| 3174 | LINC00996-001 | TCACTTCCATTTGGGAGATG |
| 3175 | LINC00996-001 | TTGGGAGATGAGGAAGTCGA |
| 3176 | LINC00996-001 | AGCTCACAGAATCAGGAAGA |
| 3177 | LINC00996-001 | GGTTCCCACAACAGCCTGGT |
| 3178 | LINC00996-001 | AGGTCTAAGCTCACAGAATC |
| 3179 | LINC00996-002 | AGCTTCCTACCGTTCCACGA |
| 3180 | LINC00996-002 | GTTGTGGGAACCGAACGATG |
| 3181 | LINC00996-002 | AGTGGCCCTCGTGGAACGGT |

| | | |
|---|---|---|
| 3182 | LINC00996-002 | GTTCCACGAGGGCCACTGAA |
| 3183 | LINC00996-002 | AAAAGGTGACAGTCTCCTTG |
| 3184 | LINC00996-002 | CAGCTTCCTACCGTTCCACG |
| 3185 | LINC00996-002 | GTTCGGTTCCCACAACAGCC |
| 3186 | LINC00998-001 | CAGCCATTACAGGAATGGCG |
| 3187 | LINC00998-001 | CGGGGCATCATAGGAGTCGG |
| 3188 | LINC00998-001 | CGCCGGGGCATCATAGGAGT |
| 3189 | LINC00998-001 | CCATTACAGGAATGGCGAGG |
| 3190 | LINC00998-001 | GGCATCATAGGAGTCGGCGG |
| 3191 | LINC00998-001 | TACAGGAATGGCGAGGAGGG |
| 3192 | LINC00998-001 | CATTACAGGAATGGCGAGGA |
| 3193 | LINC00998-002 | GAGGCCATATGGACGGTAGG |
| 3194 | LINC00998-002 | GTGGAGGCCATATGGACGGT |
| 3195 | LINC00998-002 | GAGCGTGGAGGCCATATGGA |
| 3196 | LINC00998-002 | GGAGGCCATATGGACGGTAG |
| 3197 | LINC00998-002 | TGGAGGCCATATGGACGGTA |
| 3198 | LINC00998-002 | AAAGAATTCGTACTTTGGCC |
| 3199 | LINC00998-002 | CTATCCCCCTACCGTCCATA |
| 3200 | LINC01001-001 | GCAACTAGACTGTCCACCTG |
| 3201 | LINC01001-001 | CAGGTGGACAGTCTAGTTGC |
| 3202 | LINC01001-001 | GACTGTCCACCTGGGGTGAT |
| 3203 | LINC01001-001 | TTACTGTCTCCCATCACCCC |
| 3204 | LINC01001-001 | CTGTCTCCCATCACCCCAGG |
| 3205 | LINC01001-001 | TGCAACTAGACTGTCCACCT |
| 3206 | LINC01001-002 | AGCATATGTCCTGCGAGCTT |
| 3207 | LINC01001-002 | GCTGCTCAAGTGCATTTGAA |
| 3208 | LINC01001-002 | CAGCAGCATTAGATTGTCAT |
| 3209 | LINC01001-002 | GACAGGACATGGAGCTCAGG |
| 3210 | LINC01001-002 | CAGGTGGTCATGCAAGCGAT |
| 3211 | LINC01001-002 | GCATATGTCCTGCGAGCTTA |
| 3212 | LINC01001-002 | CCACAGGGCCACAGAGCTGG |
| 3213 | LINC01002-010 | CCAGTCTTCACAGTGGTCAA |
| 3214 | LINC01002-010 | TCCCACACCAGTCTTCACAG |
| 3215 | LINC01002-010 | AGGCGGGAATGTGAGCAAAG |
| 3216 | LINC01002-010 | CTTGCGGCTCCATGGCTCAG |
| 3217 | LINC01002-010 | AGGTGAGCGGTGTCGAGTGA |
| 3218 | LINC01002-010 | GGATGCTGGTGATTTCCAGT |
| 3219 | LINC01002-010 | TGGTAGCTTGAGTCACTGTG |
| 3220 | LINC01002-012 | TCACTTGATATCAGGAGTCG |
| 3221 | LINC01002-012 | CTTGATATCAGGAGTCGAGG |
| 3222 | LINC01002-012 | ATTACTCATAACAGAGATAA |
| 3223 | LINC01002-012 | TTGATATCAGGAGTCGAGGC |
| 3224 | LINC01002-012 | TAACGGTCCAATAAGAATCC |

| | | |
|---|---|---|
| 3225 | LINC01002-012 | TCTAAAGTAACATCCTCTCA |
| 3226 | LINC01002-012 | TTAATTAAATACAACCCTAG |
| 3227 | LINC01002-014 | GAACTGTACAGGCCACCAGG |
| 3228 | LINC01002-014 | CCCAGCTCTTGCCCCCACGG |
| 3229 | LINC01002-014 | TGTACAGTTCTAGCACTGGT |
| 3230 | LINC01002-014 | AACCAGTGCTAGAACTGTAC |
| 3231 | LINC01002-014 | CTTTCTCCAGCCAAGCTCTG |
| 3232 | LINC01002-014 | ACAGGCCACCAGGAGGCAGG |
| 3233 | LINC01002-014 | GGGTGCCAGAGGCATGAGTT |
| 3234 | LINC01002-015 | GGCCTGCGGCAGTCGCCGGG |
| 3235 | LINC01002-015 | CGGGAGGCCCAACCTTGGCG |
| 3236 | LINC01002-015 | AGTCCAAAGAGGTTGTTGAG |
| 3237 | LINC01002-015 | TTCAGGACAACTTGGGCCTG |
| 3238 | LINC01002-015 | AGTGCCTACCCATCTCCTGG |
| 3239 | LINC01002-015 | GAGGCCCAACCTTGGCGTGG |
| 3240 | LINC01002-015 | GTTGTTGAGAGGCAGGAATC |
| 3241 | LINC01002-019 | TCTTTAAAACTGGAAGAGGG |
| 3242 | LINC01002-019 | GGGACCTCAGTATGTTGCCA |
| 3243 | LINC01002-019 | TCAGCCATGGCAACATACTG |
| 3244 | LINC01002-019 | CATTCTTTAAAACTGGAAGA |
| 3245 | LINC01002-019 | AGGCTATAATGAGCCATGAT |
| 3246 | LINC01002-019 | AGAAGGTTAAGAACCAGAGA |
| 3247 | LINC01002-019 | AACTGGAAGAGGGAGGCAGA |
| 3248 | LINC01003-001 | GCGGGGAATTAGGCGGTGCG |
| 3249 | LINC01003-001 | GGCGGTGCGAGGCCAATGGG |
| 3250 | LINC01003-001 | GCGGTGCGAGGCCAATGGGA |
| 3251 | LINC01003-001 | GGCACCCCGCCTCAGCCCAG |
| 3252 | LINC01003-001 | GGCGAGGGCGGGGAATTAGG |
| 3253 | LINC01003-001 | GAGGGCGGGCCGGGTCCTCT |
| 3254 | LINC01003-001 | TTAGGCGGTGCGAGGCCAAT |
| 3255 | LINC01021-002 | GACCTGGGAATCAATGTGTG |
| 3256 | LINC01021-002 | GTGAGGTGGTGCTATGTAGC |
| 3257 | LINC01021-002 | AAACTATTTGCAAAGCAAGA |
| 3258 | LINC01021-002 | CTGGGAATCAATGTGTGAGG |
| 3259 | LINC01021-002 | ATTTCTCAAGCCCCGTGTTT |
| 3260 | LINC01021-002 | TCAGTAGAGCAGGGCTCTAT |
| 3261 | LINC01021-002 | ATTGAAGGGTCAGTAGAGCA |
| 3262 | LINC01023-001 | GAGGACAGTACTCGGGCATG |
| 3263 | LINC01023-001 | GCCCGAGTACTGTCCTCAGG |
| 3264 | LINC01023-001 | TGCAGACGCCGTTCCGCCTG |
| 3265 | LINC01023-001 | CTGAGGACAGTACTCGGGCA |
| 3266 | LINC01023-001 | TGAGGACAGTACTCGGGCAT |
| 3267 | LINC01023-001 | CATGCCCGAGTACTGTCCTC |

| | | |
|---|---|---|
| 3268 | LINC01023-001 | TTCCGCCTGAGGACAGTACT |
| 3269 | LINC01055-001 | CTGAGTGGCCACCTAGACCA |
| 3270 | LINC01055-001 | CCTGAGTGGCCACCTAGACC |
| 3271 | LINC01055-001 | CCCCCTGCACTGATCCTGAG |
| 3272 | LINC01055-001 | TGTCCAGAGGCTCGACTATA |
| 3273 | LINC01055-001 | GAACCCTATAGTCGAGCCTC |
| 3274 | LINC01055-001 | GGCCACTCAGGATCAGTGCA |
| 3275 | LINC01055-001 | GTGTCCAGAGGCTCGACTAT |
| 3276 | LINC01067-001 | GCTCTTTAGAGCACACCGCT |
| 3277 | LINC01067-001 | CACAATATTTACCTTATCAG |
| 3278 | LINC01067-001 | TCAGTAGGAGATCTCACTGA |
| 3279 | LINC01067-001 | GGAGGTTTGCTTAATGAGAG |
| 3280 | LINC01067-001 | CAGCCTTGTGTTGTGAGGAG |
| 3281 | LINC01067-001 | CTCTTTAGAGCACACCGCTA |
| 3282 | LINC01067-001 | CCTCCCCTCCTCACAACACA |
| 3283 | LINC01089-003 | AAATGGCTTCCCTCATCTGT |
| 3284 | LINC01089-003 | AGGACTCACAGTTGGAAGAC |
| 3285 | LINC01089-003 | CTCTCCTTCGTGTCTGATGA |
| 3286 | LINC01089-003 | AACATTTGTCAACAGGCCCA |
| 3287 | LINC01089-003 | ATCAGCGCGCGGCTTCTATA |
| 3288 | LINC01089-003 | CAGGCCCAAGGACTCACAGT |
| 3289 | LINC01089-003 | TCTTCCAACTGTGAGTCCTT |
| 3290 | LINC01089-006 | CTTCATCAGACACGAAGGAG |
| 3291 | LINC01089-006 | ACATCCTTCATCAGACACGA |
| 3292 | LINC01089-006 | AAGGAGAGGCCAACAGATGA |
| 3293 | LINC01089-006 | CTAGTTATACAAGACAAAGA |
| 3294 | LINC01089-006 | GAACTGGATGGTAGAGTCAA |
| 3295 | LINC01089-006 | AATGGAATTAAATGGCCAAG |
| 3296 | LINC01089-006 | ATGGAATTAAATGGCCAAGT |
| 3297 | LINC01116-002 | CCTCTTTACAAACCTGAGCG |
| 3298 | LINC01116-002 | GCTCGTCTTCAGCAAAGCGA |
| 3299 | LINC01116-002 | CCCCTCTTTACAAACCTGAG |
| 3300 | LINC01116-002 | TGCTCGTCTTCAGCAAAGCG |
| 3301 | LINC01116-002 | AGGGGAGGCAAGACTTTGGT |
| 3302 | LINC01116-002 | CCCTCTTTACAAACCTGAGC |
| 3303 | LINC01116-002 | GTCTTCAGCAAAGCGAGGGC |
| 3304 | LINC01128-001 | AACCGGCGGAGACACTGGCA |
| 3305 | LINC01128-001 | AGACACGCGCTTTCAACCGG |
| 3306 | LINC01128-001 | AGGGCCAGAAACGCGCGCAG |
| 3307 | LINC01128-001 | GGCCAGAAACGCGCGCAGCG |
| 3308 | LINC01128-001 | GCCAGAAACGCGCGCAGCGG |
| 3309 | LINC01128-001 | CTTTCAACCGGCGGAGACAC |
| 3310 | LINC01128-001 | TTGAAAGCGCGTGTCTGCGT |

| | | |
|---|---|---|
| 3311 | LINC01128-004 | GGAGTGCGTTCGGTGCGCCG |
| 3312 | LINC01128-004 | GAGTGCGTTCGGTGCGCCGT |
| 3313 | LINC01128-004 | GCGGGAGCGAGAGTCGGGTG |
| 3314 | LINC01128-004 | TGGCGGGAGCGAGAGTCGGG |
| 3315 | LINC01128-004 | TTACCTAACAGGAAGTTGGG |
| 3316 | LINC01128-004 | CCGAAGAGGCTGCTACCCAG |
| 3317 | LINC01128-004 | GTGTCTGCGTCGGGTTCTGT |
| 3318 | LINC01128-016 | TGTTAGGTAAGAGGCGCGTG |
| 3319 | LINC01128-016 | AGCGCGCGTCACGCCGTGTG |
| 3320 | LINC01128-016 | CACGCCGTGTGGGGCCGAAG |
| 3321 | LINC01128-016 | CCAACTTCCTGTTAGGTAAG |
| 3322 | LINC01128-016 | GCAGCCTCTTCGGCCCCACA |
| 3323 | LINC01128-016 | CGAGCGCGCGTCACGCCGTG |
| 3324 | LINC01128-016 | GGCGGGAGCGAGAGTCGGGT |
| 3325 | LINC01138-002 | TCAAGCGCGCACACCCCTCA |
| 3326 | LINC01138-002 | CCGGAGAGCGATCACCGGCT |
| 3327 | LINC01138-002 | TGGGCGGTCACATCTGGAAA |
| 3328 | LINC01138-002 | TGCGCGCTTGAGCCGGAGCG |
| 3329 | LINC01138-002 | AGGGGTGTGCGCGCTTGAGC |
| 3330 | LINC01138-002 | TGTGCGCGCTTGAGCCGGAG |
| 3331 | LINC01138-002 | CCCAGCCGGTGATCGCTCTC |
| 3332 | LINC01138-003 | AGCTGTCGCCAAATGAACCC |
| 3333 | LINC01138-003 | GCGGTCGGAGAACTGAGAAG |
| 3334 | LINC01138-003 | CGGTCGGAGAACTGAGAAGG |
| 3335 | LINC01138-003 | TTCTCCGACCGCGACGTGTG |
| 3336 | LINC01138-003 | CCAGCCACACACGTCGCGGT |
| 3337 | LINC01138-003 | GACAGCTGCTGCAACGAACG |
| 3338 | LINC01138-003 | GTTTGCACGTCCGCATTCCG |
| 3339 | LINC01150-001 | CTTGATGAGATGCTAAACCA |
| 3340 | LINC01150-001 | CTGCAATTGTTACAGGAGAG |
| 3341 | LINC01150-001 | CCTGTATGGCCTAAAAAGCA |
| 3342 | LINC01150-001 | CTGTATGGCCTAAAAAGCAG |
| 3343 | LINC01150-001 | GGGCATGAGTAATCCACCCC |
| 3344 | LINC01150-001 | ATGAGATGCTAAACCAGGGG |
| 3345 | LINC01150-001 | CACTGCAATTGTTACAGGAG |
| 3346 | LINC01160-002 | CGGACGGGGTGGTTGCTGGG |
| 3347 | LINC01160-002 | CACCTCCCTCCCGGACGGGG |
| 3348 | LINC01160-002 | CAGACGGGGTGACTGCCGGG |
| 3349 | LINC01160-002 | CAGACGGGGCGGCTGCGAGG |
| 3350 | LINC01160-002 | CGGGGCGGCTGCGAGGCGGA |
| 3351 | LINC01160-002 | GCCCAGCAACCACCCCGTCC |
| 3352 | LINC01160-002 | CGCCCATCGTCTGAGATGTG |
| 3353 | LINC01160-003 | AAAAAATACGAAAAGCAGTG |

| | | |
|---|---|---|
| 3354 | LINC01160-003 | ATACGAAAAGCAGTGAGGCG |
| 3355 | LINC01160-003 | CTCACGTCCCAGGCGATGGG |
| 3356 | LINC01160-003 | CGCCCATCGCCTGGGACGTG |
| 3357 | LINC01160-003 | GACTGGGCAGCCAGGCAGAG |
| 3358 | LINC01160-003 | CTGGTTCACTCAGTGCTCAA |
| 3359 | LINC01160-003 | GTCTGCAATCCCCGCACCCC |
| 3360 | LINC01176-001 | CACAGTTCATTACAACTGCT |
| 3361 | LINC01176-001 | GCAAGCACACTGTCTTGCGT |
| 3362 | LINC01176-001 | AGCAACTGCTGTCTCAGCAC |
| 3363 | LINC01176-001 | CCAGCAGTTGTAATGAACTG |
| 3364 | LINC01176-001 | AAACAGCAGACAAGTGTTTG |
| 3365 | LINC01176-001 | TTGGTGCTCTTGGAGCAGGG |
| 3366 | LINC01176-001 | CATTACAACTGCTGGGAAGA |
| 3367 | LINC01184-001 | AGCATCGCCCGAAGGAATCG |
| 3368 | LINC01184-001 | CACTAGGCGCGTGGTACCCG |
| 3369 | LINC01184-001 | AAGCCCCCAGCATCGCCCGA |
| 3370 | LINC01184-001 | AAGGAATCGCGGCTGGCCTG |
| 3371 | LINC01184-001 | TCGCCCGAAGGAATCGCGGC |
| 3372 | LINC01184-001 | CAGGCCAGCCGCGATTCCTT |
| 3373 | LINC01184-001 | GGCGCGTGGTACCCGAGGCC |
| 3374 | LINC01184-003 | TGGGGAACGCACCTCGCCCG |
| 3375 | LINC01184-003 | GAGTTACGAGCGGAGATTCG |
| 3376 | LINC01184-003 | TTACGAGCGGAGATTCGCGG |
| 3377 | LINC01184-003 | TTCAGGCCCGGAGTTACGAG |
| 3378 | LINC01184-003 | GTCAGAGCCCAGAGCAAACC |
| 3379 | LINC01184-003 | GAATCTCCGCTCGTAACTCC |
| 3380 | LINC01184-003 | AACTCCGGGCCTGAACCTGG |
| 3381 | LINC01215-001 | AGATGAGGCACTAGACCACT |
| 3382 | LINC01215-001 | CAGTCTCCACCACTATTCTG |
| 3383 | LINC01215-001 | ATTCTGCCTCAGAATAGTGG |
| 3384 | LINC01215-001 | GAGATGAGGCACTAGACCAC |
| 3385 | LINC01215-001 | AGAAAACAAGAAAGTCGTGA |
| 3386 | LINC01215-001 | TCACAGCATTGTGAGCCAAT |
| 3387 | LINC01215-001 | TGCATTCTGCCTCAGAATAG |
| 3388 | LINC01232-002 | ACAAAACGCACAGTAAAGCT |
| 3389 | LINC01232-002 | AGCTAGGAAAGAATGAAGCA |
| 3390 | LINC01232-002 | GGCCAATGGGAAACATCTAG |
| 3391 | LINC01232-002 | GCACCAAGTAAGTGGCCAAT |
| 3392 | LINC01232-002 | AGCACCAAGTAAGTGGCCAA |
| 3393 | LINC01232-002 | GGGCTGAATCTTCATTTGCA |
| 3394 | LINC01232-002 | TGGGGTGAGCACCAAGTAAG |
| 3395 | LINC01237-001 | GAGGGCACCACCACCTGCAC |
| 3396 | LINC01237-001 | GGGCCCCAGTGAAGGCGCAG |

| | | |
|---|---|---|
| 3397 | LINC01237-001 | TGCCCTCTGCGCCTTCACTG |
| 3398 | LINC01237-001 | CCAAGTAACTGAATCCAGCC |
| 3399 | LINC01237-001 | GGCCCCAGTGAAGGCGCAGA |
| 3400 | LINC01237-001 | CTGGATTCAGTTACTTGGCT |
| 3401 | LINC01237-001 | CTGAATCCAGCCAGGAGCAA |
| 3402 | LINC01239-001 | TATGTTCCACTCTTAAGCCA |
| 3403 | LINC01239-001 | CTTTGACCCTGGCTTAAGAG |
| 3404 | LINC01239-001 | ACCGAGAAAACAGACTGTGA |
| 3405 | LINC01239-001 | CTATGTTCCACTCTTAAGCC |
| 3406 | LINC01239-001 | GCTGAGACACACTAGAGAAG |
| 3407 | LINC01239-001 | CTTCATCCATTTATCTTGGA |
| 3408 | LINC01239-001 | GGTCAAAGTAGAATTCTCTT |
| 3409 | LINC01273-002 | ACTGCCACCTCTAAGCTGGA |
| 3410 | LINC01273-002 | GGTAAGGTTGCTGTAGCCCA |
| 3411 | LINC01273-002 | CCCGCTTTGCGGAGTGACGC |
| 3412 | LINC01273-002 | CAGCCCCTCCAGCTTAGAGG |
| 3413 | LINC01273-002 | AGCCACTGCCACCTCTAAGC |
| 3414 | LINC01273-002 | CTGCCACCTCTAAGCTGGAG |
| 3415 | LINC01273-002 | TGCGGAGTGACGCAGGCCCT |
| 3416 | LINC01353-002 | TGTCACCACACACAGGCGCA |
| 3417 | LINC01353-002 | CCAGTCTCCATAGGGTGACA |
| 3418 | LINC01353-002 | GCGCCTGTGTGTGGTGACAG |
| 3419 | LINC01353-002 | TATTACCTTGCGCCTGTGTG |
| 3420 | LINC01353-002 | GTCAGCACCATGTCACCCTA |
| 3421 | LINC01353-002 | TGCCCGCTGTCACCACACAC |
| 3422 | LINC01353-002 | AAGGTAATACCAGTCTCCAT |
| 3423 | LINC01358-001 | GGCTCCTGCAACCTTAATGG |
| 3424 | LINC01358-001 | TGTAACACCAGAGCTGTAGA |
| 3425 | LINC01358-001 | TGTTACACTGAAAAACTAGA |
| 3426 | LINC01358-001 | GGAGCCCAGCATGTCCCCAT |
| 3427 | LINC01358-001 | GTGTAACACCAGAGCTGTAG |
| 3428 | LINC01358-001 | GAGGCCTATGGGGACATGCT |
| 3429 | LINC01358-001 | TTTACCACCATTAAGGTTGC |
| 3430 | LINC01358-007 | AGAACATGTTGGCCGAAGCA |
| 3431 | LINC01358-007 | TCCTACCTGAAAATTTCAAG |
| 3432 | LINC01358-007 | CCTACCTGAAAATTTCAAGT |
| 3433 | LINC01358-007 | ATGTGTTTGCCAGAACAATG |
| 3434 | LINC01358-007 | GGAAGACTGGGAACAAACAC |
| 3435 | LINC01358-007 | AATGCGGGAGTTGCAGAGAG |
| 3436 | LINC01358-007 | TGTGTTTGCCAGAACAATGC |
| 3437 | LINC01374-001 | AAGACAAGGCCTCAGACACG |
| 3438 | LINC01374-001 | GGCTCTGCTGACAGTAGACA |
| 3439 | LINC01374-001 | CCCTTCCGGCCGCTAAGACA |

| | | |
|---|---|---|
| 3440 | LINC01374-001 | CTAAGACAAGGCCTCAGACA |
| 3441 | LINC01374-001 | TGTCTGAGGCCTTGTCTTAG |
| 3442 | LINC01374-001 | CAGAGCCAAGCTGAGAAGGA |
| 3443 | LINC01374-001 | CCTTGTCTTAGCGGCCGGAA |
| 3444 | LINC01410-001 | CAGTAGAGCGCGAGGTGGAG |
| 3445 | LINC01410-001 | AGCACACTCTACAAATCTCG |
| 3446 | LINC01410-001 | CTCCACCTCGCGCTCTACTG |
| 3447 | LINC01410-001 | ACAGTAGAGCGCGAGGTGGA |
| 3448 | LINC01410-001 | GCAGGCCGGGGTGTGGTGGA |
| 3449 | LINC01410-001 | TTCAGCAGGCCGGGGTGTGG |
| 3450 | LINC01410-001 | CCCCCACCCAGGAAAACCAA |
| 3451 | LINC01410-002 | AACCCGCGCCTGCACCCTCG |
| 3452 | LINC01410-002 | CGGGACGGGACCGACCCCGA |
| 3453 | LINC01410-002 | GCGGGACGGGACCGACCCCG |
| 3454 | LINC01410-002 | CGCGCCTGCACCCTCGGGGT |
| 3455 | LINC01410-002 | GGGACCGACCCCGAGGGTGC |
| 3456 | LINC01410-002 | GGGTCGGTCCCGTCCCGCCC |
| 3457 | LINC01410-002 | GCAGCCCCACCCAACCCGGG |
| 3458 | LINC01410-003 | AAGAACCCTCCACCACACCC |
| 3459 | LINC01410-003 | GAGCACCACAGTAGAGCGCG |
| 3460 | LINC01410-003 | CACCACAGTAGAGCGCGAGG |
| 3461 | LINC01410-003 | CAAAGCAACCAACCAATCCA |
| 3462 | LINC01410-003 | AAGGAAAAAAACAAGTCTCA |
| 3463 | LINC01410-003 | CAAGGAAAAAAACAAGTCTC |
| 3464 | LINC01410-003 | CCACACCCCGGCCTGCTGAA |
| 3465 | LINC01431-003 | CACCGCCTGCGTCTTGGTCT |
| 3466 | LINC01431-003 | CGGCGCCACCGCCTGCGTCT |
| 3467 | LINC01431-003 | CCGGTTTCTCACTCAACCGC |
| 3468 | LINC01431-003 | ATCGGGGATGCTGGAACCTG |
| 3469 | LINC01431-003 | CCTGCGGTTGAGTGAGAAAC |
| 3470 | LINC01431-003 | GTCGGCCGTATCGGGGATGC |
| 3471 | LINC01431-003 | AAACCACGCCCAGACCCCGC |
| 3472 | LINC01480-001 | TTAAGCGACTCTTAAATAGG |
| 3473 | LINC01480-001 | AAGCAATACCTGTCTCCCGT |
| 3474 | LINC01480-001 | TCTTAAATAGGTGGCTCACC |
| 3475 | LINC01480-001 | TCAAATAGTTCAGATTGCAT |
| 3476 | LINC01480-001 | AGAAAAAATAATGATATGAT |
| 3477 | LINC01480-001 | TAGGTGGCTCACCAGGAGAA |
| 3478 | LINC01480-001 | CGGAGCTATTTGATGAAAGA |
| 3479 | LINC01480-002 | AAAGGGCTAACTCACCACAA |
| 3480 | LINC01480-002 | TTGGCCAGAACCAATGTTAG |
| 3481 | LINC01480-002 | AAGCAGTACCTGTGTCCTGT |
| 3482 | LINC01480-002 | AGCCCTTTAGAAGTCACTTG |

| | | |
|---|---|---|
| 3483 | LINC01480-002 | TATGTTAATTCCACTAACAT |
| 3484 | LINC01480-002 | TAAAGGGCTAACTCACCACA |
| 3485 | LINC01480-002 | CTTTAGAAGTCACTTGGGGT |
| 3486 | LINC01480-004 | GACTAATGTCCAGACTCCAA |
| 3487 | LINC01480-004 | AGGCAGAAGAAATATCAGCA |
| 3488 | LINC01480-004 | TTCTGGCCAATGAGTCAGGA |
| 3489 | LINC01480-004 | GGAAACAGTAGTCACAAACT |
| 3490 | LINC01480-004 | GACTACTGTTTCCTATACTG |
| 3491 | LINC01480-004 | GGGCATTAGTCATGACTACT |
| 3492 | LINC01480-004 | ACTACTGTTTCCTATACTGT |
| 3493 | LINC01481-001 | CTGGAATAAACCTCATGAGT |
| 3494 | LINC01481-001 | CAGGTGCCAGAAATCCAACA |
| 3495 | LINC01481-001 | CATAATGCCCTATAAACAGA |
| 3496 | LINC01481-001 | ACAGATGGATGTGAATGCTC |
| 3497 | LINC01481-001 | TTAACATTCTCCATCAGCTG |
| 3498 | LINC01481-001 | AGTTATCTCAAATTGGTAAG |
| 3499 | LINC01481-001 | ACTGGAATAAACCTCATGAG |
| 3500 | LINC01484-003 | CAGCTGATCTTCACAGATGT |
| 3501 | LINC01484-003 | GGCAGAGGAAGCAATCGGAT |
| 3502 | LINC01484-003 | TTCTTCAGGAAGGAAATGGG |
| 3503 | LINC01484-003 | ACAGCTGATCTTCACAGATG |
| 3504 | LINC01484-003 | ATTTGCTTCAGGTGAGCAGA |
| 3505 | LINC01484-003 | CATCTGTGAAGATCAGCTGT |
| 3506 | LINC01484-003 | CATTTGCTTCAGGTGAGCAG |
| 3507 | LINC01485-003 | CATGCCAATGTGAAACTCCG |
| 3508 | LINC01485-003 | TGGCCTTCCAGGTCATTAGG |
| 3509 | LINC01485-003 | GCAGGAAAACATGAAGAACG |
| 3510 | LINC01485-003 | CGGAGTTTCACATTGGCATG |
| 3511 | LINC01485-003 | TTTGGTTTGTGACCCTGCCA |
| 3512 | LINC01485-003 | GACTCCTCGGAGTTTCACAT |
| 3513 | LINC01485-003 | ATCAGAATGGCTAGACTCCT |
| 3514 | LINC01485-005 | GGTGAAGCATGCTTTCAAGT |
| 3515 | LINC01485-005 | GTGAAGCATGCTTTCAAGTG |
| 3516 | LINC01485-005 | ACTTGAAAGCATGCTTCACC |
| 3517 | LINC01485-005 | CGGTGAAGCATGCTTTCAAG |
| 3518 | LINC01485-005 | GCATGCTTCACCGGGCGCGG |
| 3519 | LINC01485-005 | CACTTGAAAGCATGCTTCAC |
| 3520 | LINC01485-005 | CAGGTGTGAGCCACCGCGCC |
| 3521 | LINC01534-001 | AATTCTCATAGTGCGTGGCT |
| 3522 | LINC01534-001 | ACGCACTATGAGAATTTGGA |
| 3523 | LINC01534-001 | TTCCAAATTCTCATAGTGCG |
| 3524 | LINC01534-001 | GTGCGTGGCTGGGATGAGGG |
| 3525 | LINC01534-001 | GCACACCCTGTCTGCGTCCC |

| | | |
|---|---|---|
| 3526 | LINC01534-001 | TGAGGGCGGCGCTCACGGTG |
| 3527 | LINC01534-001 | TGAGGAAGCCCCGAGCTCCC |
| 3528 | LINC01549-201 | CAAGCAACATGGCTCCAGAG |
| 3529 | LINC01549-201 | AGAGCTCAATGCTTGAGAAA |
| 3530 | LINC01549-201 | AAGGAATTCTGTCCTTTGAA |
| 3531 | LINC01549-201 | TGGAGCCATGTTGCTTGGTC |
| 3532 | LINC01549-201 | GATAAAAATAGGAAGACGAT |
| 3533 | LINC01549-201 | AACAGGTATAACATGAAAGA |
| 3534 | LINC01549-201 | CACTCTGGAGCCATGTTGCT |
| 3535 | LINC01562-001 | GTATACCAGACCCCACGCTA |
| 3536 | LINC01562-001 | GGCTCTATCAGGAACAAGCC |
| 3537 | LINC01562-001 | GCAATTAGTTCATTTGAGTG |
| 3538 | LINC01562-001 | GGGAAAAGGATGATCATACC |
| 3539 | LINC01562-001 | CAGACCCCACGCTAAGGACT |
| 3540 | LINC01562-001 | CCAGACCCCACGCTAAGGAC |
| 3541 | LINC01562-001 | CCAGTCCTTAGCGTGGGGTC |
| 3542 | LINC01569-001 | AGCGCGAGCTCCATGCCCGG |
| 3543 | LINC01569-001 | CCTTCCCGGAGAGCAGGTGA |
| 3544 | LINC01569-001 | TCCCGGAGAGCAGGTGACGG |
| 3545 | LINC01569-001 | GTCAGCGCGAGCTCCATGCC |
| 3546 | LINC01569-001 | CCGTCACCTGCTCTCCGGGA |
| 3547 | LINC01569-001 | GCCGCCGTCACCTGCTCTCC |
| 3548 | LINC01569-001 | CTGCTCTCCGGGAAGGCCCT |
| 3549 | LINC01578-001 | ACAACATGGCCTCTTCGCTG |
| 3550 | LINC01578-001 | AGCGAAGAGGCCATGTTGTG |
| 3551 | LINC01578-001 | ACATGGCCTCTTCGCTGCGG |
| 3552 | LINC01578-001 | AGGCCATGTTGTGTGGTGTG |
| 3553 | LINC01578-001 | GGCCATGTTGTGTGGTGTGT |
| 3554 | LINC01578-001 | GCCCCACACACCACACAACA |
| 3555 | LINC01578-001 | GCCATGTTGTGTGGTGTGTG |
| 3556 | LINC01578-003 | GGAGAGAGACACCACACACA |
| 3557 | LINC01578-003 | GCCATCTTGTGCCGTGTGTG |
| 3558 | LINC01578-003 | AGTAAAGCCTAGAGAGGAGA |
| 3559 | LINC01578-003 | GGCGGGGAGCTACGCCCGGA |
| 3560 | LINC01578-003 | CCAGCAGGCCCGCGGGAGTG |
| 3561 | LINC01578-003 | CGGACGGCCAGCAGGCCCGC |
| 3562 | LINC01578-003 | CCGCGGCAGCCCCACTCCCG |
| 3563 | LINC01578-004 | GCGGAACGCCTGACAGCTCG |
| 3564 | LINC01578-004 | CAGCAGCACCCCGAGCTGTC |
| 3565 | LINC01578-004 | GGCGTTCCGCCGCGGCCGCG |
| 3566 | LINC01578-004 | CGTAGCTCCCCGCCGGCCGG |
| 3567 | LINC01578-004 | CCGCGGGCCTGCTGGCCGTC |
| 3568 | LINC01578-004 | GCGAGGCCCGCCGGCCGGCG |

| | | |
|---|---|---|
| 3569 | LINC01578-004 | CCGCCGGCCGGCGGGCCTCG |
| 3570 | LINC01578-006 | CGCTGCGGTTGCCGCCTCGG |
| 3571 | LINC01578-006 | CCCAAACCCGAGTCCCGGGC |
| 3572 | LINC01578-006 | GGCTGTGACGCCGACGCGCG |
| 3573 | LINC01578-006 | GAGGCTCCCCCGGGCCCCGG |
| 3574 | LINC01578-006 | GCCCGGGGGAGCCTCCGAGG |
| 3575 | LINC01578-006 | GCGAGGCGGGCCGCGCGCGT |
| 3576 | LINC01578-006 | CGCGCCGGGTCCAGCCGCCG |
| 3577 | LINC01578-007 | AGCCATGCTGTGTAATCACA |
| 3578 | LINC01578-007 | AGAATTATCCCAAAATGCTA |
| 3579 | LINC01578-007 | GGTTCTTAAGCAGAAAACAG |
| 3580 | LINC01578-007 | ATTCAATCACCAAGTTGATC |
| 3581 | LINC01578-007 | ACCTAATTGATTGGCACATG |
| 3582 | LINC01578-007 | AAGAATTATCCCAAAATGCT |
| 3583 | LINC01578-007 | ATTATATAAGATAAATATAG |
| 3584 | LINC01578-008 | CGACGCCGAAAGCGCCGCTG |
| 3585 | LINC01578-008 | CAGCGGCGCTTTCGGCGTCG |
| 3586 | LINC01578-008 | AGCGGCGCTTTCGGCGTCGT |
| 3587 | LINC01578-008 | CTCCGAGGCGGCAACCGCAG |
| 3588 | LINC01578-008 | GTCCAGCCGCCGGGGCCCGG |
| 3589 | LINC01578-008 | TGCCGCCTCGGAGGCTCCCC |
| 3590 | LINC01578-008 | TCGGAGGCTCCCCCGGGCCC |
| 3591 | LINC01609-002 | CTGGATCACCTAAGGCAGAG |
| 3592 | LINC01609-002 | TGGATCACCTAAGGCAGAGA |
| 3593 | LINC01609-002 | GAAGACTGTATTTCCTCCTC |
| 3594 | LINC01609-002 | ACAGTTCAGCTGCTTGCACT |
| 3595 | LINC01609-002 | GGTTACAGCTGGATCACCTA |
| 3596 | LINC01609-002 | TCACCTAAGGCAGAGAGGGA |
| 3597 | LINC01609-002 | GAGAGTATCATGGTTACAGC |
| 3598 | LINC01619-006 | TTCAATTAGAAATTTCCCCG |
| 3599 | LINC01619-006 | TTTATATCTTCACTTCAACA |
| 3600 | LINC01619-006 | CAAAACATGGACACATTCAC |
| 3601 | LINC01619-006 | GACCCATAGTCTTAAAGAAC |
| 3602 | LINC01619-006 | TGTTCAATTAGAAATTTCCC |
| 3603 | LINC01619-006 | AAAAGCAACCAATCTCTCTG |
| 3604 | LINC01619-006 | ACTATGGGTCACCTGCCCCG |
| 3605 | LLNLR-246C6.1-001 | GGAAATCTACAGATCAACTG |
| 3606 | LLNLR-246C6.1-001 | AAAATACTTAGGAACAAAAG |
| 3607 | LLNLR-246C6.1-001 | AGGTCCAAGCCTGGGCACGG |
| 3608 | LLNLR-246C6.1-001 | GAACAAAAGAGGTCCAAGCC |
| 3609 | LLNLR-246C6.1-001 | AAGAGGTCCAAGCCTGGGCA |
| 3610 | LLNLR-246C6.1-001 | AACAAAAGAGGTCCAAGCCT |
| 3611 | LLNLR-246C6.1-001 | TGTAATTCCAGGACTTTGGG |

| | | |
|---|---|---|
| 3612 | MALAT1-001 | GGAAGCCTCAGCTCGCCTGA |
| 3613 | MALAT1-001 | GGCAGGAGAGGCCAGTTGCG |
| 3614 | MALAT1-001 | AGGGCAGGAGAGGCCAGTTG |
| 3615 | MALAT1-001 | ACCTGCCTTCAGGCGAGCTG |
| 3616 | MALAT1-001 | GAGACTTCTGTAAAGGACTG |
| 3617 | MALAT1-001 | GCAGCCCGAGACTTCTGTAA |
| 3618 | MALAT1-001 | CGAGACTTCTGTAAAGGACT |
| 3619 | MALAT1-002 | GCTTCCCAATTCAAACTGGG |
| 3620 | MALAT1-002 | TGTTACGGTTGGGATTGGTG |
| 3621 | MALAT1-002 | TGGCAGTGGCCTGTTACGGT |
| 3622 | MALAT1-002 | TGGCCTGTTACGGTTGGGAT |
| 3623 | MALAT1-002 | GGCAGTGGCCTGTTACGGTT |
| 3624 | MALAT1-002 | CCACCAATCCCAACCGTAAC |
| 3625 | MALAT1-002 | GGCAAATATTGGCAATTAGT |
| 3626 | MALAT1-005 | GAGCTGTGCGGTAGGCATTG |
| 3627 | MALAT1-005 | TTAGAAGAGTAGCATGAGGA |
| 3628 | MALAT1-005 | CATTGAGGCAGCCAGCGCAG |
| 3629 | MALAT1-005 | CCTCGCCCGAGCTGTGCGGT |
| 3630 | MALAT1-005 | AAGAGAAAAAAACTTATCTG |
| 3631 | MALAT1-005 | CGCAGGGGCTTCTGCTGAGG |
| 3632 | MALAT1-005 | GAGGGGGCAGGCGGAGCTTG |
| 3633 | MALAT1-008 | AATGTGAAGGACTTTCGTAA |
| 3634 | MALAT1-008 | CATTAATTGACAGCTGACCC |
| 3635 | MALAT1-008 | GTCCTAGCTTCATCAAACAC |
| 3636 | MALAT1-008 | TCCACTTCTGTGTAGCACCT |
| 3637 | MALAT1-008 | AAGAGTTAGAAGAATTTGGA |
| 3638 | MALAT1-008 | GAAGTGGATTCAGTGAATCT |
| 3639 | MALAT1-008 | ACCCAGGTGCTACACAGAAG |
| 3640 | MAPKAPK5-AS1-001 | GCGCGATCCACGGAATCCGG |
| 3641 | MAPKAPK5-AS1-001 | AGAGCCGAGAAGAACGCCAC |
| 3642 | MAPKAPK5-AS1-001 | GAGAGGTTCCATCGACTACC |
| 3643 | MAPKAPK5-AS1-001 | GGCCGAAACCCGGTAGTCGA |
| 3644 | MAPKAPK5-AS1-001 | GGCTCTAGCCTCAAAAAGCA |
| 3645 | MAPKAPK5-AS1-001 | TCTTCCCCCGCGTTGTCCTG |
| 3646 | MAPKAPK5-AS1-001 | TGAGAGGTTCCATCGACTAC |
| 3647 | MAPKAPK5-AS1-005 | GTGCCAAATCTACTAGACAT |
| 3648 | MAPKAPK5-AS1-005 | GTAGGAACAACACATGGTAC |
| 3649 | MAPKAPK5-AS1-005 | GCCCTTTGATAGGTTATCGG |
| 3650 | MAPKAPK5-AS1-005 | CAACACATGGTACCGGCACA |
| 3651 | MAPKAPK5-AS1-005 | CTAGACATTGGAGAAACAGT |
| 3652 | MAPKAPK5-AS1-005 | AGAGTAGGGAGCACAAACCT |
| 3653 | MAPKAPK5-AS1-005 | GGTAGCCGTTTGGCGTCGTG |
| 3654 | MAPKAPK5-AS1-010 | GAGTAGGGAGCACAAACCTG |

| | | |
|---|---|---|
| 3655 | MAPKAPK5-AS1-010 | CTCCACCGATAACCTATCAA |
| 3656 | MAPKAPK5-AS1-010 | GAAACAGTAGGAACAACACA |
| 3657 | MAPKAPK5-AS1-010 | GTTTGTGCACAGCCGAGAAG |
| 3658 | MAPKAPK5-AS1-010 | TTCCTGAAAGATCCATGTGC |
| 3659 | MAPKAPK5-AS1-010 | TCAAGCTAGAATGGGACCCC |
| 3660 | MAPKAPK5-AS1-010 | GGCTACCTTGTAAAGACGAA |
| 3661 | MCM10 | GCGTGCGTGTGACCCTCGGG |
| 3662 | MCM10 | GAGGGTCACACGCACGCCCG |
| 3663 | MCM10 | CCGAGGGTCACACGCACGCC |
| 3664 | MCM10 | CGGGCGTGCGTGTGACCCTC |
| 3665 | MCM10 | CCGGGCGTGCGTGTGACCCT |
| 3666 | MCM10 | CCGCGCTCCTCACCCGATGC |
| 3667 | MCM10 | CGAGGGTCACACGCACGCCC |
| 3668 | MCM2 | TTGTTGCTGTAGTGGCGGAG |
| 3669 | MCM2 | GGATCGTGGTACTGCTATGG |
| 3670 | MCM2 | CCGCCACTACAGCAACAACC |
| 3671 | MCM2 | CCTGGTTGTTGCTGTAGTGG |
| 3672 | MCM2 | AGAGGATCGTGGTACTGCTA |
| 3673 | MCM2 | TGTAGTGGCGGAGAGGATCG |
| 3674 | MCM2 | ACCAGGTTTCGCGCGAAAAG |
| 3675 | MCM3 | CCTCCACCAAAGTCGCGTGG |
| 3676 | MCM3 | CTACCTCCACCAAAGTCGCG |
| 3677 | MCM3 | CCTCCACGCGACTTTGGTGG |
| 3678 | MCM3 | TTCCCAGGATGACTCCACCC |
| 3679 | MCM3 | CTCCACATCGTCCAGCACCA |
| 3680 | MCM3 | TGCTGGACGATGTGGAGCTG |
| 3681 | MCM3 | TGGGAACCTCCACGCGACTT |
| 3682 | MCM4 | TGGGAGCGCTACTCGCCAGG |
| 3683 | MCM4 | CCGCTTGCCGACGACGCTCG |
| 3684 | MCM4 | GGCGCGCGGTTACCGTGGAA |
| 3685 | MCM4 | GCGCGGTTACCGTGGAAAGG |
| 3686 | MCM4 | GGCAAGCGGCCGCCTTTCCA |
| 3687 | MCM4 | TTTCCACGGTAACCGCGCGC |
| 3688 | MCM4 | TCGCGGACTCCGAGTCCACC |
| 3689 | MCM5 | TTTCCGCACTTCACAAACCA |
| 3690 | MCM5 | TTCCGCACTTCACAAACCAG |
| 3691 | MCM5 | TGTGAAGTGCGGAAAACCAG |
| 3692 | MCM5 | GCTCAGCCGCCGAGTTTCGC |
| 3693 | MCM5 | CGCTCAGCCGCCGAGTTTCG |
| 3694 | MCM5 | AAAACCAGAGGTGAGGCTAG |
| 3695 | MCM5 | CTCCCCTGGTTTGTGAAGTG |
| 3696 | MCM6 | AGCAAATATGGACCTCGCGG |
| 3697 | MCM6 | CTAAGCAAATATGGACCTCG |

| | | |
|---|---|---|
| 3698 | MCM6 | GTCTGGAAAAAAGCGACTTG |
| 3699 | MCM6 | TGGAAAAAAGCGACTTGTGG |
| 3700 | MCM6 | TCCATATTTGCTTAGTGCCG |
| 3701 | MCM6 | GACTTCTTGGAGGAGTAAGT |
| 3702 | MCM6 | GCGTGGCGCAGGCGAATCCT |
| 3703 | MCM7 | GGAAGCTGAGAATCTCCGCG |
| 3704 | MCM7 | AGCTGAGAATCTCCGCGCGG |
| 3705 | MCM7 | CGGAGATTCTCAGCTTCCCC |
| 3706 | MCM7 | TTCAGTGCCATCGCTGCCGA |
| 3707 | MCM7 | ACCTCTGAGCCCGCCAAGCG |
| 3708 | MCM7 | CATCGCTGCCGAGGGCCGTG |
| 3709 | MCM7 | GCCGCGCTTGGCGGGCTCAG |
| 3710 | MIAT-001 | TCGCGGGACCAGAGGGAGGG |
| 3711 | MIAT-001 | CGCTCGCGGGACCAGAGGGA |
| 3712 | MIAT-001 | TGAGATGGGAGGCGCTTGCA |
| 3713 | MIAT-001 | GCGCTCGCGGGACCAGAGGG |
| 3714 | MIAT-001 | GGCTCACAGCGCAGAACTGG |
| 3715 | MIAT-001 | CGGCTCACAGCGCAGAACTG |
| 3716 | MIAT-001 | GTCCGCAGCCTCCCTCCCTC |
| 3717 | MIATNB-008 | TTATGGTTAAAAGCATACGA |
| 3718 | MIATNB-008 | ATTAAAACATTCCTCCAGCC |
| 3719 | MIATNB-008 | AATTAAAACATTCCTCCAGC |
| 3720 | MIATNB-008 | AATTCTTCGGAAATACCAAG |
| 3721 | MIATNB-008 | TATGGTTAAAAGCATACGAT |
| 3722 | MIATNB-008 | AGCTGCTTCTAACAGCCTCA |
| 3723 | MIATNB-008 | TGTTAGAAGCAGCTTGTCAT |
| 3724 | MIR155HG-001 | GGGGAAAACACCAACTACCC |
| 3725 | MIR155HG-001 | CACGCAATGACCCACGAGAA |
| 3726 | MIR155HG-001 | AGGGGAAAACACCAACTACC |
| 3727 | MIR155HG-001 | TTCCCCTTTCCCTTTCTCGT |
| 3728 | MIR155HG-001 | TTTCCCCTTTCCCTTTCTCG |
| 3729 | MIR155HG-001 | TGACCCACGAGAAAGGGAAA |
| 3730 | MIR155HG-001 | GTCGAAAAAGCCCAGCGCCC |
| 3731 | MIR193BHG-001 | CATCACTTGGATCTCGTCTG |
| 3732 | MIR193BHG-001 | CGAGATCCAAGTGATGGGCG |
| 3733 | MIR193BHG-001 | GGAAGTACCCTTCTTCTCCA |
| 3734 | MIR193BHG-001 | CCCAGACGAGATCCAAGTGA |
| 3735 | MIR193BHG-001 | CCAGACGAGATCCAAGTGAT |
| 3736 | MIR193BHG-001 | ACGAGATCCAAGTGATGGGC |
| 3737 | MIR193BHG-001 | TCCTCCACCGTGGAGAAGAA |
| 3738 | MIR2117HG-001 | AGGATCCATTCTACAAAGCA |
| 3739 | MIR2117HG-001 | CATGTTTATAGAGAGAGCAG |
| 3740 | MIR2117HG-001 | CTGCTCTCTCTATAAACATG |

| | | |
|---|---|---|
| 3741 | MIR2117HG-001 | ATTGCCTTGAGGTGAAAGTG |
| 3742 | MIR2117HG-001 | AGGTGAAAGTGAGGTAAGGT |
| 3743 | MIR2117HG-001 | GATCTGATCTGATTGCCTTG |
| 3744 | MIR2117HG-001 | GGTGAAAGTGAGGTAAGGTG |
| 3745 | MIR222HG-002 | ACTTGGTCATCTATCACATG |
| 3746 | MIR222HG-002 | CCTGGGCTGCTGCTCAACGG |
| 3747 | MIR222HG-002 | ATCACATGTGGCTGAATCAC |
| 3748 | MIR222HG-002 | GCCTGGGCTGCTGCTCAACG |
| 3749 | MIR222HG-002 | CAGCAGCCCAGGCAGGAGAG |
| 3750 | MIR222HG-002 | TGCTGCTCAACGGGGGCTGG |
| 3751 | MIR222HG-002 | CTGCCTGGGCTGCTGCTCAA |
| 3752 | MIR22HG-001 | GTGTGGGGGTTGCTGCACGA |
| 3753 | MIR22HG-001 | CCCCACACCCAGCACCAGGT |
| 3754 | MIR22HG-001 | CCCCCACACCCAGCACCAGG |
| 3755 | MIR22HG-001 | AGCACCAGGTGGGTGTGAGC |
| 3756 | MIR22HG-001 | GTGGGTGTGAGCTGGCGACC |
| 3757 | MIR22HG-001 | ATCACTGCGTCCGCTCCCCA |
| 3758 | MIR22HG-001 | TGTGGGGGTTGCTGCACGAG |
| 3759 | MIR22HG-002 | TCCAGGCTCCGATCAGCTGG |
| 3760 | MIR22HG-002 | TCACGACGCCGCCAGCTGAT |
| 3761 | MIR22HG-002 | GGCTCCAGGCTCCGATCAGC |
| 3762 | MIR22HG-002 | CCGAGAAGAGACAGCGCCGC |
| 3763 | MIR22HG-002 | GCCGCCAGCTGATCGGAGCC |
| 3764 | MIR22HG-002 | AGACAGCGCCGCCGGCCGTG |
| 3765 | MIR22HG-002 | ACCCAGCACACACCGGCTCC |
| 3766 | MIR22HG-004 | CGGCCACACCCAGCCCTTCC |
| 3767 | MIR22HG-004 | GCGGGGGCAGGGGCGCCTCG |
| 3768 | MIR22HG-004 | CCCGCCGCCCCTTCTCCTCC |
| 3769 | MIR22HG-004 | CAGGGGCGCCTCGGGGAGAG |
| 3770 | MIR22HG-004 | GCAGGGGCGCCTCGGGGAGA |
| 3771 | MIR22HG-004 | GGGAGGAGAAGGGGCGGCGG |
| 3772 | MIR22HG-004 | CGGGAGGAGAAGGGGCGGCG |
| 3773 | MIR22HG-008 | ACGCACGAGCTGCGAATGCA |
| 3774 | MIR22HG-008 | CACCCTCCATCCCGTTACCG |
| 3775 | MIR22HG-008 | GGGTGAAAGTGGGGTCCGGG |
| 3776 | MIR22HG-008 | CGGGATGGAGGGTGAAAGTG |
| 3777 | MIR22HG-008 | ACGGGATGGAGGGTGAAAGT |
| 3778 | MIR22HG-008 | TGGAGGGTGAAAGTGGGGTC |
| 3779 | MIR22HG-008 | CTTCTCCTCCCGGAAGGGCT |
| 3780 | MIR22HG-010 | TAGCTCCTGACACACTAACC |
| 3781 | MIR22HG-010 | CACACTAACCCGGCCAGTCC |
| 3782 | MIR22HG-010 | CCACACCCAGCCCTTCCGGG |
| 3783 | MIR22HG-010 | ACTGGCCGGGTTAGTGTGTC |

| | | |
|---|---|---|
| 3784 | MIR22HG-010 | CGCCCCTTCTCCTCCCGGAA |
| 3785 | MIR22HG-010 | AGCCCTTCCGGGAGGAGAAG |
| 3786 | MIR22HG-010 | AGTCCAGGCCCCTCTCCCCG |
| 3787 | MIR4435-2HG-001 | ATGACAGAAACATACTCACC |
| 3788 | MIR4435-2HG-001 | ACTCACCAGGGAGAGGCTGG |
| 3789 | MIR4435-2HG-001 | TGACAGAAACATACTCACCA |
| 3790 | MIR4435-2HG-001 | GTGTACATCATTGGGAATGG |
| 3791 | MIR4435-2HG-001 | CATACTCACCAGGGAGAGGC |
| 3792 | MIR4435-2HG-001 | GAAACATACTCACCAGGGAG |
| 3793 | MIR4435-2HG-001 | TGGAGCCACCAGCCTCTCCC |
| 3794 | MIR4435-2HG-011 | GGAGTGCACCATGTGAAGAC |
| 3795 | MIR4435-2HG-011 | CCAGGCTACCTCCAAAGGGA |
| 3796 | MIR4435-2HG-011 | TCAAGACAGACACACACTCA |
| 3797 | MIR4435-2HG-011 | ACCACCAGAAGCCGAGAGAG |
| 3798 | MIR4435-2HG-011 | GCCTCTCTCTCGGCTTCTGG |
| 3799 | MIR4435-2HG-011 | GTTTAAGTCCAGTCTTCACA |
| 3800 | MIR4435-2HG-011 | TTCAAGACAGACACACACTC |
| 3801 | MIR4435-2HG-014 | GACCTTAGGAAGCACTGTCG |
| 3802 | MIR4435-2HG-014 | TTTACCGTGGCCCTATGAAG |
| 3803 | MIR4435-2HG-014 | CTGTCGAGGATGCAGCGCAA |
| 3804 | MIR4435-2HG-014 | CAGCTTTCCAGTGTCTCACA |
| 3805 | MIR4435-2HG-014 | ATTAAAATCACAACGAGATA |
| 3806 | MIR4435-2HG-014 | TCTTCCTCTTCATAGGGCCA |
| 3807 | MIR4435-2HG-014 | AGTAATACCTTTACTAGGAT |
| 3808 | MIR4435-2HG-019 | TGGACAGGTTGCATGTGGAA |
| 3809 | MIR4435-2HG-019 | TTGGACAGGTTGCATGTGGA |
| 3810 | MIR4435-2HG-019 | CATCTTGGACAGGTTGCATG |
| 3811 | MIR4435-2HG-019 | AGTTCACCATGAAAACCCAC |
| 3812 | MIR4435-2HG-019 | TGCATGTGGAAGGGACAGCA |
| 3813 | MIR4435-2HG-019 | GTGAACTGAGGGTTGCATCT |
| 3814 | MIR4435-2HG-019 | CTGAGGGTTGCATCTTGGAC |
| 3815 | MIR4458HG-001 | CTCCTCGTGTTACGCGTGCG |
| 3816 | MIR4458HG-001 | CCGATGAAAAACCCAAGACC |
| 3817 | MIR4458HG-001 | CCCGATGAAAAACCCAAGAC |
| 3818 | MIR4458HG-001 | CTCGTGTTACGCGTGCGCGG |
| 3819 | MIR4458HG-001 | CTGGCTGAGCCGGCTCAGTG |
| 3820 | MIR4458HG-001 | CGTGTTACGCGTGCGCGGAG |
| 3821 | MIR4458HG-001 | GCTGAGCCGGCTCAGTGAGG |
| 3822 | MIR663AHG-009 | GGGAACGACACACCACCGCT |
| 3823 | MIR663AHG-009 | CTCAGGTTCCCGAGGCCGAG |
| 3824 | MIR663AHG-009 | GCGCCTCGGTGTCCGCGCGT |
| 3825 | MIR663AHG-009 | CCAGCGCCCCCTCCTCCGGT |
| 3826 | MIR663AHG-009 | AGGTTCCCGAGGCCGAGCGG |

| | | |
|---|---|---|
| 3827 | MIR663AHG-009 | GACACCGAGGCGCCGACCGG |
| 3828 | MIR663AHG-009 | CCGAGGCGCCGACCGGAGGA |
| 3829 | MIR663AHG-029 | TACCAGGAAAAACCTGCGTG |
| 3830 | MIR663AHG-029 | CGAGTAAACCCACACACGAC |
| 3831 | MIR663AHG-029 | GGCCCTGTGAGCGATCGCGG |
| 3832 | MIR663AHG-029 | ACGGGCCCTGTGAGCGATCG |
| 3833 | MIR663AHG-029 | GCGGCAACCTTGAGCGTTCG |
| 3834 | MIR663AHG-029 | GCCCTGACCGCGAACGCTCA |
| 3835 | MIR663AHG-029 | GCCCTGTGAGCGATCGCGGT |
| 3836 | MIR663AHG-033 | GACGAGAATCACGAGCGATG |
| 3837 | MIR663AHG-033 | GACGGCGTTTGCGTGGCACG |
| 3838 | MIR663AHG-033 | TGCGGACGACGGCGTTTGCG |
| 3839 | MIR663AHG-033 | CAGCCACAAACTCAACGCCA |
| 3840 | MIR663AHG-033 | CACCGTGGGACGCTTTCCCA |
| 3841 | MIR663AHG-033 | ACAGCCACAAACTCAACGCC |
| 3842 | MIR663AHG-033 | GACGCTTTCCCAGGGCCAGG |
| 3843 | MIR663AHG-035 | GGAGAGACAAGGGGCCCCCG |
| 3844 | MIR663AHG-035 | CCTTCTTCTCGTCCCGCCCG |
| 3845 | MIR663AHG-035 | CGAGGAAGAGGAGAGACAAG |
| 3846 | MIR663AHG-035 | GGCGGACATTCGTGGCGAAT |
| 3847 | MIR663AHG-035 | TCTTCTCGTCCCGCCCGCGG |
| 3848 | MIR663AHG-035 | GGGCGAGGGGCGGACATTCG |
| 3849 | MIR663AHG-035 | CAACGCACCACCCGCAGGCG |
| 3850 | MRPL13 | GCTACTAGGAGAAGGACGTA |
| 3851 | MRPL13 | ACGTACGGTCCTGCTAGTAG |
| 3852 | MRPL13 | CGACATATTCCTCTACTAGC |
| 3853 | MRPL13 | CTTCTCCTAGTAGCCACGCC |
| 3854 | MRPL13 | TAAGGCTGAGTGACCCGGCG |
| 3855 | MRPL13 | GAATATGTCGAGTTTCTCTA |
| 3856 | MRPL13 | GGAATATGTCGAGTTTCTCT |
| 3857 | MTX1 | AGGCTGTCCAGGTCCACTGA |
| 3858 | MTX1 | TGACCAGCAGAACAGCTCCA |
| 3859 | MTX1 | GTGTTTCCGGGACAGAGGGT |
| 3860 | MTX1 | GAGCTGTTCTGCTGGTCAGG |
| 3861 | MTX1 | AGTGTTTCCGGGACAGAGGG |
| 3862 | MTX1 | GGAAGTGTTTCCGGGACAGA |
| 3863 | MTX1 | GGAGCTGTTCTGCTGGTCAG |
| 3864 | MYC | GCTGTAGTAATTCCAGCGAG |
| 3865 | MYC | GCAGCGCAGCTCTGCTCGCC |
| 3866 | MYC | GGCGAGCAGAGCTGCGCTGC |
| 3867 | MYC | CCGGCTAGGGTGGAAGAGCC |
| 3868 | MYC | TAATTCCAGCGAGAGGCAGA |
| 3869 | MYC | GTAATTCCAGCGAGAGGCAG |

| | | |
|---|---|---|
| 3870 | MYC | GAGCGGGCGGCCGGCTAGGG |
| 3871 | NDC80 | CAAATTCGAACGGCTTTGGC |
| 3872 | NDC80 | CAGTTTCCTGACACGACCCA |
| 3873 | NDC80 | TTTACAGAAATTCGGTCCCT |
| 3874 | NDC80 | TCAAATTCGAACGGCTTTGG |
| 3875 | NDC80 | GTATCTGCTAGACAGGACAG |
| 3876 | NDC80 | ACAGCGGTCATCAAAACACC |
| 3877 | NDC80 | TTGCACGGTTTACAGAAATT |
| 3878 | NDUFS2 | TCAGCGCCGCCATCTTACTC |
| 3879 | NDUFS2 | CTGCAGCCGGAGTAAGATGG |
| 3880 | NDUFS2 | TGCTGGGCTGAATCGGCAAT |
| 3881 | NDUFS2 | GCGGCGCTGAGGGCTTTGTG |
| 3882 | NDUFS2 | CTTACTCCGGCTGCAGACTG |
| 3883 | NDUFS2 | AGGGCTTTGTGCGGCTTCCG |
| 3884 | NDUFS2 | GTGCGGCTTCCGGGGCGTCG |
| 3885 | NEAT1-001 | CTGCATGGAAAATATCCGCA |
| 3886 | NEAT1-001 | AGGGCTTCAGGGGACAGACA |
| 3887 | NEAT1-001 | GCTGCATGGAAAATATCCGC |
| 3888 | NEAT1-001 | CCATGCAGCCTGCCCCACTG |
| 3889 | NEAT1-001 | AGGCAGAACAGCCACGCTCC |
| 3890 | NEAT1-001 | AGCGTGGCTGTTCTGCCTGG |
| 3891 | NEAT1-001 | GGGAGGAAGTGGCTAGCTCA |
| 3892 | NEAT1-002 | CCTCCCCCACAACTACACCC |
| 3893 | NEAT1-002 | TCCCTCCCTGTCGCTAACTC |
| 3894 | NEAT1-002 | GGCTATAAAAGCAAAAGTTG |
| 3895 | NEAT1-002 | AGCCGGAGTTAGCGACAGGG |
| 3896 | NEAT1-002 | GCCGGAGTTAGCGACAGGGA |
| 3897 | NEAT1-002 | AAAGTTGTGGCAAGTCCAGC |
| 3898 | NEAT1-002 | TCCAGCCGGAGTTAGCGACA |
| 3899 | NEAT1-004 | CCATCTAATCTCGTCTTAAG |
| 3900 | NEAT1-004 | CACTTAAGACGAGATTAGAT |
| 3901 | NEAT1-004 | ACAATTTGAAGATACTAGAT |
| 3902 | NEAT1-004 | ATTTGTTCCTCATTTGTCAC |
| 3903 | NEAT1-004 | CCACAATATAGGCATTTACA |
| 3904 | NEAT1-004 | ACGTTAAAAGCCCACAATAT |
| 3905 | NEAT1-004 | CCTTGTAAATGCCTATATTG |
| 3906 | NEAT1-005 | AGCCCTCAGCCGCGTCACCG |
| 3907 | NEAT1-005 | GCAAGGAGACTAGGTCTAGG |
| 3908 | NEAT1-005 | CTCGCTCAGCTATGCAAGAG |
| 3909 | NEAT1-005 | GAGCAATGACCCCGGTGACG |
| 3910 | NEAT1-005 | TTCCCCCTCCACCAGGCAGT |
| 3911 | NEAT1-005 | TCCCCCTCCACCAGGCAGTA |
| 3912 | NEAT1-005 | GTTCTGCCTGGGGGACCCTG |

| | | |
|---|---|---|
| 3913 | NEK2 | GGGCCCGCGACGGTTAAACG |
| 3914 | NEK2 | TCCAGGGACCAGGAACTCCA |
| 3915 | NEK2 | CGACGGTTAAACGGGGCCCA |
| 3916 | NEK2 | GCAGGTTGCGCCGCCAAGTG |
| 3917 | NEK2 | GTTAAACGGGGCCCAAGGCA |
| 3918 | NEK2 | AGGAACTCCAGGGACCTGGA |
| 3919 | NEK2 | GGCGGGTCAGTGCTGCTCGG |
| 3920 | NUF2 | CTCGCGCTCACCTACAGTGC |
| 3921 | NUF2 | ACAGGAAGACGGCGAGTTCG |
| 3922 | NUF2 | AGGTGAGCGCGAGAGGACGG |
| 3923 | NUF2 | GCACTGTAGGTGAGCGCGAG |
| 3924 | NUF2 | CGAACTCGCCGTCTTCCTGT |
| 3925 | NUF2 | CTGTCGGCGGCCGGCACTGT |
| 3926 | NUF2 | CGACAGGAAGACGGCGAGTT |
| 3927 | NUP133 | CGGCTGGGAACATGACTCCA |
| 3928 | NUP133 | GACTCCAAGGAGCAGCGACT |
| 3929 | NUP133 | AGCAGCGACTAGGACAGCGA |
| 3930 | NUP133 | TCCCAGCCGCCCCTTCTCCG |
| 3931 | NUP133 | GAGCAGCGACTAGGACAGCG |
| 3932 | NUP133 | CAGGCCAGGCTGCAACCTGA |
| 3933 | NUP133 | GACTAGGACAGCGAGGGATC |
| 3934 | NUP50-AS1-001 | TTGGGGCGTCGGCTCACGTG |
| 3935 | NUP50-AS1-001 | TTTGGGGCGTCGGCTCACGT |
| 3936 | NUP50-AS1-001 | CGACTAGCGCGCGAACATCC |
| 3937 | NUP50-AS1-001 | TGAGCCGACGCCCCAAAGCG |
| 3938 | NUP50-AS1-001 | GCGCGCTAGTCGCCGCTGAG |
| 3939 | NUP50-AS1-001 | GCGCCAGCAGCCGCGCTTTG |
| 3940 | NUP50-AS1-001 | CATCCTGGAAGACGCGTTGC |
| 3941 | OIP5 | CCACCACAAAAGTCCCCCCG |
| 3942 | OIP5 | GGGCCTGGCGTGTATTCGAA |
| 3943 | OIP5 | CCACAAAAGTCCCCCCGGGG |
| 3944 | OIP5 | TTGTGCAACGCCGCCCCGGG |
| 3945 | OIP5 | CAACGTGAGCGATGCCGCAG |
| 3946 | OIP5 | CGTTGTGCAACGCCGCCCCG |
| 3947 | OIP5 | TGCCACCACAAAAGTCCCCC |
| 3948 | OLMALINC-001 | GTCTGCCCTACCCTGGAATG |
| 3949 | OLMALINC-001 | ACCCTGGAATGGGGAAAATG |
| 3950 | OLMALINC-001 | TTCCAGGGTAGGGCAGACTG |
| 3951 | OLMALINC-001 | GAGTTCTGGAGCACTGTTCG |
| 3952 | OLMALINC-001 | TCTGGAGCACTGTTCGCGGC |
| 3953 | OLMALINC-001 | TACCTCAGTCTGCCCTACCC |
| 3954 | OLMALINC-001 | CTGTTCGCGGCCGGACTGAG |
| 3955 | OSER1-AS1-001 | GCATAGGTGACTCCACGTAG |

| | | |
|---|---|---|
| 3956 | OSER1-AS1-001 | CCTATGCAGAGGAATTCCAC |
| 3957 | OSER1-AS1-001 | CCCGTGGAATTCCTCTGCAT |
| 3958 | OSER1-AS1-001 | CCGAGCGCAAGGAGATCCCG |
| 3959 | OSER1-AS1-001 | CCCCGGGATCTCCTTGCGCT |
| 3960 | OSER1-AS1-001 | CTTGCGCTCGGTCCTCTACG |
| 3961 | OSER1-AS1-001 | AGCGCAAGGAGATCCCGGGG |
| 3962 | OSER1-AS1-002 | GAATAACAGGAAATTACGTA |
| 3963 | OSER1-AS1-002 | AGCAAATCCACGCCACTGGG |
| 3964 | OSER1-AS1-002 | AAGGCGCACGTGCTTCTGAG |
| 3965 | OSER1-AS1-002 | GGGAGCAAATCCACGCCACT |
| 3966 | OSER1-AS1-002 | TTCGGTTTGCCTCCTCCCAG |
| 3967 | OSER1-AS1-002 | AGGCGCACGTGCTTCTGAGG |
| 3968 | OSER1-AS1-002 | TAAGGCGCACGTGCTTCTGA |
| 3969 | PAXIP1-AS1-001 | ACCCAATCCCACTCCGCCCA |
| 3970 | PAXIP1-AS1-001 | TGGGAGCGGGAGCGGTTCTG |
| 3971 | PAXIP1-AS1-001 | TCGGGCTTCTTGGCCCTGGG |
| 3972 | PAXIP1-AS1-001 | GCCCAGGGCCAAGAAGCCCG |
| 3973 | PAXIP1-AS1-001 | GAGCGGTTCTGCGGCCTCCT |
| 3974 | PAXIP1-AS1-001 | CAGGGCCAAGAAGCCCGAGG |
| 3975 | PAXIP1-AS1-001 | TCCTCGGGCTTCTTGGCCCT |
| 3976 | PBK | TCTTCAAAATTGGTCCCGAG |
| 3977 | PBK | CTAGCACCAACACATACGCC |
| 3978 | PBK | CAGGAGGGTTCGAATTGCAA |
| 3979 | PBK | CGTTGCAATTCGAACCCTCC |
| 3980 | PBK | GGCGTATGTGTTGGTGCTAG |
| 3981 | PBK | CAGCTGCCGGGCGTATGTGT |
| 3982 | PBK | GAATTGCAACGGCAGCTGCC |
| 3983 | PCAT14-003 | AATGGGGAAGGTGATAACGT |
| 3984 | PCAT14-003 | AAAAGGAACAGCCCACATGA |
| 3985 | PCAT14-003 | ACTGCTTTACTACAGTAACC |
| 3986 | PCAT14-003 | AGTAAGATGTTGTTCTGCAG |
| 3987 | PCAT14-003 | GGAATTCTCTATAATTCCCA |
| 3988 | PCAT14-003 | AAAATAAAACATGGGAAATG |
| 3989 | PCAT14-003 | TGGGGAAGGTGATAACGTGG |
| 3990 | PCED1B-AS1-002 | AGCAAGAACAATAAGAGCCA |
| 3991 | PCED1B-AS1-002 | ATGTCAAAGTCTTGAAGCTT |
| 3992 | PCED1B-AS1-002 | GCAAGAACAATAAGAGCCAA |
| 3993 | PCED1B-AS1-002 | GATTAAACAATTCAGACCCT |
| 3994 | PCED1B-AS1-002 | TCTTATTGTTCTTGCTGGTG |
| 3995 | PCED1B-AS1-002 | TAGTTTCTTGCTTTAGGAAG |
| 3996 | PCED1B-AS1-002 | AGCATTTAGTTTCTTGCTTT |
| 3997 | PCED1B-AS1-004 | CTAAAAATACAAAATCAGCC |
| 3998 | PCED1B-AS1-004 | ACAAAATCAGCCGGGTGTGG |

| | | |
|---|---|---|
| 3999 | PCED1B-AS1-004 | TAATCTTGTGTTTCTGTGCA |
| 4000 | PCED1B-AS1-004 | AGCTACTCGGAGCCTGAAGC |
| 4001 | PCED1B-AS1-004 | CATGCACCCACCACCACACC |
| 4002 | PCED1B-AS1-004 | ACTAAAAATACAAAATCAGC |
| 4003 | PCED1B-AS1-004 | AATCAGCCGGGTGTGGTGGT |
| 4004 | PCED1B-AS1-006 | TGTAATCCCAGCACTCTGAG |
| 4005 | PCED1B-AS1-006 | TGAAGCAGGTAGATCACCTG |
| 4006 | PCED1B-AS1-006 | CAGGCATGAGCCACCAAGCC |
| 4007 | PCED1B-AS1-006 | TACATTTAAATAAATGAAGA |
| 4008 | PCED1B-AS1-006 | GAGGCAGGGTTTCACCATGT |
| 4009 | PCED1B-AS1-006 | CAAGACCAGCTTAGCCAACA |
| 4010 | PCED1B-AS1-006 | GCACTCTGAGAGGCTGAAGC |
| 4011 | PCED1B-AS1-007 | GGAGAAGATGACATTTCCGG |
| 4012 | PCED1B-AS1-007 | TCGTTTGCCCCAGATGCCAC |
| 4013 | PCED1B-AS1-007 | TGACATTTCCGGTGGCATCT |
| 4014 | PCED1B-AS1-007 | GACATTTCCGGTGGCATCTG |
| 4015 | PCED1B-AS1-007 | ATGACATTTCCGGTGGCATC |
| 4016 | PCED1B-AS1-009 | TACACCCAGATTACTGAGAA |
| 4017 | PCED1B-AS1-009 | GCTGAAAAGATGGTCAGAAG |
| 4018 | PCED1B-AS1-009 | CGGCTGAAAAGATGGTCAGA |
| 4019 | PCED1B-AS1-009 | GGCTGAAAAGATGGTCAGAA |
| 4020 | PCED1B-AS1-009 | CAAGCTCAAAACCAAATTCC |
| 4021 | PCED1B-AS1-009 | ACATGTGTGAAAACATGAAA |
| 4022 | PCED1B-AS1-009 | GATGGTCAGAAGGGGAAAGG |
| 4023 | PHF19 | CTGGTGGCCAGACTCCAAGT |
| 4024 | PHF19 | ACAGCGCAGAACTCACCGCG |
| 4025 | PHF19 | CTGGCCACCAGGCGCATCGG |
| 4026 | PHF19 | TCCGCCACCGATGCGCCTGG |
| 4027 | PHF19 | CCTGGTGGCCAGACTCCAAG |
| 4028 | PHF19 | AGTCTGGCCACCAGGCGCAT |
| 4029 | PHF19 | GGCCAGACTCCAAGTGGGAC |
| 4030 | PKD1P6-001 | GCGCGTGAGGATGCGCATGA |
| 4031 | PKD1P6-001 | AGCGCGTGAGGATGCGCATG |
| 4032 | PKD1P6-001 | CACGTCACTGAGGTTAGCCG |
| 4033 | PKD1P6-001 | TGAGCACGCGGGAGCGCGTG |
| 4034 | PKD1P6-001 | GAGGATGCGCATGAGGGCAG |
| 4035 | PKD1P6-001 | CAACGAGGAGCCCCTGACAC |
| 4036 | PKD1P6-001 | CGAGGAGCCCCTGACACTGG |
| 4037 | PLK1 | CGGAGGCTCTGCTCGGATCG |
| 4038 | PLK1 | TGAGTGCTGCAGTGACTGCA |
| 4039 | PLK1 | AGCGGTGCGGAGGCTCTGCT |
| 4040 | PLK1 | GACCCCGGCTTTCCCAGGGT |
| 4041 | PLK1 | ATGAGTGCTGCAGTGACTGC |

| | | |
|---|---|---|
| 4042 | PLK1 | CACGGGCACCGGCCGACCCT |
| 4043 | PLK1 | CCGGCTTTCCCAGGGTCGGC |
| 4044 | PLK4 | GTGTCAGGGAGAACTTTCCG |
| 4045 | PLK4 | GTTCTCCCTGACACCTTCCG |
| 4046 | PLK4 | CGCCGCTTGCCGTCCGAGCT |
| 4047 | PLK4 | CTAGCTCGGACGGCAAGCGG |
| 4048 | PLK4 | ACGTCACCACCAGCCTAGCT |
| 4049 | PLK4 | AGCCTAGCTCGGACGGCAAG |
| 4050 | PLK4 | CACCACCAGCCTAGCTCGGA |
| 4051 | POLA2 | CCTCCGCGGACCGTGACAGG |
| 4052 | POLA2 | GCTGAGCTTCTCAGTGACGG |
| 4053 | POLA2 | GAGGGAGCAGAAATCTCGAG |
| 4054 | POLA2 | TCCTGTCACGGTCCGCGGAG |
| 4055 | POLA2 | TCCCAAACGCCCTGAATGGA |
| 4056 | POLA2 | CTCCCAAACGCCCTGAATGG |
| 4057 | POLA2 | GGAAGCAGGACGTTCTCACC |
| 4058 | POLE2 | CAAGCCCCGCAACTTGAAGG |
| 4059 | POLE2 | TCGCAAATATGGCGCCGGAG |
| 4060 | POLE2 | CTCTCCGCCTTCAAGTTGCG |
| 4061 | POLE2 | CCAAATCGCAAATATGGCGC |
| 4062 | POLE2 | GTGGGCTCGGGCAGCTTGAA |
| 4063 | POLE2 | CAGCAAGCCCCGCAACTTGA |
| 4064 | POLE2 | CGCTCTCCGCCTTCAAGTTG |
| 4065 | POLQ | AAACTCTTCTCGGCCGATCA |
| 4066 | POLQ | CAGGCCAGGGTTCTCCCGAG |
| 4067 | POLQ | GGTTTGAGTTTGAAGACTGG |
| 4068 | POLQ | CCGCAGCTGTTGCCAGGCCA |
| 4069 | POLQ | CTCCCGAGAGGGAGGACGCT |
| 4070 | POLQ | CCTCCCTCTCGGGAGAACCC |
| 4071 | POLQ | AGATGTCCGCAGCTGTTGCC |
| 4072 | PRC1 | GCTACGCGGAGTGACATCGC |
| 4073 | PRC1 | GGGCTCAACGGCTCGCGGAG |
| 4074 | PRC1 | AGGTCCAGACCTACTCCACA |
| 4075 | PRC1 | TTGTTGCTCTCGGGGCCGTG |
| 4076 | PRC1 | CAACAACCACCCGCAAACAC |
| 4077 | PRC1 | CGGCTCGCGGAGCGGCTACG |
| 4078 | PRC1 | GTGTGGAGTAGGTCTGGACC |
| 4079 | PRKDC | GATCAGTTGATGACCGGCCA |
| 4080 | PRKDC | AGCACCGCAGCGGTCCGCAG |
| 4081 | PRKDC | CAGCAGGGAGCAACGCACAC |
| 4082 | PRKDC | TTGTCCGCTGCGGACCGCTG |
| 4083 | PRKDC | GCAGCGGTCCGCAGCGGACA |
| 4084 | PRKDC | GTGTGCGTTGCTCCCTGCTG |

| | | |
|---|---|---|
| 4085 | PRKDC | GCGGGACTCGGCGGCATGGC |
| 4086 | PSMB8-AS1-001 | CTGGACCAGGACCATCACAC |
| 4087 | PSMB8-AS1-001 | GGACCAGGACCATCACACTG |
| 4088 | PSMB8-AS1-001 | GGACAGTTAGTGCCTGGACC |
| 4089 | PSMB8-AS1-001 | TGGACCAGGACCATCACACT |
| 4090 | PSMB8-AS1-001 | GAAGCCGGTCCCCAGTGTGA |
| 4091 | PSMB8-AS1-001 | CATTCCCCCGACCCCCATAA |
| 4092 | PSMB8-AS1-001 | GGTCCCCAGTGTGATGGTCC |
| 4093 | PSMB8-AS1-004 | TCGGGGGAATGATGGGTCAA |
| 4094 | PSMB8-AS1-004 | GGTAGGGTCGTGTCATCTAA |
| 4095 | PSMB8-AS1-004 | CAGAAGACTAGAAGTCAGCC |
| 4096 | PSMB8-AS1-004 | CATCCGCTGGAAACAGGGGT |
| 4097 | PSMB8-AS1-004 | TCTTCCGAAGAAAGCGAGAA |
| 4098 | PSMB8-AS1-004 | GCTGGAAACAGGGGTGGGTA |
| 4099 | PSMB8-AS1-004 | CTAGAAGTCAGCCAGGAGCT |
| 4100 | PSMB8-AS1-005 | ATCACTTTACAAAACCAGGA |
| 4101 | PSMB8-AS1-005 | CATCACTTTACAAAACCAGG |
| 4102 | PSMB8-AS1-005 | CTTTACAAAACCAGGAGGGA |
| 4103 | PSMB8-AS1-005 | GAGCATCACTTTACAAAACC |
| 4104 | PSMB8-AS1-005 | GAGCTGGGAGTAGTGTCACG |
| 4105 | PSMB8-AS1-005 | ACTGTAACCCATCGCCAAGT |
| 4106 | PSMB8-AS1-005 | CGGGCATCCGCTGGAAACAG |
| 4107 | PSMB8-AS1-006 | CCTCGCACTTCCGTCCCTCC |
| 4108 | PSMB8-AS1-006 | TTACAGTAAGAGGTACCTCC |
| 4109 | PSMB8-AS1-006 | CTACCCACCCCTGTTTCCAG |
| 4110 | PSMB8-AS1-006 | GGCGCCTTCAAAAGCCCACT |
| 4111 | PSMB8-AS1-006 | TTCAAAAGCCCACTTGGCGA |
| 4112 | PSMB8-AS1-006 | GGGAGTAGTGTCACGCGGGG |
| 4113 | PSMB8-AS1-006 | GTAAGAGGTACCTCCAGGCC |
| 4114 | PSMD4 | TGTTGGAAAGCACTATGGTG |
| 4115 | PSMD4 | GAAAGCACTATGGTGTGGTG |
| 4116 | PSMD4 | GATGGTGTTGGAAAGCACTA |
| 4117 | PSMD4 | TGTGGTGAGGAGCTACTTCG |
| 4118 | PSMD4 | GGAGCTACTTCGGGGCAGGA |
| 4119 | PSMD4 | GAGTTGTTGTTAGGCCGTCC |
| 4120 | PSMD4 | GGTGTGGTGAGGAGCTACTT |
| 4121 | PSMG3-AS1-004 | CGCCCTTTCCTGAAAAAACG |
| 4122 | PSMG3-AS1-004 | GGGGCTCCCGGGCTTCTGAG |
| 4123 | PSMG3-AS1-004 | TTTCCTGAAAAAACGGGGTT |
| 4124 | PSMG3-AS1-004 | CCCAAGCCCTGCCGGCAGCG |
| 4125 | PSMG3-AS1-004 | CGGGTTCCGCGCTGCCGGCA |
| 4126 | PSMG3-AS1-004 | AAGGCGCCCCCAAGCCCTGC |
| 4127 | PSMG3-AS1-004 | AAAAACGGGGTTCGGCTCTA |

| | | |
|---|---|---|
| 4128 | PTCSC3-001 | GATGCAATGAGTGAGAGCTA |
| 4129 | PTCSC3-001 | TAAATAATGGCATTTCAGCT |
| 4130 | PTCSC3-001 | GCAATGAGTGAGAGCTAAGG |
| 4131 | PTCSC3-001 | ACTGTTGTGGAAAAGTCAAC |
| 4132 | PTCSC3-001 | CCTGTGAGTGAAGACTGTTG |
| 4133 | PTCSC3-001 | GTTGGTCTTTCATTAAATAA |
| 4134 | PTCSC3-001 | TGGGATGCAAGAAAAACAAC |
| 4135 | PTEN | TTGAGAGTTGAGCCGCTGTG |
| 4136 | PTEN | TCATGGCTGCAGCTTCCGAG |
| 4137 | PTEN | CAACTCTCAAACTTCCATCA |
| 4138 | PTEN | TGCGCTCAGTTCTCTCCTCT |
| 4139 | PTEN | CAGGCGAGGGAGATGAGAGA |
| 4140 | PTEN | AGCCCGGCCTCGCCTCACAG |
| 4141 | PTEN | AGTTGAGCCGCTGTGAGGCG |
| 4142 | PVT1-002 | CGAGAGTTTCCAGAAACGCA |
| 4143 | PVT1-002 | TGCGAGCAAAGATGTGCCCC |
| 4144 | PVT1-002 | CTGCGAGCAAAGATGTGCCC |
| 4145 | PVT1-002 | AATCCACTGGAAGGTGCCGG |
| 4146 | PVT1-002 | GAAATCCACTGGAAGGTGCC |
| 4147 | PVT1-002 | GGAAATCCACTGGAAGGTGC |
| 4148 | PVT1-002 | AAGATGTGCCCCGGGACCCC |
| 4149 | PVT1-004 | AGTGGTCTGGGGAATAACGC |
| 4150 | PVT1-004 | TACAGCTTGGATGTCCATGG |
| 4151 | PVT1-004 | TGGATGTCCATGGGGGACGA |
| 4152 | PVT1-004 | GACAGGCACAGCCATCTTGA |
| 4153 | PVT1-004 | GGATTTACAGTGATCTTCAG |
| 4154 | PVT1-004 | CAGTGATCTTCAGTGGTCTG |
| 4155 | PVT1-004 | ACAGGCACAGCCATCTTGAG |
| 4156 | PVT1-005 | CGTGTGGCGGCCGAGCACAT |
| 4157 | PVT1-005 | GCGTGTGGCGGCCGAGCACA |
| 4158 | PVT1-005 | GTCGTCGCCCCTCCTCGTCC |
| 4159 | PVT1-005 | CGGCCGAGCACATGGGCCCG |
| 4160 | PVT1-005 | GGCCCGCGGGCCCATGTGCT |
| 4161 | PVT1-005 | GGCCGAGCACATGGGCCCGC |
| 4162 | PVT1-005 | CGGGGCGGCCGGGACGAGGA |
| 4163 | PVT1-007 | GCTCTGATCAGTCAAGAAGG |
| 4164 | PVT1-007 | TTCAACTTGACACAGTCCTG |
| 4165 | PVT1-007 | AAAGCTCTGATCAGTCAAGA |
| 4166 | PVT1-007 | GAAAGATTCGCTATGACCAC |
| 4167 | PVT1-007 | AGCTCTGATCAGTCAAGAAG |
| 4168 | PVT1-007 | CGTGTCTGGGAGAAACCTCG |
| 4169 | PVT1-007 | AAGCTCTGATCAGTCAAGAA |
| 4170 | PVT1-009 | GGTCTAAAGCTTCGGCACAA |

| | | |
|---|---|---|
| 4171 | PVT1-009 | TGCCGAAGCTTTAGACCACG |
| 4172 | PVT1-009 | GGAGGTGAGGACGTGCCTCG |
| 4173 | PVT1-009 | CCTGTAGAAAGTTCCACTCA |
| 4174 | PVT1-009 | TGGTCTAAAGCTTCGGCACA |
| 4175 | PVT1-009 | TGCCTCGTGGTCTAAAGCTT |
| 4176 | PVT1-009 | TCAATCCTCAGAACCCTGAG |
| 4177 | PVT1-010 | CACTTACGGGTATGACTGTG |
| 4178 | PVT1-010 | TGTACCCATTGAATTCCCAG |
| 4179 | PVT1-010 | ACCACTTAACAAATCCCTCT |
| 4180 | PVT1-010 | CCACTTACGGGTATGACTGT |
| 4181 | PVT1-010 | CCGTAAGTGGAATTCCAGTT |
| 4182 | PVT1-010 | GTACCCATTGAATTCCCAGA |
| 4183 | PVT1-010 | AACCACTTAACAAATCCCTC |
| 4184 | PVT1-013 | TGCTTCACCAGGAAGAGTCG |
| 4185 | PVT1-013 | TCGGGGTCTTACATTCCATA |
| 4186 | PVT1-013 | ACGCTGGTGGAACCATGCAC |
| 4187 | PVT1-013 | ACATTTCAGGATACTAAAAG |
| 4188 | PVT1-013 | GTCGGGGTCTTACATTCCAT |
| 4189 | PVT1-013 | CCAAGCTGTAAGAATCCATG |
| 4190 | PVT1-013 | CCTCATGGATTCTTACAGCT |
| 4191 | PVT1-014 | CATCTGATGCACGTTCCATC |
| 4192 | PVT1-014 | GCACTGGAATGACACACGCC |
| 4193 | PVT1-014 | CGTGCATCAGATGCTTCACC |
| 4194 | PVT1-014 | TTTAGTATCCTGAAATGTGC |
| 4195 | PVT1-014 | TGTAAGACCCCGACTCTTCC |
| 4196 | PVT1-014 | GATCTCTGCCAACTTCCTTT |
| 4197 | PVT1-014 | TTAGTATCCTGAAATGTGCC |
| 4198 | PVT1-015 | CAATAACACACGTCAGGGCA |
| 4199 | PVT1-015 | CATCACGCCAACCACACAGA |
| 4200 | PVT1-015 | GCATCACGCCAACCACACAG |
| 4201 | PVT1-015 | GGCTGGTGTTCTCAACGAGG |
| 4202 | PVT1-015 | TGTGGCTGGTGTTCTCAACG |
| 4203 | PVT1-015 | TTTAATGAGCACCCTCTGTG |
| 4204 | PVT1-015 | TGTGTGGTTGGCGTGATGCT |
| 4205 | PVT1-017 | ACTGCATCTTAGACACAGCG |
| 4206 | PVT1-017 | AGTGTTTGAGATGACTAGCA |
| 4207 | PVT1-017 | CAGTGTTTGAGATGACTAGC |
| 4208 | PVT1-017 | CATTGATGAGATTCTGACAT |
| 4209 | PVT1-017 | ACATCAAGACAGATCAGCAC |
| 4210 | PVT1-017 | ATTATTTACCTGCTCAGCTC |
| 4211 | PVT1-017 | CTTGATGTGAATGTGAACAA |
| 4212 | PVT1-019 | CTAGACTCTGAGAACCAGGT |
| 4213 | PVT1-019 | TCCACTTATTCATTGTGCCT |

| | | |
|---|---|---|
| 4214 | PVT1-019 | TGGACTAGACTCTGAGAACC |
| 4215 | PVT1-019 | CTCCCTCCACTAGGGACAGA |
| 4216 | PVT1-019 | TAGACTCTGAGAACCAGGTC |
| 4217 | PVT1-019 | CCTCCCTCCACTAGGGACAG |
| 4218 | PVT1-019 | AGACTCTGAGAACCAGGTCG |
| 4219 | RAB30-AS1-001 | TCCCAGTACTCCTAACATGG |
| 4220 | RAB30-AS1-001 | GGTGCGCAGGAGTCAAGTGG |
| 4221 | RAB30-AS1-001 | GCACCCCTAAAACCTCTGCC |
| 4222 | RAB30-AS1-001 | TCGTCCCAGTACTCCTAACA |
| 4223 | RAB30-AS1-001 | GGAGTACTGGGACGATTCCG |
| 4224 | RAB30-AS1-001 | GCCGCCATGTTAGGAGTACT |
| 4225 | RAB30-AS1-001 | GGCCGCCATGTTAGGAGTAC |
| 4226 | RAB30-AS1-003 | AGTCAAGTGGTGGTTCAGTG |
| 4227 | RAB30-AS1-003 | GGAGTCAAGTGGTGGTTCAG |
| 4228 | RAB30-AS1-003 | AAGTGGTGGTTCAGTGGGGA |
| 4229 | RAB30-AS1-003 | CTCTGGATGCCGCTTCCAGT |
| 4230 | RAB30-AS1-003 | CGACTGGAAGCGGCATCCAG |
| 4231 | RAB30-AS1-003 | AGGGCGGGACCGACTGGAAG |
| 4232 | RAB30-AS1-003 | GCTGGGGACTGATGTAGGGC |
| 4233 | RAB30-AS1-004 | ACGATCTGGTCGGATAGCCG |
| 4234 | RAB30-AS1-004 | CAGCTCCTGAAGGGGTTGCA |
| 4235 | RAB30-AS1-004 | GAGGTGGAGGACGATCTGGT |
| 4236 | RAB30-AS1-004 | AAAATGGAGAAGGAAGCGTA |
| 4237 | RAB30-AS1-004 | AAAAATGGAGAAGGAAGCGT |
| 4238 | RAB30-AS1-004 | GCTCCTGAAGGGGTTGCATG |
| 4239 | RAB30-AS1-004 | AGCTCCTGAAGGGGTTGCAT |
| 4240 | RAB30-AS1-006 | CGATCTGGTCGGATAGCCGC |
| 4241 | RAB30-AS1-006 | CCCTACATCAGTCCCCAGCA |
| 4242 | RAB30-AS1-006 | GCACGAACCTGGGAACCAGA |
| 4243 | RAB30-AS1-006 | GGCACGAACCTGGGAACCAG |
| 4244 | RAB30-AS1-006 | ATAGCCGCGGGCACGAACCT |
| 4245 | RAB30-AS1-006 | CGAACCTGGGAACCAGAGGG |
| 4246 | RAB30-AS1-006 | GGTTCCCAGGTTCGTGCCCG |
| 4247 | RAD18 | AGCATCCTCGGGAGCGACCA |
| 4248 | RAD18 | GTCAGCCACCCACTAGCCTC |
| 4249 | RAD18 | CACTCACCTTCATGACTGCC |
| 4250 | RAD18 | AGGCCACCGAGACTCGGCCA |
| 4251 | RAD18 | TGGACTCCCTGGCCGAGTCT |
| 4252 | RAD18 | GTCGCTCCCGAGGATGCTGG |
| 4253 | RAD18 | GGAGTCCATGGTCGCTCCCG |
| 4254 | RAD21 | AGCCTTTGTAGATCTCAGAA |
| 4255 | RAD21 | ATCCATTCTGAGATCTACAA |
| 4256 | RAD21 | TTACCGAACAAGCGATGATG |

| | | |
|---|---|---|
| 4257 | RAD21 | TACCGAACAAGCGATGATGC |
| 4258 | RAD21 | GATGCGGGCCCAAGCTGCAT |
| 4259 | RAD21 | TGATGCGGGCCCAAGCTGCA |
| 4260 | RAD21 | GGCCCGGACCCATGCAGCTT |
| 4261 | RAD51 | GGAATTCTGAAAGCCGCTGG |
| 4262 | RAD51 | GCTTCCCGAGGCGTGCAGCT |
| 4263 | RAD51 | AGTTCCCAGCTGCACGCCTC |
| 4264 | RAD51 | GGAGAGTGCGGCGCTTCCCG |
| 4265 | RAD51 | GCGGGAATTCTGAAAGCCGC |
| 4266 | RAD51 | GAAGCGCCGCACTCTCCTTA |
| 4267 | RAD51 | GCTGGGAACTGCAACTCATC |
| 4268 | RAD51AP1 | TAGTTCCTGACATTCGTCGG |
| 4269 | RAD51AP1 | GTAGTTCCTGACATTCGTCG |
| 4270 | RAD51AP1 | TCCTGACATTCGTCGGGGGT |
| 4271 | RAD51AP1 | TCAGGAACTACCCACCTCAC |
| 4272 | RAD51AP1 | CCACCTCACAGGCCGCACCA |
| 4273 | RAD51AP1 | GGGTAGTTCCTGACATTCGT |
| 4274 | RAD51AP1 | TCCCACCCCCGACGAATGTC |
| 4275 | RALY-AS1-001 | GTCCGAGTGGGAGAAAGGAG |
| 4276 | RALY-AS1-001 | CGAGTGGGAGAAAGGAGCGG |
| 4277 | RALY-AS1-001 | GGCGGGTCCGAGTGGGAGAA |
| 4278 | RALY-AS1-001 | CGCCGCTCCTTTCTCCCACT |
| 4279 | RALY-AS1-001 | CCACTCGGACCCGCCGCCCC |
| 4280 | RALY-AS1-001 | TGGGAGAAAGGAGCGGCGGA |
| 4281 | RALY-AS1-001 | GGAGAAAGGAGCGGCGGAGG |
| 4282 | RBAKDN-001 | GTCCGGATCTAACAATCCCT |
| 4283 | RBAKDN-001 | TCCGGATCTAACAATCCCTG |
| 4284 | RBAKDN-001 | TTAGCCCACGGACCAGTCAC |
| 4285 | RBAKDN-001 | AGCAGCCCTGAGTTAGCCCA |
| 4286 | RBAKDN-001 | CTGGTCCGTGGGCTAACTCA |
| 4287 | RBAKDN-001 | TCTAACAATCCCTGGGGAGC |
| 4288 | RBAKDN-001 | TCCCCAGGGATTGTTAGATC |
| 4289 | RGPD4-AS1-006 | ACTGTGATTTGAGGCACCTG |
| 4290 | RGPD4-AS1-006 | AGGAATCAAAAAAGGATCAG |
| 4291 | RGPD4-AS1-006 | GGGGAAATACCACTAAATAC |
| 4292 | RGPD4-AS1-006 | ACCTGAGGAACTAAGAGCCC |
| 4293 | RGPD4-AS1-006 | TCATGACTAGAGATTAAGAT |
| 4294 | RGPD4-AS1-006 | GTACTCTAGCCTGGATGACA |
| 4295 | RGPD4-AS1-006 | AAAGACTAAACTGTGATTTG |
| 4296 | RMRP-001 | GTATCCTAGGCTACACACTG |
| 4297 | RMRP-001 | TTCCTCCCCTTTCCGCCTAG |
| 4298 | RMRP-001 | GGGGAAAGTCCCCGGACCTC |
| 4299 | RMRP-001 | ACTTTCCCCTAGGCGGAAAG |

| | | |
|---|---|---|
| 4300 | RMRP-001 | CACGTGGCACTCTCTGCCCG |
| 4301 | RMRP-001 | GACTTTCCCCTAGGCGGAAA |
| 4302 | RMRP-001 | CAGTGTGTAGCCTAGGATAC |
| 4303 | RNF115 | TGAAGCGGTGGGCCGCACCG |
| 4304 | RNF115 | GACGCGGGCTCTGAAGCGGT |
| 4305 | RNF115 | AGCGGTGGGCCGCACCGAGG |
| 4306 | RNF115 | GGCGACGACTGCAGCTCGGG |
| 4307 | RNF115 | CAGCTCGGGAGGTAGCGGCC |
| 4308 | RNF115 | GGGAGGTAGCGGCCTGGCGA |
| 4309 | RNF115 | GACTGCAGCTCGGGAGGTAG |
| 4310 | RNU12-001 | GTAAGGAAAATAACGATTCG |
| 4311 | RNU12-001 | GCGGGTAAAGGTCGCCCTCA |
| 4312 | RNU12-001 | CTAAAAAGTAGACTGACTGT |
| 4313 | RNU12-001 | TAAAAAGTAGACTGACTGTG |
| 4314 | RNU12-001 | CCTAAAAAGTAGACTGACTG |
| 4315 | RNU12-001 | GAGAATAGAATTCATTACCT |
| 4316 | RNU12-001 | ACTCATAAGTTTAAGGCATA |
| 4317 | RP1-122K4.3-001 | AACACCTGACACTTAGCAAG |
| 4318 | RP1-122K4.3-001 | AGAAAAATCAGAAACTTTGG |
| 4319 | RP1-122K4.3-001 | TTCTTAATCCTGTATTGACA |
| 4320 | RP1-122K4.3-001 | GTATCATTCCATGTCAATAC |
| 4321 | RP1-122K4.3-001 | TTGCTAAGTGTCAGGTGTTG |
| 4322 | RP1-122K4.3-001 | AAAGATAAAGAACCTCTCCT |
| 4323 | RP1-122K4.3-001 | GAATGATACTCTAGTCATCT |
| 4324 | RP1-149A16.17-001 | AGACACCTGATGAGTTCACC |
| 4325 | RP1-149A16.17-001 | AAGTGCCCCCTCGTCACCCT |
| 4326 | RP1-149A16.17-001 | GCACTGTCTCCTCAAACCTA |
| 4327 | RP1-149A16.17-001 | AAGCGTGTGCAGTATCTACA |
| 4328 | RP1-149A16.17-001 | CAGGTCTTCCCAGGGTGACG |
| 4329 | RP1-149A16.17-001 | GCCCTGCAACATGCACAATG |
| 4330 | RP1-149A16.17-001 | ACCTGATGAGTTCACCAGGA |
| 4331 | RP1-149A16.3-001 | AGGGGACTGCTGAAGCCTGG |
| 4332 | RP1-149A16.3-001 | TGGTGGACACAGCCCTTCTG |
| 4333 | RP1-149A16.3-001 | CAGGAGGACAGGAGCCATGG |
| 4334 | RP1-149A16.3-001 | CCTTACCTGGGAAGTGAGAG |
| 4335 | RP1-149A16.3-001 | TGGGAGGCCTGTCCTTACCT |
| 4336 | RP1-149A16.3-001 | CAGAAGGGCTGTGTCCACCA |
| 4337 | RP1-149A16.3-001 | CTCCAGCTCTGGACCTCAGA |
| 4338 | RP1-151F17.2-001 | GAACACTGGTCTATACCACA |
| 4339 | RP1-151F17.2-001 | TGTGTTTAGAAATCACCCAG |
| 4340 | RP1-151F17.2-001 | AATGTGTTTAGAAATCACCC |
| 4341 | RP1-151F17.2-001 | TTTATTTAGGTTGATCCCTG |
| 4342 | RP1-151F17.2-001 | TGAAAACTACACTTTAAGTA |

| | | |
|---|---|---|
| 4343 | RP1-151F17.2-001 | GAAAACTACACTTTAAGTAA |
| 4344 | RP1-151F17.2-001 | AGAACACTGGTCTATACCAC |
| 4345 | RP1-257A7.5-002 | TTTCTGTATGTACAAAACCT |
| 4346 | RP1-257A7.5-002 | CAAATTAAACTGGATCTCTG |
| 4347 | RP1-257A7.5-002 | CTGTCTGTTAATCCCCAGAC |
| 4348 | RP1-257A7.5-002 | CAAACTATGTACAGACACCC |
| 4349 | RP1-257A7.5-002 | CGTCAATCCCCTAATCTCCC |
| 4350 | RP1-257A7.5-002 | CATCTGTTAAATTAGAACCT |
| 4351 | RP1-257A7.5-002 | ATTAGAACCTGGGAGATTAG |
| 4352 | RP1-292L20.3-001 | GACAGGTTACCATTTGTACG |
| 4353 | RP1-292L20.3-001 | AGGCCTGGAAGAGTTAGGTA |
| 4354 | RP1-292L20.3-001 | CATGGAAGTATGGAGTGGAC |
| 4355 | RP1-292L20.3-001 | CAAAGACAAGTTAATAGAAG |
| 4356 | RP1-292L20.3-001 | ACACAGTTCTTGCTCAACTT |
| 4357 | RP1-292L20.3-001 | ATACAATCACCGCGTACAAA |
| 4358 | RP1-292L20.3-001 | CGGCACAGGACATGGAAGTA |
| 4359 | RP1-315G1.3-001 | TAGAGGCCCCGAGGGAGAGT |
| 4360 | RP1-315G1.3-001 | CCCTCGCCCAACTCTCCCTC |
| 4361 | RP1-315G1.3-001 | CCTCGCCCAACTCTCCCTCG |
| 4362 | RP1-315G1.3-001 | CCCCGAGGGAGAGTTGGGCG |
| 4363 | RP1-315G1.3-001 | AGAGGCCCCGAGGGAGAGTT |
| 4364 | RP1-315G1.3-001 | CCCGAGGGAGAGTTGGGCGA |
| 4365 | RP1-315G1.3-001 | ACTCTCCCTCGGGGCCTCTA |
| 4366 | RP1-78O14.1-003 | CTAAACCAGGAAGTTCAACG |
| 4367 | RP1-78O14.1-003 | ACCACTGGCTCACCTAAACC |
| 4368 | RP1-78O14.1-003 | AATATGTGTGAAGCTGCCAA |
| 4369 | RP1-78O14.1-003 | TAGAGAAAGAGATAAACCAC |
| 4370 | RP1-78O14.1-003 | TAAGGATAGCAGAGTTTATG |
| 4371 | RP1-78O14.1-003 | GAGAGAAAATGATGACACTG |
| 4372 | RP1-78O14.1-003 | TGGCAATGAAGTCAATACTA |
| 4373 | RP1-86C11.7-001 | GGGAAAACCAGACATTGACA |
| 4374 | RP1-86C11.7-001 | ACTTGGAAGGCTGAGGTAGG |
| 4375 | RP1-86C11.7-001 | CATTGACAAGGCCTGAGCAG |
| 4376 | RP1-86C11.7-001 | TTCCCACCCTCCCAGAAATG |
| 4377 | RP1-86C11.7-001 | TGTAGTTCTAGCTACTTGGA |
| 4378 | RP1-86C11.7-001 | GACCCCATTTCTGGGAGGGT |
| 4379 | RP1-86C11.7-001 | TGACAAGGCCTGAGCAGAGG |
| 4380 | RP11-1000B6.3-001 | CGAGCAGCTCGACAGCCTGG |
| 4381 | RP11-1000B6.3-001 | GCAGCTGGGCGGGGACACAG |
| 4382 | RP11-1000B6.3-001 | GGGGCTGGAGCCCAAACCCA |
| 4383 | RP11-1000B6.3-001 | GGCGGGGACACAGCGGCCTT |
| 4384 | RP11-1000B6.3-001 | CGGGCCCCCTCGCGCCGCTG |
| 4385 | RP11-1000B6.3-001 | GCGCCGCTGCGGCTGCTGCG |

| | | |
|---|---|---|
| 4386 | RP11-1000B6.3-001 | GACACAGCGGCCTTGGGTTT |
| 4387 | RP11-1007O24.3-001 | GCGGAGAAGACAAGGCCGGG |
| 4388 | RP11-1007O24.3-001 | GCGGATTCAGGCGGCACGCG |
| 4389 | RP11-1007O24.3-001 | CAGAGGCAGCGGAGAAGACA |
| 4390 | RP11-1007O24.3-001 | GCAGCGGAGAAGACAAGGCC |
| 4391 | RP11-1007O24.3-001 | CAGCCTAGCCCGGTGCCTCC |
| 4392 | RP11-1007O24.3-001 | CGGTCCCGAGGGACAGGCCT |
| 4393 | RP11-1007O24.3-001 | GCCTGGTGCGCAGCCTAGCC |
| 4394 | RP11-1017G21.6-001 | CCATCCCCAGCATATAACGA |
| 4395 | RP11-1017G21.6-001 | CATCTTCAGAATTAAAACTG |
| 4396 | RP11-1017G21.6-001 | AGATGCAGTATTAAGAGAGG |
| 4397 | RP11-1017G21.6-001 | TGAAGATGCAGTATTAAGAG |
| 4398 | RP11-1017G21.6-001 | CTTCAGAATTAAAACTGTGG |
| 4399 | RP11-1017G21.6-001 | TTCAGAATTAAAACTGTGGT |
| 4400 | RP11-1017G21.6-001 | AAACTGTGGTGGGCCAGGCG |
| 4401 | RP11-102N12.3-001 | CGACACGGCGCGTTAAGGCG |
| 4402 | RP11-102N12.3-001 | AGGGGCTCTGGAGCTCCAGT |
| 4403 | RP11-102N12.3-001 | CGCCTTAACGCGCCGTGTCG |
| 4404 | RP11-102N12.3-001 | GCGACTTCGGGAGCTCGAGC |
| 4405 | RP11-102N12.3-001 | GCGTTAAGGCGCGGCGACTT |
| 4406 | RP11-102N12.3-001 | GGAGCTCGAGCAGGTAGAGA |
| 4407 | RP11-102N12.3-001 | CGCGCCTTAACGCGCCGTGT |
| 4408 | RP11-1041F24.1-001 | AGCACTTTGTAAGACGAAGG |
| 4409 | RP11-1041F24.1-001 | CTTGTTGAAAGAAAGTTTCG |
| 4410 | RP11-1041F24.1-001 | GAGAAGAGAATAAATTGGTG |
| 4411 | RP11-1041F24.1-001 | CTTTGTAAGACGAAGGCGGG |
| 4412 | RP11-1041F24.1-001 | GCACTTTGTAAGACGAAGGC |
| 4413 | RP11-1041F24.1-001 | GGCAGAAGAATCGCAGGCTG |
| 4414 | RP11-1041F24.1-001 | CCCAGCACTTTGTAAGACGA |
| 4415 | RP11-104L21.3-001 | TTGAGAATTCCACAAAGCCT |
| 4416 | RP11-104L21.3-001 | CACTGCATGGTGGTTAGCTG |
| 4417 | RP11-104L21.3-001 | AGAAGAATAAAACATATCCC |
| 4418 | RP11-104L21.3-001 | CTTCCTGAAATGGTGTGTGA |
| 4419 | RP11-104L21.3-001 | TAATGTAAAAAAGGTACAAG |
| 4420 | RP11-104L21.3-001 | TTTCACTAATATTTGGTGAC |
| 4421 | RP11-104L21.3-001 | TACAAGAGGCCAGGGCACAG |
| 4422 | RP11-1070N10.3-001 | CTGTCACCAAGGCTTGCGCG |
| 4423 | RP11-1070N10.3-001 | ACAGGTCAGCCCGCCCCTAG |
| 4424 | RP11-1070N10.3-001 | GAGCGCGCTCAGTAGTGCCT |
| 4425 | RP11-1070N10.3-001 | GGCGGGCTGACCTGTCACCA |
| 4426 | RP11-1070N10.3-001 | GGCACTACTGAGCGCGCTCG |
| 4427 | RP11-1070N10.3-001 | GCGCTCGCGGTCGCCAGGAG |
| 4428 | RP11-1070N10.3-001 | TCACCAAGGCTTGCGCGCGG |

| | | |
|---|---|---|
| 4429 | RP11-108L7.14-001 | TGAGTAAGGACGTCTGAGGT |
| 4430 | RP11-108L7.14-001 | AGGGGAAATACCATTGGCAA |
| 4431 | RP11-108L7.14-001 | GGACTGAGTAAGGACGTCTG |
| 4432 | RP11-108L7.14-001 | TGTGTTGGGGAACTAAGGAA |
| 4433 | RP11-108L7.14-001 | AACATTTCTACCTTTGCCAA |
| 4434 | RP11-108L7.14-001 | AAATGTTCAAGCTTATGCAC |
| 4435 | RP11-108L7.14-001 | TGAGGTGGGAAGACCAGTCA |
| 4436 | RP11-108M12.3-002 | CAGTGTGTGCGCGCACTGTG |
| 4437 | RP11-108M12.3-002 | CCTCACTCGAGAAGCGCAAG |
| 4438 | RP11-108M12.3-002 | GCCTCACTCGAGAAGCGCAA |
| 4439 | RP11-108M12.3-002 | CCTTTCTGAGTCAAAGAAAG |
| 4440 | RP11-108M12.3-002 | GCTGGGAAGCTCGAACTGGG |
| 4441 | RP11-108M12.3-002 | ACTGTGCGGGAGCCGAAGCA |
| 4442 | RP11-108M12.3-002 | CCCCTTGCGCTTCTCGAGTG |
| 4443 | RP11-1094M14.11-001 | AGATTTCTGTCCCGGGCCAA |
| 4444 | RP11-1094M14.11-001 | CGGGCGGAGAAGCAAAGGAG |
| 4445 | RP11-1094M14.11-001 | CTCCGGCCTGAGGAGGATGG |
| 4446 | RP11-1094M14.11-001 | TTTGGCCCGGGACAGAAATC |
| 4447 | RP11-1094M14.11-001 | TACCTCCATCCTCCTCAGGC |
| 4448 | RP11-1094M14.11-001 | GCTGGAAGCACCTTTGGCCC |
| 4449 | RP11-1094M14.11-001 | GGGCGGAGAAGCAAAGGAGA |
| 4450 | RP11-1094M14.14-001 | GAAAACCCAACAGATGCTGG |
| 4451 | RP11-1094M14.14-001 | AGAAAACCCAACAGATGCTG |
| 4452 | RP11-1094M14.14-001 | AAAACAGATACCGGGCGCAG |
| 4453 | RP11-1094M14.14-001 | AAGAAAACCCAACAGATGCT |
| 4454 | RP11-1094M14.14-001 | TGAGGCAGGCAGATCACCTG |
| 4455 | RP11-1094M14.14-001 | CCTCAGCCTCCCAAAATGCT |
| 4456 | RP11-1094M14.14-001 | CAAGACCAGCCTGGCCAACA |
| 4457 | RP11-10A14.4-003 | GCCGTTTCATCAGAAGCCGG |
| 4458 | RP11-10A14.4-003 | GCCAGGGGGCAACTTCCTAC |
| 4459 | RP11-10A14.4-003 | TTTCATCAGAAGCCGGAGGA |
| 4460 | RP11-10A14.4-003 | TTCATCAGAAGCCGGAGGAC |
| 4461 | RP11-10A14.4-003 | GTCGCCGTTTCATCAGAAGC |
| 4462 | RP11-10A14.4-003 | TCCTCCGGCTTCTGATGAAA |
| 4463 | RP11-10A14.4-003 | GGGTCTGGGAAGGGGGTGGT |
| 4464 | RP11-10O22.1-001 | TCCCTGCTAACTCATTCTCA |
| 4465 | RP11-10O22.1-001 | GTGCTGATGTGGTGATGCAT |
| 4466 | RP11-10O22.1-001 | CCACAGAAATAATTGACAGC |
| 4467 | RP11-10O22.1-001 | TTAACTGATGTTTCAATCCA |
| 4468 | RP11-10O22.1-001 | TTACCTTATAAACAGGAGCT |
| 4469 | RP11-10O22.1-001 | GTTACTTCTTTGTGCTGATG |
| 4470 | RP11-10O22.1-001 | CCAGCTGTCAATTATTTCTG |
| 4471 | RP11-1100L3.8-002 | AGGAAGAAAGAGCTTGAGGT |

| | | |
|---|---|---|
| 4472 | RP11-1100L3.8-002 | AAGAGCTTGAGGTGGGAGCG |
| 4473 | RP11-1100L3.8-002 | AAAGGCTGCCCAGGAAGGCG |
| 4474 | RP11-1100L3.8-002 | CAGGAAGAAAGAGCTTGAGG |
| 4475 | RP11-1100L3.8-002 | GAAAGAGCTTGAGGTGGGAG |
| 4476 | RP11-1100L3.8-002 | ATACAGGAAGAAAGAGCTTG |
| 4477 | RP11-1100L3.8-002 | GGAAAGGCTGCCCAGGAAGG |
| 4478 | RP11-111F5.4-004 | CGACACTTCAGCACCAGTCG |
| 4479 | RP11-111F5.4-004 | CACTTCAGCACCAGTCGCGG |
| 4480 | RP11-111F5.4-004 | GCAGGCACCTATCTTGTGCA |
| 4481 | RP11-111F5.4-004 | GCGAGCAACCCGGGTCCCAG |
| 4482 | RP11-111F5.4-004 | AGACTCAGGGCGAGCAACCC |
| 4483 | RP11-111F5.4-004 | AATATAATTACATTGCAGCA |
| 4484 | RP11-111F5.4-004 | GAGACTCAGGGCGAGCAACC |
| 4485 | RP11-111M22.3-001 | TCGAGTTTCTTCGATCGTGA |
| 4486 | RP11-111M22.3-001 | TACCATTCCCACTTCACACG |
| 4487 | RP11-111M22.3-001 | AAATCCGAGCGTTCTCGCGT |
| 4488 | RP11-111M22.3-001 | CATTCCCACTTCACACGTGG |
| 4489 | RP11-111M22.3-001 | ATCGAGTTTCTTCGATCGTG |
| 4490 | RP11-111M22.3-001 | GCCCCCCAGATGGCATCTTG |
| 4491 | RP11-111M22.3-001 | TTGTTCCTGAAGTGCCGTCG |
| 4492 | RP11-111M22.3-006 | CGTAACCCGCAAATCGGCTA |
| 4493 | RP11-111M22.3-006 | CCCTAACCTCACTACCGTAA |
| 4494 | RP11-111M22.3-006 | GTAGTGAGGTTAGGGACGAT |
| 4495 | RP11-111M22.3-006 | AGTTGTCCGTAGCCGATTTG |
| 4496 | RP11-111M22.3-006 | CAACTCCGTAACCCGCAAAT |
| 4497 | RP11-111M22.3-006 | ACCTTTACGGTAGTGAGGTT |
| 4498 | RP11-111M22.3-006 | CTGCGACCTTTACGGTAGTG |
| 4499 | RP11-1149M10.2-001 | TATCAATCTCAGCCTTCACA |
| 4500 | RP11-1149M10.2-001 | CACATGATGAGAAGAGTCAG |
| 4501 | RP11-1149M10.2-001 | TACTTAAGATAACCCTGTGA |
| 4502 | RP11-1149M10.2-001 | GAGAAGAGTCAGTGGTCCTG |
| 4503 | RP11-1149M10.2-001 | ATACAAAAACTAGCCGGGTG |
| 4504 | RP11-1149M10.2-001 | CTTACCATGACTGCTGTACC |
| 4505 | RP11-1149M10.2-001 | TCACAACTGATCTTGGAGCC |
| 4506 | RP11-114H21.2-001 | TGTAATCCTAGCACTTTGGG |
| 4507 | RP11-114H21.2-001 | AAAAATACAAAAAATTAGCC |
| 4508 | RP11-114H21.2-001 | TACAAAAAATTAGCCTGGCG |
| 4509 | RP11-114H21.2-001 | AAATTTGCTGCCGGGCGCGG |
| 4510 | RP11-114H21.2-001 | TTATGAGATAAATAAGTTCT |
| 4511 | RP11-114H21.2-001 | ACTTACTTAAAATTTGCTGC |
| 4512 | RP11-114H21.2-001 | CGCCTGTAATCCTAGCACTT |
| 4513 | RP11-1152H15.1-003 | AGATATTCAAAATCAAAGTG |
| 4514 | RP11-1152H15.1-003 | AGGGTCCAGATGCCCCCCTA |

| | | |
|---|---|---|
| 4515 | RP11-1152H15.1-003 | CCATAAAATTTGTTGAACAC |
| 4516 | RP11-1152H15.1-003 | GACATTGGAATGACGCAACC |
| 4517 | RP11-1152H15.1-003 | AATGACGCAACCAGGAGCCA |
| 4518 | RP11-1152H15.1-003 | CCATGGGAAGCAGAAGAGGT |
| 4519 | RP11-1152H15.1-003 | CAAGGAATGCCTGGATCCAT |
| 4520 | RP11-116D17.2-001 | ATTTAGATGCTATGCTACAG |
| 4521 | RP11-116D17.2-001 | GTAAGTTGCACCAGACAAAG |
| 4522 | RP11-116D17.2-001 | AACAATTACATCAATCGGTG |
| 4523 | RP11-116D17.2-001 | TTAGATGCTATGCTACAGTG |
| 4524 | RP11-116D17.2-001 | TTTAGATGCTATGCTACAGT |
| 4525 | RP11-116D17.2-001 | CTTTGAATTCAGGATATACA |
| 4526 | RP11-116D17.2-001 | ATACAGGGACCTAACAAGTT |
| 4527 | RP11-116D17.4-001 | CGCACGGACAGAAACTGGCG |
| 4528 | RP11-116D17.4-001 | CAGCTACGTTTGAAAACCAA |
| 4529 | RP11-116D17.4-001 | ATTCTGATTCAGTAAGTACG |
| 4530 | RP11-116D17.4-001 | AGCTGCTCGTTAGAATCGCA |
| 4531 | RP11-116D17.4-001 | CCTGGAAACACCCAGACTTG |
| 4532 | RP11-116D17.4-001 | AGAATCGCACGGACAGAAAC |
| 4533 | RP11-116D17.4-001 | GAAACTGGCGTGGTTATTCT |
| 4534 | RP11-118B18.2-001 | AAAAGAGTGTCAGCAAAGGG |
| 4535 | RP11-118B18.2-001 | ACATTTGGTGCCGAAGAGCT |
| 4536 | RP11-118B18.2-001 | TGGAGTGAGGGCGAGGACAA |
| 4537 | RP11-118B18.2-001 | TGCCGAAGAGCTGGGACAGG |
| 4538 | RP11-118B18.2-001 | TGGTGCCGAAGAGCTGGGAC |
| 4539 | RP11-118B18.2-001 | AGTACCTTCTTAAGGGCAGT |
| 4540 | RP11-118B18.2-001 | TCCAAAAAGAGTGTCAGCAA |
| 4541 | RP11-120I21.2-001 | CCAGAGGACCATCGTGATCG |
| 4542 | RP11-120I21.2-001 | AGAGAAAGATGGGACACCAG |
| 4543 | RP11-120I21.2-001 | CGCAGTCTCCACGATCACGA |
| 4544 | RP11-120I21.2-001 | GACCCAAAACCAGGTTGTGG |
| 4545 | RP11-120I21.2-001 | GGAGTCAAAGGATTGAGGAA |
| 4546 | RP11-120I21.2-001 | CAGACCCAAAACCAGGTTGT |
| 4547 | RP11-120I21.2-001 | CCGGCGGAGTCAAAGGATTG |
| 4548 | RP11-1228E12.1-001 | GGTCATGCAAGTGATGGGAG |
| 4549 | RP11-1228E12.1-001 | CAGGTGGTCATGCAAGTGAT |
| 4550 | RP11-1228E12.1-001 | GTGGTCATGCAAGTGATGGG |
| 4551 | RP11-1228E12.1-001 | GATGACTCTGAAGAATGGCG |
| 4552 | RP11-1228E12.1-001 | CATATGTCCTGCGAGCTTAG |
| 4553 | RP11-1228E12.1-001 | ATGACTCTGAAGAATGGCGT |
| 4554 | RP11-1228E12.1-001 | CTCTGAAGAATGGCGTGGGA |
| 4555 | RP11-123O10.4-001 | ACCCGTCCACCCGCCGACGG |
| 4556 | RP11-123O10.4-001 | AGGGATATTCTTCTTCCAGC |
| 4557 | RP11-123O10.4-001 | GGGATATTCTTCTTCCAGCC |

| | | |
|---|---|---|
| 4558 | RP11-123O10.4-001 | CCTCTGCCTGCAAGCAGATA |
| 4559 | RP11-123O10.4-001 | TCGGGCTCGCTCTTTCTCGC |
| 4560 | RP11-123O10.4-001 | GCAGGGAGCGACACCATGCC |
| 4561 | RP11-123O10.4-001 | CGAGCCCGACAGCGAGCGCA |
| 4562 | RP11-1246C19.1-001 | TAGACTGGAATCTGTGCCTG |
| 4563 | RP11-1246C19.1-001 | GATCTGTGCTGGGAAAGATG |
| 4564 | RP11-1246C19.1-001 | GATGGGGAAGACGCCTGAGA |
| 4565 | RP11-1246C19.1-001 | GAGGGAAACTTGACTGCGCC |
| 4566 | RP11-1246C19.1-001 | ACTGGAATCTGTGCCTGCGG |
| 4567 | RP11-1246C19.1-001 | GGACAGAGAAATGGGTAGAC |
| 4568 | RP11-1246C19.1-001 | TTTCCCAGCACAGATCGCAG |
| 4569 | RP11-1250I15.3-001 | GAAAAAGAGCAGGCTTGGCT |
| 4570 | RP11-1250I15.3-001 | AGCATACTAAATCTGGGCAA |
| 4571 | RP11-1250I15.3-001 | TTGGCCACCTGAAAAAGAGC |
| 4572 | RP11-1250I15.3-001 | CTCAAGTGATCCACCCGCCG |
| 4573 | RP11-1250I15.3-001 | TCTAAGAGCATACTAAATCT |
| 4574 | RP11-1250I15.3-001 | CGCGGCGGGTGGATCACTTG |
| 4575 | RP11-1250I15.3-001 | GCAGGCTTGGCTGGGCACGA |
| 4576 | RP11-126O1.6-001 | GTCTACAGTGTACCACACCA |
| 4577 | RP11-126O1.6-001 | ATATTAACCGAGAATGATCA |
| 4578 | RP11-126O1.6-001 | CTTTGCAGGTAAGCACACTC |
| 4579 | RP11-126O1.6-001 | GGGCTGTTCATTCAGAGTGT |
| 4580 | RP11-126O1.6-001 | ATTCAAAGCTCCTGATCACT |
| 4581 | RP11-126O1.6-001 | TGAGTGTGCTTACCTGCAAA |
| 4582 | RP11-126O1.6-001 | GAATGAACAGCCCAGTGATC |
| 4583 | RP11-12G12.7-001 | GGGGCGTGCGTGAGTTCCGG |
| 4584 | RP11-12G12.7-001 | TTGACCGGCCTGGATGCACG |
| 4585 | RP11-12G12.7-001 | TGGGTTGCTGAGCCGATGCT |
| 4586 | RP11-12G12.7-001 | TGACCGGCCTGGATGCACGT |
| 4587 | RP11-12G12.7-001 | GCCGATGCTGGGAAGGTACG |
| 4588 | RP11-12G12.7-001 | ACCCCGTACCTTCCCAGCAT |
| 4589 | RP11-12G12.7-001 | TTGCTGAGCCGATGCTGGGA |
| 4590 | RP11-131K5.2-001 | CAGTGGTTGGAATAGTTTGG |
| 4591 | RP11-131K5.2-001 | ACTGGCTAACAGGCAGTGGT |
| 4592 | RP11-131K5.2-001 | ATATAGTAAATTGGTACCAG |
| 4593 | RP11-131K5.2-001 | AAGCGATTCGACAAGTCTCT |
| 4594 | RP11-131K5.2-001 | AAATTGGTACCAGTGGAGTG |
| 4595 | RP11-131K5.2-001 | AAATGAGGAATATGCAAAAG |
| 4596 | RP11-131K5.2-001 | TCAGCAACACCCCACTCCAC |
| 4597 | RP11-133F8.2-003 | CTCGCATTGATAGGTGATAG |
| 4598 | RP11-133F8.2-003 | ATTCAACGACTTTATCTAAG |
| 4599 | RP11-133F8.2-003 | ATTCACAAGTCTTATTGGAA |
| 4600 | RP11-133F8.2-003 | TTATCTAAGAGGAGTCAGTG |

| | | |
|---|---|---|
| 4601 | RP11-133F8.2-003 | TAAAAATATCTCTTTCCTGA |
| 4602 | RP11-133F8.2-003 | AGGAGTCAGTGAGGTCATGA |
| 4603 | RP11-133F8.2-003 | TCTTGATTCACAAGTCTTAT |
| 4604 | RP11-1348G14.4-001 | GCCACTGTGTGTCCGGTGCA |
| 4605 | RP11-1348G14.4-001 | TCAGGCTGGGAACCACACAC |
| 4606 | RP11-1348G14.4-001 | GCCCTGCACCGGACACACAG |
| 4607 | RP11-1348G14.4-001 | TGCCTGCCAGCCCCAGTGTG |
| 4608 | RP11-1348G14.4-001 | GGTGCCAAGGTTGTCAGGCT |
| 4609 | RP11-1348G14.4-001 | GTTGAATGCCACTGTGTGTC |
| 4610 | RP11-1348G14.4-001 | GTGCAGGGCTGCCTGCCACG |
| 4611 | RP11-134G8.5-001 | ATTTATCAACATGCACGCGG |
| 4612 | RP11-134G8.5-001 | GAGTTACCCTAACCTCCCAG |
| 4613 | RP11-134G8.5-001 | GTAAGCTCCTATACCCCACT |
| 4614 | RP11-134G8.5-001 | AGCTCCTATACCCCACTGGG |
| 4615 | RP11-134G8.5-001 | GTTACCCTAACCTCCCAGTG |
| 4616 | RP11-134G8.5-001 | TAAATTTATCAACATGCACG |
| 4617 | RP11-134G8.5-001 | AATTTATCAACATGCACGCG |
| 4618 | RP11-134N1.2-001 | TAGCAACTGAAAGGGGTCCG |
| 4619 | RP11-134N1.2-001 | TCCAAAGCGGTGAGTAACCT |
| 4620 | RP11-134N1.2-001 | GATAGAATAGCAACTGAAAG |
| 4621 | RP11-134N1.2-001 | GGTCCGGGATTTCTCCAAAG |
| 4622 | RP11-134N1.2-001 | TCCGAGGTTACTCACCGCTT |
| 4623 | RP11-134N1.2-001 | TCACCGCTTTGGAGAAATCC |
| 4624 | RP11-134N1.2-001 | CAGACCCAAAGAGTGTGCAG |
| 4625 | RP11-138E2.1-001 | AATAATAATAAATTAAGAGG |
| 4626 | RP11-138E2.1-001 | TGTAATCCCAGCTACTCAGG |
| 4627 | RP11-138E2.1-001 | ACAGTCTCGCTCTGTCACCC |
| 4628 | RP11-138E2.1-001 | AATAATAATAATAAATTAAG |
| 4629 | RP11-138E2.1-001 | TGAACCCGGGAGGCAGAAGT |
| 4630 | RP11-138E2.1-001 | CAAGAGAATCGCTTGAACCC |
| 4631 | RP11-138E2.1-001 | GCAAGAGAATCGCTTGAACC |
| 4632 | RP11-138I1.3-001 | GAGATGGGGTTTCACCATGT |
| 4633 | RP11-138I1.3-001 | ATCTTAATTAGCTTCTCAGT |
| 4634 | RP11-138I1.3-001 | CAAAAATACAAAAATTAGCC |
| 4635 | RP11-138I1.3-001 | CAAAAATTAGCCGGGTGTGG |
| 4636 | RP11-138I1.3-001 | ACCTCTAATTAGGTCTTAAG |
| 4637 | RP11-138I1.3-001 | CAAGTACACGCCACCACACC |
| 4638 | RP11-138I1.3-001 | GCCCCTTAAGACCTAATTAG |
| 4639 | RP11-142G1.3-001 | AACACAAACTAACTCCCATG |
| 4640 | RP11-142G1.3-001 | AAAACACAAACTAACTCCCA |
| 4641 | RP11-142G1.3-001 | AAACACAAACTAACTCCCAT |
| 4642 | RP11-142G1.3-001 | ACACATACTGAGTAGAGATG |
| 4643 | RP11-142G1.3-001 | AAGGAGTGATGGAACTGCAA |

| | | |
|---|---|---|
| 4644 | RP11-142G1.3-001 | AAACACATACTGAGTAGAGA |
| 4645 | RP11-142G1.3-001 | AAGAAAAAGCCTTACCAATA |
| 4646 | RP11-147L13.11-001 | AAACTACAATCATCTTCCTG |
| 4647 | RP11-147L13.11-001 | TAAGTTGAATGCAGTCCTAG |
| 4648 | RP11-147L13.11-001 | GGTCACTTGCACATAGGTCT |
| 4649 | RP11-147L13.11-001 | GATGACGGACTGTAGTTGAT |
| 4650 | RP11-147L13.11-001 | GGATGACGGACTGTAGTTGA |
| 4651 | RP11-147L13.11-001 | AAGTTGAATGCAGTCCTAGT |
| 4652 | RP11-147L13.11-001 | GACTGTAGTTGATGGGTTAG |
| 4653 | RP11-147L13.12-001 | TCCTGCCTCGTGATTGCAGG |
| 4654 | RP11-147L13.12-001 | TTACCTTGAATCTCATCATG |
| 4655 | RP11-147L13.12-001 | GCCACCTGCAATCACGAGGC |
| 4656 | RP11-147L13.12-001 | ATGAGCCACCTGCAATCACG |
| 4657 | RP11-147L13.12-001 | ACAGAATCCAATCAAAGATT |
| 4658 | RP11-147L13.12-001 | ACCCAGCAAAATAACTACAA |
| 4659 | RP11-147L13.12-001 | ATAACTACAATGGAGAGATC |
| 4660 | RP11-147L13.13-001 | TCCAAAACTGATGAAATGGA |
| 4661 | RP11-147L13.13-001 | GTGAATGTGGCCCGGCCCAG |
| 4662 | RP11-147L13.13-001 | ATCCAAAACTGATGAAATGG |
| 4663 | RP11-147L13.13-001 | TGAGCACACAGCCTTGGGCA |
| 4664 | RP11-147L13.13-001 | AGGGGAATGTGAATGTGGCC |
| 4665 | RP11-147L13.13-001 | TGATGAAATGGAGGGTCCTC |
| 4666 | RP11-147L13.13-001 | AATGGAGGGTCCTCTGGGCC |
| 4667 | RP11-147L13.15-001 | CGGCTGAAGCCGATCCTGAG |
| 4668 | RP11-147L13.15-001 | GCTGAAGCCGATCCTGAGTG |
| 4669 | RP11-147L13.15-001 | GCTGGGGCCCCACTCAGGAT |
| 4670 | RP11-147L13.15-001 | AGGCTCATTCCGAATTGCTG |
| 4671 | RP11-147L13.15-001 | AAGGCTCATTCCGAATTGCT |
| 4672 | RP11-147L13.15-001 | CGGGAGAGGGATGAAACAAG |
| 4673 | RP11-147L13.15-001 | GGGCCCCGGTCGGCAGAAGC |
| 4674 | RP11-14N9.2-001 | AAGGGAAATAGGACCAGGTG |
| 4675 | RP11-14N9.2-001 | GGAGTGGAATCGGTAAGTGT |
| 4676 | RP11-14N9.2-001 | AGGAGTGGAATCGGTAAGTG |
| 4677 | RP11-14N9.2-001 | TTGAATAAATTAATGTACTG |
| 4678 | RP11-14N9.2-001 | TAGAAAAGGGAAATAGGACC |
| 4679 | RP11-14N9.2-001 | AAGACCCGCTTACACTCAAA |
| 4680 | RP11-14N9.2-001 | TATAGCTTGAGTCTAGAAAA |
| 4681 | RP11-156P1.3-001 | CTCAGCCGGGCGCTCCCTCG |
| 4682 | RP11-156P1.3-001 | CGAGGGAGCGCCCGGCTGAG |
| 4683 | RP11-156P1.3-001 | GAGGCTGGGGGAGCAGCCGT |
| 4684 | RP11-156P1.3-001 | GGCAGAGGAGGCGGCCCCGA |
| 4685 | RP11-156P1.3-001 | AGGCAGAGGAGGCGGCCCCG |
| 4686 | RP11-156P1.3-001 | CCCAGGGGCACCAGAGCCAA |

| | | |
|---|---|---|
| 4687 | RP11-156P1.3-001 | GGGAGCGCCCGGCTGAGAGG |
| 4688 | RP11-156P1.3-003 | AGTGGGCAAGGATGATGGTG |
| 4689 | RP11-156P1.3-003 | ACTCGTGCTCCCACCCCCAG |
| 4690 | RP11-156P1.3-003 | GGATGATGGTGAGGCAGAGG |
| 4691 | RP11-156P1.3-003 | GCACGAGTGGGCAAGGATGA |
| 4692 | RP11-156P1.3-003 | CACTCGTGCTCCCACCCCCA |
| 4693 | RP11-156P1.3-003 | GTTGGCTCTGGTGCCCCTGG |
| 4694 | RP11-156P1.3-003 | CCACTCGTGCTCCCACCCCC |
| 4695 | RP11-159D12.2-001 | CTCACCACAGCCTGAACGTG |
| 4696 | RP11-159D12.2-001 | TTCTCACCACAGCCTGAACG |
| 4697 | RP11-159D12.2-001 | CATTTGATATATCGAGTACT |
| 4698 | RP11-159D12.2-001 | AAACAAAGACCTCTCTAATG |
| 4699 | RP11-159D12.2-001 | CCATTTGATATATCGAGTAC |
| 4700 | RP11-159D12.2-001 | AAGGAGAGTCCACATTAGAG |
| 4701 | RP11-159D12.2-001 | TCTTTAGCTGTTTCTCCCAG |
| 4702 | RP11-159G9.5-001 | TGGCCACAAGCTCTTCTCCG |
| 4703 | RP11-159G9.5-001 | GGAGGAAGAGAGCGACCAAG |
| 4704 | RP11-159G9.5-001 | GGGCGGCGAGAGCCTCATGG |
| 4705 | RP11-159G9.5-001 | CGCTCTCTTCCTCCGCCATG |
| 4706 | RP11-159G9.5-001 | GACCAAGAGGCCGAACGCCT |
| 4707 | RP11-159G9.5-001 | AAGCTCTTCTCCGAGGCGTT |
| 4708 | RP11-159G9.5-001 | AGGGGCTGGTACGCGCTGGG |
| 4709 | RP11-159G9.5-002 | TGAACGCATACCCGAAGCAA |
| 4710 | RP11-159G9.5-002 | AAGGAATTTAACAACAACAG |
| 4711 | RP11-159G9.5-002 | GTATGCGTTCACAAGCAATT |
| 4712 | RP11-159G9.5-002 | CGGAATAGCTGATCTTCTAC |
| 4713 | RP11-159G9.5-002 | TAGTCTCAAACAATACCTCC |
| 4714 | RP11-159G9.5-002 | TTCAACATGACCTTAAGTTA |
| 4715 | RP11-159G9.5-002 | AGAAGATCAGCTATTCCGGG |
| 4716 | RP11-15H20.6-004 | AGAGCGAGCTGTGAAACCGG |
| 4717 | RP11-15H20.6-004 | GTGGCTTCCGGCATTTGGTG |
| 4718 | RP11-15H20.6-004 | GTGGCGGGAGTGAGATACAA |
| 4719 | RP11-15H20.6-004 | GACCGATCCCAAGCTGTGGC |
| 4720 | RP11-15H20.6-004 | GGGGCTGGGCATCCCGAGAG |
| 4721 | RP11-15H20.6-004 | CTCCCGCCACAGCTTGGGAT |
| 4722 | RP11-15H20.6-004 | GCTGGGAAAAGGTGAGTGTG |
| 4723 | RP11-15H20.6-011 | GAGGCTGTGACAGTCACCGG |
| 4724 | RP11-15H20.6-011 | AGAGGCTGTGACAGTCACCG |
| 4725 | RP11-15H20.6-011 | GAGACTCAGGTTGGCGACCC |
| 4726 | RP11-15H20.6-011 | AAGAGGCTGTGACAGTCACC |
| 4727 | RP11-15H20.6-011 | ACAGTCACCGGGGGCTCCCG |
| 4728 | RP11-15H20.6-011 | CCCAGAACCTGCAGATCACA |
| 4729 | RP11-15H20.6-011 | CCTGCAGATCACAGGGCAAG |

| | | |
|---|---|---|
| 4730 | RP11-15J22.8-001 | AATGCGCTAAAGCAATCTGG |
| 4731 | RP11-15J22.8-001 | AATGTGAAGAAAGCTGGAGT |
| 4732 | RP11-15J22.8-001 | GAATGCGCTAAAGCAATCTG |
| 4733 | RP11-15J22.8-001 | CGGGAGGGTTTGAGCCCAAG |
| 4734 | RP11-15J22.8-001 | GCTAAAGCAATCTGGGGGCA |
| 4735 | RP11-15J22.8-001 | AGAATGCGCTAAAGCAATCT |
| 4736 | RP11-15J22.8-001 | CAGAATGCGCTAAAGCAATC |
| 4737 | RP11-160E2.6-001 | GTAGAGCACCGAAAACCCCG |
| 4738 | RP11-160E2.6-001 | CTGTCATTGAGCCATCAAGA |
| 4739 | RP11-160E2.6-001 | GGGAGAGAACGCGGTCTGAG |
| 4740 | RP11-160E2.6-001 | AAAACGCTACCTCTCTTCCT |
| 4741 | RP11-160E2.6-001 | AACGCTACCTCTCTTCCTCG |
| 4742 | RP11-160E2.6-001 | CCGAAAACCCCGAGGAAGAG |
| 4743 | RP11-160E2.6-001 | CCGTTAAGGCTATACTTTCA |
| 4744 | RP11-160H22.5-004 | CCGAGGCAGCAATGTCAGGA |
| 4745 | RP11-160H22.5-004 | ACTACCCCACCCCCAGGTAA |
| 4746 | RP11-160H22.5-004 | TCCTGACATTGCTGCCTCGG |
| 4747 | RP11-160H22.5-004 | AAAGCCACTACCCCACCCCC |
| 4748 | RP11-160H22.5-004 | GGGTCGGCGCCCTTACCTGG |
| 4749 | RP11-160H22.5-004 | GCCGAGGCAGCAATGTCAGG |
| 4750 | RP11-160H22.5-004 | AGCAGCCACAGCGTAGCAGC |
| 4751 | RP11-160O5.1-001 | CGCCTCCCCGTCGACCACCC |
| 4752 | RP11-160O5.1-001 | CCAGCCCTGGGTGGTCGACG |
| 4753 | RP11-160O5.1-001 | GCGCCTAGTGAGGGTCAGGG |
| 4754 | RP11-160O5.1-001 | GCCCTGGGTGGTCGACGGGG |
| 4755 | RP11-160O5.1-001 | TAGTGAGGGTCAGGGAGGCC |
| 4756 | RP11-160O5.1-001 | CCCAGCCCTGGGTGGTCGAC |
| 4757 | RP11-160O5.1-001 | CCCCAGCCCTGGGTGGTCGA |
| 4758 | RP11-162A12.2-001 | GCGGGGCCTTAGCGGACGGA |
| 4759 | RP11-162A12.2-001 | GGGCCGGCAGCCCTAGAGCT |
| 4760 | RP11-162A12.2-001 | CGGGGCCTTAGCGGACGGAG |
| 4761 | RP11-162A12.2-001 | GGCAGCCCTAGAGCTCGGCC |
| 4762 | RP11-162A12.2-001 | AGGCCGAGCTCTAGGGCTGC |
| 4763 | RP11-162A12.2-001 | CCCCGCCAGGCCGAGCTCTA |
| 4764 | RP11-162A12.2-001 | AGGGGAGCGCGGGGGTCACC |
| 4765 | RP11-166O4.6-001 | ACTCGCAAACTTCACAAGTG |
| 4766 | RP11-166O4.6-001 | AAAAAATTAAATTCCTGGCC |
| 4767 | RP11-166O4.6-001 | TGAAGTTTGAGGCCAGAAGT |
| 4768 | RP11-166O4.6-001 | GCCTCAAACTTCACTTTCAG |
| 4769 | RP11-166O4.6-001 | TCCTTGCTGCATAATAAAAG |
| 4770 | RP11-166O4.6-001 | ATTAAATTCCTGGCCGGGCG |
| 4771 | RP11-166O4.6-001 | TCCACTGAAAGTGAAGTTTG |
| 4772 | RP11-166P13.3-002 | CGGGACGGTACAGCAACCGG |

| | | |
|---|---|---|
| 4773 | RP11-166P13.3-002 | GGTTGCTGTACCGTCCCGCC |
| 4774 | RP11-166P13.3-002 | GCAGGGAATCAAGGTTCCTC |
| 4775 | RP11-166P13.3-002 | CCGTCATCTCTCCGGTGCCC |
| 4776 | RP11-166P13.3-002 | GGGCGGGACGGTACAGCAAC |
| 4777 | RP11-166P13.3-002 | GCTTGCGGGGCAGGGAATCA |
| 4778 | RP11-166P13.3-002 | CTCTCCGGTGCCCGGGAGAG |
| 4779 | RP11-167N5.5-001 | CTATAGCCACTAAGTACTCA |
| 4780 | RP11-167N5.5-001 | TGAGTACTTAGTGGCTATAG |
| 4781 | RP11-167N5.5-001 | AGGGGCTTTAAGCTGGAAAG |
| 4782 | RP11-167N5.5-001 | TTAAGCTGGAAAGTGGCCAG |
| 4783 | RP11-167N5.5-001 | CCTGAGTACTTAGTGGCTAT |
| 4784 | RP11-167N5.5-001 | AAGTGGCCAGGGGAGCATGA |
| 4785 | RP11-167N5.5-001 | TTGTAGCCCTGAGTACTTAG |
| 4786 | RP11-16E12.2-001 | ACAGATGACATGAAAACCCA |
| 4787 | RP11-16E12.2-001 | AAGTAACTTGAAAAGTTGGT |
| 4788 | RP11-16E12.2-001 | TGCAGGATCCTGGTAGAAGG |
| 4789 | RP11-16E12.2-001 | AAAGTAACTTGAAAAGTTGG |
| 4790 | RP11-16E12.2-001 | CTGCAGGATCCTGGTAGAAG |
| 4791 | RP11-16E12.2-001 | AGGATCCTGCAGCAGCTGAA |
| 4792 | RP11-16E12.2-001 | AGTTTGAAAGAAGGATGAGC |
| 4793 | RP11-16E12.2-003 | AGCACCAGGAAGGGTGCCCT |
| 4794 | RP11-16E12.2-003 | TCAACTGCAGAAATGTCTGC |
| 4795 | RP11-16E12.2-003 | CCCTGGGCTGAAGTCTGCAT |
| 4796 | RP11-16E12.2-003 | ACAGGCAGGCTCTGTGCCGT |
| 4797 | RP11-16E12.2-003 | CCCATGCAGACTTCAGCCCA |
| 4798 | RP11-16E12.2-003 | GGGTGGTGTCCCAAATCTGA |
| 4799 | RP11-16E12.2-003 | TGCTGCTCCACCATCAGATT |
| 4800 | RP11-16E12.2-004 | TGGGTACACCAGCCAGCCCA |
| 4801 | RP11-16E12.2-004 | TTGGGACACCACCCCCCCCC |
| 4802 | RP11-16E12.2-004 | TGGTGTCCCAAATCTGATGG |
| 4803 | RP11-16E12.2-004 | AATCTGATGGTGGAGCAGCA |
| 4804 | RP11-16E12.2-004 | AGCAGCAGGGGATCCCCTGT |
| 4805 | RP11-16E12.2-004 | ACGGCACAGAGCCTGCCTGT |
| 4806 | RP11-16E12.2-004 | GTGGGCTGGCTGGTGTACCC |
| 4807 | RP11-16N2.1-002 | AATATTCTTAGAAAGAACAC |
| 4808 | RP11-16N2.1-002 | TAATATTCTTAGAAAGAACA |
| 4809 | RP11-16N2.1-002 | GTGGAACATATGGTTCAGGT |
| 4810 | RP11-16N2.1-002 | AGTGGAACATATGGTTCAGG |
| 4811 | RP11-16N2.1-002 | AAGTGTTTGGCAAGTAGTGC |
| 4812 | RP11-16N2.1-002 | AGTGCTGGTTATTTCAGCCT |
| 4813 | RP11-16N2.1-002 | GGACTGATACAATCATCTAT |
| 4814 | RP11-171L9.1-002 | TAATTCTTGTACCTCAACCC |
| 4815 | RP11-171L9.1-002 | TAGAGCTCACTGATTGCCCA |

| | | |
|---|---|---|
| 4816 | RP11-171L9.1-002 | AGTTGATGAAAATGAAACAC |
| 4817 | RP11-171L9.1-002 | CTAGAGCTCACTGATTGCCC |
| 4818 | RP11-171L9.1-002 | TCAACTGCAAACTTGGATGC |
| 4819 | RP11-171L9.1-002 | AATTCTTGTACCTCAACCCT |
| 4820 | RP11-171L9.1-002 | ACCCTAAGAAAAATAATCTG |
| 4821 | RP11-172F10.1-002 | TGGGATATAATCTCGTGGTG |
| 4822 | RP11-172F10.1-002 | CACTAGGGAGCGCCACACAG |
| 4823 | RP11-172F10.1-002 | AGCAATCAGCCAGACTCCGT |
| 4824 | RP11-172F10.1-002 | ACTGACCTGCGCCCACTGTG |
| 4825 | RP11-172F10.1-002 | ACTAGGGAGCGCCACACAGT |
| 4826 | RP11-172F10.1-002 | TATCCCACACCTGGCTTGGA |
| 4827 | RP11-172F10.1-002 | TAGCAATCAGCCAGACTCCG |
| 4828 | RP11-173D3.1-001 | GTTCACCTCAGCACAGACCC |
| 4829 | RP11-173D3.1-001 | TATCTCCGAAGTCTGTGCTG |
| 4830 | RP11-173D3.1-001 | CAGAGCGTAGAAAGGCATCG |
| 4831 | RP11-173D3.1-001 | GTTCTCCTCAGCACAGACTT |
| 4832 | RP11-173D3.1-001 | TGTTCACCTCAGCACAGACC |
| 4833 | RP11-173D3.1-001 | GCACTATCCAGGCTGCACTG |
| 4834 | RP11-173D3.1-001 | CAAAGCGAGGCAGTGCCCCA |
| 4835 | RP11-180M15.6-001 | TGACATCTCAGACAGAGATG |
| 4836 | RP11-180M15.6-001 | ATAGTGATATGGAACAGAGA |
| 4837 | RP11-180M15.6-001 | CAGATGTTGGAAGAGTGTGG |
| 4838 | RP11-180M15.6-001 | AGATGTTGGAAGAGTGTGGA |
| 4839 | RP11-180M15.6-001 | CGAATTCATTGCTAAAGCAT |
| 4840 | RP11-180M15.6-001 | AGGCAGATGTTGGAAGAGTG |
| 4841 | RP11-180M15.6-001 | TGTCTGAGATGTCATCAGCC |
| 4842 | RP11-182L21.6-001 | GGAGCGGGAGCCGAGCCTCG |
| 4843 | RP11-182L21.6-001 | GGGGTCGGGAGGCACTGCCG |
| 4844 | RP11-182L21.6-001 | TCGGGAGGCACTGCCGGGGA |
| 4845 | RP11-182L21.6-001 | GCGCCGTGCCCCGGGCCGCG |
| 4846 | RP11-182L21.6-001 | CGGCGCCGCCTGCCCCTCCC |
| 4847 | RP11-182L21.6-001 | GCGGAGGGCAGGGGGTCGGG |
| 4848 | RP11-182L21.6-001 | GAGCCGAGCCTCGCGGCCCG |
| 4849 | RP11-184M15.2-002 | GGCAGATTGGCAAACAGTGT |
| 4850 | RP11-184M15.2-002 | GGGCCCACAGTGCGAGCCAG |
| 4851 | RP11-184M15.2-002 | TGGGCCCACAGTGCGAGCCA |
| 4852 | RP11-184M15.2-002 | AGGCAGATTGGCAAACAGTG |
| 4853 | RP11-184M15.2-002 | GCTTCAGGTATGCTCTTCAG |
| 4854 | RP11-184M15.2-002 | TGAGCTTCAGGGGAATCAGG |
| 4855 | RP11-184M15.2-002 | GATGGGAAATGGCTGACAGC |
| 4856 | RP11-18H7.1-001 | TCTTCACCAATCACCGAAGG |
| 4857 | RP11-18H7.1-001 | TTCTCTTCACCAATCACCGA |
| 4858 | RP11-18H7.1-001 | TCCAGCCAATCACAAAGCGG |

| | | |
|---|---|---|
| 4859 | RP11-18H7.1-001 | GGTGATTGGTGAAGAGAAGG |
| 4860 | RP11-18H7.1-001 | GCTGGTGAAACTGCACGGCT |
| 4861 | RP11-18H7.1-001 | CGAAGGAGGTACGCTCCCAG |
| 4862 | RP11-18H7.1-001 | CGGTGATTGGTGAAGAGAAG |
| 4863 | RP11-196G11.2-001 | CCTGCTCAAAGACCTTCTAT |
| 4864 | RP11-196G11.2-001 | TATGGGTCCTGACAGCCCAC |
| 4865 | RP11-196G11.2-001 | AGCCCACAGGAAAAGTTCAA |
| 4866 | RP11-196G11.2-001 | AGGATTGAGCCTGGCATCCG |
| 4867 | RP11-196G11.2-001 | TCCGAGGCCACTTGGCCATC |
| 4868 | RP11-196G11.2-001 | GCCTGGCATCCGAGGCCACT |
| 4869 | RP11-196G11.2-001 | AAAGTTCAAAGGATTGAGCC |
| 4870 | RP11-206L10.9-009 | ATTCCAACAGGCCAAATATG |
| 4871 | RP11-206L10.9-009 | TCCTAGCTCTGTTCACGGAA |
| 4872 | RP11-206L10.9-009 | CACACCTGCTACCTAGAACT |
| 4873 | RP11-206L10.9-009 | CCCTTTCCGTGAACAGAGCT |
| 4874 | RP11-206L10.9-009 | TCGCCCCATATTTGGCCTGT |
| 4875 | RP11-206L10.9-009 | TCTGTTCACGGAAAGGGCCT |
| 4876 | RP11-206L10.9-009 | GATTCCAACAGGCCAAATAT |
| 4877 | RP11-20D14.6-002 | CCAGATAAGAGAATGAAACC |
| 4878 | RP11-20D14.6-002 | AGTGTGGAAGGGGACCCAAG |
| 4879 | RP11-20D14.6-002 | TACAGCTCATAAAAGCAGCA |
| 4880 | RP11-20D14.6-002 | TTGGGTCCCCTTCCACACTG |
| 4881 | RP11-20D14.6-002 | CGTTCCTCCCGGCGGGCTCG |
| 4882 | RP11-20D14.6-002 | AGTGAAGCTGCAGGCCTTCG |
| 4883 | RP11-20D14.6-002 | CAGGCTGCCCCAGCCAGCAG |
| 4884 | RP11-211G23.2-001 | TTTAGAAAGTCTTCTTTCAG |
| 4885 | RP11-211G23.2-001 | GCACATAGGGAGATGAGTGA |
| 4886 | RP11-211G23.2-001 | GGAACCAAAAAGGTAATCGC |
| 4887 | RP11-211G23.2-001 | TTAGAAAGTCTTCTTTCAGA |
| 4888 | RP11-211G23.2-001 | TGGGGATGGTAACAGTGTCT |
| 4889 | RP11-211G23.2-001 | TACCATCCCCATTACAGCTG |
| 4890 | RP11-211G23.2-001 | CAGTGTCTAGGGGTACTGCT |
| 4891 | RP11-212P7.2-001 | GGGCGTCCTGCGACTGCCCG |
| 4892 | RP11-212P7.2-001 | GCCCGAGGTCGGAGGCCCGG |
| 4893 | RP11-212P7.2-001 | GTCCTGCGACTGCCCGAGGT |
| 4894 | RP11-212P7.2-001 | GGGCAGTCGCAGGACGCCCC |
| 4895 | RP11-212P7.2-001 | CTGCGACTGCCCGAGGTCGG |
| 4896 | RP11-212P7.2-001 | ACTGCCCGAGGTCGGAGGCC |
| 4897 | RP11-212P7.2-001 | CTCCGACCTCGGGCAGTCGC |
| 4898 | RP11-214N9.1-001 | GAGCCCCAGCTAGACGTGAC |
| 4899 | RP11-214N9.1-001 | CCCCAGCTAGACGTGACGGG |
| 4900 | RP11-214N9.1-001 | CCGCCCGTCACGTCTAGCTG |
| 4901 | RP11-214N9.1-001 | GGACTCATACGTCTGCGTCC |

| | | |
|---|---|---|
| 4902 | RP11-214N9.1-001 | CGAGCCCCAGCTAGACGTGA |
| 4903 | RP11-214N9.1-001 | CGAACACGGCAGCGGACCAG |
| 4904 | RP11-214N9.1-001 | TCATACGTCTGCGTCCAGGA |
| 4905 | RP11-218E20.3-001 | ACCTGCACAAAATAATACCC |
| 4906 | RP11-218E20.3-001 | GAATATCTGTTGCAGCCAAG |
| 4907 | RP11-218E20.3-001 | ACCATGCTTCTCTGTCCCAG |
| 4908 | RP11-218E20.3-001 | CAAAACAGAACTGTGTGCAA |
| 4909 | RP11-218E20.3-001 | GGATATGAAAAGAGAAGGCA |
| 4910 | RP11-218E20.3-001 | AAAACAGAACTGTGTGCAAA |
| 4911 | RP11-218E20.3-001 | TTAGATCAATGTCAGCATAA |
| 4912 | RP11-225B17.2-001 | TAATTACCATAAAACAAAGG |
| 4913 | RP11-225B17.2-001 | ATATAATTACCATAAAACAA |
| 4914 | RP11-225B17.2-001 | TTACAAAATAGATGCATTGG |
| 4915 | RP11-225B17.2-001 | TTTACAAAATAGATGCATTG |
| 4916 | RP11-225B17.2-001 | AATTACCATAAAACAAAGGA |
| 4917 | RP11-225B17.2-001 | CATTTACAAAATAGATGCAT |
| 4918 | RP11-225B17.2-001 | TAAATGCTGTTTCACATATG |
| 4919 | RP11-227G15.10-001 | TGTTAACATTTGATCACCAA |
| 4920 | RP11-227G15.10-001 | TCACCCAGGCTAGAGTGCAG |
| 4921 | RP11-227G15.10-001 | GCACCACTGCACTCTAGCCT |
| 4922 | RP11-227G15.10-001 | GATAGAAACATTACTGCATA |
| 4923 | RP11-227G15.10-001 | GAGGATCGCTTGAGCCTGGG |
| 4924 | RP11-227G15.10-001 | TATAGTCCCAGCTACTCAGG |
| 4925 | RP11-227G15.10-001 | ACTCAGGAGGCTGAAGTGGG |
| 4926 | RP11-227G15.2-001 | AAAAAGGATTTGTTCAGACA |
| 4927 | RP11-227G15.2-001 | GAATACAAATGTTAGGCTTG |
| 4928 | RP11-227G15.2-001 | CTTGTGTGCTGTACACACAG |
| 4929 | RP11-227G15.2-001 | TGTTTGTCATGTATGAAGAA |
| 4930 | RP11-227G15.2-001 | TAAAACTTTAAAAGCTGGCT |
| 4931 | RP11-227G15.2-001 | TTGTTGGGAATACAAATGTT |
| 4932 | RP11-227G15.2-001 | GCCAAGGCGGGTGGATCACA |
| 4933 | RP11-227G15.9-001 | GAGCTTTAGCTTGGTCAACA |
| 4934 | RP11-227G15.9-001 | CTTGTCTCTTTCCCTATGTG |
| 4935 | RP11-227G15.9-001 | TAAGATACCTATCTAAAGCC |
| 4936 | RP11-227G15.9-001 | TTTATCTTCTGATAATCTTG |
| 4937 | RP11-227G15.9-001 | AAAATGAACTACCTAAGACT |
| 4938 | RP11-227G15.9-001 | AAGGAAAAGAGATTTAGGCC |
| 4939 | RP11-227G15.9-001 | TGATAACAGACTAATGCAGA |
| 4940 | RP11-23E10.3-001 | CTGTTACTGATGCTGGAGTG |
| 4941 | RP11-23E10.3-001 | AAGAGGTATCTTGATGAACA |
| 4942 | RP11-23E10.3-001 | TTACTGATGCTGGAGTGAGG |
| 4943 | RP11-23E10.3-001 | AAAAGAGTGTAATATCATAT |
| 4944 | RP11-23E10.3-001 | ATAAATAGAATAATTTGAAG |

| | | |
|---|---|---|
| 4945 | RP11-23E10.3-001 | TGAACAAGGAGTCATTAGAA |
| 4946 | RP11-23E10.3-001 | CAAAAATTAGCCAGGCATGG |
| 4947 | RP11-242D8.1-001 | AAAGAGCCTAGCATCTCTCG |
| 4948 | RP11-242D8.1-001 | GGAAGGACCGAACACCTTGG |
| 4949 | RP11-242D8.1-001 | CCAGAGCCCCGAGAGATGCT |
| 4950 | RP11-242D8.1-001 | GTCACCAGGGGAGGACTGGG |
| 4951 | RP11-242D8.1-001 | AGAAAGAGCCTAGCATCTCT |
| 4952 | RP11-242D8.1-001 | AACTGTGGAACCCCTTTCCT |
| 4953 | RP11-242D8.1-001 | GAAAGAGCCTAGCATCTCTC |
| 4954 | RP11-242D8.1-002 | GGAAATCAAGACACAGTCAG |
| 4955 | RP11-242D8.1-002 | TCTGAGAGGCTATTGCTCAG |
| 4956 | RP11-242D8.1-002 | CCGTGGCAACAGTAAAGCGT |
| 4957 | RP11-242D8.1-002 | TCTTGATTTCCTATTCTGAG |
| 4958 | RP11-242D8.1-002 | GCTACTCACCGGATGAAGTG |
| 4959 | RP11-242D8.1-002 | TCCGTGGCAACAGTAAAGCG |
| 4960 | RP11-242D8.1-002 | CCATAGCGCGGGCGAGTGTG |
| 4961 | RP11-242D8.1-003 | GGGCGCTTTACGGCGAACTC |
| 4962 | RP11-242D8.1-003 | CGCCTCTACGCTTCCAGTTG |
| 4963 | RP11-242D8.1-003 | CTGGAAGCGTAGAGGCGAGA |
| 4964 | RP11-242D8.1-003 | ACTGGAAGCGTAGAGGCGAG |
| 4965 | RP11-242D8.1-003 | AGGAAGCTAAGGCCGCGTTG |
| 4966 | RP11-242D8.1-003 | AGCCGCAACTGGAAGCGTAG |
| 4967 | RP11-242D8.1-003 | GTGACGTAATAAGCCGCAAC |
| 4968 | RP11-245D16.4-001 | CCTAGTCCTGCTATGTCGTG |
| 4969 | RP11-245D16.4-001 | TTCATCCTGGACGTGCGATG |
| 4970 | RP11-245D16.4-001 | CCCCACGACATAGCAGGACT |
| 4971 | RP11-245D16.4-001 | AACCTAGTCCTGCTATGTCG |
| 4972 | RP11-245D16.4-001 | TCATCCTGGACGTGCGATGG |
| 4973 | RP11-245D16.4-001 | CTCAGTCCCCACGACATAGC |
| 4974 | RP11-245D16.4-001 | CTTCATCCTGGACGTGCGAT |
| 4975 | RP11-246A10.1-001 | TTGCGCCTACCAGAAATGAG |
| 4976 | RP11-246A10.1-001 | GCAAATCTCTCTACTCACCT |
| 4977 | RP11-246A10.1-001 | GGGCTCTCGAACTTTCCCAG |
| 4978 | RP11-246A10.1-001 | GAATCACAAGCTGGGCCACT |
| 4979 | RP11-246A10.1-001 | GAGAGCCCTAATCCTGCCAG |
| 4980 | RP11-246A10.1-001 | CGCAAATCTCTCTACTCACC |
| 4981 | RP11-246A10.1-001 | GAAAGGTTATTTGTCCTATT |
| 4982 | RP11-252E2.2-001 | GTTTGAGCTCGCAACCAACA |
| 4983 | RP11-252E2.2-001 | TCAGCCTGAAACCCATCCCT |
| 4984 | RP11-252E2.2-001 | GCAACCAACAGGGAAGATGA |
| 4985 | RP11-252E2.2-001 | AAAGGAAGTCCCAAATTATG |
| 4986 | RP11-252E2.2-001 | ATGATTACTAAGCTTTCTAG |
| 4987 | RP11-252E2.2-001 | TCTGACTTGATCAAATACAA |

| | | |
|---|---|---|
| 4988 | RP11-252E2.2-001 | CGCAACCAACAGGGAAGATG |
| 4989 | RP11-253I19.3-001 | AGATGAAACCCTCAGCAACC |
| 4990 | RP11-253I19.3-001 | GTAATCACAAACAAATGGCC |
| 4991 | RP11-253I19.3-001 | CAGGTGAAAATTATAATATG |
| 4992 | RP11-253I19.3-001 | CAAGATTTCCAGGTTGCTGA |
| 4993 | RP11-253I19.3-001 | TCAAGATTTCCAGGTTGCTG |
| 4994 | RP11-253I19.3-001 | AAATAGAATATTCCCCTCAC |
| 4995 | RP11-253I19.3-001 | TGTGGTCCCGCCTACTCGGG |
| 4996 | RP11-254F7.2-001 | GGCGTGACCTTGAGAGTACA |
| 4997 | RP11-254F7.2-001 | TGCAAAGTCAAAGCAGACGA |
| 4998 | RP11-254F7.2-001 | TAGACAGCATTCCTTCTGCG |
| 4999 | RP11-254F7.2-001 | GTTGGCATGTTCCACGCAGA |
| 5000 | RP11-254F7.2-001 | TGTCTAGAAACAGAAGAGAG |
| 5001 | RP11-254F7.2-001 | TTGCAAAGTCAAAGCAGACG |
| 5002 | RP11-254F7.2-001 | AAGTCAAAGCAGACGAGGGT |
| 5003 | RP11-255M2.3-001 | CGGGAGGTGTTCCTGCTGCG |
| 5004 | RP11-255M2.3-001 | AGGAACACCTCCCGGTGCCG |
| 5005 | RP11-255M2.3-001 | CGCGCAGCAGGAACACCTCC |
| 5006 | RP11-255M2.3-001 | GCTCCGGCCCCGGCACCGGG |
| 5007 | RP11-255M2.3-001 | GCGCGGGATCAAGGCCCAGG |
| 5008 | RP11-255M2.3-001 | GGGAGGTGTTCCTGCTGCGC |
| 5009 | RP11-255M2.3-001 | CGTTGCGGACTGCCAGGCTC |
| 5010 | RP11-258F22.1-001 | TGCATAGACACCTCTTTGGG |
| 5011 | RP11-258F22.1-001 | TTACCAGAGCAACAGTACAC |
| 5012 | RP11-258F22.1-001 | GGAGTGCAATGGCATGACCT |
| 5013 | RP11-258F22.1-001 | ACTCCAGCCTGGGCAACAAG |
| 5014 | RP11-258F22.1-001 | TTGCCCAGGCTGGAGTGCAA |
| 5015 | RP11-258F22.1-001 | GGAGAATTGCTTGAACCGGG |
| 5016 | RP11-258F22.1-001 | GAACCGGGAGGCAGAGATTG |
| 5017 | RP11-259O2.3-001 | GCATCATGACAATTCACCCG |
| 5018 | RP11-259O2.3-001 | CATCATGACAATTCACCCGT |
| 5019 | RP11-259O2.3-001 | GGAAATCGCATAGTTTCCCA |
| 5020 | RP11-259O2.3-001 | CTGATCTTTCCAAAAGACGG |
| 5021 | RP11-259O2.3-001 | TTTCTTTATTTAAGGACCCA |
| 5022 | RP11-259O2.3-001 | GCATTTAAACTGAAAATGTG |
| 5023 | RP11-259O2.3-001 | GAATTGTCATGATGCTGTTG |
| 5024 | RP11-263K4.5-001 | TTCTTCTATTGCTGTCAAGC |
| 5025 | RP11-263K4.5-001 | TGCATTTACTAGATTAGAGG |
| 5026 | RP11-263K4.5-001 | TTGCAAGACAGATTAAGTTA |
| 5027 | RP11-263K4.5-001 | TATTAACTAATATAGGTAAG |
| 5028 | RP11-263K4.5-001 | CATGAAATCTGTAATTGAAG |
| 5029 | RP11-263K4.5-001 | ATTAACTAATATAGGTAAGT |
| 5030 | RP11-263K4.5-001 | TTAACTAATATAGGTAAGTG |

| | | |
|---|---|---|
| 5031 | RP11-266K4.9-001 | GCCAGGACTGTCCTAGAACT |
| 5032 | RP11-266K4.9-001 | CCATTCTGCAAGGCAGAGGG |
| 5033 | RP11-266K4.9-001 | ACTGTCCTAGAACTTGGCAG |
| 5034 | RP11-266K4.9-001 | GGCCCATTCTGCAAGGCAGA |
| 5035 | RP11-266K4.9-001 | TGCAGATGGCAAGGCCTGTG |
| 5036 | RP11-266K4.9-001 | CTTCAGCGACCCTCACAGCC |
| 5037 | RP11-266K4.9-001 | TTCACTAACAGGTGACCTCG |
| 5038 | RP11-266K4.9-002 | TAGCATACTGAATCATCACA |
| 5039 | RP11-266K4.9-002 | CGGGCCTCAAGATTGTCACG |
| 5040 | RP11-266K4.9-002 | CATGCCACGTGACAATCTTG |
| 5041 | RP11-266K4.9-002 | GACATACCAACAGCTTTCAC |
| 5042 | RP11-266K4.9-002 | TCGATGAATGGAATCCTGTG |
| 5043 | RP11-266K4.9-002 | TTGAGGCCCGTGAAAGCTGT |
| 5044 | RP11-266K4.9-002 | GGCCGGGTGTGGCTATGAAG |
| 5045 | RP11-266K4.9-003 | ATAGATACAGAAAACCACAA |
| 5046 | RP11-266K4.9-003 | TGGTGGCCAGAGGCCGGGTG |
| 5047 | RP11-266K4.9-003 | GTCAGTGGTGGCCAGAGGCC |
| 5048 | RP11-266K4.9-003 | GGCGATGGAGCACAGGTCAG |
| 5049 | RP11-266K4.9-003 | ACACTCATCTCAGGCCACAC |
| 5050 | RP11-266K4.9-003 | TCATAGCCACACCCGGCCTC |
| 5051 | RP11-266K4.9-003 | AGACAACGGAATTCTGGCGA |
| 5052 | RP11-268P4.5-001 | CATTTCAGTCTCACCTCATG |
| 5053 | RP11-268P4.5-001 | TACATTCAGGGATCCCCATG |
| 5054 | RP11-268P4.5-001 | ATTTATTTACTTGGCCGTGG |
| 5055 | RP11-268P4.5-001 | CCATTTCAGTCTCACCTCAT |
| 5056 | RP11-268P4.5-001 | ATATGTGAAAACTAATAACC |
| 5057 | RP11-268P4.5-001 | AGCATTCCAAGAAGCTGTCC |
| 5058 | RP11-268P4.5-001 | ATTACAGGTATGTGCCACCA |
| 5059 | RP11-269F21.1-001 | AATTTATACTCATACCATCA |
| 5060 | RP11-269F21.1-001 | TGGCAACAATCATACCCTGA |
| 5061 | RP11-269F21.1-001 | CCAAACGACATCAAGGGGCA |
| 5062 | RP11-269F21.1-001 | GACACAGCCAAACGACATCA |
| 5063 | RP11-269F21.1-001 | ACACAGCCAAACGACATCAA |
| 5064 | RP11-269F21.1-001 | GGTGCTGCCCTTGACATGTG |
| 5065 | RP11-269F21.1-001 | CACAGCCAAACGACATCAAG |
| 5066 | RP11-26L16.1-001 | ATTGACAAACCCTTACCAGA |
| 5067 | RP11-26L16.1-001 | ATGGTAAAGCCAGTTGGGAA |
| 5068 | RP11-26L16.1-001 | CCAGATCAAGTTTGACTGAA |
| 5069 | RP11-26L16.1-001 | AAATCACATTTAAGGCTCAA |
| 5070 | RP11-26L16.1-001 | AGGCAAACACCATTCCCAAC |
| 5071 | RP11-26L16.1-001 | CTGGCTTTACCATCTGGTAA |
| 5072 | RP11-26L16.1-001 | ACCAGATGGTAAAGCCAGTT |
| 5073 | RP11-271C24.2-001 | TCTGCTACCCGAAGCCCCAA |

| | | |
|---|---|---|
| 5074 | RP11-271C24.2-001 | CCGCACTCACTTCCAACCCT |
| 5075 | RP11-271C24.2-001 | TCAGATGTCCACCTTTCCCA |
| 5076 | RP11-271C24.2-001 | CTCACTTCCAACCCTGGGAA |
| 5077 | RP11-271C24.2-001 | AAGCCCCAAAGGCTCTGAGT |
| 5078 | RP11-271C24.2-001 | AGCCCCAAAGGCTCTGAGTA |
| 5079 | RP11-271C24.2-001 | ACTTCCAACCCTGGGAAAGG |
| 5080 | RP11-275I4.2-002 | TCAGGTTACGAAGAGCACTA |
| 5081 | RP11-275I4.2-002 | CCATATGTTCCCTGGCTGAG |
| 5082 | RP11-275I4.2-002 | CTGGCTGAGAGGTGACTCAG |
| 5083 | RP11-275I4.2-002 | TGACATTGCCATATGTTCCC |
| 5084 | RP11-275I4.2-002 | GAGAACAGGCTGCAGCTTCG |
| 5085 | RP11-275I4.2-002 | GCACTAAGGCAGCTTTGAAG |
| 5086 | RP11-275I4.2-002 | TTCTAAAAAACATGCTGTAG |
| 5087 | RP11-277P12.10-001 | AGCAAAGAACAGGAGTACAG |
| 5088 | RP11-277P12.10-001 | CAGGAAAGGGAGATAAGGGT |
| 5089 | RP11-277P12.10-001 | AGGAAAGGGAGATAAGGGTG |
| 5090 | RP11-277P12.10-001 | GAGCAAAGAACAGGAGTACA |
| 5091 | RP11-277P12.10-001 | TCAGGAAAGGGAGATAAGGG |
| 5092 | RP11-277P12.10-001 | GTCCGAAAAGAGTCAGGAAA |
| 5093 | RP11-277P12.10-001 | TGAGGACCTGAGGTCATAGG |
| 5094 | RP11-284N8.3-001 | TTGAGTGTGCAATAAGGCTC |
| 5095 | RP11-284N8.3-001 | ATGAAATAGCTAGAAAGGAG |
| 5096 | RP11-284N8.3-001 | TTTCCTGCATAAGAAAATCT |
| 5097 | RP11-284N8.3-001 | AATGAAATAGCTAGAAAGGA |
| 5098 | RP11-284N8.3-001 | GATTAATGAAATAGCTAGAA |
| 5099 | RP11-284N8.3-001 | ATAGATCAGTGAAAATTCTA |
| 5100 | RP11-284N8.3-001 | AAAATGTTGAGTGTGCAATA |
| 5101 | RP11-28H5.2-001 | AAAAAAAGCTTTATACAGCT |
| 5102 | RP11-28H5.2-001 | TTCAAATATGCAATCTGGAC |
| 5103 | RP11-28H5.2-001 | TCAGAAGTTCAAGACCAGCC |
| 5104 | RP11-28H5.2-001 | CAGACTTCAAATATGCAATC |
| 5105 | RP11-28H5.2-001 | AAACCTGCAAAGCTAGAATT |
| 5106 | RP11-28H5.2-001 | CCACTTTGGGAGGCTGAGGA |
| 5107 | RP11-28H5.2-001 | TGTAATCCCACCACTTTGGG |
| 5108 | RP11-290F5.1-001 | TGAGGCAGGTGGATCACCTG |
| 5109 | RP11-290F5.1-001 | CAAGACCAGGCTGACCAACA |
| 5110 | RP11-290F5.1-001 | GAGGTCAAGAGTTCAAGACC |
| 5111 | RP11-290F5.1-001 | GAAAAACGGGAGGAAAAGAA |
| 5112 | RP11-290F5.1-001 | ACAAAATTAGCCGGGCGTGG |
| 5113 | RP11-290F5.1-001 | GGTCTTGAACTCTTGACCTC |
| 5114 | RP11-290F5.1-001 | ACTAAAAATACAAAATTAGC |
| 5115 | RP11-290F5.1-002 | AGTGTCTTCCTCTACTGCAG |
| 5116 | RP11-290F5.1-002 | CAGGATGTCTGTCACACCTC |

| | | |
|---|---|---|
| 5117 | RP11-290F5.1-002 | GTAGAGGAAGACACTCCAGA |
| 5118 | RP11-290F5.1-002 | TGGGAAAGCAACATATGTAG |
| 5119 | RP11-290F5.1-002 | CACACCTCAGGAAACCTGCC |
| 5120 | RP11-290F5.1-002 | TATTCCTGGCAGGTTTCCTG |
| 5121 | RP11-290F5.1-002 | GTGTGACAGACATCCTGAAG |
| 5122 | RP11-290F5.1-003 | AAACCTGCCAGGAATACCAA |
| 5123 | RP11-290F5.1-003 | TCCAGTCTCACTTGCAGCTG |
| 5124 | RP11-290F5.1-003 | CCATACCTTCCAGAGTAGTT |
| 5125 | RP11-290F5.1-003 | TGGGGTTCTAACTGTGAATT |
| 5126 | RP11-290F5.1-003 | ATATGCATTTGCAAGAGATT |
| 5127 | RP11-290F5.1-003 | CCCATACCTTCCAGAGTAGT |
| 5128 | RP11-290F5.1-003 | CAAAGGGAGTGAAAAATCCA |
| 5129 | RP11-296K13.4-001 | TTTAAAAATTAGGCCAGGTG |
| 5130 | RP11-296K13.4-001 | GCACTTTGGGAGTCTGAGGT |
| 5131 | RP11-296K13.4-001 | AAACAAACAAAAACGCTGAC |
| 5132 | RP11-296K13.4-001 | CAGGCATGAACCACCACACC |
| 5133 | RP11-296K13.4-001 | TCCAGCACTTTGGGAGTCTG |
| 5134 | RP11-296K13.4-001 | AAAAATTAGGCCAGGTGTGG |
| 5135 | RP11-296K13.4-001 | ATACAAAAATTAGCCAGGTG |
| 5136 | RP11-297N6.4-001 | CCCGGCCAATCAGTCGCGGA |
| 5137 | RP11-297N6.4-001 | GGGTAAGCGGAAGAAAACAA |
| 5138 | RP11-297N6.4-001 | AGGGCACCAATCCCGCTCGT |
| 5139 | RP11-297N6.4-001 | ACACTCTAGAAGCTCATTGG |
| 5140 | RP11-297N6.4-001 | CCGAGAAAGAGGCCGACGAG |
| 5141 | RP11-297N6.4-001 | ATAACACTCTAGAAGCTCAT |
| 5142 | RP11-297N6.4-001 | CGAGAAAGAGGCCGACGAGC |
| 5143 | RP11-2B6.3-001 | CCATACCCACCTCGCGCTCG |
| 5144 | RP11-2B6.3-001 | CGAGCGCGAGGTGGGTATGG |
| 5145 | RP11-2B6.3-001 | GCTCGCGGAGAGGGCCGTCG |
| 5146 | RP11-2B6.3-001 | TCGCGGCCGTGGGAATGCCC |
| 5147 | RP11-2B6.3-001 | GGGCATTCCCACGGCCGCGA |
| 5148 | RP11-2B6.3-001 | GAGAGGGCCGTCGCGGCCGT |
| 5149 | RP11-2B6.3-001 | GGTGCTCTCCCGGCAGCGAC |
| 5150 | RP11-304C12.5-001 | AGCAAGCTGGAAGCCTGCAG |
| 5151 | RP11-304C12.5-001 | GCAAAATAAAAGATTAGGGT |
| 5152 | RP11-304C12.5-001 | CTGTTTCTTTACTGTAACTG |
| 5153 | RP11-304C12.5-001 | TTAAAAGGGAAATTCTGCTG |
| 5154 | RP11-304C12.5-001 | GCTAGGTATGCCATTTGTAT |
| 5155 | RP11-304C12.5-001 | ACAGCTCAAAAAACTGGTTA |
| 5156 | RP11-304C12.5-001 | AATAGAAAAGGGCTACCTGG |
| 5157 | RP11-304L19.13-002 | AGTTTGTGGGAGTCGCGCTG |
| 5158 | RP11-304L19.13-002 | GAGGAAGCGTTAAAGAGTCA |
| 5159 | RP11-304L19.13-002 | TAAAGAGTCAAGGCAGTTTG |

| | | |
|---|---|---|
| 5160 | RP11-304L19.13-002 | ACTCTTTAACGCTTCCTCGC |
| 5161 | RP11-304L19.13-002 | AGTCGCGCTGGGGACGTTCA |
| 5162 | RP11-304L19.13-002 | TGGCCCAATGGGCGCCAGCG |
| 5163 | RP11-304L19.13-002 | CAGTTTGTGGGAGTCGCGCT |
| 5164 | RP11-306O13.1-001 | CTGTACATATTGCTAAACTG |
| 5165 | RP11-306O13.1-001 | TACAGGAAGTAAAGACCCTC |
| 5166 | RP11-306O13.1-001 | CATATTGCTAAACTGAGGGA |
| 5167 | RP11-306O13.1-001 | TGTACATATTGCTAAACTGA |
| 5168 | RP11-306O13.1-001 | GTTGGCAAAAATCAGTGGTT |
| 5169 | RP11-306O13.1-001 | AGTTCCCTCTCCTGTACAGA |
| 5170 | RP11-306O13.1-001 | GTCTAAATTAATCACTAGTA |
| 5171 | RP11-307E17.8-001 | AACCACCGGCAACCGATGCT |
| 5172 | RP11-307E17.8-001 | CCGGCAACCGATGCTGGGAG |
| 5173 | RP11-307E17.8-001 | CATCGGTTGCCGGTGGTTCG |
| 5174 | RP11-307E17.8-001 | GCCATGAGAGCTAAGGCCGT |
| 5175 | RP11-307E17.8-001 | CTCCCAGCATCGGTTGCCGG |
| 5176 | RP11-307E17.8-001 | CCATGAGAGCTAAGGCCGTT |
| 5177 | RP11-307E17.8-001 | CGCAGCGCGAGCGGAAAGGG |
| 5178 | RP11-308B16.2-001 | GCTGTGCTCGAGTTCTCGCG |
| 5179 | RP11-308B16.2-001 | GTCCCAAGCCCTGCCCCGCG |
| 5180 | RP11-308B16.2-001 | AGGTCCCAAGCCCTGCCCCG |
| 5181 | RP11-308B16.2-001 | GGCGCCGAGCGAGGGCTGTG |
| 5182 | RP11-308B16.2-001 | GCTTCCCCCCGCCCCCGCCG |
| 5183 | RP11-308B16.2-001 | GTCGATGGGACTGGGCATCG |
| 5184 | RP11-308B16.2-001 | CTTCCCCCCGCCCCCGCCGT |
| 5185 | RP11-309L24.4-001 | CCGACTGAAAAGGCGCCTCT |
| 5186 | RP11-309L24.4-001 | GGTGCCCAGCCCTGTCCCAG |
| 5187 | RP11-309L24.4-001 | GAAAAGGCGCCTCTGGGACA |
| 5188 | RP11-309L24.4-001 | CCCCAAGTCAGCCCCTTCCA |
| 5189 | RP11-309L24.4-001 | CAGGGCTGGGCACCTTGGAA |
| 5190 | RP11-309L24.4-001 | TGAAAAGGCGCCTCTGGGAC |
| 5191 | RP11-309L24.4-001 | CTTGGAAGGGGCTGACTTGG |
| 5192 | RP11-30K9.6-001 | CCAACAATGCCTCAGGCCGA |
| 5193 | RP11-30K9.6-001 | CGGCCTGAGGCATTGTTGGG |
| 5194 | RP11-30K9.6-001 | GGCTTCTTTAAAGGGAAGGC |
| 5195 | RP11-30K9.6-001 | CCCAACAATGCCTCAGGCCG |
| 5196 | RP11-30K9.6-001 | GTGCGGCTTCTTTAAAGGGA |
| 5197 | RP11-30K9.6-001 | GGTCGCGTGTGGCCCACCCT |
| 5198 | RP11-30K9.6-001 | ACACGCGACCCAGCGCTAAA |
| 5199 | RP11-313D6.3-001 | GTACTACAATGACAGGTGAG |
| 5200 | RP11-313D6.3-001 | ACAGGCACAATAAATGAACA |
| 5201 | RP11-313D6.3-001 | TACTACAATGACAGGTGAGT |
| 5202 | RP11-313D6.3-001 | CAGATACTGGCCACACCAGG |

| | | |
|---|---|---|
| 5203 | RP11-313D6.3-001 | CAAAGATTCTCAGCAGTTGG |
| 5204 | RP11-313D6.3-001 | GGTGAGTGGGTCAGGCACAG |
| 5205 | RP11-313D6.3-001 | ATCTTAAGTACTACAATGAC |
| 5206 | RP11-314D7.2-001 | TCACACATGTTGAAAACAGA |
| 5207 | RP11-314D7.2-001 | TGAAAACAGAAGGACTCGCC |
| 5208 | RP11-314D7.2-001 | TCAAAAGTATATTCACCAAG |
| 5209 | RP11-314D7.2-001 | GAAGGACTCGCCAGGCGCAG |
| 5210 | RP11-314D7.2-001 | AGCACTTTAGGAGGCCGAAG |
| 5211 | RP11-314D7.2-001 | TGTAATCCCAGCACTTTAGG |
| 5212 | RP11-314D7.2-001 | ATATTAGAAGAAAAACCTCT |
| 5213 | RP11-317N12.1-002 | CCTCACCCTCCAAACTTCAG |
| 5214 | RP11-317N12.1-002 | GAAAGTTCCCCAAAACTTTG |
| 5215 | RP11-317N12.1-002 | GTGTTCCCCTGAAGTTTGGA |
| 5216 | RP11-317N12.1-002 | GGGAAAGTTCCCCAAAACTT |
| 5217 | RP11-317N12.1-002 | AGAGGTTGAGAAGAATGAGC |
| 5218 | RP11-317N12.1-002 | TTTACTCTTTGCATGCCCTG |
| 5219 | RP11-317N12.1-002 | TCCTCACCCTCCAAACTTCA |
| 5220 | RP11-318E3.9-001 | CAATTTCCTAAAGCGCCCAC |
| 5221 | RP11-318E3.9-001 | GGGCGCTTTAGGAAATTGGT |
| 5222 | RP11-318E3.9-001 | TTGTTTAGTGCGTTGCCGGT |
| 5223 | RP11-318E3.9-001 | CTTGTTTAGTGCGTTGCCGG |
| 5224 | RP11-318E3.9-001 | ATCGGCTTCCCAATGAAACC |
| 5225 | RP11-318E3.9-001 | GCGTTGCCGGTGGGCGCTTT |
| 5226 | RP11-318E3.9-001 | CTCCTTGTTTAGTGCGTTGC |
| 5227 | RP11-318G21.4-001 | TTAGTGCTTGCATGTTGAGT |
| 5228 | RP11-318G21.4-001 | GCACTAATCCTCTAACATGC |
| 5229 | RP11-318G21.4-001 | TTTCACCTAACAAATAGACA |
| 5230 | RP11-318G21.4-001 | TTCACCTAACAAATAGACAT |
| 5231 | RP11-318G21.4-001 | CAGGATTTCTAAAGGCAGAA |
| 5232 | RP11-318G21.4-001 | AGAAACTCTCACATAGAGTT |
| 5233 | RP11-318G21.4-001 | CTGTTCACTGAAATATGCCT |
| 5234 | RP11-321F8.4-001 | TATTAACAGTAGTGACTACT |
| 5235 | RP11-321F8.4-001 | CTGGGGATCACCGAGATACA |
| 5236 | RP11-321F8.4-001 | GTATTAACAGTAGTGACTAC |
| 5237 | RP11-321F8.4-001 | ACAACTATGAAAACATCACA |
| 5238 | RP11-321F8.4-001 | GAGATACAGGGTATTAGGGA |
| 5239 | RP11-321F8.4-001 | ATCCCTAATACCCTGTATCT |
| 5240 | RP11-321F8.4-001 | ATTAACAGTAGTGACTACTG |
| 5241 | RP11-321L2.2-001 | TAAAAACGTTAGACAGACTC |
| 5242 | RP11-321L2.2-001 | AATTGAGTGATAAATATGTG |
| 5243 | RP11-321L2.2-001 | CAGAACATCAGCAAAACGCC |
| 5244 | RP11-321L2.2-001 | CTTGATGTAGATGCTTATAC |
| 5245 | RP11-321L2.2-001 | TAGACAGACTCCGGGCACAG |

| | | |
|---|---|---|
| 5246 | RP11-321L2.2-001 | AAAAACGTTAGACAGACTCC |
| 5247 | RP11-321L2.2-001 | TAAAATGTTGAAGCAATCTC |
| 5248 | RP11-322E11.2-001 | TGAATTATTCTTCTTATATG |
| 5249 | RP11-322E11.2-001 | TTGCAGAATTACAGGAGTAC |
| 5250 | RP11-322E11.2-001 | GGTGATTGCCAGGGACTGGG |
| 5251 | RP11-322E11.2-001 | ATTGCCAGGGACTGGGCGGA |
| 5252 | RP11-322E11.2-001 | AAGTAGAATGGTGATTGCCA |
| 5253 | RP11-322E11.2-001 | AATGGTGATTGCCAGGGACT |
| 5254 | RP11-322E11.2-001 | CTCCCCCTCCGCCCAGTCCC |
| 5255 | RP11-323P17.2-002 | TTATGTAGATGACTCCCAAG |
| 5256 | RP11-323P17.2-002 | CAGATCATGTCATATCACAC |
| 5257 | RP11-323P17.2-002 | TTTGGTATAAGTCTCCAACC |
| 5258 | RP11-323P17.2-002 | CTTATGTAGATGACTCCCAA |
| 5259 | RP11-323P17.2-002 | GCTTATGTAGATGACTCCCA |
| 5260 | RP11-323P17.2-002 | GCAGCAACAGTCAAACACAG |
| 5261 | RP11-323P17.2-002 | AGATCATGTCATATCACACA |
| 5262 | RP11-325F22.2-001 | AACTATAACACTCACGGCGA |
| 5263 | RP11-325F22.2-001 | GAACTATAACACTCACGGCG |
| 5264 | RP11-325F22.2-001 | GCACAGCTGCATATCCTCAG |
| 5265 | RP11-325F22.2-001 | AGTGAAGCCTATCAGTAAGG |
| 5266 | RP11-325F22.2-001 | ATATGCAGCTGTGCACCTGG |
| 5267 | RP11-325F22.2-001 | TGCACAGCTGCATATCCTCA |
| 5268 | RP11-325F22.2-001 | ACACTCACGGCGAGGGTCCA |
| 5269 | RP11-325F22.2-002 | TACAGCTCACAAAGGCAATG |
| 5270 | RP11-325F22.2-002 | TTTAAGAACTATAACACTCA |
| 5271 | RP11-325F22.2-002 | GAGTGTTACAGCTCCTAAGG |
| 5272 | RP11-325F22.2-002 | TTTGTGAGCTGTAACACTCA |
| 5273 | RP11-325F22.2-002 | GTGAGTGTTACAGCTCACAA |
| 5274 | RP11-325F22.2-002 | AACGAACAACTCCAGACGCG |
| 5275 | RP11-325F22.2-002 | CTGCAGCTTCACTCCTGAGC |
| 5276 | RP11-326C3.12-003 | GGGACCTCAGAGCAGAGCAG |
| 5277 | RP11-326C3.12-003 | GAAACAGGAGCCCCCACCTG |
| 5278 | RP11-326C3.12-003 | GGAAACAGGAGCCCCCACCT |
| 5279 | RP11-326C3.12-003 | GTTTCCCCTGCTCTGCTCTG |
| 5280 | RP11-326C3.12-003 | AGCCCCCACCTGGGGAGAGG |
| 5281 | RP11-326C3.12-003 | GGGGACCTCAGAGCAGAGCA |
| 5282 | RP11-326C3.12-003 | CAGAGCAGAGCAGGGGAAAC |
| 5283 | RP11-326C3.14-001 | CCCAGCACTCCCTTAACACA |
| 5284 | RP11-326C3.14-001 | TCCCAGCACTCCCTTAACAC |
| 5285 | RP11-326C3.14-001 | TTATCTAAGAAGTACAGTGA |
| 5286 | RP11-326C3.14-001 | TTATCCCATTACCTTCAGTG |
| 5287 | RP11-326C3.14-001 | TAAACTCATTCCAAAAACAA |
| 5288 | RP11-326C3.14-001 | TTATCCCATAACCCTCACTG |

| | | |
|---|---|---|
| 5289 | RP11-326C3.14-001 | CCCTGTGTTAAGGGAGTGCT |
| 5290 | RP11-326G21.1-001 | AATGATGATGGTCAATAGGG |
| 5291 | RP11-326G21.1-001 | ATAAAAGAAAGTCTGATGCA |
| 5292 | RP11-326G21.1-001 | TTGAAATCAGTGGTGTAATG |
| 5293 | RP11-326G21.1-001 | AAAAATGATGATGGTCAATA |
| 5294 | RP11-326G21.1-001 | GATGCAGGGAAAGGGGAAGT |
| 5295 | RP11-326G21.1-001 | AAAGTCTGATGCAGGGAAAG |
| 5296 | RP11-326G21.1-001 | AAAAAGGACTGGAAAGACAT |
| 5297 | RP11-326I11.3-001 | AGCAGCGGCTCGTGAGTCCG |
| 5298 | RP11-326I11.3-001 | GACGTTCTAGCCGCACGTGC |
| 5299 | RP11-326I11.3-001 | TTCGCTTCCCCAGGTCGCGG |
| 5300 | RP11-326I11.3-001 | GGACGTTCTAGCCGCACGTG |
| 5301 | RP11-326I11.3-001 | AGTCCGCGGCTCGCACCCAG |
| 5302 | RP11-326I11.3-001 | CCCAGGGGCGCCCGCACGTG |
| 5303 | RP11-326I11.3-001 | GCGGCGGGAGCAACAAGCAG |
| 5304 | RP11-326N17.2-001 | AGCCAAAAGAACAAAACTGG |
| 5305 | RP11-326N17.2-001 | TGACTTCAAACTATACTACA |
| 5306 | RP11-326N17.2-001 | CTACAGTAACCAAAACATCA |
| 5307 | RP11-326N17.2-001 | CCTGACAAAAACAAGCAATG |
| 5308 | RP11-326N17.2-001 | GATATGTGGCATTATTTCTG |
| 5309 | RP11-326N17.2-001 | AAAACAGAGATATAGATCAA |
| 5310 | RP11-326N17.2-001 | ATATGTGGCATTATTTCTGA |
| 5311 | RP11-332H14.2-001 | GAGAGAAGGAGCCCGGCATG |
| 5312 | RP11-332H14.2-001 | CACGGACCAGAGAAAAGAGG |
| 5313 | RP11-332H14.2-001 | CACTCTGAGTCGCCGCATGC |
| 5314 | RP11-332H14.2-001 | TCCCACGGACCAGAGAAAAG |
| 5315 | RP11-332H14.2-001 | CCCGCTTGCAAAAGCTCCCA |
| 5316 | RP11-332H14.2-001 | CTCGCTGCTGGTTCGGGTGT |
| 5317 | RP11-332H14.2-001 | CGCTGCTGGTTCGGGTGTTG |
| 5318 | RP11-338H14.1-001 | CAGACAGGGCGGTTGCCAGG |
| 5319 | RP11-338H14.1-001 | GCCCGGCAGCCACCCCATCA |
| 5320 | RP11-338H14.1-001 | TCACCTCCCAGACAGGGTCG |
| 5321 | RP11-338H14.1-001 | CAGCCGCGACCCTGTCTGGG |
| 5322 | RP11-338H14.1-001 | CAGGGCGGTTGCCAGGCGGA |
| 5323 | RP11-338H14.1-001 | GCTGACCCCCATCTCCCTCC |
| 5324 | RP11-338H14.1-001 | CCCACATCTCAGACGATGGG |
| 5325 | RP11-33B1.3-001 | GGTGGACCATCCGAAGTCAG |
| 5326 | RP11-33B1.3-001 | GAAGGATTCCTTGAGCCAGA |
| 5327 | RP11-33B1.3-001 | ATGAGCAAAATCGCCAGGCG |
| 5328 | RP11-33B1.3-001 | AGCTACCGGGGACGCTGACG |
| 5329 | RP11-33B1.3-001 | TTCCTTGAGCCAGAAGGTCA |
| 5330 | RP11-33B1.3-001 | AGCCTTGACCTTCTGGCTCA |
| 5331 | RP11-33B1.3-001 | GCAAAATCGCCAGGCGCGGT |

| | | |
|---|---|---|
| 5332 | RP11-346D14.1-001 | AAATAATGGAATCTAGCCTG |
| 5333 | RP11-346D14.1-001 | AGCCCTACACTTACCAAGCG |
| 5334 | RP11-346D14.1-001 | AATGATCTTAGATCCACGCT |
| 5335 | RP11-346D14.1-001 | GAAATAATGGAATCTAGCCT |
| 5336 | RP11-346D14.1-001 | GATCCACGCTTGGTAAGTGT |
| 5337 | RP11-346D14.1-001 | AAGCGTGGATCTAAGATCAT |
| 5338 | RP11-346D14.1-001 | AGATCATTGGAAATGCTGAA |
| 5339 | RP11-347C12.10-001 | TTCAAGTCACCCTTCCCCCA |
| 5340 | RP11-347C12.10-001 | GTCCGTCATTCAAGCAGTAG |
| 5341 | RP11-347C12.10-001 | AGAATTCCTGGACTCTGACT |
| 5342 | RP11-347C12.10-001 | AGTGACGTCAGTCACGTGGG |
| 5343 | RP11-347C12.10-001 | TCAGTCACGTGGGAGGACAA |
| 5344 | RP11-347C12.10-001 | GGAGGACAATGGTGAGAAGA |
| 5345 | RP11-347C12.10-001 | CTCACGCCCAGTCAGAGTCC |
| 5346 | RP11-347I19.8-001 | GATCCAATCAGTGAGCTGTT |
| 5347 | RP11-347I19.8-001 | AGGGGTACAGTCCAGCCAGG |
| 5348 | RP11-347I19.8-001 | GGGGTACAGTCCAGCCAGGT |
| 5349 | RP11-347I19.8-001 | GCACCCAACAGCTCACTGAT |
| 5350 | RP11-347I19.8-001 | CAGCTCACTGATTGGATCCA |
| 5351 | RP11-347I19.8-001 | AGCTGTTGGGTGCTCAGCTG |
| 5352 | RP11-347I19.8-001 | ACAGCTCACTGATTGGATCC |
| 5353 | RP11-348P10.2-001 | ACTAACCTAAGTAACAATAG |
| 5354 | RP11-348P10.2-001 | AAGAAATTGGGGGTTAACTG |
| 5355 | RP11-348P10.2-001 | AGATCATGGCTTGTTCAGAG |
| 5356 | RP11-348P10.2-001 | TTTCTGAGATGCACATGCAT |
| 5357 | RP11-348P10.2-001 | ATAGGAAAGAAAGAAATTGG |
| 5358 | RP11-348P10.2-001 | GAAGTGTGAAATGGGTAGTG |
| 5359 | RP11-348P10.2-001 | TTGTAGAAAATGTAGACTGC |
| 5360 | RP11-34P13.13-001 | CCCAGAACCTGCCAATGCTG |
| 5361 | RP11-34P13.13-001 | ACTGAGAGTACAGAAGATAA |
| 5362 | RP11-34P13.13-001 | AGTACAGAAGATAAAGGAAA |
| 5363 | RP11-34P13.13-001 | AGTTACCACATGCTCTGCAG |
| 5364 | RP11-34P13.13-001 | TCCCAGAACCTGCCAATGCT |
| 5365 | RP11-34P13.13-001 | TGCTGGGGAACGGAAAATGA |
| 5366 | RP11-34P13.13-001 | TTGGCAGGTTCTGGGACTGG |
| 5367 | RP11-34P13.13-002 | ACTTGTATCTGTAAGGTCGG |
| 5368 | RP11-34P13.13-002 | TAAACTTGTATCTGTAAGGT |
| 5369 | RP11-34P13.13-002 | ACAAAAACACAAGGTTAAGA |
| 5370 | RP11-34P13.13-002 | AACTTGTATCTGTAAGGTCG |
| 5371 | RP11-34P13.13-002 | TCTGTCATCCAGGCTGGCAG |
| 5372 | RP11-34P13.13-002 | AATAAGACTTTACAGCAGCC |
| 5373 | RP11-34P13.13-002 | TCCCAGCACTTTGGCAGCAG |
| 5374 | RP11-350N15.5-001 | AGCTAGCGTCTGGACGACTA |

| | | |
|---|---|---|
| 5375 | RP11-350N15.5-001 | AGCCCCAGCGTCCCGAACCA |
| 5376 | RP11-350N15.5-001 | GCGCGAACCAGGAACGCCCG |
| 5377 | RP11-350N15.5-001 | GTCCAGACGCTAGCTGCCGC |
| 5378 | RP11-350N15.5-001 | CAGCTAGCGTCTGGACGACT |
| 5379 | RP11-350N15.5-001 | CTCCGGAGGAAGCGGCGAGT |
| 5380 | RP11-350N15.5-001 | GAGGAAGCGGCGAGTCGGCG |
| 5381 | RP11-351I21.11-001 | ATAACGACCCTCAAAAGCGG |
| 5382 | RP11-351I21.11-001 | AGCAGGATTGCATACCTAGG |
| 5383 | RP11-351I21.11-001 | AAGATAACGACCCTCAAAAG |
| 5384 | RP11-351I21.11-001 | TAGGCGGCCTCGGAACTCAG |
| 5385 | RP11-351I21.11-001 | ATTTGCTGAGACACACCAAA |
| 5386 | RP11-351I21.11-001 | TCAGGGGGAGAACCATGGAA |
| 5387 | RP11-351I21.11-001 | CTCAGCAGGATTGCATACCT |
| 5388 | RP11-352M15.2-001 | TTCCCCTAAGAGGACCGATG |
| 5389 | RP11-352M15.2-001 | TCCCCTAAGAGGACCGATGG |
| 5390 | RP11-352M15.2-001 | TCCCCCATCGGTCCTCTTAG |
| 5391 | RP11-352M15.2-001 | CTTCCCCTAAGAGGACCGAT |
| 5392 | RP11-352M15.2-001 | ACTTCCCCTAAGAGGACCGA |
| 5393 | RP11-352M15.2-001 | ACATCTCCACTTCCCCTAAG |
| 5394 | RP11-352M15.2-001 | TAAGAGGACCGATGGGGGAA |
| 5395 | RP11-356C4.3-001 | ACTGTCCGAACAGTGAACTT |
| 5396 | RP11-356C4.3-001 | ACTTCGGAGATCCACCGGAG |
| 5397 | RP11-356C4.3-001 | GCAGGATGCTCAGGTGCTCG |
| 5398 | RP11-356C4.3-001 | AGTGAACTTCGGAGATCCAC |
| 5399 | RP11-356C4.3-001 | GAACTTCGGAGATCCACCGG |
| 5400 | RP11-356C4.3-001 | AACTTCGGAGATCCACCGGA |
| 5401 | RP11-356C4.3-001 | AAACCAGCTGAAGCCGGGCG |
| 5402 | RP11-360F5.3-001 | TAGAATATGAGGCCTCCGAG |
| 5403 | RP11-360F5.3-001 | AGGCCTCATATTCTAGGTGA |
| 5404 | RP11-360F5.3-001 | ACATTTCCAACCTGTGCATG |
| 5405 | RP11-360F5.3-001 | CTTCCGTCACCTAGAATATG |
| 5406 | RP11-360F5.3-001 | CATTTCCAACCTGTGCATGG |
| 5407 | RP11-360F5.3-001 | TAATTGAAAGAAGAGAGCAT |
| 5408 | RP11-360F5.3-001 | CTCAGGGTGGCCCTGACATG |
| 5409 | RP11-364P22.2-001 | GCTGAACTCCATCCTCCCGG |
| 5410 | RP11-364P22.2-001 | GAGGAGCGGACGGCCCCGCG |
| 5411 | RP11-364P22.2-001 | GCCGGGAGGATGGAGTTCAG |
| 5412 | RP11-364P22.2-001 | CCATCCTCCCGGCGGTCGGG |
| 5413 | RP11-364P22.2-001 | GCCGCCGCCCGACCGCCGGG |
| 5414 | RP11-364P22.2-001 | TGGAGGAGCGGACGGCCCCG |
| 5415 | RP11-364P22.2-001 | CCCGCTGAACTCCATCCTCC |
| 5416 | RP11-370F5.4-001 | ATGGAGGATGTGTACCAACA |
| 5417 | RP11-370F5.4-001 | TTGTAAACACTCTTCCATGT |

| | | |
|---|---|---|
| 5418 | RP11-370F5.4-001 | CACACTGTAAATGTTAGTCA |
| 5419 | RP11-370F5.4-001 | ACTGTAAATGTTAGTCATGG |
| 5420 | RP11-370F5.4-001 | GATTACCATACATCTGGGTG |
| 5421 | RP11-370F5.4-001 | ATAATTCCAGTATGCATACT |
| 5422 | RP11-370F5.4-001 | GAGACATAAGGTGAGTTTGT |
| 5423 | RP11-378J18.9-001 | AAGGGAATCTTCAGCTTAGG |
| 5424 | RP11-378J18.9-001 | TCAGAGCTACTCACAATGAA |
| 5425 | RP11-378J18.9-001 | CCACTCAGCACCCATAGTCC |
| 5426 | RP11-378J18.9-001 | CTGGACTATGGGTGCTGAGT |
| 5427 | RP11-378J18.9-001 | AGGACAGTACTTGTTCTCAA |
| 5428 | RP11-378J18.9-001 | GGTCCCCATGCCTGGACTAT |
| 5429 | RP11-378J18.9-001 | TTGTCGCAGGTCAGGTGAGG |
| 5430 | RP11-378J18.9-004 | TGAGAACAAGTACTGTCCTG |
| 5431 | RP11-378J18.9-004 | AAGAAAGGCCATGAAACCGC |
| 5432 | RP11-378J18.9-004 | GCACCCATAGTCCAGGCATG |
| 5433 | RP11-378J18.9-004 | TAAATGACATCTTATCAAGA |
| 5434 | RP11-378J18.9-004 | CATTTGTGGTTCTTGCGAAG |
| 5435 | RP11-378J18.9-004 | AGCACCCATAGTCCAGGCAT |
| 5436 | RP11-378J18.9-004 | TCTGGGGCTCAGGTGTGTTG |
| 5437 | RP11-378J18.9-005 | ACTTGTGGGGAACACTTAGG |
| 5438 | RP11-378J18.9-005 | AAGACTTGTGGGGAACACTT |
| 5439 | RP11-378J18.9-005 | ACTTAGGAGGGAAAGTTACT |
| 5440 | RP11-378J18.9-005 | AGTGCATGCATCAAGTGCAA |
| 5441 | RP11-378J18.9-005 | CTTGTGGGGAACACTTAGGA |
| 5442 | RP11-378J18.9-005 | ACAGAATTCAAAGACTTGTG |
| 5443 | RP11-378J18.9-005 | AACAGAATTCAAAGACTTGT |
| 5444 | RP11-379B18.5-017 | TCACAGCCGAATTCTACCAG |
| 5445 | RP11-379B18.5-017 | GCTTATTGACCATGATCACG |
| 5446 | RP11-379B18.5-017 | ATTCTACCAGAGGTATAAGG |
| 5447 | RP11-379B18.5-017 | AAAAAGAGTCCAGGACCAGA |
| 5448 | RP11-379B18.5-017 | CCTCATAAAACAAGTTAGGG |
| 5449 | RP11-379B18.5-017 | TTGTTTGGAATAGTTTCAGA |
| 5450 | RP11-379B18.5-017 | AATTCGGCTGTGAATCCATC |
| 5451 | RP11-37B2.1-005 | GAGGCGAGAGCCAGGCGGGA |
| 5452 | RP11-37B2.1-005 | CTGGCTCTCGCCTCGGCCTG |
| 5453 | RP11-37B2.1-005 | AGGCCGAGGCGAGAGCCAGG |
| 5454 | RP11-37B2.1-005 | GGAGAGGGGCAGCCCGCTCG |
| 5455 | RP11-37B2.1-005 | GCGACCCGGGCGGAGGATCC |
| 5456 | RP11-37B2.1-005 | GGCCGAGGCGAGAGCCAGGC |
| 5457 | RP11-37B2.1-005 | AGGATCCCGGCCGCAGGCCG |
| 5458 | RP11-37B2.1-007 | AGTGTACAACCTCTTAAGTG |
| 5459 | RP11-37B2.1-007 | CCTGCGCAGGAAGCGATTTG |
| 5460 | RP11-37B2.1-007 | GGGGAGTGTTGTGAATTCCG |

| | | |
|---|---|---|
| 5461 | RP11-37B2.1-007 | GGCTTTATTAGGGGGCACCG |
| 5462 | RP11-37B2.1-007 | GCGCAGGCGCGGAGTCGGAA |
| 5463 | RP11-37B2.1-007 | CCTCTTAAGTGTGGAGCCCT |
| 5464 | RP11-37B2.1-007 | GGGAGTGTTGTGAATTCCGA |
| 5465 | RP11-381N20.2-001 | TTAAAGAAGAAAACTCGACC |
| 5466 | RP11-381N20.2-001 | ATTAAAGAAGAAAACTCGAC |
| 5467 | RP11-381N20.2-001 | GAAAACTCGACCGGGCGCGG |
| 5468 | RP11-381N20.2-001 | TGTAATCCCAACACTTTGGG |
| 5469 | RP11-381N20.2-001 | GAAGAAAACTCGACCGGGCG |
| 5470 | RP11-381N20.2-001 | GCCAAGGCAGGCGGATCACG |
| 5471 | RP11-381N20.2-001 | TTCTTCTTTAATCTGGTATG |
| 5472 | RP11-383M4.6-001 | CCAACTGGCAGCGCGCCTCG |
| 5473 | RP11-383M4.6-001 | GCCAGTTGGGATGAAAATGG |
| 5474 | RP11-383M4.6-001 | TTGGGGGCTCCCAGGCCGCG |
| 5475 | RP11-383M4.6-001 | TGGGATGAAAATGGCGGATG |
| 5476 | RP11-383M4.6-001 | GGGCTGTTGAGATCGGTTGG |
| 5477 | RP11-383M4.6-001 | CTGCCCAGGCTGGAGTGCAG |
| 5478 | RP11-383M4.6-001 | GCTGCCAGTTGGGATGAAAA |
| 5479 | RP11-385D13.3-001 | ATAAGAAAGGATGAGCATGG |
| 5480 | RP11-385D13.3-001 | ACTATAAGAAAGGATGAGCA |
| 5481 | RP11-385D13.3-001 | CAGAAACTACAGAGGTCAGA |
| 5482 | RP11-385D13.3-001 | GGAGTGCAGTGACACAATCA |
| 5483 | RP11-385D13.3-001 | ATCCTTCAAAACTATAAGAA |
| 5484 | RP11-385D13.3-001 | AAAAAATACAACAAACAGCT |
| 5485 | RP11-385D13.3-001 | GCAGGAGGATCACTTGAGTC |
| 5486 | RP11-385F7.1-001 | CCAAGCTGTTCCGAGCCTCC |
| 5487 | RP11-385F7.1-001 | CGTCGCCGCCGCGGAAAGTG |
| 5488 | RP11-385F7.1-001 | TGGGCTCGCTCTCCCCCCGG |
| 5489 | RP11-385F7.1-001 | GTCGCCTCAGATGGATCAAA |
| 5490 | RP11-385F7.1-001 | AAGCTGTTCCGAGCCTCCGG |
| 5491 | RP11-385F7.1-001 | AGCTGTTCCGAGCCTCCGGG |
| 5492 | RP11-385F7.1-001 | CAAGCTGTTCCGAGCCTCCG |
| 5493 | RP11-38L15.2-002 | CGTCCCCACGGCTCTCCTAG |
| 5494 | RP11-38L15.2-002 | TCCCCACGGCTCTCCTAGCG |
| 5495 | RP11-38L15.2-002 | GCTGACTGACAAACTTCCCG |
| 5496 | RP11-38L15.2-002 | GGGACGGAGCCTGCAGACGG |
| 5497 | RP11-38L15.2-002 | CCCCACAGTGCCCAGACCCC |
| 5498 | RP11-38L15.2-002 | CAGACGGCGGTGTCCCCGCT |
| 5499 | RP11-38L15.2-002 | TCCCCACAGTGCCCAGACCC |
| 5500 | RP11-38L15.2-005 | CTCTGAATCTTCAATACACA |
| 5501 | RP11-38L15.2-005 | TATCCTGACATGGTAAAGCA |
| 5502 | RP11-38L15.2-005 | GATCACCTATGGGATGGTTG |
| 5503 | RP11-38L15.2-005 | CGCTGACTGCCAACAAGGAG |

| | | |
|---|---|---|
| 5504 | RP11-38L15.2-005 | CTGCCAACAAGGAGTGGAAC |
| 5505 | RP11-38L15.2-005 | TTCTGCGCTGACTGCCAACA |
| 5506 | RP11-38L15.2-005 | AGGCCAGAGGAAGAAATTGC |
| 5507 | RP11-390P24.1-001 | GTGATGGGACTAATGTGGAG |
| 5508 | RP11-390P24.1-001 | CTCCAGCCATCCAGTCCTCA |
| 5509 | RP11-390P24.1-001 | GACTAATGTGGAGAGGCCTG |
| 5510 | RP11-390P24.1-001 | TGACAGAGGAGGCTAAGCCC |
| 5511 | RP11-390P24.1-001 | GGAGCCTGAGTTGACAGAGG |
| 5512 | RP11-390P24.1-001 | GCCATCCAGTCCTCAGGGAA |
| 5513 | RP11-390P24.1-001 | TTAGCCTCCTCTGTCAACTC |
| 5514 | RP11-395A13.2-001 | GCTGGGTGGCTAACGCTCGA |
| 5515 | RP11-395A13.2-001 | AAGGCTGGTATCCAAGGCTG |
| 5516 | RP11-395A13.2-001 | CTGGTATCCAAGGCTGCGGC |
| 5517 | RP11-395A13.2-001 | AGGCTGCGGCCGGTCAGGCG |
| 5518 | RP11-395A13.2-001 | ACCCAGCATTGAGCTGCCAG |
| 5519 | RP11-395A13.2-001 | TGCTTAGGGAGAACAGCCGC |
| 5520 | RP11-395A13.2-001 | GCTGGCAGCTCAATGCTGGG |
| 5521 | RP11-395G23.3-001 | GCTCTTGGTAGCTTCCACGC |
| 5522 | RP11-395G23.3-001 | GGCTCTTGGTAGCTTCCACG |
| 5523 | RP11-395G23.3-001 | AGTTTCCGGACCTGCCCGCG |
| 5524 | RP11-395G23.3-001 | GCATGTCTGCAAGCGCTCAG |
| 5525 | RP11-395G23.3-001 | TTGGTAGCTTCCACGCGGGC |
| 5526 | RP11-395G23.3-001 | AGCTTCCACGCGGGCAGGTC |
| 5527 | RP11-395G23.3-001 | CTGCAAGCGCTCAGCGGCGC |
| 5528 | RP11-395G23.3-002 | GGCTGAGCCCATTCACCTCG |
| 5529 | RP11-395G23.3-002 | CGGCCACAGGAGCTCAGCGC |
| 5530 | RP11-395G23.3-002 | CCCATTCACCTCGCGGCCAC |
| 5531 | RP11-395G23.3-002 | CGAGTGCGCCCCTCTCGGCG |
| 5532 | RP11-395G23.3-002 | GGCGAGTGCGCCCCTCTCGG |
| 5533 | RP11-395G23.3-002 | GCGCCGGCGCTGAGCTCCTG |
| 5534 | RP11-395G23.3-002 | GGCGGCGAGTGCGCCCCTCT |
| 5535 | RP11-395I14.2-001 | TATTAACTCACACGATCCCA |
| 5536 | RP11-395I14.2-001 | GTCTATAAAGGTGTCGCCAA |
| 5537 | RP11-395I14.2-001 | CGATCCCAAGGTCCCACAGT |
| 5538 | RP11-395I14.2-001 | ACTTTGAGTCCAATGTTCGA |
| 5539 | RP11-395I14.2-001 | ACACCTTTATAGACACACCC |
| 5540 | RP11-395I14.2-001 | AGTAGGCCGTCCGCAAGCTG |
| 5541 | RP11-395I14.2-001 | CCAGCACAGGAGAAGGATGT |
| 5542 | RP11-396N11.1-001 | GACAACATGAATGCCCCTGG |
| 5543 | RP11-396N11.1-001 | AAAAATGTTGGCGGGCACAG |
| 5544 | RP11-396N11.1-001 | TGTAGCTCCAACTACTCGGA |
| 5545 | RP11-396N11.1-001 | GTAGAATCTAAAAATGTTGG |
| 5546 | RP11-396N11.1-001 | TTAAACATAATGTCCTCCAG |

| | | |
|---|---|---|
| 5547 | RP11-396N11.1-001 | AAGGTTGAGGCAGAATGGCG |
| 5548 | RP11-396N11.1-001 | TAGAATCTAAAAATGTTGGC |
| 5549 | RP11-397A16.1-003 | TGAGTCAAAGAAAGGAGTGA |
| 5550 | RP11-397A16.1-003 | CTCGTAGGGTCCTATGCCCA |
| 5551 | RP11-397A16.1-003 | GGAAAAGCGCAGTATTCTGG |
| 5552 | RP11-397A16.1-003 | AGCAATCAGCGAGACTCCCT |
| 5553 | RP11-397A16.1-003 | GCCTGGAAGCTGGAACTGGG |
| 5554 | RP11-397A16.1-003 | GAAAAGCGCAGTATTCTGGT |
| 5555 | RP11-397A16.1-003 | AGGACCCTACGAGCCAGGTG |
| 5556 | RP11-398H6.1-001 | CAAGTCGACAAGAGATCCCG |
| 5557 | RP11-398H6.1-001 | GAGCGCACAAGCTTACCGTA |
| 5558 | RP11-398H6.1-001 | CCTACATCTGGTGCCCAACG |
| 5559 | RP11-398H6.1-001 | AGGCTGACTGTAGACGTCCT |
| 5560 | RP11-398H6.1-001 | CCTACAAGAAAACCATGGCC |
| 5561 | RP11-398H6.1-001 | AAGAAAACCATGGCCGGGCG |
| 5562 | RP11-398H6.1-001 | GACGTCTACAGTCAGCCTTA |
| 5563 | RP11-3B12.3-001 | AGGTCCCCCTATCCAAGTCA |
| 5564 | RP11-3B12.3-001 | GGTGCCGTGACTTGGATAGG |
| 5565 | RP11-3B12.3-001 | CTGGTCGGTCTGAGGACGTG |
| 5566 | RP11-3B12.3-001 | TGAGGACGTGAGGTCGTAGG |
| 5567 | RP11-3B12.3-001 | GTCTGAGGACGTGAGGTCGT |
| 5568 | RP11-3B12.3-001 | TGGTGCCGTGACTTGGATAG |
| 5569 | RP11-3B12.3-001 | GCTGACTCTTCGGACTCAGC |
| 5570 | RP11-3P17.5-001 | GAAGCTCTATGACATTGATG |
| 5571 | RP11-3P17.5-001 | CCGAATCAGGATGTTGACCT |
| 5572 | RP11-3P17.5-001 | CCAAGGTCAACATCCTGATT |
| 5573 | RP11-3P17.5-001 | CTATGACATTGATGTGGCCA |
| 5574 | RP11-3P17.5-001 | GAAGAAGGCATATGTTCGAC |
| 5575 | RP11-3P17.5-001 | TATATTCAGAATTAGGCAGC |
| 5576 | RP11-3P17.5-001 | ATGGCAGACTCAGTGGTCAG |
| 5577 | RP11-403I13.8-002 | CGCCGGGACCAACCGCCGTG |
| 5578 | RP11-403I13.8-002 | AGGAAACTGGAATCCAGCAG |
| 5579 | RP11-403I13.8-002 | CACGGCGGTTGGTCCCGGCG |
| 5580 | RP11-403I13.8-002 | GCCGGGACCAACCGCCGTGG |
| 5581 | RP11-403I13.8-002 | ATCACCAATCCGTCGCCAGA |
| 5582 | RP11-403I13.8-002 | CACCAATCCGTCGCCAGAAG |
| 5583 | RP11-403I13.8-002 | CCCACGGCGGTTGGTCCCGG |
| 5584 | RP11-410L14.2-001 | AAACACTCGAGATCCGGAAG |
| 5585 | RP11-410L14.2-001 | CCCAAGACGGACCTCCACAT |
| 5586 | RP11-410L14.2-001 | CCAAGACGGACCTCCACATC |
| 5587 | RP11-410L14.2-001 | GATCCGGAAGTGGCGCTCCG |
| 5588 | RP11-410L14.2-001 | TCCACATCGGGAACCGCCGC |
| 5589 | RP11-410L14.2-001 | AGCCGCAGCCACCTTCTCCC |

| | | |
|---|---|---|
| 5590 | RP11-410L14.2-001 | GGGAACCGCCGCCGGCGCAG |
| 5591 | RP11-411H5.1-002 | AGACACAGTGTTCCATCCAG |
| 5592 | RP11-411H5.1-002 | CTGTGTCTTCTCATGGATGA |
| 5593 | RP11-411H5.1-002 | ATCCAGAGGATGTAGCATCA |
| 5594 | RP11-411H5.1-002 | AAATCTTAGGATATTGTCTT |
| 5595 | RP11-411H5.1-002 | AAGATCAATGCACTGGCATC |
| 5596 | RP11-411H5.1-002 | TTGATGCTACATCCTCTGGA |
| 5597 | RP11-411H5.1-002 | GAAGTCCAAGATCAATGCAC |
| 5598 | RP11-428O18.6-001 | AAGGCAGCTTCCGGCGAAGG |
| 5599 | RP11-428O18.6-001 | GCGTATGTTTAGACGTGTGG |
| 5600 | RP11-428O18.6-001 | GGTGCGTATGTTTAGACGTG |
| 5601 | RP11-428O18.6-001 | AAACATACGCACCTGCTAGT |
| 5602 | RP11-428O18.6-001 | GCCACATTGCTTTCCCAAAG |
| 5603 | RP11-428O18.6-001 | ACATTTGCGCACCTACTAGC |
| 5604 | RP11-428O18.6-001 | GCAGGGAAAACTGCGGCTGG |
| 5605 | RP11-42O15.3-001 | TCTTCCTCATAGGAAAAATG |
| 5606 | RP11-42O15.3-001 | GACTTTCTAAGTTCAAAACC |
| 5607 | RP11-42O15.3-001 | CGAGGCGTGTGGATCACCTG |
| 5608 | RP11-42O15.3-001 | CTATTAAAGGCCAGGTGCTG |
| 5609 | RP11-42O15.3-001 | CCTCGACCTCCCAAAGTGCT |
| 5610 | RP11-42O15.3-001 | TAAGTTTAAGGTCAGCTAGC |
| 5611 | RP11-42O15.3-001 | ATGAAAGCCTATTAAAGGCC |
| 5612 | RP11-430B1.2-002 | CTAAGCCACAGTTTGCCCGC |
| 5613 | RP11-430B1.2-002 | AACTGTGGCTTAGGGATGCT |
| 5614 | RP11-430B1.2-002 | CCGGAGGAGAGAGGCTGACG |
| 5615 | RP11-430B1.2-002 | GTCACACTCCCTGCCTCTCC |
| 5616 | RP11-430B1.2-002 | ACACTCCCTGCCTCTCCCGG |
| 5617 | RP11-430B1.2-002 | GATGCTGGGTGATTTGCCCG |
| 5618 | RP11-430B1.2-002 | CTGCCTCTCCCGGAGGAGAG |
| 5619 | RP11-430C7.5-001 | GTGCACTGGGACATGATACA |
| 5620 | RP11-430C7.5-001 | ATGCGTGTGTATAAAATAGG |
| 5621 | RP11-430C7.5-001 | TCATGAAAACAATTTAACGT |
| 5622 | RP11-430C7.5-001 | AATGCGTGTGTATAAAATAG |
| 5623 | RP11-430C7.5-001 | ATATCATACTGCATTGTATG |
| 5624 | RP11-430C7.5-001 | GTCATGAAAACAATTTAACG |
| 5625 | RP11-430C7.5-001 | GCTTTCTGTTAGCATCCCAG |
| 5626 | RP11-434D9.2-001 | AGGTGTGGGATGTAATTTCG |
| 5627 | RP11-434D9.2-001 | ACGAAATTACATCCCACACC |
| 5628 | RP11-434D9.2-001 | GAGCCCACCACAGCTCAGGG |
| 5629 | RP11-434D9.2-001 | CCTCATTTGGGAAGCGCAAG |
| 5630 | RP11-434D9.2-001 | CAAACTGCAAGGCAGCAGCG |
| 5631 | RP11-434D9.2-001 | ATTACATCCCACACCTGGCT |
| 5632 | RP11-434D9.2-001 | GCTGGGAAGCTCCAACTGGG |

| | | |
|---|---|---|
| 5633 | RP11-434H6.7-001 | CATGTAAGGTAACGTACCAA |
| 5634 | RP11-434H6.7-001 | GGCACTTACATGGATGCTAG |
| 5635 | RP11-434H6.7-001 | GAGGCACTTACATGGATGCT |
| 5636 | RP11-434H6.7-001 | ATCCTAATCTCTGAAACCGT |
| 5637 | RP11-434H6.7-001 | CAGTTAAGAAGCTTGAGATA |
| 5638 | RP11-434H6.7-001 | TAAGAAGCTTGAGATAGGGC |
| 5639 | RP11-434H6.7-001 | AGGCACTTACATGGATGCTA |
| 5640 | RP11-439L18.2-001 | ATCAGCAGACTGTCTCTCGA |
| 5641 | RP11-439L18.2-001 | CTTGTTAGATCTCATGTCAG |
| 5642 | RP11-439L18.2-001 | CAGCTGTTTACAGCGTAAGT |
| 5643 | RP11-439L18.2-001 | GGAAGCAGTTCATGGAACCC |
| 5644 | RP11-439L18.2-001 | GGATTGGCCAAATCACTGTC |
| 5645 | RP11-439L18.2-001 | AATCACTGTCAGGTAAATGA |
| 5646 | RP11-439L18.2-001 | AAATCACTGTCAGGTAAATG |
| 5647 | RP11-443N24.3-001 | CAGGCATTTCAAGAGACAAG |
| 5648 | RP11-443N24.3-001 | TTTGGTTGATATCTTCTCAG |
| 5649 | RP11-443N24.3-001 | AAAGTACAATTCCACCCATC |
| 5650 | RP11-443N24.3-001 | CATTTCAAGAGACAAGAGGC |
| 5651 | RP11-443N24.3-001 | AGTCTGTCATTTCTAACATG |
| 5652 | RP11-443N24.3-001 | CCAGATCACCTCTGCAGCAC |
| 5653 | RP11-443N24.3-001 | TGGGAGCTCTGCCTGATGGG |
| 5654 | RP11-446N19.1-001 | CTCAATGAGCTATGGACCTG |
| 5655 | RP11-446N19.1-001 | CCACAAAGACGTTTCATGTG |
| 5656 | RP11-446N19.1-001 | GGCTCAGACTCAATGAGCTA |
| 5657 | RP11-446N19.1-001 | CCTCACATGAAACGTCTTTG |
| 5658 | RP11-446N19.1-001 | CACAAAGACGTTTCATGTGA |
| 5659 | RP11-446N19.1-001 | ATAAAGTTCCAACAGGTGCT |
| 5660 | RP11-446N19.1-001 | AGAAAAATCCTGCTGAATCC |
| 5661 | RP11-44F14.2-002 | GTCTAGAAGTACCTAACAGT |
| 5662 | RP11-44F14.2-002 | CAAGGGCTTAGGTAACAGGG |
| 5663 | RP11-44F14.2-002 | TTCCCCTAAGACATAAATAC |
| 5664 | RP11-44F14.2-002 | AAACAAGGGCTTAGGTAACA |
| 5665 | RP11-44F14.2-002 | TTTGATAAGCACCTACTGTT |
| 5666 | RP11-44F14.2-002 | TCTGGTGCCTTGGTCACTCG |
| 5667 | RP11-44F14.2-002 | TCACTTTATTCCTGAAACCC |
| 5668 | RP11-44L9.1-001 | AATGAGAACGCGGAGCTACG |
| 5669 | RP11-44L9.1-001 | GGCCTATTTGTTCAGCGTCG |
| 5670 | RP11-44L9.1-001 | AGGCCCTTGTAGGCAGCTGA |
| 5671 | RP11-44L9.1-001 | GAGAGTTCCTCAAGAGAGCC |
| 5672 | RP11-44L9.1-001 | AGAGAGTTCCTCAAGAGAGC |
| 5673 | RP11-44L9.1-001 | TTGGTCCCTACCCTGGCTCC |
| 5674 | RP11-44L9.1-001 | AGGGTAGGGACCAAAAGGGC |
| 5675 | RP11-44N12.2-001 | TAACACCATCTAGGTAGCCT |

| | | |
|---|---|---|
| 5676 | RP11-44N12.2-001 | CAAAATGATTATGGTAACCT |
| 5677 | RP11-44N12.2-001 | GGTAACCTAGGCTACCTAGA |
| 5678 | RP11-44N12.2-001 | TGAATGTCATAACACCATCT |
| 5679 | RP11-44N12.2-001 | GAGCTTTGAAGACAGATATC |
| 5680 | RP11-44N12.2-001 | ATGGCCTCAAAGAACATTCA |
| 5681 | RP11-44N12.2-001 | AAAAAGAAATGGTCGCTTGT |
| 5682 | RP11-452C8.1-001 | ATGGAGTAGCTGGGAAACGT |
| 5683 | RP11-452C8.1-001 | AGAACATAACAGAAAGCCTG |
| 5684 | RP11-452C8.1-001 | GTTCTGATGATGGTAAGCAG |
| 5685 | RP11-452C8.1-001 | ACTCCATCTCCCCACACCGC |
| 5686 | RP11-452C8.1-001 | ATAACAGAAAGCCTGCGGTG |
| 5687 | RP11-452C8.1-001 | TTGCTCTGATTTGTGCCATG |
| 5688 | RP11-452C8.1-001 | AACAGAAAGCCTGCGGTGTG |
| 5689 | RP11-452F19.3-007 | AAAATGTAATGAGATTACTG |
| 5690 | RP11-452F19.3-007 | GAAATATGTTCTCAGATACT |
| 5691 | RP11-452F19.3-007 | GAGATTACTGAGGACCTGGA |
| 5692 | RP11-452F19.3-007 | GAGATGGGAAACTGCAACTA |
| 5693 | RP11-452F19.3-007 | TGAGATTACTGAGGACCTGG |
| 5694 | RP11-452F19.3-007 | TAATGAGATTACTGAGGACC |
| 5695 | RP11-452F19.3-007 | TCCCTATAGGAGTTGAGAAG |
| 5696 | RP11-452L6.1-001 | TGTGGGATGTCTTGGTACGG |
| 5697 | RP11-452L6.1-001 | TCTAAGCGCTAGACAGGACG |
| 5698 | RP11-452L6.1-001 | CCCAGAGCTAAGCAGAAGCA |
| 5699 | RP11-452L6.1-001 | GAGCCTGGCGAGACACACAC |
| 5700 | RP11-452L6.1-001 | CTCTAAGCGCTAGACAGGAC |
| 5701 | RP11-452L6.1-001 | ACTCTAAGCGCTAGACAGGA |
| 5702 | RP11-452L6.1-001 | CACACACAGGCTGGTGCTGT |
| 5703 | RP11-454K7.1-001 | ATTACGTTCAAATTGTAGGG |
| 5704 | RP11-454K7.1-001 | AATTACGTTCAAATTGTAGG |
| 5705 | RP11-454K7.1-001 | ATTTCAGCACTAAATCATGA |
| 5706 | RP11-454K7.1-001 | AAGGCTGGATTCTTGAAGCC |
| 5707 | RP11-454K7.1-001 | GAAATTCAACATGTCAGTCT |
| 5708 | RP11-454K7.1-001 | AAATTCAACATGTCAGTCTG |
| 5709 | RP11-454K7.1-001 | CTGTGATGAGCCTGAACATC |
| 5710 | RP11-454P21.1-004 | CAAGTTGAAATAGAATTGAT |
| 5711 | RP11-454P21.1-004 | CCATCAATTCTATTTCAACT |
| 5712 | RP11-454P21.1-004 | TAGAATTGATGGGATACTTT |
| 5713 | RP11-454P21.1-004 | CCAAGTTGAAATAGAATTGA |
| 5714 | RP11-454P21.1-004 | AAAAAGAATAAATAAATAGC |
| 5715 | RP11-454P21.1-004 | AAGTTCCAGGATACATGTGC |
| 5716 | RP11-454P21.1-004 | GATGTGCAGGTTTGTTACGT |
| 5717 | RP11-458F8.4-001 | CCAGGCCCAGCACACTCTCG |
| 5718 | RP11-458F8.4-001 | GCAAGCCTCCCCTGTCCTGG |

| | | |
|---|---|---|
| 5719 | RP11-458F8.4-001 | GCCGCCACGAAGGAAAAGCT |
| 5720 | RP11-458F8.4-001 | GGGAGACAGCCACCAGGACA |
| 5721 | RP11-458F8.4-001 | GACAGCCACCAGGACAGGGG |
| 5722 | RP11-458F8.4-001 | GGAGACAGCCACCAGGACAG |
| 5723 | RP11-458F8.4-001 | CTGGGCCTGGAGAGTAGGCC |
| 5724 | RP11-468E2.10-001 | ATGCTATAAACAAACCTGGG |
| 5725 | RP11-468E2.10-001 | GATATGCTATAAACAAACCT |
| 5726 | RP11-468E2.10-001 | CTGGGAGGAGATGTAAGGGA |
| 5727 | RP11-468E2.10-001 | AGATATGCTATAAACAAACC |
| 5728 | RP11-468E2.10-001 | AAACCTGGGAGGAGATGTAA |
| 5729 | RP11-468E2.10-001 | CAAAAAAAAAGGATAAGGCC |
| 5730 | RP11-468E2.10-001 | GAATCGCTTGAACTCGGAGG |
| 5731 | RP11-46D6.1-002 | CTGGGGCTGAAATTCAAACG |
| 5732 | RP11-46D6.1-002 | GGGAACAGGGAGCATTCACA |
| 5733 | RP11-46D6.1-002 | GGGCTGAAATTCAAACGGGG |
| 5734 | RP11-46D6.1-002 | AAAAACAGAGGGGCTCTGCA |
| 5735 | RP11-46D6.1-002 | CCCTGTTCCCCCAAAAACAG |
| 5736 | RP11-46D6.1-002 | CCTGTTCCCCCAAAAACAGA |
| 5737 | RP11-46D6.1-002 | GGTCTTGGGAACAAAGGCCT |
| 5738 | RP11-473I1.6-001 | AGTCACCACATGTCATCACG |
| 5739 | RP11-473I1.6-001 | GAGGCCTTGTACTGAAAGCG |
| 5740 | RP11-473I1.6-001 | ACCACATGTCATCACGTGGA |
| 5741 | RP11-473I1.6-001 | TGATGACATGTGGTGACTCA |
| 5742 | RP11-473I1.6-001 | CATCTTGCTCATCATAGAGG |
| 5743 | RP11-473I1.6-001 | GCCTTCCACGTGATGACATG |
| 5744 | RP11-473I1.6-001 | GTGATGACATGTGGTGACTC |
| 5745 | RP11-473M20.16-001 | ACAAAATCTCACGCCCAACG |
| 5746 | RP11-473M20.16-001 | AAGGGGATCCTGAAGCCAGT |
| 5747 | RP11-473M20.16-001 | CAAAATCTCACGCCCAACGT |
| 5748 | RP11-473M20.16-001 | AAAATCTCACGCCCAACGTG |
| 5749 | RP11-473M20.16-001 | GGATCCTGAAGCCAGTTGGT |
| 5750 | RP11-473M20.16-001 | AAAACGCTCCACCAACCAAC |
| 5751 | RP11-473M20.16-001 | CAAATCGGGAATTGTTGGTC |
| 5752 | RP11-474N24.6-001 | CCAGGTATTAGGGTCCATCC |
| 5753 | RP11-474N24.6-001 | CCTGGATGGACCCTAATACC |
| 5754 | RP11-474N24.6-001 | CGAAACAGCAGTCTAGAAGT |
| 5755 | RP11-474N24.6-001 | CATGTGCGATCCAGGTATTA |
| 5756 | RP11-474N24.6-001 | CTGGATCGCACATGAAAACT |
| 5757 | RP11-474N24.6-001 | CTGAAATTCCTGGGAGTGAG |
| 5758 | RP11-474N24.6-001 | GCCAGGCTAGCAATTCCAGC |
| 5759 | RP11-47A8.5-001 | AGGGCTGCTAGATCCGGACG |
| 5760 | RP11-47A8.5-001 | GGATCTAGCAGCCCTCGGCG |
| 5761 | RP11-47A8.5-001 | CGTCCGGATCTAGCAGCCCT |

| | | |
|---|---|---|
| 5762 | RP11-47A8.5-001 | CGAGGGCTGCTAGATCCGGA |
| 5763 | RP11-47A8.5-001 | GAGGGCTGCTAGATCCGGAC |
| 5764 | RP11-47A8.5-001 | GCGCCGAGGGCTGCTAGATC |
| 5765 | RP11-47A8.5-001 | GCGGGGCCAGGAGGCGAGGG |
| 5766 | RP11-480A16.1-001 | GATACTTCCCACTAATAACG |
| 5767 | RP11-480A16.1-001 | TGTATCCAATGATAGCTCAG |
| 5768 | RP11-480A16.1-001 | TGTTGCCAGAGACTGAACTG |
| 5769 | RP11-480A16.1-001 | TTACAGAAAAAGCAGCAGTC |
| 5770 | RP11-480A16.1-001 | ATGTATCCAATGATAGCTCA |
| 5771 | RP11-480A16.1-001 | AAAGGCTCCTCGTTATTAGT |
| 5772 | RP11-480A16.1-001 | AGAAAAGATGACTGCCCCTG |
| 5773 | RP11-482G13.1-003 | CAGCTCCTCAGGAACGACTG |
| 5774 | RP11-482G13.1-003 | CAGTCGTTCCTGAGGAGCTG |
| 5775 | RP11-482G13.1-003 | GAGCTGAGGAGAAGGAGGCT |
| 5776 | RP11-482G13.1-003 | TGGCTCTCTAGGAACACCTG |
| 5777 | RP11-482G13.1-003 | TTCCTGAGGAGCTGAGGAGA |
| 5778 | RP11-482G13.1-003 | CTGAGGAGCTGAGGAGAAGG |
| 5779 | RP11-482G13.1-003 | CTGGCTCTCTAGGAACACCT |
| 5780 | RP11-482M8.1-002 | CGTCGTCGTCGTCGCCCGGG |
| 5781 | RP11-482M8.1-002 | TTCCCCGCGCCCGTGCCCAG |
| 5782 | RP11-482M8.1-002 | CGTCGTCGTCGTCGTCGCCC |
| 5783 | RP11-482M8.1-002 | TCGTCGTCGTCGTCGTCGCC |
| 5784 | RP11-482M8.1-002 | GCCGCGCACCGGCCGCCGCT |
| 5785 | RP11-482M8.1-002 | CGTCGTCGTCGCCCGGGAGG |
| 5786 | RP11-482M8.1-002 | CAGTGACAGACCCTGCGGCG |
| 5787 | RP11-486O12.2-001 | AAAGCAACCAAGCATGCCGA |
| 5788 | RP11-486O12.2-001 | CTTGAACGACGGGTTCCTGT |
| 5789 | RP11-486O12.2-001 | TGGGTTCCGGGATCCCTTGG |
| 5790 | RP11-486O12.2-001 | GATTCACAAAGCTCAGCGAG |
| 5791 | RP11-486O12.2-001 | CTGGGTTCCGGGATCCCTTG |
| 5792 | RP11-486O12.2-001 | CAACAGGAACCCGTCGTTCA |
| 5793 | RP11-486O12.2-001 | CCCAAGGGATCCCGGAACCC |
| 5794 | RP11-488C13.5-001 | GCTGGTCTTACTACATAGGG |
| 5795 | RP11-488C13.5-001 | GAAGACAAAAGCACCAACAG |
| 5796 | RP11-488C13.5-001 | TTGGCTGGTCTTACTACATA |
| 5797 | RP11-488C13.5-001 | GTCTTACTACATAGGGAGGT |
| 5798 | RP11-488C13.5-001 | TCTGGTTTCAGGGCTCCCAG |
| 5799 | RP11-488C13.5-001 | ACAAAAGCACCAACAGAGGT |
| 5800 | RP11-488C13.5-001 | CTTGGCTGGTCTTACTACAT |
| 5801 | RP11-488L18.10-001 | TTACATACCATGACTAAGCA |
| 5802 | RP11-488L18.10-001 | GAAAATACCCTGCTTAGTCA |
| 5803 | RP11-488L18.10-001 | ATTCCCTATATTGTCCAGAA |
| 5804 | RP11-488L18.10-001 | ATTACATACCATGACTAAGC |

| | | |
|---|---|---|
| 5805 | RP11-488L18.10-001 | TGGAAAGACTGCATAAAGCC |
| 5806 | RP11-488L18.10-001 | TACTTGAAATAGAAAAAGTT |
| 5807 | RP11-488L18.10-001 | AAAGAATACAAAAAGGTATA |
| 5808 | RP11-489E7.1-001 | CCCACGACCCACGAATCCAG |
| 5809 | RP11-489E7.1-001 | CTCAGGACCCAACTTCACAT |
| 5810 | RP11-489E7.1-001 | ATACAATACTACGATACATC |
| 5811 | RP11-489E7.1-001 | TCATTAAAGGAGGATCTGGT |
| 5812 | RP11-489E7.1-001 | CTCTGGATTCGTGGGTCGTG |
| 5813 | RP11-489E7.1-001 | GTAACAGCTCTCATTAAAGG |
| 5814 | RP11-489E7.1-001 | AGTTGGGTCCTGAGAATAGG |
| 5815 | RP11-489O18.1-001 | GAGTGGGAGCCAAAACAGGT |
| 5816 | RP11-489O18.1-001 | CCCACACACAGCATCAAGGG |
| 5817 | RP11-489O18.1-001 | CACACAGCATCAAGGGAGGG |
| 5818 | RP11-489O18.1-001 | CCACACACAGCATCAAGGGA |
| 5819 | RP11-489O18.1-001 | TGTCCCACACACAGCATCAA |
| 5820 | RP11-489O18.1-001 | CAAAAATGTGACTTGTCCTA |
| 5821 | RP11-489O18.1-001 | AATGTGACTTGTCCTAGGGC |
| 5822 | RP11-48B3.3-001 | ATGTGACCATTAGCCAGGCG |
| 5823 | RP11-48B3.3-001 | AGGACAGATAACATAGTATG |
| 5824 | RP11-48B3.3-001 | TGACCATTAGCCAGGCGTGG |
| 5825 | RP11-48B3.3-001 | GAGCCACCACGCCTGGCTAA |
| 5826 | RP11-48B3.3-001 | CAGGTGTGAGCCACCACGCC |
| 5827 | RP11-48B3.3-001 | AGCACTTTGGGAGGCCAAGG |
| 5828 | RP11-48B3.3-001 | AAAAATACAAAAAATTAGCA |
| 5829 | RP11-493L12.4-002 | CTGGGATCATATCGGATGGG |
| 5830 | RP11-493L12.4-002 | ACTCCTGGGATCATATCGGA |
| 5831 | RP11-493L12.4-002 | ACTTATGTATCTGCCATACG |
| 5832 | RP11-493L12.4-002 | CTCCTGGGATCATATCGGAT |
| 5833 | RP11-493L12.4-002 | GATTACAGCTCCAGACAGAG |
| 5834 | RP11-493L12.4-002 | AAGTCAGGTATGCCACTCCT |
| 5835 | RP11-493L12.4-002 | ATCCGATATGATCCCAGGAG |
| 5836 | RP11-494G9.1-001 | CACAGGTCCTGATATGTGAG |
| 5837 | RP11-494G9.1-001 | ATATGTGAGAGGTATTCACC |
| 5838 | RP11-494G9.1-001 | AGCTGTCGTGATGATGGATG |
| 5839 | RP11-494G9.1-001 | GAGCTGTCGTGATGATGGAT |
| 5840 | RP11-494G9.1-001 | GTGAATACCTCTCACATATC |
| 5841 | RP11-494G9.1-001 | CTCTGGACCCTCTTTCCCAG |
| 5842 | RP11-494G9.1-001 | CTGGAGAGCTGTCGTGATGA |
| 5843 | RP11-495K9.5-001 | GAGTCGACTGTCAGCCACAG |
| 5844 | RP11-495K9.5-001 | AGTCGACTGTCAGCCACAGA |
| 5845 | RP11-495K9.5-001 | AGAAGTACCCTTGAGACTCA |
| 5846 | RP11-495K9.5-001 | GTCGACTGTCAGCCACAGAG |
| 5847 | RP11-495K9.5-001 | GTCACTGCCCTGAGTCTCAA |

| | | |
|---|---|---|
| 5848 | RP11-495K9.5-001 | GAGAAGTACCCTTGAGACTC |
| 5849 | RP11-495K9.5-001 | GACTCAGGGCAGTGACTCCA |
| 5850 | RP11-496H1.2-001 | TGTGGAGCTGAAGACAACGA |
| 5851 | RP11-496H1.2-001 | AGACAACGAAGGAGCCAGTG |
| 5852 | RP11-496H1.2-001 | TGGCTGTTCTGTGAGACTCG |
| 5853 | RP11-496H1.2-001 | CCGCTGTTCAAATCTGAGGG |
| 5854 | RP11-496H1.2-001 | GCTAAAACACCTCTCCCACC |
| 5855 | RP11-496H1.2-001 | GGGCTGAAGACCTGGAGAGG |
| 5856 | RP11-496H1.2-001 | GTGGTGGAGATGGAGCCTTG |
| 5857 | RP11-498P14.5-001 | TCAGCAGGACCCACTAGCCA |
| 5858 | RP11-498P14.5-001 | GGCTACTTACGACGGAAGCT |
| 5859 | RP11-498P14.5-001 | TTACGACGGAAGCTCGGTTG |
| 5860 | RP11-498P14.5-001 | GACCCACTAGCCAAGGAAAG |
| 5861 | RP11-498P14.5-001 | CTTACGACGGAAGCTCGGTT |
| 5862 | RP11-498P14.5-001 | AGGACCCACTAGCCAAGGAA |
| 5863 | RP11-498P14.5-001 | ACTTACGACGGAAGCTCGGT |
| 5864 | RP11-501E14.1-001 | ACATGTTTATCTAAGGTAAG |
| 5865 | RP11-501E14.1-001 | GTGGGAGTTAGGACTGCCAG |
| 5866 | RP11-501E14.1-001 | TTAGGACTGCCAGTGGTGCA |
| 5867 | RP11-501E14.1-001 | CAGACTCTTCCATGCACCAC |
| 5868 | RP11-501E14.1-001 | AGCAGTATGGAAGCTTCTGG |
| 5869 | RP11-501E14.1-001 | TGCTGCTTTATGGTGATAGA |
| 5870 | RP11-501E14.1-001 | GGCAGTCCTAACTCCCACTT |
| 5871 | RP11-504P24.9-001 | GGTGGACCAGCCGAAGTCAG |
| 5872 | RP11-504P24.9-001 | AGCCTTGACCTTCGGGCTCA |
| 5873 | RP11-504P24.9-001 | GAAGGATTCCTTGAGCCCGA |
| 5874 | RP11-504P24.9-001 | GGAGTGAGTTGGTGCAAACA |
| 5875 | RP11-504P24.9-001 | TTCCTTGAGCCCGAAGGTCA |
| 5876 | RP11-504P24.9-001 | AGCTACCGGGGAGGCTGACA |
| 5877 | RP11-504P24.9-001 | ACCGGGGAGGCTGACAGGGA |
| 5878 | RP11-505C13.1-001 | AACTAGAAGCTGCTGGCACA |
| 5879 | RP11-505C13.1-001 | GATCACATTACCTTCCTGAT |
| 5880 | RP11-505C13.1-001 | CTACACAGACTGAAAACGAT |
| 5881 | RP11-505C13.1-001 | CAGTCTGTGTAGTCTCTGAA |
| 5882 | RP11-505C13.1-001 | TCCACAAAACCCTATCAGGA |
| 5883 | RP11-505C13.1-001 | TAAATCCACAAAACCCTATC |
| 5884 | RP11-505C13.1-001 | AATTCATCTCACTGGAAATC |
| 5885 | RP11-506M13.3-002 | CATATTACTCTGTTTGTACA |
| 5886 | RP11-506M13.3-002 | TGTACAAACAGAGTAATATG |
| 5887 | RP11-506M13.3-002 | ATAAACCAGGATGCATTAGC |
| 5888 | RP11-506M13.3-002 | TATACTAAGATGGATAGGCA |
| 5889 | RP11-506M13.3-002 | TGTGCTTTGTTGTATACATT |
| 5890 | RP11-506M13.3-002 | AATAAAGAGAAAGCGAATAA |

| | | |
|---|---|---|
| 5891 | RP11-506M13.3-002 | AACTGACAGCAGCAGGAATG |
| 5892 | RP11-517P14.2-001 | GGCCCCGGAAAGGCGAAAGG |
| 5893 | RP11-517P14.2-001 | GAGGAGCCAAAAGAGGCTCT |
| 5894 | RP11-517P14.2-001 | GCACACGCCTGGCCCCGGAA |
| 5895 | RP11-517P14.2-001 | TTCACAGAGCTGCACACGCC |
| 5896 | RP11-517P14.2-001 | CTCCTCCTTTCGCCTTTCCG |
| 5897 | RP11-517P14.2-001 | GCGAAAGGAGGAGCCAAAAG |
| 5898 | RP11-517P14.2-001 | CGAGCGGCTGAGGCGAGAGG |
| 5899 | RP11-51J9.5-001 | TATGCAGGGCTAAAGTCCGC |
| 5900 | RP11-51J9.5-001 | ATTTGGAACAGCCATTGACT |
| 5901 | RP11-51J9.5-001 | GGTCTACCCATCTCTATGCA |
| 5902 | RP11-51J9.5-001 | CTTTAGCCCTGCATAGAGAT |
| 5903 | RP11-51J9.5-001 | CTATGCAGGGCTAAAGTCCG |
| 5904 | RP11-51J9.5-001 | GGGTCTACCCATCTCTATGC |
| 5905 | RP11-51J9.5-001 | GCAGGGCTAAAGTCCGCGGG |
| 5906 | RP11-525A16.4-002 | CCAGTACGCACCATGCTGGG |
| 5907 | RP11-525A16.4-002 | AAAACGCAACCCCCCCAGCA |
| 5908 | RP11-525A16.4-002 | CCCAGCATGGTGCGTACTGG |
| 5909 | RP11-525A16.4-002 | CAGTACGCACCATGCTGGGG |
| 5910 | RP11-525A16.4-002 | TGGTGCGTACTGGAGGTATG |
| 5911 | RP11-525A16.4-002 | CAGGCAGAGGAAACTGCTTG |
| 5912 | RP11-525A16.4-002 | ACCTCCAGTACGCACCATGC |
| 5913 | RP11-526A4.1-003 | ATGGGTGCGGCACACCAACA |
| 5914 | RP11-526A4.1-003 | TATGTATACATGTGCCATGT |
| 5915 | RP11-526A4.1-003 | AAATGACGAGTTAATGGGTG |
| 5916 | RP11-526A4.1-003 | GGTACATGTGCACAACGTGC |
| 5917 | RP11-526A4.1-003 | CCAATATCCCTGATGAACAT |
| 5918 | RP11-526A4.1-003 | TGAAGCCAACTTGATCATGG |
| 5919 | RP11-526A4.1-003 | CTTTGGTATCAGGATGATGT |
| 5920 | RP11-529E15.1-002 | ATGGGAATGATATTCAAGAG |
| 5921 | RP11-529E15.1-002 | TCAACCTGTGAAGATCGATT |
| 5922 | RP11-529E15.1-002 | TTCCTACTGCACCATGCCAT |
| 5923 | RP11-529E15.1-002 | CAAGAGTGGTGACATGACTG |
| 5924 | RP11-529E15.1-002 | AGAGGATGCAGAAATCTGTG |
| 5925 | RP11-529E15.1-002 | TGAATATCATTCCCATGGCA |
| 5926 | RP11-529E15.1-002 | TGGCATGGTGCAGTAGGAAC |
| 5927 | RP11-530C5.1-001 | GGTGGCTACCGGACCAGCAG |
| 5928 | RP11-530C5.1-001 | CCACTCCTCCAGGAAGTCTG |
| 5929 | RP11-530C5.1-001 | AGACTTCCTGGAGGAGTGGA |
| 5930 | RP11-530C5.1-001 | CAGCACCACAGACTTCCTGG |
| 5931 | RP11-530C5.1-001 | GTTTCGCCTTCCACTCCTCC |
| 5932 | RP11-530C5.1-001 | AGTCTGTGGTGCTGCCGCCG |
| 5933 | RP11-530C5.1-001 | CGGACCAGCAGCGGCCTCGG |

| | | |
|---|---|---|
| 5934 | RP11-533E19.5-001 | CAAAGCCAATACTTAATGGG |
| 5935 | RP11-533E19.5-001 | AGCTGCAACTACTCTGACAG |
| 5936 | RP11-533E19.5-001 | GTGCTTTGAGAGGCTCTCTG |
| 5937 | RP11-533E19.5-001 | GAGCTGCAACTACTCTGACA |
| 5938 | RP11-533E19.5-001 | AGTTTCCTCCCATTAAGTAT |
| 5939 | RP11-533E19.5-001 | TGCTTTAACAGTGCTTTGAG |
| 5940 | RP11-533E19.5-001 | ACAGTGACAGCAGTGGGTAA |
| 5941 | RP11-533E19.7-001 | CAACAACTATGGCGGGCGAC |
| 5942 | RP11-533E19.7-001 | GCTACGTCAACAACTATGGC |
| 5943 | RP11-533E19.7-001 | TAAAGCTACGTCAACAACTA |
| 5944 | RP11-533E19.7-001 | GGGGTGTGTACGTCACCCCC |
| 5945 | RP11-533E19.7-001 | GGGTGTGTACGTCACCCCCA |
| 5946 | RP11-533E19.7-001 | CCTCTCGGATGGGGGCCCTG |
| 5947 | RP11-533E19.7-001 | CCCTCTCGGATGGGGGCCCT |
| 5948 | RP11-538D16.2-001 | TGTAGGTGGTGTGACCTGGG |
| 5949 | RP11-538D16.2-001 | GCATGTAGGTGGTGTGACCT |
| 5950 | RP11-538D16.2-001 | CCCTGGCCAGCGCATGTAGG |
| 5951 | RP11-538D16.2-001 | GTCACACCACCTACATGCGC |
| 5952 | RP11-538D16.2-001 | CCACCTACATGCGCTGGCCA |
| 5953 | RP11-538D16.2-001 | GTACCCTGGCCAGCGCATGT |
| 5954 | RP11-538D16.2-001 | CCGCAGCTGTGTCAGAAGCC |
| 5955 | RP11-54C4.3-001 | TGGCCAATCGCAACGCCGGA |
| 5956 | RP11-54C4.3-001 | CGAGAGGCCAAGACAGCCTG |
| 5957 | RP11-54C4.3-001 | CCTGTGGCCAATCGCAACGC |
| 5958 | RP11-54C4.3-001 | CAGCCTCACAACCGCCCAGA |
| 5959 | RP11-54C4.3-001 | ACAGCCTCACAACCGCCCAG |
| 5960 | RP11-54C4.3-001 | GCTCCTTCCGGCGTTGCGAT |
| 5961 | RP11-54C4.3-001 | GAAGGAGCTGATAGACACGG |
| 5962 | RP11-54G14.1-001 | AATGGCAGATAAGCCGAGTG |
| 5963 | RP11-54G14.1-001 | GTGCAGGATGGAATGCAGAG |
| 5964 | RP11-54G14.1-001 | GCATCACTACATGTGCAGGA |
| 5965 | RP11-54G14.1-001 | GGCAGATAAGCCGAGTGCGG |
| 5966 | RP11-54G14.1-001 | AAAAATTAGGGAGGGCATGG |
| 5967 | RP11-54G14.1-001 | TGCAGGATGGAATGCAGAGT |
| 5968 | RP11-54G14.1-001 | GTAATCCCAGCAGTTTGGGA |
| 5969 | RP11-551L14.6-001 | TCTATCCTGTCAGATAACCA |
| 5970 | RP11-551L14.6-001 | CAGGATAGAAGAGAAGTGGG |
| 5971 | RP11-551L14.6-001 | CGATCGGGTTCCTCGGATGG |
| 5972 | RP11-551L14.6-001 | GGACGGACATCTCAGATCAT |
| 5973 | RP11-551L14.6-001 | TGGGAAGTGCTGGCCTGAGC |
| 5974 | RP11-551L14.6-001 | AGTGCTGGCCTGAGCAGGCA |
| 5975 | RP11-551L14.6-001 | TGCTGGAGAAAAAGTTGAAG |
| 5976 | RP11-553L6.5-001 | GCATTCGCATCGGATCAAGA |

| | | |
|---|---|---|
| 5977 | RP11-553L6.5-001 | GAACATTACTGCATTCGCAT |
| 5978 | RP11-553L6.5-001 | TCAAATGGATGGCCACGTTA |
| 5979 | RP11-553L6.5-001 | AGATAGCAAACACATCAATA |
| 5980 | RP11-553L6.5-001 | GTGGCCATCCATTTGAGAAG |
| 5981 | RP11-553L6.5-001 | TGGCCATCCATTTGAGAAGT |
| 5982 | RP11-553L6.5-001 | AAGGTGCTCAAAGAAAAAAG |
| 5983 | RP11-555J4.3-001 | GCTTACTGGTTGGATTCCAG |
| 5984 | RP11-555J4.3-001 | GGGAACACAGGAGGTACAGT |
| 5985 | RP11-555J4.3-001 | TCTGATAACAGCTTACTGGT |
| 5986 | RP11-555J4.3-001 | AGCTTACTGGTTGGATTCCA |
| 5987 | RP11-555J4.3-001 | TTGGCAATGGTATCCAGGAG |
| 5988 | RP11-555J4.3-001 | CAGAACCAGAGCTACTCCCC |
| 5989 | RP11-555J4.3-001 | GCTTGTTGGCAATGGTATCC |
| 5990 | RP11-561B11.6-001 | AATTTCAGGACTGGGTGTGG |
| 5991 | RP11-561B11.6-001 | GAAATTAACATTTGAGACAG |
| 5992 | RP11-561B11.6-001 | TTGAGACAGTGGACTGGGAG |
| 5993 | RP11-561B11.6-001 | AGTCGAAAAGCTGAAGAACT |
| 5994 | RP11-561B11.6-001 | TTACACCAGTGGTTTGTCAG |
| 5995 | RP11-561B11.6-001 | ACTTGGAGTCCAATGTTCGA |
| 5996 | RP11-561B11.6-001 | AACATTTGAGACAGTGGACT |
| 5997 | RP11-562A8.5-001 | AGTGTGTTGTGACAGGACAC |
| 5998 | RP11-562A8.5-001 | TACCTACAGTGTGTTGTGAC |
| 5999 | RP11-562A8.5-001 | GTCCTGTCACAACACACTGT |
| 6000 | RP11-562A8.5-001 | AAAATAATCTAAGGCAATCC |
| 6001 | RP11-562A8.5-001 | ACACAGGAAAGCCCCTAACC |
| 6002 | RP11-562A8.5-001 | TATTTAGTTAGTCACTCTGC |
| 6003 | RP11-562A8.5-001 | CTAGATCACAAGCTTTGGGA |
| 6004 | RP11-563N4.1-001 | GAGCTCTCCGAGAGATTGGT |
| 6005 | RP11-563N4.1-001 | AAGCGTTGAGGTTAAGGGCA |
| 6006 | RP11-563N4.1-001 | TCAGGTCCCGACCAATCTCT |
| 6007 | RP11-563N4.1-001 | GCTCTAGCCGCCTTCCGCCA |
| 6008 | RP11-563N4.1-001 | GGTCGGGACCTGAAGCGTTG |
| 6009 | RP11-563N4.1-001 | GCTGTCCCTGCACTCCGTGG |
| 6010 | RP11-563N4.1-001 | AGCTCTCCGAGAGATTGGTC |
| 6011 | RP11-565N2.2-001 | ATGGATGATTAGAGAATGAG |
| 6012 | RP11-565N2.2-001 | AATTGTCGCCATGTAAGTCT |
| 6013 | RP11-565N2.2-001 | GTATGAATCCCAGACTTACA |
| 6014 | RP11-565N2.2-001 | ACAGTCCTGTGGTCAATGCT |
| 6015 | RP11-565N2.2-001 | TGTGGTCAATGCTCGGGTCA |
| 6016 | RP11-565N2.2-001 | ATGACCCGAGCATTGACCAC |
| 6017 | RP11-565N2.2-001 | CAGTCCTGTGGTCAATGCTC |
| 6018 | RP11-568N6.1-001 | ATAGACGTCTTCCCGCACTC |
| 6019 | RP11-568N6.1-001 | CTGAACGTTAACCGGCTGAA |

| | | |
|---|---|---|
| 6020 | RP11-568N6.1-001 | GTTAACGTTCAGCTCAAGAC |
| 6021 | RP11-568N6.1-001 | GGAGAGCAAGAGACTTAATC |
| 6022 | RP11-568N6.1-001 | CTCAACCGGCGCCTGAGTGC |
| 6023 | RP11-568N6.1-001 | CCTCAACCGGCGCCTGAGTG |
| 6024 | RP11-568N6.1-001 | CCGCACTCAGGCGCCGGTTG |
| 6025 | RP11-571E6.3-001 | CTGCTGGGTTACATTCCCCG |
| 6026 | RP11-571E6.3-001 | CAGAAAAAGCCAAACCTGCT |
| 6027 | RP11-571E6.3-001 | CCAAGAGTGCCCAACCCTCG |
| 6028 | RP11-571E6.3-001 | TGTTCATGGTTGAGGGAACA |
| 6029 | RP11-571E6.3-001 | TTTAGAGATAAGAGTAACAT |
| 6030 | RP11-571E6.3-001 | TGCTGGGTTACATTCCCCGA |
| 6031 | RP11-571E6.3-001 | GACCAAGAGTGCCCAACCCT |
| 6032 | RP11-572O17.1-001 | AACAATAGACGGCGCGTACG |
| 6033 | RP11-572O17.1-001 | CGGGTACTGACTGCTTCCCA |
| 6034 | RP11-572O17.1-001 | AAACAATAGACGGCGCGTAC |
| 6035 | RP11-572O17.1-001 | TGCAGCGGCCACGGGACCGT |
| 6036 | RP11-572O17.1-001 | AAGCAGTCAGTACCCGCCAC |
| 6037 | RP11-572O17.1-001 | AAAACAATAGACGGCGCGTA |
| 6038 | RP11-572O17.1-001 | GACTGCTTCCCACGGTCCCG |
| 6039 | RP11-573D15.2-001 | CCCCAGAACGCGGCTTGTGG |
| 6040 | RP11-573D15.2-001 | CCCAGAACGCGGCTTGTGGG |
| 6041 | RP11-573D15.2-001 | CCCCCACAAGCCGCGTTCTG |
| 6042 | RP11-573D15.2-001 | GAGGCCACGGCCCCAGAACG |
| 6043 | RP11-573D15.2-001 | GGCCCCAGAACGCGGCTTGT |
| 6044 | RP11-573D15.2-001 | GCCCCAGAACGCGGCTTGTG |
| 6045 | RP11-573D15.2-001 | CGGCCCCAGAACGCGGCTTG |
| 6046 | RP11-573G6.10-001 | GCTCTTCTACGTCCCGCGCG |
| 6047 | RP11-573G6.10-001 | CGCCCTGGCCGGGAAAGGTG |
| 6048 | RP11-573G6.10-001 | CGGGACGTAGAAGAGCGCCC |
| 6049 | RP11-573G6.10-001 | GTCCCGCGCGCGGACAGCCG |
| 6050 | RP11-573G6.10-001 | GCGCCCTGGCCGGGAAAGGT |
| 6051 | RP11-573G6.10-001 | CGTAGAAGAGCGCCCTGGCC |
| 6052 | RP11-573G6.10-001 | ACGTAGAAGAGCGCCCTGGC |
| 6053 | RP11-574K11.29-002 | ATTGACACACCTCTACAGCT |
| 6054 | RP11-574K11.29-002 | GAACTCTTGCTTATTTGCCA |
| 6055 | RP11-574K11.29-002 | CCAACTGGTGAAGTACTGAG |
| 6056 | RP11-574K11.29-002 | AGAGGTGTGTCAATCCCAAC |
| 6057 | RP11-574K11.29-002 | GATTGACACACCTCTACAGC |
| 6058 | RP11-574K11.29-002 | AGACAGCAGCACACAGAACG |
| 6059 | RP11-574K11.29-002 | CCTCTCAGTACTTCACCAGT |
| 6060 | RP11-577B7.1-001 | GTTATACAACCTATGATCAG |
| 6061 | RP11-577B7.1-001 | GATCATAGGTTGTATAACAG |
| 6062 | RP11-577B7.1-001 | AAGCAATGTAAGAAAAAAGG |

| | | |
|---|---|---|
| 6063 | RP11-577B7.1-001 | AGGCATGGTGTGAGCTTGAG |
| 6064 | RP11-577B7.1-001 | TCGATGTGACCTCTGATCAT |
| 6065 | RP11-577B7.1-001 | AAAAAAGGAGGCGGGCATGG |
| 6066 | RP11-577B7.1-001 | GGAGGCAGGAGGATCACTTG |
| 6067 | RP11-57P19.1-001 | GGACTGTGACAGGAGATATG |
| 6068 | RP11-57P19.1-001 | AAGATAAACAAAAACCACTC |
| 6069 | RP11-57P19.1-001 | CAATAGTTCAGGACTGTGAC |
| 6070 | RP11-57P19.1-001 | CCTTTATAAAGGAGATTGAG |
| 6071 | RP11-57P19.1-001 | GACTGTGACAGGAGATATGG |
| 6072 | RP11-57P19.1-001 | CAAGACCAGTCTGGCCAACG |
| 6073 | RP11-57P19.1-001 | GAGACAGGGTTTCACCACGT |
| 6074 | RP11-582E3.6-001 | AGTCATGTCGTCTGCAAACA |
| 6075 | RP11-582E3.6-001 | CAGTCATGTCGTCTGCAAAC |
| 6076 | RP11-582E3.6-001 | AGCAACTTCAGCAAAGTCTC |
| 6077 | RP11-582E3.6-001 | TGAAGTTGCTTATCAGCTTA |
| 6078 | RP11-582E3.6-001 | CCAACTTACAAGGGACGTGA |
| 6079 | RP11-582E3.6-001 | ACTTAGGAATCCAACTTACA |
| 6080 | RP11-582E3.6-001 | ACTGCTCAAGGAAATAAAAG |
| 6081 | RP11-589M4.3-001 | AAATATCAGATTTAGCATCC |
| 6082 | RP11-589M4.3-001 | GAGAATGGCGTGAACCCGGG |
| 6083 | RP11-589M4.3-001 | ATAAAGAAATAGATATGGCC |
| 6084 | RP11-589M4.3-001 | TGTAGTCCCAGCTACTCGGG |
| 6085 | RP11-589M4.3-001 | CAGGAGAATGGCGTGAACCC |
| 6086 | RP11-589M4.3-001 | ATAGATATGGCCAGGTGCGA |
| 6087 | RP11-589M4.3-001 | AATGGCGTGAACCCGGGAGG |
| 6088 | RP11-589P10.5-001 | TAATCTGGAACCTACTGGAG |
| 6089 | RP11-589P10.5-001 | GAAGGAGCAGGAAAGAGCTT |
| 6090 | RP11-589P10.5-001 | AAGGAGCAGGAAAGAGCTTG |
| 6091 | RP11-589P10.5-001 | AAAATTAATCTGGAACCTAC |
| 6092 | RP11-589P10.5-001 | GCAGCCCCTTCCGCTCCAGT |
| 6093 | RP11-589P10.5-001 | TTGGGGATAAAAAATTAATC |
| 6094 | RP11-589P10.5-001 | TGAAGGAGCAGGAAAGAGCT |
| 6095 | RP11-616L12.1-001 | TCCCCGCAAATTACCACAGG |
| 6096 | RP11-616L12.1-001 | ACCTCCTGTGGTAATTTGCG |
| 6097 | RP11-616L12.1-001 | ATCTCCCCGCAAATTACCAC |
| 6098 | RP11-616L12.1-001 | TAAATAACTATACATGAGTG |
| 6099 | RP11-616L12.1-001 | AACTATACATGAGTGTGGAC |
| 6100 | RP11-616L12.1-001 | CTCCCTATTATTAAGGGTGC |
| 6101 | RP11-616L12.1-001 | CACCAGCACCCTTAATAATA |
| 6102 | RP11-61K9.3-001 | CGGTCGGGTTGTCTCCCCGC |
| 6103 | RP11-61K9.3-001 | GAATCTCGGGCTCCTGACCC |
| 6104 | RP11-61K9.3-001 | CTGGCGGACCCGGTAGCTGG |
| 6105 | RP11-61K9.3-001 | CCTGGCGGACCCGGTAGCTG |

| | | |
|---|---|---|
| 6106 | RP11-61K9.3-001 | CGGACCCGGTAGCTGGGGGC |
| 6107 | RP11-61K9.3-001 | CTACCGGGTCCGCCAGGGTC |
| 6108 | RP11-61K9.3-001 | CCCCAGCTACCGGGTCCGCC |
| 6109 | RP11-620J15.3-001 | AGATAACGGTCAATCGCCTC |
| 6110 | RP11-620J15.3-001 | TGTTCTGGGAGGAGTTACGC |
| 6111 | RP11-620J15.3-001 | GCTGTTACCAGGACAACCGG |
| 6112 | RP11-620J15.3-001 | CGGCTCCAAACCGCAGATAA |
| 6113 | RP11-620J15.3-001 | AAGACGCACGGTGTTCTGGG |
| 6114 | RP11-620J15.3-001 | TCAATCGCCTCCGGTTGTCC |
| 6115 | RP11-620J15.3-001 | GATAAGACGCACGGTGTTCT |
| 6116 | RP11-620J15.3-002 | GGGTACGACTCAGATCCCCG |
| 6117 | RP11-620J15.3-002 | AGGCGATTGACCGTTATCTG |
| 6118 | RP11-620J15.3-002 | GGATGCGCTAAAGGGTCGTG |
| 6119 | RP11-620J15.3-002 | AAACTAGAGAAGCGACAGTG |
| 6120 | RP11-620J15.3-002 | GATGCGCTAAAGGGTCGTGA |
| 6121 | RP11-620J15.3-002 | TGAAATGAAGTGTGGCCCAG |
| 6122 | RP11-620J15.3-002 | GGAGAGGCAGAGATATTCAG |
| 6123 | RP11-624L4.1-006 | GTTGCCATAGTGCACTGAGC |
| 6124 | RP11-624L4.1-006 | AAAGCCAGCTCAGTGCACTA |
| 6125 | RP11-624L4.1-006 | GCACTATGGCAACACACAGC |
| 6126 | RP11-624L4.1-006 | AGACAGTGGATGCAGGCCAG |
| 6127 | RP11-624L4.1-006 | GAATTATTTGACTCTAGTCA |
| 6128 | RP11-624L4.1-006 | CTTTCTAATACCAAACTCAG |
| 6129 | RP11-624L4.1-006 | GGCAACACACAGCAGGAGAA |
| 6130 | RP11-630C16.2-001 | GGGTTTGAACACAATCTGCA |
| 6131 | RP11-630C16.2-001 | GAACACAATCTGCAAGGAAG |
| 6132 | RP11-630C16.2-001 | TTACTGACACAATTAGCAAT |
| 6133 | RP11-630C16.2-001 | AGAGGAAACCAGCAGCAGCG |
| 6134 | RP11-630C16.2-001 | GAATCAAAGGCTGGACGCCG |
| 6135 | RP11-630C16.2-001 | TGTAAAATAGAATCAAAGGC |
| 6136 | RP11-630C16.2-001 | TTTACTGACACAATTAGCAA |
| 6137 | RP11-631N16.4-001 | TCTTTAGCTGTAGGTTGGAG |
| 6138 | RP11-631N16.4-001 | TTGGGAAAGAGCTAAGCCAT |
| 6139 | RP11-631N16.4-001 | GTGTGAGCGATCTGCAGAAC |
| 6140 | RP11-631N16.4-001 | CCAACCTTTAAAATGCATCG |
| 6141 | RP11-631N16.4-001 | ATCTGCAGAACTGGCATTAG |
| 6142 | RP11-631N16.4-001 | TTACTTCATTCTTTAGCTGT |
| 6143 | RP11-631N16.4-001 | CTTCAAGAAAAGGAATCCAA |
| 6144 | RP11-63B13.1-001 | CAGCGTGTTCCAATCAGAGG |
| 6145 | RP11-63B13.1-001 | TCTGTGGAAACCGCCCTGTA |
| 6146 | RP11-63B13.1-001 | TGGAGCCAAGACTGCGTCTG |
| 6147 | RP11-63B13.1-001 | TCTTGGCTCCACCCTCACCG |
| 6148 | RP11-63B13.1-001 | GTGGAGCCAAGACTGCGTCT |

| | | |
|---|---|---|
| 6149 | RP11-63B13.1-001 | CCAGTATTAGCCCTGGGCCG |
| 6150 | RP11-63B13.1-001 | CTGTGGAAACCGCCCTGTAT |
| 6151 | RP11-651P23.5-001 | ATTCCAAATAGAATCCAGGG |
| 6152 | RP11-651P23.5-001 | TCAGCCTGAAATGGTAAAGT |
| 6153 | RP11-651P23.5-001 | GAAATTCCAAATAGAATCCA |
| 6154 | RP11-651P23.5-001 | GCATAAACAAACATGAGAGT |
| 6155 | RP11-651P23.5-001 | ATCAGCCTGAAATGGTAAAG |
| 6156 | RP11-651P23.5-001 | AGATGATGAAGGTGGTATGC |
| 6157 | RP11-651P23.5-001 | TAACCTCCCTGGATTCTATT |
| 6158 | RP11-66B24.2-001 | AGTCTCATAGCTGCCGTGAG |
| 6159 | RP11-66B24.2-001 | TTCTGTCGCGAACCCTCTCA |
| 6160 | RP11-66B24.2-001 | CGCGACAGAATCACCGCGAA |
| 6161 | RP11-66B24.2-001 | TCGCGACAGAATCACCGCGA |
| 6162 | RP11-66B24.2-001 | CCCGGATCGTCGGGAGAGGT |
| 6163 | RP11-66B24.2-001 | CCCACCTCTCCCGACGATCC |
| 6164 | RP11-66B24.2-001 | ACTGCTGCGCTCCCCAGACG |
| 6165 | RP11-670E13.6-001 | CGATCCCCAAGAGCTCCCGC |
| 6166 | RP11-670E13.6-001 | CCGATCCCCAAGAGCTCCCG |
| 6167 | RP11-670E13.6-001 | CCCAAGAGCTCCCGCGGGCT |
| 6168 | RP11-670E13.6-001 | CCAGCCCGCGGGAGCTCTTG |
| 6169 | RP11-670E13.6-001 | CCGGGGGCTAAGGCCGAGCC |
| 6170 | RP11-670E13.6-001 | CCCAGCCCGCGGGAGCTCTT |
| 6171 | RP11-670E13.6-001 | CCGCGGGAGCTCTTGGGGAT |
| 6172 | RP11-671J11.7-001 | AACAGATCACAAGAAGATTG |
| 6173 | RP11-671J11.7-001 | CAAGACAGCTACTTAGTGGT |
| 6174 | RP11-671J11.7-001 | CCACAGTGCTGGGATTACAG |
| 6175 | RP11-671J11.7-001 | AGCACTGTGGGGGTCCAAAG |
| 6176 | RP11-671J11.7-001 | GATCACAAGAAGATTGAGGT |
| 6177 | RP11-671J11.7-001 | CAACCACTAAGTAGCTGTCT |
| 6178 | RP11-671J11.7-001 | AAGAAGATTGAGGTTGGGTG |
| 6179 | RP11-67L2.2-001 | GCCCAGACTTCCTGACACTG |
| 6180 | RP11-67L2.2-001 | ATGCTCACATTATAGTCACG |
| 6181 | RP11-67L2.2-001 | TCTGCATGGCTGAAGACATG |
| 6182 | RP11-67L2.2-001 | CACGAGGAAGTCTTGTTGGC |
| 6183 | RP11-67L2.2-001 | TAGTCACGAGGAAGTCTTGT |
| 6184 | RP11-67L2.2-001 | AGACTTCCTGACACTGAGGT |
| 6185 | RP11-67L2.2-001 | AAGTCTTGTTGGCCGGACAT |
| 6186 | RP11-681H18.2-001 | AATCTCATCTAAAAAAAAGG |
| 6187 | RP11-681H18.2-001 | GATTCTCAAAGACTTCGATA |
| 6188 | RP11-681H18.2-001 | CTCACTACTTAATACTGCCA |
| 6189 | RP11-681H18.2-001 | ATATGCTTCCCAGACACACA |
| 6190 | RP11-681H18.2-001 | AAATGTATCCATGTGTGTCT |
| 6191 | RP11-681H18.2-001 | ATACTGCCATGGTGAAAATG |

| | | |
|---|---|---|
| 6192 | RP11-681H18.2-001 | CTGCCATGGTGAAAATGTGG |
| 6193 | RP11-686D22.4-001 | CCCTTAAAATCACGAACTGG |
| 6194 | RP11-686D22.4-001 | GGATTACAAGACTTCGTATG |
| 6195 | RP11-686D22.4-001 | CACGAACTGGTGGTATAGTG |
| 6196 | RP11-686D22.4-001 | CATGGCAACTCTCTACAGGT |
| 6197 | RP11-686D22.4-001 | ACTACCAGGCCCAGGATGTG |
| 6198 | RP11-686D22.4-001 | AGTGCATGGCAACTCTCTAC |
| 6199 | RP11-686D22.4-001 | AACATAACCGATTAGGTCAT |
| 6200 | RP11-691G17.1-001 | CCTGGGAAGTCTTCACAAGA |
| 6201 | RP11-691G17.1-001 | TTCATTTAGTTCATTAGGCG |
| 6202 | RP11-691G17.1-001 | ATGAACTAAATGAAGCACCA |
| 6203 | RP11-691G17.1-001 | AATGAACTAAATGAAGCACC |
| 6204 | RP11-691G17.1-001 | TCAAGTCTCTTCAAATACCT |
| 6205 | RP11-691G17.1-001 | TAACACAGAGAAGGAATTGA |
| 6206 | RP11-691G17.1-001 | GGAAGTCTTCACAAGAAGGA |
| 6207 | RP11-692C24.2-001 | ATAATATTTACTATTACTAG |
| 6208 | RP11-692C24.2-001 | CAATCAGATACACATAACCT |
| 6209 | RP11-692C24.2-001 | TAATAATATTTACTATTACT |
| 6210 | RP11-692C24.2-001 | GAATTACTTAAATATAGGAA |
| 6211 | RP11-692C24.2-001 | GCCTCACTTTAAGAGATCCT |
| 6212 | RP11-692C24.2-001 | CCCACCAAGATGAAAAAGGT |
| 6213 | RP11-692C24.2-001 | CTCCAGATAACAACTTTCCC |
| 6214 | RP11-697M17.2-001 | GGGGTTTCACCGTGTTAGCC |
| 6215 | RP11-697M17.2-001 | TTTCACCGTGTTAGCCAGGA |
| 6216 | RP11-697M17.2-001 | GCCGAGGCGGGCGGATCACG |
| 6217 | RP11-697M17.2-001 | CAAGCGTGAGCCACCGCGCC |
| 6218 | RP11-697M17.2-001 | ACCTCGTGATCCGCCCGCCT |
| 6219 | RP11-697M17.2-001 | CGCTTGTAATCCCAGCACTT |
| 6220 | RP11-697M17.2-001 | TACAAAAAAAAAAATTAGCC |
| 6221 | RP11-6N17.4-001 | GTTTACGCGACACTTCTAAG |
| 6222 | RP11-6N17.4-001 | AGGGGCGACTTTGTACAGTG |
| 6223 | RP11-6N17.4-001 | ACCTGGGCGAGACTGGACTT |
| 6224 | RP11-6N17.4-001 | CAATGTATATGCTATTTCTG |
| 6225 | RP11-6N17.4-001 | TAAAAGCTCCTGCCACACCT |
| 6226 | RP11-6N17.4-001 | AGTCCAGTCTCGCCCAGGTG |
| 6227 | RP11-6N17.4-001 | CCCTAAGTCCAGTCTCGCCC |
| 6228 | RP11-701H24.4-001 | GTCTGTTCAGGCTTAGAATG |
| 6229 | RP11-701H24.4-001 | AACAAGAAATTTCCTATACA |
| 6230 | RP11-701H24.4-001 | TTATGGGTGATGGGGGTCAC |
| 6231 | RP11-701H24.4-001 | CCCCTGTTGGAGTAATGTGC |
| 6232 | RP11-701H24.4-001 | ACAAGAAATTTCCTATACAT |
| 6233 | RP11-701H24.4-001 | TGGGGGTCACAGGAGAGTGG |
| 6234 | RP11-701H24.4-001 | GAGTGGTGGCTCCCATGTAT |

| | | |
|---|---|---|
| 6235 | RP11-703G6.1-001 | GTTGAATAAGCAGGATCCAA |
| 6236 | RP11-703G6.1-001 | TTCTACAAATGTATAATGGT |
| 6237 | RP11-703G6.1-001 | TGTTGAATAAGCAGGATCCA |
| 6238 | RP11-703G6.1-001 | GGAGAAAGTGTTGAATAAGC |
| 6239 | RP11-703G6.1-001 | TCCATAGAGTGTACAACATC |
| 6240 | RP11-703G6.1-001 | GTCTTCCTTAATGTAAACTA |
| 6241 | RP11-703G6.1-001 | CCATTAAAGAAGTCATAGAG |
| 6242 | RP11-706O15.1-004 | GCCCTCCGAGGCTGCATCCG |
| 6243 | RP11-706O15.1-004 | ACCTGAGTGCGTCCCCATGA |
| 6244 | RP11-706O15.1-004 | AGGCTGCATCCGCGGAAGCC |
| 6245 | RP11-706O15.1-004 | CTCGGGGAGCCGGGCTTCCG |
| 6246 | RP11-706O15.1-004 | GCCTTCATGGGGACGCACTC |
| 6247 | RP11-706O15.1-004 | ATGGGGACGCACTCAGGTCC |
| 6248 | RP11-706O15.1-004 | GCCCGGCTCCCCGAGCGCTC |
| 6249 | RP11-706O15.1-006 | ATAAATGCGACCGGGCGCGG |
| 6250 | RP11-706O15.1-006 | TTAAAGCAATAAATGCGACC |
| 6251 | RP11-706O15.1-006 | ACTCCAGCCTGGGCGACAGA |
| 6252 | RP11-706O15.1-006 | CACCTTGTTAGCCAGGAGGA |
| 6253 | RP11-706O15.1-006 | ATACCACTGCACTCCAGCCT |
| 6254 | RP11-706O15.1-006 | TCGCCCAGGCTGGAGTGCAG |
| 6255 | RP11-706O15.1-006 | CACTCCAGCCTGGGCGACAG |
| 6256 | RP11-706O15.1-008 | CAGATGCAAACATTACCTGC |
| 6257 | RP11-706O15.1-008 | TTGGTGAGAGAGTCAGCACG |
| 6258 | RP11-706O15.1-008 | GTGAGAGAGTCAGCACGCGG |
| 6259 | RP11-706O15.1-008 | CCCGGGGCTGTCCGTGCCAG |
| 6260 | RP11-706O15.1-008 | ACTCTCTCACCAAGCAGGAG |
| 6261 | RP11-706O15.1-008 | TGCTGACTCTCTCACCAAGC |
| 6262 | RP11-706O15.1-008 | GAGCGGGCCCCTCCCTCACG |
| 6263 | RP11-706O15.3-002 | CCCGGGCTTGTAGAGCCAGG |
| 6264 | RP11-706O15.3-002 | GGGCTTGTAGAGCCAGGCGG |
| 6265 | RP11-706O15.3-002 | CGGGCTTGTAGAGCCAGGCG |
| 6266 | RP11-706O15.3-002 | CGCTGCTACCGAGCTGCAGG |
| 6267 | RP11-706O15.3-002 | CATTCCCAGTCTCGCGCCCG |
| 6268 | RP11-706O15.3-002 | GAGGCCGCGGGCGCGAGACT |
| 6269 | RP11-706O15.3-002 | CGGCCCGGGCTTGTAGAGCC |
| 6270 | RP11-707G18.1-001 | GAGCCCGGGGAGACTCCCAA |
| 6271 | RP11-707G18.1-001 | CTCGTATTTAAAGAGCCCTT |
| 6272 | RP11-707G18.1-001 | GAGCCCTTGGGAGTCTCCCC |
| 6273 | RP11-707G18.1-001 | GAGTCTCCCCGGGCTCGGCA |
| 6274 | RP11-707G18.1-001 | AGAGCCCTTGGGAGTCTCCC |
| 6275 | RP11-707G18.1-001 | CGAGCCCGGGGAGACTCCCA |
| 6276 | RP11-707G18.1-001 | CTTGGGAGTCTCCCCGGGCT |
| 6277 | RP11-70P17.1-001 | AGGAGCTCCCGATGTGTCAG |

| | | |
|---|---|---|
| 6278 | RP11-70P17.1-001 | CTTCAGGTGCCCTAGTTGGA |
| 6279 | RP11-70P17.1-001 | GGCTGAGGCTTCTAACGGCA |
| 6280 | RP11-70P17.1-001 | TAGAAGCCTCAGCCCAAGGT |
| 6281 | RP11-70P17.1-001 | TTAGAAGCCTCAGCCCAAGG |
| 6282 | RP11-70P17.1-001 | GGGCTGAGGCTTCTAACGGC |
| 6283 | RP11-70P17.1-001 | TCAGCCCAAGGTGGGAGACA |
| 6284 | RP11-713M6.2-001 | TTCAGATCACTTACAAAAGT |
| 6285 | RP11-713M6.2-001 | CTTTGGTTTAATCAGATACA |
| 6286 | RP11-713M6.2-001 | TGATTATTAATATTCATTTG |
| 6287 | RP11-713M6.2-001 | TACAAGGACAAGTAGACTTG |
| 6288 | RP11-713M6.2-001 | TCATTTGGGGCCATGTGAAA |
| 6289 | RP11-713M6.2-001 | GTTCAGATCACTTACAAAAG |
| 6290 | RP11-713M6.2-001 | ACCAAAAAGATATATAGAAA |
| 6291 | RP11-723O4.9-001 | ACAACACTAGGCACTTGTGG |
| 6292 | RP11-723O4.9-001 | TAAAAGGGTAGCACAACACT |
| 6293 | RP11-723O4.9-001 | GCGGTTACACATGCCAACAT |
| 6294 | RP11-723O4.9-001 | CACAGTCACGACACTGAACA |
| 6295 | RP11-723O4.9-001 | AGCACAACACTAGGCACTTG |
| 6296 | RP11-723O4.9-001 | CATGGGTGGCAAAATCGTCC |
| 6297 | RP11-723O4.9-001 | GTTCAGTGTCGTGACTGTGG |
| 6298 | RP11-727A23.5-004 | ACTTGGCTGGAGAGGCAGCA |
| 6299 | RP11-727A23.5-004 | TGTACTCGTTTCCAAGGCGA |
| 6300 | RP11-727A23.5-004 | GGGAAATGACCTCAGCACGT |
| 6301 | RP11-727A23.5-004 | GCCAAGTGGCTGGAGTCGGG |
| 6302 | RP11-727A23.5-004 | CTGCTGCCTCTCCAGCCAAG |
| 6303 | RP11-727A23.5-004 | TCCAGCCAAGTGGCTGGAGT |
| 6304 | RP11-727A23.5-004 | TGCCTCTCCAGCCAAGTGGC |
| 6305 | RP11-727A23.5-009 | TTGACCAAGATCACACAAGT |
| 6306 | RP11-727A23.5-009 | GATCACACAAGTAGGAGTTG |
| 6307 | RP11-727A23.5-009 | TGTAGTAGATAAATTTACTG |
| 6308 | RP11-727A23.5-009 | TTAAAGGATACACACATGAC |
| 6309 | RP11-727A23.5-009 | ATGACAGGAAGGATTAGAAG |
| 6310 | RP11-727A23.5-009 | AAAGCAGACTCCGGGTGCAG |
| 6311 | RP11-727A23.5-009 | AGGATACACACATGACAGGA |
| 6312 | RP11-72M17.1-003 | CTGTACCCTGATTTACTGAG |
| 6313 | RP11-72M17.1-003 | CCAACCAGCACATCGGACAG |
| 6314 | RP11-72M17.1-003 | CAACCAGCACATCGGACAGT |
| 6315 | RP11-72M17.1-003 | AAAATCAGTAATATTAACAC |
| 6316 | RP11-72M17.1-003 | AGTGGTATGATCTGTGTAGC |
| 6317 | RP11-72M17.1-003 | AACTCAGCCAACCAGCACAT |
| 6318 | RP11-72M17.1-003 | GCACATCGGACAGTGGGAGG |
| 6319 | RP11-731F5.2-001 | CCTGCACAGGAAGGCTTCTG |
| 6320 | RP11-731F5.2-001 | TGGCTCTCACTCATGGTGCA |

| | | |
|---|---|---|
| 6321 | RP11-731F5.2-001 | CTCAGAAGCCTTCCTGTGCA |
| 6322 | RP11-731F5.2-001 | GGGTCCCAGGGCTGGCCGAG |
| 6323 | RP11-731F5.2-001 | GTGCTCCAGAAGGAACAAGA |
| 6324 | RP11-731F5.2-001 | TGCTCCAGAAGGAACAAGAT |
| 6325 | RP11-731F5.2-001 | CCTCAGAAGCCTTCCTGTGC |
| 6326 | RP11-731F5.2-002 | AAGAACAGGATTGAAACCGG |
| 6327 | RP11-731F5.2-002 | TTCTGTGTCACAGGGTAAGG |
| 6328 | RP11-731F5.2-002 | TGTTCTTAGTCTTTGCAGCG |
| 6329 | RP11-731F5.2-002 | CAGGAAGTAAGCGGTGCCGC |
| 6330 | RP11-731F5.2-002 | TTTCTGTGTCACAGGGTAAG |
| 6331 | RP11-731F5.2-002 | CTGGTGGTTTCTGTGTCACA |
| 6332 | RP11-731F5.2-002 | AGGGAACTTCAGGAAGTAAG |
| 6333 | RP11-734K23.9-001 | TTGGCCTCATAGAATGAGTT |
| 6334 | RP11-734K23.9-003 | ACTGCTTTGGGTAGTGTCTG |
| 6335 | RP11-734K23.9-003 | TAGACTAGCTGGACACATGC |
| 6336 | RP11-734K23.9-003 | GTGAAAGACGGCTGTGCTCT |
| 6337 | RP11-734K23.9-003 | TGGCTGAAGGATAGACTAGC |
| 6338 | RP11-734K23.9-003 | TCATCTCTCTGTCATGAGTC |
| 6339 | RP11-734K23.9-003 | AAGAGCACAGCCGTCTTTCA |
| 6340 | RP11-734K23.9-003 | CTGGACACATGCTGGCTGTG |
| 6341 | RP11-735B13.2-001 | GATGAACCAAGTACCCCAGT |
| 6342 | RP11-735B13.2-001 | GCAGGGCAGGGAAGCACAAG |
| 6343 | RP11-735B13.2-001 | ACAGACTACCTGGAAAAACG |
| 6344 | RP11-735B13.2-001 | AGCAGGGCAGGGAAGCACAA |
| 6345 | RP11-735B13.2-001 | AAGCAGGGCAGGGAAGCACA |
| 6346 | RP11-735B13.2-001 | GCGTGATCAATGCAGAAGAA |
| 6347 | RP11-735B13.2-001 | ACAAAGGGTGAGCCGAAGCA |
| 6348 | RP11-757O6.1-001 | GCTCCTTATGGAGAAAGAGT |
| 6349 | RP11-757O6.1-001 | GACTCTTGACCTCGCCTTGT |
| 6350 | RP11-757O6.1-001 | AGCTCCTTATGGAGAAAGAG |
| 6351 | RP11-757O6.1-001 | GAGGTCTGAGGGTTTGAAGT |
| 6352 | RP11-757O6.1-001 | AGCACTCTTCCTACAAGGCG |
| 6353 | RP11-757O6.1-001 | CTGCCAGAACTACTTGACTG |
| 6354 | RP11-757O6.1-001 | TGGCAGTTATTTGAGGTCTG |
| 6355 | RP11-760H22.2-001 | ATGTCATCAGTGCTCTAACG |
| 6356 | RP11-760H22.2-001 | CAGATGCTTAGAGTCTACTT |
| 6357 | RP11-760H22.2-001 | TGCTCTAACGGGGCATAGGG |
| 6358 | RP11-760H22.2-001 | CTAAAAACCCACCTCAAGGG |
| 6359 | RP11-760H22.2-001 | TGTCTAAAAACCCACCTCAA |
| 6360 | RP11-760H22.2-001 | TTGTCTAAAAACCCACCTCA |
| 6361 | RP11-760H22.2-001 | GGCAAACAGACATGTAACAG |
| 6362 | RP11-769N22.1-001 | CCTCTGCTCAACCTCCACTC |
| 6363 | RP11-769N22.1-001 | TTAAACATCAGCTCAGTGAG |

| | | |
|---|---|---|
| 6364 | RP11-769N22.1-001 | TGTTTAAAACAATTCCTGAG |
| 6365 | RP11-769N22.1-001 | AGTGTGTAAGATGCCAGCTG |
| 6366 | RP11-769N22.1-001 | TTAAAACAATTCCTGAGTGG |
| 6367 | RP11-769N22.1-001 | GACTGTTATAGTTATAATCT |
| 6368 | RP11-769N22.1-001 | TACACATGCTCATGCTGCTG |
| 6369 | RP11-775C24.5-001 | CAGAGGAACAAACATTCTGT |
| 6370 | RP11-775C24.5-001 | GGTTCGGTAAAAAGAGGCTG |
| 6371 | RP11-775C24.5-001 | GAACAAGGTTCGGTAAAAAG |
| 6372 | RP11-775C24.5-001 | CAGGAAGACAGCATATGGCA |
| 6373 | RP11-775C24.5-001 | AGGTTCGGTAAAAAGAGGCT |
| 6374 | RP11-775C24.5-001 | TGAAGTGACTTCCGAAGATC |
| 6375 | RP11-775C24.5-001 | GTAAAAAGAGGCTGGGGCGT |
| 6376 | RP11-77H9.8-001 | TCGAACGTAAGCCCGACGCT |
| 6377 | RP11-77H9.8-001 | CTAACAGCACTGCCTAGCGT |
| 6378 | RP11-77H9.8-001 | TAACAGCACTGCCTAGCGTC |
| 6379 | RP11-77H9.8-001 | GGAAGAGCGCTTCCGAGCCA |
| 6380 | RP11-77H9.8-001 | AGTGCTGTTAGAAAGTGATT |
| 6381 | RP11-77H9.8-001 | GCTCCGGTGCGCTCCCCCGG |
| 6382 | RP11-77H9.8-001 | CTCCGGTGCGCTCCCCCGGA |
| 6383 | RP11-783K16.13-001 | CATGGGCCCCCAACCACAGG |
| 6384 | RP11-783K16.13-001 | CTTATGGTCCTGACCTCCTG |
| 6385 | RP11-783K16.13-001 | TCCTGACCTCCTGTGGTTGG |
| 6386 | RP11-783K16.13-001 | GGGCATGGGCCCCCAACCAC |
| 6387 | RP11-783K16.13-001 | CTCCAGTCCCAGTAGGGCTC |
| 6388 | RP11-783K16.13-001 | AGTCCCAGTAGGGCTCAGGC |
| 6389 | RP11-783K16.13-001 | GTCCTGACCTCCTGTGGTTG |
| 6390 | RP11-791G15.2-001 | AGAAGTGGATTGCAGAAGTG |
| 6391 | RP11-791G15.2-001 | AAGCAAAGAGAATGTTACCT |
| 6392 | RP11-791G15.2-001 | TCAGAAGTGGATTGCAGAAG |
| 6393 | RP11-791G15.2-001 | ACAGTAGTGGGCCAATCAGG |
| 6394 | RP11-791G15.2-001 | CTTCTGCAATCCACTTCTGA |
| 6395 | RP11-791G15.2-001 | TAAGCAAAGAGAATGTTACC |
| 6396 | RP11-791G15.2-001 | CAGAAGTGGATTGCAGAAGT |
| 6397 | RP11-802F5.1-001 | CGGGGCGGTTGCCAGGCAGA |
| 6398 | RP11-802F5.1-001 | TCACTTCCTAGATGGGATGG |
| 6399 | RP11-802F5.1-001 | GCCATCCCATCTAGGAAGTG |
| 6400 | RP11-802F5.1-001 | TGCCCATCGTCTGGGATGTG |
| 6401 | RP11-802F5.1-001 | AGCCCATCGTCTGAGATGTG |
| 6402 | RP11-802F5.1-001 | GCCGCCCCGTCTGAGAAGTG |
| 6403 | RP11-802F5.1-001 | TCCTCACTTCTCAGACGGGG |
| 6404 | RP11-804H8.6-002 | AACACAAAGGCTGTACACAG |
| 6405 | RP11-804H8.6-002 | TTCAAAGTAAGTTAACACAA |
| 6406 | RP11-804H8.6-002 | CAGTCATGGTGGTATGCGCC |

| | | |
|---|---|---|
| 6407 | RP11-804H8.6-002 | AGATAAAGTTAACACAGAGA |
| 6408 | RP11-804H8.6-002 | TGAGGCAGGCAGATGACTTG |
| 6409 | RP11-804H8.6-002 | CCAGAAGTTCAAGACCAGCC |
| 6410 | RP11-804H8.6-002 | CACTTGTAATCCCAGCACTT |
| 6411 | RP11-817O13.8-001 | CTGGCCCCAAAAAGCCTGCG |
| 6412 | RP11-817O13.8-001 | GCCGCCCCGTTTCCGCCTGG |
| 6413 | RP11-817O13.8-001 | CTGGCCCCCAGGCGGAAACG |
| 6414 | RP11-817O13.8-001 | TGGCCCCAAAAAGCCTGCGT |
| 6415 | RP11-817O13.8-001 | CCGGCCAGCTGGCCCCCAGG |
| 6416 | RP11-817O13.8-001 | GGCCGGTGGGCACATGCTGG |
| 6417 | RP11-817O13.8-001 | GGCCTCCAGCATGTGCCCAC |
| 6418 | RP11-81H14.1-001 | TTTGGTCACCTACATGTCAA |
| 6419 | RP11-81H14.1-001 | CCACATGCTATGATGCAGTG |
| 6420 | RP11-81H14.1-001 | CCACACTGCATCATAGCATG |
| 6421 | RP11-81H14.1-001 | AGCCTGAGAGCAAATAAGGG |
| 6422 | RP11-81H14.1-001 | TAAGACCACCTTTGACATGT |
| 6423 | RP11-81H14.1-001 | TTACAATGAGCTCAAGTGTA |
| 6424 | RP11-81H14.1-001 | TTTACAATGAGCTCAAGTGT |
| 6425 | RP11-81H14.2-001 | AGCACAACCAAAAGGCACCA |
| 6426 | RP11-81H14.2-001 | AGGTACTGTACCATCTTCAC |
| 6427 | RP11-81H14.2-001 | GTGAAGTGTGAATCCTTGAG |
| 6428 | RP11-81H14.2-001 | TTGTTCTACACATTCAGGCA |
| 6429 | RP11-81H14.2-001 | ACTTTCTGATATCTTCTAAG |
| 6430 | RP11-81H14.2-001 | TGTAGAACAAGAACAGCTGG |
| 6431 | RP11-81H14.2-001 | AGCAATCAACAGCAAAGATC |
| 6432 | RP11-81H14.2-002 | ACCATGTAAAAATGTGACTC |
| 6433 | RP11-81H14.2-002 | ACAGATAAGCAAAGACCTCC |
| 6434 | RP11-81H14.2-002 | GACTCAAGCAACTCAGAGTT |
| 6435 | RP11-81H14.2-002 | CTTTCTGGTGCTCTGCCTGG |
| 6436 | RP11-81H14.2-002 | CAAGGATTCACACTTCACAC |
| 6437 | RP11-81H14.2-002 | GCTCTGCCTGCAGGTCAAGA |
| 6438 | RP11-81H14.2-002 | AAGGAGCCAGTATCTCCAGG |
| 6439 | RP11-81H14.2-003 | CAGTGATTGAGTTATGCTGT |
| 6440 | RP11-81H14.2-003 | AGGGCAGATCAGTCACCCTG |
| 6441 | RP11-81H14.2-003 | GAGGCAGATAAAATTCCCCA |
| 6442 | RP11-81H14.2-003 | ACAGTGATTGAGTTATGCTG |
| 6443 | RP11-81H14.2-003 | AAGGGCAGATCAGTCACCCT |
| 6444 | RP11-81H14.2-003 | TGAGGCAGATAAAATTCCCC |
| 6445 | RP11-81H14.2-003 | GAAGGGCAGATCAGTCACCC |
| 6446 | RP11-81H14.2-004 | TAAGAGAGCTAGACAAACGC |
| 6447 | RP11-81H14.2-004 | CTCCTGAATAGGTATTGGAA |
| 6448 | RP11-81H14.2-004 | ACAACTGCAGAGAAACTCTC |
| 6449 | RP11-81H14.2-004 | TGGAGGCTGTTGTGAGCATG |

| | | |
|---|---|---|
| 6450 | RP11-81H14.2-004 | AGAGGTAATAATAACTAATG |
| 6451 | RP11-81H14.2-004 | AAGGAAGAGCACCTGGTACA |
| 6452 | RP11-81H14.2-004 | TGCCTTTCCAATACCTATTC |
| 6453 | RP11-81H14.2-005 | TATAATTCCAGCATGACACC |
| 6454 | RP11-81H14.2-005 | AGCTACTGAGAGATAAACTG |
| 6455 | RP11-81H14.2-005 | CTAGTAGCTTTGGATGTGAG |
| 6456 | RP11-81H14.2-005 | AGCACTGAGGAAGTTGGGCA |
| 6457 | RP11-81H14.2-005 | CAGCTACTGAGAGATAAACT |
| 6458 | RP11-81H14.2-005 | GTCACAAACTCCCATGAAGT |
| 6459 | RP11-81H14.2-005 | ATCCAAAGCTACTAGCACTG |
| 6460 | RP11-81H14.2-006 | AAAAGGTCCTTCTTACACGG |
| 6461 | RP11-81H14.2-006 | GGAGGACTTAGAATCATGGT |
| 6462 | RP11-81H14.2-006 | TCAATATCACGGAATAGCAT |
| 6463 | RP11-81H14.2-006 | GGCGGCAGCAAGAGAAACTG |
| 6464 | RP11-81H14.2-006 | GGGAGGACTTAGAATCATGG |
| 6465 | RP11-81H14.2-006 | ATTGGACTTACAGTTCCACG |
| 6466 | RP11-81H14.2-006 | GAATAACCTAAAGACTCTAC |
| 6467 | RP11-81H3.2-001 | CCCCTTACTCAACCCAACCA |
| 6468 | RP11-81H3.2-001 | TGAAGGCTGTGCGGACACTG |
| 6469 | RP11-81H3.2-001 | GTAAGGGGCCAAACCCCTGG |
| 6470 | RP11-81H3.2-001 | GCCTGCGCCGGAGAGGACCG |
| 6471 | RP11-81H3.2-001 | CCGTGGTTGGGTTGAGTAAG |
| 6472 | RP11-81H3.2-001 | ACCGTGGTTGGGTTGAGTAA |
| 6473 | RP11-81H3.2-001 | GGACACTGGGGGCCCGCCAG |
| 6474 | RP11-848P1.3-001 | AAAAAATATTAGATTGACAA |
| 6475 | RP11-848P1.3-001 | AAAAATACAATAACAGGGCA |
| 6476 | RP11-848P1.3-001 | AATACAATAACAGGGCATGG |
| 6477 | RP11-848P1.3-001 | TGATATACTCAGAAAGGACT |
| 6478 | RP11-848P1.3-001 | GTTTACTGATATACTCAGAA |
| 6479 | RP11-848P1.3-001 | TATTTGTGGGACTGAAGCGA |
| 6480 | RP11-848P1.3-001 | CTGATATACTCAGAAAGGAC |
| 6481 | RP11-863P13.2-001 | TAGCCTGGGCACTACACAGG |
| 6482 | RP11-863P13.2-001 | AAATTGGAGGCCAAGCATGG |
| 6483 | RP11-863P13.2-001 | CGTAAATTGGAGGCCAAGCA |
| 6484 | RP11-863P13.2-001 | GATTAGCCTGGGCACTACAC |
| 6485 | RP11-863P13.2-001 | CACGTGTAATCCCAGCACTT |
| 6486 | RP11-863P13.2-001 | TGTAATCCCAGCACTTCGGA |
| 6487 | RP11-863P13.2-001 | AACAAAATACCGTAAATTGG |
| 6488 | RP11-865I6.2-001 | TAAAATGACTGAGACAACAT |
| 6489 | RP11-865I6.2-001 | CCAGGAAGAATAGACAGACA |
| 6490 | RP11-865I6.2-001 | TGAGGGGAGAAACTCACCAA |
| 6491 | RP11-865I6.2-001 | AGTGTCTATTGTTCATGGTT |
| 6492 | RP11-865I6.2-001 | GAATCTGAACAGACAGCCTT |

| | | |
|---|---|---|
| 6493 | RP11-865I6.2-001 | ATCTAATAGTAATAAACTGA |
| 6494 | RP11-865I6.2-001 | AAGGGTCCTGTCCTGTACCC |
| 6495 | RP11-867G23.3-001 | GAGGGAGGCGCACAGCACCA |
| 6496 | RP11-867G23.3-001 | GATGACAAAGAAGATGAGGG |
| 6497 | RP11-867G23.3-001 | CCAGATGACAAAGAAGATGA |
| 6498 | RP11-867G23.3-001 | GCCAGATGACAAAGAAGATG |
| 6499 | RP11-867G23.3-001 | GGTGAGCATGTAGCAGAACG |
| 6500 | RP11-867G23.3-001 | TGAGGGAGGCGCACAGCACC |
| 6501 | RP11-867G23.3-001 | CATGTAGCAGAACGCGGCGA |
| 6502 | RP11-87N24.3-001 | AATTTCAGCTGAAACAAACC |
| 6503 | RP11-87N24.3-001 | ATAGATACCCTGGTTTGAGT |
| 6504 | RP11-87N24.3-001 | ATTATATCCCACTCAAACCA |
| 6505 | RP11-87N24.3-001 | CCAATGTGTCATAGATACCC |
| 6506 | RP11-87N24.3-001 | ATCTATGACACATTGGAACA |
| 6507 | RP11-87N24.3-001 | CCAGGGTATCTATGACACAT |
| 6508 | RP11-87N24.3-001 | ATGACACATTGGAACATGGC |
| 6509 | RP11-883G14.4-001 | GAGAAGGAGATTCCACGGTA |
| 6510 | RP11-883G14.4-001 | AGAAGGAGATTCCACGGTAA |
| 6511 | RP11-883G14.4-001 | AAAAGTAATAGCTTATGCCC |
| 6512 | RP11-883G14.4-001 | AAGGTACTGTTTGAAAGAGA |
| 6513 | RP11-883G14.4-001 | GAAGGGACAGGAGCTTAAGA |
| 6514 | RP11-883G14.4-001 | CATTTGGGGGGCCCTTACCG |
| 6515 | RP11-883G14.4-001 | TGAAAGAGAAGGAGATTCCA |
| 6516 | RP11-890B15.2-001 | CCTGTGTCCACTGAGAACTG |
| 6517 | RP11-890B15.2-001 | TGTGGATGTGGAATATGCCC |
| 6518 | RP11-890B15.2-001 | GACACAGGGTCTGATGCCCG |
| 6519 | RP11-890B15.2-001 | CCACAGTTCTCAGTGGACAC |
| 6520 | RP11-890B15.2-001 | TCCACATCCACAGTTCTCAG |
| 6521 | RP11-890B15.2-001 | AGCCGTGCTCCAGTAAGTTA |
| 6522 | RP11-890B15.2-001 | CACAGTTCTCAGTGGACACA |
| 6523 | RP11-89K11.1-001 | GCCGCCGTCGCCACGCACCG |
| 6524 | RP11-89K11.1-001 | GCCGTCGCCACGCACCGAGG |
| 6525 | RP11-89K11.1-001 | GTGGCGACGGCGGCCAAGAG |
| 6526 | RP11-89K11.1-001 | CCGTCGCCACGCACCGAGGT |
| 6527 | RP11-89K11.1-001 | ACCTCGGTGCGTGGCGACGG |
| 6528 | RP11-89K11.1-001 | GCGTGGCGACGGCGGCCAAG |
| 6529 | RP11-89K11.1-001 | CGGCGGCCAAGAGGGGCCAG |
| 6530 | RP11-89K11.1-002 | GAAAACGGCCTCGTTGACGG |
| 6531 | RP11-89K11.1-002 | GATCGCTCAAACAGCACCAG |
| 6532 | RP11-89K11.1-002 | AATGAAAACGGCCTCGTTGA |
| 6533 | RP11-89K11.1-002 | TGTTTGTGCCTTACAGTGAC |
| 6534 | RP11-89K11.1-002 | CTCGTTGACGGAGGAAAAGT |
| 6535 | RP11-89K11.1-002 | TGTCACAAGAAAAGAAGGAA |

| | | |
|---|---|---|
| 6536 | RP11-89K11.1-002 | GTTTGCAGCCTGTCACTGTA |
| 6537 | RP11-90C4.1-003 | GAAGGACTCGCTCATCCATC |
| 6538 | RP11-90C4.1-003 | CCAGTGACTCATTTCACTCA |
| 6539 | RP11-90C4.1-003 | GGTGGAAAGAACTCTAGTTC |
| 6540 | RP11-90C4.1-003 | ACTAGAGCAGTGTTGTATGG |
| 6541 | RP11-90C4.1-003 | TAGCTGTGAAATATGTCCCT |
| 6542 | RP11-90C4.1-003 | AGTGCTAAATAAATACCTGA |
| 6543 | RP11-90C4.1-003 | CCTTGAGTGAAATGAGTCAC |
| 6544 | RP11-91G21.1-001 | CGTGCACTGGCCATCTAGGG |
| 6545 | RP11-91G21.1-001 | GTGACTCCCAGGAAAACTAG |
| 6546 | RP11-91G21.1-001 | GAAAACTAGCGGAGTGAGCG |
| 6547 | RP11-91G21.1-001 | AAAATAGGACTGGAGCTGAC |
| 6548 | RP11-91G21.1-001 | TAAGAGGGTGAAGTATCAGC |
| 6549 | RP11-91G21.1-001 | CTTGGTCGCGTTATTGGGTC |
| 6550 | RP11-91G21.1-001 | TCCGGGAATTCAGAGAATCA |
| 6551 | RP11-92G12.3-001 | TGCCTCCCACCCACGCCCGG |
| 6552 | RP11-92G12.3-001 | CCGTGACGTGAAGCGCCTCG |
| 6553 | RP11-92G12.3-001 | CGTTGCCTCCCACCCACGCC |
| 6554 | RP11-92G12.3-001 | ACGTGAAGCGCCTCGGGGTG |
| 6555 | RP11-92G12.3-001 | CGGTGCCGCCGGGCGTGGGT |
| 6556 | RP11-92G12.3-001 | TGGGTGGGAGGCAACGCCCG |
| 6557 | RP11-92G12.3-001 | TGCCGCCGGGCGTGGGTGGG |
| 6558 | RP11-92K15.3-001 | CTGCAATGTGGTGACCGGCG |
| 6559 | RP11-92K15.3-001 | CCTTGGGGTTCCCCAAGCCA |
| 6560 | RP11-92K15.3-001 | CCCGCCGGTCACCACATTGC |
| 6561 | RP11-92K15.3-001 | GTTGCCTGCAATGTGGTGAC |
| 6562 | RP11-92K15.3-001 | CTTGGGGAACCCCAAGGTGT |
| 6563 | RP11-92K15.3-001 | GAGAAGAGTTGAGCCCTTCT |
| 6564 | RP11-92K15.3-001 | CCCTGGCTTGGGGAACCCCA |
| 6565 | RP11-932O9.9-001 | AGAAATTAAAATGTATTCAG |
| 6566 | RP11-932O9.9-001 | CACAAATACTCGGCTGAATG |
| 6567 | RP11-932O9.9-001 | TGAGGAAGAGCACAAATACT |
| 6568 | RP11-932O9.9-001 | GCAAAAGACTGCATGCACTT |
| 6569 | RP11-932O9.9-001 | CTATGGATTAAATCACACAC |
| 6570 | RP11-932O9.9-001 | ATTTAAGCTGAAGGTCAGTC |
| 6571 | RP11-932O9.9-001 | CACTGGCATATTTAAGCTGA |
| 6572 | RP11-944C7.1-002 | ATCATATGGCAACATCACTG |
| 6573 | RP11-944C7.1-002 | GTTGTACAAAGATTCATCTG |
| 6574 | RP11-944C7.1-002 | ATGTTGCCATATGATGTCCT |
| 6575 | RP11-944C7.1-002 | AAAAAAAAAAGATTCATCTG |
| 6576 | RP11-944C7.1-002 | GATTTGAGAGATTCCCAGAA |
| 6577 | RP11-944C7.1-002 | TGTTGTACAAAGATTCATCT |
| 6578 | RP11-944C7.1-002 | AGCCCTGGCGTTCAGAAGAC |

| | | |
|---|---|---|
| 6579 | RP11-960L18.1-001 | ACCCAGACACTGAGTACCAG |
| 6580 | RP11-960L18.1-001 | GCCACTGGTACTCAGTGTCT |
| 6581 | RP11-960L18.1-001 | GGTACTCAGTGTCTGGGTGG |
| 6582 | RP11-960L18.1-001 | GTCGGGCCTTCTCCCTCCAG |
| 6583 | RP11-960L18.1-001 | CTGGTACTCAGTGTCTGGGT |
| 6584 | RP11-960L18.1-001 | ACTGGTACTCAGTGTCTGGG |
| 6585 | RP11-960L18.1-001 | AGCCACTGGTACTCAGTGTC |
| 6586 | RP11-96H19.1-001 | CCGACCCCCACACACCCGGG |
| 6587 | RP11-96H19.1-001 | GGAGGCGTTGGCAGGTACCT |
| 6588 | RP11-96H19.1-001 | ACGATGAGGTGCGGAGGCGT |
| 6589 | RP11-96H19.1-001 | GACCCGACCCCCACACACCC |
| 6590 | RP11-96H19.1-001 | CCCGCCGCCCGGGTGTGTGG |
| 6591 | RP11-96H19.1-001 | TGAGGTGCGGAGGCGTTGGC |
| 6592 | RP11-96H19.1-001 | GCCCCGCCGCCCGGGTGTGT |
| 6593 | RP11-96H19.1-002 | TCCTGCAAGAGCAGAGTCAT |
| 6594 | RP11-96H19.1-002 | GGCGTGCAGTTGGTTCCGTA |
| 6595 | RP11-96H19.1-002 | CCACGCTCACCGAAGCTTCA |
| 6596 | RP11-96H19.1-002 | TTATTTCAAAAATACTGCAA |
| 6597 | RP11-96H19.1-002 | AGTCTTAATGGGGAACCATA |
| 6598 | RP11-96H19.1-002 | CCTGAAGCTTCGGTGAGCGT |
| 6599 | RP11-96H19.1-002 | CGTGGGATTTGGCGTGCAGT |
| 6600 | RP11-973H7.5-001 | TCATGATAGAGATTTGAGTG |
| 6601 | RP11-973H7.5-001 | CGTGGCAAGGACTCGGAATG |
| 6602 | RP11-973H7.5-001 | TCGGAATGTGGTGAAGACAG |
| 6603 | RP11-973H7.5-001 | GCCATCCCAAGGCCGTGGCA |
| 6604 | RP11-973H7.5-001 | GACTTTGCACAGCCATCCCA |
| 6605 | RP11-973H7.5-001 | ATAGAGATTTGAGTGAGGCT |
| 6606 | RP11-973H7.5-001 | CACATTCCGAGTCCTTGCCA |
| 6607 | RP11-98O2.1-002 | ACTTCTCAGAGATCAACTGT |
| 6608 | RP11-98O2.1-002 | CACTTCTCAGAGATCAACTG |
| 6609 | RP11-98O2.1-002 | AGGATACCAGCCAAGCCTAA |
| 6610 | RP11-98O2.1-002 | CTTCCTTTAGAAATTGAAAG |
| 6611 | RP11-98O2.1-002 | TCTGAGAAGTGAGTAGCTGG |
| 6612 | RP11-98O2.1-002 | CCTTTAGAAATTGAAAGAGG |
| 6613 | RP11-98O2.1-002 | AATTGAAAGAGGAGGAGAGG |
| 6614 | RP11-9M16.2-001 | AGGAAACCACCCGCGCTCGG |
| 6615 | RP11-9M16.2-001 | TCCAGGAAACCACCCGCGCT |
| 6616 | RP11-9M16.2-001 | CCGCCAGCAGGTGAGACTCG |
| 6617 | RP11-9M16.2-001 | CCCCGAGTCTCACCTGCTGG |
| 6618 | RP11-9M16.2-001 | GGTGGTTTCCTGGAGTTGAG |
| 6619 | RP11-9M16.2-001 | CCAGCAGGTGAGACTCGGGG |
| 6620 | RP11-9M16.2-001 | TTTGTTTGCCGCTCAACTCC |
| 6621 | RP13-188A5.1-001 | GTAATGAAAAGAGATGCGGT |

| | | |
|---|---|---|
| 6622 | RP13-188A5.1-001 | CAAAGTAATGAAAAGAGATG |
| 6623 | RP13-188A5.1-001 | TAATGAAAAGAGATGCGGTG |
| 6624 | RP13-188A5.1-001 | GCACCCAAACTTCTTCCGGC |
| 6625 | RP13-188A5.1-001 | CCCAAACTTCTTCCGGCAGG |
| 6626 | RP13-188A5.1-001 | AGTAATGAAAAGAGATGCGG |
| 6627 | RP13-188A5.1-001 | CACCCAAACTTCTTCCGGCA |
| 6628 | RP13-20L14.1-001 | CTCAAACGTAGTCACGCCTG |
| 6629 | RP13-20L14.1-001 | ACTTCTTAATAGACCAGGCG |
| 6630 | RP13-20L14.1-001 | TTCAAACTTCTTAATAGACC |
| 6631 | RP13-20L14.1-001 | GCACAGCACTCACCGACCCC |
| 6632 | RP13-20L14.1-001 | CCCAGGCGTGACTACGTTTG |
| 6633 | RP13-20L14.1-001 | TCTTAATAGACCAGGCGTGG |
| 6634 | RP13-20L14.1-001 | AACGTAGTCACGCCTGGGGT |
| 6635 | RP13-212L9.1-001 | CTTGTAGGGAGGACGATTGG |
| 6636 | RP13-212L9.1-001 | GTGGTGCAGAGAGATTCTCT |
| 6637 | RP13-212L9.1-001 | GCACCACGTATTCCAGCTGG |
| 6638 | RP13-212L9.1-001 | TCTGCACCACGTATTCCAGC |
| 6639 | RP13-212L9.1-001 | CCACCTCCAGCTGGAATACG |
| 6640 | RP13-212L9.1-001 | AACTTGGTGTCCTTGTAGGG |
| 6641 | RP13-212L9.1-001 | CACCAATCGTCCTCCCTACA |
| 6642 | RP13-216E22.4-001 | ATGCCATCACTATACATAGT |
| 6643 | RP13-216E22.4-001 | AATCCTACTATGTATAGTGA |
| 6644 | RP13-216E22.4-001 | CATTAAGGAATGGCCGGGCG |
| 6645 | RP13-216E22.4-001 | TTAAACATTAAGGAATGGCC |
| 6646 | RP13-216E22.4-001 | TAAGGAATGGCCGGGCGCGG |
| 6647 | RP13-216E22.4-001 | TTAGGAGTTCCAGACCAGCC |
| 6648 | RP13-216E22.4-001 | GTTAAACATTAAGGAATGGC |
| 6649 | RP13-259N13.2-001 | AAAAAATACAAATAATTAGC |
| 6650 | RP13-259N13.2-001 | AAATAATTAGCCGGTTGTGG |
| 6651 | RP13-259N13.2-001 | TATGAGACTCAGTTCACTGA |
| 6652 | RP13-259N13.2-001 | TGTAATCCCAGCTACTCAGA |
| 6653 | RP13-259N13.2-001 | GAGAATTGCTTGAATCCGGG |
| 6654 | RP13-259N13.2-001 | CCCAGCTACTCAGAAGGCTG |
| 6655 | RP13-259N13.2-001 | GGAGTGCAGTGGTATGATCT |
| 6656 | RP13-297E16.4-001 | CATCTCGTCATGGACAAGGT |
| 6657 | RP13-297E16.4-001 | TCCACATCACAGGCACAGGT |
| 6658 | RP13-297E16.4-001 | CACATTCTAAAGGTGACTGT |
| 6659 | RP13-297E16.4-001 | GAACAGTGGACACTGTGTTG |
| 6660 | RP13-297E16.4-001 | GGAAAATAAATATAAACCCA |
| 6661 | RP13-297E16.4-001 | GTGAACAGTGGACACTGTGT |
| 6662 | RP13-297E16.4-001 | GTCCACTGTTCACATTCTAA |
| 6663 | RP13-516M14.1-001 | AGCTGTAAAACGGATGACTT |
| 6664 | RP13-516M14.1-001 | CTTTCACCGCAGAATCCAGG |

| | | |
|---|---|---|
| 6665 | RP13-516M14.1-001 | TAAAACGGATGACTTAGGCC |
| 6666 | RP13-516M14.1-001 | GTTGATCTGCCTGACGAGAT |
| 6667 | RP13-516M14.1-001 | AACCCCTGAGCAGAACGCCA |
| 6668 | RP13-516M14.1-001 | TGACGAGATGGGCTGATGCA |
| 6669 | RP13-516M14.1-001 | AGAACGTCAAACATCTGGCA |
| 6670 | RP3-324O17.8-001 | ATGAGCGGAAGAGCCACTCG |
| 6671 | RP3-324O17.8-001 | GCGTACAATCAGCATCTCTG |
| 6672 | RP3-324O17.8-001 | CGCTCATGTAAATACGTCCT |
| 6673 | RP3-324O17.8-001 | CTAATCATCCAGAGTGACCT |
| 6674 | RP3-324O17.8-001 | AAATGCGCAGCACAATCTGT |
| 6675 | RP3-324O17.8-001 | CTAGGACGTATTTACATGAG |
| 6676 | RP3-324O17.8-001 | TACAATCAGCATCTCTGTGG |
| 6677 | RP3-388N13.3-001 | AAACGCTTACCTACTTTGTG |
| 6678 | RP3-388N13.3-001 | CAATTATTCTATGACTCAGT |
| 6679 | RP3-388N13.3-001 | CCAATTATTCTATGACTCAG |
| 6680 | RP3-388N13.3-001 | GTGTTATGCAATATGCAGCT |
| 6681 | RP3-388N13.3-001 | TTATTTAAACCGCACAAAGT |
| 6682 | RP3-388N13.3-001 | TGTGCGTGCAAGAAGAACAA |
| 6683 | RP3-388N13.3-001 | GCGTGCAAGAAGAACAAAGG |
| 6684 | RP3-395M20.12-001 | TAGCGCAGGGCCCTCCTGCG |
| 6685 | RP3-395M20.12-001 | CCCTGCCTGGCCCCGCAGGA |
| 6686 | RP3-395M20.12-001 | GGACAGCAGGGAGAGCAGAA |
| 6687 | RP3-395M20.12-001 | CCCCTGCCTGGCCCCGCAGG |
| 6688 | RP3-395M20.12-001 | GGGAGAGCAGAAAGGTGGCA |
| 6689 | RP3-395M20.12-001 | CAGCAGGGAGAGCAGAAAGG |
| 6690 | RP3-395M20.12-001 | GGGGCCAGGCAGGGGCGGGT |
| 6691 | RP3-460G2.2-001 | CCTCATCTGAAATGTAGGTG |
| 6692 | RP3-460G2.2-001 | CTTGAACAAAGTATTACAAG |
| 6693 | RP3-460G2.2-001 | TTGAACAAAGTATTACAAGA |
| 6694 | RP3-460G2.2-001 | ATTCTGAAAACATTTAGTGA |
| 6695 | RP3-460G2.2-001 | GCTTCTTCCATTCATAAGCT |
| 6696 | RP3-460G2.2-001 | GACATTCTTTCAGATGCCAG |
| 6697 | RP3-460G2.2-001 | CGTCTTGAAACCAGTCAAGT |
| 6698 | RP3-465N24.6-001 | CGGGTCTGCGACTTGCCGGT |
| 6699 | RP3-465N24.6-001 | GGTGGCGGCAACGACCCTGT |
| 6700 | RP3-465N24.6-001 | CCGGGTCTGCGACTTGCCGG |
| 6701 | RP3-465N24.6-001 | CTGACGGGCTGGTGCCCACA |
| 6702 | RP3-465N24.6-001 | GGGTGGCGGCAACGACCCTG |
| 6703 | RP3-465N24.6-001 | TCTGACGGGCTGGTGCCCAC |
| 6704 | RP3-465N24.6-001 | GACGGCCCCAGCTCCGAGCC |
| 6705 | RP4-535B20.1-001 | TAAAAACAGAAAAATGGCGA |
| 6706 | RP4-535B20.1-001 | TTCTTCCCGCTATTTGCATG |
| 6707 | RP4-535B20.1-001 | GATTATCTAATGACGCAGCA |

| | | |
|---|---|---|
| 6708 | RP4-535B20.1-001 | ATAATCCTCATGCAAATAGC |
| 6709 | RP4-535B20.1-001 | AAATGGCGACGGTCACGTTG |
| 6710 | RP4-535B20.1-001 | GATAATCCTCATGCAAATAG |
| 6711 | RP4-535B20.1-001 | GCAAATAGCGGGAAGAACAA |
| 6712 | RP4-563E14.1-001 | TTCTCACCTGGGATTCAGAG |
| 6713 | RP4-563E14.1-001 | TAAAACGTCCTGACGCTGTC |
| 6714 | RP4-563E14.1-001 | CCTCTCCAAGAACCGGAAGC |
| 6715 | RP4-563E14.1-001 | CCGCTGCCTCTCTGAATCCC |
| 6716 | RP4-563E14.1-001 | CCTGGGATTCAGAGAGGCAG |
| 6717 | RP4-563E14.1-001 | AGAGGCAGCGGCCAAGGACA |
| 6718 | RP4-563E14.1-001 | ACGCTGCCCGGTGGCCTCGG |
| 6719 | RP4-616B8.5-001 | CAAATCTGACAGTAGACAGA |
| 6720 | RP4-616B8.5-001 | CACCCCAAGACTGATGCAGG |
| 6721 | RP4-616B8.5-001 | ATTACTGGTTTCACTCCCAG |
| 6722 | RP4-616B8.5-001 | GACCACCTGCATCAGTCTTG |
| 6723 | RP4-616B8.5-001 | AAGCACCCCAAGACTGATGC |
| 6724 | RP4-616B8.5-001 | GATTACTGGTTTCACTCCCA |
| 6725 | RP4-616B8.5-001 | CCAAATCTGACAGTAGACAG |
| 6726 | RP4-665J23.1-005 | TAGAGGCCTCCCTTCCGAGG |
| 6727 | RP4-665J23.1-005 | CGGTAGAGGCCTCCCTTCCG |
| 6728 | RP4-665J23.1-005 | GCTGGGGCTCCGCTCCCCGG |
| 6729 | RP4-665J23.1-005 | CGGGCGCAGGACTGCAGGCT |
| 6730 | RP4-665J23.1-005 | CCACGCGCGGGCTCCCGCCG |
| 6731 | RP4-665J23.1-005 | ACCACGCGCGGGCTCCCGCC |
| 6732 | RP4-665J23.1-005 | CCGAGGAGGCGGTGCTGGGG |
| 6733 | RP4-665J23.4-001 | AGGTACTTCTTACATAGCGG |
| 6734 | RP4-665J23.4-001 | GAAAGGTACTTCTTACATAG |
| 6735 | RP4-665J23.4-001 | GGAGGCCTTAGAATTGTGGC |
| 6736 | RP4-665J23.4-001 | ATAGCGGTGGCAAGAGAATG |
| 6737 | RP4-665J23.4-001 | GGCCTTAGAATTGTGGCAGG |
| 6738 | RP4-665J23.4-001 | AGAGCTGATGGGTTTATCAG |
| 6739 | RP4-665J23.4-001 | CTGGGGAGGCCTTAGAATTG |
| 6740 | RP4-669B10.3-002 | AGGGCTACCTGAAAGACCGG |
| 6741 | RP4-669B10.3-002 | ATGGGATTGTAATTGTACTT |
| 6742 | RP4-669B10.3-002 | TAGTAACATGATCTCTGTGT |
| 6743 | RP4-669B10.3-002 | CCCATCCCACAAAAAGGTCA |
| 6744 | RP4-669B10.3-002 | CCATCCCACAAAAAGGTCAA |
| 6745 | RP4-669B10.3-002 | GAATTTGTGGAGAGGATGCA |
| 6746 | RP4-669B10.3-002 | TTGTATCTGAATTTGTGGAG |
| 6747 | RP4-669L17.10-004 | TACACACACAGTAAAAATCT |
| 6748 | RP4-669L17.10-004 | GTCAATTTACATCAAGGGTA |
| 6749 | RP4-669L17.10-004 | GTGCTAAGATTATAGCTGGG |
| 6750 | RP4-669L17.10-004 | AAAGTGCTAAGATTATAGCT |

| | | |
|---|---|---|
| 6751 | RP4-669L17.10-004 | TTACTGTGAGTCATGGTGAA |
| 6752 | RP4-669L17.10-004 | TCACAGTAATCTTACTTCTG |
| 6753 | RP4-669L17.10-004 | ACCCTTGATGTAAATTGACA |
| 6754 | RP4-669L17.10-005 | CAGGTGGGAACATCCAGTTG |
| 6755 | RP4-669L17.10-005 | TGAGGCAGTAGGCTCATCTG |
| 6756 | RP4-669L17.10-005 | GGAGGCTGTTGTGAGGCAGT |
| 6757 | RP4-669L17.10-005 | GGCAGTAGGCTCATCTGCGG |
| 6758 | RP4-669L17.10-005 | ACCAAGCTCATGACTCACAA |
| 6759 | RP4-669L17.10-005 | ATGTTCCCACCTGAGCGTGA |
| 6760 | RP4-669L17.10-005 | ATAGGTATTAGACTCTCATA |
| 6761 | RP4-669L17.10-014 | CAGTTTGCGCTATCCCATCC |
| 6762 | RP4-669L17.10-014 | TAGCGCAAACTGCCCGGGAA |
| 6763 | RP4-669L17.10-014 | TCCCATTGCACCTGATGCAT |
| 6764 | RP4-669L17.10-014 | CATGAAAGACTTGTGAATCC |
| 6765 | RP4-669L17.10-014 | ATAGCGCAAACTGCCCGGGA |
| 6766 | RP4-669L17.10-014 | TCCCATGCATCAGGTGCAAT |
| 6767 | RP4-669L17.10-014 | TTCCCATGCATCAGGTGCAA |
| 6768 | RP4-669L17.10-015 | ACCTTCATAACATACTTGAG |
| 6769 | RP4-669L17.10-015 | GGAGGTAGACCCACACAGAT |
| 6770 | RP4-669L17.10-015 | ATGTTAGGAAGGGTCCCAGG |
| 6771 | RP4-669L17.10-015 | AGGAGGTAGACCCACACAGA |
| 6772 | RP4-669L17.10-015 | GCATAGGTTTGGTTTCCCAG |
| 6773 | RP4-669L17.10-015 | TCTGTGTGGGTCTACCTCCT |
| 6774 | RP4-669L17.10-015 | TGAAGGTTATATGTTAGGAA |
| 6775 | RP4-669P10.16-001 | ACTCAGCCCACCCCACATCA |
| 6776 | RP4-669P10.16-001 | AAGGGAGGTGACTGTTGTGA |
| 6777 | RP4-669P10.16-001 | GGGGTGTCATATATGAAGGG |
| 6778 | RP4-669P10.16-001 | GGACTGCTGTCCCTGATGTG |
| 6779 | RP4-669P10.16-001 | GAGGGCAGGACTCAGCCTGG |
| 6780 | RP4-669P10.16-001 | GGCCTGAAATATGGGCCTCC |
| 6781 | RP4-669P10.16-001 | CTGCTGTCCCTGATGTGGGG |
| 6782 | RP4-669P10.20-001 | CCAGACAGCAGCATCCCTGA |
| 6783 | RP4-669P10.20-001 | AGGTGGCCAGTCCTGTGGTG |
| 6784 | RP4-669P10.20-001 | TCCAGACAGCAGCATCCCTG |
| 6785 | RP4-669P10.20-001 | GTTGAGAGGTGTCAACGTGC |
| 6786 | RP4-669P10.20-001 | AGTCCTGTGGTGAGGTTGAG |
| 6787 | RP4-669P10.20-001 | CAGAGAGGTGGCCAGTCCTG |
| 6788 | RP4-669P10.20-001 | ACACCTCTCAACCTCACCAC |
| 6789 | RP4-680D5.9-001 | TGAGGCCCTTGGGATACGTG |
| 6790 | RP4-680D5.9-001 | AGAGGGATAAGGATAAGGAG |
| 6791 | RP4-680D5.9-001 | AAGGATAAGGAGTGGGAATG |
| 6792 | RP4-680D5.9-001 | GAGGGATAAGGATAAGGAGT |
| 6793 | RP4-680D5.9-001 | ATACGTGGGGCTGAAGAAGA |

| | | |
|---|---|---|
| 6794 | RP4-680D5.9-001 | GGAGTGGGAATGGGGCATTG |
| 6795 | RP4-680D5.9-001 | GATACGTGGGGCTGAAGAAG |
| 6796 | RP4-717I23.3-002 | GGAAGGGCGGCAGCCACACC |
| 6797 | RP4-717I23.3-002 | CTGCTGGCAGCGCGAACGGG |
| 6798 | RP4-717I23.3-002 | GAAGCTGCTGGCAGCGCGAA |
| 6799 | RP4-717I23.3-002 | CGCCCTTCCCCCACCCACGC |
| 6800 | RP4-717I23.3-002 | CCTTCCCCCACCCACGCCGG |
| 6801 | RP4-717I23.3-002 | AAGCTGCTGGCAGCGCGAAC |
| 6802 | RP4-717I23.3-002 | CCGTAGCAGGCGGAAGCTGC |
| 6803 | RP4-717I23.3-015 | CTTGCCAGGACCGTAGCAGG |
| 6804 | RP4-717I23.3-015 | CCTTCACAGGGACCCTAGCC |
| 6805 | RP4-717I23.3-015 | GTGCTTGCCAGGACCGTAGC |
| 6806 | RP4-717I23.3-015 | TGTGAAGGAATGGGATCCCG |
| 6807 | RP4-717I23.3-015 | GGCGCGCGCTTGCCTGGCTA |
| 6808 | RP4-717I23.3-015 | TACGGTCCTGGCAAGCACCT |
| 6809 | RP4-717I23.3-015 | TAGGGTCCCTGTGAAGGAAT |
| 6810 | RP4-717I23.3-023 | TCCACCCTTACGCTGTGCTG |
| 6811 | RP4-717I23.3-023 | CCCTCCCCAGCACAGCGTAA |
| 6812 | RP4-717I23.3-023 | GTATAGGAGTTGTACTGGAA |
| 6813 | RP4-717I23.3-023 | TCCCCAGCACAGCGTAAGGG |
| 6814 | RP4-717I23.3-023 | CTTGGAAAGATGGCAAGTAT |
| 6815 | RP4-717I23.3-023 | TGCCATCTTTCCAAGCCCAA |
| 6816 | RP4-717I23.3-023 | AGGAGTTGTACTGGAACGGC |
| 6817 | RP4-758J18.13-001 | GCTCTGCACAACACTTTGGG |
| 6818 | RP4-758J18.13-001 | GGCTGGAGTGCAATGAATGG |
| 6819 | RP4-758J18.13-001 | GGCCCCAGTAATCCCAGCTG |
| 6820 | RP4-758J18.13-001 | CCATCTTAAAAAAAAAGAGG |
| 6821 | RP4-758J18.13-001 | TCACCCACGCTAGAGTGCAG |
| 6822 | RP4-758J18.13-001 | GAGAATTGCTTGAACCCGGG |
| 6823 | RP4-758J18.13-001 | GCTGGAGTGCAATGAATGGC |
| 6824 | RP4-761J14.10-001 | CATACTAAAGCCGAATGTCA |
| 6825 | RP4-761J14.10-001 | CTAAAGCCGAATGTCAGGGT |
| 6826 | RP4-761J14.10-001 | AAATTCCCAAATGTGGCAGT |
| 6827 | RP4-761J14.10-001 | CAATTCACAAGGAATATGGC |
| 6828 | RP4-761J14.10-001 | TATCTGGAATGACTGATAAG |
| 6829 | RP4-761J14.10-001 | TCATCTCCAACCCTGACATT |
| 6830 | RP4-761J14.10-001 | AATTCACAAGGAATATGGCA |
| 6831 | RP5-1057J7.7-001 | CATTATCCACAGACGTGCAG |
| 6832 | RP5-1057J7.7-001 | GTTGGTCCCTCAGATGTATG |
| 6833 | RP5-1057J7.7-001 | ACATTATCCACAGACGTGCA |
| 6834 | RP5-1057J7.7-001 | ATAAATGCCACATACATCTG |
| 6835 | RP5-1057J7.7-001 | ATTATCCACAGACGTGCAGG |
| 6836 | RP5-1057J7.7-001 | AATATAGGTGAGAATTCACT |

| | | |
|---|---|---|
| 6837 | RP5-1057J7.7-001 | GCACAGATTTCCACACCAGA |
| 6838 | RP5-1068E13.7-001 | CCTCTTAACAGGATTGACCA |
| 6839 | RP5-1068E13.7-001 | TGGATCCTGGAACTAAACTG |
| 6840 | RP5-1068E13.7-001 | AGTTGACCCTTGATGAACCA |
| 6841 | RP5-1068E13.7-001 | GATTGACCATGGTTCATCAA |
| 6842 | RP5-1068E13.7-001 | CCTGAATCTCCTGACTATGG |
| 6843 | RP5-1068E13.7-001 | CACACTCCTCCACCATAGTC |
| 6844 | RP5-1068E13.7-001 | AGATTCAGGGAAAGTTGGCA |
| 6845 | RP5-1085F17.3-001 | AGGCACTTCTTGGCCAGGAG |
| 6846 | RP5-1085F17.3-001 | TTAAAGTAGCTGCTGAACAC |
| 6847 | RP5-1085F17.3-001 | GCCGAGGCAGGTGGATCACA |
| 6848 | RP5-1085F17.3-001 | GAAACAGGATTTCACCATGT |
| 6849 | RP5-1085F17.3-001 | ACACAAAAATTAGCCAGGCA |
| 6850 | RP5-1085F17.3-001 | CAAAAATTAGCCAGGCACGG |
| 6851 | RP5-1085F17.3-001 | TAAAAACACAAAAATTAGCC |
| 6852 | RP5-115904.1-001 | GGTCGTGGGTCCCAGCCCAG |
| 6853 | RP5-115904.1-001 | TGCGCGTCCTCCAGGCGTGG |
| 6854 | RP5-115904.1-001 | GGTCCAGGTCACGGGCCGCA |
| 6855 | RP5-115904.1-001 | GGCCCGTGACCTGGACCACT |
| 6856 | RP5-115904.1-001 | GAGCCCGGACGCGGGCAAGA |
| 6857 | RP5-115904.1-001 | AGCCCAGTGGTCCAGGTCAC |
| 6858 | RP5-115904.1-001 | GTGACCTGGACCACTGGGCT |
| 6859 | RP5-1186P10.2-001 | TTAAAGAAAGGGCAGAAGTG |
| 6860 | RP5-1186P10.2-001 | CTGCCCTTTCTTTAAGACAT |
| 6861 | RP5-1186P10.2-001 | AAACGCTTTGGCTTGGAGCA |
| 6862 | RP5-1186P10.2-001 | AACGCTTTGGCTTGGAGCAG |
| 6863 | RP5-1186P10.2-001 | TTGCTTCAAACGCTTTGGCT |
| 6864 | RP5-1186P10.2-001 | GACCCTATGTCTTAAAGAAA |
| 6865 | RP5-1186P10.2-001 | TCTTAAAGAAAGGGCAGAAG |
| 6866 | RP5-1198O20.4-013 | CACTGCAGATCTCGAGCAGG |
| 6867 | RP5-1198O20.4-013 | GGCCGGGGGTGGGCTCTCCA |
| 6868 | RP5-1198O20.4-013 | AGCCCTGGAGAGCCCACCCC |
| 6869 | RP5-1198O20.4-013 | GGGCAGGGGCCTAGCAGCCC |
| 6870 | RP5-1198O20.4-013 | CTGGAGAGCCCACCCCCGGC |
| 6871 | RP5-1198O20.4-013 | TCTGCAGTGGCTGCCGGGTG |
| 6872 | RP5-1198O20.4-013 | GTGGGCTCTCCAGGGCTGCT |
| 6873 | RP5-832C2.5-001 | TGAGGACTATGGGCCCCACG |
| 6874 | RP5-832C2.5-001 | GCTTGCCGGAGATAGCATTG |
| 6875 | RP5-832C2.5-001 | AATGCTATCTCCGGCAAGCG |
| 6876 | RP5-832C2.5-001 | TAGCATTGAGGGCCCTTCCG |
| 6877 | RP5-832C2.5-001 | AGGGCCCTCAATGCTATCTC |
| 6878 | RP5-832C2.5-001 | CTTGCCGGAGATAGCATTGA |
| 6879 | RP5-832C2.5-001 | GCGTTCCCTCGGTCCCCGTG |

| | | |
|---|---|---|
| 6880 | RP5-843L14.1-002 | ACCGTTGTGTCTAAACCATG |
| 6881 | RP5-843L14.1-002 | CCACCGTTGTGTCTAAACCA |
| 6882 | RP5-843L14.1-002 | CGAGATTGGCCAGAATCTGG |
| 6883 | RP5-843L14.1-002 | TCAGGTCCAGGTGAGAATTG |
| 6884 | RP5-843L14.1-002 | ACTGAGAGAATTTAGCTAGG |
| 6885 | RP5-843L14.1-002 | CACCGTTGTGTCTAAACCAT |
| 6886 | RP5-843L14.1-002 | AGAGGGAAAAAAGCCGAGAT |
| 6887 | RP5-887A10.1-001 | TATGTTATTAATGCCCAGCA |
| 6888 | RP5-887A10.1-001 | AAAATGAAGCATACAGTTGT |
| 6889 | RP5-887A10.1-001 | CCATATGCTTATCAATTATG |
| 6890 | RP5-887A10.1-001 | TAAAATGAAGCATACAGTTG |
| 6891 | RP5-887A10.1-001 | TGCTGGGCATTAATAACATA |
| 6892 | RP5-887A10.1-001 | AGCATATTGCAGAAACTCTC |
| 6893 | RP5-887A10.1-001 | TATTAATGCCCAGCAAGGCA |
| 6894 | RP5-902P8.10-001 | CCTTCTGGTAAGCAGGACAC |
| 6895 | RP5-902P8.10-001 | AGAAGGCCACAGATGCGGGT |
| 6896 | RP5-902P8.10-001 | GTGCTGGCCACACCTGTCCG |
| 6897 | RP5-902P8.10-001 | TACCAGAAGGCCACAGATGC |
| 6898 | RP5-902P8.10-001 | TTACCAGAAGGCCACAGATG |
| 6899 | RP5-902P8.10-001 | GGGGTCTTGGCAAGACGTGC |
| 6900 | RP5-902P8.10-001 | CAGAAGGCCACAGATGCGGG |
| 6901 | RP5-902P8.10-002 | GCCATCGGCTCTATGGCTAG |
| 6902 | RP5-902P8.10-002 | CCTATTCCTGAAGCTGCCAG |
| 6903 | RP5-902P8.10-002 | CCTGTGTCCTGCTTACCAGA |
| 6904 | RP5-902P8.10-002 | GCCCTCAGAGGGCAATCTCA |
| 6905 | RP5-902P8.10-002 | CCAGAGGAAGTCCCTCCGGA |
| 6906 | RP5-902P8.10-002 | TCTGTGGCCTTCTGGTAAGC |
| 6907 | RP5-902P8.10-002 | GCCAGAGGAAGTCCCTCCGG |
| 6908 | RP5-902P8.12-001 | GGGACACACCGTGCTCCCAG |
| 6909 | RP5-902P8.12-001 | GTGACAACACAAGAACAGAG |
| 6910 | RP5-902P8.12-001 | GGGGACACACCGTGCTCCCA |
| 6911 | RP5-902P8.12-001 | GGTGACAACACAAGAACAGA |
| 6912 | RP5-902P8.12-001 | GGGGCCCACGATTCCTCTAA |
| 6913 | RP5-902P8.12-001 | CGGTGACAACACAAGAACAG |
| 6914 | RP5-902P8.12-001 | AGGGGCCCACGATTCCTCTA |
| 6915 | RP5-903G2.2-001 | ACGTCACTCCCGAAGAGAAG |
| 6916 | RP5-903G2.2-001 | TGGCCGGCCCAGGAGAGAGG |
| 6917 | RP5-903G2.2-001 | ATGCCTAGCTCCGCGCCGCG |
| 6918 | RP5-903G2.2-001 | CTGCGGCTGTTGCTGGAGGA |
| 6919 | RP5-903G2.2-001 | ACGTCGTCCTCCTCTCTCCT |
| 6920 | RP5-903G2.2-001 | GACGCCCCAGCTCCGCCGAC |
| 6921 | RP5-903G2.2-001 | GCGGCCTGTCGGCGGAGCTG |
| 6922 | RP5-921G16.2-001 | GGAAAGCTGAGATAAACCAC |

| | | |
|---|---|---|
| 6923 | RP5-921G16.2-001 | TAGAGTCAGTGCTCTTTGTG |
| 6924 | RP5-921G16.2-001 | AGTGGGATGAGGCCTGTGAC |
| 6925 | RP5-921G16.2-001 | GCAGCTTTCCCGCAATTCCC |
| 6926 | RP5-921G16.2-001 | ACAGGTCACCAGGGAATTGC |
| 6927 | RP5-921G16.2-001 | GCGGGAAAGCTGCCAGTCAC |
| 6928 | RP5-921G16.2-001 | TACCAGGCAGCCAGTGATCA |
| 6929 | RP5-943J3.2-001 | CGGCGAGCGACTGCTAGATG |
| 6930 | RP5-943J3.2-001 | CTCGCCACACAGACCCCGTC |
| 6931 | RP5-943J3.2-001 | CGCCGCCAGTCTCCAACCCC |
| 6932 | RP5-943J3.2-001 | CTGCAGCGCCTGACCGGACG |
| 6933 | RP5-943J3.2-001 | GGACGGGGTCTGTGTGGCGA |
| 6934 | RP5-943J3.2-001 | ATGCGGAGCAGGTGCTTGGG |
| 6935 | RP5-943J3.2-001 | CCTGCAGCGCCTGACCGGAC |
| 6936 | RP5-967N21.11-001 | AAATACCCAAAATATAAACA |
| 6937 | RP5-967N21.11-001 | GACTCGCTCTGTACCTAGTC |
| 6938 | RP5-967N21.11-001 | TATATATTCCCTGGGCTGGT |
| 6939 | RP5-967N21.11-001 | AGGTACAGAGCGAGTCTATG |
| 6940 | RP5-967N21.11-001 | CAAGGCAAGAGAACTGTTGA |
| 6941 | RP5-967N21.11-001 | CAAGAGAACTGTTGAAGGTC |
| 6942 | RP5-967N21.11-001 | ACCAGCCCAGGGAATATATA |
| 6943 | RP5-968D22.3-001 | AAACCAGGAGAAGATCCACC |
| 6944 | RP5-968D22.3-001 | ACCAAAAATTAGCCAGGCGT |
| 6945 | RP5-968D22.3-001 | GAGAATGGCGTGAACCCAGG |
| 6946 | RP5-968D22.3-001 | AATATGTTGTTGGCCAGGCG |
| 6947 | RP5-968D22.3-001 | CTAAATCACTTAAGTTTCTG |
| 6948 | RP5-968D22.3-001 | TGAATGAAAAAATATGTTGT |
| 6949 | RP5-968D22.3-001 | AAAATAACCAAAAATTAGCC |
| 6950 | RP5-973N23.4-001 | TACTCCAGAGCAATTCAAGA |
| 6951 | RP5-973N23.4-001 | CTACTCCAGAGCAATTCAAG |
| 6952 | RP5-973N23.4-001 | CAAAAATTAGCCGGACATGG |
| 6953 | RP5-973N23.4-001 | TGCTGTGACCATAGATAGTG |
| 6954 | RP5-973N23.4-001 | GAGAATTGCTTGAAACCGGG |
| 6955 | RP5-973N23.4-001 | AAAATATAAAAAACAGAAGT |
| 6956 | RP5-973N23.4-001 | CAGGAGAATTGCTTGAAACC |
| 6957 | RP5-999L4.2-001 | GAAAAGCCCATGAACAGGGT |
| 6958 | RP5-999L4.2-001 | ATGGAGTTAGATCATTATGG |
| 6959 | RP5-999L4.2-001 | GGAAAAGCCCATGAACAGGG |
| 6960 | RP5-999L4.2-001 | AATGGAGTTAGATCATTATG |
| 6961 | RP5-999L4.2-001 | CAGGGAAAAGCCCATGAACA |
| 6962 | RP5-999L4.2-001 | CATTATGGGGGCAGTGCTCA |
| 6963 | RP5-999L4.2-001 | ATAATGGAGTTAGATCATTA |
| 6964 | RPARP-AS1-001 | GTGGGCGACGACTCTGATAG |
| 6965 | RPARP-AS1-001 | GACTCTGATAGAGGGAAAAG |

| | | |
|---|---|---|
| 6966 | RPARP-AS1-001 | TGGGCGACGACTCTGATAGA |
| 6967 | RPARP-AS1-001 | TGATAGAGGGAAAAGAGGAA |
| 6968 | RPARP-AS1-001 | GAGCGGTGGGAAAGGGGGGT |
| 6969 | RPARP-AS1-001 | TCGGCTTGGCAGGAAGGCGG |
| 6970 | RPARP-AS1-001 | CCCTCGGCTTGGCAGGAAGG |
| 6971 | RPP30 | GAATCAGACCAACACCTCGG |
| 6972 | RPP30 | TAGGAATCAGACCAACACCT |
| 6973 | RPP30 | ACGGTCATGGGACTTCAGCA |
| 6974 | RPP30 | GCGAGCGGGTTCTGACCTGA |
| 6975 | RPP30 | TGCAGATTTGGACCTGCGAG |
| 6976 | RPP30 | TCAGGTCAGAACCCGCTCGC |
| 6977 | RPP30 | TCTGACCTGAAGGCTCTGCG |
| 6978 | SCAMP1-AS1-001 | ACTCGGTCGCAGCTAAAGCC |
| 6979 | SCAMP1-AS1-001 | GCAGCTAAAGCCCGGAGCGT |
| 6980 | SCAMP1-AS1-001 | CGCAGCTAAAGCCCGGAGCG |
| 6981 | SCAMP1-AS1-001 | AGCGTGGGATCCGGGCCCGG |
| 6982 | SCAMP1-AS1-001 | GGGCCCGGATCCCACGCTCC |
| 6983 | SCAMP1-AS1-001 | GTTCGCCATCCCGTTCCTCC |
| 6984 | SCAMP1-AS1-001 | AAGCCCGGAGCGTGGGATCC |
| 6985 | SERPINB9P1-001 | TTGAGGAGGAGTACTAAGAG |
| 6986 | SERPINB9P1-001 | TGAGGAGGAGTACTAAGAGT |
| 6987 | SERPINB9P1-001 | GGACTGCCAGTACTATGTTG |
| 6988 | SERPINB9P1-001 | CTGCCAGTACTATGTTGAGG |
| 6989 | SERPINB9P1-001 | TGTCTTGTTCCCATTCTCAG |
| 6990 | SERPINB9P1-001 | ACTCCTCCTCAACATAGTAC |
| 6991 | SERPINB9P1-001 | GGTTTCATATCAGGGATGCA |
| 6992 | SETDB1 | GACCCCAGACTCACAACTCA |
| 6993 | SETDB1 | CGAAAGCGAAGGGATAAGGG |
| 6994 | SETDB1 | ACCCCAGACTCACAACTCAG |
| 6995 | SETDB1 | ACCCCTGAGTTGTGAGTCTG |
| 6996 | SETDB1 | ACGGAAAAGAGCGAAAGCGA |
| 6997 | SETDB1 | CGGAAAAGAGCGAAAGCGAA |
| 6998 | SETDB1 | GAGCGAAAGCGAAGGGATAA |
| 6999 | SNAPIN | GAAGCCCGAAACCTGCCGCG |
| 7000 | SNAPIN | CCCTGCCCCCGATACAGCGG |
| 7001 | SNAPIN | TGGTTCCGCCGCTGTATCGG |
| 7002 | SNAPIN | GGCTTCAGGACAATTCGTGA |
| 7003 | SNAPIN | AGGACAATTCGTGATGGCGG |
| 7004 | SNAPIN | CAGGACAATTCGTGATGGCG |
| 7005 | SNAPIN | CTGGTTCCGCCGCTGTATCG |
| 7006 | SNHG1-001 | GAGGCTTAAAAGTAGCAGTG |
| 7007 | SNHG1-001 | TGCCTGGCCAGGGCGACTGG |
| 7008 | SNHG1-001 | GGCGGATAAGGTCTTGTGCG |

| | | |
|---|---|---|
| 7009 | SNHG1-001 | AGGTCTTGTGCGTGGCCTCG |
| 7010 | SNHG1-001 | ACCTTATCCGCCAGTCGCCC |
| 7011 | SNHG1-001 | GAAGGACAAAATGGCGATGG |
| 7012 | SNHG1-001 | GCCAGGGCGACTGGCGGATA |
| 7013 | SNHG1-004 | TGCAGGATTTACGCGCACGT |
| 7014 | SNHG1-004 | TGCGCGTAAATCCTGCAGAC |
| 7015 | SNHG1-004 | ACTGTACGCAACAAATGTCA |
| 7016 | SNHG1-004 | ACACCTCAGACAAATCAATA |
| 7017 | SNHG1-004 | GGGCCCTATTGATTTGTCTG |
| 7018 | SNHG1-004 | AGTGTGTGGAAGTTGCTACG |
| 7019 | SNHG1-004 | TACTGTACGCAACAAATGTC |
| 7020 | SNHG1-006 | CTGGAACATTACTGTAACGC |
| 7021 | SNHG1-006 | ACTACTGAGAAGTGAGAGAG |
| 7022 | SNHG1-006 | CCACAAGAGCTTACTGGTGA |
| 7023 | SNHG1-006 | CTTCACCAGTAAGCTCTTGT |
| 7024 | SNHG1-006 | CGCTGGCTTTGCATAAAGAT |
| 7025 | SNHG1-006 | AGAGCTTACTGGTGAAGGAA |
| 7026 | SNHG1-006 | TTCAGCCCACAAGAGCTTAC |
| 7027 | SNHG1-021 | TAAGACATTAGGTCAACTCA |
| 7028 | SNHG1-021 | ATAAGACATTAGGTCAACTC |
| 7029 | SNHG1-021 | CAGGAGCCCAATAAGACATT |
| 7030 | SNHG1-021 | CTTATTGGGCTCCTGTCTGC |
| 7031 | SNHG1-021 | AGTTGACCTAATGTCTTATT |
| 7032 | SNHG1-021 | GAGTTGACCTAATGTCTTAT |
| 7033 | SNHG1-021 | ACTGAACTCTCTCTTTCTGA |
| 7034 | SNHG1-025 | GATTAAATACCATGCCGCAG |
| 7035 | SNHG1-025 | AATGAGAACGAATCTCCCCG |
| 7036 | SNHG1-025 | AAATGAGAACGAATCTCCCC |
| 7037 | SNHG1-025 | AAAATGAGAACGAATCTCCC |
| 7038 | SNHG1-025 | GCTCGTGGTAAGTGGCTTCG |
| 7039 | SNHG1-025 | CCAGGCAGCTACAACACAGA |
| 7040 | SNHG1-025 | CCAAATCCCGTGGGCCTCTG |
| 7041 | SNHG11-006 | CGGGTACAGGGGTAACCGCG |
| 7042 | SNHG11-006 | GTACAGGGGTAACCGCGCGG |
| 7043 | SNHG11-006 | ACTCATCCGGAGCGCCAGGA |
| 7044 | SNHG11-006 | CGGAGCGCCAGGACGGATGG |
| 7045 | SNHG11-006 | ATCCGGAGCGCCAGGACGGA |
| 7046 | SNHG11-006 | GTACCCGCCGCCATCCGTCC |
| 7047 | SNHG11-006 | GACGGATGGCGGCGGGTACA |
| 7048 | SNHG15-001 | TTGGGGTAGATGACGTCACG |
| 7049 | SNHG15-001 | GCGTCTGCGTGTCAACTCAG |
| 7050 | SNHG15-001 | TCGCTTGGTAGGGTGTGCTG |
| 7051 | SNHG15-001 | GCGTAGAGTTTAGGGTCCTG |

| | | |
|---|---|---|
| 7052 | SNHG15-001 | CCCTACCAAGCGAGCGCCCG |
| 7053 | SNHG15-001 | TAGATGACGTCACGCGGCCG |
| 7054 | SNHG15-001 | GCCGCGGGCGCTCGCTTGGT |
| 7055 | SNHG15-002 | ACTTCGTAGTCGCAGCGGCG |
| 7056 | SNHG15-002 | TCCGTACTTCGTAGTCGCAG |
| 7057 | SNHG15-002 | GAATGGTCAGGCAACACGTG |
| 7058 | SNHG15-002 | AAGTACGGAGTACGGAGTAC |
| 7059 | SNHG15-002 | AGGAATGGTCAGGCAACACG |
| 7060 | SNHG15-002 | CGACTACGAAGTACGGAGTA |
| 7061 | SNHG15-002 | GGAATGGTCAGGCAACACGT |
| 7062 | SNHG15-004 | TGCTACTTCCTCTCAAGAGG |
| 7063 | SNHG15-004 | CGCGGTGACGTCGAGGTGCG |
| 7064 | SNHG15-004 | GGGACTACGCGGTGACGTCG |
| 7065 | SNHG15-004 | GACCATTCCTGAGCCCAGGT |
| 7066 | SNHG15-004 | AAGAGGCGGTTGAGCGCGTT |
| 7067 | SNHG15-004 | GCCACTGAACCCAATCCCCC |
| 7068 | SNHG15-004 | GCTGTGGCCCCGGCCACGCA |
| 7069 | SNHG15-006 | TACCCCAAACGCTGTGGCCC |
| 7070 | SNHG15-006 | AGCCCAGGTAGGGGTCCTGG |
| 7071 | SNHG15-006 | TCCCGCCCCGAAGCCGTGCG |
| 7072 | SNHG15-006 | CCCCTACCTGGGCTCAGGAA |
| 7073 | SNHG15-006 | TCCTGGGGGATTGGGTTCAG |
| 7074 | SNHG15-006 | CCCCGAAGCCGTGCGTGGCC |
| 7075 | SNHG15-006 | GCCTGACCATTCCTGAGCCC |
| 7076 | SNHG15-007 | GCGCTCAACCGCCTCTTGAG |
| 7077 | SNHG15-007 | TAGCACGTGTAGTGAAGGCT |
| 7078 | SNHG15-007 | ACTGGTGCCCCCACTCACTG |
| 7079 | SNHG15-007 | ACGTGCTACTTCCTCTCAAG |
| 7080 | SNHG15-007 | GGAAGTAGCACGTGTAGTGA |
| 7081 | SNHG15-007 | GTGGGGGCACCAGTGATTCT |
| 7082 | SNHG15-007 | GAGGCTGGGCCGCAGTGAGT |
| 7083 | SNHG16-002 | AGTATGTTAGTGTCGTGTCA |
| 7084 | SNHG16-002 | AAGCCGGCCCTATGACGCCG |
| 7085 | SNHG16-002 | TGATGGTAAGGCTGCCTGGG |
| 7086 | SNHG16-002 | CCGAGAAGACTTGCGCGCAG |
| 7087 | SNHG16-002 | GATGGTAAGGCTGCCTGGGT |
| 7088 | SNHG16-002 | GAGAAAATGTTACTATGTGA |
| 7089 | SNHG16-002 | TGACACGACACTAACATACT |
| 7090 | SNHG16-003 | TCATAGGGCCGGCTTCTGCG |
| 7091 | SNHG16-003 | CATAGGGCCGGCTTCTGCGG |
| 7092 | SNHG16-003 | CGTCATAGGGCCGGCTTCTG |
| 7093 | SNHG16-003 | GCGCTCTCCCCCGCAGAAGC |
| 7094 | SNHG16-003 | TGCCGAGAAGACTTGCGCGC |

| | | |
|---|---|---|
| 7095 | SNHG16-003 | AGCCCGTCCTCGGCGTCATA |
| 7096 | SNHG16-003 | GTCATAGGGCCGGCTTCTGC |
| 7097 | SNHG16-007 | GTGCTCAACACCGACTGGAG |
| 7098 | SNHG16-007 | GAAGACATGGCCACTCCAGT |
| 7099 | SNHG16-007 | CGAAGTGGTGACGATAATCT |
| 7100 | SNHG16-007 | GGCCATGTCTTCCAGCCACG |
| 7101 | SNHG16-007 | TGATGATGGTAAGGCTGCCT |
| 7102 | SNHG16-007 | CCCTCCTAACCACCCTATGA |
| 7103 | SNHG16-007 | TCGGTTTCTCCCTCATAGGG |
| 7104 | SNHG17-001 | TCTGCAGATTTCGAGAGATG |
| 7105 | SNHG17-001 | GCGATGGCGAAGGACGGCGA |
| 7106 | SNHG17-001 | CGAGGGACGGCGAAGGACTG |
| 7107 | SNHG17-001 | GCGGTGGCGTGCGATGGCGA |
| 7108 | SNHG17-001 | TGGCGTGCGATGGCGAAGGA |
| 7109 | SNHG17-001 | TGTCTGCAGATTTCGAGAGA |
| 7110 | SNHG17-001 | GTCTGCAGATTTCGAGAGAT |
| 7111 | SNHG17-003 | CAGGATCCATGCTCTCCCAG |
| 7112 | SNHG17-003 | GCAGAGAACTTGAGCTCTAG |
| 7113 | SNHG17-003 | TGAGCTCTAGAGGACTGAGC |
| 7114 | SNHG17-003 | AGCATGGATCCTGAGGTCCC |
| 7115 | SNHG17-003 | GCAGGTTGTGAGGCCAGAGG |
| 7116 | SNHG17-003 | TCTGCAGGTTGTGAGGCCAG |
| 7117 | SNHG17-003 | CCTGAGGTCCCAGGTAAGTG |
| 7118 | SNHG17-008 | AGCGTAGCTTCCTTGTCGTG |
| 7119 | SNHG17-008 | GGCTCCACTCTCACCATCGG |
| 7120 | SNHG17-008 | AGGACTGAGGCCACACGACA |
| 7121 | SNHG17-008 | GAGACACTGCACCAAGGACA |
| 7122 | SNHG17-008 | CGGCTCCACTCTCACCATCG |
| 7123 | SNHG17-008 | TCCACAGCAGGGCGCATGGT |
| 7124 | SNHG17-008 | TGAGTCCACAGCAGGGCGCA |
| 7125 | SNHG18-002 | CAGCCTCCCCAGTGCGAGTG |
| 7126 | SNHG18-002 | GCTCGGCCGCACTCGCACTG |
| 7127 | SNHG18-002 | CGGCCGCACTCGCACTGGGG |
| 7128 | SNHG18-002 | GGACCCCCACCTTCGCTCCT |
| 7129 | SNHG18-002 | GGAACTCCTGGGGTGCAGCG |
| 7130 | SNHG18-002 | AGGCTGGAATGGAACTCCTG |
| 7131 | SNHG18-002 | ATGGAACTCCTGGGGTGCAG |
| 7132 | SNHG19-001 | GACGAACTGAGCCACGAGCG |
| 7133 | SNHG19-001 | TTAACAAACCTCTTCCCCCT |
| 7134 | SNHG19-001 | CGTCCCAAGATCGTAGCACA |
| 7135 | SNHG19-001 | GACTCGACCCCGAAAAGACA |
| 7136 | SNHG19-001 | GGCCCTCAAAGCCACGCTCG |
| 7137 | SNHG19-001 | GGACTCGACCCCGAAAAGAC |

| | | |
|---|---|---|
| 7138 | SNHG19-001 | TCGCCATGTGCTACGATCTT |
| 7139 | SNHG20-002 | GAAGCCGTCCCCGAAATCCC |
| 7140 | SNHG20-002 | CCACGGCGCCCGGGATTTCG |
| 7141 | SNHG20-002 | AGAAGCCGTCCCCGAAATCC |
| 7142 | SNHG20-002 | CCCCGAAATCCCGGGCGCCG |
| 7143 | SNHG20-002 | CGAAATCCCGGGCGCCGTGG |
| 7144 | SNHG20-002 | AATCCCGGGCGCCGTGGAGG |
| 7145 | SNHG20-002 | AAATCCCGGGCGCCGTGGAG |
| 7146 | SNHG23-005 | TGATCATCCTGTAGCGACGG |
| 7147 | SNHG23-005 | AATGTAATTCAGTACAACTG |
| 7148 | SNHG23-005 | GATTGATCATCCTGTAGCGA |
| 7149 | SNHG23-005 | GTCAATACCACCGTCGCTAC |
| 7150 | SNHG23-005 | TTCGCAGGGTTGATGTTCAT |
| 7151 | SNHG23-005 | ATACCTAAAGAAAATCAAAG |
| 7152 | SNHG23-005 | TGGTATTGACTGTTCCCCAG |
| 7153 | SNHG25-002 | AGGAATACAACCTGACCTCC |
| 7154 | SNHG25-002 | GGTCAGGTTGTATTCCTGGC |
| 7155 | SNHG25-002 | AGGTCAGGTTGTATTCCTGG |
| 7156 | SNHG25-002 | ACCCAGCCAGACGACCCGCC |
| 7157 | SNHG25-002 | CTCCCGAGCCACGACCTCCG |
| 7158 | SNHG25-002 | TGTATTCCTGGCGGGTCGTC |
| 7159 | SNHG25-002 | CCTGCAGGTTCCGGGAGGTC |
| 7160 | SNHG5-001 | GGCCTCGTTCTGACTCCGGG |
| 7161 | SNHG5-001 | GAGAGTTGAAAGCTTGCACG |
| 7162 | SNHG5-001 | CTTCACTAAGATGTCTTCAC |
| 7163 | SNHG5-001 | GCCGCTCAGGCACATGCCGG |
| 7164 | SNHG5-001 | GAGGCTATATAAGGAGCTAC |
| 7165 | SNHG5-001 | AGCCTCCCGGAGTCAGAACG |
| 7166 | SNHG5-001 | GGACGGCCTCGTTCTGACTC |
| 7167 | SNHG5-003 | AAATGGCGAAGGCCTTCCGT |
| 7168 | SNHG5-003 | AAAAACCACCTTAATTGGGG |
| 7169 | SNHG5-003 | ACCCTCCGCCCCAATTAAGG |
| 7170 | SNHG5-003 | TTTAAAACAACGCCGACGGA |
| 7171 | SNHG5-003 | CGGAAAAACCACCTTAATTG |
| 7172 | SNHG5-003 | GGCCTTTACTAAAATGGCGA |
| 7173 | SNHG5-003 | ACCACCTTAATTGGGGCGGA |
| 7174 | SNHG5-004 | AACAGCTCAGTTTCAATACA |
| 7175 | SNHG5-004 | TTCGGATTAATGAAGGGGAA |
| 7176 | SNHG5-004 | GAACAGCTCAGTTTCAATAC |
| 7177 | SNHG5-004 | GTGCAATTCGGATTAATGAA |
| 7178 | SNHG5-004 | AGGCGGCGTTGTGTGCAATT |
| 7179 | SNHG5-011 | GGCCTTCGCGAGCGTCTGGG |
| 7180 | SNHG5-011 | GCTAGTATCCGAGCAAAAGA |

| | | |
|---|---|---|
| 7181 | SNHG5-011 | GTCGGCCTTCGCGAGCGTCT |
| 7182 | SNHG5-011 | ACCCGCCCAGACGCTCGCGA |
| 7183 | SNHG5-011 | CAGGATTAAAGCTCTTAACT |
| 7184 | SNHG5-011 | AGTATCCGAGCAAAAGATGG |
| 7185 | SNHG5-011 | TGGTAGGTGAGTGGGTATTG |
| 7186 | SNHG5-012 | CGACTCAACCTCAGTACATG |
| 7187 | SNHG5-012 | AGGAACCGCTGATCGGCCAC |
| 7188 | SNHG5-012 | GCTTGCACGTGGGAAACGTG |
| 7189 | SNHG5-012 | TGGCGTGAGGAACCGCTGAT |
| 7190 | SNHG5-012 | AGAGTTGAAAGCTTGCACGT |
| 7191 | SNHG5-012 | GAGCTACTGGCTGCGCACTT |
| 7192 | SNHG5-012 | GAGGAACCGCTGATCGGCCA |
| 7193 | SNHG5-013 | ATTGAAACTGAGCTGTTCGA |
| 7194 | SNHG5-013 | TCAGTTTCAATACAGGGTCG |
| 7195 | SNHG5-013 | TGCAATTCGGATTAATGAAG |
| 7196 | SNHG5-013 | CTTAAATACCGATTTCCCGC |
| 7197 | SNHG5-013 | GAAACTGAGCTGTTCGAAGG |
| 7198 | SNHG5-013 | ATTCGGATTAATGAAGGGGA |
| 7199 | SNHG5-013 | AGACAGCGCCATTGTTCCTG |
| 7200 | SNHG6-003 | CGGGAAAGGAAGAGCGACAT |
| 7201 | SNHG6-003 | ACAGCTCTTAGGCACTCCGG |
| 7202 | SNHG6-003 | CGGAGGAACCCGGCACGTAG |
| 7203 | SNHG6-003 | AACCCGGCACGTAGAGGAGT |
| 7204 | SNHG6-003 | AAGACAGCTCTTAGGCACTC |
| 7205 | SNHG6-003 | GACATCGGAAGACAGCTCTT |
| 7206 | SNHG6-003 | GCCGCGTACCGGAAGAGGCG |
| 7207 | SNHG6-005 | AGCCGTTAGTCATGCCGGTG |
| 7208 | SNHG6-005 | CGAAGAGCCGTTAGTCATGC |
| 7209 | SNHG6-005 | CGTTAGTCATGCCGGTGTGG |
| 7210 | SNHG6-005 | CACCACACCGGCATGACTAA |
| 7211 | SNHG6-005 | GCCCGTAGCTGGGCTCTGCG |
| 7212 | SNHG6-005 | CTCCGCCGCCGCCACCACAC |
| 7213 | SNHG6-005 | TAGTCATGCCGGTGTGGTGG |
| 7214 | SNHG8-001 | AACTGATGACGCTATTCCGG |
| 7215 | SNHG8-001 | TCAGGATATTGGTAGGTCGG |
| 7216 | SNHG8-001 | CTTTCACATTCGGGAAGCGT |
| 7217 | SNHG8-001 | CGACGCTTCCCGAATGTGAA |
| 7218 | SNHG8-001 | TATAAGAGAGCGTGTGCCGA |
| 7219 | SNHG8-001 | GAACTGATGACGCTATTCCG |
| 7220 | SNHG8-001 | TTCCCGAATGTGAAAGGCCG |
| 7221 | SNHG8-003 | TGAGTGCTCATTAGGTCGCC |
| 7222 | SNHG8-003 | CCCTGAACCCCGGCGTGCGT |
| 7223 | SNHG8-003 | CCCACGCACGCCGGGGTTCA |

| | | |
|---|---|---|
| 7224 | SNHG8-003 | CCCACGGCCCACGCACGCCG |
| 7225 | SNHG8-003 | CAGCCCTGGACCCTGAACCC |
| 7226 | SNHG8-003 | TCAGGGTCCAGGGCTGCACG |
| 7227 | SNHG8-003 | CGCGGACTTGAGTGCTCATT |
| 7228 | SNHG8-004 | GGAAAGGTTCCGTGCACTCG |
| 7229 | SNHG8-004 | TGCACGGAACCTTTCCCCGA |
| 7230 | SNHG8-004 | AGCGCAGTCAGGATATTGGT |
| 7231 | SNHG8-004 | GTGCACGGAACCTTTCCCCG |
| 7232 | SNHG8-004 | GCGTGGGCCGTGGGCCCTCG |
| 7233 | SNHG8-004 | ACGCCGGGGTTCAGGGTCCA |
| 7234 | SNHG8-004 | GGCCCACGGCCCACGCACGC |
| 7235 | SNHG8-005 | GCTCAAACTGACGGTTCTCG |
| 7236 | SNHG8-005 | CACCATCTGGCTCAAACTGA |
| 7237 | SNHG8-005 | TGACGGTTCTCGGGGTGCTA |
| 7238 | SNHG8-005 | GCGACAGAGTAGACTCAAGA |
| 7239 | SNHG8-005 | GTTCAGCCGTCTGTGTCATG |
| 7240 | SNHG8-005 | AGATGGCCACATGACACAGA |
| 7241 | SNHG8-005 | GACAGCTTTGCCAATAAACA |
| 7242 | SNHG9-001 | CGAAAAGCGACTATAAACGC |
| 7243 | SNHG9-001 | AGAGCACTCAGAGCCCGGTG |
| 7244 | SNHG9-001 | TTCGGGTGACGTAGATTCCC |
| 7245 | SNHG9-001 | GAGCACTCAGAGCCCGGTGA |
| 7246 | SNHG9-001 | TGACGTAGATTCCCGGGCCG |
| 7247 | SNHG9-001 | TTTCGGGTGACGTAGATTCC |
| 7248 | SNHG9-001 | TGCCCGTCCGGCCCCAGCCG |
| 7249 | SNHG9-002 | TCGGGTGGATTCCCGAGCCG |
| 7250 | SNHG9-002 | TCGGGAATCCACCCGAAGAG |
| 7251 | SNHG9-002 | GTTTATAGCCACTCTTCGGG |
| 7252 | SNHG9-002 | TGAAGAGGAGAGCGCAATGG |
| 7253 | SNHG9-002 | AGGAGAGCGCAATGGAGGCG |
| 7254 | SNHG9-002 | AAGTGAAGAGGAGAGCGCAA |
| 7255 | SNHG9-002 | TTGCGCTCTCCTCTTCACTT |
| 7256 | SP1 | GCTGAGGGACAAGCTCAAGG |
| 7257 | SP1 | AGCTGAGGGACAAGCTCAAG |
| 7258 | SP1 | GCTGACGAGGCAAGCGAACC |
| 7259 | SP1 | GCAGCTGAGGGACAAGCTCA |
| 7260 | SP1 | AAGCGAACCCGGACCGGACA |
| 7261 | SP1 | GAAAAACGCGGACGCTGACG |
| 7262 | SP1 | CCGGCCCCCCCCAACCCCCC |
| 7263 | SPACA6P-011 | CCTCCAGGGCTCGATCAGCT |
| 7264 | SPACA6P-011 | TCGATCAGCTCGGCATCCCG |
| 7265 | SPACA6P-011 | GCAAGCAGGAACAGATTGCT |
| 7266 | SPACA6P-011 | CCGAGCTGATCGAGCCCTGG |

| | | |
|---|---|---|
| 7267 | SPACA6P-011 | ATGCCGAGCTGATCGAGCCC |
| 7268 | SPACA6P-011 | GTTCCGGGAAGTGCTGCGCT |
| 7269 | SPACA6P-011 | GCCAGCAGCTCGCCCAGGCT |
| 7270 | SRP14-AS1-002 | ATAGTATCTACCGCGCAGTG |
| 7271 | SRP14-AS1-002 | CGCTTCACGACCACACTGCG |
| 7272 | SRP14-AS1-002 | GCACATAGAGCGTGTTCAGT |
| 7273 | SRP14-AS1-002 | TTTATATTCAGCTGTATGCT |
| 7274 | SRP14-AS1-002 | TTTCTCGCATGTGACGACCC |
| 7275 | SRP14-AS1-002 | AGAGTTTAAAGGATCTCTAC |
| 7276 | SRP14-AS1-002 | TCACTCCGTCACACTGGCTG |
| 7277 | TCL6-002 | AGCTGTACCACTCACCGCGA |
| 7278 | TCL6-002 | CTGCAGACCTTCGCGGTGAG |
| 7279 | TCL6-002 | TACAGCTCATAAAAGCAGCG |
| 7280 | TCL6-002 | TCGGGTCCCCTTCCACACTG |
| 7281 | TCL6-002 | AGCAAGACCACGAGCCCACC |
| 7282 | TCL6-002 | AGTGTGGAAGGGGACCCGAG |
| 7283 | TCL6-002 | AACACATCTGAACATCAGAA |
| 7284 | TK1 | ACGAACCCGAGTACTCTCCA |
| 7285 | TK1 | GGGAACCAGGGGCTTACTGC |
| 7286 | TK1 | GGGGAGCCAGGCAGCACAGT |
| 7287 | TK1 | CGGGTTCGTGAACTTCCCGG |
| 7288 | TK1 | CGGGAACCAGGGGCTTACTG |
| 7289 | TK1 | ACCAGGGGCTTACTGCGGGA |
| 7290 | TK1 | ACTCGGGTTCGTGAACTTCC |
| 7291 | TOP2A | CCCCTTCACGACCGTCACCA |
| 7292 | TOP2A | TCCATGGTGACGGTCGTGAA |
| 7293 | TOP2A | CTGAAAGCGACTAAACAGGC |
| 7294 | TOP2A | TGGTGACACTTCCATGGTGA |
| 7295 | TOP2A | CTGCAATGGTGACACTTCCA |
| 7296 | TOP2A | TTCCATGGTGACGGTCGTGA |
| 7297 | TOP2A | TGGAGAAGCGGCTTGGTCGG |
| 7298 | TOPBP1 | GTTCTCCAGCGTTTGCTTTG |
| 7299 | TOPBP1 | TCCAGCGTTTGCTTTGGGGT |
| 7300 | TOPBP1 | GGGTAGGGGCGGCGCCGAGT |
| 7301 | TOPBP1 | CCAGCGTTTGCTTTGGGGTA |
| 7302 | TOPBP1 | CAGCGTTTGCTTTGGGGTAG |
| 7303 | TOPBP1 | CGTTTGCTTTGGGGTAGGGG |
| 7304 | TOPBP1 | GGTAGGGGCGGCGCCGAGTC |
| 7305 | TP53TG1-001 | CAAGACGCCCACTGGAGACA |
| 7306 | TP53TG1-001 | CGCAGCCCAGAATAGAGCCG |
| 7307 | TP53TG1-001 | ACGCCCACTGGAGACAGGGT |
| 7308 | TP53TG1-001 | CCCGCAGCCCAGAATAGAGC |
| 7309 | TP53TG1-001 | GGACCCGACCCTGTCTCCAG |

| | | |
|---|---|---|
| 7310 | TP53TG1-001 | CCGGGGCCCAAGACGCCCAC |
| 7311 | TP53TG1-001 | TTGGGCCCCGGCTCTATTCT |
| 7312 | TP53TG1-003 | AATGAGCTGTCCTAACTCTG |
| 7313 | TP53TG1-003 | ACGAAGGTACCCAACCCTCT |
| 7314 | TP53TG1-003 | CTTCCTGCATCTATCCCAGA |
| 7315 | TP53TG1-003 | TGAGCTGTCCTAACTCTGCG |
| 7316 | TP53TG1-003 | CAACCCTCTGGGATAGATGC |
| 7317 | TP53TG1-003 | AGAGGGTTGGGTACCTTCGT |
| 7318 | TP53TG1-003 | ATGAGCTGTCCTAACTCTGC |
| 7319 | TPTEP1-009 | GGAAATGCGAGAGTCTGAGC |
| 7320 | TPTEP1-009 | AAATGCGAGAGTCTGAGCCG |
| 7321 | TPTEP1-009 | GGCGGGAGCCCCCCAAGAGG |
| 7322 | TPTEP1-009 | GGCGGCGGGAGCCCCCCAAG |
| 7323 | TPTEP1-009 | TGCGAGAGTCTGAGCCGGGG |
| 7324 | TPTEP1-009 | GATCCCGGTCCTCCTCTTGG |
| 7325 | TPTEP1-009 | ATCCCGGTCCTCCTCTTGGG |
| 7326 | TPX2 | TCGTTGTCAGATCTGAGGCG |
| 7327 | TPX2 | GCGCTTCGTTGTCAGATCTG |
| 7328 | TPX2 | CAGCCGTTCCAAACTCCGGG |
| 7329 | TPX2 | CTTCAGCCGTTCCAAACTCC |
| 7330 | TPX2 | ACTTCAGCCGTTCCAAACTC |
| 7331 | TPX2 | CGTTCCAAACTCCGGGAGGA |
| 7332 | TPX2 | AAGAAAAGAGGGAGCAGCTA |
| 7333 | TRAM2-AS1-003 | GCGGTGGAGAAAGCAGGCCG |
| 7334 | TRAM2-AS1-003 | GTGGAGAAAGCAGGCCGCGG |
| 7335 | TRAM2-AS1-003 | GGCCTGCCCGGACAGGAAGT |
| 7336 | TRAM2-AS1-003 | CCGCTCGCGGTGGAGAAAGC |
| 7337 | TRAM2-AS1-003 | GCTGGGCCCACTTCCTGTCC |
| 7338 | TRAM2-AS1-003 | AGCTGGGCCCACTTCCTGTC |
| 7339 | TRAM2-AS1-003 | AGGCCTGCCCGGACAGGAAG |
| 7340 | TRIM52-AS1-002 | CAGCACTAGCCCAGCTGGAG |
| 7341 | TRIM52-AS1-002 | AGCACTAGCCCAGCTGGAGC |
| 7342 | TRIM52-AS1-002 | CCCCCAGCCCACCCAACTCC |
| 7343 | TRIM52-AS1-002 | GCGCGGGCCGGGAGTTGGGT |
| 7344 | TRIM52-AS1-002 | ACTAGCCCAGCTGGAGCGGG |
| 7345 | TRIM52-AS1-002 | GGCCGCGCGGGCCGGGAGTT |
| 7346 | TRIM52-AS1-002 | CCGCGCCGCCCGCTCCAGCT |
| 7347 | TRIP13 | GAGCAGTGGGGACCCAACCA |
| 7348 | TRIP13 | GCGCGGCAGATTCGAAGCTA |
| 7349 | TRIP13 | GCAGATTCGAAGCTAGGGCG |
| 7350 | TRIP13 | CGGCGACGCTGGGCGTGAGG |
| 7351 | TRIP13 | TGGCGGCGACGCTGGGCGTG |
| 7352 | TRIP13 | GCGGCTGTGGCGGCGACGCT |

| | | |
|---|---|---|
| 7353 | TRIP13 | GGCTGAGGCAGCGGCTGTGG |
| 7354 | TRUB1 | AGCCCAGACCTGTTTCATCG |
| 7355 | TRUB1 | CAGGACCGGATCCGAAGCCA |
| 7356 | TRUB1 | CAGCAGCTTGGTAGCCAAAG |
| 7357 | TRUB1 | AAGTATGGCCGCTTCTGAGG |
| 7358 | TRUB1 | CCAGACCTGTTTCATCGTGG |
| 7359 | TRUB1 | ACCAAGCTGCTGTCCTTGAG |
| 7360 | TRUB1 | CAAAAGTATGGCCGCTTCTG |
| 7361 | TTK | TCGGAGCCAGGTATGCGACG |
| 7362 | TTK | TCCCCAGCGCAGCTTTCTGT |
| 7363 | TTK | ACCTACAGAAAGCTGCGCTG |
| 7364 | TTK | GTTCTGTCTTACCGGTGCAC |
| 7365 | TTK | TACCTACAGAAAGCTGCGCT |
| 7366 | TTK | ATTCAAACGTGTTTGCGGAA |
| 7367 | TTK | ACCGGTGCACCGGCGCACTG |
| 7368 | U91328.19-001 | GCTAGAAAACAAGCCTCGCG |
| 7369 | U91328.19-001 | GCGCTAGAAAACAAGCCTCG |
| 7370 | U91328.19-001 | AGAAAACAAGCCTCGCGGGG |
| 7371 | U91328.19-001 | CCAGGCTCACACCAGGCCTG |
| 7372 | U91328.19-001 | ACGGACCCCAGGCTCACACC |
| 7373 | U91328.19-001 | CAGGGCGACTGCAAGCTGTA |
| 7374 | U91328.19-001 | GCTCACACCAGGCCTGCGGG |
| 7375 | UBAP2L | GCGGGGCCGGGTATTGTCCG |
| 7376 | UBAP2L | GGCCCGGGGTGTCAGTGTGG |
| 7377 | UBAP2L | AAGGGGCCGCGGACAATACC |
| 7378 | UBAP2L | TCGCGGATTTAAAGGGGCCG |
| 7379 | UBAP2L | AGGTAAAGTTGGGAGCGCAG |
| 7380 | UBAP2L | GAGGAGACTGAGGTAAAGTT |
| 7381 | UBAP2L | GGAGGAGACTGAGGTAAAGT |
| 7382 | UBE2T | CAGCAACGCGCGGACCAACG |
| 7383 | UBE2T | GCGCGTTGCTGCGTTGTGAG |
| 7384 | UBE2T | CCTCACAACGCAGCAACGCG |
| 7385 | UBE2T | AGTGTACCTTCTTTCAGATG |
| 7386 | UBE2T | CCGCGCGTTGCTGCGTTGTG |
| 7387 | UBE2T | CCATCTGAAAGAAGGTACAC |
| 7388 | UBE2T | GAAAGAAGGTACACTGGAGA |
| 7389 | UBE2V2 | CCGGCCCGGGAACCGACCTG |
| 7390 | UBE2V2 | GGGGCAGAGCCGGGTCGGAG |
| 7391 | UBE2V2 | CGGGCTGCAGGAGAAGATGG |
| 7392 | UBE2V2 | GGGCGCCAGCAGCAGACGGA |
| 7393 | UBE2V2 | GAGCGGAAAATCACGCAGCC |
| 7394 | UBE2V2 | GGCGCCAGCAGCAGACGGAG |
| 7395 | UBE2V2 | AAAATCACGCAGCCCGGCCC |

| | | |
|---|---|---|
| 7396 | URB1-AS1-001 | TGTACCGGTTGTCCCGGTCA |
| 7397 | URB1-AS1-001 | GTTGTCCCGGTCACGGCATG |
| 7398 | URB1-AS1-001 | GCTCGCCCCATGCCGTGACC |
| 7399 | URB1-AS1-001 | CTTGGGGATCCAGAGTCTTG |
| 7400 | URB1-AS1-001 | GTTTGCTGTACCGGTTGTCC |
| 7401 | URB1-AS1-001 | ACCGGTACAGCAAACTCACA |
| 7402 | URB1-AS1-001 | CGGTCACGGCATGGGGCGAG |
| 7403 | USP1 | CGGAGTGCTAAAGACCCTAG |
| 7404 | USP1 | AAAACGCGCCAAGTTCCCCT |
| 7405 | USP1 | TCTTTAGCACTCCGCCACCG |
| 7406 | USP1 | CTTTAGCACTCCGCCACCGA |
| 7407 | USP1 | CGGTTCAGGCGTTCGGCGAG |
| 7408 | USP1 | TTTAGCACTCCGCCACCGAG |
| 7409 | USP1 | ACGCGCCAAGTTCCCCTCGG |
| 7410 | USP21 | CTGCGGCTAAAGCTGCAGCC |
| 7411 | USP21 | GGCGGGAGCGATCTGGGCAA |
| 7412 | USP21 | ACGGTGAAGCCGGTGGCCGG |
| 7413 | USP21 | GGCGGGACCAACAAAGATGG |
| 7414 | USP21 | CCATCTTTGTTGGTCCCGCC |
| 7415 | USP21 | AGCGATCTGGGCAACGGCTG |
| 7416 | USP21 | CCCAGATCGCTCCCGCCGCA |
| 7417 | USP37 | AGTAGGGATACCGAACCCAA |
| 7418 | USP37 | CCCTGACTAGGGGTAGCTAG |
| 7419 | USP37 | AGGGATACCGAACCCAAAGG |
| 7420 | USP37 | CTACCCCTAGTCAGGGAGAG |
| 7421 | USP37 | CCGCTAGCTACCCCTAGTCA |
| 7422 | USP37 | GGGATACCGAACCCAAAGGA |
| 7423 | USP37 | CAGCCGCTCTCCCTGACTAG |
| 7424 | WDR76 | AAGCCGAAAGATGTCCAGGT |
| 7425 | WDR76 | CCGTTTCTCACCTTCATCTG |
| 7426 | WDR76 | AGCCGAAAGATGTCCAGGTC |
| 7427 | WDR76 | CAGACAGCGACCCCAGATGA |
| 7428 | WDR76 | TAAGAAGCCGAAAGATGTCC |
| 7429 | WDR76 | GATGAAGGTGAGAAACGGAC |
| 7430 | WDR76 | TCTTTCGGCTTCTTAGCGGC |
| 7431 | XRCC5 | ACCGGAAGCGAGTTGCGACA |
| 7432 | XRCC5 | TGAGGACCGGCAACATGGTG |
| 7433 | XRCC5 | GAAGCGAGTTGCGACACGGC |
| 7434 | XRCC5 | GACCGGCAACATGGTGCGGT |
| 7435 | XRCC5 | GCGCCTGAGGACCGGCAACA |
| 7436 | XRCC5 | CGACCAAAGCGCCTGAGGAC |
| 7437 | XRCC5 | CGACACGGCAGGTTCCCGCC |
| 7438 | XX-C2158C6.3-001 | AGGCTGTAGAATGTGCACGG |

| | | |
|---|---|---|
| 7439 | XX-C2158C6.3-001 | ATTGGCCAGTGCAAGTCACA |
| 7440 | XX-C2158C6.3-001 | GAGGCTGTAGAATGTGCACG |
| 7441 | XX-C2158C6.3-001 | TGAGCAGACCTGCATGTCTG |
| 7442 | XX-C2158C6.3-001 | AACAAGTCTGATGTGCACAG |
| 7443 | XX-C2158C6.3-001 | TTGGCTCTGGTTCCTTCGTG |
| 7444 | XX-C2158C6.3-001 | CAGAGGCTGTAGAATGTGCA |
| 7445 | XXbac-BPG181B23.7-001 | CTTTGTATGTACACACACAG |
| 7446 | XXbac-BPG181B23.7-001 | TTGCAAGCAATGCCTAGGTG |
| 7447 | XXbac-BPG181B23.7-001 | TGTTGCAAGCAATGCCTAGG |
| 7448 | XXbac-BPG181B23.7-001 | ACTTTGTATGTACACACACA |
| 7449 | XXbac-BPG181B23.7-001 | GACTGTTGCAAGCAATGCCT |
| 7450 | XXbac-BPG181B23.7-001 | CCTAGGAAACATCCCCACCT |
| 7451 | XXbac-BPG181B23.7-001 | TAGGCATTGCTTGCAACAGT |
| 7452 | XXyac-YX155B6.5-008 | AGTTAGAACATTAAATACAC |
| 7453 | XXyac-YX155B6.5-008 | ATACTATCAAGTACAGCACG |
| 7454 | XXyac-YX155B6.5-008 | CCTCTGATGTTTCATCAGCG |
| 7455 | XXyac-YX155B6.5-008 | GATTAAAACTGGAAAGTCTG |
| 7456 | XXyac-YX155B6.5-008 | TTCCTCTGATGTTTCATCAG |
| 7457 | XXyac-YX155B6.5-008 | GATGTTTCATCAGCGGGGAC |
| 7458 | XXyac-YX155B6.5-008 | AGTGCCACATTAAGATAACA |
| 7459 | YBX1 | GATCGAACTAGCGAGAATGG |
| 7460 | YBX1 | GCCATTCTCGCTAGTTCGAT |
| 7461 | YBX1 | ATCGAACTAGCGAGAATGGC |
| 7462 | YBX1 | TGTGATGGTAACTAGGCCGG |
| 7463 | YBX1 | GATGGTAACTAGGCCGGCGG |
| 7464 | YBX1 | TCGGTAGCGGGAGCGGAGAG |
| 7465 | YBX1 | AGTTCGATCGGTAGCGGGAG |
| 7466 | YTHDF3-AS1-001 | CGGCCGCCGTGACTTCAGCA |
| 7467 | YTHDF3-AS1-001 | CTTCCTTCTATCCCGCTCCG |
| 7468 | YTHDF3-AS1-001 | GCAGAGTGGCGTCAGCTGGA |
| 7469 | YTHDF3-AS1-001 | GCAGAAAAGACGCTGAAAGT |
| 7470 | YTHDF3-AS1-001 | AGGGCCCTGCAGCCTCGGAG |
| 7471 | YTHDF3-AS1-001 | CAGCTGACGCCACTCTGCCA |
| 7472 | YTHDF3-AS1-001 | CAGCCTCGGAGCGGGATAGA |
| 7473 | YY1AP1 | GCCGCTGAAGGTACGCGTGA |
| 7474 | YY1AP1 | CCCCTCAGACCCTGTCAAGC |
| 7475 | YY1AP1 | CGGCTTGACAGGGTCTGAGG |
| 7476 | YY1AP1 | CCGGCTTGACAGGGTCTGAG |
| 7477 | YY1AP1 | CGCCGCTGAAGGTACGCGTG |
| 7478 | YY1AP1 | GCGCTGGAGCCGGCTTGACA |
| 7479 | YY1AP1 | AGAGGAGGTGGAGAAGGCTT |
| 7480 | ZNF273 | TCTATGTCCTCTGCTCCTAG |
| 7481 | ZNF273 | TCCTAGAGGTCCACCTTCTG |

| | | |
|---|---|---|
| 7482 | ZNF273 | GCCACAGAAGGTGGACCTCT |
| 7483 | ZNF273 | GCAGGTAACGGGGCCACAGA |
| 7484 | ZNF273 | GGTAACGGGGCCACAGAAGG |
| 7485 | ZNF273 | CCCAATACCTGCAGGTAACG |
| 7486 | ZNF273 | CTCCCAATACCTGCAGGTAA |
| 7487 | ZNF330 | ACAGCAGTAAATCCAACTCG |
| 7488 | ZNF330 | GACAGCCGATTCCGGACACT |
| 7489 | ZNF330 | GTAAATCCAACTCGTGGACC |
| 7490 | ZNF330 | AGTTGGATTTACTGCTGTCG |
| 7491 | ZNF330 | GATTTACTGCTGTCGCGGGT |
| 7492 | ZNF330 | AAAGAAGGTACTAACAGCCA |
| 7493 | ZNF330 | GAAGGTACTAACAGCCAGGG |
| 7494 | ZNF674-AS1-001 | CTGTCATAACCACTGAAGCG |
| 7495 | ZNF674-AS1-001 | TGTCATAACCACTGAAGCGC |
| 7496 | ZNF674-AS1-001 | AACAAAGCCTGTGGCCGATG |
| 7497 | ZNF674-AS1-001 | ACGCGCAGTGGCTTTGAAGG |
| 7498 | ZNF674-AS1-001 | AGGCGGGGCAACAAAGCCTG |
| 7499 | ZNF674-AS1-001 | GTGGCCGATGTGGCGCTCCT |
| 7500 | ZNF674-AS1-001 | TTATGACAGTGTTCGCTCTA |

| Seq# | Gene Symbol | sgRNA |
|---|---|---|
| 1 | Non_Targeting_Human | CTGCATGGGGCGCGAATCA |
| 2 | Non_Targeting_Human | TGCACCCGGCTAGGACCGG |
| 3 | Non_Targeting_Human | TGTGCAACCTCCGCCGTTG |
| 4 | Non_Targeting_Human | CCCGAGCAGTGGCTCGCTA |
| 5 | Non_Targeting_Human | GTCAGGTAGAGGGATTGAG |
| 6 | Non_Targeting_Human | GAGGACGATCGTACTCCAG |
| 7 | Non_Targeting_Human | TATGGAGGGCTGGATCTGC |
| 8 | Non_Targeting_Human | CCTTGGCTAAACCGCTCCC |
| 9 | Non_Targeting_Human | GAGTCGGGTAAATAGACAA |
| 10 | Non_Targeting_Human | AGTGGACGACCCTGTCAGG |
| 11 | Non_Targeting_Human | TATCGGTCTGTACGGGTAG |
| 12 | Non_Targeting_Human | TGGTACGGGGCCGGCCTGT |
| 13 | Non_Targeting_Human | CAGTCGCCGCGCATCGGGC |
| 14 | Non_Targeting_Human | AGAGGTCGGATACGCTTCC |
| 15 | Non_Targeting_Human | GCCCTACATCGCTAGGCTA |
| 16 | Non_Targeting_Human | CGCCGGTTCCGTCGACTAG |
| 17 | Non_Targeting_Human | AGCAAGGTTCGGCAAAGCG |
| 18 | Non_Targeting_Human | CCAGTACTGACAGCCTTAT |
| 19 | Non_Targeting_Human | GTCGCCGTAGGGTCCGCCA |
| 20 | Non_Targeting_Human | CTAGAATGGTTGCGCGAAC |
| 21 | Non_Targeting_Human | AGCCTAACGATCGGACCGA |
| 22 | Non_Targeting_Human | CCGCTCCATCAGACTTTCT |
| 23 | Non_Targeting_Human | CTACGCCCGGGGGAAAAGA |
| 24 | Non_Targeting_Human | CAGGAGCTAAGGGTCCCGT |
| 25 | Non_Targeting_Human | TACTCCCCGAATGCAGCGG |
| 26 | Non_Targeting_Human | AGGCCGCAAACCGCGTGCG |
| 27 | Non_Targeting_Human | AGCCTCTTGGTTCCATGGA |
| 28 | Non_Targeting_Human | CTCCCCCGCTTCACTTGGG |
| 29 | Non_Targeting_Human | GTGGCCTTTGCAATTGGCG |
| 30 | Non_Targeting_Human | CGCTAGGAAACCAACAAGG |
| 31 | Non_Targeting_Human | GGTGGAGTGCAGCCACAAA |
| 32 | Non_Targeting_Human | TTCCGATGGTTAAGCACGA |
| 33 | Non_Targeting_Human | TGCGTACCACACGACACAT |
| 34 | Non_Targeting_Human | AGAACTCCCGGCAATTAGG |
| 35 | Non_Targeting_Human | TCATTCACGTTGGGTTTTC |
| 36 | Non_Targeting_Human | ATCTCCGGGGCGTTAGGCC |
| 37 | Non_Targeting_Human | CAGCGAGCAGCTTTTACGA |
| 38 | Non_Targeting_Human | ACGTGTGTTACGGGGGGAA |
| 39 | Non_Targeting_Human | GAGAGGAAGTCGTCGGAAA |
| 40 | Non_Targeting_Human | CGTACGACAATACGCGCGA |
| 41 | Non_Targeting_Human | GACTGGAGAGTGCCCGGCC |
| 42 | Non_Targeting_Human | CAATTCCCCGCAATCCAGC |

| | | |
|---|---|---|
| 43 | Non_Targeting_Human | TCAAAAGCAACGTACGCAA |
| 44 | Non_Targeting_Human | TACCAGCACGGAGCGGCAG |
| 45 | Non_Targeting_Human | CAGGCAGCCGCTGAAGCAT |
| 46 | Non_Targeting_Human | AGGGGTCGTCGGGCAACCC |
| 47 | Non_Targeting_Human | TCACGGGCGTGCTCGAGTC |
| 48 | Non_Targeting_Human | TACATTGCTCTGGGGCGTA |
| 49 | Non_Targeting_Human | GTGGTGACAAGTCCTGAGG |
| 50 | Non_Targeting_Human | ACACAGGCCGTTAAAAAGA |
| 51 | Non_Targeting_Human | GTGATCAAATGCCTATGAT |
| 52 | Non_Targeting_Human | GAGCTAGCGGAATGCATGG |
| 53 | Non_Targeting_Human | GTGTGACCTCGTCTGGCAG |
| 54 | Non_Targeting_Human | CGTGATGGGCACGAGTAAG |
| 55 | Non_Targeting_Human | GGCCTGGACGAGCCTAAAA |
| 56 | Non_Targeting_Human | GATCGCGTCTAGTGCAGAT |
| 57 | Non_Targeting_Human | AACCTGAGACGCTAAGTGA |
| 58 | Non_Targeting_Human | CCCCCTACGACAGGTACAT |
| 59 | Non_Targeting_Human | CGGCGAGGGAGAATGCCCT |
| 60 | Non_Targeting_Human | CCACAGGGAAGCGAATACG |
| 61 | Non_Targeting_Human | GGGAAGCGTCGCGCCGGAA |
| 62 | Non_Targeting_Human | ATGTCGTAGGAGAAGGACG |
| 63 | Non_Targeting_Human | CCCAAAGGGCGGCAAGCAG |
| 64 | Non_Targeting_Human | TGAGACACCGTACACGATT |
| 65 | Non_Targeting_Human | CGCCCTGGGCCCCACCAGC |
| 66 | Non_Targeting_Human | GCGGCGACGCACGCTACCT |
| 67 | Non_Targeting_Human | GCAGAGGCAAGGGCATAAA |
| 68 | Non_Targeting_Human | GCTCATGCGCGCGAACGCC |
| 69 | Non_Targeting_Human | CCTCCCCCATGCTGAGTAA |
| 70 | Non_Targeting_Human | TGTTTAGCCGCGGGCGGAC |
| 71 | Non_Targeting_Human | GCCTCGGGCGTACATTAAG |
| 72 | Non_Targeting_Human | TCAGCGGGCCGCATCTTGT |
| 73 | Non_Targeting_Human | GGCGCTTTGAGACGATGGG |
| 74 | Non_Targeting_Human | CGGCTCCATGGGCGGCCGT |
| 75 | Non_Targeting_Human | GGTGGGTCACGCGGAAGGG |
| 76 | Non_Targeting_Human | GATCGCTTCAATCACCGAG |
| 77 | Non_Targeting_Human | GAAGGTGCTGAATGTGCCG |
| 78 | Non_Targeting_Human | GAAACGACGGAGTATTCCG |
| 79 | Non_Targeting_Human | AAGCACCCATCGCCGGCTG |
| 80 | Non_Targeting_Human | GTCTAGTATCTTGAGCGTT |
| 81 | Non_Targeting_Human | TGCCACTCACGAGTTGACT |
| 82 | Non_Targeting_Human | TCCCGTTGAGAGAGGACAC |
| 83 | Non_Targeting_Human | ACGTTGAGGGACTGTCCTA |
| 84 | Non_Targeting_Human | ATCCTGACGGGGACTACAT |
| 85 | Non_Targeting_Human | GGTTGCGCGGGGTGGCGTA |

| | | |
|---|---|---|
| 86 | Non_Targeting_Human | CATCACCCGATGTACCGAG |
| 87 | Non_Targeting_Human | TACCGATTCCCGGCCAAGA |
| 88 | Non_Targeting_Human | ATAACCACAAGGCGCTCGA |
| 89 | Non_Targeting_Human | AGGACCCAGTAGACCGGCT |
| 90 | Non_Targeting_Human | GGGTGAGGGTCCAATTCGG |
| 91 | Non_Targeting_Human | TAGCTATGCGGTAGGACGG |
| 92 | Non_Targeting_Human | TAGACGTGCAAAAGCCGCA |
| 93 | Non_Targeting_Human | GATTAACCCAGGGCTTTGA |
| 94 | Non_Targeting_Human | TGCTAATGGGGAAAGCCGG |
| 95 | Non_Targeting_Human | CGATGGGACAGACGCCGGA |
| 96 | Non_Targeting_Human | CCGGGGCCTTGGAAAACAC |
| 97 | Non_Targeting_Human | TGACAGGATGCGTTCACGG |
| 98 | Non_Targeting_Human | CGACATCGGTTACATGTGG |
| 99 | Non_Targeting_Human | GTCGATGTGGGGACGGCAC |
| 100 | Non_Targeting_Human | TGCGCAAACCGGATCGGGG |
| 101 | Non_Targeting_Human | GCAAGATAGACGCTACACA |
| 102 | Non_Targeting_Human | CGACAATTGCAGCTGGACC |
| 103 | Non_Targeting_Human | ATCCGTCACTGACCCTTCC |
| 104 | Non_Targeting_Human | CGCAGCGGTTGTCAAACGA |
| 105 | Non_Targeting_Human | AGACTCTAGGTCCAGGAGT |
| 106 | Non_Targeting_Human | GGGAAAGAGAATCGCCCTA |
| 107 | Non_Targeting_Human | AAACTCCGCAGGCCAAGTA |
| 108 | Non_Targeting_Human | GGCAGGGCTACGATGCAGG |
| 109 | Non_Targeting_Human | GCCGGGTATTCCTGAGGGG |
| 110 | Non_Targeting_Human | TCGGTTGGACAATTTCCCG |
| 111 | Non_Targeting_Human | GCGCGTGGTTGCATTAGAC |
| 112 | Non_Targeting_Human | TAGACAGCAGGGGCGTTTG |
| 113 | Non_Targeting_Human | GTAAGGAAAAACGGCGGTG |
| 114 | Non_Targeting_Human | TTCAGGGTGGCCGACAGGC |
| 115 | Non_Targeting_Human | ATGAGTTCGCGAGGCCGTA |
| 116 | Non_Targeting_Human | CTTCTAGGAGGCACTACGG |
| 117 | Non_Targeting_Human | CAGGAGCTCAACGACGGTG |
| 118 | Non_Targeting_Human | AGCTCTTTGACGGACTGTG |
| 119 | Non_Targeting_Human | GTATCGGGTGGCGTTAATT |
| 120 | Non_Targeting_Human | GCGCCACGAGGATCAAAGC |
| 121 | Non_Targeting_Human | CGATCTGGTTGCACAACTA |
| 122 | Non_Targeting_Human | AGGGCCCGCTCGTCCTCAG |
| 123 | Non_Targeting_Human | GAGATCAGGATGATCGTAC |
| 124 | Non_Targeting_Human | GCTAGAGTGGCGCAGGGAG |
| 125 | Non_Targeting_Human | GGCGGGTCGGACATCGCCT |
| 126 | Non_Targeting_Human | TTCTACACATTGGAAATAA |
| 127 | Non_Targeting_Human | GGCTAATCCCGGTGAAAAA |
| 128 | Non_Targeting_Human | CGCCAGACCGGCCAGCGAG |

| | | |
|---|---|---|
| 129 | Non_Targeting_Human | TCGGTTCGAGCGAAACGTG |
| 130 | Non_Targeting_Human | GAGAGTCGGGAGGGATCGT |
| 131 | Non_Targeting_Human | CGCGTCGTTACAGGGGCGG |
| 132 | Non_Targeting_Human | CACGCCACGAGACGTTGGA |
| 133 | Non_Targeting_Human | CGAGGAGCAGTTCAACACC |
| 134 | Non_Targeting_Human | TGACGTTGTGAGGACGGTC |
| 135 | Non_Targeting_Human | CCCACCCGAACGGAGGCGG |
| 136 | Non_Targeting_Human | GCCGGGCGCATATCCTCAC |
| 137 | Non_Targeting_Human | GCCATGCGGAAGATGTGGA |
| 138 | Non_Targeting_Human | ACTGTTTGCGCGTGTAGAG |
| 139 | Non_Targeting_Human | CACACAGGCCTAGCGATCT |
| 140 | Non_Targeting_Human | CCCTGGCAAACGAAGGGGG |
| 141 | Non_Targeting_Human | AGGCGTCAGTGAGCGGTGT |
| 142 | Non_Targeting_Human | TGTAACTGCCACTCCTAGG |
| 143 | Non_Targeting_Human | CCCGGGGGGGGTCGTAGGT |
| 144 | Non_Targeting_Human | TGTCCAGTTCCGTAGGATG |
| 145 | Non_Targeting_Human | CCTTGCCGCGACGAAAACT |
| 146 | Non_Targeting_Human | GACCTGAGTCGTGACACAG |
| 147 | Non_Targeting_Human | AAGCACTATACGGACGGAG |
| 148 | Non_Targeting_Human | GTATGGAGGATCTTCGTGG |
| 149 | Non_Targeting_Human | CGCGCTGACTCGTTACGAG |
| 150 | Non_Targeting_Human | CGGGCGCTCTATCTTGCAG |
| 151 | Non_Targeting_Human | CGATTTGTCCCGGCAAGGG |
| 152 | Non_Targeting_Human | CGTATACCGGGTCCAAGAG |
| 153 | Non_Targeting_Human | CCTCATATGCTACTTTATG |
| 154 | Non_Targeting_Human | CCCCTCGTTGAAGCCGGCT |
| 155 | Non_Targeting_Human | GACTACTCGTGGCGCTTGC |
| 156 | Non_Targeting_Human | GCGGCGGAGGCCCCGCCAA |
| 157 | Non_Targeting_Human | GGCCTCATATAGAACACTG |
| 158 | Non_Targeting_Human | CTCGTAAGGAGGATATGCG |
| 159 | Non_Targeting_Human | ATGTTGTATCAGCCAGTAC |
| 160 | Non_Targeting_Human | TGGACGAGACGGCGGACGG |
| 161 | Non_Targeting_Human | CGGGCGTTAACGCGTGCAA |
| 162 | Non_Targeting_Human | TGGGCGGCTCGGAGAACTG |
| 163 | Non_Targeting_Human | CGATGCGCCGACCCGTCAA |
| 164 | Non_Targeting_Human | AGTCACGCGGCTCCCATCG |
| 165 | Non_Targeting_Human | GACGGGGACCCGGTTGGCA |
| 166 | Non_Targeting_Human | AGCCCCGCCTGGGTACGCG |
| 167 | Non_Targeting_Human | CCCGATGCCGCAGTCCATA |
| 168 | Non_Targeting_Human | AAAGCGGGATAACTTGAGG |
| 169 | Non_Targeting_Human | GCGCCGGTTAACAATAACT |
| 170 | Non_Targeting_Human | GATGGTTTCGCGTACATAG |
| 171 | Non_Targeting_Human | CCTGCGCGACAAGCGGACG |

| | | |
|---|---|---|
| 172 | Non_Targeting_Human | CAGGGGCCTGCGCCAGCGG |
| 173 | Non_Targeting_Human | GCAGGTACAACGAGCGTCA |
| 174 | Non_Targeting_Human | GCGCCCCAGTCGGCCCCTA |
| 175 | Non_Targeting_Human | ACAATACTGTCGATGGCAT |
| 176 | Non_Targeting_Human | GCGTGTCCTGAGCCGTGAA |
| 177 | Non_Targeting_Human | CTACGCCGACACAAAGATG |
| 178 | Non_Targeting_Human | GACCCTGACATCAAGACAG |
| 179 | Non_Targeting_Human | CCATGCCTCGTACAGGAGA |
| 180 | Non_Targeting_Human | CTATTAAGGTTATGGCCGA |
| 181 | Non_Targeting_Human | TATGGCTAAGGGCGGGCGT |
| 182 | Non_Targeting_Human | TACCGCACGAAATGCTCCG |
| 183 | Non_Targeting_Human | GGGGAATCTGCAAGACCAT |
| 184 | Non_Targeting_Human | TCGGAGTAATGACCACAGC |
| 185 | Non_Targeting_Human | GGAGCTCCGGATTGCCGAA |
| 186 | Non_Targeting_Human | CCACCCGGTGGGTGACGGG |
| 187 | Non_Targeting_Human | AGACGTGGCACGTCTCGAC |
| 188 | Non_Targeting_Human | AAAAATGAGCCATGGGCGG |
| 189 | Non_Targeting_Human | ACCGGTCGGCGGACGTGGG |
| 190 | Non_Targeting_Human | TGCGCGCCGCCCAATTCTT |
| 191 | Non_Targeting_Human | TGTGGCTCGGCCAGCGATA |
| 192 | Non_Targeting_Human | TGTACGGCACCCGTGGCAA |
| 193 | Non_Targeting_Human | GGGCGGGGGCCGGGCGGAT |
| 194 | Non_Targeting_Human | TGCCAGCTTGTGGTGTCGT |
| 195 | Non_Targeting_Human | GCCGTAACGGTGTCACACT |
| 196 | Non_Targeting_Human | GAGATGTCCAGGCGTACAG |
| 197 | Non_Targeting_Human | CGTGGGCTTACTAAGCAGC |
| 198 | Non_Targeting_Human | TGCACACGGCAGTAGAAAA |
| 199 | Non_Targeting_Human | GAGTTGCAGCCGTCACGAG |
| 200 | Non_Targeting_Human | ACCGCCGCGCGGCCTAACG |
| 201 | Non_Targeting_Human | TCAGCGGAGATGTGCACTG |
| 202 | Non_Targeting_Human | CTACCCCTGATTGCCAGCC |
| 203 | Non_Targeting_Human | GAGGCCTGTACGCGGAGGC |
| 204 | Non_Targeting_Human | GTCGGGCGCATCACGCGGG |
| 205 | Non_Targeting_Human | TTCGCACCGGCCACCACAC |
| 206 | Non_Targeting_Human | ATGGCATACACTTCGCGAG |
| 207 | Non_Targeting_Human | GCCTGGCCCTCGGCAACTG |
| 208 | Non_Targeting_Human | GGAGCCAAGCCGATGCACA |
| 209 | Non_Targeting_Human | CGCCCGGAAACGTTAACCT |
| 210 | Non_Targeting_Human | GTTCGCTTACACGCCGTCC |
| 211 | Non_Targeting_Human | ACACTGAGGGCCGCAGCCG |
| 212 | Non_Targeting_Human | AACAGTTTAAGGAGCACCG |
| 213 | Non_Targeting_Human | CAACAAGCGCGCGTAGGTA |
| 214 | Non_Targeting_Human | AGCGCCGACAAGCTGACGC |

| | | |
|---|---|---|
| 215 | Non_Targeting_Human | TCCGCCGAGTCTAAACTTC |
| 216 | Non_Targeting_Human | GCGTTGGCTTAAGCTGTAG |
| 217 | Non_Targeting_Human | TGAACTCAAAAATCCCGAC |
| 218 | Non_Targeting_Human | GCCTGGCGTGAAACGGCCG |
| 219 | Non_Targeting_Human | TGACAGCCCCGTAACTGAG |
| 220 | Non_Targeting_Human | TCAAACGCGTGCACCTTGA |
| 221 | Non_Targeting_Human | ATTGACCGGACGGTGTCGC |
| 222 | Non_Targeting_Human | GCGGAACCGACGGTCACTG |
| 223 | Non_Targeting_Human | AGCAGAACGGATGCGCGTC |
| 224 | Non_Targeting_Human | ACGAAGGGCAAAATTCCAG |
| 225 | Non_Targeting_Human | AAGCATAATGGACGGAAGT |
| 226 | Non_Targeting_Human | GAGGGGCGAGCCGTCGCCG |
| 227 | Non_Targeting_Human | CGTGACATGCCGATCCCTG |
| 228 | Non_Targeting_Human | GGAGTGCCAGGCCGAGAGG |
| 229 | Non_Targeting_Human | GTCCAGCTAATATTGCAGG |
| 230 | Non_Targeting_Human | CCCAGCGAAGCGGGGACCT |
| 231 | Non_Targeting_Human | TCGTCTGGAGGTGCGACGG |
| 232 | Non_Targeting_Human | AACGGTATAGCGGCATGCG |
| 233 | Non_Targeting_Human | GGGCCCCGGAGGCAGAACG |
| 234 | Non_Targeting_Human | GAGCCCCGACGCGCGCGTC |
| 235 | Non_Targeting_Human | TTACCTCAGATATGCGGGG |
| 236 | Non_Targeting_Human | ACTTGCGACCCCGAAGAAT |
| 237 | Non_Targeting_Human | ACGCCCCATGCTGAAACCG |
| 238 | Non_Targeting_Human | TGCGGGCCCGCTTCTGCAG |
| 239 | Non_Targeting_Human | ATTGTCACTTTAGATCTGT |
| 240 | Non_Targeting_Human | GCAGCGACCGGCATGCCGC |
| 241 | Non_Targeting_Human | CGAGCGGTTGCCGCCCTGG |
| 242 | Non_Targeting_Human | TTGAATTACAGTTCGACGG |
| 243 | Non_Targeting_Human | TATGGGCGCAAGGGTACAG |
| 244 | Non_Targeting_Human | GCGCTTGGTGCGCGACACG |
| 245 | Non_Targeting_Human | CGGGCAGCGATAGTGGCTG |
| 246 | Non_Targeting_Human | CGGCATTGACATAGGGTCA |
| 247 | Non_Targeting_Human | CGGTGGGCTCCCGGACCGC |
| 248 | Non_Targeting_Human | AACAAGTAACCCAGGCGCC |
| 249 | Non_Targeting_Human | CGGGTGGACCATGACTACG |
| 250 | Non_Targeting_Human | CGAGCCTAGAACCCCGCTC |
| 251 | Non_Targeting_Human | ACCACGCCGGCCCCGTTGA |
| 252 | Non_Targeting_Human | CGACACCGGTAAAGGCACG |
| 253 | Non_Targeting_Human | CCCCCCAGGAGGGCATTTT |
| 254 | Non_Targeting_Human | TACTTGGGGGAGTCGGTGA |
| 255 | Non_Targeting_Human | TCGGGTAACAAGAGGAATC |
| 256 | Non_Targeting_Human | TTTGACGCGCGTCATCCAG |
| 257 | Non_Targeting_Human | GTAGCAGTTACAGGCTCGC |

| | | |
|---|---|---|
| 258 | Non_Targeting_Human | CGTAGCGCCCTCACAGTGG |
| 259 | Non_Targeting_Human | AAGGATGTGTCCCGAATTG |
| 260 | Non_Targeting_Human | TGGGATCGTGCGCGCCCCA |
| 261 | Non_Targeting_Human | GCTATACCACGTTCAGTCA |
| 262 | Non_Targeting_Human | TGCCCGCACGAAAGTGGGT |
| 263 | Non_Targeting_Human | TATGGCGGGGATGCACGAA |
| 264 | Non_Targeting_Human | TCGGTAAGCGACCCCATAG |
| 265 | Non_Targeting_Human | TTAGCGCGCTAGTATATAG |
| 266 | Non_Targeting_Human | CGGACGAACCAGGCTAAAG |
| 267 | Non_Targeting_Human | AGCATCATTGATCTTCCGG |
| 268 | Non_Targeting_Human | CTAGGAGTCGGATCTACTG |
| 269 | Non_Targeting_Human | CACACTCCGCGGGGAAGTG |
| 270 | Non_Targeting_Human | ATACGCTACGTAGCGAAAA |
| 271 | Non_Targeting_Human | GCGCGATCATGTGATAAGT |
| 272 | Non_Targeting_Human | AACGTCGCGGGCGCCCGGA |
| 273 | Non_Targeting_Human | AATCCCCTACTGGGATCAC |
| 274 | Non_Targeting_Human | CCCACGGAGTAGCGCACCG |
| 275 | Non_Targeting_Human | CAAAACTCAGCTACCAGGT |
| 276 | Non_Targeting_Human | GCCGGCCCGGACCCCTAGT |
| 277 | Non_Targeting_Human | GGTGCCCACAGCGCCGTCA |
| 278 | Non_Targeting_Human | GTACTGCGAGGCCGTTATG |
| 279 | Non_Targeting_Human | TAATCGAATCAGGGCTCAC |
| 280 | Non_Targeting_Human | TGTTAACACCCCAGCCACT |
| 281 | Non_Targeting_Human | GATCGTAGACAATAAGCGG |
| 282 | Non_Targeting_Human | GGCCGACGGATAGGAGGGA |
| 283 | Non_Targeting_Human | CACGGACGCGATGACACGC |
| 284 | Non_Targeting_Human | TCAGGAAGAGCGCGGGAGT |
| 285 | Non_Targeting_Human | TTTTGCCCGCGGCGTAACG |
| 286 | Non_Targeting_Human | GTGCCTCACTTGTTCGACG |
| 287 | Non_Targeting_Human | GTACAAGTGCCCACAAGTG |
| 288 | Non_Targeting_Human | CCCCTGACCGGCACCCTCG |
| 289 | Non_Targeting_Human | CCGCGCCCTGGGGTCGAGA |
| 290 | Non_Targeting_Human | CGCGCTGAGGTACATGGGG |
| 291 | Non_Targeting_Human | GCGAACGTCGACCAGCCCT |
| 292 | Non_Targeting_Human | AGCTTCCCGGGTCGACCGC |
| 293 | Non_Targeting_Human | TTAGCCGGACAAGAACGAT |
| 294 | Non_Targeting_Human | TCAGAGAGATAAGCTATCG |
| 295 | Non_Targeting_Human | CGTGGGCGTGCGGACTCAG |
| 296 | Non_Targeting_Human | GTCGCCGTGCGCCTACCGA |
| 297 | Non_Targeting_Human | CGGACTAGGCGTAGCGGGT |
| 298 | Non_Targeting_Human | CGAGTATGTCGCCTTGCAG |
| 299 | Non_Targeting_Human | GACGGAACGCGGACAGCAA |
| 300 | Non_Targeting_Human | AAGTAGCTGAGTCATCACG |

| | | |
|---|---|---|
| 301 | Non_Targeting_Human | CTCCGACCCCCGAGATGAG |
| 302 | Non_Targeting_Human | GGGCGAGACGACCGAGCAA |
| 303 | Non_Targeting_Human | GACCGATCGAGTATACCGT |
| 304 | Non_Targeting_Human | ATTGTTCAGGCGAGGGTGT |
| 305 | Non_Targeting_Human | ACGGCGCGTGCGCTTAGTG |
| 306 | Non_Targeting_Human | TATCCTCTACCTGCTACGT |
| 307 | Non_Targeting_Human | CGACAGCGGCTCGCCTGCG |
| 308 | Non_Targeting_Human | CTCCCCGCAGGGTTAGTCG |
| 309 | Non_Targeting_Human | CAGCATTTGAACACGGAAG |
| 310 | Non_Targeting_Human | CAAAAACTTGTGCAGCTAG |
| 311 | Non_Targeting_Human | CTTACCACCGACGACCAAT |
| 312 | Non_Targeting_Human | TGCGGTGCCGAGCCCTATG |
| 313 | Non_Targeting_Human | AACGGGGGGGCCCTTCGG |
| 314 | Non_Targeting_Human | CTTGTAACCTGGGACGGCT |
| 315 | Non_Targeting_Human | ATCGACAGTCTAGCGGCTG |
| 316 | Non_Targeting_Human | GGTCGACCACCGCCCTCTG |
| 317 | Non_Targeting_Human | GCGCAGACGCCGACCAGAG |
| 318 | Non_Targeting_Human | TGAGGCAGTCTGAGTGCGA |
| 319 | Non_Targeting_Human | GAGGATACCCGGGGAACCT |
| 320 | Non_Targeting_Human | GGCACGTTCGCGAAAACGA |
| 321 | Non_Targeting_Human | ACAAGTCTAGGGCCTCCCC |
| 322 | Non_Targeting_Human | AGAGGGCATGGCGCATGCA |
| 323 | Non_Targeting_Human | CTGGGCCTGGCAAATTGAA |
| 324 | Non_Targeting_Human | GTCAGGGGTTTGGTCGGCA |
| 325 | Non_Targeting_Human | CCGCGTTTTACGCGACCGA |
| 326 | Non_Targeting_Human | GTTCTCGCGGCTGTGCGGT |
| 327 | Non_Targeting_Human | GCAGGGCCCAGTCGTGCAG |
| 328 | Non_Targeting_Human | TCGGTACGGCGGTTGATCG |
| 329 | Non_Targeting_Human | ACTGGCCACCGGCACCTGG |
| 330 | Non_Targeting_Human | GCCCCGGACGTCCTGACCC |
| 331 | Non_Targeting_Human | AAAGGATGGTGATATCTCG |
| 332 | Non_Targeting_Human | AGGGAAAATTGACGTTCGG |
| 333 | Non_Targeting_Human | GGAACGACTATGACCGCCA |
| 334 | Non_Targeting_Human | CCTGGCGAACCCGACGTCT |
| 335 | Non_Targeting_Human | ACAGAGGAAGGTGCCGTAT |
| 336 | Non_Targeting_Human | TATTCGTACCGGGCAGCAG |
| 337 | Non_Targeting_Human | GCCAGATCCCGCCCAGCAG |
| 338 | Non_Targeting_Human | GTGATGGGGCACGACATCG |
| 339 | Non_Targeting_Human | GGAGCCGGAAGGCCGATGG |
| 340 | Non_Targeting_Human | CCCGCGCTGCGGGACGCAG |
| 341 | Non_Targeting_Human | GCTAAAAAACCCCTTCCCC |
| 342 | Non_Targeting_Human | CGAATGAGGGGGACGGACT |
| 343 | Non_Targeting_Human | CAGCAGGCGCGATGACCCT |

| | | |
|---|---|---|
| 344 | Non_Targeting_Human | ACAGGATCAGTGAGCTAAA |
| 345 | Non_Targeting_Human | ATCGCGCGGCACGCGTCAG |
| 346 | Non_Targeting_Human | TTGGTCGGTCACCACTGGG |
| 347 | Non_Targeting_Human | GCTTCTCCAGTCAACGGTT |
| 348 | Non_Targeting_Human | GGCGCATGACAGCCAGGCG |
| 349 | Non_Targeting_Human | AGTCGGTGAACGAGGCCGG |
| 350 | Non_Targeting_Human | TCGGGAGCATACCAAGTGT |
| 351 | Non_Targeting_Human | GCGAGGCGGGAATTCACCA |
| 352 | Non_Targeting_Human | GGCAGACTGATGAAGACGT |
| 353 | Non_Targeting_Human | GGTAGGCCCCGAGTCGCCC |
| 354 | Non_Targeting_Human | GTACTCGCCGGAACGCGAC |
| 355 | Non_Targeting_Human | TATGAGAAATACGCGCGGC |
| 356 | Non_Targeting_Human | GGATCAGGAGGGACGTCGC |
| 357 | Non_Targeting_Human | TCGAGGACGCACCCCATAA |
| 358 | Non_Targeting_Human | ACGGTCGTTGAATCGGCTA |
| 359 | Non_Targeting_Human | CCTACCTCGATGAAGCGTG |
| 360 | Non_Targeting_Human | GTACCAGCGGGGTTTTAGA |
| 361 | Non_Targeting_Human | ACCGCGGGCGTAAGCCCCC |
| 362 | Non_Targeting_Human | GGAAGAGCGAGATCGGGTT |
| 363 | Non_Targeting_Human | CTTCGGCCTCCTAGTGCAG |
| 364 | Non_Targeting_Human | TGAATGAGTGCCCACCCGG |
| 365 | Non_Targeting_Human | CTTAGACCCTCCTGGACTC |
| 366 | Non_Targeting_Human | GGCGTAACGCGGTGGCACA |
| 367 | Non_Targeting_Human | GCGAAGTGCAACCGATGGG |
| 368 | Non_Targeting_Human | GCGGTGCGGGGGATCGAAG |
| 369 | Non_Targeting_Human | CGCTGGTAAAGGCGGAAAG |
| 370 | Non_Targeting_Human | CGAACGTTCAGGATTCCAC |
| 371 | Non_Targeting_Human | TCGAAGTACGGTGTCCCTA |
| 372 | Non_Targeting_Human | TAAACCACGGGCAACGTCT |
| 373 | Non_Targeting_Human | GTAGTGCGCTTCCCCGAAG |
| 374 | Non_Targeting_Human | GGGCAGACCAGGCTCGAGC |
| 375 | Non_Targeting_Human | TCCCCGCGACGGGAATGAA |
| 376 | Non_Targeting_Human | GAATCCATTCGGGCTGCGA |
| 377 | Non_Targeting_Human | AACGACTTGCGTTCATGCC |
| 378 | Non_Targeting_Human | AATATCAAAGGTCTTACGG |
| 379 | Non_Targeting_Human | GGGGTGGGCGCTCTCTCCA |
| 380 | Non_Targeting_Human | CGCGATCAGTAGGACAGTT |
| 381 | Non_Targeting_Human | CGGTGGGGCCGCTCACTAG |
| 382 | Non_Targeting_Human | AGCTGGCACCTCTTAGCGA |
| 383 | Non_Targeting_Human | GTCCCGCATGGGACGGTGG |
| 384 | Non_Targeting_Human | CGACACACGCGACTTCAGA |
| 385 | Non_Targeting_Human | CCCGCATTCAGCACCAGGC |
| 386 | Non_Targeting_Human | GAATATCGCCTGTCGACCA |

| | | |
|---|---|---|
| 387 | Non_Targeting_Human | GACTAGGGAGGCGACACGG |
| 388 | Non_Targeting_Human | GCTTGTTATACACGTCAGT |
| 389 | Non_Targeting_Human | GGCGAGCTGCCATCGTCGA |
| 390 | Non_Targeting_Human | CGCACTATACTAAAGGCGT |
| 391 | Non_Targeting_Human | CCGAGAAGCAGGCTCCAAC |
| 392 | Non_Targeting_Human | AGGGGTGGCTAAGTCCGGC |
| 393 | Non_Targeting_Human | CCCCTCGATTCCCGAACAG |
| 394 | Non_Targeting_Human | CGACCCAACTCTGTGCGTC |
| 395 | Non_Targeting_Human | TTACGTCGATCGATAAAAT |
| 396 | Non_Targeting_Human | GTGAGGCGCGGTACATGGA |
| 397 | Non_Targeting_Human | TGAACGCATGCGGGAGCGT |
| 398 | Non_Targeting_Human | GAGCCGACGTGAGAACGAT |
| 399 | Non_Targeting_Human | GCCATGAGACGTTATGCAG |
| 400 | Non_Targeting_Human | CACATGAAAGTCCAAAAAT |
| 401 | Non_Targeting_Human | GCGCCGGACTGGACCTCGA |
| 402 | Non_Targeting_Human | GAGACAACAGCCGGCTCCC |
| 403 | Non_Targeting_Human | ATGTGATCGAGGGGCACTC |
| 404 | Non_Targeting_Human | TTGCGCAGTGGCGAAATGG |
| 405 | Non_Targeting_Human | TAGGGTTGCGAGGGGTGCC |
| 406 | Non_Targeting_Human | GCGGGCAGCAGGCGCGAGA |
| 407 | Non_Targeting_Human | GGCGGCGCACCGAGTCGAC |
| 408 | Non_Targeting_Human | GCTTGACCGGCTTCCCAGG |
| 409 | Non_Targeting_Human | AGTAAGATTGCACGATGCG |
| 410 | Non_Targeting_Human | GGGGATCGGAGTCCCCCGG |
| 411 | Non_Targeting_Human | CCACGACGTCGGTACTCGA |
| 412 | Non_Targeting_Human | GTGATCATCGCACTGGCAG |
| 413 | Non_Targeting_Human | CAGCGAACACGGACGTCTG |
| 414 | Non_Targeting_Human | TCGCCTCGGAGCAAACCGT |
| 415 | Non_Targeting_Human | ACCCCCGCCGAGAGTTGGT |
| 416 | Non_Targeting_Human | CAGTATTTCCCTTCAAGCG |
| 417 | Non_Targeting_Human | AACCTAAAGGGTATAGTTT |
| 418 | Non_Targeting_Human | CCCGTTTAGGTCATAACCG |
| 419 | Non_Targeting_Human | AACTGCTGGCGTCTGACGC |
| 420 | Non_Targeting_Human | TCCGTGCCCCCGGAGGAAT |
| 421 | Non_Targeting_Human | ACTTAGCCTATTTCTTGCG |
| 422 | Non_Targeting_Human | CAGCAACTGCACTCGCATG |
| 423 | Non_Targeting_Human | CAGCGCCCGCAATGGGCTA |
| 424 | Non_Targeting_Human | TCTCGTTGCATGCTTCGCG |
| 425 | Non_Targeting_Human | GACCGGCCAGCGAAACCAT |
| 426 | Non_Targeting_Human | CACGGAGCGGGGTAAAGAG |
| 427 | Non_Targeting_Human | TTGGCTCCTGCTTCCCCAT |
| 428 | Non_Targeting_Human | CCCGGAACGCTGGTACCCC |
| 429 | Non_Targeting_Human | TGGGGCGAATCAGGTGTAG |

| | | |
|---|---|---|
| 430 | Non_Targeting_Human | TTCGGGAAGCGGGGACGCT |
| 431 | Non_Targeting_Human | ATCATCCGTTTAAACAAAT |
| 432 | Non_Targeting_Human | TCGCAAGTGCATTCCCTGG |
| 433 | Non_Targeting_Human | AGGGTACGTGCTCACCCCC |
| 434 | Non_Targeting_Human | GCTAATTCCGAGGTCGTAT |
| 435 | Non_Targeting_Human | GTTATGCCGTTACTATTAA |
| 436 | Non_Targeting_Human | TGTCCTCACTGGCCGAACG |
| 437 | Non_Targeting_Human | GTGCCGTTGCTGAACTAAG |
| 438 | Non_Targeting_Human | CTGAACTTCGGTAAAGGCG |
| 439 | Non_Targeting_Human | GAGCCAATCTATGGTCGAC |
| 440 | Non_Targeting_Human | GAAGGACCCGGCGCCACTT |
| 441 | Non_Targeting_Human | GCCGTGACAATCACAACTA |
| 442 | Non_Targeting_Human | CGGGCCCGACTCAGTAGCC |
| 443 | Non_Targeting_Human | CCGGACGTCGCTCTTGGTG |
| 444 | Non_Targeting_Human | GTTGGCGGTCAGCGGTGAT |
| 445 | Non_Targeting_Human | CACGGCTCATCGAACTTGG |
| 446 | Non_Targeting_Human | GGCGAAAAGCGAGCCTAGG |
| 447 | Non_Targeting_Human | AAGAGTCGCCTAATTCCGT |
| 448 | Non_Targeting_Human | GGATGGGCGCGCGTGGGGA |
| 449 | Non_Targeting_Human | AGGTCGCGCGCCGGCGTAG |
| 450 | Non_Targeting_Human | CACGCCATGTTGCGAGAAC |
| 451 | Non_Targeting_Human | TCTGGGGCCACGAAACGTT |
| 452 | Non_Targeting_Human | AGGCAGAGCCATGTGGCGG |
| 453 | Non_Targeting_Human | CTCACCGTAGAGACGGCAC |
| 454 | Non_Targeting_Human | CCGCATTGGGAGATGAAAG |
| 455 | Non_Targeting_Human | TAGGACTGAAAACGGATCG |
| 456 | Non_Targeting_Human | CGTTTGACTCACCGAATTG |
| 457 | Non_Targeting_Human | TACGGCGTGCGCCGGCACG |
| 458 | Non_Targeting_Human | TAACATATCAGACGGTCCA |
| 459 | Non_Targeting_Human | ACGGCTCCGTGTGTACCGC |
| 460 | Non_Targeting_Human | TCTGCGGACGCCGTACTAG |
| 461 | Non_Targeting_Human | TGGAGACGCGACTACTACA |
| 462 | Non_Targeting_Human | TCTGTGTATAACTCGGAGG |
| 463 | Non_Targeting_Human | ACATTATTAAGGCCAGCGG |
| 464 | Non_Targeting_Human | GGAAGTTCGGCCCGCACTA |
| 465 | Non_Targeting_Human | AGCCCTGACCCCCACCGGG |
| 466 | Non_Targeting_Human | TGCAGTCAAAGGGCATAAG |
| 467 | Non_Targeting_Human | TCCGACCGCGCTGAAGCAA |
| 468 | Non_Targeting_Human | ACGTCAGTATAGACTGCAA |
| 469 | Non_Targeting_Human | CACGCGCTTAAGTTAATAT |
| 470 | Non_Targeting_Human | GGCCCACGGATTTACGGTG |
| 471 | Non_Targeting_Human | AGGGGCGCACATCGATGTG |
| 472 | Non_Targeting_Human | GGCCCGGGCCTAACCCCCG |

| | | |
|---|---|---|
| 473 | Non_Targeting_Human | GACACGGCGTCGTCATTGA |
| 474 | Non_Targeting_Human | GGTCTAGCTATGAAATTAT |
| 475 | Non_Targeting_Human | GATGGTGCCGGGAACCCGT |
| 476 | Non_Targeting_Human | CGATGACGGGTCCAGACGC |
| 477 | Non_Targeting_Human | CGCGTCAACACGCCATTTA |
| 478 | Non_Targeting_Human | CTTGGCCCCGGGTGACTGG |
| 479 | Non_Targeting_Human | AGGATGGCACCAGATGGGG |
| 480 | Non_Targeting_Human | ACGTGTCTGGCCAGCTAGG |
| 481 | Non_Targeting_Human | GCGACTCCTCACCTGTACC |
| 482 | Non_Targeting_Human | TAGCGCGTTGCGTTCTGAC |
| 483 | Non_Targeting_Human | GGCTATCCAAGGTGCGCAC |
| 484 | Non_Targeting_Human | TGGCGCCCGTAAGAACAAG |
| 485 | Non_Targeting_Human | CGCAGACTCGATTGCCCAC |
| 486 | Non_Targeting_Human | GGTTGCGGGGGGCTCTTTT |
| 487 | Non_Targeting_Human | CCCGACGTAGCACCAATGG |
| 488 | Non_Targeting_Human | AGTGCACCCATACCGAACG |
| 489 | Non_Targeting_Human | GTGCTTCCGGTGAGCCGAC |
| 490 | Non_Targeting_Human | GGGTCGTGGACACACGGTG |
| 491 | Non_Targeting_Human | CCCAAAGTGAGGACGCATT |
| 492 | Non_Targeting_Human | CTTGCGACACACTCTCCAG |
| 493 | Non_Targeting_Human | CTACCGGCGGTCCGCCTCG |
| 494 | Non_Targeting_Human | AGGTGGCGGACGCCCGCAT |
| 495 | Non_Targeting_Human | CGCTGACCCCTAGAACGAG |
| 496 | Non_Targeting_Human | AATCTCCCCGAGAAGAGTC |
| 497 | Non_Targeting_Human | GAATCCCGGGCGCGTGAAA |
| 498 | Non_Targeting_Human | AGAGGGCCCGACTAGGCCG |
| 499 | Non_Targeting_Human | CCGCCAACGAGACATACCT |
| 500 | Non_Targeting_Human | CTACAGTGTCGGTACTCAG |
| 501 | Non_Targeting_Human | GTCGGTGGGGAACACACCG |
| 502 | Non_Targeting_Human | TAAGCTGGACGCAAATGCC |
| 503 | Non_Targeting_Human | CGGCCCGCGAAGCGGGGGT |
| 504 | Non_Targeting_Human | ACAGCACGCAGACACCTAC |
| 505 | Non_Targeting_Human | GGCACTAATGTGGGCGATT |
| 506 | Non_Targeting_Human | TAAGGACGACCCGTCACGG |
| 507 | Non_Targeting_Human | GGGCATAGGCTTGACCGAG |
| 508 | Non_Targeting_Human | GACGGCCCAGAGTACAATA |
| 509 | Non_Targeting_Human | AGGGTACGCGCAAGAGTAG |
| 510 | Non_Targeting_Human | TCCTAGAGGCTCTATAGCC |
| 511 | Non_Targeting_Human | ATTGCGCAAGCCAGGTTAC |
| 512 | Non_Targeting_Human | CGCGTCTTGATGAATGTTG |
| 513 | Non_Targeting_Human | TTTCGATTCGCATGGGGTC |
| 514 | Non_Targeting_Human | TCACGCGAGTTCAACAACG |
| 515 | Non_Targeting_Human | TCGAACCGCGGCCGTACCG |

| | | |
|---|---|---|
| 516 | Non_Targeting_Human | TCAATTCAAACCCACTCGG |
| 517 | Non_Targeting_Human | CAAGAGCTATTCGCTTCGG |
| 518 | Non_Targeting_Human | CTTATCGGGGGCAAAAACG |
| 519 | Non_Targeting_Human | CCGCGTTACGCCGCTACGG |
| 520 | Non_Targeting_Human | TGGGAGCAGATCAAGACTC |
| 521 | Non_Targeting_Human | CATGGGAGGACACCCTCGC |
| 522 | Non_Targeting_Human | ACATCCGCAGAACCGGACT |
| 523 | Non_Targeting_Human | GGCCAGGCAAGCGTTTCCG |
| 524 | Non_Targeting_Human | TTACTACAGGAGCCGAAGG |
| 525 | Non_Targeting_Human | AGGCGGCGCTAACCTCGGG |
| 526 | Non_Targeting_Human | CCAGAAGATCCAAAGGCCC |
| 527 | Non_Targeting_Human | CGGGTGAGGTCCCACCAGA |
| 528 | Non_Targeting_Human | GCTAGTGCACAACATTTGC |
| 529 | Non_Targeting_Human | ACGGGAACCAAGAAGCGGC |
| 530 | Non_Targeting_Human | TGCCCCGCGCATACGACAT |
| 531 | Non_Targeting_Human | AGTCAGAGATGTGTTCCTG |
| 532 | Non_Targeting_Human | TGGAACCGCGACAGGCCCG |
| 533 | Non_Targeting_Human | CGTGAGGAATCTTAAGAGT |
| 534 | Non_Targeting_Human | GTCCCAGGAGCCGCCGTTA |
| 535 | Non_Targeting_Human | TAGATCTCAGCGCGAGTCA |
| 536 | Non_Targeting_Human | CCCGGACAGGAAAATCGCA |
| 537 | Non_Targeting_Human | CAGACCAGTGCGTGCCGAG |
| 538 | Non_Targeting_Human | GGACTAAGGGCGCCTAAGA |
| 539 | Non_Targeting_Human | CGCGAGCCGGCCCTCGTCT |
| 540 | Non_Targeting_Human | AGAGGCGAGTCGTCCTGAT |
| 541 | Non_Targeting_Human | CGGATGGGCCCGATAGCTG |
| 542 | Non_Targeting_Human | CCCCGGCGGACAAAACCGG |
| 543 | Non_Targeting_Human | GACGTAGCACCTCGGCGGT |
| 544 | Non_Targeting_Human | GCAGCCGCGAGTCGTCCCA |
| 545 | Non_Targeting_Human | GGGCACGCGCGCGTCTGAT |
| 546 | Non_Targeting_Human | TTAATCAATCATCGCGTAT |
| 547 | Non_Targeting_Human | GTGGCCGAGGACTAGAAAG |
| 548 | Non_Targeting_Human | CCATGGAGACAGCGCCGGT |
| 549 | Non_Targeting_Human | GGTCGACGAAGATCCGATG |
| 550 | Non_Targeting_Human | CGAGGTGGCGGGGGACGGT |
| 551 | Non_Targeting_Human | TCCTTCCCGGCGGCCGTCT |
| 552 | Non_Targeting_Human | GCCGATCTAGGGTGTGTAG |
| 553 | Non_Targeting_Human | GTTGGACGCGCCCCCCTCG |
| 554 | Non_Targeting_Human | CGTGCGCGCGGATGGGCCG |
| 555 | Non_Targeting_Human | CCGATGGTCTTGTACTACA |
| 556 | Non_Targeting_Human | CTCCGGCGACGACAGGCGC |
| 557 | Non_Targeting_Human | CCGGCAGTTGGGCAGTTTG |
| 558 | Non_Targeting_Human | CCTGGTGCAGTAGACAGCG |

| | | |
|---|---|---|
| 559 | Non_Targeting_Human | CGTCAGAGATCGGGGGAG |
| 560 | Non_Targeting_Human | GGCACATGAGCCTACCATG |
| 561 | Non_Targeting_Human | GTACCGGTGGCGATAAGGA |
| 562 | Non_Targeting_Human | ACGCCGCATGGCCGATTGT |
| 563 | Non_Targeting_Human | ACCGCTGCCCTAACCACTG |
| 564 | Non_Targeting_Human | GTATCGTGCTGGCCGTGGA |
| 565 | Non_Targeting_Human | TAGTCCCGCGGGTCCGAAC |
| 566 | Non_Targeting_Human | GTCGGGCGCTCCATCACCG |
| 567 | Non_Targeting_Human | GAGGGGCGGACGACGTCCG |
| 568 | Non_Targeting_Human | TCGGGCGCCAGTCAGCGAG |
| 569 | Non_Targeting_Human | TGCGGGGCAGTGGGCACGC |
| 570 | Non_Targeting_Human | TGTGGTCGCTTGCAAGTCC |
| 571 | Non_Targeting_Human | GATGAAAGCGGGGGTCGGT |
| 572 | Non_Targeting_Human | GCGCACCCTCGGAAGCGAG |
| 573 | Non_Targeting_Human | ATAGCCAGCCGCCCAAAAT |
| 574 | Non_Targeting_Human | CCGGCCGCTGCCGTCGTCT |
| 575 | Non_Targeting_Human | AGCCGGGGCTCGATAGAGA |
| 576 | Non_Targeting_Human | GCGACACGTATCCCGTAAG |
| 577 | Non_Targeting_Human | GGACAAGCGCGCATGTAAC |
| 578 | Non_Targeting_Human | ATACACACGTATAGCCCCG |
| 579 | Non_Targeting_Human | CTCCGCGCACTGATAAGGA |
| 580 | Non_Targeting_Human | GAGCGGACTATCAACTCAG |
| 581 | Non_Targeting_Human | GCCTATCCACCCCCAGCCG |
| 582 | Non_Targeting_Human | TTCGCAGTTAGGGAAGCGA |
| 583 | Non_Targeting_Human | CGGAGGTCGCTCTGAGCGT |
| 584 | Non_Targeting_Human | TCACGGTGGGGTGGTGCTT |
| 585 | Non_Targeting_Human | AAACATAGGTATCAAGAGG |
| 586 | Non_Targeting_Human | TACGCGAATATAGCCGGGT |
| 587 | Non_Targeting_Human | AGCGGCCCGCCTTAAATAG |
| 588 | Non_Targeting_Human | TCCCCGCAAGCAGCTAGGG |
| 589 | Non_Targeting_Human | CCGGTGGTCCTATCCAGTC |
| 590 | Non_Targeting_Human | CTAGCGGCTAGAGCTACAG |
| 591 | Non_Targeting_Human | GCGGCTGCGGTAAGGTTCG |
| 592 | Non_Targeting_Human | CGGGCGCCACCGAAACCAG |
| 593 | Non_Targeting_Human | CACGGACCCTATTCTAATA |
| 594 | Non_Targeting_Human | GTCCGCTACCCAAAAACAT |
| 595 | Non_Targeting_Human | TATAAGAGATCCTTAATAC |
| 596 | Non_Targeting_Human | GCCGTTAGAGGGACCTTGT |
| 597 | Non_Targeting_Human | CACTCCTCCGTTGCGCAGG |
| 598 | Non_Targeting_Human | CGGTACGCAATCGGTCCAG |
| 599 | Non_Targeting_Human | TCAACGGAAGTCCGAGGCG |
| 600 | Non_Targeting_Human | AGCGGGCGACACAGATGGT |
| 601 | Non_Targeting_Human | CAACGTAGATCGCCTCCTG |

| | | |
|---|---|---|
| 602 | Non_Targeting_Human | ACCAGGGTATGGCAGCTAG |
| 603 | Non_Targeting_Human | GAGCCAAAGCACGAGCGGG |
| 604 | Non_Targeting_Human | GACGGCGGGGAGGATAGAT |
| 605 | Non_Targeting_Human | TCAGACCACGTGCGACGCT |
| 606 | Non_Targeting_Human | GGCGGACTTACTAGTTCCT |
| 607 | Non_Targeting_Human | GACGTAGCGACAGGGGCAT |
| 608 | Non_Targeting_Human | GGATGGCAGCGGCCCGACA |
| 609 | Non_Targeting_Human | AACCGCGCGAGTAGGGGGG |
| 610 | Non_Targeting_Human | GTCTTAGAAGGACCGTTCG |
| 611 | Non_Targeting_Human | TAGTGAGCCGCCTGCCAAC |
| 612 | Non_Targeting_Human | GGCTGTCCAGGTAAAAATA |
| 613 | Non_Targeting_Human | CTAACGTCTGAGTACACGG |
| 614 | Non_Targeting_Human | GGTGGCCCGACCCACCCAT |
| 615 | Non_Targeting_Human | AGCGGCGCAAGGGTTAGCT |
| 616 | Non_Targeting_Human | GGGAGGTTCAAGGTTAGGA |
| 617 | Non_Targeting_Human | AACATGGCCTACCAGAGCA |
| 618 | Non_Targeting_Human | CGCACCAGAGCAGTTGAAA |
| 619 | Non_Targeting_Human | GGGGCCGCCAACAAATCTC |
| 620 | Non_Targeting_Human | GTTGGAGGGCTTCATGCGG |
| 621 | Non_Targeting_Human | GTGTTTAAGAGCGAACCGC |
| 622 | Non_Targeting_Human | ACGGCGTCCGCGCGGTTTT |
| 623 | Non_Targeting_Human | AGAGCGTGATGAATGGGCC |
| 624 | Non_Targeting_Human | GCCGTTCTCCCAAACGGCC |
| 625 | Non_Targeting_Human | CCTAGGCTTGTATGAGAGG |
| 626 | Non_Targeting_Human | AAAGCGCCTTGACAATCAG |
| 627 | Non_Targeting_Human | CCGTCAAATGCACGCAGGG |
| 628 | Non_Targeting_Human | TGCTGACACCGGCTATGCG |
| 629 | Non_Targeting_Human | GCACGGCGCCCCAGTAAAT |
| 630 | Non_Targeting_Human | CTCACAGCATGATGGGTAA |
| 631 | Non_Targeting_Human | CGCGCCCCTGACTAGAGCG |
| 632 | Non_Targeting_Human | CCCTGGGTCCAACCGGTCC |
| 633 | Non_Targeting_Human | GAGTTAAGGCCTCGTCTAG |
| 634 | Non_Targeting_Human | ACGTGCACGTGTGCCGTAA |
| 635 | Non_Targeting_Human | GTCCGACGCGACACAGTTA |
| 636 | Non_Targeting_Human | TAGGTGTCCAGATCACCCT |
| 637 | Non_Targeting_Human | GCCATCCTCATCACGGGTG |
| 638 | Non_Targeting_Human | CTATCCGACGACACGACGG |
| 639 | Non_Targeting_Human | AGCACGAGAGGTAACGGTA |
| 640 | Non_Targeting_Human | GTCGGTTCGTGTGAATCAC |
| 641 | Non_Targeting_Human | GGTCACTTGAAGGTGCGCA |
| 642 | Non_Targeting_Human | ACGCAGAGACGCGGAACTG |
| 643 | Non_Targeting_Human | GTCTGCTCAAATCTAGGTG |
| 644 | Non_Targeting_Human | GCGTCTGCACGCTTGCCCA |

| | | |
|---|---|---|
| 645 | Non_Targeting_Human | GACTAAACAGACATCGACT |
| 646 | Non_Targeting_Human | TCAATCTGGGCGCTCTAGT |
| 647 | Non_Targeting_Human | ACCATCCTTGTAATCGAAG |
| 648 | Non_Targeting_Human | GATACCGCCCAGTTCTATT |
| 649 | Non_Targeting_Human | GATCAGACGGCCGGCGAAT |
| 650 | Non_Targeting_Human | GGCGACTCACACCTCCCGA |
| 651 | Non_Targeting_Human | CAACAGCACGCTAAACAAA |
| 652 | Non_Targeting_Human | TATCTTGGCAGCTCCCAGA |
| 653 | Non_Targeting_Human | GACCGGCAAGGATCCCGGG |
| 654 | Non_Targeting_Human | TGCTGGCCCCGGCCGTGCT |
| 655 | Non_Targeting_Human | AGGACCCCCCCACGCTCAG |
| 656 | Non_Targeting_Human | GTAGAACGGAACGCGGACG |
| 657 | Non_Targeting_Human | TCGCCTGTCGTGCAAAGAG |
| 658 | Non_Targeting_Human | ACAGACACCCACTCAACGG |
| 659 | Non_Targeting_Human | CCGGCTACCGGCCGTGAAC |
| 660 | Non_Targeting_Human | GAACCTGATGGGCACGCGA |
| 661 | Non_Targeting_Human | CCCCGCTGGGATTGCCGAC |
| 662 | Non_Targeting_Human | GTCGTCAGGAAACAGACGT |
| 663 | Non_Targeting_Human | CCGGCCCGGGGCGACTCCA |
| 664 | Non_Targeting_Human | AGGGGCCCTGGTCAACGAT |
| 665 | Non_Targeting_Human | ACTCCATCGCACAGTACAA |
| 666 | Non_Targeting_Human | ACCAGGCGGAGGAAGTTAT |
| 667 | Non_Targeting_Human | CTGGGTGAACAGTTCCCAG |
| 668 | Non_Targeting_Human | CGCCGGAATAGCACATCAA |
| 669 | Non_Targeting_Human | GGAAAATAGAAAGGCCGAG |
| 670 | Non_Targeting_Human | TTCATAGAAGGAATTGCGG |
| 671 | Non_Targeting_Human | GGCTGTGTGACGTGGCCGA |
| 672 | Non_Targeting_Human | CGGACGTATCACCCACTTC |
| 673 | Non_Targeting_Human | CGGAAGGCTAGGTGCAACT |
| 674 | Non_Targeting_Human | ACTTTTCCTCGTATACTGT |
| 675 | Non_Targeting_Human | ACAGTCACCCGGACCTTGC |
| 676 | Non_Targeting_Human | GCTGTAACTCCCTAGATGG |
| 677 | Non_Targeting_Human | GAGCCCCTCGACAGACCAA |
| 678 | Non_Targeting_Human | TGCCGATAGGGTGTAGCCA |
| 679 | Non_Targeting_Human | TATCTACTCCCCGGCGGCC |
| 680 | Non_Targeting_Human | TACGGCGCTGCCACGTATA |
| 681 | Non_Targeting_Human | CCATTGAGGTTTGAGACAG |
| 682 | Non_Targeting_Human | ATCCGCAGATGGAGGGGAT |
| 683 | Non_Targeting_Human | ATAGCATGGATGCGTGGTG |
| 684 | Non_Targeting_Human | ATGGGGGCACTCGACACCG |
| 685 | Non_Targeting_Human | GCTCGGCGATCGGCCCCAG |
| 686 | Non_Targeting_Human | GCAGACCGTCCCTGATCAA |
| 687 | Non_Targeting_Human | GAGGAACCCAATTCAAAAG |

| | | |
|---|---|---|
| 688 | Non_Targeting_Human | ACGTCAAGGCCTGTCAGAG |
| 689 | Non_Targeting_Human | CACGTCCGCCGGATCAAAA |
| 690 | Non_Targeting_Human | CCCAGGCAACGGCCAGAGT |
| 691 | Non_Targeting_Human | CTATCGGTGCCAAAAACAA |
| 692 | Non_Targeting_Human | CGGCAGGAACGAACCGGAT |
| 693 | Non_Targeting_Human | TGCAGGTCAGGAACAATAG |
| 694 | Non_Targeting_Human | CGATGGCACAGTGCTCCTT |
| 695 | Non_Targeting_Human | CGGCCCGGCCTGTACCACC |
| 696 | Non_Targeting_Human | GAACCGCGGCATCTGCCGG |
| 697 | Non_Targeting_Human | ACGGCTGTCCTGTCAACAG |
| 698 | Non_Targeting_Human | ATGGGCAAGAATACGCCAG |
| 699 | Non_Targeting_Human | TCAAGAAGTGAACGCGCCG |
| 700 | Non_Targeting_Human | GAGAATCGCTACGAGACGA |
| 701 | Non_Targeting_Human | GCCCTGCACCGTGCGATTG |
| 702 | Non_Targeting_Human | AACCGGCAAGCCTGTCGAA |
| 703 | Non_Targeting_Human | CAATAGGTCGTCGGACGCC |
| 704 | Non_Targeting_Human | TAAAGTTGTCGGCAATTGC |
| 705 | Non_Targeting_Human | TATATGGCCGCCTTACCAC |
| 706 | Non_Targeting_Human | CCTGGTGGGGGACGTTCCA |
| 707 | Non_Targeting_Human | TCCCAGGCTCTCCACTATG |
| 708 | Non_Targeting_Human | GCTTCTGACCGCTGCCTGG |
| 709 | Non_Targeting_Human | CATCCGCCAAACAGCCCGA |
| 710 | Non_Targeting_Human | TGACTTTACTGGCCATCAA |
| 711 | Non_Targeting_Human | CATGAGCGTATCTAACATG |
| 712 | Non_Targeting_Human | ATCGCGGAGGTCAACCCGC |
| 713 | Non_Targeting_Human | ACGGCGAATGCAAGGAGGG |
| 714 | Non_Targeting_Human | TCGGAGCATCCCCACTCCG |
| 715 | Non_Targeting_Human | ACAGTTCCCCACGACGAAG |
| 716 | Non_Targeting_Human | TTGATCAAGCGCAGCGTTA |
| 717 | Non_Targeting_Human | GCGGCTGCGACGACACCAC |
| 718 | Non_Targeting_Human | GAGGGCGAGAGTTTACTTG |
| 719 | Non_Targeting_Human | ACCTACTGCGCCAGTGGGG |
| 720 | Non_Targeting_Human | GGAGCGCGGGCGTCAGTCA |
| 721 | Non_Targeting_Human | CCGTTGTCCAACCCATCAA |
| 722 | Non_Targeting_Human | CCGTTCACAGTCGGGGGCA |
| 723 | Non_Targeting_Human | TGGGGTTGCCGTCACAGAG |
| 724 | Non_Targeting_Human | GGACGCCGGGGCAAAAGCG |
| 725 | Non_Targeting_Human | AGCAGCCCTCATAGTGCCG |
| 726 | Non_Targeting_Human | GCTCGCCGTCAGGCGTGGT |
| 727 | Non_Targeting_Human | CGGGTACAGGGACTTACTC |
| 728 | Non_Targeting_Human | TAACGCTTGCGTGGGACGG |
| 729 | Non_Targeting_Human | GGGTCGAATCCACACGGTA |
| 730 | Non_Targeting_Human | TCGGATGAACCGCTAGAAC |

| | | |
|---|---|---|
| 731 | Non_Targeting_Human | GGGAGGCGTCTGAAACTTG |
| 732 | Non_Targeting_Human | ATCGACGCTGATTAGGGCG |
| 733 | Non_Targeting_Human | AGTGGAGTCAACGACATAG |
| 734 | Non_Targeting_Human | GCTGCGTGCGCCGCGCACG |
| 735 | Non_Targeting_Human | GGGCGCACGTTGGACGTGA |
| 736 | Non_Targeting_Human | CTGTCCACGACTGTTTCAA |
| 737 | Non_Targeting_Human | TGCGGGGGCATGGCCCCGC |
| 738 | Non_Targeting_Human | TGTGCCACTGTGGGACCGG |
| 739 | Non_Targeting_Human | TATTCGGGTAGCGGGTCGC |
| 740 | Non_Targeting_Human | TTGTACGGCTGGGTATCGG |
| 741 | Non_Targeting_Human | AGCCCCCGAACATGAAGAA |
| 742 | Non_Targeting_Human | AACCATGCGTGGAAAGTGT |
| 743 | Non_Targeting_Human | TGATTGGGACAGATGCATT |
| 744 | Non_Targeting_Human | TGTCGTAAAGACACCGGGG |
| 745 | Non_Targeting_Human | CGCCTCGCGAGCGATGTAA |
| 746 | Non_Targeting_Human | ATGAGGGGTCGGCCAACGA |
| 747 | Non_Targeting_Human | CCCGCGCGACGCTAGCAAA |
| 748 | Non_Targeting_Human | AAGGCGCTAGTCAAAAGGA |
| 749 | Non_Targeting_Human | GTGGGAGAAGCACCGTATG |
| 750 | Non_Targeting_Human | TAGGACGGGGTACTAAGTG |
| 751 | Non_Targeting_Human | ACCGCGGGGGGCAATCGGA |
| 752 | Non_Targeting_Human | GCGCGGGCCCCATAAAAAC |
| 753 | Non_Targeting_Human | AGGTGCTTAGATCGCCTCG |
| 754 | Non_Targeting_Human | GGCCCCTAGCGCTCAACAT |
| 755 | Non_Targeting_Human | GGTGATGCGGACAGGCCCG |
| 756 | Non_Targeting_Human | TAGTTGTGTAACACCTACG |
| 757 | Non_Targeting_Human | CGAACTCGCTGAGAAACTA |
| 758 | Non_Targeting_Human | AGGCATCCCCTGATAAGAA |
| 759 | Non_Targeting_Human | GAAAGTCCGTTCCCACCGA |
| 760 | Non_Targeting_Human | TGCCTCAGGCCCACAACGG |
| 761 | Non_Targeting_Human | CTACCGATACTCGGGAGGA |
| 762 | Non_Targeting_Human | CTCTTGCTTCCTTTATCTA |
| 763 | Non_Targeting_Human | GCGTGTGTCGGGCGAGCCT |
| 764 | Non_Targeting_Human | GCTGTATATCGGCGCCCCG |
| 765 | Non_Targeting_Human | AGGCCTTCACCTCGTGCAA |
| 766 | Non_Targeting_Human | GCTCGTTGGACAAACGTAG |
| 767 | Non_Targeting_Human | GTAGACGCGGTGAACGTGG |
| 768 | Non_Targeting_Human | TTGTGGCCGCGTGCGGTCG |
| 769 | Non_Targeting_Human | GGAGGCGCATACGGGTCGG |
| 770 | Non_Targeting_Human | GGGGTCAGTTGACGAGTAC |
| 771 | Non_Targeting_Human | GCGCGAGCACAGATAACTT |
| 772 | Non_Targeting_Human | GGTTACCCGCAAGCGAGCG |
| 773 | Non_Targeting_Human | GTGATAGAGCGTGGCCACA |

| | | |
|---|---|---|
| 774 | Non_Targeting_Human | CCCCCCGATCGCGTGGGAG |
| 775 | Non_Targeting_Human | GTAGTAGGATCCACCCCCG |
| 776 | Non_Targeting_Human | CTTGGTTCGAGGCGAGATT |
| 777 | Non_Targeting_Human | CGCGTGGGCACGCACCCCC |
| 778 | Non_Targeting_Human | CAGCATCTACCAGTAGGAG |
| 779 | Non_Targeting_Human | CTAGTTATGCCACTCTGCC |
| 780 | Non_Targeting_Human | AAGTTAAGAGATCGACTTC |
| 781 | Non_Targeting_Human | CCACCGTGTGAACCGCCCA |
| 782 | Non_Targeting_Human | CGGATGCGCAACCCGGTTG |
| 783 | Non_Targeting_Human | ATGAGTAGCCTCGGAGCGG |
| 784 | Non_Targeting_Human | TACCGCGCCGCGACCACGG |
| 785 | Non_Targeting_Human | GCCCCTCCAGTGGGTGCGA |
| 786 | Non_Targeting_Human | CTTGCCTAGGCGCTACCAC |
| 787 | Non_Targeting_Human | ATATCCCGGTGGGGTTCTC |
| 788 | Non_Targeting_Human | CTGCCCACCCGTAAGGCCC |
| 789 | Non_Targeting_Human | ACATGCACGCATTGGCCTG |
| 790 | Non_Targeting_Human | ACTGGGGAGGCAGTCGCAA |
| 791 | Non_Targeting_Human | CACGACGCTCCGGCACGCG |
| 792 | Non_Targeting_Human | ACCAAACCAAATGAAGGAC |
| 793 | Non_Targeting_Human | ATGGTCCTGGGCAAAGCTA |
| 794 | Non_Targeting_Human | CGGAATAGAAGCATGCCCC |
| 795 | Non_Targeting_Human | CCGGCGAGAAGCTCGTGAG |
| 796 | Non_Targeting_Human | GGACGGGAAAGACTGGTCC |
| 797 | Non_Targeting_Human | GCAGGGTTACTCGTACGGA |
| 798 | Non_Targeting_Human | CTACAACGCTGTTGTACAC |
| 799 | Non_Targeting_Human | ACGCTCAGGCGAATCTGAG |
| 800 | Non_Targeting_Human | CGGAGGTTGAGGGCCAAGC |
| 801 | Non_Targeting_Human | ACACACTGACCGTCATAAC |
| 802 | Non_Targeting_Human | CGGACGTTCGCAACTCACC |
| 803 | Non_Targeting_Human | CACTTCGAATAGTAATAGG |
| 804 | Non_Targeting_Human | GACCCAACGGTAGATGCTG |
| 805 | Non_Targeting_Human | GAGACAATGGGAACCCGAA |
| 806 | Non_Targeting_Human | TGCGCGACATCCACAGGGG |
| 807 | Non_Targeting_Human | CAGAGGTCGGTCAAGGCGC |
| 808 | Non_Targeting_Human | TGCGTGGACGCATCTGCAC |
| 809 | Non_Targeting_Human | CGCGGCGCATGCGAGAACT |
| 810 | Non_Targeting_Human | GCACTGACGGACTCACCGT |
| 811 | Non_Targeting_Human | GAGTGCTGAACCACTGCGT |
| 812 | Non_Targeting_Human | ACGGCTGCCAGGATCTCTA |
| 813 | Non_Targeting_Human | AGGTTTCGAGATAACTACA |
| 814 | Non_Targeting_Human | GGACTGACGGCGGGAAAGA |
| 815 | Non_Targeting_Human | GCGCCCGGTACATCTCCCT |
| 816 | Non_Targeting_Human | GGGATCGATCCGTTGATGG |

| | | |
|---|---|---|
| 817 | Non_Targeting_Human | TCGGTGTCGAAGGACGCCG |
| 818 | Non_Targeting_Human | GGCCATTGATCGACTGCTA |
| 819 | Non_Targeting_Human | TGCACACGGACGGACCCGA |
| 820 | Non_Targeting_Human | AACGGAGCTGTAGAAACTT |
| 821 | Non_Targeting_Human | CGCCCGCTCGGCCGAGCAA |
| 822 | Non_Targeting_Human | CACTGTCAAGTGAAATGCG |
| 823 | Non_Targeting_Human | TCTGGGACGATACCAATTT |
| 824 | Non_Targeting_Human | TGTGCTGCAGCGACGTGAT |
| 825 | Non_Targeting_Human | CCGAGCGTTAGGACGCCGG |
| 826 | Non_Targeting_Human | AGCTTAGATTACAGCCCAC |
| 827 | Non_Targeting_Human | GGAACCCTTCTAACCCCCT |
| 828 | Non_Targeting_Human | GGCGCGAGGTACGTACGTG |
| 829 | Non_Targeting_Human | GAAGGGGACAGCACTGACG |
| 830 | Non_Targeting_Human | GTATGATAACCGCACCTAA |
| 831 | Non_Targeting_Human | TAGGCATCCGGGTCCAGAG |
| 832 | Non_Targeting_Human | AAAGTCGGGAGAATAAACC |
| 833 | Non_Targeting_Human | GAGGGCCAGTCAACTCGTA |
| 834 | Non_Targeting_Human | GGGCCCTATTGGAACACCG |
| 835 | Non_Targeting_Human | CGTGCCGAGCCCCGTTACA |
| 836 | Non_Targeting_Human | CATCCAACGCGCATTTGAC |
| 837 | Non_Targeting_Human | GCGCGCGCACGAAAATCAA |
| 838 | Non_Targeting_Human | CCCCGAGGACCGGCGGCAG |
| 839 | Non_Targeting_Human | GAGCGGAATGCTAGTGCTA |
| 840 | Non_Targeting_Human | AGCTAGGGCAGCGCATGCA |
| 841 | Non_Targeting_Human | CGCAATTCACCCCTAGGTG |
| 842 | Non_Targeting_Human | TCCTCCCCTCTGCGACCTT |
| 843 | Non_Targeting_Human | CGTTGGCGCAGTAATGAGC |
| 844 | Non_Targeting_Human | CGCTAGACGAAGGAAGTTA |
| 845 | Non_Targeting_Human | TAGGGCAGAACGCGACCGA |
| 846 | Non_Targeting_Human | TAGTTCCGCCAGCATTTAG |
| 847 | Non_Targeting_Human | GAGGCAGACTAATGTAAGG |
| 848 | Non_Targeting_Human | CGAGTCTGCAGACTGGGCA |
| 849 | Non_Targeting_Human | GCGCTGACTCCGAGCTCCG |
| 850 | Non_Targeting_Human | TAGGACAACCTTACGTCTT |
| 851 | Non_Targeting_Human | GCCACAATAGGCGGTGCTC |
| 852 | Non_Targeting_Human | GGGGATTCACGGGTCAATG |
| 853 | Non_Targeting_Human | GAACCCGAACAGCACGCGT |
| 854 | Non_Targeting_Human | CGGCCACAGCTCACCGCCC |
| 855 | Non_Targeting_Human | GGTCGATGGAGTTCGCCCG |
| 856 | Non_Targeting_Human | CGGTTAGGGCCACAAAACA |
| 857 | Non_Targeting_Human | GAAACGACAGGGAACCCGT |
| 858 | Non_Targeting_Human | TCGGGCGAATAACTCCGCC |
| 859 | Non_Targeting_Human | GTCTATGGTCCGCACACAG |

| | | |
|---|---|---|
| 860 | Non_Targeting_Human | TGCAATAGCGGAATTACCG |
| 861 | Non_Targeting_Human | GTTGGCCTACCGAGGAGTT |
| 862 | Non_Targeting_Human | GGACGTCATGCGCCCGCTT |
| 863 | Non_Targeting_Human | TAGTTGATCCGCGCAGCTC |
| 864 | Non_Targeting_Human | TCGGACCAAGCAGGCAGAA |
| 865 | Non_Targeting_Human | GCCGCCATCCTCACCCGTG |
| 866 | Non_Targeting_Human | ATTGTCCGATTGTAGTTAC |
| 867 | Non_Targeting_Human | CACAGGTTATTCCGAACGA |
| 868 | Non_Targeting_Human | GCGCTACCGGAAGCTTTTG |
| 869 | Non_Targeting_Human | GTGCAGCGCGACGAAAGGA |
| 870 | Non_Targeting_Human | CAGCACCGGGGCAGAAAGT |
| 871 | Non_Targeting_Human | CCCCTCGTCTTGGAAGCGA |
| 872 | Non_Targeting_Human | GGCTTGGCGCCGCCAGCTT |
| 873 | Non_Targeting_Human | ATGCGACGTACGGGAAGGG |
| 874 | Non_Targeting_Human | TAGGATGCGGGCTATACAC |
| 875 | Non_Targeting_Human | CGTAGAAGGGTATGTAGAA |
| 876 | Non_Targeting_Human | TCTGCCCCTTGATTAGGCG |
| 877 | Non_Targeting_Human | GGCTCCAGTGTAAAACCCG |
| 878 | Non_Targeting_Human | CCCAGAACAGGTGGAGTTC |
| 879 | Non_Targeting_Human | AGCAGAACGGGTCGGATTG |
| 880 | Non_Targeting_Human | AAGCGACTTTACTCTCCGC |
| 881 | Non_Targeting_Human | GTGGAATGATTCAGCTATG |
| 882 | Non_Targeting_Human | ACTGAAGAGGGCCTGGATA |
| 883 | Non_Targeting_Human | GAAGGGATTATACGCCGCC |
| 884 | Non_Targeting_Human | CGGAGTGTCGAGCGTGCTG |
| 885 | Non_Targeting_Human | AGGAGGGGTGTGCCTGACG |
| 886 | Non_Targeting_Human | CAGCATCCATGTGTAACGA |
| 887 | Non_Targeting_Human | ATGAGCGCGAACGCTTAGG |
| 888 | Non_Targeting_Human | CCGGCCGGCGTACCCGCAC |
| 889 | Non_Targeting_Human | GACCCAGAAAGTTCGAGGG |
| 890 | Non_Targeting_Human | TCTACTGCCGCACCGGGGT |
| 891 | Non_Targeting_Human | CGCCAGGGCGAACCTTCCC |
| 892 | Non_Targeting_Human | CACCAAGTGAGCGATCAGA |
| 893 | Non_Targeting_Human | TTTCCCGTGACGGGAGCGA |
| 894 | Non_Targeting_Human | AACACGATCACCACAGGGG |
| 895 | Non_Targeting_Human | GGCCCGAGCCTCATGACGA |
| 896 | Non_Targeting_Human | CAGACCATGCGCCACCCTG |
| 897 | Non_Targeting_Human | AGGAACGCGCAAATTACAC |
| 898 | Non_Targeting_Human | CGACTAGCCGGCACGTGGA |
| 899 | Non_Targeting_Human | ACGCACCGAGGGACATAGG |
| 900 | Non_Targeting_Human | TGACCGGGAGTCCAGACTG |
| 901 | Non_Targeting_Human | TTTCGACGGCACGGACAGA |
| 902 | Non_Targeting_Human | CGGAAACCGGTGCGTGGCA |

| | | |
|---|---|---|
| 903 | Non_Targeting_Human | GCGCCCGGCGTGCATCTCG |
| 904 | Non_Targeting_Human | TTCCTTTTAATGATAATAG |
| 905 | Non_Targeting_Human | AGGTGGGGCCGACTCCGGG |
| 906 | Non_Targeting_Human | AAAAGCATGCTATACGGCG |
| 907 | Non_Targeting_Human | CGCCCCGCCCTCAGGACGG |
| 908 | Non_Targeting_Human | TTTGTAGCGAAGTTAATTC |
| 909 | Non_Targeting_Human | CAGATCACCCGACGGGAAG |
| 910 | Non_Targeting_Human | CACCGAGAACAATTGACCC |
| 911 | Non_Targeting_Human | CACGCGTATCCGTAGGTTG |
| 912 | Non_Targeting_Human | ATGCGTTGCCGGGAGGTGA |
| 913 | Non_Targeting_Human | GAAGAGAAGCCAAACTCAC |
| 914 | Non_Targeting_Human | CATTCTACTAACAGCTTAG |
| 915 | Non_Targeting_Human | CTCTCATCCCTACAGTCAA |
| 916 | Non_Targeting_Human | CAGCTTGCGCAAATTCCAA |
| 917 | Non_Targeting_Human | CCGAGCGGCAGGGTGCCGC |
| 918 | Non_Targeting_Human | AAGTAGAGATGAAGGCCTA |
| 919 | Non_Targeting_Human | GCAGGGGAGGCGGAACTCA |
| 920 | Non_Targeting_Human | ATGCGGCGAACTGCCGTAA |
| 921 | Non_Targeting_Human | CCGTGCGCTGCACTACGCC |
| 922 | Non_Targeting_Human | ATTGAGATACGTTGGACAG |
| 923 | Non_Targeting_Human | AACAGCTCCCGAGAGGTCT |
| 924 | Non_Targeting_Human | TTAGGGCCGGGCCAGATGG |
| 925 | Non_Targeting_Human | AACTCCCGGCCACGTTATG |
| 926 | Non_Targeting_Human | TCCGACGGGTGCCGTGTCT |
| 927 | Non_Targeting_Human | GGGGAAAGTCGGGGAATGA |
| 928 | Non_Targeting_Human | GCCATCAAGAACGTCAACC |
| 929 | Non_Targeting_Human | AGATGGTATGGGCCACGGC |
| 930 | Non_Targeting_Human | TGGACGGTACATCGGCACA |
| 931 | Non_Targeting_Human | GTGCCGACGAAATTCGTCG |
| 932 | Non_Targeting_Human | CGTCGATACCGTCTCTCGC |
| 933 | Non_Targeting_Human | CTAGCCTCGCGCCGCTGGC |
| 934 | Non_Targeting_Human | GTTGGCTGTACTGGGCCAA |
| 935 | Non_Targeting_Human | AAGGGCTCGGAGTGCATGG |
| 936 | Non_Targeting_Human | GACTCCCGAGCGGGCACTC |
| 937 | Non_Targeting_Human | CGGAAGCTAGCGTCGTAGG |
| 938 | Non_Targeting_Human | ATGGCCGCAGCATGTGTAG |
| 939 | Non_Targeting_Human | GACACAGTCAGCGTTGGGC |
| 940 | Non_Targeting_Human | TAACTGCTACGGCGCCCAA |
| 941 | Non_Targeting_Human | AGTGAATATGGGTATAGCG |
| 942 | Non_Targeting_Human | TACCCAACACCCCTCTAGC |
| 943 | Non_Targeting_Human | CGCACCAGAAGACAGCAGG |
| 944 | Non_Targeting_Human | CTGGGCCCGTTTTACCGGG |
| 945 | Non_Targeting_Human | TCACCCGTTAAATATTGAA |

| | | |
|---|---|---|
| 946 | Non_Targeting_Human | AACGAGAACCGCGTTCTAT |
| 947 | Non_Targeting_Human | CCGCGCTCTGCAAGGAAGT |
| 948 | Non_Targeting_Human | TAGGGTCTGCCGTACACGC |
| 949 | Non_Targeting_Human | GACGATATTAAGGAAGCCA |
| 950 | Non_Targeting_Human | GGGATTGGGTGTGACCGCA |
| 951 | Non_Targeting_Human | CAACGGGTTCGGGTGAGAG |
| 952 | Non_Targeting_Human | GAAAGTGCGTTATCACCCG |
| 953 | Non_Targeting_Human | ACAGCGTGTAGTTTACCGA |
| 954 | Non_Targeting_Human | GCCTGTGACGCCCGCGGAC |
| 955 | Non_Targeting_Human | AAGCCTCCGGCGGCCTCCG |
| 956 | Non_Targeting_Human | TGCATGACAGGATCTCTCG |
| 957 | Non_Targeting_Human | GACCGGCGCCGTGACGAAC |
| 958 | Non_Targeting_Human | GACATATTCGGCCCGGCAT |
| 959 | Non_Targeting_Human | TCCGGCGCGGAATGCGAAA |
| 960 | Non_Targeting_Human | GTTGAACCGCCCCCGCCGA |
| 961 | Non_Targeting_Human | CCAGGCGCATAGTAATCGA |
| 962 | Non_Targeting_Human | GGCGTCTCCGAGATCTACT |
| 963 | Non_Targeting_Human | GAGTTGCGGAACGCCGCAG |
| 964 | Non_Targeting_Human | CATTGAGAGATGACCCAGG |
| 965 | Non_Targeting_Human | ACCTCTGGTTTAATAACGG |
| 966 | Non_Targeting_Human | CTCATGTCTAAGGCGGCAG |
| 967 | Non_Targeting_Human | TGACGGAGGTGCGAAGCGT |
| 968 | Non_Targeting_Human | TCACCAGCGCATAAGCGCA |
| 969 | Non_Targeting_Human | CGCGGGGCCGAATATAAGG |
| 970 | Non_Targeting_Human | GCTACCCTTAGCACCGAAA |
| 971 | Non_Targeting_Human | CGCCCCAGGGTTGATGCGA |
| 972 | Non_Targeting_Human | GCCGACGAGAGGCCTCTTT |
| 973 | Non_Targeting_Human | GAAATCACTCTGCGTTCAG |
| 974 | Non_Targeting_Human | GTAGAGGCGGACGGCTGAG |
| 975 | Non_Targeting_Human | ATGGGCTTCGGGATCTCAC |
| 976 | Non_Targeting_Human | CGCACGTGCGCCCTCTAAA |
| 977 | Non_Targeting_Human | CTACGGGGCCTCTGTGGTC |
| 978 | Non_Targeting_Human | ATAGGTGGCAGCGGCCCAT |
| 979 | Non_Targeting_Human | TAGCGCGATGCACTATAGG |
| 980 | Non_Targeting_Human | ATTCCCAATGTCGTTGGCG |
| 981 | Non_Targeting_Human | GTTGTGCCGAGATGTCAAG |
| 982 | Non_Targeting_Human | CACAAGGACGCAAAAGACG |
| 983 | Non_Targeting_Human | CTTGAATGGCTCCACCTAA |
| 984 | Non_Targeting_Human | ACCTTGACCGCCCTGTCGA |
| 985 | Non_Targeting_Human | GTGATCGCGGGCCTCATAC |
| 986 | Non_Targeting_Human | CACTAGGGAGCGCGGCAGG |
| 987 | Non_Targeting_Human | CAATGTCTAGGGCGAGCTG |
| 988 | Non_Targeting_Human | TAGTGGGCGACGGCATGCG |

| | | |
|---|---|---|
| 989 | Non_Targeting_Human | GTCAGGGGCCGGATTTCGC |
| 990 | Non_Targeting_Human | TACGCACCTGGCGGGCGTG |
| 991 | Non_Targeting_Human | TCCTCTTCGCGTGCACCGC |
| 992 | Non_Targeting_Human | CTTTGCCCACAGGACACGA |
| 993 | Non_Targeting_Human | CGTCGCCAGTCACTCACAG |
| 994 | Non_Targeting_Human | AAAGGAGGTCTAGCGGCAA |
| 995 | Non_Targeting_Human | ACTACGCGAGGATGTCGAG |
| 996 | Non_Targeting_Human | GAGTTCCGTGCCGCCTGGT |
| 997 | Non_Targeting_Human | GTGGCACAGCGCTGGCCAG |
| 998 | Non_Targeting_Human | TGGGCAGCTGGGCGGAGCG |
| 999 | Non_Targeting_Human | ACCATTCCCATGTTCGTAA |
| 1000 | Non_Targeting_Human | CCTCGCCTGCTCACATTGG |
| 1001 | Non_Targeting_Human | AGCCCACCGACGAAAACCG |
| 1002 | Non_Targeting_Human | AGGGCAACGAATAGTTCCG |
| 1003 | Non_Targeting_Human | TCAGGGGTGCTAACATGGA |
| 1004 | Non_Targeting_Human | GCGCGGCCAGCAGGCAATT |
| 1005 | Non_Targeting_Human | GGCGTGCAACAGGTGGACT |
| 1006 | Non_Targeting_Human | AGGGTATGAGAAATCCGAG |
| 1007 | Non_Targeting_Human | TGACCCCAGGCGCCGCGCG |
| 1008 | Non_Targeting_Human | GAACAGCGCTGGATCGCAC |
| 1009 | Non_Targeting_Human | CAGCAATGAAGCGTCCTAA |
| 1010 | Non_Targeting_Human | CCCAGGAGTGCAATTGGGC |
| 1011 | Non_Targeting_Human | CCGGGCTTGGTCGGGGCGT |
| 1012 | Non_Targeting_Human | GCCTTTGTAGGCTTCGCCA |
| 1013 | Non_Targeting_Human | GATCAACGGCGGTGCGTGC |
| 1014 | Non_Targeting_Human | GCTGCGTCCCGCATCATAG |
| 1015 | Non_Targeting_Human | TCGAACCCGGGGTTCCACT |
| 1016 | Non_Targeting_Human | TGAGTAGAAGGCGTACCCA |
| 1017 | Non_Targeting_Human | CTTGATGTAAAGCGTACGA |
| 1018 | Non_Targeting_Human | TCAACCAACCCTCCGGCTC |
| 1019 | Non_Targeting_Human | AGAGGCCATCACGTTATGG |
| 1020 | Non_Targeting_Human | TTCGCATGGGGAACAACTG |
| 1021 | Non_Targeting_Human | TAGTCGCTGCAACCCTGGA |
| 1022 | Non_Targeting_Human | GCAAGTGCCTAGACTATAC |
| 1023 | Non_Targeting_Human | CGGTAGAGGGCCCTCACTA |
| 1024 | Non_Targeting_Human | TACGGCATGGCGCACTGCG |
| 1025 | Non_Targeting_Human | CATGGCGTAGCCAACGCAA |
| 1026 | Non_Targeting_Human | CTGATACCGTAAATGACCC |
| 1027 | Non_Targeting_Human | ACACGGTGGCGTACGGCGA |
| 1028 | Non_Targeting_Human | ACGCCCCTGCAAGCGTGGT |
| 1029 | Non_Targeting_Human | GCTGCCCACACGGCCATCA |
| 1030 | Non_Targeting_Human | GTCCTGAGGCCCTAGGCGA |
| 1031 | Non_Targeting_Human | CAGGTGAGCCTGTGTCGGA |

| | | |
|---|---|---|
| 1032 | Non_Targeting_Human | CGCCCGTCGCCCAAACCTT |
| 1033 | Non_Targeting_Human | TACTCCGCCAGGCGTCCGC |
| 1034 | Non_Targeting_Human | GGCCCTAGGCTTTGAGCTA |
| 1035 | Non_Targeting_Human | AGCCCTCCTCCATTACGGA |
| 1036 | Non_Targeting_Human | GAGTCGGGTCCTAAAGCTA |
| 1037 | Non_Targeting_Human | AGCGGGCATCAGAAGCGCA |
| 1038 | Non_Targeting_Human | CTTGCCGCGCTTCGATCCG |
| 1039 | Non_Targeting_Human | CGATCCCGTCAACCGTCGC |
| 1040 | Non_Targeting_Human | CTGGCGTCCACAGGAAGCT |
| 1041 | Non_Targeting_Human | ATGTCCACATGGCGTTGTC |
| 1042 | Non_Targeting_Human | TGAGGTTCACCTCACGGAT |
| 1043 | Non_Targeting_Human | CGTGACGGCCCGCAACGCG |
| 1044 | Non_Targeting_Human | GTGCTAGGCCGCGAAACTT |
| 1045 | Non_Targeting_Human | GTATTCCTAATCGTCGTTC |
| 1046 | Non_Targeting_Human | TGTCTGTGACCTAGGGTGG |
| 1047 | Non_Targeting_Human | CGTCGCTACAGCGAATTGG |
| 1048 | Non_Targeting_Human | TTACCAACGGACGAAGGCT |
| 1049 | Non_Targeting_Human | CGGCGTGTACGCAAATGGC |
| 1050 | Non_Targeting_Human | CAGGGTATAGAGGCCATAT |
| 1051 | Non_Targeting_Human | GAATTAGACGGAGCAGCGC |
| 1052 | Non_Targeting_Human | GTCCGCTTTTAAGCGGCTT |
| 1053 | Non_Targeting_Human | AAGTTCGGGCCCAGGTAGG |
| 1054 | Non_Targeting_Human | CGTTCCAAAGAACCACGGG |
| 1055 | Non_Targeting_Human | GTACGACACAAGGTATTAT |
| 1056 | Non_Targeting_Human | GCACAGCATCAGTACCCGA |
| 1057 | Non_Targeting_Human | GCAGTCCACTATAAAGATG |
| 1058 | Non_Targeting_Human | AAAAATCTTACTGGCAGGA |
| 1059 | Non_Targeting_Human | GGTAGATAGCACCGCTTAC |
| 1060 | Non_Targeting_Human | CATGCTCCCCCTCGCCCTT |
| 1061 | Non_Targeting_Human | GCCGGCGCCAACAGCAAGA |
| 1062 | Non_Targeting_Human | AGTATCCTGCGCACGGCCT |
| 1063 | Non_Targeting_Human | ACCGGAAAGTCAGATGTAC |
| 1064 | Non_Targeting_Human | AAGCTGTCGCATTCTGGTG |
| 1065 | Non_Targeting_Human | TTATTCATCGATCCTTGTC |
| 1066 | Non_Targeting_Human | GATCGCCTCCGTTTACCTA |
| 1067 | Non_Targeting_Human | TTGCCAATGAAGAATGGTG |
| 1068 | Non_Targeting_Human | CTTGGCGAGACGTGAGAAG |
| 1069 | Non_Targeting_Human | TAACGCCCGACGACCTGCC |
| 1070 | Non_Targeting_Human | GGGAGTGTGGCCGCCCTCT |
| 1071 | Non_Targeting_Human | CAGCCAGCGGGGTATCCTA |
| 1072 | Non_Targeting_Human | CTGAACCTCGTGGGCGCCC |
| 1073 | Non_Targeting_Human | CGAGCTGGGAGTGAAGCTC |
| 1074 | Non_Targeting_Human | GAAACGGGAGTGTCCGTTT |

| | | |
|---|---|---|
| 1075 | Non_Targeting_Human | AGGTGCCCCGTCGCCGGGG |
| 1076 | Non_Targeting_Human | ATCCGGGGATGACCACCTG |
| 1077 | Non_Targeting_Human | GCCCGGAACGGACGACCAC |
| 1078 | Non_Targeting_Human | GGACGAAGCAGTCCGAGCG |
| 1079 | Non_Targeting_Human | GTGGCTGAACTGGGACCAA |
| 1080 | Non_Targeting_Human | AACGCACTCCTCCGGACGC |
| 1081 | Non_Targeting_Human | ACTGCCTCGCGATTGACTG |
| 1082 | Non_Targeting_Human | GGGCACGCACTTTTACTGG |
| 1083 | Non_Targeting_Human | GTCCGAGACAGGGGCACGA |
| 1084 | Non_Targeting_Human | CAATATCGGGACTTATACT |
| 1085 | Non_Targeting_Human | GGGCAATGGGGTGTTTGAG |
| 1086 | Non_Targeting_Human | AGCGCCTCAAACCAAGTAA |
| 1087 | Non_Targeting_Human | CCCTCAGTTACCCAAGAAG |
| 1088 | Non_Targeting_Human | CTCGGACCTTACGCACCTC |
| 1089 | Non_Targeting_Human | TCAACCCGCTTTCCACGCA |
| 1090 | Non_Targeting_Human | AGCTGGGACGTCGGGACAC |
| 1091 | Non_Targeting_Human | GAACAGTGTATGCCTGTGG |
| 1092 | Non_Targeting_Human | TGTTCGGTTAGACAGTGCC |
| 1093 | Non_Targeting_Human | ACTGTTGTGAGTATGCCGA |
| 1094 | Non_Targeting_Human | CGGTGCACGAGCAGCACCG |
| 1095 | Non_Targeting_Human | GCATACTGACAGCCGCACG |
| 1096 | Non_Targeting_Human | CAGTAGATGGAATCAGGCG |
| 1097 | Non_Targeting_Human | GGCTCGGCCCTATACGTTC |
| 1098 | Non_Targeting_Human | ACCTACCCGGGAAGGGGCT |
| 1099 | Non_Targeting_Human | CGGCAGCGGCCCACGAGAA |
| 1100 | Non_Targeting_Human | AGTAATCCAAGGTGGGTGA |
| 1101 | Non_Targeting_Human | AACGTGCCCCCGGTCGTCG |
| 1102 | Non_Targeting_Human | GGAGATCGAGTGCGCGTGG |
| 1103 | Non_Targeting_Human | CGAGTTGGGTAGGCATAAG |
| 1104 | Non_Targeting_Human | TGGCGACGGCTAGTTCTGC |
| 1105 | Non_Targeting_Human | ACGACGCGCTGCTGGCAAA |
| 1106 | Non_Targeting_Human | CGAGTGTCTCTATTACCAA |
| 1107 | Non_Targeting_Human | CCACGTTGACCCTCTGCGA |
| 1108 | Non_Targeting_Human | CACCGGTAGGAGACACGCT |
| 1109 | Non_Targeting_Human | GTCGGAACTGGTGGTGCAA |
| 1110 | Non_Targeting_Human | CCGCACATACAGAAAAGTA |
| 1111 | Non_Targeting_Human | CCGTCTACAAGGGGGCTAG |
| 1112 | Non_Targeting_Human | CAGGCCTGGCCACAAGGAT |
| 1113 | Non_Targeting_Human | ACAGCCCGAGGGTGAATGG |
| 1114 | Non_Targeting_Human | GGCCCTCGCATGACGGCGA |
| 1115 | Non_Targeting_Human | GTCCGGCGTCGGCCGCTAT |
| 1116 | Non_Targeting_Human | ATAGGCTGCCTGTGATTAA |
| 1117 | Non_Targeting_Human | GTACGCCTGCGGTCGAGTA |

| | | |
|---|---|---|
| 1118 | Non_Targeting_Human | AACGGCGGGTACAAGACCC |
| 1119 | Non_Targeting_Human | GGGGAAGCAGTCTTTAGTC |
| 1120 | Non_Targeting_Human | GCCAGTTCTCCGAGCCCAG |
| 1121 | Non_Targeting_Human | AGGAGGCCCCGCCGGATTT |
| 1122 | Non_Targeting_Human | ATACGCTCCGGGATGGGCA |
| 1123 | Non_Targeting_Human | GCGCCTTCCGAGCGGTAGA |
| 1124 | Non_Targeting_Human | GGCGCGCGGCGGCACCTAG |
| 1125 | Non_Targeting_Human | TTCGCGAGCGGAAAGTACA |
| 1126 | Non_Targeting_Human | TGCGAGCAATCGCAGTAAC |
| 1127 | Non_Targeting_Human | CGGAGCGCCCGTCTTGAGG |
| 1128 | Non_Targeting_Human | CTGTTTCCCACAAACAAGA |
| 1129 | Non_Targeting_Human | GGAGTTCGACCATGCGTTG |
| 1130 | Non_Targeting_Human | GTTACCCCGTTGAGCCGGA |
| 1131 | Non_Targeting_Human | GGCTAGGTTGCGCGGTCGA |
| 1132 | Non_Targeting_Human | TGGGCCGCTCGGCGGTGAA |
| 1133 | Non_Targeting_Human | GGCAGGCGCGCTAGAGAGG |
| 1134 | Non_Targeting_Human | ACGACCAGGCCCACGCAAG |
| 1135 | Non_Targeting_Human | CCGACCTAGCACGTTAGTG |
| 1136 | Non_Targeting_Human | CGCCTCGGTACGGAGCGAG |
| 1137 | Non_Targeting_Human | AAAGGGCGAACTCTATGAT |
| 1138 | Non_Targeting_Human | TCGCGCGAAATATGTTCAG |
| 1139 | Non_Targeting_Human | GGCTTCTGGCCGCTAGTAC |
| 1140 | Non_Targeting_Human | GTCCCGGGAACGGCGCTCA |
| 1141 | Non_Targeting_Human | GCTGGCGTTGCCAACGAAC |
| 1142 | Non_Targeting_Human | CAGTCGCAGGCCGCCAGCG |
| 1143 | Non_Targeting_Human | CCCTCACCGCGTGCGCTCA |
| 1144 | Non_Targeting_Human | CCGTGTATCGGGTAGCTGG |
| 1145 | Non_Targeting_Human | ATGCGGCGGGCGGAAGTTC |
| 1146 | Non_Targeting_Human | AACCAGCCGCGCCAGTCGG |
| 1147 | Non_Targeting_Human | CTCGAGGGAGAGACTAGCT |
| 1148 | Non_Targeting_Human | TTAGGCTCTTCACCAGCTT |
| 1149 | Non_Targeting_Human | CCTCCGGGTCAGCCAAAGG |
| 1150 | Non_Targeting_Human | GGCCCACAGCGACGTTGGG |
| 1151 | Non_Targeting_Human | CTTGAGGTTCCGAAATCGG |
| 1152 | Non_Targeting_Human | TGTTTTGCCCAGTAGGAGC |
| 1153 | Non_Targeting_Human | ATGTAAGGGCGAATTAGTG |
| 1154 | Non_Targeting_Human | CGGACCATCATGCCGGTGG |
| 1155 | Non_Targeting_Human | ACACCCGGGCAACCCTCCC |
| 1156 | Non_Targeting_Human | CGCTTCTCGCAAGCAGGCA |
| 1157 | Non_Targeting_Human | GTCTAGCTACAGGAGTCAG |
| 1158 | Non_Targeting_Human | CTCCTCGGACTAGACGAGA |
| 1159 | Non_Targeting_Human | GTGTAGACCTTCGCAATGG |
| 1160 | Non_Targeting_Human | TCTCTAGCGCGCTTAATGG |

| | | |
|---|---|---|
| 1161 | Non_Targeting_Human | CCTAGGGGTTTCTAAGGA |
| 1162 | Non_Targeting_Human | TGTCTAACACCACCGATAG |
| 1163 | Non_Targeting_Human | CGTAGAAGGATGGGCAGCG |
| 1164 | Non_Targeting_Human | GCGAAGCAATAACACACGG |
| 1165 | Non_Targeting_Human | TAACTTATAGACACCGCTG |
| 1166 | Non_Targeting_Human | GGAATAGCGAGGCATCGCC |
| 1167 | Non_Targeting_Human | TTCCGGGGATGCGTACCCA |
| 1168 | Non_Targeting_Human | TACGACCACTCGGTGATGG |
| 1169 | Non_Targeting_Human | CACAGGCAACAAAGTTAGT |
| 1170 | Non_Targeting_Human | CGGGCGCCGCGAAATGCGA |
| 1171 | Non_Targeting_Human | TACCAGCCACCAGTAGCAG |
| 1172 | Non_Targeting_Human | CATCATTGGAGATCCTCGA |
| 1173 | Non_Targeting_Human | TAGATTGACATCCAGCGCA |
| 1174 | Non_Targeting_Human | ATAACGGCCTGCGTTTTGG |
| 1175 | Non_Targeting_Human | GAATCGATGAAGTAACTGG |
| 1176 | Non_Targeting_Human | GAACAGGAGAAGGGAACGT |
| 1177 | Non_Targeting_Human | TAGCCGGTCGATAGCTGGG |
| 1178 | Non_Targeting_Human | CGTCAGGGCCCACCGCCGC |
| 1179 | Non_Targeting_Human | TTGCCGAATCTTTCGCGGA |
| 1180 | Non_Targeting_Human | CACAGCTTTCGGGGTAACT |
| 1181 | Non_Targeting_Human | GCGGTGATACCTCACCCGG |
| 1182 | Non_Targeting_Human | CAGAGGGCACCTAGACAAT |
| 1183 | Non_Targeting_Human | GGATTGTGGCCCGACGCGA |
| 1184 | Non_Targeting_Human | CTGTGTCAACTGGCACCGA |
| 1185 | Non_Targeting_Human | CGGAGTCCTGAGCACGTTG |
| 1186 | Non_Targeting_Human | GTACTCCGAACAGCCGCTA |
| 1187 | Non_Targeting_Human | AGCTGCGGTCGGTGGCCGG |
| 1188 | Non_Targeting_Human | CGGTGTCGTCTCGCCCAAA |
| 1189 | Non_Targeting_Human | CGTCACGTCAGCCGTACGT |
| 1190 | Non_Targeting_Human | TGTGGAAACCATTGGCCGG |
| 1191 | Non_Targeting_Human | CCCTCAAGGGCACGTTTAG |
| 1192 | Non_Targeting_Human | GAAGACGGAGAACGGCATG |
| 1193 | Non_Targeting_Human | GTGCCCGCCTGGCCGGGAA |
| 1194 | Non_Targeting_Human | TCTGAGATGACCTCGGCCG |
| 1195 | Non_Targeting_Human | CGGCGGTAGCAGGCTCGCA |
| 1196 | Non_Targeting_Human | CAACGCGTGCTGAGGGGGG |
| 1197 | Non_Targeting_Human | TGCCTCCCGGGTCGTACAG |
| 1198 | Non_Targeting_Human | GCGCCTGTCGCCCAGTGCA |
| 1199 | Non_Targeting_Human | CCTACGGGAAGAGACGCGA |
| 1200 | Non_Targeting_Human | TGAAAAGAAGCCGATCGCC |
| 1201 | Non_Targeting_Human | CTTCACCAATAGGAACCCG |
| 1202 | Non_Targeting_Human | CGGATTCCCCCCACGGCGA |
| 1203 | Non_Targeting_Human | TTTCACGAGGGGCACGACC |

| | | |
|---|---|---|
| 1204 | Non_Targeting_Human | GCTCCCGCGATCGGGGACA |
| 1205 | Non_Targeting_Human | CGACGCCAACGTACCTCTT |
| 1206 | Non_Targeting_Human | CAGACACGGACGTTCTGAG |
| 1207 | Non_Targeting_Human | GGTGGCCTGAGCTAGTACA |
| 1208 | Non_Targeting_Human | GAATTGTCAGCGGGTCGAC |
| 1209 | Non_Targeting_Human | GCACCGACGGAAAAGTTTT |
| 1210 | Non_Targeting_Human | CTGCCAACGGATCCCCGAG |
| 1211 | Non_Targeting_Human | GGGACAGCTATCGGCTCAG |
| 1212 | Non_Targeting_Human | ACGTGATGTATTAAGCGGG |
| 1213 | Non_Targeting_Human | GCACCCCATTCGCCCAGGG |
| 1214 | Non_Targeting_Human | GTGCTTTGCGGGAAACCAA |
| 1215 | Non_Targeting_Human | AGGCAATGAGTCCCTACTA |
| 1216 | Non_Targeting_Human | ACAGGGACTCGGCAAGAGA |
| 1217 | Non_Targeting_Human | GGTCTGGGATCGCATCGCT |
| 1218 | Non_Targeting_Human | CACGGTGCTCGCGCAAACT |
| 1219 | Non_Targeting_Human | CCGCCCGTTGAAACAAGGG |
| 1220 | Non_Targeting_Human | CGCTTGCGGAAATGGAAGG |
| 1221 | Non_Targeting_Human | CGGAACTGACAGGAAACAT |
| 1222 | Non_Targeting_Human | GCTGCGCCGTCGAGAGTAG |
| 1223 | Non_Targeting_Human | CTAGGTACCAATGCAACGG |
| 1224 | Non_Targeting_Human | GCTCCAGCGCACCCGTCCG |
| 1225 | Non_Targeting_Human | CGTGCCAAGGGGTTAAGGC |
| 1226 | Non_Targeting_Human | GGCGCCTCCTGATTATCAC |
| 1227 | Non_Targeting_Human | CCCCTGGCGCAGGCCACGG |
| 1228 | Non_Targeting_Human | CAGGCTTCCCTGCGCAGGG |
| 1229 | Non_Targeting_Human | CCCCGCGCCCACCGTCGAG |
| 1230 | Non_Targeting_Human | GCCCGAACGAAGCGCTCCG |
| 1231 | Non_Targeting_Human | TCATACGGATAACCTCCGG |
| 1232 | Non_Targeting_Human | GGAGCCCCGGGCGAATCAT |
| 1233 | Non_Targeting_Human | GCCCGACGATGCAGCCAGA |
| 1234 | Non_Targeting_Human | TCCCAAACGTCTCTCAACA |
| 1235 | Non_Targeting_Human | GATAAAAAAACATCAGCCC |
| 1236 | Non_Targeting_Human | CACTGGGAGGACCGCCTGC |
| 1237 | Non_Targeting_Human | GGGCCGGCGTGCCCCCTCG |
| 1238 | Non_Targeting_Human | AACAGACCCAGCACCAAGC |
| 1239 | Non_Targeting_Human | CGGACTCGAGGCTCTCCCG |
| 1240 | Non_Targeting_Human | TCCAATTCCTGGCTAGCCG |
| 1241 | Non_Targeting_Human | CGTGGGGCGGCCTCAGCAT |
| 1242 | Non_Targeting_Human | CCCAGCGGAGAGCAACGGG |
| 1243 | Non_Targeting_Human | CGGGCTACACGCATAATGG |
| 1244 | Non_Targeting_Human | CCAGCCTTCCGGCTAGAAC |
| 1245 | Non_Targeting_Human | GGCGTCGAGGATCGGGCGC |
| 1246 | Non_Targeting_Human | ATACTCTGCAAGACAACAG |

| | | |
|---|---|---|
| 1247 | Non_Targeting_Human | TGAGATCCGGCGCGTGAAG |
| 1248 | Non_Targeting_Human | CCGAAAAAGTAGGGGAATA |
| 1249 | Non_Targeting_Human | TACACAGAGGCGCCGGCCG |
| 1250 | Non_Targeting_Human | GGATCTCCGAGACGCGTGG |
| 1251 | Non_Targeting_Human | TGCGAGGTGCATACAGTAG |
| 1252 | Non_Targeting_Human | GCCGGCACGTAAAGCACCC |
| 1253 | Non_Targeting_Human | GGAAGCCGTCCGTACGAAT |
| 1254 | Non_Targeting_Human | CGTACGTATGACGACGGGG |
| 1255 | Non_Targeting_Human | AAAGGTGGCAGACAGCTCT |
| 1256 | Non_Targeting_Human | GACCCCTGCGCAATTGATT |
| 1257 | Non_Targeting_Human | GCAATCCGTGTACGTGTCA |
| 1258 | Non_Targeting_Human | CCCAGGTGACTTAAGGCTG |
| 1259 | Non_Targeting_Human | CATAAGAGTATCTTCCTGA |
| 1260 | Non_Targeting_Human | AGAGTCGTAATGTGCAGGG |
| 1261 | Non_Targeting_Human | TTGGCAGAGCAGCGGCCAA |
| 1262 | Non_Targeting_Human | GGAGGTCCGAACAAACGAA |
| 1263 | Non_Targeting_Human | AGGAAGCGCGCGCCATCGG |
| 1264 | Non_Targeting_Human | GATGCCGCTCATTTAGCTT |
| 1265 | Non_Targeting_Human | TCCCGCGCAGTCATGACCG |
| 1266 | Non_Targeting_Human | TGCTCTACGCAGAAGCGCC |
| 1267 | Non_Targeting_Human | TGCTGGCGGTATTGGGATA |
| 1268 | Non_Targeting_Human | TCGCCAGCGCAAGAGCCTA |
| 1269 | Non_Targeting_Human | GAACGCTTTATATACTTAA |
| 1270 | Non_Targeting_Human | TCGCAGTTTGCCCCGGGTC |
| 1271 | Non_Targeting_Human | TTCCCGTGGCTCTGCACCA |
| 1272 | Non_Targeting_Human | CCGCCTGTAATCACCCGCG |
| 1273 | Non_Targeting_Human | AGTACTGGTCTGACCACTG |
| 1274 | Non_Targeting_Human | TTCCCCCATTGAATACTTA |
| 1275 | Non_Targeting_Human | CTGAAGCCCGGCGTTTCTT |
| 1276 | Non_Targeting_Human | CTAACAAATACATCGGCAG |
| 1277 | Non_Targeting_Human | CGCTAAACACACTGCATGA |
| 1278 | Non_Targeting_Human | GTCGGCCTGAACGGCACTG |
| 1279 | Non_Targeting_Human | GTTCAGCCAAAATCCCCCG |
| 1280 | Non_Targeting_Human | CAGCGCCGCAGAGAACTAC |
| 1281 | Non_Targeting_Human | GAGATGAGTCGTAGCGAGA |
| 1282 | Non_Targeting_Human | CCCCCTCGCTAGAGAACAA |
| 1283 | Non_Targeting_Human | TCGGAGGCCGAGGACGGAA |
| 1284 | Non_Targeting_Human | CGTGTGCCCCGCTCCGAAC |
| 1285 | Non_Targeting_Human | CACCCGGCGGGTGCCTCTT |
| 1286 | Non_Targeting_Human | GCGCGGACCCGCTAGCGGA |
| 1287 | Non_Targeting_Human | GGGCGGGTCAAGGAAACTT |
| 1288 | Non_Targeting_Human | GGTTACCGGTAACTCAATC |
| 1289 | Non_Targeting_Human | GTGCTGCCCCTACGGCCCG |

| | | |
|---|---|---|
| 1290 | Non_Targeting_Human | GAACCGCGACGCCGGGGTG |
| 1291 | Non_Targeting_Human | CCCCGAGTTAAGGGATTGC |
| 1292 | Non_Targeting_Human | ATGTGGCTGCCGCCATCGG |
| 1293 | Non_Targeting_Human | GTACGAATCAGCAGTGCGC |
| 1294 | Non_Targeting_Human | GCACGCAGACCGTACGGTG |
| 1295 | Non_Targeting_Human | GCGGATAACGCATATGAAC |
| 1296 | Non_Targeting_Human | GATACACAAAGCCTAGTGT |
| 1297 | Non_Targeting_Human | TGGGTCCCGGACGAATAAG |
| 1298 | Non_Targeting_Human | ATTGAAGCCAAGGACCCAT |
| 1299 | Non_Targeting_Human | GGGACTTACCGTACACCAG |
| 1300 | Non_Targeting_Human | TGGGCGGCACTGCGTTAGG |
| 1301 | Non_Targeting_Human | CGAGCAAGCAGGCGCCACA |
| 1302 | Non_Targeting_Human | TGCACAGTGCAGCCATCGA |
| 1303 | Non_Targeting_Human | GTACAATGTCTTGTGGTCA |
| 1304 | Non_Targeting_Human | CAGGCCGAGTGCCTATCTG |
| 1305 | Non_Targeting_Human | GCACCTTAGCTGTCCACGG |
| 1306 | Non_Targeting_Human | TCAGCCGTCCCGCACACAG |
| 1307 | Non_Targeting_Human | GCCGTGTCTGGACCCGGGG |
| 1308 | Non_Targeting_Human | GCGAGGCCGAGACAAGTGG |
| 1309 | Non_Targeting_Human | CCGAAGCTCAAGCGTAAGA |
| 1310 | Non_Targeting_Human | GGGCCGGGCGGCGACTGGT |
| 1311 | Non_Targeting_Human | CGACTAGCACTAGGAACGC |
| 1312 | Non_Targeting_Human | CGAGCCCCAGTGAAATCGA |
| 1313 | Non_Targeting_Human | AGGGTGGCAAGGCCAGACC |
| 1314 | Non_Targeting_Human | CGTAGGCGGACTCACAACT |
| 1315 | Non_Targeting_Human | AAAAGACCGCTCGGGCATA |
| 1316 | Non_Targeting_Human | GGGGACCCTCCAGAAAAAG |
| 1317 | Non_Targeting_Human | CGTGTCGCAAGCGATGCAA |
| 1318 | Non_Targeting_Human | CGATCCGGGGCGGACCCAA |
| 1319 | Non_Targeting_Human | ATGCGAACTGCACGGAACG |
| 1320 | Non_Targeting_Human | AGGTACGCCAGCCAACATC |
| 1321 | Non_Targeting_Human | CCATGAGGGCGAGATTACG |
| 1322 | Non_Targeting_Human | GTCGCGTACCGGGGTGGGG |
| 1323 | Non_Targeting_Human | TCATCGCCTCGTCAGAACC |
| 1324 | Non_Targeting_Human | GCTAGGGCTCATCTCAGAG |
| 1325 | Non_Targeting_Human | CAGGGCGCACCGGAGGAGG |
| 1326 | Non_Targeting_Human | GAAGTGGACTGCACTGCAA |
| 1327 | Non_Targeting_Human | GAACCGCGGGACCTCTCCG |
| 1328 | Non_Targeting_Human | GTTGAGGGGTAGGCGTTTG |
| 1329 | Non_Targeting_Human | CTGATGCTGAACCTCACGG |
| 1330 | Non_Targeting_Human | CAAGAGAACTCAAGTAGAG |
| 1331 | Non_Targeting_Human | CCGGACAGGGGGGTTCGGG |
| 1332 | Non_Targeting_Human | CCGATCTCGGTCCCCCATA |

| | | |
|---|---|---|
| 1333 | Non_Targeting_Human | GCGCGAAGAGTCCCGCCGG |
| 1334 | Non_Targeting_Human | TCTTCGGCCCCTCCCATCG |
| 1335 | Non_Targeting_Human | AGATCAATATTATGCAAGA |
| 1336 | Non_Targeting_Human | GGACGCTGGGTACGAGAGG |
| 1337 | Non_Targeting_Human | GCCCTTACACTCTCAAATA |
| 1338 | Non_Targeting_Human | ACCGTGCGTCCACAAACAG |
| 1339 | Non_Targeting_Human | ATAGGACTCCAACCAACGA |
| 1340 | Non_Targeting_Human | GAAGCGGTAGTCTGCAGCA |
| 1341 | Non_Targeting_Human | GGAAAGCACAGGAGCGCAT |
| 1342 | Non_Targeting_Human | GTGGGCCCGTCCAGAGCGG |
| 1343 | Non_Targeting_Human | CGCCCCACTCGTCAAGTTC |
| 1344 | Non_Targeting_Human | AGTAACACAGGACGAGCTT |
| 1345 | Non_Targeting_Human | TCCGACCCGGGCCGTACAG |
| 1346 | Non_Targeting_Human | CGGGTATTGTCCAATGTCG |
| 1347 | Non_Targeting_Human | CACCGGGAAACATCTTCCG |
| 1348 | Non_Targeting_Human | GCGGGAGGCGCGCCATCAG |
| 1349 | Non_Targeting_Human | CCGTATGTGTGTGGGCCAA |
| 1350 | Non_Targeting_Human | CGCCCGGACGGGACCGAAG |
| 1351 | Non_Targeting_Human | CGCGCGCCGCCGGCGTTAA |
| 1352 | Non_Targeting_Human | GGGGTACCACGAGGTGCAT |
| 1353 | Non_Targeting_Human | GCACCGTAATATAACGCTA |
| 1354 | Non_Targeting_Human | TTGGGCGTGGCTCTACTAG |
| 1355 | Non_Targeting_Human | AGGTACTGGTATGAAAGTG |
| 1356 | Non_Targeting_Human | TCCCGGGATCAAGCCGAGG |
| 1357 | Non_Targeting_Human | GGATTGGCTAAGAGGGTTC |
| 1358 | Non_Targeting_Human | CCGTCGTCGCATACGGGAA |
| 1359 | Non_Targeting_Human | TCCGCGTGGAAAAAAACGT |
| 1360 | Non_Targeting_Human | TCATAGCGAGACGGTGAAG |
| 1361 | Non_Targeting_Human | GACGGGGCACCGTGCAAGG |
| 1362 | Non_Targeting_Human | GTCAGGACGCCACAAGTGC |
| 1363 | Non_Targeting_Human | CGGGTGCTCCAACGAAGCG |
| 1364 | Non_Targeting_Human | AGCGTTCTGTGCAACCCGA |
| 1365 | Non_Targeting_Human | GACGAGCACTGCGAGAGCG |
| 1366 | Non_Targeting_Human | AGATTCGCACCGAACACGG |
| 1367 | Non_Targeting_Human | GCGATGGCCGGAGTTTCTC |
| 1368 | Non_Targeting_Human | GACGATTGGTTGAGGCCGG |
| 1369 | Non_Targeting_Human | GAGTGAGGTCGTGGACGTG |
| 1370 | Non_Targeting_Human | AAGACAGCCAGCAACGACG |
| 1371 | Non_Targeting_Human | TACCAAAAGGGGTTCACAG |
| 1372 | Non_Targeting_Human | GAACCCCGTCGGTGTAGCG |
| 1373 | Non_Targeting_Human | GCCTTTGAATGCAACCGCG |
| 1374 | Non_Targeting_Human | AGCGGAGCTGGAGTAATAA |
| 1375 | Non_Targeting_Human | GACCGGCTTACCGAGTACT |

| | | |
|---|---|---|
| 1376 | Non_Targeting_Human | TGGTTGGGCGGGTAGACGT |
| 1377 | Non_Targeting_Human | CCGTGCGGACATCATGGAG |
| 1378 | Non_Targeting_Human | TGGCGCCTTCGGACTACGT |
| 1379 | Non_Targeting_Human | CGCGGGGTGATAGAAGCTG |
| 1380 | Non_Targeting_Human | GCATAGGTATTGTGGCGCC |
| 1381 | Non_Targeting_Human | TATGGCGGGCGTAGAGCGT |
| 1382 | Non_Targeting_Human | ACACTCAGAGGCCGCGGCG |
| 1383 | Non_Targeting_Human | AGGAACAGAAATTGAGCAT |
| 1384 | Non_Targeting_Human | GGCTCGCCGTGGCTCGATA |
| 1385 | Non_Targeting_Human | GGAGCGGGCTCGTATTCAG |
| 1386 | Non_Targeting_Human | GTTTTCGGACGAGGTGGAA |
| 1387 | Non_Targeting_Human | TAGAGCCAGGGCCCCCGAC |
| 1388 | Non_Targeting_Human | CTCGTGCCCCCGAAAAGAG |
| 1389 | Non_Targeting_Human | TAAGCGGTGGTGAATTCTA |
| 1390 | Non_Targeting_Human | GGCGCAGGGGCCGAGTACG |
| 1391 | Non_Targeting_Human | CGAGGCGACGAGGCCGGGT |
| 1392 | Non_Targeting_Human | TCGGCGCGACACCCTAAAC |
| 1393 | Non_Targeting_Human | CCCATGGGACCGGTCTGCT |
| 1394 | Non_Targeting_Human | GATGTAGAGGGACACCCAG |
| 1395 | Non_Targeting_Human | TGGTCAATTAACGCTGCAA |
| 1396 | Non_Targeting_Human | CAGGTGATGCCCCGGATTG |
| 1397 | Non_Targeting_Human | GCAGAATGTGTGCGTGCCA |
| 1398 | Non_Targeting_Human | AGCTCCCCCGAGCGGTGGA |
| 1399 | Non_Targeting_Human | CCATGGTGAGGACCACGGT |
| 1400 | Non_Targeting_Human | AAAGCGGTCAAGGCATTAA |
| 1401 | Non_Targeting_Human | CCGAATGAGAGATTCCCTG |
| 1402 | Non_Targeting_Human | TCGATCGGCTCCTTATGAG |
| 1403 | Non_Targeting_Human | GTCCTAGTCGGGCACTACT |
| 1404 | Non_Targeting_Human | GTCGTAACCCAGCTCGATC |
| 1405 | Non_Targeting_Human | GCGGCCGGTGCAACGCTAT |
| 1406 | Non_Targeting_Human | CGCACCTTGGTCTATTGAT |
| 1407 | Non_Targeting_Human | CGGCGAAGAGGAACAATGG |
| 1408 | Non_Targeting_Human | CTGCAAGTGTACTTATCAA |
| 1409 | Non_Targeting_Human | CACCACTGGCATGCTTGAA |
| 1410 | Non_Targeting_Human | CTAGCCAGGGGACTAGCGC |
| 1411 | Non_Targeting_Human | TGCGCGCTTATCGGTCCCA |
| 1412 | Non_Targeting_Human | GTGCCGCATATTGTGGACC |
| 1413 | Non_Targeting_Human | ACCGGGAAGGCGTGAAGGG |
| 1414 | Non_Targeting_Human | CAGGCCGCCCTGCTAAGAA |
| 1415 | Non_Targeting_Human | TGGCATCCATCGTAGAGTG |
| 1416 | Non_Targeting_Human | GTCTGCAGGCGTGCTGTTG |
| 1417 | Non_Targeting_Human | CGCACATACCTATATTACG |
| 1418 | Non_Targeting_Human | GGGCCAAACAACCCCGCGA |

| | | |
|---|---|---|
| 1419 | Non_Targeting_Human | CAAATGCGGCTGTCACCAG |
| 1420 | Non_Targeting_Human | TTTCATATGGGGACAGCAT |
| 1421 | Non_Targeting_Human | TGACGGAGACTAGCCTCTG |
| 1422 | Non_Targeting_Human | TCGCGGCCTTAGGATGTAC |
| 1423 | Non_Targeting_Human | CCGCCATAGCCGGATCGAG |
| 1424 | Non_Targeting_Human | TCGTTGGCAGAGTTCGCGT |
| 1425 | Non_Targeting_Human | GAGCAAGACGGTCAGGTGA |
| 1426 | Non_Targeting_Human | GGGAACGCGACGGAACCCA |
| 1427 | Non_Targeting_Human | GGCGGGCTTGGGGTTCGTC |
| 1428 | Non_Targeting_Human | AGCCGACTAGAACCGCTTT |
| 1429 | Non_Targeting_Human | CTTCGCGGTCGATATGAGG |
| 1430 | Non_Targeting_Human | CCTCTCATCAGGGTTCTGA |
| 1431 | Non_Targeting_Human | AGTAGAGGCCCCCGAGCTC |
| 1432 | Non_Targeting_Human | CCAAACCACATCGTGGGGG |
| 1433 | Non_Targeting_Human | CGCTCTGCGGTCCCGGATA |
| 1434 | Non_Targeting_Human | CCGGGGCCGACATTCCGCT |
| 1435 | Non_Targeting_Human | AGGCGCAAGCGGACGTGTG |
| 1436 | Non_Targeting_Human | GACCTGCCGGTGGAACTTG |
| 1437 | Non_Targeting_Human | GTGAAGCATCGCGACAACT |
| 1438 | Non_Targeting_Human | TTTTAGCCCCCACAGTCCT |
| 1439 | Non_Targeting_Human | TCCAGTGCGCAGACGGGAG |
| 1440 | Non_Targeting_Human | CGTTCAGATGCGTCTTGGC |
| 1441 | Non_Targeting_Human | CGACTCATATACACGGCGA |
| 1442 | Non_Targeting_Human | CTGCAACCGACGACCGGCC |
| 1443 | Non_Targeting_Human | CTGCGGGCGCGGACAGTGC |
| 1444 | Non_Targeting_Human | CGTCCGAGGTACTGAATAA |
| 1445 | Non_Targeting_Human | ATACGTGGGCGTGGCGCGG |
| 1446 | Non_Targeting_Human | AGGAAAACGCCGGGTGAAA |
| 1447 | Non_Targeting_Human | ATAACATCTGGCCAGGTAG |
| 1448 | Non_Targeting_Human | CGAATTCGGGGTCTTCCAG |
| 1449 | Non_Targeting_Human | CGAGCTGAACCGGGGCCCT |
| 1450 | Non_Targeting_Human | GCTAGAACCCACACTCTTA |
| 1451 | Non_Targeting_Human | GGGCCTAAAGCGTGGCCTC |
| 1452 | Non_Targeting_Human | AATGCAGCGCTCCGGCCGA |
| 1453 | Non_Targeting_Human | GCACCAAGGGAGTACGAAG |
| 1454 | Non_Targeting_Human | CGTATGGCGCTCAATCGCA |
| 1455 | Non_Targeting_Human | CCCCGTACTGCCGCTTCGG |
| 1456 | Non_Targeting_Human | ACATTAGCCAGATGAAAGA |
| 1457 | Non_Targeting_Human | GGTAATAGGACGACCCGAG |
| 1458 | Non_Targeting_Human | CGACAGCACAAATCGGTCA |
| 1459 | Non_Targeting_Human | CGTTGCTATCGGGCATAGG |
| 1460 | Non_Targeting_Human | GCGTGCTTAACTCCACCCG |
| 1461 | Non_Targeting_Human | GACGTTACTGAGGAAAATT |

| | | |
|---|---|---|
| 1462 | Non_Targeting_Human | CGCCTCTAAACTATCGCAA |
| 1463 | Non_Targeting_Human | CCCAGCGCATACAGGGCTA |
| 1464 | Non_Targeting_Human | CAAATGACAACACATGCGA |
| 1465 | Non_Targeting_Human | CCCGGAAGCGACATACGAC |
| 1466 | Non_Targeting_Human | ACCCCACCGAACCCGGTGT |
| 1467 | Non_Targeting_Human | GCCCATACGTCGTTCTTAT |
| 1468 | Non_Targeting_Human | TGTACAGCGCGGGCCGGAT |
| 1469 | Non_Targeting_Human | TAACAGGGCCCACTACGAC |
| 1470 | Non_Targeting_Human | CTGCAACGTGGTGCTGCCG |
| 1471 | Non_Targeting_Human | AGGCGTAGCGTAATGGGTG |
| 1472 | Non_Targeting_Human | ACCTCGGAGGGGAATCCGT |
| 1473 | Non_Targeting_Human | CTTACGAGGGGGCGCTGCT |
| 1474 | Non_Targeting_Human | GCCGAGGTCACCGCAGCAT |
| 1475 | Non_Targeting_Human | CGTCCATTGAGGGGTCCCA |
| 1476 | Non_Targeting_Human | CTGCAGGCCACCCGTGGCG |
| 1477 | Non_Targeting_Human | CCCGTTCGACGTTTACCGC |
| 1478 | Non_Targeting_Human | GTACTAGCCACCCCTATGC |
| 1479 | Non_Targeting_Human | ACTACGGATACTCGTGTGC |
| 1480 | Non_Targeting_Human | CTCAGGCGAAACCTCAAAG |
| 1481 | Non_Targeting_Human | TCTCTTTAGGAGGGCGAGA |
| 1482 | Non_Targeting_Human | TGCGAACTACACCAAACGA |
| 1483 | Non_Targeting_Human | TTGGCTCCTTCTTATCATG |
| 1484 | Non_Targeting_Human | ACTCAGTTGTTCTTTCGAA |
| 1485 | Non_Targeting_Human | CCGGGCCCCAAGGAATCAT |
| 1486 | Non_Targeting_Human | AGTACTGATGTGACGATGG |
| 1487 | Non_Targeting_Human | CAGTCTGCGCTATGACACC |
| 1488 | Non_Targeting_Human | CGGATGATGAGCGACACTA |
| 1489 | Non_Targeting_Human | GGCTTCACGCTGGCCTCAA |
| 1490 | Non_Targeting_Human | AGGCTTTCAGACAACTTAG |
| 1491 | Non_Targeting_Human | TAGGAAGCTCACGGCCGCA |
| 1492 | Non_Targeting_Human | CAGAGCATGCCGGTCGAGG |
| 1493 | Non_Targeting_Human | GAGTTGGAGTGACCAGCGA |
| 1494 | Non_Targeting_Human | CAGATGCAAACCCGTATGT |
| 1495 | Non_Targeting_Human | CGGGCGCAGCGCCGACCAC |
| 1496 | Non_Targeting_Human | TGGGGCGAATTAGCATGCC |
| 1497 | Non_Targeting_Human | GCACACGTGAGGTACAACA |
| 1498 | Non_Targeting_Human | AACGAAGACGAACGCTGCG |
| 1499 | Non_Targeting_Human | ACCTGTAGACCCGACTGAG |
| 1500 | Non_Targeting_Human | CTTCAACGGTCAACCTAGA |
| 1501 | Non_Targeting_Human | GGCCGGGGATGGTTAATCC |
| 1502 | Non_Targeting_Human | CTTTATTACCGCAAGGAGA |
| 1503 | Non_Targeting_Human | CCCGTGCACACCGCGGAAG |
| 1504 | Non_Targeting_Human | AGGTAGGCGTGCGCACGAA |

| | | |
|---|---|---|
| 1505 | Non_Targeting_Human | CCGCATGAACGACAAACCG |
| 1506 | Non_Targeting_Human | TGATAGCACTCGCCATCGC |
| 1507 | Non_Targeting_Human | CTACCCCCGGCAGCTTATT |
| 1508 | Non_Targeting_Human | ACGCTCCGACGCGGCCCGA |
| 1509 | Non_Targeting_Human | GTGTTCGAGGAACAGGCGA |
| 1510 | Non_Targeting_Human | TGGCACACAACCCGCGCGA |
| 1511 | Non_Targeting_Human | TCCGTGTTGACGTTTCCCT |
| 1512 | Non_Targeting_Human | GACTCCCTCGCCCTGGTGA |
| 1513 | Non_Targeting_Human | CGGGGCATTTAGCCCGCGT |
| 1514 | Non_Targeting_Human | GCGAAGGCGGTCACACCAG |
| 1515 | Non_Targeting_Human | ACCAACACAGCCGCCACGA |
| 1516 | Non_Targeting_Human | GTGGCAGGAAGCGGTTTAA |
| 1517 | Non_Targeting_Human | TTTCAGAAGGCGGCCATAT |
| 1518 | Non_Targeting_Human | CGCTAGCCGCAGAAGAGGC |
| 1519 | Non_Targeting_Human | GCGGGGGCCGTCCAAAAGG |
| 1520 | Non_Targeting_Human | TCTGATTTGGCCATGGCGG |
| 1521 | Non_Targeting_Human | TTCGCAGAAAGTGGCCTAA |
| 1522 | Non_Targeting_Human | GAGGGACACCGGTCAGGGA |
| 1523 | Non_Targeting_Human | CCAACACGCCCAATCAGCA |
| 1524 | Non_Targeting_Human | TTCGCCTATTTCTAACTTC |
| 1525 | Non_Targeting_Human | CGAATTAAGTGGCGCACCG |
| 1526 | Non_Targeting_Human | ACGGCCGAGTCAAATCCCA |
| 1527 | Non_Targeting_Human | CGTGCTGCGCGAACGGCCG |
| 1528 | Non_Targeting_Human | CTTAATGCGGAACACGAGA |
| 1529 | Non_Targeting_Human | GTAAATCCACGCGACGGAG |
| 1530 | Non_Targeting_Human | ATCGAAAGGTCCTCTTCTA |
| 1531 | Non_Targeting_Human | CGGGCGGCTGCACCCGGAA |
| 1532 | Non_Targeting_Human | GGTGGGGAGGCCTTGCCAC |
| 1533 | Non_Targeting_Human | GCGGCACAGGTACACCCAT |
| 1534 | Non_Targeting_Human | AGCGGTCCTGGAGGCGTGT |
| 1535 | Non_Targeting_Human | CGCGGAGTTCATCAGGGGA |
| 1536 | Non_Targeting_Human | GGGGCCACGCAGTACAGGC |
| 1537 | Non_Targeting_Human | GACGAAAGGCGCTGGTAGT |
| 1538 | Non_Targeting_Human | AGCCGATAATGGTCCTTTG |
| 1539 | Non_Targeting_Human | GCCCCGCGGACCAAAATTG |
| 1540 | Non_Targeting_Human | CGGGGATGTCCACACTCGG |
| 1541 | Non_Targeting_Human | GTTGTTGCGCCGCGCTCAC |
| 1542 | Non_Targeting_Human | CCGATTCGCTAATAATCGG |
| 1543 | Non_Targeting_Human | TAAACCGGAGTCCTCGCCG |
| 1544 | Non_Targeting_Human | TAGGGCCTAGCGAAAGCCG |
| 1545 | Non_Targeting_Human | CCGCAAAGGGCCGTTGATT |
| 1546 | Non_Targeting_Human | GCTCAATCCAGCGCCGATC |
| 1547 | Non_Targeting_Human | GAGAATTTCAAGCATATGC |

| | | |
|---|---|---|
| 1548 | Non_Targeting_Human | GACCGGTGGGTCGTGCCCA |
| 1549 | Non_Targeting_Human | GGCTAGGCACGGTAGCCTG |
| 1550 | Non_Targeting_Human | GTTACCTTAGAGCCCAGGC |
| 1551 | Non_Targeting_Human | GCATCGCAACTGGTGGTCG |
| 1552 | Non_Targeting_Human | CGGTTGCGGAGGACGAGTA |
| 1553 | Non_Targeting_Human | CGGCGCCACAGGAAAGCAT |
| 1554 | Non_Targeting_Human | AAGCTAAGGAAGATACCGG |
| 1555 | Non_Targeting_Human | CGTAAGCAGCCCATGGTTG |
| 1556 | Non_Targeting_Human | GCATCCCCAGGTCTTTCGT |
| 1557 | Non_Targeting_Human | TCTAAGGCATGTCGGCCAT |
| 1558 | Non_Targeting_Human | AAGCGTACACTCATTCAAG |
| 1559 | Non_Targeting_Human | AGTACGTCGGCTCCGGCCG |
| 1560 | Non_Targeting_Human | GATCACTGGCCGCAAGTAT |
| 1561 | Non_Targeting_Human | GCGCTCCCACCGATAAAGT |
| 1562 | Non_Targeting_Human | CACGATCGACGAAGATCGG |
| 1563 | Non_Targeting_Human | CGAGGCGGGGGCTTCATTC |
| 1564 | Non_Targeting_Human | CTGACGTCGGGCACTCGAA |
| 1565 | Non_Targeting_Human | AAGAATGTGAGAGCCGAAA |
| 1566 | Non_Targeting_Human | CCGCAGCGTAGCGACGCGC |
| 1567 | Non_Targeting_Human | ATTCCCCCAGGATGGTGAG |
| 1568 | Non_Targeting_Human | TTAGGGCGTAAGGGTTCGA |
| 1569 | Non_Targeting_Human | GTAGTTACAGATAAACCTG |
| 1570 | Non_Targeting_Human | TGGGGGCATCTAGTAAGTG |
| 1571 | Non_Targeting_Human | CCTTACCCCGTGATAGCGC |
| 1572 | Non_Targeting_Human | CCACGGCCTTCATCGCAGG |
| 1573 | Non_Targeting_Human | ACCTCTAAAGTGCCAGATA |
| 1574 | Non_Targeting_Human | AACCTTGGGAGCAGCGAGG |
| 1575 | Non_Targeting_Human | ATAATATCAAGCTGACGGG |
| 1576 | Non_Targeting_Human | TAAGTCCAACGGATCGTTG |
| 1577 | Non_Targeting_Human | ATCCTCGCCTCAGAACCAG |
| 1578 | Non_Targeting_Human | GAAACAATCGGAAGCCTAG |
| 1579 | Non_Targeting_Human | TCGGAACCTTTATACCACA |
| 1580 | Non_Targeting_Human | ACCCGGTACACGGACACGG |
| 1581 | Non_Targeting_Human | GGCCGGTCAATCAACTTTC |
| 1582 | Non_Targeting_Human | CCGCCAGCGGTCAGAGGGA |
| 1583 | Non_Targeting_Human | CCTGGCACCGCCCGAACCC |
| 1584 | Non_Targeting_Human | CTTGCGACTTACCCCGTCG |
| 1585 | Non_Targeting_Human | CAGCTACTTCCGTTTCGCG |
| 1586 | Non_Targeting_Human | ATGGGCGGGGTGCCTACTA |
| 1587 | Non_Targeting_Human | GAGTGCACGGATTAAGTTG |
| 1588 | Non_Targeting_Human | ACCCTAGGCGGCCCCGTGG |
| 1589 | Non_Targeting_Human | CCTGAACCGACCTACGAGG |
| 1590 | Non_Targeting_Human | CAACACCCCGTGCAAGTTA |

| | | |
|---|---|---|
| 1591 | Non_Targeting_Human | GCACGATCCGTGTGAAGGT |
| 1592 | Non_Targeting_Human | GTTCCCGCGTAGCACAAGG |
| 1593 | Non_Targeting_Human | TGGCGGCTGGAGCATCTTT |
| 1594 | Non_Targeting_Human | CCCACAGGAGGCCCCGCAG |
| 1595 | Non_Targeting_Human | GTCAGGCCCGGCTTAGTAG |
| 1596 | Non_Targeting_Human | ACGTACTTGGCCACCTAAA |
| 1597 | Non_Targeting_Human | CCATGGCCTGTCAATAGAA |
| 1598 | Non_Targeting_Human | CCGTAGCGGATGGAGGCGT |
| 1599 | Non_Targeting_Human | CTGGGGCAAAAGGCGACTA |
| 1600 | Non_Targeting_Human | GTTGGGCCCCAGTTAGTCG |
| 1601 | Non_Targeting_Human | GAAGGTCGTAAGCCCAACG |
| 1602 | Non_Targeting_Human | GCGCCAGGGCTGCGCGACA |
| 1603 | Non_Targeting_Human | TTATCAGGGTGGCCATCTT |
| 1604 | Non_Targeting_Human | CGGGGAGTTCCGGGTCTGC |
| 1605 | Non_Targeting_Human | CAGCCCCCAGAGAAAGAGC |
| 1606 | Non_Targeting_Human | GCTCCAGTTCCGCATGCCA |
| 1607 | Non_Targeting_Human | AGCCTCCCAGTATGAAGTT |
| 1608 | Non_Targeting_Human | GGAGCCGCAGCTAAAGATG |
| 1609 | Non_Targeting_Human | TCTGAAACGGGTGCGAGAA |
| 1610 | Non_Targeting_Human | CTCACTCGCCCAAGCCGTC |
| 1611 | Non_Targeting_Human | CGGGCGAAAGTCAGCACGT |
| 1612 | Non_Targeting_Human | GCGCCTGGTTCGAACCGTG |
| 1613 | Non_Targeting_Human | TCGCACCGGGCGTGATACG |
| 1614 | Non_Targeting_Human | CACGACGTGCAGAGTCAGA |
| 1615 | Non_Targeting_Human | ACATCTTCCTGCCTGCGGT |
| 1616 | Non_Targeting_Human | GTGGCGCGACCGTTCCTCG |
| 1617 | Non_Targeting_Human | CGGCGTTTTGACCAGCCGG |
| 1618 | Non_Targeting_Human | GACTCTCGATAGGAACCGC |
| 1619 | Non_Targeting_Human | AACGAGGCCCCCCCGAGGC |
| 1620 | Non_Targeting_Human | CTGGGTTGCCGCGAGGGTG |
| 1621 | Non_Targeting_Human | CCTCGCGCCCAGTATGGAG |
| 1622 | Non_Targeting_Human | TGGAGTAGTCGGCCCCGGG |
| 1623 | Non_Targeting_Human | GTTTACTCGGCAAACCACG |
| 1624 | Non_Targeting_Human | TCTACACCACCGTCTGTCG |
| 1625 | Non_Targeting_Human | TTAAACGCCGCTATTTTGA |
| 1626 | Non_Targeting_Human | CGATGACCACGATGCGCTG |
| 1627 | Non_Targeting_Human | ACGGGCGGGATCGTGGTAG |
| 1628 | Non_Targeting_Human | GGCCGACGAACCCACTAGT |
| 1629 | Non_Targeting_Human | CGCAGCGGGGTCCGAGTCA |
| 1630 | Non_Targeting_Human | AGATCCCAGGCCCGCGACA |
| 1631 | Non_Targeting_Human | CTACCGGTGTCCGAATCGC |
| 1632 | Non_Targeting_Human | GCCCCTGAGCCTAACTGGC |
| 1633 | Non_Targeting_Human | ACAACCCCAGTAAATCGGG |

| | | |
|---|---|---|
| 1634 | Non_Targeting_Human | GTACCCGCTAAGGAGCGCG |
| 1635 | Non_Targeting_Human | AGGTCCGGATCCCGTGACC |
| 1636 | Non_Targeting_Human | GGTCTGGGACGGCCCCGTT |
| 1637 | Non_Targeting_Human | AACGCGGCCATTACCCGAG |
| 1638 | Non_Targeting_Human | CACAGGGCTACGCATACGA |
| 1639 | Non_Targeting_Human | GTCTTTCTCCATTAGTCCA |
| 1640 | Non_Targeting_Human | CCAGGGCGATGAGCCAGCA |
| 1641 | Non_Targeting_Human | GTCGGTCAAAGCGCTGCGG |
| 1642 | Non_Targeting_Human | TCAGTATAAGAACCCTGAG |
| 1643 | Non_Targeting_Human | GGTCACACACCGCCGGAGG |
| 1644 | Non_Targeting_Human | CAGAAGAGTAAGAACTCGT |
| 1645 | Non_Targeting_Human | CATATAGGCTACCCTTAAG |
| 1646 | Non_Targeting_Human | CGAGGACCATGCATGGACA |
| 1647 | Non_Targeting_Human | AGATTCCGTAGGGACGCAC |
| 1648 | Non_Targeting_Human | ATGCACGAAGGGTGCAGTC |
| 1649 | Non_Targeting_Human | ACGCGAAGCCCGCAAAGAG |
| 1650 | Non_Targeting_Human | GAGCGTACGACAATGCGGC |
| 1651 | Non_Targeting_Human | CGCACGCGGACGGGCCTGC |
| 1652 | Non_Targeting_Human | TATGTACAGCAGCGCGTGA |
| 1653 | Non_Targeting_Human | GGCGCGTCCGCGAGCCTAG |
| 1654 | Non_Targeting_Human | TTGCGACTGCGGTCATCTG |
| 1655 | Non_Targeting_Human | TTGCGGAGAGTTATCGGCA |
| 1656 | Non_Targeting_Human | CGCCGGTCGTTTTGTCAGA |
| 1657 | Non_Targeting_Human | GCGAAAGGCGGTCCATACG |
| 1658 | Non_Targeting_Human | AGGACCAGTGCTTGGCCCG |
| 1659 | Non_Targeting_Human | CCTGCGCCTATACGCGGAG |
| 1660 | Non_Targeting_Human | TGACAATCCGGCTCACGCG |
| 1661 | Non_Targeting_Human | AGTTTGGTTCCAATGCTGG |
| 1662 | Non_Targeting_Human | CGCTAGGGACTCCGGTGGG |
| 1663 | Non_Targeting_Human | TAGGTATTCCGTGAAGGGG |
| 1664 | Non_Targeting_Human | TAAGAGCGGCCGACAATGA |
| 1665 | Non_Targeting_Human | CCACGACGGTTAAAACCGG |
| 1666 | Non_Targeting_Human | GTAGAGTTGTGCGTGCCTT |
| 1667 | Non_Targeting_Human | ATCTCTTTGCTGGGCCGCA |
| 1668 | Non_Targeting_Human | CTCGGCAGCTAGCTGAGAT |
| 1669 | Non_Targeting_Human | AATAGGTGCCAGTCGGCTA |
| 1670 | Non_Targeting_Human | GGGGATCTTAGTCCTTACG |
| 1671 | Non_Targeting_Human | ACTCGGCCCGATAGGCGAA |
| 1672 | Non_Targeting_Human | GATGTGACAAACTTTGTCA |
| 1673 | Non_Targeting_Human | TGACCGGGGGTGCGAGTGG |
| 1674 | Non_Targeting_Human | GTACCACACAGGGACGCAC |
| 1675 | Non_Targeting_Human | AGCATGGGGGGATCGCGCG |
| 1676 | Non_Targeting_Human | TACGCAGGTGAGGTGGTCA |

| | | |
|---|---|---|
| 1677 | Non_Targeting_Human | CGATCGACAGGCAGGAAAG |
| 1678 | Non_Targeting_Human | GGGATAGCGCGCAAGGAAG |
| 1679 | Non_Targeting_Human | CAAGCTCGGCGGAAAGGAC |
| 1680 | Non_Targeting_Human | GGTGGTCACGGGTCCCTGG |
| 1681 | Non_Targeting_Human | CTAAAGCCATCGGAGAAAC |
| 1682 | Non_Targeting_Human | CCCTACCAGTGCCACAAGG |
| 1683 | Non_Targeting_Human | AGCGGGCGAGCACCAGCAT |
| 1684 | Non_Targeting_Human | CCTGCTGGGCGGTTCGACA |
| 1685 | Non_Targeting_Human | CCTGCCCGCGAGACACAAA |
| 1686 | Non_Targeting_Human | CTCGAACGCCACCAAGTCG |
| 1687 | Non_Targeting_Human | GAACGGACAAACCTCAAAG |
| 1688 | Non_Targeting_Human | CGGACCCAGCTAGATAACT |
| 1689 | Non_Targeting_Human | GTGCCGAGGCGTAGACGGT |
| 1690 | Non_Targeting_Human | TCCCGACGGGTTAACCTAC |
| 1691 | Non_Targeting_Human | AGCGCAAACCGCCCTAGTA |
| 1692 | Non_Targeting_Human | CCGCCACGTAGCGGCTCAG |
| 1693 | Non_Targeting_Human | AGATGATGTAAGAAGGCGG |
| 1694 | Non_Targeting_Human | GGTTGCACAAAGTTGCGAT |
| 1695 | Non_Targeting_Human | GTGAGTGCGTCAATCACTG |
| 1696 | Non_Targeting_Human | CGCCGGGATGTTGTTCCCC |
| 1697 | Non_Targeting_Human | GGGCGGCGTCGCTTCACGG |
| 1698 | Non_Targeting_Human | GGGGCCGCAGTGTACGACG |
| 1699 | Non_Targeting_Human | AACAGCGCCGAGCCATGCC |
| 1700 | Non_Targeting_Human | CGCCCTTCCGTAGGTGTGC |
| 1701 | Non_Targeting_Human | AATCTGAGGAAGGTTTAGG |
| 1702 | Non_Targeting_Human | GAGTCGAGTCCGAGAACTA |
| 1703 | Non_Targeting_Human | GCCGCAGCGTAGATGGATG |
| 1704 | Non_Targeting_Human | CCGCGACGTCACAAATTGC |
| 1705 | Non_Targeting_Human | GGTCAGGGCAGACGCGAGA |
| 1706 | Non_Targeting_Human | GGCCAGGGCCCATCTCAAC |
| 1707 | Non_Targeting_Human | AATAACCGGAGCAAGTGCT |
| 1708 | Non_Targeting_Human | GCCGAAGGATCACTACACA |
| 1709 | Non_Targeting_Human | GCGGACGTGCTACTCTCCA |
| 1710 | Non_Targeting_Human | CCGTATGGGGTTCGTGTCC |
| 1711 | Non_Targeting_Human | CCGCGGGACGCCGGAAGGG |
| 1712 | Non_Targeting_Human | GCGATCTTCAATTCACGTG |
| 1713 | Non_Targeting_Human | CGGCGGTGGGCGATCGACA |
| 1714 | Non_Targeting_Human | TGTTGTGAGAGCATCCGGA |
| 1715 | Non_Targeting_Human | GAGCTCGCCGCGATCAGGG |
| 1716 | Non_Targeting_Human | GGCTGACGTGGATCCCAGC |
| 1717 | Non_Targeting_Human | GCGAGAGGAAGCATCGCTT |
| 1718 | Non_Targeting_Human | AGGGTGATGAAGGGCCGAG |
| 1719 | Non_Targeting_Human | AAAAAATACCCCTCCGCAG |

| | | |
|---|---|---|
| 1720 | Non_Targeting_Human | CAAAGGGGGAACTCAACGC |
| 1721 | Non_Targeting_Human | GGGAGTCAGCAAGCCGCGA |
| 1722 | Non_Targeting_Human | GTTGGAGCCGCTCGCTGCA |
| 1723 | Non_Targeting_Human | CTGCAGCATATTGTGCAGG |
| 1724 | Non_Targeting_Human | GCATTGGGCCCGACCTTTA |
| 1725 | Non_Targeting_Human | ACTGCGCGATGCTGTATCA |
| 1726 | Non_Targeting_Human | CGCCGGACGATGTCACTGG |
| 1727 | Non_Targeting_Human | GCGCTCTCAGCCAGGCAAC |
| 1728 | Non_Targeting_Human | GTCCGACGATCTGTGGTTG |
| 1729 | Non_Targeting_Human | TTCCGCCCTAGGCAGACTC |
| 1730 | Non_Targeting_Human | GCAGAACTAGAGGCGGTGG |
| 1731 | Non_Targeting_Human | GAGAAAGGTGCACTGACGA |
| 1732 | Non_Targeting_Human | ACCCGTCCGTCGGTGGCTC |
| 1733 | Non_Targeting_Human | GGGCGGGACGGTCCGTGGG |
| 1734 | Non_Targeting_Human | GCGCATGACTGACCCGCGA |
| 1735 | Non_Targeting_Human | GTGATATCCTACGGGATTA |
| 1736 | Non_Targeting_Human | CAAGTAGGTCCCGGGTCGA |
| 1737 | Non_Targeting_Human | CGACACGGACCGGACTCCG |
| 1738 | Non_Targeting_Human | CCAAAATTCGAGCAAGTAT |
| 1739 | Non_Targeting_Human | CGGAATTTAGGGCGTCTTA |
| 1740 | Non_Targeting_Human | CAGTCAACTCAAAAGACTA |
| 1741 | Non_Targeting_Human | CAACTGGTAGGACCACCGA |
| 1742 | Non_Targeting_Human | GCGAGTTCGACCCTAAAAG |
| 1743 | Non_Targeting_Human | CATATGACGAACAAAGATC |
| 1744 | Non_Targeting_Human | CCCCCCACGGCGCGTGGTG |
| 1745 | Non_Targeting_Human | GGCATCGGGCAGACGGGAT |
| 1746 | Non_Targeting_Human | TCGGAACCTGATACTGGCG |
| 1747 | Non_Targeting_Human | GGACGAGTAACCTGCCGGG |
| 1748 | Non_Targeting_Human | AGAATCAGGCGACAGAAGA |
| 1749 | Non_Targeting_Human | TTGCCTGTCGGACCGGGTG |
| 1750 | Non_Targeting_Human | GGCCCTTGAGGCGGGCCGA |
| 1751 | Non_Targeting_Human | AACGTTCGCGCGGTCACAT |
| 1752 | Non_Targeting_Human | TCATTCGCGAGCAGGCGCG |
| 1753 | Non_Targeting_Human | AACAGGGATAGCACTACAC |
| 1754 | Non_Targeting_Human | TTGTCGAGGGAGAAGTTAA |
| 1755 | Non_Targeting_Human | CGGGGGACGTCGACGAGAA |
| 1756 | Non_Targeting_Human | GTCGACCGGGTGCACACTG |
| 1757 | Non_Targeting_Human | GCCGTCTACGAATAGTAGG |
| 1758 | Non_Targeting_Human | CTCATCGCGGTCCTCTAAG |
| 1759 | Non_Targeting_Human | GATGCGAAGGAGGCATCGG |
| 1760 | Non_Targeting_Human | TCCTTGGCCGGGAGGAGCT |
| 1761 | Non_Targeting_Human | TCGACAACACACATATAAG |
| 1762 | Non_Targeting_Human | GATCGGGCTGCTGTTGCAG |

| | | |
|---|---|---|
| 1763 | Non_Targeting_Human | ACTTCGAGCGTGAGATGTG |
| 1764 | Non_Targeting_Human | CCAATCCCGAGGTTATGTG |
| 1765 | Non_Targeting_Human | TTGCCCCGGTTGGCATGCT |
| 1766 | Non_Targeting_Human | TTCCGCCTGGATAGTGCGG |
| 1767 | Non_Targeting_Human | CTCAAGGTAGACTTCTCCG |
| 1768 | Non_Targeting_Human | TGGGTCTGAGGCACGCAAC |
| 1769 | Non_Targeting_Human | AGCTTTCAGCCACTCGTGC |
| 1770 | Non_Targeting_Human | ATTAACGTGACGCTGCTTG |
| 1771 | Non_Targeting_Human | CCACGCTTGACCGCTCCGT |
| 1772 | Non_Targeting_Human | ACTGCTTCAGTCGCGTGGT |
| 1773 | Non_Targeting_Human | TCTGGGGGGCCTACAGCCA |
| 1774 | Non_Targeting_Human | GTCCTATAAGCACCCCCAG |
| 1775 | Non_Targeting_Human | GTCCCATCTGGACTCTGCG |
| 1776 | Non_Targeting_Human | GGCGACATCGGCCATGTGC |
| 1777 | Non_Targeting_Human | AAGACCACCGTAACGAGAG |
| 1778 | Non_Targeting_Human | CGGACGCGAATGAAAGTCA |
| 1779 | Non_Targeting_Human | AGGGAGGAATCGGAGACTG |
| 1780 | Non_Targeting_Human | GTTACCGGGGCGTCGTGTG |
| 1781 | Non_Targeting_Human | TCGTAGTCGGGCACTAGGG |
| 1782 | Non_Targeting_Human | GCCACAGCATAGATTGATT |
| 1783 | Non_Targeting_Human | GAGACAGGGACCTATACAA |
| 1784 | Non_Targeting_Human | GTGCCCGAGGTTGCGCCAG |
| 1785 | Non_Targeting_Human | GGAACCCAGGGTATAAGTA |
| 1786 | Non_Targeting_Human | ACCGCTGGGAGCACTGTAA |
| 1787 | Non_Targeting_Human | CTAACCGAGGCGGCATAGG |
| 1788 | Non_Targeting_Human | GGAACCAGGGTAGGATAAG |
| 1789 | Non_Targeting_Human | CGGGGTTGAAGGTCGGGGC |
| 1790 | Non_Targeting_Human | ATTCGTCTGCCTGCTGCAA |
| 1791 | Non_Targeting_Human | CATGCAGGGTTCCAGAAAA |
| 1792 | Non_Targeting_Human | CCCACTCTGGGCACGTAGG |
| 1793 | Non_Targeting_Human | CCACGGCAAAGCCCCCCTG |
| 1794 | Non_Targeting_Human | GTCGAGTAAGAGCAGGCTC |
| 1795 | Non_Targeting_Human | AGCGCGCTAGGGTCCCTGT |
| 1796 | Non_Targeting_Human | CGAGTACCGGTCCGCGGAT |
| 1797 | Non_Targeting_Human | ACTGCCTCCGTGGCGGCGG |
| 1798 | Non_Targeting_Human | TAGCTTGCACTCCCCTTGT |
| 1799 | Non_Targeting_Human | GACACCACCTAGGGGGACC |
| 1800 | Non_Targeting_Human | TGTGACCGAGAAGAAGCGG |
| 1801 | Non_Targeting_Human | CGCAGGTTGGGCGACCGAA |
| 1802 | Non_Targeting_Human | GGTACCGCAAGTGCGGCGT |
| 1803 | Non_Targeting_Human | ATAAGTGCACCCGCTATAA |
| 1804 | Non_Targeting_Human | GATTAGGGCGCACACGTGT |
| 1805 | Non_Targeting_Human | TAGGCCCAGCGAATAGGCC |

| | | |
|---|---|---|
| 1806 | Non_Targeting_Human | ATGGGTCTGTGGGGACCAT |
| 1807 | Non_Targeting_Human | GTCAAAGTCTCACACACGA |
| 1808 | Non_Targeting_Human | GTTGTGGCGCACCGCATGG |
| 1809 | Non_Targeting_Human | CTCCTAGGCGCGTTTTCCG |
| 1810 | Non_Targeting_Human | GTCGGTAAACCACACCCTC |
| 1811 | Non_Targeting_Human | CCGGGAATAAAGGATCGGC |
| 1812 | Non_Targeting_Human | GTCATTGTGACGACAAGGT |
| 1813 | Non_Targeting_Human | AGGCAAGCGAGGCCCGCCT |
| 1814 | Non_Targeting_Human | CGGAGCTTAGAATGGGATA |
| 1815 | Non_Targeting_Human | CTCATGCGCGTGCTCGTCC |
| 1816 | Non_Targeting_Human | TAGCTAAGTTATGCAGCGT |
| 1817 | Non_Targeting_Human | CCGCGACAGCGCAGAATGG |
| 1818 | Non_Targeting_Human | TTGTCGCGGAGTCGATGCC |
| 1819 | Non_Targeting_Human | GTTTCAGGATCATACAGGC |
| 1820 | Non_Targeting_Human | ACAAACGGGGATTTGCCGG |
| 1821 | Non_Targeting_Human | CGCTTCCGTGAGGACTCAT |
| 1822 | Non_Targeting_Human | CCGGGCACTAATGCAACTG |
| 1823 | Non_Targeting_Human | GGCGCTCCGCAGCCAGTCG |
| 1824 | Non_Targeting_Human | TTGGGGCTAGGATCACGTA |
| 1825 | Non_Targeting_Human | AGCCGGCCCGTGGGCGCTT |
| 1826 | Non_Targeting_Human | CCGTCCGCGGTGTGGCCAG |
| 1827 | Non_Targeting_Human | GCGGCGCCATAGTAGTTAG |
| 1828 | Non_Targeting_Human | GCAGTCAAGGTCAGAGGCG |
| 1829 | Non_Targeting_Human | CCCGGCCATGATCCCGCCC |
| 1830 | Non_Targeting_Human | GGGGATCATCCTTGACAGG |
| 1831 | Non_Targeting_Human | TCTCGTATTACTAGACGGG |
| 1832 | Non_Targeting_Human | CTACATGGCGTCCCTGACT |
| 1833 | Non_Targeting_Human | ACAGCGGGCACGAAGGCGT |
| 1834 | Non_Targeting_Human | AGTATTGTGAAATCCGCCG |
| 1835 | Non_Targeting_Human | ACGGACCGCGCCAAGGGAG |
| 1836 | Non_Targeting_Human | TATGGGCCGCTCTTCCCGA |
| 1837 | Non_Targeting_Human | CGAGGCAACGAGTGCTGTA |
| 1838 | Non_Targeting_Human | TCGGCAAAAAACACGGCAG |
| 1839 | Non_Targeting_Human | GGCGGGGGCAACAATCCTG |
| 1840 | Non_Targeting_Human | CGGACCTAAGGTCCCGCTT |
| 1841 | Non_Targeting_Human | ACCCGAACGCTAGTTACAG |
| 1842 | Non_Targeting_Human | GGCCCACCCCCGCAATCGG |
| 1843 | Non_Targeting_Human | CCGCCTCCAAATGCGCTGC |
| 1844 | Non_Targeting_Human | GGGCGGGATACGGCAAGTT |
| 1845 | Non_Targeting_Human | CCAATGCCGCCGCGGTCGT |
| 1846 | Non_Targeting_Human | ACCTCGCTGCTCCGACCGG |
| 1847 | Non_Targeting_Human | GGGCCGAGTAGAAATCGCC |
| 1848 | Non_Targeting_Human | CAGCTGGGGCGCAACGAGG |

| | | |
|---|---|---|
| 1849 | Non_Targeting_Human | GGGGACTTGCTCCAGGACG |
| 1850 | Non_Targeting_Human | CGGCGACGAATGGATCCGG |
| 1851 | Non_Targeting_Human | GACGGTGCGGCGTGAAGCT |
| 1852 | Non_Targeting_Human | CGAACGATCCGGGAACCAG |
| 1853 | Non_Targeting_Human | CCAAATGTTCTGCACCTTA |
| 1854 | Non_Targeting_Human | AGAAGGAGCGAGGCTAAAG |
| 1855 | Non_Targeting_Human | CGCCTCTCAGGGCGGATAG |
| 1856 | Non_Targeting_Human | AGAAGCCAGTGGACAGCGC |
| 1857 | Non_Targeting_Human | CCCCCCGTGAGTCCATCCA |
| 1858 | Non_Targeting_Human | GGGAGGACGGCACCAAGTC |
| 1859 | Non_Targeting_Human | GCCCGCCGACAAAGGTTGG |
| 1860 | Non_Targeting_Human | GCCCGGGGAGCCTAGCTTG |
| 1861 | Non_Targeting_Human | AAGGCGCCTCGTATAAGGC |
| 1862 | Non_Targeting_Human | GACACGGGCGAAGGGATAA |
| 1863 | Non_Targeting_Human | CATCGGAATACACGCACCG |
| 1864 | Non_Targeting_Human | GACACACCGCCCCAAAATC |
| 1865 | Non_Targeting_Human | TTCGGCGTGACGCACACCG |
| 1866 | Non_Targeting_Human | TGTTCACACGGTGCAGGGG |
| 1867 | Non_Targeting_Human | TCAGGCCCGGTGCGCCCGA |
| 1868 | Non_Targeting_Human | TGGCACCGGGGCTCGTCAG |
| 1869 | Non_Targeting_Human | GGGATTTCGGATAGCGCGG |
| 1870 | Non_Targeting_Human | GCGGTCGACAGGAGATCCC |
| 1871 | Non_Targeting_Human | TACAGCTGGAAGGAAAGCC |
| 1872 | Non_Targeting_Human | GGGCGAAGCGCCACATCAG |
| 1873 | Non_Targeting_Human | AGCAGCGCGCGATGTACGG |
| 1874 | Non_Targeting_Human | GTCGCGGCGGGACATTCAC |
| 1875 | Non_Targeting_Human | GAGGGGCAGCCAGATAAGC |
| 1876 | Non_Targeting_Human | CGGCCAAAGGTGGACGTGA |
| 1877 | Non_Targeting_Human | GGATCGGGACTTCGCACGG |
| 1878 | Non_Targeting_Human | TATACCCCTGAAGAGACGG |
| 1879 | Non_Targeting_Human | CCGTAAGCGTGCAAAAATT |
| 1880 | Non_Targeting_Human | ACAGTAGAAGTCTCCGCGG |
| 1881 | Non_Targeting_Human | GGGCGGGAGGCACGAAAGC |
| 1882 | Non_Targeting_Human | TCGGCCGCCCAGGACTAAC |
| 1883 | Non_Targeting_Human | CTAATGTAAGAGGCCTACG |
| 1884 | Non_Targeting_Human | CGGAGGGGGGCCGTCAGTA |
| 1885 | Non_Targeting_Human | CCCACGGCGTCTGATTATC |
| 1886 | Non_Targeting_Human | GATTTGAGCCCACCGTAGG |
| 1887 | Non_Targeting_Human | GATTCTCGCCCCCGAAGAT |
| 1888 | Non_Targeting_Human | TAGTAGCGCAGCGACCCCG |
| 1889 | Non_Targeting_Human | TACTCATGGTCCCGGATCG |
| 1890 | Non_Targeting_Human | CTACTGTGCTCGGGAACGG |
| 1891 | Non_Targeting_Human | CGCCGAGGGTTCTATGCTG |

| | | |
|---|---|---|
| 1892 | Non_Targeting_Human | CCTGCGGGCCGAAGGCGCA |
| 1893 | Non_Targeting_Human | CCACGCGCCTCCGATAGGG |
| 1894 | Non_Targeting_Human | TATTGCTCCAGCCACAAAA |
| 1895 | Non_Targeting_Human | TGCGGTACCGCGGCCGCTC |
| 1896 | Non_Targeting_Human | GCCCCGAAAAATGCGCTGA |
| 1897 | Non_Targeting_Human | TCTGGATATAACGATGGCG |
| 1898 | Non_Targeting_Human | ATCCCTGCCACGGGCGGAA |
| 1899 | Non_Targeting_Human | AACGCTGTCTGCGGGCCAG |
| 1900 | Non_Targeting_Human | TCTGGACGGGGCGATGCGC |
| 1901 | Non_Targeting_Human | CGAAACCGAATCCTCTCAG |
| 1902 | Non_Targeting_Human | CGCGGGGCAAAAGGGCAGG |
| 1903 | Non_Targeting_Human | CGTGGGATGAGTAACACAG |
| 1904 | Non_Targeting_Human | CGAAGCCCACGCGGAGCCT |
| 1905 | Non_Targeting_Human | AACGGTGGTGCTTGTGTTT |
| 1906 | Non_Targeting_Human | CTGCGCCGTGTGTAAGGCG |
| 1907 | Non_Targeting_Human | AAGAGGACGGGCAAGTCTG |
| 1908 | Non_Targeting_Human | AGCGTGGCGTCGCAGTTGG |
| 1909 | Non_Targeting_Human | GTGTGCGGACCGGTCAGGC |
| 1910 | Non_Targeting_Human | CGCGGGCCCTATCTCCAGG |
| 1911 | Non_Targeting_Human | GTTGTGCGGGCAGATGAAA |
| 1912 | Non_Targeting_Human | CTGCACGAGTGGAATTGAC |
| 1913 | Non_Targeting_Human | CCTCTGCCGGTCATCGCCG |
| 1914 | Non_Targeting_Human | TGGCGGTCACTTCTTTATA |
| 1915 | Non_Targeting_Human | AGCCGGCAGGTCGCTTCCA |
| 1916 | Non_Targeting_Human | CTCAGAAAGTCGAAACGTC |
| 1917 | Non_Targeting_Human | TTCCTGCGCAATTCCGGCG |
| 1918 | Non_Targeting_Human | ACCGAGCCAGGGGTCGCAA |
| 1919 | Non_Targeting_Human | GGACGGGAATACAGCGCCC |
| 1920 | Non_Targeting_Human | GCTGGCACGAACGAGAGAC |
| 1921 | Non_Targeting_Human | GGCCTACGGAAGCGCGGTT |
| 1922 | Non_Targeting_Human | ATAGGTGCCACAGCTCGCA |
| 1923 | Non_Targeting_Human | AAGCCCTAGTGACGCAGCA |
| 1924 | Non_Targeting_Human | CGCATGGTATTGGCTTCCG |
| 1925 | Non_Targeting_Human | ATGCCCCCCCACTAAGAGG |
| 1926 | Non_Targeting_Human | GGACTCACGGTGTCATCGG |
| 1927 | Non_Targeting_Human | CGCGGCCGTGCTGCAACGC |
| 1928 | Non_Targeting_Human | CCAACGCTAGATATAAACA |
| 1929 | Non_Targeting_Human | AACCCGCCGGCCCCGTTCG |
| 1930 | Non_Targeting_Human | CCAATATCGCACGCGTCGC |
| 1931 | Non_Targeting_Human | CCTGGCGGAGCAATTCTCC |
| 1932 | Non_Targeting_Human | CGACGCGCCGCCCGGCGTT |
| 1933 | Non_Targeting_Human | ACCGGCACAGTAAACATAC |
| 1934 | Non_Targeting_Human | CTTCTCCGGGCCACCATCC |

| | | |
|---|---|---|
| 1935 | Non_Targeting_Human | GAGCTTTCTTAGCACGGAG |
| 1936 | Non_Targeting_Human | GGCACCGCCGTTAAAAGTT |
| 1937 | Non_Targeting_Human | CCGAACGATGACCATTCTG |
| 1938 | Non_Targeting_Human | GTCGCCCCGCGAAGTTCAA |
| 1939 | Non_Targeting_Human | CGCTCAAACCTGCCTACCA |
| 1940 | Non_Targeting_Human | CATTCTAGAAATCGTAGTA |
| 1941 | Non_Targeting_Human | GAGCGGTATGGGTAACCGA |
| 1942 | Non_Targeting_Human | GGCTGCCTGCGGAAACCAC |
| 1943 | Non_Targeting_Human | CCTATGTAGCGGTCTGAAG |
| 1944 | Non_Targeting_Human | GACCTGCGCGCGCCCGTTT |
| 1945 | Non_Targeting_Human | AAAAGCGGCGCAGTATTTG |
| 1946 | Non_Targeting_Human | CTCGCAAGCTAAGAAACCG |
| 1947 | Non_Targeting_Human | CGTCGGGCTGGAATAGAGG |
| 1948 | Non_Targeting_Human | TGGGGAACTGATGAAGAAT |
| 1949 | Non_Targeting_Human | CGCTGCCAGACCCAATCGG |
| 1950 | Non_Targeting_Human | CTACCCGGGTAGTGGCGAG |
| 1951 | Non_Targeting_Human | TTATTGGGTCTGCCACCGT |
| 1952 | Non_Targeting_Human | AGGGACGTCGAACGATGCC |
| 1953 | Non_Targeting_Human | TCGGGACTGACTGCTTAGG |
| 1954 | Non_Targeting_Human | GGGCTTGCGGGGGAGCCGC |
| 1955 | Non_Targeting_Human | GCGGCCCTAGCAACAACAG |
| 1956 | Non_Targeting_Human | GATACTCAGCTGTTTAGGG |
| 1957 | Non_Targeting_Human | AAGAGCAGGGCCGAAGTAG |
| 1958 | Non_Targeting_Human | GACAGAAACGACTCCGGAA |
| 1959 | Non_Targeting_Human | CCCAGCGCCAGGCTTAAAG |
| 1960 | Non_Targeting_Human | ATGCGAGGGACGGAGTGGA |
| 1961 | Non_Targeting_Human | CAGGCATCGCGACCAGTGA |
| 1962 | Non_Targeting_Human | CCAGTCGGGCCAGCTTGGC |
| 1963 | Non_Targeting_Human | CGGCAAGGCGTGACAGGTC |
| 1964 | Non_Targeting_Human | TGGCGTGGAGGCTACTAAT |
| 1965 | Non_Targeting_Human | GGGTACCTAAGGGTGGATA |
| 1966 | Non_Targeting_Human | AACCTGACGGTGCAATGTT |
| 1967 | Non_Targeting_Human | TAGACGGAATAAGCAACTG |
| 1968 | Non_Targeting_Human | CAATGCCGCATCTTCAGAG |
| 1969 | Non_Targeting_Human | TAACCCGCCGCGGGAATGC |
| 1970 | Non_Targeting_Human | TTACCCCGGTGGCCGGAAG |
| 1971 | Non_Targeting_Human | GGTAGCTAGACTCTCGCCC |
| 1972 | Non_Targeting_Human | CGGCAGCAAGAATGCGTTA |
| 1973 | Non_Targeting_Human | ACTCCACGGACGTAGTGGT |
| 1974 | Non_Targeting_Human | GAACGCCTCCCGTTAGCGG |
| 1975 | Non_Targeting_Human | CAAGGCACCGCATGCGATT |
| 1976 | Non_Targeting_Human | TTGATCTGGGCGGAGCTTC |
| 1977 | Non_Targeting_Human | GCTTGGTCGTTGAAAACAT |

| | | |
|---|---|---|
| 1978 | Non_Targeting_Human | ATTCGGGGCGCGGGGAGAA |
| 1979 | Non_Targeting_Human | GGACAGGGCGTCAGAGAAG |
| 1980 | Non_Targeting_Human | AACCCCGTCGCAACCTAAG |
| 1981 | Non_Targeting_Human | AGAATTGGGGAAGTACCCG |
| 1982 | Non_Targeting_Human | AACGCGGGCAACCAGCTTG |
| 1983 | Non_Targeting_Human | GTCGCCGATGAGTTCCGTG |
| 1984 | Non_Targeting_Human | GCCTCGGTATTCGGGAAGG |
| 1985 | Non_Targeting_Human | CAATTACCAGCGGGACTTA |
| 1986 | Non_Targeting_Human | TCGAGCCCCCAAACACCCG |
| 1987 | Non_Targeting_Human | TGATGGAGCGTCTACGTAG |
| 1988 | Non_Targeting_Human | GAGCCATGACGGCGACGGA |
| 1989 | Non_Targeting_Human | CGAATGAGATGGGAAAAGG |
| 1990 | Non_Targeting_Human | ACCCGCCGCAACGCGTGCG |
| 1991 | Non_Targeting_Human | GGCCGGGGCGAACTAGCAC |
| 1992 | Non_Targeting_Human | CGGGAAGTATATGACACCG |
| 1993 | Non_Targeting_Human | CGATGCCGATACGCCAAGG |
| 1994 | Non_Targeting_Human | GCTGCTCTGCCTAGCAAGT |
| 1995 | Non_Targeting_Human | CCAGGATCAAGCCGTGACA |
| 1996 | Non_Targeting_Human | GCGCAGCCTCGTTGTTTTA |
| 1997 | Non_Targeting_Human | CGAAGTGTACAGGGACCTT |
| 1998 | Non_Targeting_Human | TAGCAGGGGGGTACTTCAG |
| 1999 | Non_Targeting_Human | TGGAGTCCGCCTCTGTCGG |
| 2000 | Non_Targeting_Human | GAGGCGGACGCTAGAGTCC |
| 2001 | Non_Targeting_Human | AAACGATATGAGCGCACTG |
| 2002 | Non_Targeting_Human | CTGTTAAAACGACCTTAAC |
| 2003 | Non_Targeting_Human | GCGCCCGTTGAGAGCGCAA |
| 2004 | Non_Targeting_Human | GGGAACTTCAACGAGACGG |
| 2005 | Non_Targeting_Human | TGCGATAGGCCGCAGTCAG |
| 2006 | Non_Targeting_Human | CACGGAGTGCCGAATGTGC |
| 2007 | Non_Targeting_Human | GAACGACTTAGTAGACCTA |
| 2008 | Non_Targeting_Human | TGCCAGGGCAGATGCGCAA |
| 2009 | Non_Targeting_Human | CGTGAAGTGAGCCAACAGG |
| 2010 | Non_Targeting_Human | TCTCCGTGGGAGACGATCA |
| 2011 | Non_Targeting_Human | GTTGGGTCAAGCGAGCCCG |
| 2012 | Non_Targeting_Human | CTACTTACTGCGTACGCCT |
| 2013 | Non_Targeting_Human | CTGCCGTGCCGTGTGATAA |
| 2014 | Non_Targeting_Human | GCGGAGCCATATTCGTTAT |
| 2015 | Non_Targeting_Human | TCGGAGGCTAATCGCAACT |
| 2016 | Non_Targeting_Human | GCGGAGGTTAGTCATTGAG |
| 2017 | Non_Targeting_Human | ATGCAAGGTCCAACTTCGG |
| 2018 | Non_Targeting_Human | TATGAGATCACGAGCTTAT |
| 2019 | Non_Targeting_Human | CACCTATGTTAATTCCTAA |
| 2020 | Non_Targeting_Human | TGGCTGAGAACGCGCAGCC |

| | | |
|---|---|---|
| 2021 | Non_Targeting_Human | TCCAAGTATCCAGCCTTGG |
| 2022 | Non_Targeting_Human | AATGCCGCTCGCAGAGCGG |
| 2023 | Non_Targeting_Human | GGACACTGACCAATTGTGC |
| 2024 | Non_Targeting_Human | CGACTCTCGGGGGGATAAG |
| 2025 | Non_Targeting_Human | CCCGGGGGCGAGATGAGCT |
| 2026 | Non_Targeting_Human | GCAGGGGTGCTTACTGCGG |
| 2027 | Non_Targeting_Human | GCCGTGCGCCGTACAGGTT |
| 2028 | Non_Targeting_Human | GGGAGCACGTCACCCTTCG |
| 2029 | Non_Targeting_Human | ATGGTCACTCTGACGATCA |
| 2030 | Non_Targeting_Human | CGCCGGCCAATGCGCGAAC |
| 2031 | Non_Targeting_Human | GGGAAAGGGCGCAAACTGG |
| 2032 | Non_Targeting_Human | ACATCCTAGTTTGACAAAA |
| 2033 | Non_Targeting_Human | TTGGGAGTAGGCGGGACTT |
| 2034 | Non_Targeting_Human | CTACGGGGGCGCCAAATTT |
| 2035 | Non_Targeting_Human | CAGTGCAGGATTAAGACCG |
| 2036 | Non_Targeting_Human | GCAACGACGCGCGTCTGAG |
| 2037 | Non_Targeting_Human | GCACGCCCAGCAGCAGCGA |
| 2038 | Non_Targeting_Human | CCCGGATGACGGCGATGCT |
| 2039 | Non_Targeting_Human | TGGGAAGTCAAATGCGATT |
| 2040 | Non_Targeting_Human | GCTTTCCCGAAATGTAACG |
| 2041 | Non_Targeting_Human | GGAGTAGAGTAGAACTGCT |
| 2042 | Non_Targeting_Human | CTTAAAGTGCGCTCCCGCG |
| 2043 | Non_Targeting_Human | TGCAAAGCAGAGCCTAATC |
| 2044 | Non_Targeting_Human | CAACCGGACACCTAAAGCC |
| 2045 | Non_Targeting_Human | CTTGACACGACGCAAGCCA |
| 2046 | Non_Targeting_Human | ATAGGGGATGAGCAGGTTG |
| 2047 | Non_Targeting_Human | TCGTGGGGACTCTTAATGG |
| 2048 | Non_Targeting_Human | CCTCAACTTGCGCCAGGAT |
| 2049 | Non_Targeting_Human | CGCCCGTGACGCAGTAAAC |
| 2050 | Non_Targeting_Human | CCGACGCGAGAGGCACAGA |
| 2051 | Non_Targeting_Human | AGGGTAGGTTTCGGAATAC |
| 2052 | Non_Targeting_Human | GGTCGCACTGGGAACAACA |
| 2053 | Non_Targeting_Human | TCTGTGCCGACGCCGGTAG |
| 2054 | Non_Targeting_Human | GCCAGGCACTCGCATAGAA |
| 2055 | Non_Targeting_Human | GTGCTTTCAGGTCGCGGAT |
| 2056 | Non_Targeting_Human | GGGGAGCGACACTAAGTAC |
| 2057 | Non_Targeting_Human | GGCGACGGGGTGTCTGGGG |
| 2058 | Non_Targeting_Human | GCGGTGGCGCAGAATACAG |
| 2059 | Non_Targeting_Human | CCCCGCTGGCTCGGCGTCT |
| 2060 | Non_Targeting_Human | TCGAGCCGCGGCGTGCGCG |
| 2061 | Non_Targeting_Human | CAAAGGCTGAGGAGCGGTG |
| 2062 | Non_Targeting_Human | GAGGGGTCTTTGGCGTAAA |
| 2063 | Non_Targeting_Human | TGGCGTCGGACCCAGGGAT |

| | | |
|---|---|---|
| 2064 | Non_Targeting_Human | CCTCCCAGTGCGCAGCGGA |
| 2065 | Non_Targeting_Human | ATCAAAATCGACGCGTCTA |
| 2066 | Non_Targeting_Human | AATTCGGCGGCGTGGCCGT |
| 2067 | Non_Targeting_Human | CGGGGGCCACGGCGGTAAG |
| 2068 | Non_Targeting_Human | CCCCCCTCCTGCCTTAATA |
| 2069 | Non_Targeting_Human | CGGGGGAGTGGCCTCGGAT |
| 2070 | Non_Targeting_Human | CCTATGCCGGTCAATATTG |
| 2071 | Non_Targeting_Human | CGAGGGTTGCTGTAGAGAG |
| 2072 | Non_Targeting_Human | ATGCGCGGATCGTAGGCGA |
| 2073 | Non_Targeting_Human | ACTAGAGCCGGGCTAACCG |
| 2074 | Non_Targeting_Human | CGTCCGTCGAAATTATCCT |
| 2075 | Non_Targeting_Human | GAAGCGCCAACCTCACGTC |
| 2076 | Non_Targeting_Human | TCGCCGCGTGCGATATCAT |
| 2077 | Non_Targeting_Human | TGCCATGCCGGGTATACTT |
| 2078 | Non_Targeting_Human | AGTTCGTCCAGCGCCGCTG |
| 2079 | Non_Targeting_Human | GAGGTGCCGTTGTCGTTGC |
| 2080 | Non_Targeting_Human | CCATAGCGACCGGACCACG |
| 2081 | Non_Targeting_Human | ACATGTGACTGCCCACGAG |
| 2082 | Non_Targeting_Human | ACACTGCGAGGCGACCCGG |
| 2083 | Non_Targeting_Human | CGAGGGCTAGTGCGAATCC |
| 2084 | Non_Targeting_Human | GGACCCCATAGCGTGAGCT |
| 2085 | Non_Targeting_Human | GCTCGACGCCGCAAGAATG |
| 2086 | Non_Targeting_Human | GTCCACCATCAATACTAAA |
| 2087 | Non_Targeting_Human | CGCCCCGACTCGGAAATAA |
| 2088 | Non_Targeting_Human | AGCTAAAATCGATTTGGCC |
| 2089 | Non_Targeting_Human | AGGCAAGTCCTCGGCGGAA |
| 2090 | Non_Targeting_Human | GCCCTGTGGCACACGCTGG |
| 2091 | Non_Targeting_Human | GAACTCGTAACGGACTGGG |
| 2092 | Non_Targeting_Human | GCCGCACGATAGTGTCCTA |
| 2093 | Non_Targeting_Human | GTTCGAGCATGCATCGCTA |
| 2094 | Non_Targeting_Human | GTGTGTCCTGGGGCGATAC |
| 2095 | Non_Targeting_Human | GGGGAGGGTTCCGCGAAGC |
| 2096 | Non_Targeting_Human | GCCCTCGAGTGGTGCCCAA |
| 2097 | Non_Targeting_Human | ACGGTACTTGCTGCAGCAT |
| 2098 | Non_Targeting_Human | CCACGGTGCGAGGACGATA |
| 2099 | Non_Targeting_Human | TCAAGTGGAGAAAATACGA |
| 2100 | Non_Targeting_Human | TTCGCGGCGGACCGAGCGG |
| 2101 | Non_Targeting_Human | TGGAGTGGTATGCGTAGTA |
| 2102 | Non_Targeting_Human | AGCGAAAGGTCGGAGGCAA |
| 2103 | Non_Targeting_Human | CACCCCGGTTGCACGACCC |
| 2104 | Non_Targeting_Human | GCTAGGTTAAGAGGGCTGA |
| 2105 | Non_Targeting_Human | GTCAGGGGGGTACCTAGAA |
| 2106 | Non_Targeting_Human | CCGTTACTGAGGCAGCACG |

| | | |
|---|---|---|
| 2107 | Non_Targeting_Human | GGAGCGGGCTTCCCTACGT |
| 2108 | Non_Targeting_Human | GCCCATGCTTGGTCGCATG |
| 2109 | Non_Targeting_Human | AGTTGACACAGCGCGCGGG |
| 2110 | Non_Targeting_Human | TCACGAGCTGGAGCCCAAG |
| 2111 | Non_Targeting_Human | TGGAAATCCACCCGTTCGC |
| 2112 | Non_Targeting_Human | AGCGTCCGTCCGCCTGGAT |
| 2113 | Non_Targeting_Human | GGGATGACGTCGGTACTTA |
| 2114 | Non_Targeting_Human | GTTATAAGCGGCAATACTG |
| 2115 | Non_Targeting_Human | CAGTCGCGGCGCACGTAAA |
| 2116 | Non_Targeting_Human | CCCGGCCCCGAGCACCGAG |
| 2117 | Non_Targeting_Human | ACACAACAGCACGGGTCCG |
| 2118 | Non_Targeting_Human | TAGGTGCAAGAGCTGACAT |
| 2119 | Non_Targeting_Human | TTCTGTCCCATGACGGCGC |
| 2120 | Non_Targeting_Human | AGTCCGACTCACCATTAGG |
| 2121 | Non_Targeting_Human | GCCCGAAGAGCGTCGGAAG |
| 2122 | Non_Targeting_Human | CTACCACATCACGTGAGCG |
| 2123 | Non_Targeting_Human | GGCATGCCGAAATCAAAAG |
| 2124 | Non_Targeting_Human | GTAGGCATGGCGGCCGACT |
| 2125 | Non_Targeting_Human | GCCTGAAGCCCTAAGCCTA |
| 2126 | Non_Targeting_Human | AAGCGAATGGTTATGAATG |
| 2127 | Non_Targeting_Human | AGGACTACCGGTCTGCCAG |
| 2128 | Non_Targeting_Human | GTGCGGCGCGGCGAGGCAC |
| 2129 | Non_Targeting_Human | GAGGCGCGATAAAGTCCGA |
| 2130 | Non_Targeting_Human | AGCGGCGGTAGCTCCAAGC |
| 2131 | Non_Targeting_Human | GTCCGGCGGCTTACCAGGT |
| 2132 | Non_Targeting_Human | AGCGCGCTAGGGGCCTTGT |
| 2133 | Non_Targeting_Human | GGTAGGCTCGCACTGCCAG |
| 2134 | Non_Targeting_Human | ACCTTGACATCCGGGTCCA |
| 2135 | Non_Targeting_Human | ACAGTCTACGGGCAGACTA |
| 2136 | Non_Targeting_Human | GCCCTTCTAGCCGAGCATA |
| 2137 | Non_Targeting_Human | CCGACATGGGGCCCCCAAC |
| 2138 | Non_Targeting_Human | CCTGGTGTTAGGCCACCGG |
| 2139 | Non_Targeting_Human | CTGGGATTAAGGCCTGTAG |
| 2140 | Non_Targeting_Human | GCTCGGCAACGGGGTCTGA |
| 2141 | Non_Targeting_Human | CCACTGTGACGCCGTGTCC |
| 2142 | Non_Targeting_Human | TCTTGGTGGGGAACTACGG |
| 2143 | Non_Targeting_Human | GTTAAACGAAGCCACCCGC |
| 2144 | Non_Targeting_Human | GCCCGCCTAACGCAGAACG |
| 2145 | Non_Targeting_Human | GCTTCATGCGCCGACCTGG |
| 2146 | Non_Targeting_Human | CAGGCAGCGGGGTAGCGCT |
| 2147 | Non_Targeting_Human | TTCCCCTGCCGCAACCGGG |
| 2148 | Non_Targeting_Human | CGGCGCGGGGACCAAGCAC |
| 2149 | Non_Targeting_Human | CGACAGGGACGGCGGGTCA |

| | | |
|---|---|---|
| 2150 | Non_Targeting_Human | GGACGCGCCGAGCGCACCT |
| 2151 | Non_Targeting_Human | GCTGGAGGAGCGGCACCGC |
| 2152 | Non_Targeting_Human | ATGCAGACGCTGCGTCGGG |
| 2153 | Non_Targeting_Human | CCGCCACTACAGCTATACG |
| 2154 | Non_Targeting_Human | GGGTACGGCGCGCCAACCT |
| 2155 | Non_Targeting_Human | GCTAATCGGTGACAGTCAC |
| 2156 | Non_Targeting_Human | GGACACCTTCACATAAGGA |
| 2157 | Non_Targeting_Human | GTTCGCAACCGACAGAGCG |
| 2158 | Non_Targeting_Human | CGGGGCCAGCGGTTTCCAA |
| 2159 | Non_Targeting_Human | CAGTGCCCACGACGTCACG |
| 2160 | Non_Targeting_Human | AGGCAACCGCTCTAGCGCG |
| 2161 | Non_Targeting_Human | ACGCGAAGTCGCGCACTCG |
| 2162 | Non_Targeting_Human | GGAGGTGAGCGCTCCGCTT |
| 2163 | Non_Targeting_Human | GCAGTAGGACGTTAGGAGT |
| 2164 | Non_Targeting_Human | AACTGCTAGAAGTTCGTCC |
| 2165 | Non_Targeting_Human | GGTCCGGTGTCAGAGAGCG |
| 2166 | Non_Targeting_Human | GCAGTGGCGCAGGCTCTTG |
| 2167 | Non_Targeting_Human | GTGTGCCGTGGCTGGAAAA |
| 2168 | Non_Targeting_Human | GGACCGAGTAAGGGCATCG |
| 2169 | Non_Targeting_Human | CCGCGCGGAGATCCGGCGC |
| 2170 | Non_Targeting_Human | TGGTCGCGCTTGTCCAAAG |
| 2171 | Non_Targeting_Human | CTGTATGATAGTAACTGGA |
| 2172 | Non_Targeting_Human | GAGGATCAGAACAATCGTT |
| 2173 | Non_Targeting_Human | GAGCGGCTAGCGCATATGC |
| 2174 | Non_Targeting_Human | ACCGGTGCGCCGTCGCGTG |
| 2175 | Non_Targeting_Human | GGGGAGGCCTCGGCGCTAG |
| 2176 | Non_Targeting_Human | TCCCCACCGCAGTCGTCTA |
| 2177 | Non_Targeting_Human | TGGCTGCGCAACATCGGAA |
| 2178 | Non_Targeting_Human | AGGGGTCTTTTGTAACATG |
| 2179 | Non_Targeting_Human | GGCGAGGCTAACAACATCT |
| 2180 | Non_Targeting_Human | CGCTAACGAGGAGCTTCGG |
| 2181 | Non_Targeting_Human | CCTCTCTCCCGCCCGCCAG |
| 2182 | Non_Targeting_Human | AAGCGAGTAGGATTACTTA |
| 2183 | Non_Targeting_Human | AGCACCTCACGCGGAAGAA |
| 2184 | Non_Targeting_Human | GCCAAACGCGCTAAAACGA |
| 2185 | Non_Targeting_Human | GCCTGTCAGAGCACCCACG |
| 2186 | Non_Targeting_Human | AACCCGGTCAAATTGATGA |
| 2187 | Non_Targeting_Human | GGCGGAGGCGTAAGTGGTT |
| 2188 | Non_Targeting_Human | AGACGGAAAGCACTAGGTC |
| 2189 | Non_Targeting_Human | GCGGCGCCTGTAAGCCCCT |
| 2190 | Non_Targeting_Human | GGCGTGCTTGACAACGGTG |
| 2191 | Non_Targeting_Human | GGGTGGGACACTGCTCTGC |
| 2192 | Non_Targeting_Human | GGGTTTAGCGGATGGACCT |

| | | |
|---|---|---|
| 2193 | Non_Targeting_Human | TGGTGGTGATAATGACATA |
| 2194 | Non_Targeting_Human | CCATCTGATCTTAACGTGG |
| 2195 | Non_Targeting_Human | GAATACCTTAGGCCGGCAC |
| 2196 | Non_Targeting_Human | GCGCCAATACAGAAACCAC |
| 2197 | Non_Targeting_Human | AGCGGCCTCAGCCGCACTA |
| 2198 | Non_Targeting_Human | GACGGTGCGTGCGGTGTGC |
| 2199 | Non_Targeting_Human | ACGGGGCGGGGCATCTGAC |
| 2200 | Non_Targeting_Human | CATCGCTCCGGCAACTGGA |
| 2201 | Non_Targeting_Human | GACGTCACGTGACCTCAAT |
| 2202 | Non_Targeting_Human | CAGCGACGTGGCCGCCCAG |
| 2203 | Non_Targeting_Human | GCAATACCACGGCTATCCG |
| 2204 | Non_Targeting_Human | GCTCTCCCGAAGGAGTGCT |
| 2205 | Non_Targeting_Human | TGGCCTTAAAACGCTAGTG |
| 2206 | Non_Targeting_Human | CACAGACGCGGAATCCAAA |
| 2207 | Non_Targeting_Human | CCGCCGGGTATCGACGAAA |
| 2208 | Non_Targeting_Human | CAAGCCCGCATACGAGCAA |
| 2209 | Non_Targeting_Human | GTTTAGTGCGGCGGACCGT |
| 2210 | Non_Targeting_Human | AGCATCTAAGGCGCGGTAC |
| 2211 | Non_Targeting_Human | CGCCGAACGGAGTTTCCCG |
| 2212 | Non_Targeting_Human | CGACACTCCCGTCGTACCT |
| 2213 | Non_Targeting_Human | CCCCTGGCGGATACCCCAG |
| 2214 | Non_Targeting_Human | GAGCGGTGATAGCTTAATG |
| 2215 | Non_Targeting_Human | GCGGTGACGTAGTCTAAAG |
| 2216 | Non_Targeting_Human | ATCTTCGATTTACCTGGCC |
| 2217 | Non_Targeting_Human | CCAGGAAACGGTTTGGGCG |
| 2218 | Non_Targeting_Human | GGGTATACGGCGCTGCGAG |
| 2219 | Non_Targeting_Human | TGCGAGGGGCCCCCCTAAT |
| 2220 | Non_Targeting_Human | ACCGTCTGGGTGGAGGGCG |
| 2221 | Non_Targeting_Human | GTAAGCGCCTAACAATGGA |
| 2222 | Non_Targeting_Human | GCTGGGGTTACGGACCGGA |
| 2223 | Non_Targeting_Human | GGAGGGTCTACCCTACCAA |
| 2224 | Non_Targeting_Human | ACGGCGAGATCAATTGAGT |
| 2225 | Non_Targeting_Human | TCGTCTCCTGCGGTACCAA |
| 2226 | Non_Targeting_Human | GGCAGAAGCTTACAATCCG |
| 2227 | Non_Targeting_Human | AGCGTTTCTTGGCACCGCT |
| 2228 | Non_Targeting_Human | GCGGGCGCAAGACGTGGCA |
| 2229 | Non_Targeting_Human | TCGGATCCTCGGAGTGCGG |
| 2230 | Non_Targeting_Human | CGCGTAGTAGCTGCAACAG |
| 2231 | Non_Targeting_Human | GCCGGGTGTCTCGGGGCCT |
| 2232 | Non_Targeting_Human | GGTCCGGAGGTTATACCAA |
| 2233 | Non_Targeting_Human | CGAAGCCCTTGCCGTTCAA |
| 2234 | Non_Targeting_Human | GCGACGAAGATGGTGCGCC |
| 2235 | Non_Targeting_Human | CCACCTTCGAAGTCCGTAT |

| | | |
|---|---|---|
| 2236 | Non_Targeting_Human | AGGGCCCGCGCCGAAGGGC |
| 2237 | Non_Targeting_Human | TGTGGAGGCAGACTTCGCA |
| 2238 | Non_Targeting_Human | GCAGCCTGATGCGTTAGAG |
| 2239 | Non_Targeting_Human | CCCACAATCAACGGCGTAC |
| 2240 | Non_Targeting_Human | TTGGCCTTGGTCGGGAGAA |
| 2241 | Non_Targeting_Human | CCGTTGACGCAGAAACTAA |
| 2242 | Non_Targeting_Human | TCGGCGTAGGAAATCCAGG |
| 2243 | Non_Targeting_Human | CCAGTACCATGATACTTCG |
| 2244 | Non_Targeting_Human | GACACCCGTGTAGAAGCGT |
| 2245 | Non_Targeting_Human | GTTCCGGACGGAGTGAGGA |
| 2246 | AC007383.3-002 | CCTCGGGCCGGTAGGCCCCA |
| 2247 | AC007383.3-002 | CCGTGGGGCCTACCGGCCCG |
| 2248 | AC007383.3-002 | CCTACCGGCCCGAGGCCTAC |
| 2249 | AC007383.3-002 | GGCGCCGGTAGGCCTCGGGC |
| 2250 | AC007383.3-002 | GCGGCTGCCGTGGGGCCTAC |
| 2251 | AC007383.3-002 | CCGGTAGGCCTCGGGCCGGT |
| 2252 | AC007383.3-002 | GGCAGGGCGCGGCTGCCGTG |
| 2253 | AC009120.10-001 | ATTACAGGCATGAGCCACCA |
| 2254 | AC009120.10-001 | TGAGGTGGGTGGATCACCTG |
| 2255 | AC009120.10-001 | ACAGGCATGCACACCACGCC |
| 2256 | AC009120.10-001 | TGTAATCCCAGCACTTTGGG |
| 2257 | AC009120.10-001 | CCTCAGCCTCCCAAAGTGCT |
| 2258 | AC009120.10-001 | TGCCTGTAATCCCAGCACTT |
| 2259 | AC009120.10-001 | AATACACAATTAGCCAGGCG |
| 2260 | AC009158.1-007 | ACAAACTAGGTAGTGAACAC |
| 2261 | AC009158.1-007 | GAATCAGACAGGATGAGAGG |
| 2262 | AC009158.1-007 | CATCAACACATGAATCAGAC |
| 2263 | AC009158.1-007 | ACCTTGGTGACAAGGAATGG |
| 2264 | AC009158.1-007 | CTGAGTTAGGCAATTGCTTG |
| 2265 | AC009158.1-007 | GAGGATAAGGCTAAAGACAG |
| 2266 | AC009158.1-007 | GCTACCTTGGTGACAAGGAA |
| 2267 | AC015849.16-002 | AATATTCACATATTTAACTG |
| 2268 | AC015849.16-002 | ATTCAGCAGCTAAATACCCC |
| 2269 | AC015849.16-002 | CGGAAGTTCATCATTTCCAG |
| 2270 | AC015849.16-002 | AAACGAACTTACAATGTCTG |
| 2271 | AC015849.16-002 | TGGGCTCTTGATAGATTTGG |
| 2272 | AC015849.16-002 | CTGAGTTGGAGATGGAGTCA |
| 2273 | AC015849.16-002 | GCGGAAGTTCATCATTTCCA |
| 2274 | AC017002.2-001 | CCTGTAGTGTGACTGTCAGG |
| 2275 | AC017002.2-001 | ACACTACAGGACAGTAACCA |
| 2276 | AC017002.2-001 | CACTACAGGACAGTAACCAT |
| 2277 | AC017002.2-001 | ACTACAGGACAGTAACCATG |
| 2278 | AC017002.2-001 | GCTTCACTTTGGCTTCTCAG |

| | | |
|---|---|---|
| 2279 | AC017002.2-001 | TTGGAAGAGACTGCAGTGCT |
| 2280 | AC017002.2-001 | CAGGACAGTAACCATGGGGC |
| 2281 | AC021224.1-201 | CACCATATCTACCACCAGCA |
| 2282 | AC021224.1-201 | CGTGCTGGTGGTAGATATGG |
| 2283 | AC021224.1-201 | TGGTAGATATGGTGGCAGTG |
| 2284 | AC021224.1-201 | TTTGGTGGCAGCCGTGCTGG |
| 2285 | AC021224.1-201 | AGCCGTGCTGGTGGTAGATA |
| 2286 | AC021224.1-201 | AGAAACTCTGGCCCCTATGG |
| 2287 | AC021224.1-201 | AGCAGCAGTAGCTGTGTCAG |
| 2288 | AC022182.1-003 | TCCCGCCACACCGCCCACCA |
| 2289 | AC022182.1-003 | AGGGAATGCGAAGGGCGCCG |
| 2290 | AC022182.1-003 | GGGATGTGCCCGCCTGCCCC |
| 2291 | AC022182.1-003 | CGGGATGTGCCCGCCTGCCC |
| 2292 | AC022182.1-003 | CCGCGGGCCCTGCGCGCCCG |
| 2293 | AC022182.1-003 | CCGCCTGCCCCGGGCGCGCA |
| 2294 | AC022182.1-003 | CCCTGCGCGCCCGGGGCAGG |
| 2295 | AC034187.2-001 | ATGACATGGAAATTCAGACC |
| 2296 | AC034187.2-001 | AGAGAAAAGAGATTCTACTG |
| 2297 | AC034187.2-001 | ATACTATTTATAGAATGACA |
| 2298 | AC034187.2-001 | AAAAATAGAGAAATATGGCT |
| 2299 | AC034187.2-001 | GAGATTCTACTGTGGCCACC |
| 2300 | AC034187.2-001 | CAAAAATTAGCTGGGTGTGG |
| 2301 | AC034187.2-001 | ATATGCAAATATATTCTCAC |
| 2302 | AC068286.1-001 | CCGAGTGCCTCCCTCCACCG |
| 2303 | AC068286.1-001 | TCATTGACTTCTCACAACAG |
| 2304 | AC068286.1-001 | AGTGGAGTGATGTACCAAAG |
| 2305 | AC068286.1-001 | TCCAAGTTTCCTTCTAGGCA |
| 2306 | AC068286.1-001 | ATCAGATGCTGTGTTCTCTC |
| 2307 | AC068286.1-001 | ACTGTTGTGAGAAGTCAATG |
| 2308 | AC068286.1-001 | ACAGCATCTGATTTCCTCTT |
| 2309 | AC092415.1-002 | CAGGAACAAATAGGACTGGG |
| 2310 | AC092415.1-002 | ACAATGACACCATGTTTGCT |
| 2311 | AC092415.1-002 | CTTACAAAGGAATGAGCAAG |
| 2312 | AC092415.1-002 | AAATCAGGAACAAATAGGAC |
| 2313 | AC092415.1-002 | TTTATCTTCTGGGAATCTGG |
| 2314 | AC092415.1-002 | CTAATTACAGTGAAGAGATG |
| 2315 | AC092415.1-002 | AATTTCAACTCTGCTTACCT |
| 2316 | AC131056.5-001 | AAATATTCTAAAATTGATTG |
| 2317 | AC131056.5-001 | TATTCTAAAATTGATTGTGG |
| 2318 | AC131056.5-001 | ATTGTGGTGGCCAGGCACAG |
| 2319 | AC131056.5-001 | TCCAGTCTGGCGAGAGAGTG |
| 2320 | AC131056.5-001 | CGAGACTAGCCTGACCAACA |
| 2321 | AC131056.5-001 | CGAGGCGGGCGGATCACCTG |

| | | |
|---|---|---|
| 2322 | AC131056.5-001 | GCACTTTGAGAGGCCGAGGC |
| 2323 | AC159540.1-001 | TAATTGATATGTGTATGTTG |
| 2324 | AC159540.1-001 | TTTGCATTGATGTTCATCGA |
| 2325 | AC159540.1-001 | ACTATTTGAAAATATTGAGG |
| 2326 | AC159540.1-001 | GGAACTATTTGAAAATATTG |
| 2327 | AC159540.1-001 | TGGCCTCATAGAATGAGTTG |
| 2328 | AC159540.1-001 | ATTTGAAAATATTGAGGAGG |
| 2329 | AC159540.1-001 | GTTGGCCTCATAGAATGAGT |
| 2330 | AC159540.2-001 | CATTTCCAAAGCCAAGCACG |
| 2331 | AC159540.2-001 | TCCAAAGCCAAGCACGTGGT |
| 2332 | AC159540.2-001 | GACAACAAATGCTTGCTCCT |
| 2333 | AC159540.2-001 | CTTCATCAGTCAGACAACAT |
| 2334 | AC159540.2-001 | AGAGATGTACAAAGTGCCAT |
| 2335 | AC159540.2-001 | AAAATGTAGGGCTTCAATGT |
| 2336 | AC159540.2-001 | GCCATGGGATCACCAAGCAG |
| 2337 | AF011889.2-001 | GGGTCCTTCAGCTCCTCGGG |
| 2338 | AF011889.2-001 | TGTGTCCTCCCTACCACCCG |
| 2339 | AF011889.2-001 | CCTACCACCCGAGGAGCTGA |
| 2340 | AF011889.2-001 | TCGGGTGGTAGGGAGGACAC |
| 2341 | AF011889.2-001 | TGGTAGGGAGGACACAGGCT |
| 2342 | AF011889.2-001 | GGTAGGGAGGACACAGGCTG |
| 2343 | AF011889.2-001 | AGGGAGGACACAGGCTGGGG |
| 2344 | AP001058.3-001 | GGCCCGGGGAGCATGGGTCA |
| 2345 | AP001058.3-001 | CCATGCTCCCCGGGCCCCTG |
| 2346 | AP001058.3-001 | TCGCCCTGACCCATGCTCCC |
| 2347 | AP001058.3-001 | CTCAGGGGCCCGGGGAGCAT |
| 2348 | AP001058.3-001 | CAGGGCGAGTGGCTGCTCCC |
| 2349 | AP001058.3-001 | GGTCCTCCTGGGTGCCTCAG |
| 2350 | AP001058.3-001 | CGCCCTGACCCATGCTCCCC |
| 2351 | AP006621.5-003 | CGTTCGTACCCGGTCCGGGT |
| 2352 | AP006621.5-003 | CTCGGATGCCGAACCCGCCG |
| 2353 | AP006621.5-003 | TGCGCGTTCGTACCCGGTCC |
| 2354 | AP006621.5-003 | CTGCGCGTTCGTACCCGGTC |
| 2355 | AP006621.5-003 | AGGCGCGACCTGCGACCGCC |
| 2356 | AP006621.5-003 | GGCGCGACCTGCGACCGCCC |
| 2357 | AP006621.5-003 | GAACCCGCCGGGGCCAAAGC |
| 2358 | bP-21264C1.2-002 | CTAGAACCCGTTTCACTGCC |
| 2359 | bP-21264C1.2-002 | AGAGGAACCCGTGGGCCTCA |
| 2360 | bP-21264C1.2-002 | CTAGAGGCGCAGGAGCAGGT |
| 2361 | bP-21264C1.2-002 | CAGACGCGTGGGCACATTGC |
| 2362 | bP-21264C1.2-002 | GAAACGGGTTCTAGAGGCGC |
| 2363 | bP-21264C1.2-002 | GAGCAGGTAGGAGAGGCCGG |
| 2364 | bP-21264C1.2-002 | CGGGATCCCCTGAGGCCCAC |

| | | |
|---|---|---|
| 2365 | C1orf132-001 | TGCGCAAACAGGCTCTCCCC |
| 2366 | C1orf132-001 | CCCGAGATGCGCAGCCGCTG |
| 2367 | C1orf132-001 | CGCATCTCGGGCCTGCAGCG |
| 2368 | C1orf132-001 | CCGCAGCGGCTGCGCATCTC |
| 2369 | C1orf132-001 | TGCGCATCTCGGGCCTGCAG |
| 2370 | C1orf132-001 | CCCGCCAAGCGCCCCGCTGC |
| 2371 | C1orf132-001 | GCCTGCAGCGGGGCGCTTGG |
| 2372 | CECR7-001 | CAGTCCGCAGGACCCCCTCG |
| 2373 | CECR7-001 | CTGCGTCAGCATCCGCGAGG |
| 2374 | CECR7-001 | GCATCCGCGAGGGGGTCCTG |
| 2375 | CECR7-001 | GCTGACGCAGAATGGAGTTG |
| 2376 | CECR7-001 | ATTCTGCGTCAGCATCCGCG |
| 2377 | CECR7-001 | ACGCAGAATGGAGTTGAGGT |
| 2378 | CECR7-001 | TTCTGCGTCAGCATCCGCGA |
| 2379 | CECR7-003 | TCCCCCTCCACTTACCGCCG |
| 2380 | CECR7-003 | AGGCCTCGGCGGTAAGTGGA |
| 2381 | CECR7-003 | TGGTGGCTTCCGTTTGTTGG |
| 2382 | CECR7-003 | GAGGAGGTGGCAGGCCTCGG |
| 2383 | CECR7-003 | TGGCTTCCGTTTGTTGGTGG |
| 2384 | CECR7-003 | CCACCTCCTCCACCAACAAA |
| 2385 | CECR7-003 | ATGTGGTGGCTTCCGTTTGT |
| 2386 | CH507-513H4.4-001 | TACCCGCGCCTGCCGGCGTA |
| 2387 | CH507-513H4.4-001 | TACCCTACGCCGGCAGGCGC |
| 2388 | CH507-513H4.4-001 | AACGGGTTACCCGCGCCTGC |
| 2389 | CH507-513H4.4-001 | TGTGCCTACCCTACGCCGGC |
| 2390 | CH507-513H4.4-001 | CCCCGATCCCCATCACGAAT |
| 2391 | CH507-513H4.4-001 | CCATCACGAATGGGGTTCAA |
| 2392 | CH507-513H4.4-001 | GCGACGGGCGGTGTGTACAA |
| 2393 | CH507-513H4.6-001 | CAATCGGTAGTAGCGACGGG |
| 2394 | CH507-513H4.6-001 | AGCGTGTGCCTACCCTACGC |
| 2395 | CH507-513H4.6-001 | CATCCAATCGGTAGTAGCGA |
| 2396 | CH507-513H4.6-001 | CCCATTCGTGATGGGGATCG |
| 2397 | CH507-513H4.6-001 | CGCCCGTCGCTACTACCGAT |
| 2398 | CH507-513H4.6-001 | CGTCGCTACTACCGATTGGA |
| 2399 | CH507-513H4.6-001 | CGCGCCTGCCGGCGTAGGGT |
| 2400 | CITF22-1A6.3-001 | GTGTGCACATGTGCACTGTG |
| 2401 | CITF22-1A6.3-001 | GTGTGCACGTGTGCACTGTG |
| 2402 | CITF22-1A6.3-001 | ACATGTGCACTGTGTATGCA |
| 2403 | CITF22-1A6.3-001 | GTGCACTGTGTGTGGATGCA |
| 2404 | CITF22-1A6.3-001 | GTGCACGTGTGCACGGTGTG |
| 2405 | CITF22-1A6.3-001 | GTGCACGTGTGCACTGTGTG |
| 2406 | CITF22-1A6.3-001 | ATGGTGTGTGCACGTGTGCA |
| 2407 | CRNDE-004 | GTTACCGACCGTTAGCGCCA |

| | | |
|---|---|---|
| 2408 | CRNDE-004 | TTACCGACCGTTAGCGCCAG |
| 2409 | CRNDE-004 | CTGGCGCTAACGGTCGGTAA |
| 2410 | CRNDE-004 | GGGGCTCGCGCCAGCCACGA |
| 2411 | CRNDE-004 | CGCGGACGCTCCCTCGTGGC |
| 2412 | CRNDE-004 | TGGCTGGCGCGAGCCCCCGG |
| 2413 | CRNDE-004 | TGGCGCGAGCCCCCGGCGGG |
| 2414 | CTB-58E17.1-001 | CCGCCGCCCTATGCGTGCCA |
| 2415 | CTB-58E17.1-001 | CTGCCCCAGTCCCTTCGCCG |
| 2416 | CTB-58E17.1-001 | GTCCCGCGGCGAAGGGACTG |
| 2417 | CTB-58E17.1-001 | CGCGCGGCCCTGGCACGCAT |
| 2418 | CTB-58E17.1-001 | CTCGCGCGTCCCGCGGCGAA |
| 2419 | CTB-58E17.1-001 | TCGCCGCGGGACGCGCGAGA |
| 2420 | CTB-58E17.1-001 | TCCGCCGCCCTATGCGTGCC |
| 2421 | CTB-61M7.2-001 | AAATAGGGTAATCCTCCTGT |
| 2422 | CTB-61M7.2-001 | AATTTGTAGATGAACACAGC |
| 2423 | CTB-61M7.2-001 | GAAAGTCATAATCCTACAGG |
| 2424 | CTB-61M7.2-001 | AGTGAAAGTCATAATCCTAC |
| 2425 | CTB-61M7.2-001 | TTGTAGATGAACACAGCAGG |
| 2426 | CTB-61M7.2-001 | CTTCCCCAAATATAAATGGT |
| 2427 | CTB-61M7.2-001 | TTACCAACCATTTATATTTG |
| 2428 | CTC-444N24.8-001 | GCTTTGTGGTAAACGCGCTG |
| 2429 | CTC-444N24.8-001 | CTTTGTGGTAAACGCGCTGA |
| 2430 | CTC-444N24.8-001 | AGAAAAGACGGAGAATGTCT |
| 2431 | CTC-444N24.8-001 | GTAAATGGCCATACACAGCA |
| 2432 | CTC-444N24.8-001 | AGTAACATGACAGAAATTAG |
| 2433 | CTC-444N24.8-001 | TACTCAAACCAGACACATTG |
| 2434 | CTC-444N24.8-001 | GTAACATGACAGAAATTAGG |
| 2435 | CTD-2054N24.2-001 | AAAACACGCGAGCATCTCAG |
| 2436 | CTD-2054N24.2-001 | CGCCTGGGCTCAAACAACGC |
| 2437 | CTD-2054N24.2-001 | GTTTGAGCCCAGGCGCGGAA |
| 2438 | CTD-2054N24.2-001 | GCGTTGTTTGAGCCCAGGCG |
| 2439 | CTD-2054N24.2-001 | TGTTTGAGCCCAGGCGCGGA |
| 2440 | CTD-2054N24.2-001 | CATGGGTCCCAAGAGGGCTC |
| 2441 | CTD-2054N24.2-001 | CTCCAGCGTTGTTTGAGCCC |
| 2442 | CTD-2192J16.26-001 | CAGGCATGAGCCACTGCGCC |
| 2443 | CTD-2192J16.26-001 | CAAGACCAGCCTGACCAACA |
| 2444 | CTD-2192J16.26-001 | AAAAAATTAGCCGAGCATGG |
| 2445 | CTD-2192J16.26-001 | GAGATGAGGTTTCACCATGT |
| 2446 | CTD-2192J16.26-001 | CCAGGTGTGGCCGGGCGCAG |
| 2447 | CTD-2192J16.26-001 | CTTTGGGAGGCTGAGGCAAG |
| 2448 | CTD-2192J16.26-001 | TGTAATCCCAGCTACTCGAG |
| 2449 | CTD-2227E11.1-001 | ACTGACCCACTATAGAGAGC |
| 2450 | CTD-2227E11.1-001 | GACTGCTCTCTCATCCCTAG |

| | | |
|---|---|---|
| 2451 | CTD-2227E11.1-001 | AAGCTCATAAAACCAAAACA |
| 2452 | CTD-2227E11.1-001 | CAAATTGTAGTCCTGTTCCA |
| 2453 | CTD-2227E11.1-001 | CAGGGCAATGTCCATGGAAC |
| 2454 | CTD-2227E11.1-001 | CATGGAACAGGACTACAATT |
| 2455 | CTD-2227E11.1-001 | AGTGGAATGGAGGTTTGGGT |
| 2456 | CTD-2232E5.2-001 | ACCTCGTGATCCACCCACCT |
| 2457 | CTD-2232E5.2-001 | CAGGTGTGAGTCACCAAGCC |
| 2458 | CTD-2232E5.2-001 | GACCATCCTGGCTAACACAG |
| 2459 | CTD-2232E5.2-001 | TACATTCTTAGGGCCAGGCT |
| 2460 | CTD-2232E5.2-001 | GCCGAGGTGGGTGGATCACG |
| 2461 | CTD-2232E5.2-001 | TCAGGAGTCCAAGACCATCC |
| 2462 | CTD-2232E5.2-001 | TATGAAAAATTAGCCAGGTG |
| 2463 | CTD-2270L9.4-003 | ACTTAAGAGTTGGCTCCGGG |
| 2464 | CTD-2270L9.4-003 | GCGAGCCTGCGAGAGCACCG |
| 2465 | CTD-2270L9.4-003 | AGCACCGAGGCTAGCCTCCG |
| 2466 | CTD-2270L9.4-003 | GGCGTCCAGAACTTAAGAGT |
| 2467 | CTD-2270L9.4-003 | CAGAACTTAAGAGTTGGCTC |
| 2468 | CTD-2270L9.4-003 | GTTGGCTCCGGGCGGCAACG |
| 2469 | CTD-2270L9.4-003 | GTTTCTGGGGTTCGCCTCGG |
| 2470 | CTD-2270P14.1-001 | AAGCTGAACTCCAGCAACGG |
| 2471 | CTD-2270P14.1-001 | GACTCTGAGGAGGCCCCGAG |
| 2472 | CTD-2270P14.1-001 | CCTAAGCTGAACTCCAGCAA |
| 2473 | CTD-2270P14.1-001 | CGTTGCTGGAGTTCAGCTTA |
| 2474 | CTD-2270P14.1-001 | GCTGGAGTTCAGCTTAGGGA |
| 2475 | CTD-2270P14.1-001 | CCGAAGCGCTCCGCCGCTCG |
| 2476 | CTD-2270P14.1-001 | CCGTTGCTGGAGTTCAGCTT |
| 2477 | CTD-2313J17.6-001 | AGGATTACCTAAGATAACGC |
| 2478 | CTD-2313J17.6-001 | CTGAACGTTTGTCTTGAGCA |
| 2479 | CTD-2313J17.6-001 | CTTGCTCAAGACAAACGTTC |
| 2480 | CTD-2313J17.6-001 | GAAAGACAAGAGCAGCTCCA |
| 2481 | CTD-2313J17.6-001 | CCCAGCGCCACCCTGCACTG |
| 2482 | CTD-2313J17.6-001 | AGCAAGGCCTGCGTTATCTT |
| 2483 | CTD-2313J17.6-001 | CCCGGGTCCGCAGTGCAGGG |
| 2484 | CTD-2315E11.1-001 | GACCCACCGGGACCCGCGAG |
| 2485 | CTD-2315E11.1-001 | GGGACCCGCGAGCGGGATCA |
| 2486 | CTD-2315E11.1-001 | CCCAGCCCTGATCCCGCTCG |
| 2487 | CTD-2315E11.1-001 | CTGATCCCGCTCGCGGGTCC |
| 2488 | CTD-2315E11.1-001 | TCCCGCTCGCGGGTCCCGGT |
| 2489 | CTD-2315E11.1-001 | ATCCCGCTCGCGGGTCCCGG |
| 2490 | CTD-2315E11.1-001 | GCGAGCGGGATCAGGGCTGG |
| 2491 | CTD-2325M2.1-001 | TGGGGGTTGCTGTTTGAAAG |
| 2492 | CTD-2325M2.1-001 | CAAGTGATTTCCATGTCAAG |
| 2493 | CTD-2325M2.1-001 | AGCCTGTTCACCACTTGACA |

| | | |
|---|---|---|
| 2494 | CTD-2325M2.1-001 | CTAAGAAGAAAAGTAAATTG |
| 2495 | CTD-2325M2.1-001 | GTAAATTGGGGAAAGAGCAT |
| 2496 | CTD-2325M2.1-001 | GGTTGCTGTTTGAAAGAGGA |
| 2497 | CTD-2325M2.1-001 | CACTTGACATGGAAATCACT |
| 2498 | CTD-2619J13.14-002 | TGGTCCGCAGAAACGAACGA |
| 2499 | CTD-2619J13.14-002 | CGTTCGTTTCTGCGGACCAG |
| 2500 | CTD-2619J13.14-002 | GCGGCAGTGGGATCATTGGG |
| 2501 | CTD-2619J13.14-002 | CGCGCCTTCGTTCGTTTCTG |
| 2502 | CTD-2619J13.14-002 | TGGGATCATTGGGCGGTGGG |
| 2503 | CTD-2619J13.14-002 | CAGTGGGATCATTGGGCGGT |
| 2504 | CTD-2619J13.14-002 | TTTGCGGCAGTGGGATCATT |
| 2505 | CTD-3075F15.1-001 | CAGCCACAAGGCCCTTCGCT |
| 2506 | CTD-3075F15.1-001 | GGGCCCAGCGAAGGGCCTTG |
| 2507 | CTD-3075F15.1-001 | CTCGCGCCTAGGCGGCCGGT |
| 2508 | CTD-3075F15.1-001 | CCCAATAGGAAACAGCCACA |
| 2509 | CTD-3075F15.1-001 | TCCCCGCCCTCGCGCCTAGG |
| 2510 | CTD-3075F15.1-001 | CCTTCGCTGGGCCCGCCCAC |
| 2511 | CTD-3075F15.1-001 | ACAGCCACAAGGCCCTTCGC |
| 2512 | CYP4F35P-001 | AAACATGTCACGTTCTGGGA |
| 2513 | CYP4F35P-001 | GAAACATGTCACGTTCTGGG |
| 2514 | CYP4F35P-001 | AGTGCTTCTGAAAGGAAAGG |
| 2515 | CYP4F35P-001 | CAAAAGGCTCACTGTCCCAA |
| 2516 | CYP4F35P-001 | CCTTTACGGTCTGTAGGGCA |
| 2517 | CYP4F35P-001 | TGAGAAACATGTCACGTTCT |
| 2518 | CYP4F35P-001 | GGGCTGGAGAATCCGAGGGC |
| 2519 | EBLN3-005 | CAAACAGAAAGCACCAAACA |
| 2520 | EBLN3-005 | CAGGGACCAAAACGCGTAGA |
| 2521 | EBLN3-005 | GCAAACAGAAAGCACCAAAC |
| 2522 | EBLN3-005 | ACAGGGACCAAAACGCGTAG |
| 2523 | EBLN3-005 | GGGAGGGGCGAGGACAAATG |
| 2524 | EBLN3-005 | CGAGGACAAATGAGGTAAGG |
| 2525 | EBLN3-005 | AAACGCGTAGAGGGACGGGG |
| 2526 | EBLN3-006 | GGAGGGTAGGCAGGGGTCCC |
| 2527 | EBLN3-006 | CGGCGCGCAGCTGAGGGACC |
| 2528 | EBLN3-006 | GACCCGCAGCCCGGCCGCGT |
| 2529 | EBLN3-006 | CGCAGCCCGGCCGCGTGGGG |
| 2530 | EBLN3-006 | GCGGCCGGGCTGCGGGTCTA |
| 2531 | EBLN3-006 | TAGGCAGGGGTCCCAGGCCG |
| 2532 | EBLN3-006 | CGGCCGGGCTGCGGGTCTAG |
| 2533 | EPB41L4A-AS1-003 | CAGTCGACTCACCAGACTCA |
| 2534 | EPB41L4A-AS1-003 | GGGTCCTCTGGGATGGCGCC |
| 2535 | EPB41L4A-AS1-003 | TCTGGGATGGCGCCTGGCTG |
| 2536 | EPB41L4A-AS1-003 | CTCACGCAGGGGCGCCGGGC |

| | | |
|---|---|---|
| 2537 | EPB41L4A-AS1-003 | CACGCAGGGGCGCCGGGCGG |
| 2538 | EPB41L4A-AS1-003 | GGCGCCTGGCTGTGGCCGCG |
| 2539 | EPB41L4A-AS1-003 | GGTGGCCGCGTTCCCCCGCC |
| 2540 | FAM95B1-001 | CCCTTCCTAATCATACCAGA |
| 2541 | FAM95B1-001 | ATGAGATGTTTCATATGGCA |
| 2542 | FAM95B1-001 | CCTTCTGGTATGATTAGGAA |
| 2543 | FAM95B1-001 | ATGTTTCATATGGCAGGGGT |
| 2544 | FAM95B1-001 | CATAGTGGAAGTTTCTCAAG |
| 2545 | FAM95B1-001 | AGGAAGGGCCTGGTCAGTGT |
| 2546 | FAM95B1-001 | TGAGATGTTTCATATGGCAG |
| 2547 | FIRRE-001 | AATAAAAAGAAAAATTGGCT |
| 2548 | FIRRE-001 | GCATTTATTTAAAAAGATGT |
| 2549 | FIRRE-001 | TATTTAAAAAGATGTAGGCC |
| 2550 | FIRRE-001 | ATGCCACTGCACTCTAGTCT |
| 2551 | FIRRE-001 | TGTAACCCCAGAACTTTGGG |
| 2552 | FIRRE-001 | TTCAGCCTCCCAAAGTTCTG |
| 2553 | FIRRE-001 | GAAAAATTATTTGATGGCGT |
| 2554 | FLJ20021-002 | GCGGCTTAACCCAGCCGACA |
| 2555 | FLJ20021-002 | ACAAGGCACGCTTGCCAAAG |
| 2556 | FLJ20021-002 | CAAGTGCAGAGGGTGTCTCC |
| 2557 | FLJ20021-002 | GGAGAGTGGGCAAGGCGTCG |
| 2558 | FLJ20021-002 | GTGTGAGTAGCCAAGTGCAG |
| 2559 | FLJ20021-002 | TGTGAGTAGCCAAGTGCAGA |
| 2560 | FLJ20021-002 | GCACGCTTGCCAAAGAGGCG |
| 2561 | GLIDR-001 | AAAAGGCCTCTGAATAAGCA |
| 2562 | GLIDR-001 | GTTTCATATGCACTGGCATG |
| 2563 | GLIDR-001 | GATTCTCAACCTGAGATCTC |
| 2564 | GLIDR-001 | TTCTCAAACCCTGAGATCTC |
| 2565 | GLIDR-001 | ATTCTCAACCTGAGATCTCA |
| 2566 | GLIDR-001 | TTTATTCCTTGCTTATTCAG |
| 2567 | GLIDR-001 | TATGTGAAATTTCGTGTTTA |
| 2568 | GLIDR-003 | CGACACTTCAGCACCAGTGG |
| 2569 | GLIDR-003 | CGCCGACACTTCAGCACCAG |
| 2570 | GLIDR-003 | GCAGGCACCTATCTTGTGCA |
| 2571 | GLIDR-003 | GCCATCTCCGCCCACAGTGG |
| 2572 | GLIDR-003 | AGTGGCGGTGGCCACCACTG |
| 2573 | GLIDR-003 | CGCCACTGGTGCTGAAGTGT |
| 2574 | GLIDR-003 | CACTTCAGCACCAGTGGCGG |
| 2575 | GS1-124K5.3-002 | CTGAACGGATCCTTCCGCCT |
| 2576 | GS1-124K5.3-002 | GGCCTTCGGCGACGTCACAG |
| 2577 | GS1-124K5.3-002 | GGCCGGCGCTATTACGACGC |
| 2578 | GS1-124K5.3-002 | CGCGCCTGTGACGTCAGCTG |
| 2579 | GS1-124K5.3-002 | TTCCGGCGTCGTAATAGCGC |

| | | |
|---|---|---|
| 2580 | GS1-124K5.3-002 | GGCGCTATTACGACGCCGGA |
| 2581 | GS1-124K5.3-002 | ACTATGTCCAGGCTCGGCCG |
| 2582 | GS1-124K5.4-001 | GACGCATGGCCACAGACTGC |
| 2583 | GS1-124K5.4-001 | CACCGCCTGCTGACACTGAC |
| 2584 | GS1-124K5.4-001 | TGCCGGTCAGTGTCAGCAGG |
| 2585 | GS1-124K5.4-001 | GGTCAGTGTCAGCAGGCGGT |
| 2586 | GS1-124K5.4-001 | TTGTAGCTTATTGGTGTCGG |
| 2587 | GS1-124K5.4-001 | AGGGTCCGCAGGCCCAGACC |
| 2588 | GS1-124K5.4-001 | GACTGCCGGTCAGTGTCAGC |
| 2589 | GS1-124K5.4-002 | GACACCAATAAGCTACAAGG |
| 2590 | GS1-124K5.4-002 | CTGCGCTACTTCCCAGCCTG |
| 2591 | GS1-124K5.4-002 | GGCAGTAGTAGTACAGGCAG |
| 2592 | GS1-124K5.4-002 | GCCGACACCAATAAGCTACA |
| 2593 | GS1-124K5.4-002 | AGCGCAGGCAGTAGTAGTAC |
| 2594 | GS1-124K5.4-002 | TGCGGCGTAGGGCGCAGCAA |
| 2595 | GS1-124K5.4-002 | ACTTCCCAGCCTGCGGCGTA |
| 2596 | HOTAIRM1-001 | AAAACGGTGATCCATCACTG |
| 2597 | HOTAIRM1-001 | ACTGCGCGTCACCTAGACGG |
| 2598 | HOTAIRM1-001 | TTTAGTGGCATTTAAATCCC |
| 2599 | HOTAIRM1-001 | TGATCCATCACTGCGGAAGC |
| 2600 | HOTAIRM1-001 | TGACGCGCAGTCGCCCCCCC |
| 2601 | HOTAIRM1-001 | ACCTAGACGGCGGAGCGCCG |
| 2602 | HOTAIRM1-001 | CCGCCGCCTAGGCTGCCTGG |
| 2603 | HOTAIRM1-004 | TTCGCTCCAGCACTCCAAAT |
| 2604 | HOTAIRM1-004 | CGCCCCGCCAGGCCTCCGGG |
| 2605 | HOTAIRM1-004 | GGCGACTGCAAAGGCCGATT |
| 2606 | HOTAIRM1-004 | CAAAGGCCGATTTGGAGTGC |
| 2607 | HOTAIRM1-004 | TCCCAGCCCCCACCTCCCGG |
| 2608 | HOTAIRM1-004 | CACCTCCCGGAGGCCTGGCG |
| 2609 | HOTAIRM1-004 | CCCACCTCCCGGAGGCCTGG |
| 2610 | IQCH-AS1-002 | CCTGCCCGAGTTACTAATGA |
| 2611 | IQCH-AS1-002 | CGTCATTAGTAACTCGGGCA |
| 2612 | IQCH-AS1-002 | GAGTTACTAATGACGGCGTG |
| 2613 | IQCH-AS1-002 | CACGCCGTCATTAGTAACTC |
| 2614 | IQCH-AS1-002 | GTCATTAGTAACTCGGGCAG |
| 2615 | IQCH-AS1-002 | CAAAACCAGCTCCTTGACGC |
| 2616 | IQCH-AS1-002 | TCACGCCGTCATTAGTAACT |
| 2617 | IRF4 | CCGAGTCCAGGGCGAGGTAA |
| 2618 | IRF4 | TCGGAGCTGAGGGCAGCGGT |
| 2619 | IRF4 | TGCCCTCAGCTCCGAGTCCA |
| 2620 | IRF4 | GCGAGGTAAGGGCTGGAGTC |
| 2621 | IRF4 | TCAGCTCCGAGTCCAGGGCG |
| 2622 | IRF4 | CCCTTACCTCGCCCTGGACT |

| | | |
|---|---|---|
| 2623 | IRF4 | TCCGAGTCCAGGGCGAGGTA |
| 2624 | JPX-012 | AAAATAAAGTTAGGCCCAAG |
| 2625 | JPX-012 | ACGACCCATATATCCCTCCT |
| 2626 | JPX-012 | TGCAGAAGGTAGCTCCTAGG |
| 2627 | JPX-012 | GCAGAAGGTAGCTCCTAGGA |
| 2628 | JPX-012 | CTAGGAACAGCCTAACCTCT |
| 2629 | JPX-012 | CTAATAAGTCTGGACATGAA |
| 2630 | JPX-012 | CTGTTCCTAGGAAAGCTCTG |
| 2631 | KANTR-001 | CCCGCTGACCTAGGGTTGAG |
| 2632 | KANTR-001 | TGGGGCGGCCTCTCAACCCT |
| 2633 | KANTR-001 | CTCCGCAACCCGCTGACCTA |
| 2634 | KANTR-001 | GCCTCTCAACCCTAGGTCAG |
| 2635 | KANTR-001 | GCCCCAGTAAAAGCTCGCGC |
| 2636 | KANTR-001 | CCTCTCAACCCTAGGTCAGC |
| 2637 | KANTR-001 | GGCCGGCCGAAGCGCGCGAG |
| 2638 | LINC00106-001 | TCAGGCTTCATCGGACGGCC |
| 2639 | LINC00106-001 | GGTGAGGTTGGCTTTGTGCG |
| 2640 | LINC00106-001 | TTCAGGCTTCATCGGACGGC |
| 2641 | LINC00106-001 | GGGGTGAGGTTGGCTTTGTG |
| 2642 | LINC00106-001 | CCGGGAAGGAATCAGAGCAG |
| 2643 | LINC00106-001 | CCATCCATTCCAGAAGCATT |
| 2644 | LINC00106-001 | GCATTTGGCAAGCACATCCT |
| 2645 | LINC00152-003 | GAAACATACTCACCAAGGAG |
| 2646 | LINC00152-003 | ACTCACCAAGGAGAGGCTGG |
| 2647 | LINC00152-003 | CATACTCACCAAGGAGAGGC |
| 2648 | LINC00152-003 | TGGAGCCACCAGCCTCTCCT |
| 2649 | LINC00152-003 | CAGCACAGTTCCTGGGAAGG |
| 2650 | LINC00152-003 | TTCCCACAGCCCAGACTCCA |
| 2651 | LINC00152-003 | GTTGCTTCTGCAGTTTGCCT |
| 2652 | LINC00152-005 | CACAAGTATGAAGAGAATGT |
| 2653 | LINC00152-005 | AATGTCGGGAGAGGAAGTGG |
| 2654 | LINC00152-005 | ACAAGTATGAAGAGAATGTC |
| 2655 | LINC00152-005 | TATGAAGAGAATGTCGGGAG |
| 2656 | LINC00152-005 | AAATTGACATTCCAGACAAG |
| 2657 | LINC00152-005 | TCACAGCACAGTTCCTGGGA |
| 2658 | LINC00152-005 | TGAAGATCTGAAGACAGGCA |
| 2659 | LINC00273-001 | AGCCCGGCGTCCCAGTCGCG |
| 2660 | LINC00273-001 | GGCGTCCCAGTCGCGAGGAT |
| 2661 | LINC00273-001 | CCGCGCCCATCCTCGCGACT |
| 2662 | LINC00273-001 | TCGCGAGGATGGGCGCGGCA |
| 2663 | LINC00273-001 | CCCAGTCGCGAGGATGGGCG |
| 2664 | LINC00273-001 | CGGCGTCCCAGTCGCGAGGA |
| 2665 | LINC00273-001 | ACGCCGGGCTCATCCTCGCG |

| | | |
|---|---|---|
| 2666 | LINC00324-001 | ACCACTCCCACTGCTCACCT |
| 2667 | LINC00324-001 | AGGAGGCAGTACCCTAAGAG |
| 2668 | LINC00324-001 | CCTGAATCTCCCAGAGCCAA |
| 2669 | LINC00324-001 | GCCAAAGGAGCTCTTGGGTG |
| 2670 | LINC00324-001 | AGCTCTTGGGTGTGGAACTC |
| 2671 | LINC00324-001 | GGGAGATTCAGGGACGGGGA |
| 2672 | LINC00324-001 | CCAGAGCCAAAGGAGCTCTT |
| 2673 | LINC00352-001 | GAAATCTTGGAGCATACGGG |
| 2674 | LINC00352-001 | TGTATATGTGCGCGTATGTG |
| 2675 | LINC00352-001 | TGGAAATCTTGGAGCATACG |
| 2676 | LINC00352-001 | TCACAAAACATAAGCACTGT |
| 2677 | LINC00352-001 | GGAAATCTTGGAGCATACGG |
| 2678 | LINC00352-001 | TATGTATATGTGCGCGTATG |
| 2679 | LINC00352-001 | GTGCGCGTATGTGGGGATGT |
| 2680 | LINC00467-001 | GCGCTGTGACGTTCCCTACG |
| 2681 | LINC00467-001 | GACAACCTCAACCGGACCGA |
| 2682 | LINC00467-001 | AAGGTGCTGAACAACCAGTG |
| 2683 | LINC00467-001 | GTTGTCAAGTCGGACCAAAC |
| 2684 | LINC00467-001 | TGTGACGTTCCCTACGCGGT |
| 2685 | LINC00467-001 | GTGACGTTCCCTACGCGGTC |
| 2686 | LINC00467-001 | GTTGTTCAGCACCTTCGGTC |
| 2687 | LINC00494-004 | GCTTGCCCACCTCCTCAGAG |
| 2688 | LINC00494-004 | GGGAAGAGCACCCCAAGCAG |
| 2689 | LINC00494-004 | GAAGGGCAAGGCCGTGTCAG |
| 2690 | LINC00494-004 | TGAGGAGGTGGGCAAGCCCT |
| 2691 | LINC00494-004 | GGAGGCCTCTCTGAGGAGGT |
| 2692 | LINC00494-004 | TGCTTGGGGTGCTCTTCCCA |
| 2693 | LINC00494-004 | ACCTGCTGTGCCTCTGCTTG |
| 2694 | LINC00511-002 | GCGCTGGTATCCAGAGGACG |
| 2695 | LINC00511-002 | TGCCTAACAGAGGGTCCTTG |
| 2696 | LINC00511-002 | AGAACGACAAATGCCAGAGG |
| 2697 | LINC00511-002 | GGTGCGGCGCTGGTATCCAG |
| 2698 | LINC00511-002 | CGCAGAACGACAAATGCCAG |
| 2699 | LINC00511-002 | TCGTTCTGCGCTTCTCCGCA |
| 2700 | LINC00511-002 | GAGGCGGTGACCGCGTCCTC |
| 2701 | LINC00657-001 | CTGCCAAGTCAGTTCCGGTC |
| 2702 | LINC00657-001 | GAACGCAGCCCGCTCCTCCA |
| 2703 | LINC00657-001 | AGAACGCAGCCCGCTCCTCC |
| 2704 | LINC00657-001 | CCGGGGCCGGAGGGCCTGGA |
| 2705 | LINC00657-001 | GAGGGCCTGGAGGGCCCTGG |
| 2706 | LINC00657-001 | ACCCGCCGGCCCGGGGCCGG |
| 2707 | LINC00657-001 | CCAGGGCCCTCCAGGCCCTC |
| 2708 | LINC00869-001 | TTTCGTGGCGAAGCAGCAGG |

| | | |
|---|---|---|
| 2709 | LINC00869-001 | AGGCGTTGTCGGTGACATCA |
| 2710 | LINC00869-001 | TTGTCGGTGACATCACGGAG |
| 2711 | LINC00869-001 | ATGGCTCCGGGATGCGCGTG |
| 2712 | LINC00869-001 | TGTCGGTGACATCACGGAGA |
| 2713 | LINC00869-001 | AGAGCAGACGGGGATCCGGA |
| 2714 | LINC00869-001 | CACTTGCCTCACGCGCATCC |
| 2715 | LINC00936-001 | GCTCCGCAGAGCGGCACACA |
| 2716 | LINC00936-001 | GGCTCCGCAGAGCGGCACAC |
| 2717 | LINC00936-001 | GCGCCCTGTGTGCCGCTCTG |
| 2718 | LINC00936-001 | AGGTGGCTGCACCCCGCCCG |
| 2719 | LINC00936-001 | GCGGCTGCTGTTGCGGACGG |
| 2720 | LINC00936-001 | CTGTTGCGGACGGCGGCGTG |
| 2721 | LINC00936-001 | CTGCCCGTGCCGAGCGAGCG |
| 2722 | LINC00957-001 | GGACTTCATCACCAAGCTCG |
| 2723 | LINC00957-001 | CACCAAGCTCGTGGACATCG |
| 2724 | LINC00957-001 | CGCCACGCCCTGGCGCACGG |
| 2725 | LINC00957-001 | CTGCGCCACGCCCTGGCGCA |
| 2726 | LINC00957-001 | TGGCGCACGGTGGGCTGCAG |
| 2727 | LINC00957-001 | CTCCTCGATGTCCACGAGCT |
| 2728 | LINC00957-001 | CTGCAGCCCACCGTGCGCCA |
| 2729 | LINC00969-039 | GGCGACTACAGACGTGTCGG |
| 2730 | LINC00969-039 | CGGCGACTACAGACGTGTCG |
| 2731 | LINC00969-039 | GGCGGCGACTACAGACGTGT |
| 2732 | LINC00969-039 | GGCGCTGGCGCTGGCCAAGG |
| 2733 | LINC00969-039 | TGTCGCGGTTGCCAAGCGCT |
| 2734 | LINC00969-039 | TTGGCAACCGCGACAGGCCC |
| 2735 | LINC00969-039 | GCGGCGACTACAGACGTGTC |
| 2736 | LINC00969-044 | GCATTCCCACCAGCTACGAG |
| 2737 | LINC00969-044 | ACCGTGCATTATAACATGGA |
| 2738 | LINC00969-044 | CGGCATTCCCACCAGCTACG |
| 2739 | LINC00969-044 | CCATGTTATAATGCACGGTG |
| 2740 | LINC00969-044 | GTCCATGTTATAATGCACGG |
| 2741 | LINC00969-044 | GGCATTCCCACCAGCTACGA |
| 2742 | LINC00969-044 | ACCTGCCCCTCGTAGCTGGT |
| 2743 | LINC00969-129 | CAGCTTTGTGTGGATGTACT |
| 2744 | LINC00969-129 | TGGAAGAAGCATAAGAACAT |
| 2745 | LINC00969-129 | TCCAATTAAACCAAATGCTG |
| 2746 | LINC00969-129 | CTCCAATTAAACCAAATGCT |
| 2747 | LINC00969-129 | GATGCAGGTGCAGCTTTGTG |
| 2748 | LINC00969-129 | CTTTGTGTGGATGTACTGGG |
| 2749 | LINC00969-129 | GACAAATTGAGATTTGCTGA |
| 2750 | LINC00996-001 | AGTGACTACCCCGACCTACC |
| 2751 | LINC00996-001 | CACAACAGCCTGGTAGGTCG |

| | | |
|---|---|---|
| 2752 | LINC00996-001 | TCACTTCCATTTGGGAGATG |
| 2753 | LINC00996-001 | TTGGGAGATGAGGAAGTCGA |
| 2754 | LINC00996-001 | AGCTCACAGAATCAGGAAGA |
| 2755 | LINC00996-001 | GGTTCCCACAACAGCCTGGT |
| 2756 | LINC00996-001 | AGGTCTAAGCTCACAGAATC |
| 2757 | LINC00996-002 | AGCTTCCTACCGTTCCACGA |
| 2758 | LINC00996-002 | GTTGTGGGAACCGAACGATG |
| 2759 | LINC00996-002 | AGTGGCCCTCGTGGAACGGT |
| 2760 | LINC00996-002 | GTTCCACGAGGGCCACTGAA |
| 2761 | LINC00996-002 | AAAAGGTGACAGTCTCCTTG |
| 2762 | LINC00996-002 | CAGCTTCCTACCGTTCCACG |
| 2763 | LINC00996-002 | GTTCGGTTCCCACAACAGCC |
| 2764 | LINC01002-019 | TCTTTAAAACTGGAAGAGGG |
| 2765 | LINC01002-019 | GGGACCTCAGTATGTTGCCA |
| 2766 | LINC01002-019 | TCAGCCATGGCAACATACTG |
| 2767 | LINC01002-019 | CATTCTTTAAAACTGGAAGA |
| 2768 | LINC01002-019 | AGGCTATAATGAGCCATGAT |
| 2769 | LINC01002-019 | AGAAGGTTAAGAACCAGAGA |
| 2770 | LINC01002-019 | AACTGGAAGAGGGAGGCAGA |
| 2771 | LINC01003-001 | GCGGGGAATTAGGCGGTGCG |
| 2772 | LINC01003-001 | GGCGGTGCGAGGCCAATGGG |
| 2773 | LINC01003-001 | GCGGTGCGAGGCCAATGGGA |
| 2774 | LINC01003-001 | GGCACCCCGCCTCAGCCCAG |
| 2775 | LINC01003-001 | GGCGAGGGCGGGGAATTAGG |
| 2776 | LINC01003-001 | GAGGGCGGGCCGGGTCCTCT |
| 2777 | LINC01003-001 | TTAGGCGGTGCGAGGCCAAT |
| 2778 | LINC01089-006 | CTTCATCAGACACGAAGGAG |
| 2779 | LINC01089-006 | ACATCCTTCATCAGACACGA |
| 2780 | LINC01089-006 | AAGGAGAGGCCAACAGATGA |
| 2781 | LINC01089-006 | CTAGTTATACAAGACAAAGA |
| 2782 | LINC01089-006 | GAACTGGATGGTAGAGTCAA |
| 2783 | LINC01089-006 | AATGGAATTAAATGGCCAAG |
| 2784 | LINC01089-006 | ATGGAATTAAATGGCCAAGT |
| 2785 | LINC01128-004 | GGAGTGCGTTCGGTGCGCCG |
| 2786 | LINC01128-004 | GAGTGCGTTCGGTGCGCCGT |
| 2787 | LINC01128-004 | GCGGGAGCGAGAGTCGGGTG |
| 2788 | LINC01128-004 | TGGCGGGAGCGAGAGTCGGG |
| 2789 | LINC01128-004 | TTACCTAACAGGAAGTTGGG |
| 2790 | LINC01128-004 | CCGAAGAGGCTGCTACCCAG |
| 2791 | LINC01128-004 | GTGTCTGCGTCGGGTTCTGT |
| 2792 | LINC01128-016 | TGTTAGGTAAGAGGCGCGTG |
| 2793 | LINC01128-016 | AGCGCGCGTCACGCCGTGTG |
| 2794 | LINC01128-016 | CACGCCGTGTGGGGCCGAAG |

| | | |
|---|---|---|
| 2795 | LINC01128-016 | CCAACTTCCTGTTAGGTAAG |
| 2796 | LINC01128-016 | GCAGCCTCTTCGGCCCCACA |
| 2797 | LINC01128-016 | CGAGCGCGCGTCACGCCGTG |
| 2798 | LINC01128-016 | GGCGGGAGCGAGAGTCGGGT |
| 2799 | LINC01138-002 | TCAAGCGCGCACACCCCTCA |
| 2800 | LINC01138-002 | CCGGAGAGCGATCACCGGCT |
| 2801 | LINC01138-002 | TGGGCGGTCACATCTGGAAA |
| 2802 | LINC01138-002 | TGCGCGCTTGAGCCGGAGCG |
| 2803 | LINC01138-002 | AGGGGTGTGCGCGCTTGAGC |
| 2804 | LINC01138-002 | TGTGCGCGCTTGAGCCGGAG |
| 2805 | LINC01138-002 | CCCAGCCGGTGATCGCTCTC |
| 2806 | LINC01234-006 | TGCCAACCTAAATCCAAGTG |
| 2807 | LINC01234-006 | ATTACTTCATGGAGCCCTGA |
| 2808 | LINC01234-006 | TTGAACACAATTGCTTCCAG |
| 2809 | LINC01234-006 | TTCCTGATTCCACTCCGTCA |
| 2810 | LINC01234-006 | TCTAGTAAAGAAGAAATGGT |
| 2811 | LINC01234-006 | CAACCTAAATCCAAGTGTGG |
| 2812 | LINC01234-006 | CACCACACTTGGATTTAGGT |
| 2813 | LINC01353-002 | TGTCACCACACACAGGCGCA |
| 2814 | LINC01353-002 | CCAGTCTCCATAGGGTGACA |
| 2815 | LINC01353-002 | GCGCCTGTGTGTGGTGACAG |
| 2816 | LINC01353-002 | TATTACCTTGCGCCTGTGTG |
| 2817 | LINC01353-002 | GTCAGCACCATGTCACCCTA |
| 2818 | LINC01353-002 | TGCCCGCTGTCACCACACAC |
| 2819 | LINC01353-002 | AAGGTAATACCAGTCTCCAT |
| 2820 | LINC01410-001 | CAGTAGAGCGCGAGGTGGAG |
| 2821 | LINC01410-001 | AGCACACTCTACAAATCTCG |
| 2822 | LINC01410-001 | CTCCACCTCGCGCTCTACTG |
| 2823 | LINC01410-001 | ACAGTAGAGCGCGAGGTGGA |
| 2824 | LINC01410-001 | GCAGGCCGGGGTGTGGTGGA |
| 2825 | LINC01410-001 | TTCAGCAGGCCGGGGTGTGG |
| 2826 | LINC01410-001 | CCCCCACCCAGGAAAACCAA |
| 2827 | LINC01410-002 | AACCCGCGCCTGCACCCTCG |
| 2828 | LINC01410-002 | CGGGACGGGACCGACCCCGA |
| 2829 | LINC01410-002 | GCGGGACGGGACCGACCCCG |
| 2830 | LINC01410-002 | CGCGCCTGCACCCTCGGGGT |
| 2831 | LINC01410-002 | GGGACCGACCCCGAGGGTGC |
| 2832 | LINC01410-002 | GGGTCGGTCCCGTCCCGCCC |
| 2833 | LINC01410-002 | GCAGCCCCACCCAACCCGGG |
| 2834 | LINC01481-001 | CTGGAATAAACCTCATGAGT |
| 2835 | LINC01481-001 | CAGGTGCCAGAAATCCAACA |
| 2836 | LINC01481-001 | CATAATGCCCTATAAACAGA |
| 2837 | LINC01481-001 | ACAGATGGATGTGAATGCTC |

| | | |
|---|---|---|
| 2838 | LINC01481-001 | TTAACATTCTCCATCAGCTG |
| 2839 | LINC01481-001 | AGTTATCTCAAATTGGTAAG |
| 2840 | LINC01481-001 | ACTGGAATAAACCTCATGAG |
| 2841 | LINC01534-001 | AATTCTCATAGTGCGTGGCT |
| 2842 | LINC01534-001 | ACGCACTATGAGAATTTGGA |
| 2843 | LINC01534-001 | TTCCAAATTCTCATAGTGCG |
| 2844 | LINC01534-001 | GTGCGTGGCTGGGATGAGGG |
| 2845 | LINC01534-001 | GCACACCCTGTCTGCGTCCC |
| 2846 | LINC01534-001 | TGAGGGCGGCGCTCACGGTG |
| 2847 | LINC01534-001 | TGAGGAAGCCCCGAGCTCCC |
| 2848 | LINC01549-201 | CAAGCAACATGGCTCCAGAG |
| 2849 | LINC01549-201 | AGAGCTCAATGCTTGAGAAA |
| 2850 | LINC01549-201 | AAGGAATTCTGTCCTTTGAA |
| 2851 | LINC01549-201 | TGGAGCCATGTTGCTTGGTC |
| 2852 | LINC01549-201 | GATAAAAATAGGAAGACGAT |
| 2853 | LINC01549-201 | AACAGGTATAACATGAAAGA |
| 2854 | LINC01549-201 | CACTCTGGAGCCATGTTGCT |
| 2855 | LINC01562-001 | GTATACCAGACCCCACGCTA |
| 2856 | LINC01562-001 | GGCTCTATCAGGAACAAGCC |
| 2857 | LINC01562-001 | GCAATTAGTTCATTTGAGTG |
| 2858 | LINC01562-001 | GGGAAAAGGATGATCATACC |
| 2859 | LINC01562-001 | CAGACCCCACGCTAAGGACT |
| 2860 | LINC01562-001 | CCAGACCCCACGCTAAGGAC |
| 2861 | LINC01562-001 | CCAGTCCTTAGCGTGGGGTC |
| 2862 | LINC01569-001 | AGCGCGAGCTCCATGCCCGG |
| 2863 | LINC01569-001 | CCTTCCCGGAGAGCAGGTGA |
| 2864 | LINC01569-001 | TCCCGGAGAGCAGGTGACGG |
| 2865 | LINC01569-001 | GTCAGCGCGAGCTCCATGCC |
| 2866 | LINC01569-001 | CCGTCACCTGCTCTCCGGGA |
| 2867 | LINC01569-001 | GCCGCCGTCACCTGCTCTCC |
| 2868 | LINC01569-001 | CTGCTCTCCGGGAAGGCCCT |
| 2869 | LINC01578-004 | GCGGAACGCCTGACAGCTCG |
| 2870 | LINC01578-004 | CAGCAGCACCCCGAGCTGTC |
| 2871 | LINC01578-004 | GGCGTTCCGCCGCGGCCGCG |
| 2872 | LINC01578-004 | CGTAGCTCCCCGCCGGCCGG |
| 2873 | LINC01578-004 | CCGCGGGCCTGCTGGCCGTC |
| 2874 | LINC01578-004 | GCGAGGCCCGCCGGCCGGCG |
| 2875 | LINC01578-004 | CCGCCGGCCGGCGGGCCTCG |
| 2876 | LINC01578-006 | CGCTGCGGTTGCCGCCTCGG |
| 2877 | LINC01578-006 | CCCAAACCCGAGTCCCGGGC |
| 2878 | LINC01578-006 | GGCTGTGACGCCGACGCGCG |
| 2879 | LINC01578-006 | GAGGCTCCCCCGGGCCCCGG |
| 2880 | LINC01578-006 | GCCCGGGGGAGCCTCCGAGG |

| | | |
|---|---|---|
| 2881 | LINC01578-006 | GCGAGGCGGGCCGCGCGCGT |
| 2882 | LINC01578-006 | CGCGCCGGGTCCAGCCGCCG |
| 2883 | LINC01578-007 | AGCCATGCTGTGTAATCACA |
| 2884 | LINC01578-007 | AGAATTATCCCAAAATGCTA |
| 2885 | LINC01578-007 | GGTTCTTAAGCAGAAAACAG |
| 2886 | LINC01578-007 | ATTCAATCACCAAGTTGATC |
| 2887 | LINC01578-007 | ACCTAATTGATTGGCACATG |
| 2888 | LINC01578-007 | AAGAATTATCCCAAAATGCT |
| 2889 | LINC01578-007 | ATTATATAAGATAAATATAG |
| 2890 | LINC01578-008 | CGACGCCGAAAGCGCCGCTG |
| 2891 | LINC01578-008 | CAGCGGCGCTTTCGGCGTCG |
| 2892 | LINC01578-008 | AGCGGCGCTTTCGGCGTCGT |
| 2893 | LINC01578-008 | CTCCGAGGCGGCAACCGCAG |
| 2894 | LINC01578-008 | GTCCAGCCGCCGGGGCCCGG |
| 2895 | LINC01578-008 | TGCCGCCTCGGAGGCTCCCC |
| 2896 | LINC01578-008 | TCGGAGGCTCCCCCGGGCCC |
| 2897 | MIAT-001 | TCGCGGGACCAGAGGGAGGG |
| 2898 | MIAT-001 | CGCTCGCGGGACCAGAGGGA |
| 2899 | MIAT-001 | TGAGATGGGAGGCGCTTGCA |
| 2900 | MIAT-001 | GCGCTCGCGGGACCAGAGGG |
| 2901 | MIAT-001 | GGCTCACAGCGCAGAACTGG |
| 2902 | MIAT-001 | CGGCTCACAGCGCAGAACTG |
| 2903 | MIAT-001 | GTCCGCAGCCTCCCTCCCTC |
| 2904 | MIR17HG-001 | CTGCGCTTTACTACGACCGG |
| 2905 | MIR17HG-001 | GGTCGTAGTAAAGCGCAGGC |
| 2906 | MIR17HG-001 | CTCCGGTCGTAGTAAAGCGC |
| 2907 | MIR17HG-001 | CGGTCGTAGTAAAGCGCAGG |
| 2908 | MIR17HG-001 | CGCCTGCGCTTTACTACGAC |
| 2909 | MIR17HG-001 | TTACTACGACCGGAGGCCCC |
| 2910 | MIR17HG-001 | CGTAGTAAAGCGCAGGCGGG |
| 2911 | MIR17HG-003 | GGAGGGAAACCTGTTGTGTG |
| 2912 | MIR17HG-003 | GGTTTCCCTCCGTCGCCGGA |
| 2913 | MIR17HG-003 | AGGTTTCCCTCCGTCGCCGG |
| 2914 | MIR17HG-003 | AACAGGTTTCCCTCCGTCGC |
| 2915 | MIR17HG-003 | CCCGCCCCTCCGGCGACGGA |
| 2916 | MIR17HG-003 | GTTTCCCTCCGTCGCCGGAG |
| 2917 | MIR17HG-003 | GCCCGCCCCTCCGGCGACGG |
| 2918 | MIR17HG-NEW-1 | CAAAACGCATGCCCGAGCAG |
| 2919 | MIR17HG-NEW-1 | GAGCAGGCCCCTAATTACCT |
| 2920 | MIR17HG-NEW-1 | GCAGCTGTGAAACCTCTGCT |
| 2921 | MIR17HG-NEW-1 | TGGGTTTCCGAGGTAATTAG |
| 2922 | MIR17HG-NEW-1 | GTTTACACGCGGCCAGGTTA |
| 2923 | MIR17HG-NEW-1 | TGTTTCGATGTTAAAGTCAC |

| | | |
|---|---|---|
| 2924 | MIR17HG-NEW-1 | CTCTGCTCGGGCATGCGTTT |
| 2925 | MIR17HG-NEW-2 | CCCTATTCCAGTCATACACG |
| 2926 | MIR17HG-NEW-2 | GAAAAGCTACTGGTGCAGTT |
| 2927 | MIR17HG-NEW-2 | GTTAGGTCCACGTGTATGAC |
| 2928 | MIR17HG-NEW-2 | AAAAACTAACTCATTAAGTA |
| 2929 | MIR17HG-NEW-2 | GTTTCTTAGAGAGTAAAAGT |
| 2930 | MIR17HG-NEW-2 | GCCAGTTTCCAAAAATGTAC |
| 2931 | MIR17HG-NEW-2 | CCACGTGTATGACTGGAATA |
| 2932 | MIR17HG-NEW-3 | AGGAAATCTTCACATCCACG |
| 2933 | MIR17HG-NEW-3 | CATCCAACTAGCACTGGAAG |
| 2934 | MIR17HG-NEW-3 | TTTCCTCTAAAAGGTACACA |
| 2935 | MIR17HG-NEW-3 | CCTTAAGAATTCTTTACAGA |
| 2936 | MIR17HG-NEW-3 | ACCAACCATCCAACTAGCAC |
| 2937 | MIR17HG-NEW-3 | CCAACTAGCACTGGAAGTGG |
| 2938 | MIR17HG-NEW-3 | TGCTAAGTGGAAGCCAGAAG |
| 2939 | MIR17HG-NEW-4 | CCTCGCCCGAGGGCGCGAAG |
| 2940 | MIR17HG-NEW-4 | GGTAGTCGTCCCCACCCCCT |
| 2941 | MIR17HG-NEW-4 | AGTGGCGCGAAGGCGCAGGT |
| 2942 | MIR17HG-NEW-4 | TCGCGCCACTTCGCGCCCTC |
| 2943 | MIR17HG-NEW-4 | GCGAAGTGGCGCGAAGGCGC |
| 2944 | MIR17HG-NEW-4 | GAGGGCGCGAAGTGGCGCGA |
| 2945 | MIR17HG-NEW-4 | GTGGCGCGAAGGCGCAGGTC |
| 2946 | MIR17HG-NEW-5 | TAAAGCTGCTTGCAAAGTGT |
| 2947 | MIR17HG-NEW-5 | TCAACTGAGCAGTAAAGGTA |
| 2948 | MIR17HG-NEW-5 | GAATAATCTGAAGTCTCAAG |
| 2949 | MIR17HG-NEW-5 | GCCTCTCAACTGAGCAGTAA |
| 2950 | MIR17HG-NEW-5 | AATAATCTGAAGTCTCAAGT |
| 2951 | MIR17HG-NEW-5 | TCAGTGTGTCTTCAAACTAC |
| 2952 | MIR17HG-NEW-5 | ACCTTTACTGCTCAGTTGAG |
| 2953 | MIR17HG-NEW-6 | CCCCCCAACCCGTTTCCCGG |
| 2954 | MIR17HG-NEW-6 | GGGTCAGAGTCAGGCCCCGC |
| 2955 | MIR17HG-NEW-6 | GAGTCAGGCCCCGCCGGACG |
| 2956 | MIR17HG-NEW-6 | GCCGCCGGGAAACGGGTTGG |
| 2957 | MIR17HG-NEW-6 | CCCCCGAGCCCGGCCCAGAG |
| 2958 | MIR17HG-NEW-6 | AGGCCTCGGGCCGCGTGCGA |
| 2959 | MIR17HG-NEW-6 | CGCGGAGCCCCCGCCCCTCT |
| 2960 | MIR17HG-NEW-7 | CGGCAGCACATGTCGCACAA |
| 2961 | MIR17HG-NEW-7 | CGGCTACGCGGAGAATCGCA |
| 2962 | MIR17HG-NEW-7 | GGCAGCACATGTCGCACAAG |
| 2963 | MIR17HG-NEW-7 | ATTCTCCGCGTAGCCGCCAG |
| 2964 | MIR17HG-NEW-7 | GCGGCTACGCGGAGAATCGC |
| 2965 | MIR17HG-NEW-7 | GATTCTCCGCGTAGCCGCCA |
| 2966 | MIR17HG-NEW-7 | TCTGACCCGCCGCCCCCTGG |

| | | |
|---|---|---|
| 2967 | MIR17HG-NEW-8 | TGTTCCTAAACTGCAGCAAA |
| 2968 | MIR17HG-NEW-8 | CCTTGTGCGACATGTGCTGC |
| 2969 | MIR17HG-NEW-8 | CGGCCCGGGCTCCATGAGCG |
| 2970 | MIR17HG-NEW-8 | CTGTTCCTAAACTGCAGCAA |
| 2971 | MIR17HG-NEW-8 | GCAGTTTAGGAACAGAAGAC |
| 2972 | MIR17HG-NEW-8 | TTAGGAACAGAAGACCGGGC |
| 2973 | MIR17HG-NEW-8 | CCGGGCTCCATGAGCGTGGC |
| 2974 | MIR17HG-NEW-9 | TGGTTATTTAGTGACAAACG |
| 2975 | MIR17HG-NEW-9 | TAAAAAGTTCCGGCTGGACA |
| 2976 | MIR17HG-NEW-9 | CCTTTCTATTCAGGAATCCA |
| 2977 | MIR17HG-NEW-9 | TTTCTTAAAGAACACATTAG |
| 2978 | MIR17HG-NEW-9 | AAATTTAAGGAACTTAGCCT |
| 2979 | MIR17HG-NEW-9 | AAAGTTCCGGCTGGACACGG |
| 2980 | MIR17HG-NEW-9 | TATGCTTAAAAAGTTCCGGC |
| 2981 | MIR222HG-002 | ACTTGGTCATCTATCACATG |
| 2982 | MIR222HG-002 | CCTGGGCTGCTGCTCAACGG |
| 2983 | MIR222HG-002 | ATCACATGTGGCTGAATCAC |
| 2984 | MIR222HG-002 | GCCTGGGCTGCTGCTCAACG |
| 2985 | MIR222HG-002 | CAGCAGCCCAGGCAGGAGAG |
| 2986 | MIR222HG-002 | TGCTGCTCAACGGGGGCTGG |
| 2987 | MIR222HG-002 | CTGCCTGGGCTGCTGCTCAA |
| 2988 | MIR22HG-004 | CGGCCACACCCAGCCCTTCC |
| 2989 | MIR22HG-004 | GCGGGGGCAGGGGCGCCTCG |
| 2990 | MIR22HG-004 | CCCGCCGCCCCTTCTCCTCC |
| 2991 | MIR22HG-004 | CAGGGGCGCCTCGGGGAGAG |
| 2992 | MIR22HG-004 | GCAGGGGCGCCTCGGGGAGA |
| 2993 | MIR22HG-004 | GGGAGGAGAAGGGGCGGCGG |
| 2994 | MIR22HG-004 | CGGGAGGAGAAGGGGCGGCG |
| 2995 | MIR4435-2HG-001 | ATGACAGAAACATACTCACC |
| 2996 | MIR4435-2HG-001 | ACTCACCAGGGAGAGGCTGG |
| 2997 | MIR4435-2HG-001 | TGACAGAAACATACTCACCA |
| 2998 | MIR4435-2HG-001 | GTGTACATCATTGGGAATGG |
| 2999 | MIR4435-2HG-001 | CATACTCACCAGGGAGAGGC |
| 3000 | MIR4435-2HG-001 | GAAACATACTCACCAGGGAG |
| 3001 | MIR4435-2HG-001 | TGGAGCCACCAGCCTCTCCC |
| 3002 | MIR663AHG-009 | GGGAACGACACACCACCGCT |
| 3003 | MIR663AHG-009 | CTCAGGTTCCCGAGGCCGAG |
| 3004 | MIR663AHG-009 | GCGCCTCGGTGTCCGCGCGT |
| 3005 | MIR663AHG-009 | CCAGCGCCCCCTCCTCCGGT |
| 3006 | MIR663AHG-009 | AGGTTCCCGAGGCCGAGCGG |
| 3007 | MIR663AHG-009 | GACACCGAGGCGCCGACCGG |
| 3008 | MIR663AHG-009 | CCGAGGCGCCGACCGGAGGA |
| 3009 | MIR663AHG-035 | GGAGAGACAAGGGGCCCCCG |

| | | |
|---|---|---|
| 3010 | MIR663AHG-035 | CCTTCTTCTCGTCCCGCCCG |
| 3011 | MIR663AHG-035 | CGAGGAAGAGGAGAGACAAG |
| 3012 | MIR663AHG-035 | GGCGGACATTCGTGGCGAAT |
| 3013 | MIR663AHG-035 | TCTTCTCGTCCCGCCCGCGG |
| 3014 | MIR663AHG-035 | GGGCGAGGGGCGGACATTCG |
| 3015 | MIR663AHG-035 | CAACGCACCACCCGCAGGCG |
| 3016 | MYC | GCTGTAGTAATTCCAGCGAG |
| 3017 | MYC | GCAGCGCAGCTCTGCTCGCC |
| 3018 | MYC | GGCGAGCAGAGCTGCGCTGC |
| 3019 | MYC | CCGGCTAGGGTGGAAGAGCC |
| 3020 | MYC | TAATTCCAGCGAGAGGCAGA |
| 3021 | MYC | GTAATTCCAGCGAGAGGCAG |
| 3022 | MYC | GAGCGGGCGGCCGGCTAGGG |
| 3023 | NOVMYEL1-1 | GTATATACCCAGTAATGGGA |
| 3024 | NOVMYEL1-1 | ATTACTGGGTATATACCCAA |
| 3025 | NOVMYEL1-1 | TACCCAGTAATGGGACGGCT |
| 3026 | NOVMYEL1-1 | ATGCACACATATGTTTATTG |
| 3027 | NOVMYEL1-1 | TCATTGTTGGACATTTGGGT |
| 3028 | NOVMYEL1-1 | AATGTGGCGCATATACACCA |
| 3029 | NOVMYEL1-1 | ATACCCAGTAATGGGACGGC |
| 3030 | NOVMYEL1-2 | TGTAGTCCCAGCCACCCAGG |
| 3031 | NOVMYEL1-2 | AAAAAATTAGCTGGGCGTGG |
| 3032 | NOVMYEL1-2 | TGTAGGTTGCCTGTTTGAGA |
| 3033 | NOVMYEL1-2 | TCCTGCCTCAGCCTCCTGGG |
| 3034 | NOVMYEL1-2 | AAAAATATAAAAAATTAGCT |
| 3035 | NOVMYEL1-2 | TATAAAAAATTAGCTGGGCG |
| 3036 | NOVMYEL1-2 | CAGGAGAATCATTTGAACCC |
| 3037 | NOVMYEL1-6 | CAGACTCTATTAACTCAGGT |
| 3038 | NOVMYEL1-6 | ACCACAGACTCTATTAACTC |
| 3039 | NOVMYEL1-6 | ACAGACTCTATTAACTCAGG |
| 3040 | NOVMYEL1-6 | AGAGTCTGTGGTAAGCATTG |
| 3041 | NOVMYEL1-6 | ACCTGAGTTAATAGAGTCTG |
| 3042 | NOVMYEL1-6 | TGGGATGTGGGAGGAAACCA |
| 3043 | NOVMYEL1-6 | AAGAGAAACAATGTTATTCC |
| 3044 | NOVMYEL2-1 | AAATGGCAGGAAACCGTTAC |
| 3045 | NOVMYEL2-1 | ATTGTTTCCTCAACTGCATG |
| 3046 | NOVMYEL2-1 | TCTGCTTACCCATCCTGCAG |
| 3047 | NOVMYEL2-1 | GGGATTCAAACCTCCACTGC |
| 3048 | NOVMYEL2-1 | TTCAAACCTCCACTGCAGGA |
| 3049 | NOVMYEL2-1 | TCAAACCTCCACTGCAGGAT |
| 3050 | NOVMYEL2-1 | GGTAAGCAGACAGCTGATGC |
| 3051 | NOVMYEL2-11 | TATTCCTTTAGATCCCAGAG |
| 3052 | NOVMYEL2-11 | TCATGCCTCACAGGGCCCGT |

| | | |
|---|---|---|
| 3053 | NOVMYEL2-11 | AGATAACTCTTCCTTTCCCA |
| 3054 | NOVMYEL2-11 | ACGCATGCTCAACCAGCTGG |
| 3055 | NOVMYEL2-11 | GACACGCATGCTCAACCAGC |
| 3056 | NOVMYEL2-11 | CGCATGCTCAACCAGCTGGT |
| 3057 | NOVMYEL2-11 | AGAGGGCATCATGCCTCACA |
| 3058 | NOVMYEL2-7 | AAATGCCATCAAATAAGTCA |
| 3059 | NOVMYEL2-7 | CAGACTTTGGCCAAGTATAG |
| 3060 | NOVMYEL2-7 | TGTAATCCCAACACTTTGGG |
| 3061 | NOVMYEL2-7 | AGAGATCTGTTTCTATTCAT |
| 3062 | NOVMYEL2-7 | CAGGCATGAGCCACTATACT |
| 3063 | NOVMYEL2-7 | TGCCTGTAATCCCAACACTT |
| 3064 | NOVMYEL2-7 | ACACTTTGGGAGGCCGAGGA |
| 3065 | NOVMYEL3-3 | CTTGGACCCAGCTTACCCAA |
| 3066 | NOVMYEL3-3 | ACTTGGACCCAGCTTACCCA |
| 3067 | NOVMYEL3-3 | TACATGGATGACCTTCTGCT |
| 3068 | NOVMYEL3-3 | AGTCAGGTGCGCAAAAGGGA |
| 3069 | NOVMYEL3-3 | GTCCTCCAGGTGCTGAAGCA |
| 3070 | NOVMYEL3-3 | AGGGGAGTTCTCAAACCCTT |
| 3071 | NOVMYEL3-3 | TAGGGGAGTTCTCAAACCCT |
| 3072 | NOVMYEL3-5 | GAAATGCAAAGCCCAGTACA |
| 3073 | NOVMYEL3-5 | ATTTGAGTAGACTCTGTCAG |
| 3074 | NOVMYEL3-5 | TTTGAGTAGACTCTGTCAGA |
| 3075 | NOVMYEL3-5 | AAATGTGATACAAGAAGTGA |
| 3076 | NOVMYEL3-5 | ATAAAAAATTCAGGAAACTG |
| 3077 | NOVMYEL3-5 | AGGAAGCAGAAAACCAACCC |
| 3078 | NOVMYEL3-5 | AATGTGATACAAGAAGTGAA |
| 3079 | NOVMYEL4-2 | AAAAAGTTGACTCATTTGGA |
| 3080 | NOVMYEL4-2 | CTCCCATATAATCTGAGGTA |
| 3081 | NOVMYEL4-2 | AAAAGTTGACTCATTTGGAA |
| 3082 | NOVMYEL4-2 | AAAAAGGCAAACCACAAACA |
| 3083 | NOVMYEL4-2 | ATCCCTACCTCAGATTATAT |
| 3084 | NOVMYEL4-2 | GATCCAGTTTATTTGTTATG |
| 3085 | NOVMYEL4-2 | AAGTTGACTCATTTGGAAGG |
| 3086 | NOVMYEL4-3 | GGAAGAGTGATTGCAAAGTG |
| 3087 | NOVMYEL4-3 | AATCCCGGCACATGGAGTGA |
| 3088 | NOVMYEL4-3 | CAGGAAGAGTGATTGCAAAG |
| 3089 | NOVMYEL4-3 | CTGAGAAGTACAAGACCCAG |
| 3090 | NOVMYEL4-3 | TTCTCTCTGAATAGACCTCT |
| 3091 | NOVMYEL4-3 | AGGAAGAGTGATTGCAAAGT |
| 3092 | NOVMYEL4-3 | CTCTGGGTCTTGTACTTCTC |
| 3093 | NOVMYEL5-5 | CTATTAGCCAATCAGTTGGT |
| 3094 | NOVMYEL5-5 | TAGTCTATTAGCCAATCAGT |
| 3095 | NOVMYEL5-5 | ACTTCTGCCTACCAACTGAT |

| | | |
|---|---|---|
| 3096 | NOVMYEL5-5 | CAGAAGAAGCCAACGAATCT |
| 3097 | NOVMYEL5-5 | AGCAGGGTTGGCTCCTTACA |
| 3098 | NOVMYEL5-5 | GCTCCTTACAAGGACTATGA |
| 3099 | NOVMYEL5-5 | AGATAAAGGTGCAGCAGGGT |
| 3100 | NOVMYEL5-6 | TGCAGATGCACGCAGGTCTG |
| 3101 | NOVMYEL5-6 | AATTTACACTAAATAAACAC |
| 3102 | NOVMYEL5-6 | GTGAGTGCCACAGCATGTGA |
| 3103 | NOVMYEL5-6 | GGTCAGATATCAAGCTGTAA |
| 3104 | NOVMYEL5-6 | ATTGCAGATGCACGCAGGTC |
| 3105 | NOVMYEL5-6 | TGCTAATTCTCCTGTATACT |
| 3106 | NOVMYEL5-6 | TCTGCAGCCTTCACATGCTG |
| 3107 | NOVMYEL7-1 | GCGGGCGCGAAGCGCACCTG |
| 3108 | NOVMYEL7-1 | TCCTCTCATGGGCTTTCTGC |
| 3109 | NOVMYEL7-1 | CGGCTTCTGGGTAACCCTAG |
| 3110 | NOVMYEL7-1 | GGTGCGCTTCGCGCCCGCTA |
| 3111 | NOVMYEL7-1 | GTGTCTTACCTTCCTCTCAT |
| 3112 | NOVMYEL7-1 | AGCGGGCGCGAAGCGCACCT |
| 3113 | NOVMYEL7-1 | AGGTGCGCTTCGCGCCCGCT |
| 3114 | NOVMYEL7-5 | GATGGGCGGATCACGAGGTC |
| 3115 | NOVMYEL7-5 | ATCTCGGCCTCCCAAAATGC |
| 3116 | NOVMYEL7-5 | TCCCAAAATGCTGGGATTAC |
| 3117 | NOVMYEL7-5 | TGGCTGTAATCCCAGCTACT |
| 3118 | NOVMYEL7-5 | GCAGGACAATCGCTTGAACC |
| 3119 | NOVMYEL7-5 | CAGGACAATCGCTTGAACCC |
| 3120 | NOVMYEL7-5 | TGCCTGTAATCCCAGCATTT |
| 3121 | NOVMYEL7-6 | AATACTGTGACATAAAATAG |
| 3122 | NOVMYEL7-6 | TACTAAATGTTAAGCAACAC |
| 3123 | NOVMYEL7-6 | TATTTGTATAGGCCTGGAAG |
| 3124 | NOVMYEL7-6 | CTACTAAATGTTAAGCAACA |
| 3125 | NOVMYEL7-6 | TTTATTTATTTGTATAGGCC |
| 3126 | NOVMYEL7-6 | ATCAGTGTTATACACAGAAT |
| 3127 | NOVMYEL7-6 | TCCCTTACAATCCTTGCCAT |
| 3128 | NOVMYEL8-1 | GCACTCTGTTCACTCAACAA |
| 3129 | NOVMYEL8-1 | ACATGGGATTTCCCGCCCAG |
| 3130 | NOVMYEL8-1 | AGCACTCTGTTCACTCAACA |
| 3131 | NOVMYEL8-1 | CCATTGCATGAGCTTCCGCT |
| 3132 | NOVMYEL8-1 | GAATCCCCCACTCACAACAT |
| 3133 | NOVMYEL8-1 | GGAAATCCCATGTTGTGAGT |
| 3134 | NOVMYEL8-1 | GTGCTGACTCACTGAGAACC |
| 3135 | NOVMYEL8-2 | TTGACCAGGTCACCTACACG |
| 3136 | NOVMYEL8-2 | CTGAGAAGCTCAAGTCACAT |
| 3137 | NOVMYEL8-2 | TGAGAAGCTCAAGTCACATG |
| 3138 | NOVMYEL8-2 | CTTGTTAGAATGCTACCTCC |

| | | |
|---|---|---|
| 3139 | NOVMYEL8-2 | GAAGCTCAAGTCACATGGGG |
| 3140 | NOVMYEL8-2 | TAGCATTCTAACAAGACATC |
| 3141 | NOVMYEL8-2 | TTTGGCTGAGACTGTTGACC |
| 3142 | NOVMYEL9-1 | GAAGATAGTAGAAGACACAA |
| 3143 | NOVMYEL9-1 | ATAAAGAATGAGGACCTGCG |
| 3144 | NOVMYEL9-1 | ATTATAAATAGGTTATGATT |
| 3145 | NOVMYEL9-1 | GAATAAAGAATGAGGACCTG |
| 3146 | NOVMYEL9-1 | AATAAAGAATGAGGACCTGC |
| 3147 | NOVMYEL9-1 | TGTTTGGTAAATATTATGGA |
| 3148 | NOVMYEL9-2 | AAAAAAATTAGCCAGCGTGG |
| 3149 | NOVMYEL9-2 | TCCCAGCTACTCTGAGACTG |
| 3150 | NOVMYEL9-2 | TAAAAATACAGAAATTAGCC |
| 3151 | NOVMYEL9-2 | AGCTACTCTGAGACTGAGGC |
| 3152 | NOVMYEL9-2 | TGTTATCCCAGATATTCTGG |
| 3153 | NOVMYEL9-2 | GTTATCCCAGATATTCTGGA |
| 3154 | NOVMYEL9-2 | GAGAATCACTTGAACCAGGA |
| 3155 | NOVMYEL9-3 | CCTGAGAGCTTCCATCAGTG |
| 3156 | NOVMYEL9-3 | GCATCTTCGTGCAGATACAG |
| 3157 | NOVMYEL9-3 | TATTCCGGGAACCACACTGA |
| 3158 | NOVMYEL9-3 | CTGGAGGATCTCCATGACAG |
| 3159 | NOVMYEL9-3 | GAAGTGCATCTCCCCTGTCA |
| 3160 | NOVMYEL9-3 | AGCTGGAGGATCTCCATGAC |
| 3161 | NOVMYEL9-3 | GCTGGAGGATCTCCATGACA |
| 3162 | NUP50-AS1-001 | TTGGGGCGTCGGCTCACGTG |
| 3163 | NUP50-AS1-001 | TTTGGGGCGTCGGCTCACGT |
| 3164 | NUP50-AS1-001 | CGACTAGCGCGCGAACATCC |
| 3165 | NUP50-AS1-001 | TGAGCCGACGCCCCAAAGCG |
| 3166 | NUP50-AS1-001 | GCGCGCTAGTCGCCGCTGAG |
| 3167 | NUP50-AS1-001 | GCGCCAGCAGCCGCGCTTTG |
| 3168 | NUP50-AS1-001 | CATCCTGGAAGACGCGTTGC |
| 3169 | PCAT14-003 | AATGGGGAAGGTGATAACGT |
| 3170 | PCAT14-003 | AAAAGGAACAGCCCACATGA |
| 3171 | PCAT14-003 | ACTGCTTTACTACAGTAACC |
| 3172 | PCAT14-003 | AGTAAGATGTTGTTCTGCAG |
| 3173 | PCAT14-003 | GGAATTCTCTATAATTCCCA |
| 3174 | PCAT14-003 | AAAATAAAACATGGGAAATG |
| 3175 | PCAT14-003 | TGGGGAAGGTGATAACGTGG |
| 3176 | PCED1B-AS1-002 | AGCAAGAACAATAAGAGCCA |
| 3177 | PCED1B-AS1-002 | ATGTCAAAGTCTTGAAGCTT |
| 3178 | PCED1B-AS1-002 | GCAAGAACAATAAGAGCCAA |
| 3179 | PCED1B-AS1-002 | GATTAAACAATTCAGACCCT |
| 3180 | PCED1B-AS1-002 | TCTTATTGTTCTTGCTGGTG |
| 3181 | PCED1B-AS1-002 | TAGTTTCTTGCTTTAGGAAG |

| | | |
|---|---|---|
| 3182 | PCED1B-AS1-002 | AGCATTTAGTTTCTTGCTTT |
| 3183 | PCED1B-AS1-004 | CTAAAAATACAAAATCAGCC |
| 3184 | PCED1B-AS1-004 | ACAAAATCAGCCGGGTGTGG |
| 3185 | PCED1B-AS1-004 | TAATCTTGTGTTTCTGTGCA |
| 3186 | PCED1B-AS1-004 | AGCTACTCGGAGCCTGAAGC |
| 3187 | PCED1B-AS1-004 | CATGCACCCACCACCACACC |
| 3188 | PCED1B-AS1-004 | ACTAAAAATACAAAATCAGC |
| 3189 | PCED1B-AS1-004 | AATCAGCCGGGTGTGGTGGT |
| 3190 | PCED1B-AS1-007 | GGAGAAGATGACATTTCCGG |
| 3191 | PCED1B-AS1-007 | TCGTTTGCCCCAGATGCCAC |
| 3192 | PCED1B-AS1-007 | TGACATTTCCGGTGGCATCT |
| 3193 | PCED1B-AS1-007 | GACATTTCCGGTGGCATCTG |
| 3194 | PCED1B-AS1-007 | ATGACATTTCCGGTGGCATC |
| 3195 | PCED1B-AS1-009 | TACACCCAGATTACTGAGAA |
| 3196 | PCED1B-AS1-009 | GCTGAAAAGATGGTCAGAAG |
| 3197 | PCED1B-AS1-009 | CGGCTGAAAAGATGGTCAGA |
| 3198 | PCED1B-AS1-009 | GGCTGAAAAGATGGTCAGAA |
| 3199 | PCED1B-AS1-009 | CAAGCTCAAAACCAAATTCC |
| 3200 | PCED1B-AS1-009 | ACATGTGTGAAAACATGAAA |
| 3201 | PCED1B-AS1-009 | GATGGTCAGAAGGGGAAAGG |
| 3202 | PTEN | TTGAGAGTTGAGCCGCTGTG |
| 3203 | PTEN | AGCCCGGCCTCGCCTCACAG |
| 3204 | PTEN | AGTTGAGCCGCTGTGAGGCG |
| 3205 | PTEN | TCATCTCCCTCGCCTGAGCC |
| 3206 | PTEN | GCGAGGCCGGGCTCAGGCGA |
| 3207 | PTEN | GAGCCGCTGTGAGGCGAGGC |
| 3208 | PTEN | AGCCGCTGTGAGGCGAGGCC |
| 3209 | PVT1-005 | CGTGTGGCGGCCGAGCACAT |
| 3210 | PVT1-005 | GCGTGTGGCGGCCGAGCACA |
| 3211 | PVT1-005 | GTCGTCGCCCCTCCTCGTCC |
| 3212 | PVT1-005 | CGGCCGAGCACATGGGCCCG |
| 3213 | PVT1-005 | GGCCCGCGGGCCCATGTGCT |
| 3214 | PVT1-005 | GGCCGAGCACATGGGCCCGC |
| 3215 | PVT1-005 | CGGGGCGGCCGGGACGAGGA |
| 3216 | RAB30-AS1-006 | CGATCTGGTCGGATAGCCGC |
| 3217 | RAB30-AS1-006 | CCCTACATCAGTCCCCAGCA |
| 3218 | RAB30-AS1-006 | GCACGAACCTGGGAACCAGA |
| 3219 | RAB30-AS1-006 | GGCACGAACCTGGGAACCAG |
| 3220 | RAB30-AS1-006 | ATAGCCGCGGGCACGAACCT |
| 3221 | RAB30-AS1-006 | CGAACCTGGGAACCAGAGGG |
| 3222 | RAB30-AS1-006 | GGTTCCCAGGTTCGTGCCCG |
| 3223 | RALY-AS1-001 | GTCCGAGTGGGAGAAAGGAG |
| 3224 | RALY-AS1-001 | CGAGTGGGAGAAAGGAGCGG |

| | | |
|---|---|---|
| 3225 | RALY-AS1-001 | GGCGGGTCCGAGTGGGAGAA |
| 3226 | RALY-AS1-001 | CGCCGCTCCTTTCTCCCACT |
| 3227 | RALY-AS1-001 | CCACTCGGACCCGCCGCCCC |
| 3228 | RALY-AS1-001 | TGGGAGAAAGGAGCGGCGGA |
| 3229 | RALY-AS1-001 | GGAGAAAGGAGCGGCGGAGG |
| 3230 | RP1-315G1.3-001 | TAGAGGCCCCGAGGGAGAGT |
| 3231 | RP1-315G1.3-001 | CCCTCGCCCAACTCTCCCTC |
| 3232 | RP1-315G1.3-001 | CCTCGCCCAACTCTCCCTCG |
| 3233 | RP1-315G1.3-001 | CCCCGAGGGAGAGTTGGGCG |
| 3234 | RP1-315G1.3-001 | AGAGGCCCCGAGGGAGAGTT |
| 3235 | RP1-315G1.3-001 | CCCGAGGGAGAGTTGGGCGA |
| 3236 | RP1-315G1.3-001 | ACTCTCCCTCGGGGCCTCTA |
| 3237 | RP11-1070N10.3-001 | CTGTCACCAAGGCTTGCGCG |
| 3238 | RP11-1070N10.3-001 | ACAGGTCAGCCCGCCCCTAG |
| 3239 | RP11-1070N10.3-001 | GAGCGCGCTCAGTAGTGCCT |
| 3240 | RP11-1070N10.3-001 | GGCGGGCTGACCTGTCACCA |
| 3241 | RP11-1070N10.3-001 | GGCACTACTGAGCGCGCTCG |
| 3242 | RP11-1070N10.3-001 | GCGCTCGCGGTCGCCAGGAG |
| 3243 | RP11-1070N10.3-001 | TCACCAAGGCTTGCGCGCGG |
| 3244 | RP11-1100L3.8-002 | AGGAAGAAAGAGCTTGAGGT |
| 3245 | RP11-1100L3.8-002 | AAGAGCTTGAGGTGGGAGCG |
| 3246 | RP11-1100L3.8-002 | AAAGGCTGCCCAGGAAGGCG |
| 3247 | RP11-1100L3.8-002 | CAGGAAGAAAGAGCTTGAGG |
| 3248 | RP11-1100L3.8-002 | GAAAGAGCTTGAGGTGGGAG |
| 3249 | RP11-1100L3.8-002 | ATACAGGAAGAAAGAGCTTG |
| 3250 | RP11-1100L3.8-002 | GGAAAGGCTGCCCAGGAAGG |
| 3251 | RP11-1152H15.1-003 | AGATATTCAAAATCAAAGTG |
| 3252 | RP11-1152H15.1-003 | AGGGTCCAGATGCCCCCCTA |
| 3253 | RP11-1152H15.1-003 | CCATAAAATTTGTTGAACAC |
| 3254 | RP11-1152H15.1-003 | GACATTGGAATGACGCAACC |
| 3255 | RP11-1152H15.1-003 | AATGACGCAACCAGGAGCCA |
| 3256 | RP11-1152H15.1-003 | CCATGGGAAGCAGAAGAGGT |
| 3257 | RP11-1152H15.1-003 | CAAGGAATGCCTGGATCCAT |
| 3258 | RP11-1250I15.3-001 | GAAAAAGAGCAGGCTTGGCT |
| 3259 | RP11-1250I15.3-001 | AGCATACTAAATCTGGGCAA |
| 3260 | RP11-1250I15.3-001 | TTGGCCACCTGAAAAAGAGC |
| 3261 | RP11-1250I15.3-001 | CTCAAGTGATCCACCCGCCG |
| 3262 | RP11-1250I15.3-001 | TCTAAGAGCATACTAAATCT |
| 3263 | RP11-1250I15.3-001 | CGCGGCGGGTGGATCACTTG |
| 3264 | RP11-1250I15.3-001 | GCAGGCTTGGCTGGGCACGA |
| 3265 | RP11-12G12.7-001 | GGGGCGTGCGTGAGTTCCGG |
| 3266 | RP11-12G12.7-001 | TTGACCGGCCTGGATGCACG |
| 3267 | RP11-12G12.7-001 | TGGGTTGCTGAGCCGATGCT |

| | | |
|---|---|---|
| 3268 | RP11-12G12.7-001 | TGACCGGCCTGGATGCACGT |
| 3269 | RP11-12G12.7-001 | GCCGATGCTGGGAAGGTACG |
| 3270 | RP11-12G12.7-001 | ACCCCGTACCTTCCCAGCAT |
| 3271 | RP11-12G12.7-001 | TTGCTGAGCCGATGCTGGGA |
| 3272 | RP11-142G1.3-001 | AACACAAACTAACTCCCATG |
| 3273 | RP11-142G1.3-001 | AAAACACAAACTAACTCCCA |
| 3274 | RP11-142G1.3-001 | AAACACAAACTAACTCCCAT |
| 3275 | RP11-142G1.3-001 | ACACATACTGAGTAGAGATG |
| 3276 | RP11-142G1.3-001 | AAGGAGTGATGGAACTGCAA |
| 3277 | RP11-142G1.3-001 | AAACACATACTGAGTAGAGA |
| 3278 | RP11-142G1.3-001 | AAGAAAAAGCCTTACCAATA |
| 3279 | RP11-147L13.12-001 | TCCTGCCTCGTGATTGCAGG |
| 3280 | RP11-147L13.12-001 | TTACCTTGAATCTCATCATG |
| 3281 | RP11-147L13.12-001 | GCCACCTGCAATCACGAGGC |
| 3282 | RP11-147L13.12-001 | ATGAGCCACCTGCAATCACG |
| 3283 | RP11-147L13.12-001 | ACAGAATCCAATCAAAGATT |
| 3284 | RP11-147L13.12-001 | ACCCAGCAAAATAACTACAA |
| 3285 | RP11-147L13.12-001 | ATAACTACAATGGAGAGATC |
| 3286 | RP11-14N9.2-001 | AAGGGAAATAGGACCAGGTG |
| 3287 | RP11-14N9.2-001 | GGAGTGGAATCGGTAAGTGT |
| 3288 | RP11-14N9.2-001 | AGGAGTGGAATCGGTAAGTG |
| 3289 | RP11-14N9.2-001 | TTGAATAAATTAATGTACTG |
| 3290 | RP11-14N9.2-001 | TAGAAAAGGGAAATAGGACC |
| 3291 | RP11-14N9.2-001 | AAGACCCGCTTACACTCAAA |
| 3292 | RP11-14N9.2-001 | TATAGCTTGAGTCTAGAAAA |
| 3293 | RP11-156P1.3-003 | AGTGGGCAAGGATGATGGTG |
| 3294 | RP11-156P1.3-003 | ACTCGTGCTCCCACCCCCAG |
| 3295 | RP11-156P1.3-003 | GGATGATGGTGAGGCAGAGG |
| 3296 | RP11-156P1.3-003 | GCACGAGTGGGCAAGGATGA |
| 3297 | RP11-156P1.3-003 | CACTCGTGCTCCCACCCCCA |
| 3298 | RP11-156P1.3-003 | GTTGGCTCTGGTGCCCCTGG |
| 3299 | RP11-156P1.3-003 | CCACTCGTGCTCCCACCCCC |
| 3300 | RP11-159G9.5-001 | TGGCCACAAGCTCTTCTCCG |
| 3301 | RP11-159G9.5-001 | GGAGGAAGAGAGCGACCAAG |
| 3302 | RP11-159G9.5-001 | GGGCGGCGAGAGCCTCATGG |
| 3303 | RP11-159G9.5-001 | CGCTCTCTTCCTCCGCCATG |
| 3304 | RP11-159G9.5-001 | GACCAAGAGGCCGAACGCCT |
| 3305 | RP11-159G9.5-001 | AAGCTCTTCTCCGAGGCGTT |
| 3306 | RP11-159G9.5-001 | AGGGGCTGGTACGCGCTGGG |
| 3307 | RP11-15J22.8-001 | AATGCGCTAAAGCAATCTGG |
| 3308 | RP11-15J22.8-001 | AATGTGAAGAAAGCTGGAGT |
| 3309 | RP11-15J22.8-001 | GAATGCGCTAAAGCAATCTG |
| 3310 | RP11-15J22.8-001 | CGGGAGGGTTTGAGCCCAAG |

| | | |
|---|---|---|
| 3311 | RP11-15J22.8-001 | GCTAAAGCAATCTGGGGGCA |
| 3312 | RP11-15J22.8-001 | AGAATGCGCTAAAGCAATCT |
| 3313 | RP11-15J22.8-001 | CAGAATGCGCTAAAGCAATC |
| 3314 | RP11-160O5.1-001 | CGCCTCCCCGTCGACCACCC |
| 3315 | RP11-160O5.1-001 | CCAGCCCTGGGTGGTCGACG |
| 3316 | RP11-160O5.1-001 | GCGCCTAGTGAGGGTCAGGG |
| 3317 | RP11-160O5.1-001 | GCCCTGGGTGGTCGACGGGG |
| 3318 | RP11-160O5.1-001 | TAGTGAGGGTCAGGGAGGCC |
| 3319 | RP11-160O5.1-001 | CCCAGCCCTGGGTGGTCGAC |
| 3320 | RP11-160O5.1-001 | CCCCAGCCCTGGGTGGTCGA |
| 3321 | RP11-166P13.3-002 | CGGGACGGTACAGCAACCGG |
| 3322 | RP11-166P13.3-002 | GGTTGCTGTACCGTCCCGCC |
| 3323 | RP11-166P13.3-002 | GCAGGGAATCAAGGTTCCTC |
| 3324 | RP11-166P13.3-002 | CCGTCATCTCTCCGGTGCCC |
| 3325 | RP11-166P13.3-002 | GGGCGGGACGGTACAGCAAC |
| 3326 | RP11-166P13.3-002 | GCTTGCGGGGCAGGGAATCA |
| 3327 | RP11-166P13.3-002 | CTCTCCGGTGCCCGGGAGAG |
| 3328 | RP11-182L21.6-001 | GGAGCGGGAGCCGAGCCTCG |
| 3329 | RP11-182L21.6-001 | GGGGTCGGGAGGCACTGCCG |
| 3330 | RP11-182L21.6-001 | TCGGGAGGCACTGCCGGGGA |
| 3331 | RP11-182L21.6-001 | GCGCCGTGCCCCGGGCCGCG |
| 3332 | RP11-182L21.6-001 | CGGCGCCGCCTGCCCCTCCC |
| 3333 | RP11-182L21.6-001 | GCGGAGGGCAGGGGGTCGGG |
| 3334 | RP11-182L21.6-001 | GAGCCGAGCCTCGCGGCCCG |
| 3335 | RP11-225B17.2-001 | TAATTACCATAAAACAAAGG |
| 3336 | RP11-225B17.2-001 | ATATAATTACCATAAAACAA |
| 3337 | RP11-225B17.2-001 | TTACAAAATAGATGCATTGG |
| 3338 | RP11-225B17.2-001 | TTTACAAAATAGATGCATTG |
| 3339 | RP11-225B17.2-001 | AATTACCATAAAACAAAGGA |
| 3340 | RP11-225B17.2-001 | CATTTACAAAATAGATGCAT |
| 3341 | RP11-225B17.2-001 | TAAATGCTGTTTCACATATG |
| 3342 | RP11-242D8.1-003 | GGGCGCTTTACGGCGAACTC |
| 3343 | RP11-242D8.1-003 | CGCCTCTACGCTTCCAGTTG |
| 3344 | RP11-242D8.1-003 | CTGGAAGCGTAGAGGCGAGA |
| 3345 | RP11-242D8.1-003 | ACTGGAAGCGTAGAGGCGAG |
| 3346 | RP11-242D8.1-003 | AGGAAGCTAAGGCCGCGTTG |
| 3347 | RP11-242D8.1-003 | AGCCGCAACTGGAAGCGTAG |
| 3348 | RP11-242D8.1-003 | GTGACGTAATAAGCCGCAAC |
| 3349 | RP11-277P12.10-001 | AGCAAAGAACAGGAGTACAG |
| 3350 | RP11-277P12.10-001 | CAGGAAAGGGAGATAAGGGT |
| 3351 | RP11-277P12.10-001 | AGGAAAGGGAGATAAGGGTG |
| 3352 | RP11-277P12.10-001 | GAGCAAAGAACAGGAGTACA |
| 3353 | RP11-277P12.10-001 | TCAGGAAAGGGAGATAAGGG |

| | | |
|---|---|---|
| 3354 | RP11-277P12.10-001 | GTCCGAAAAGAGTCAGGAAA |
| 3355 | RP11-277P12.10-001 | TGAGGACCTGAGGTCATAGG |
| 3356 | RP11-284N8.3-001 | TTGAGTGTGCAATAAGGCTC |
| 3357 | RP11-284N8.3-001 | ATGAAATAGCTAGAAAGGAG |
| 3358 | RP11-284N8.3-001 | TTTCCTGCATAAGAAAATCT |
| 3359 | RP11-284N8.3-001 | AATGAAATAGCTAGAAAGGA |
| 3360 | RP11-284N8.3-001 | GATTAATGAAATAGCTAGAA |
| 3361 | RP11-284N8.3-001 | ATAGATCAGTGAAAATTCTA |
| 3362 | RP11-284N8.3-001 | AAAATGTTGAGTGTGCAATA |
| 3363 | RP11-28H5.2-001 | AAAAAAAGCTTTATACAGCT |
| 3364 | RP11-28H5.2-001 | TTCAAATATGCAATCTGGAC |
| 3365 | RP11-28H5.2-001 | TCAGAAGTTCAAGACCAGCC |
| 3366 | RP11-28H5.2-001 | CAGACTTCAAATATGCAATC |
| 3367 | RP11-28H5.2-001 | AAACCTGCAAAGCTAGAATT |
| 3368 | RP11-28H5.2-001 | CCACTTTGGGAGGCTGAGGA |
| 3369 | RP11-28H5.2-001 | TGTAATCCCACCACTTTGGG |
| 3370 | RP11-2B6.3-001 | CCATACCCACCTCGCGCTCG |
| 3371 | RP11-2B6.3-001 | CGAGCGCGAGGTGGGTATGG |
| 3372 | RP11-2B6.3-001 | GCTCGCGGAGAGGGCCGTCG |
| 3373 | RP11-2B6.3-001 | TCGCGGCCGTGGGAATGCCC |
| 3374 | RP11-2B6.3-001 | GGGCATTCCCACGGCCGCGA |
| 3375 | RP11-2B6.3-001 | GAGAGGGCCGTCGCGGCCGT |
| 3376 | RP11-2B6.3-001 | GGTGCTCTCCCGGCAGCGAC |
| 3377 | RP11-304L19.13-002 | AGTTTGTGGGAGTCGCGCTG |
| 3378 | RP11-304L19.13-002 | GAGGAAGCGTTAAAGAGTCA |
| 3379 | RP11-304L19.13-002 | TAAAGAGTCAAGGCAGTTTG |
| 3380 | RP11-304L19.13-002 | ACTCTTTAACGCTTCCTCGC |
| 3381 | RP11-304L19.13-002 | AGTCGCGCTGGGGACGTTCA |
| 3382 | RP11-304L19.13-002 | TGGCCCAATGGGCGCCAGCG |
| 3383 | RP11-304L19.13-002 | CAGTTTGTGGGAGTCGCGCT |
| 3384 | RP11-308B16.2-001 | GCTGTGCTCGAGTTCTCGCG |
| 3385 | RP11-308B16.2-001 | GTCCCAAGCCCTGCCCCGCG |
| 3386 | RP11-308B16.2-001 | AGGTCCCAAGCCCTGCCCCG |
| 3387 | RP11-308B16.2-001 | GGCGCCGAGCGAGGGCTGTG |
| 3388 | RP11-308B16.2-001 | GCTTCCCCCCGCCCCCGCCG |
| 3389 | RP11-308B16.2-001 | GTCGATGGGACTGGGCATCG |
| 3390 | RP11-308B16.2-001 | CTTCCCCCCGCCCCCGCCGT |
| 3391 | RP11-317N12.1-002 | CCTCACCCTCCAAACTTCAG |
| 3392 | RP11-317N12.1-002 | GAAAGTTCCCCAAAACTTTG |
| 3393 | RP11-317N12.1-002 | GTGTTCCCCTGAAGTTTGGA |
| 3394 | RP11-317N12.1-002 | GGGAAAGTTCCCCAAAACTT |
| 3395 | RP11-317N12.1-002 | AGAGGTTGAGAAGAATGAGC |
| 3396 | RP11-317N12.1-002 | TTTACTCTTTGCATGCCCTG |

| | | |
|---|---|---|
| 3397 | RP11-317N12.1-002 | TCCTCACCCTCCAAACTTCA |
| 3398 | RP11-322E11.2-001 | TGAATTATTCTTCTTATATG |
| 3399 | RP11-322E11.2-001 | TTGCAGAATTACAGGAGTAC |
| 3400 | RP11-322E11.2-001 | GGTGATTGCCAGGGACTGGG |
| 3401 | RP11-322E11.2-001 | ATTGCCAGGGACTGGGCGGA |
| 3402 | RP11-322E11.2-001 | AAGTAGAATGGTGATTGCCA |
| 3403 | RP11-322E11.2-001 | AATGGTGATTGCCAGGGACT |
| 3404 | RP11-322E11.2-001 | CTCCCCCTCCGCCCAGTCCC |
| 3405 | RP11-385F7.1-001 | CCAAGCTGTTCCGAGCCTCC |
| 3406 | RP11-385F7.1-001 | CGTCGCCGCCGCGGAAAGTG |
| 3407 | RP11-385F7.1-001 | TGGGCTCGCTCTCCCCCCGG |
| 3408 | RP11-385F7.1-001 | GTCGCCTCAGATGGATCAAA |
| 3409 | RP11-385F7.1-001 | AAGCTGTTCCGAGCCTCCGG |
| 3410 | RP11-385F7.1-001 | AGCTGTTCCGAGCCTCCGGG |
| 3411 | RP11-385F7.1-001 | CAAGCTGTTCCGAGCCTCCG |
| 3412 | RP11-395G23.3-002 | GGCTGAGCCCATTCACCTCG |
| 3413 | RP11-395G23.3-002 | CGGCCACAGGAGCTCAGCGC |
| 3414 | RP11-395G23.3-002 | CCCATTCACCTCGCGGCCAC |
| 3415 | RP11-395G23.3-002 | CGAGTGCGCCCCTCTCGGCG |
| 3416 | RP11-395G23.3-002 | GGCGAGTGCGCCCCTCTCGG |
| 3417 | RP11-395G23.3-002 | GCGCCGGCGCTGAGCTCCTG |
| 3418 | RP11-395G23.3-002 | GGCGGCGAGTGCGCCCCTCT |
| 3419 | RP11-446N19.1-001 | CTCAATGAGCTATGGACCTG |
| 3420 | RP11-446N19.1-001 | CCACAAAGACGTTTCATGTG |
| 3421 | RP11-446N19.1-001 | GGCTCAGACTCAATGAGCTA |
| 3422 | RP11-446N19.1-001 | CCTCACATGAAACGTCTTTG |
| 3423 | RP11-446N19.1-001 | CACAAAGACGTTTCATGTGA |
| 3424 | RP11-446N19.1-001 | ATAAAGTTCCAACAGGTGCT |
| 3425 | RP11-446N19.1-001 | AGAAAAATCCTGCTGAATCC |
| 3426 | RP11-452C8.1-001 | ATGGAGTAGCTGGGAAACGT |
| 3427 | RP11-452C8.1-001 | AGAACATAACAGAAAGCCTG |
| 3428 | RP11-452C8.1-001 | GTTCTGATGATGGTAAGCAG |
| 3429 | RP11-452C8.1-001 | ACTCCATCTCCCCACACCGC |
| 3430 | RP11-452C8.1-001 | ATAACAGAAAGCCTGCGGTG |
| 3431 | RP11-452C8.1-001 | TTGCTCTGATTTGTGCCATG |
| 3432 | RP11-452C8.1-001 | AACAGAAAGCCTGCGGTGTG |
| 3433 | RP11-46D6.1-002 | CTGGGGCTGAAATTCAAACG |
| 3434 | RP11-46D6.1-002 | GGGAACAGGGAGCATTCACA |
| 3435 | RP11-46D6.1-002 | GGGCTGAAATTCAAACGGGG |
| 3436 | RP11-46D6.1-002 | AAAAACAGAGGGGCTCTGCA |
| 3437 | RP11-46D6.1-002 | CCCTGTTCCCCCAAAAACAG |
| 3438 | RP11-46D6.1-002 | CCTGTTCCCCCAAAAACAGA |
| 3439 | RP11-46D6.1-002 | GGTCTTGGGAACAAAGGCCT |

| | | |
|---|---|---|
| 3440 | RP11-474N24.6-001 | CCAGGTATTAGGGTCCATCC |
| 3441 | RP11-474N24.6-001 | CCTGGATGGACCCTAATACC |
| 3442 | RP11-474N24.6-001 | CGAAACAGCAGTCTAGAAGT |
| 3443 | RP11-474N24.6-001 | CATGTGCGATCCAGGTATTA |
| 3444 | RP11-474N24.6-001 | CTGGATCGCACATGAAAACT |
| 3445 | RP11-474N24.6-001 | CTGAAATTCCTGGGAGTGAG |
| 3446 | RP11-474N24.6-001 | GCCAGGCTAGCAATTCCAGC |
| 3447 | RP11-482M8.1-002 | CGTCGTCGTCGTCGCCCGGG |
| 3448 | RP11-482M8.1-002 | TTCCCCGCGCCCGTGCCCAG |
| 3449 | RP11-482M8.1-002 | CGTCGTCGTCGTCGTCGCCC |
| 3450 | RP11-482M8.1-002 | TCGTCGTCGTCGTCGTCGCC |
| 3451 | RP11-482M8.1-002 | GCCGCGCACCGGCCGCCGCT |
| 3452 | RP11-482M8.1-002 | CGTCGTCGTCGCCCGGGAGG |
| 3453 | RP11-482M8.1-002 | CAGTGACAGACCCTGCGGCG |
| 3454 | RP11-493L12.4-002 | CTGGGATCATATCGGATGGG |
| 3455 | RP11-493L12.4-002 | ACTCCTGGGATCATATCGGA |
| 3456 | RP11-493L12.4-002 | ACTTATGTATCTGCCATACG |
| 3457 | RP11-493L12.4-002 | CTCCTGGGATCATATCGGAT |
| 3458 | RP11-493L12.4-002 | GATTACAGCTCCAGACAGAG |
| 3459 | RP11-493L12.4-002 | AAGTCAGGTATGCCACTCCT |
| 3460 | RP11-493L12.4-002 | ATCCGATATGATCCCAGGAG |
| 3461 | RP11-495P10.1-001 | TTCCTCAACAAGCTACAGCA |
| 3462 | RP11-495P10.1-001 | TGCCATGCTGTAGCTTGTTG |
| 3463 | RP11-495P10.1-001 | CAGGTGATTGGTCAGGGAGT |
| 3464 | RP11-495P10.1-001 | CCTTGGACCAAAACAAGTGG |
| 3465 | RP11-495P10.1-001 | AGAAATATAGTAGGTCCCAA |
| 3466 | RP11-495P10.1-001 | AAGGGCAGAAATCAGCATCC |
| 3467 | RP11-495P10.1-001 | AAATATAGTAGGTCCCAAAG |
| 3468 | RP11-495P10.1-003 | GCAAGTCTGGGACCGCTGGT |
| 3469 | RP11-495P10.1-003 | CGACTCTCCGGTCCGACCAG |
| 3470 | RP11-495P10.1-003 | CCAGACTTGCTCCCACGGCA |
| 3471 | RP11-495P10.1-003 | CGGTCCCAGACTTGCTCCCA |
| 3472 | RP11-495P10.1-003 | GGGAGCAAGTCTGGGACCGC |
| 3473 | RP11-495P10.1-003 | CGAAAGAGTTCCCGTGCCGT |
| 3474 | RP11-495P10.1-003 | AACACACAGCTGCGACTCTC |
| 3475 | RP11-495P10.1-004 | GGCTCTGATAGGAATGCCAG |
| 3476 | RP11-495P10.1-004 | AGACTCTAAGGTACTGTCTG |
| 3477 | RP11-495P10.1-004 | GTACCTTAGAGTCTAAGCTT |
| 3478 | RP11-495P10.1-004 | TAGCCACAAAAACAAGACAA |
| 3479 | RP11-495P10.1-004 | CCAGACCATTACCTGGCACT |
| 3480 | RP11-495P10.1-004 | GAGTCACAATCAGAATTAGA |
| 3481 | RP11-495P10.1-004 | GCTAAGGATTCTCAGAGTTG |
| 3482 | RP11-573G6.10-001 | GCTCTTCTACGTCCCGCGCG |

| | | |
|---|---|---|
| 3483 | RP11-573G6.10-001 | CGCCCTGGCCGGGAAAGGTG |
| 3484 | RP11-573G6.10-001 | CGGGACGTAGAAGAGCGCCC |
| 3485 | RP11-573G6.10-001 | GTCCCGCGCGCGGACAGCCG |
| 3486 | RP11-573G6.10-001 | GCGCCCTGGCCGGGAAAGGT |
| 3487 | RP11-573G6.10-001 | CGTAGAAGAGCGCCCTGGCC |
| 3488 | RP11-573G6.10-001 | ACGTAGAAGAGCGCCCTGGC |
| 3489 | RP11-57P19.1-001 | GGACTGTGACAGGAGATATG |
| 3490 | RP11-57P19.1-001 | AAGATAAACAAAAACCACTC |
| 3491 | RP11-57P19.1-001 | CAATAGTTCAGGACTGTGAC |
| 3492 | RP11-57P19.1-001 | CCTTTATAAAGGAGATTGAG |
| 3493 | RP11-57P19.1-001 | GACTGTGACAGGAGATATGG |
| 3494 | RP11-57P19.1-001 | CAAGACCAGTCTGGCCAACG |
| 3495 | RP11-57P19.1-001 | GAGACAGGGTTTCACCACGT |
| 3496 | RP11-630C16.2-001 | GGGTTTGAACACAATCTGCA |
| 3497 | RP11-630C16.2-001 | GAACACAATCTGCAAGGAAG |
| 3498 | RP11-630C16.2-001 | TTACTGACACAATTAGCAAT |
| 3499 | RP11-630C16.2-001 | AGAGGAAACCAGCAGCAGCG |
| 3500 | RP11-630C16.2-001 | GAATCAAAGGCTGGACGCCG |
| 3501 | RP11-630C16.2-001 | TGTAAAATAGAATCAAAGGC |
| 3502 | RP11-630C16.2-001 | TTTACTGACACAATTAGCAA |
| 3503 | RP11-670E13.6-001 | CGATCCCCAAGAGCTCCCGC |
| 3504 | RP11-670E13.6-001 | CCGATCCCCAAGAGCTCCCG |
| 3505 | RP11-670E13.6-001 | CCCAAGAGCTCCCGCGGGCT |
| 3506 | RP11-670E13.6-001 | CCAGCCCGCGGGAGCTCTTG |
| 3507 | RP11-670E13.6-001 | CCGGGGGCTAAGGCCGAGCC |
| 3508 | RP11-670E13.6-001 | CCCAGCCCGCGGGAGCTCTT |
| 3509 | RP11-670E13.6-001 | CCGCGGGAGCTCTTGGGGAT |
| 3510 | RP11-67L2.2-001 | GCCCAGACTTCCTGACACTG |
| 3511 | RP11-67L2.2-001 | ATGCTCACATTATAGTCACG |
| 3512 | RP11-67L2.2-001 | TCTGCATGGCTGAAGACATG |
| 3513 | RP11-67L2.2-001 | CACGAGGAAGTCTTGTTGGC |
| 3514 | RP11-67L2.2-001 | TAGTCACGAGGAAGTCTTGT |
| 3515 | RP11-67L2.2-001 | AGACTTCCTGACACTGAGGT |
| 3516 | RP11-67L2.2-001 | AAGTCTTGTTGGCCGGACAT |
| 3517 | RP11-6N17.4-001 | GTTTACGCGACACTTCTAAG |
| 3518 | RP11-6N17.4-001 | AGGGGCGACTTTGTACAGTG |
| 3519 | RP11-6N17.4-001 | ACCTGGGCGAGACTGGACTT |
| 3520 | RP11-6N17.4-001 | CAATGTATATGCTATTTCTG |
| 3521 | RP11-6N17.4-001 | TAAAAGCTCCTGCCACACCT |
| 3522 | RP11-6N17.4-001 | AGTCCAGTCTCGCCCAGGTG |
| 3523 | RP11-6N17.4-001 | CCCTAAGTCCAGTCTCGCCC |
| 3524 | RP11-703G6.1-001 | GTTGAATAAGCAGGATCCAA |
| 3525 | RP11-703G6.1-001 | TTCTACAAATGTATAATGGT |

| | | |
|---|---|---|
| 3526 | RP11-703G6.1-001 | TGTTGAATAAGCAGGATCCA |
| 3527 | RP11-703G6.1-001 | GGAGAAAGTGTTGAATAAGC |
| 3528 | RP11-703G6.1-001 | TCCATAGAGTGTACAACATC |
| 3529 | RP11-703G6.1-001 | GTCTTCCTTAATGTAAACTA |
| 3530 | RP11-703G6.1-001 | CCATTAAAGAAGTCATAGAG |
| 3531 | RP11-706O15.1-006 | ATAAATGCGACCGGGCGCGG |
| 3532 | RP11-706O15.1-006 | TTAAAGCAATAAATGCGACC |
| 3533 | RP11-706O15.1-006 | ACTCCAGCCTGGGCGACAGA |
| 3534 | RP11-706O15.1-006 | CACCTTGTTAGCCAGGAGGA |
| 3535 | RP11-706O15.1-006 | ATACCACTGCACTCCAGCCT |
| 3536 | RP11-706O15.1-006 | TCGCCCAGGCTGGAGTGCAG |
| 3537 | RP11-706O15.1-006 | CACTCCAGCCTGGGCGACAG |
| 3538 | RP11-706O15.3-002 | CCCGGGCTTGTAGAGCCAGG |
| 3539 | RP11-706O15.3-002 | GGGCTTGTAGAGCCAGGCGG |
| 3540 | RP11-706O15.3-002 | CGGGCTTGTAGAGCCAGGCG |
| 3541 | RP11-706O15.3-002 | CGCTGCTACCGAGCTGCAGG |
| 3542 | RP11-706O15.3-002 | CATTCCCAGTCTCGCGCCCG |
| 3543 | RP11-706O15.3-002 | GAGGCCGCGGGCGCGAGACT |
| 3544 | RP11-706O15.3-002 | CGGCCCGGGCTTGTAGAGCC |
| 3545 | RP11-734K23.9-001 | TTGGCCTCATAGAATGAGTT |
| 3546 | RP11-734K23.9-001 | TGAGGCAACCTTGCATCCTG |
| 3547 | RP11-734K23.9-001 | TGCTTCATCCCCAGGATGCA |
| 3548 | RP11-734K23.9-003 | ACTGCTTTGGGTAGTGTCTG |
| 3549 | RP11-734K23.9-003 | TAGACTAGCTGGACACATGC |
| 3550 | RP11-734K23.9-003 | GTGAAAGACGGCTGTGCTCT |
| 3551 | RP11-734K23.9-003 | TGGCTGAAGGATAGACTAGC |
| 3552 | RP11-734K23.9-003 | TCATCTCTCTGTCATGAGTC |
| 3553 | RP11-734K23.9-003 | AAGAGCACAGCCGTCTTTCA |
| 3554 | RP11-734K23.9-003 | CTGGACACATGCTGGCTGTG |
| 3555 | RP11-760H22.2-001 | ATGTCATCAGTGCTCTAACG |
| 3556 | RP11-760H22.2-001 | CAGATGCTTAGAGTCTACTT |
| 3557 | RP11-760H22.2-001 | TGCTCTAACGGGGCATAGGG |
| 3558 | RP11-760H22.2-001 | CTAAAAACCCACCTCAAGGG |
| 3559 | RP11-760H22.2-001 | TGTCTAAAAACCCACCTCAA |
| 3560 | RP11-760H22.2-001 | TTGTCTAAAAACCCACCTCA |
| 3561 | RP11-760H22.2-001 | GGCAAACAGACATGTAACAG |
| 3562 | RP11-783K16.13-001 | CATGGGCCCCCAACCACAGG |
| 3563 | RP11-783K16.13-001 | CTTATGGTCCTGACCTCCTG |
| 3564 | RP11-783K16.13-001 | TCCTGACCTCCTGTGGTTGG |
| 3565 | RP11-783K16.13-001 | GGGCATGGGCCCCCAACCAC |
| 3566 | RP11-783K16.13-001 | CTCCAGTCCCAGTAGGGCTC |
| 3567 | RP11-783K16.13-001 | AGTCCCAGTAGGGCTCAGGC |
| 3568 | RP11-783K16.13-001 | GTCCTGACCTCCTGTGGTTG |

| | | |
|---|---|---|
| 3569 | RP11-81H14.1-001 | TTTGGTCACCTACATGTCAA |
| 3570 | RP11-81H14.1-001 | CCACATGCTATGATGCAGTG |
| 3571 | RP11-81H14.1-001 | CCACACTGCATCATAGCATG |
| 3572 | RP11-81H14.1-001 | AGCCTGAGAGCAAATAAGGG |
| 3573 | RP11-81H14.1-001 | TAAGACCACCTTTGACATGT |
| 3574 | RP11-81H14.1-001 | TTACAATGAGCTCAAGTGTA |
| 3575 | RP11-81H14.1-001 | TTTACAATGAGCTCAAGTGT |
| 3576 | RP11-81H3.2-001 | CCCCTTACTCAACCCAACCA |
| 3577 | RP11-81H3.2-001 | TGAAGGCTGTGCGGACACTG |
| 3578 | RP11-81H3.2-001 | GTAAGGGGCCAAACCCCTGG |
| 3579 | RP11-81H3.2-001 | GCCTGCGCCGGAGAGGACCG |
| 3580 | RP11-81H3.2-001 | CCGTGGTTGGGTTGAGTAAG |
| 3581 | RP11-81H3.2-001 | ACCGTGGTTGGGTTGAGTAA |
| 3582 | RP11-81H3.2-001 | GGACACTGGGGGCCCGCCAG |
| 3583 | RP11-883G14.4-001 | GAGAAGGAGATTCCACGGTA |
| 3584 | RP11-883G14.4-001 | AGAAGGAGATTCCACGGTAA |
| 3585 | RP11-883G14.4-001 | AAAAGTAATAGCTTATGCCC |
| 3586 | RP11-883G14.4-001 | AAGGTACTGTTTGAAAGAGA |
| 3587 | RP11-883G14.4-001 | GAAGGGACAGGAGCTTAAGA |
| 3588 | RP11-883G14.4-001 | CATTTGGGGGGCCCTTACCG |
| 3589 | RP11-883G14.4-001 | TGAAAGAGAAGGAGATTCCA |
| 3590 | RP11-89K11.1-001 | GCCGCCGTCGCCACGCACCG |
| 3591 | RP11-89K11.1-001 | GCCGTCGCCACGCACCGAGG |
| 3592 | RP11-89K11.1-001 | GTGGCGACGGCGGCCAAGAG |
| 3593 | RP11-89K11.1-001 | CCGTCGCCACGCACCGAGGT |
| 3594 | RP11-89K11.1-001 | ACCTCGGTGCGTGGCGACGG |
| 3595 | RP11-89K11.1-001 | GCGTGGCGACGGCGGCCAAG |
| 3596 | RP11-89K11.1-001 | CGGCGGCCAAGAGGGGCCAG |
| 3597 | RP11-92G12.3-001 | TGCCTCCCACCCACGCCCGG |
| 3598 | RP11-92G12.3-001 | CCGTGACGTGAAGCGCCTCG |
| 3599 | RP11-92G12.3-001 | CGTTGCCTCCCACCCACGCC |
| 3600 | RP11-92G12.3-001 | ACGTGAAGCGCCTCGGGGTG |
| 3601 | RP11-92G12.3-001 | CGGTGCCGCCGGGCGTGGGT |
| 3602 | RP11-92G12.3-001 | TGGGTGGGAGGCAACGCCCG |
| 3603 | RP11-92G12.3-001 | TGCCGCCGGGCGTGGGTGGG |
| 3604 | RP11-92K15.3-001 | CTGCAATGTGGTGACCGGCG |
| 3605 | RP11-92K15.3-001 | CCTTGGGGTTCCCCAAGCCA |
| 3606 | RP11-92K15.3-001 | CCCGCCGGTCACCACATTGC |
| 3607 | RP11-92K15.3-001 | GTTGCCTGCAATGTGGTGAC |
| 3608 | RP11-92K15.3-001 | CTTGGGGAACCCCAAGGTGT |
| 3609 | RP11-92K15.3-001 | GAGAAGAGTTGAGCCCTTCT |
| 3610 | RP11-92K15.3-001 | CCCTGGCTTGGGGAACCCCA |
| 3611 | RP11-96H19.1-001 | CCGACCCCCACACACCCGGG |

| | | |
|---|---|---|
| 3612 | RP11-96H19.1-001 | GGAGGCGTTGGCAGGTACCT |
| 3613 | RP11-96H19.1-001 | ACGATGAGGTGCGGAGGCGT |
| 3614 | RP11-96H19.1-001 | GACCCGACCCCCACACACCC |
| 3615 | RP11-96H19.1-001 | CCCGCCGCCCGGGTGTGTGG |
| 3616 | RP11-96H19.1-001 | TGAGGTGCGGAGGCGTTGGC |
| 3617 | RP11-96H19.1-001 | GCCCCGCCGCCCGGGTGTGT |
| 3618 | RP13-188A5.1-001 | GTAATGAAAAGAGATGCGGT |
| 3619 | RP13-188A5.1-001 | CAAAGTAATGAAAAGAGATG |
| 3620 | RP13-188A5.1-001 | TAATGAAAAGAGATGCGGTG |
| 3621 | RP13-188A5.1-001 | GCACCCAAACTTCTTCCGGC |
| 3622 | RP13-188A5.1-001 | CCCAAACTTCTTCCGGCAGG |
| 3623 | RP13-188A5.1-001 | AGTAATGAAAAGAGATGCGG |
| 3624 | RP13-188A5.1-001 | CACCCAAACTTCTTCCGGCA |
| 3625 | RP13-212L9.1-001 | CTTGTAGGGAGGACGATTGG |
| 3626 | RP13-212L9.1-001 | GTGGTGCAGAGAGATTCTCT |
| 3627 | RP13-212L9.1-001 | GCACCACGTATTCCAGCTGG |
| 3628 | RP13-212L9.1-001 | TCTGCACCACGTATTCCAGC |
| 3629 | RP13-212L9.1-001 | CCACCTCCAGCTGGAATACG |
| 3630 | RP13-212L9.1-001 | AACTTGGTGTCCTTGTAGGG |
| 3631 | RP13-212L9.1-001 | CACCAATCGTCCTCCCTACA |
| 3632 | RP4-665J23.1-005 | TAGAGGCCTCCCTTCCGAGG |
| 3633 | RP4-665J23.1-005 | CGGTAGAGGCCTCCCTTCCG |
| 3634 | RP4-665J23.1-005 | GCTGGGGCTCCGCTCCCCGG |
| 3635 | RP4-665J23.1-005 | CGGGCGCAGGACTGCAGGCT |
| 3636 | RP4-665J23.1-005 | CCACGCGCGGGCTCCCGCCG |
| 3637 | RP4-665J23.1-005 | ACCACGCGCGGGCTCCCGCC |
| 3638 | RP4-665J23.1-005 | CCGAGGAGGCGGTGCTGGGG |
| 3639 | RP4-669L17.10-004 | TACACACACAGTAAAAATCT |
| 3640 | RP4-669L17.10-004 | GTCAATTTACATCAAGGGTA |
| 3641 | RP4-669L17.10-004 | GTGCTAAGATTATAGCTGGG |
| 3642 | RP4-669L17.10-004 | AAAGTGCTAAGATTATAGCT |
| 3643 | RP4-669L17.10-004 | TTACTGTGAGTCATGGTGAA |
| 3644 | RP4-669L17.10-004 | TCACAGTAATCTTACTTCTG |
| 3645 | RP4-669L17.10-004 | ACCCTTGATGTAAATTGACA |
| 3646 | RP4-669P10.16-001 | ACTCAGCCCACCCCACATCA |
| 3647 | RP4-669P10.16-001 | AAGGGAGGTGACTGTTGTGA |
| 3648 | RP4-669P10.16-001 | GGGGTGTCATATATGAAGGG |
| 3649 | RP4-669P10.16-001 | GGACTGCTGTCCCTGATGTG |
| 3650 | RP4-669P10.16-001 | GAGGGCAGGACTCAGCCTGG |
| 3651 | RP4-669P10.16-001 | GGCCTGAAATATGGGCCTCC |
| 3652 | RP4-669P10.16-001 | CTGCTGTCCCTGATGTGGGG |
| 3653 | RP4-680D5.9-001 | TGAGGCCCTTGGGATACGTG |
| 3654 | RP4-680D5.9-001 | AGAGGGATAAGGATAAGGAG |

| | | |
|---|---|---|
| 3655 | RP4-680D5.9-001 | AAGGATAAGGAGTGGGAATG |
| 3656 | RP4-680D5.9-001 | GAGGGATAAGGATAAGGAGT |
| 3657 | RP4-680D5.9-001 | ATACGTGGGGCTGAAGAAGA |
| 3658 | RP4-680D5.9-001 | GGAGTGGGAATGGGGCATTG |
| 3659 | RP4-680D5.9-001 | GATACGTGGGGCTGAAGAAG |
| 3660 | RP5-1159O4.1-001 | GGTCGTGGGTCCCAGCCCAG |
| 3661 | RP5-1159O4.1-001 | TGCGCGTCCTCCAGGCGTGG |
| 3662 | RP5-1159O4.1-001 | GGTCCAGGTCACGGGCCGCA |
| 3663 | RP5-1159O4.1-001 | GGCCCGTGACCTGGACCACT |
| 3664 | RP5-1159O4.1-001 | GAGCCCGGACGCGGGCAAGA |
| 3665 | RP5-1159O4.1-001 | AGCCCAGTGGTCCAGGTCAC |
| 3666 | RP5-1159O4.1-001 | GTGACCTGGACCACTGGGCT |
| 3667 | RP5-903G2.2-001 | ACGTCACTCCCGAAGAGAAG |
| 3668 | RP5-903G2.2-001 | TGGCCGGCCCAGGAGAGAGG |
| 3669 | RP5-903G2.2-001 | ATGCCTAGCTCCGCGCCGCG |
| 3670 | RP5-903G2.2-001 | CTGCGGCTGTTGCTGGAGGA |
| 3671 | RP5-903G2.2-001 | ACGTCGTCCTCCTCTCTCCT |
| 3672 | RP5-903G2.2-001 | GACGCCCCAGCTCCGCCGAC |
| 3673 | RP5-903G2.2-001 | GCGGCCTGTCGGCGGAGCTG |
| 3674 | RP5-968D22.3-001 | AAACCAGGAGAAGATCCACC |
| 3675 | RP5-968D22.3-001 | ACCAAAAATTAGCCAGGCGT |
| 3676 | RP5-968D22.3-001 | GAGAATGGCGTGAACCCAGG |
| 3677 | RP5-968D22.3-001 | AATATGTTGTTGGCCAGGCG |
| 3678 | RP5-968D22.3-001 | CTAAATCACTTAAGTTTCTG |
| 3679 | RP5-968D22.3-001 | TGAATGAAAAAATATGTTGT |
| 3680 | RP5-968D22.3-001 | AAAATAACCAAAAATTAGCC |
| 3681 | SERPINB9P1-001 | TTGAGGAGGAGTACTAAGAG |
| 3682 | SERPINB9P1-001 | TGAGGAGGAGTACTAAGAGT |
| 3683 | SERPINB9P1-001 | GGACTGCCAGTACTATGTTG |
| 3684 | SERPINB9P1-001 | CTGCCAGTACTATGTTGAGG |
| 3685 | SERPINB9P1-001 | TGTCTTGTTCCCATTCTCAG |
| 3686 | SERPINB9P1-001 | ACTCCTCCTCAACATAGTAC |
| 3687 | SERPINB9P1-001 | GGTTTCATATCAGGGATGCA |
| 3688 | SNHG1-001 | GAGGCTTAAAAGTAGCAGTG |
| 3689 | SNHG1-001 | TGCCTGGCCAGGGCGACTGG |
| 3690 | SNHG1-001 | GGCGGATAAGGTCTTGTGCG |
| 3691 | SNHG1-001 | AGGTCTTGTGCGTGGCCTCG |
| 3692 | SNHG1-001 | ACCTTATCCGCCAGTCGCCC |
| 3693 | SNHG1-001 | GAAGGACAAAATGGCGATGG |
| 3694 | SNHG1-001 | GCCAGGGCGACTGGCGGATA |
| 3695 | SNHG15-004 | TGCTACTTCCTCTCAAGAGG |
| 3696 | SNHG15-004 | CGCGGTGACGTCGAGGTGCG |
| 3697 | SNHG15-004 | GGGACTACGCGGTGACGTCG |

| | | |
|---|---|---|
| 3698 | SNHG15-004 | GACCATTCCTGAGCCCAGGT |
| 3699 | SNHG15-004 | AAGAGGCGGTTGAGCGCGTT |
| 3700 | SNHG15-004 | GCCACTGAACCCAATCCCCC |
| 3701 | SNHG15-004 | GCTGTGGCCCCGGCCACGCA |
| 3702 | SNHG16-003 | TCATAGGGCCGGCTTCTGCG |
| 3703 | SNHG16-003 | CATAGGGCCGGCTTCTGCGG |
| 3704 | SNHG16-003 | CGTCATAGGGCCGGCTTCTG |
| 3705 | SNHG16-003 | GCGCTCTCCCCCGCAGAAGC |
| 3706 | SNHG16-003 | TGCCGAGAAGACTTGCGCGC |
| 3707 | SNHG16-003 | AGCCCGTCCTCGGCGTCATA |
| 3708 | SNHG16-003 | GTCATAGGGCCGGCTTCTGC |
| 3709 | SNHG6-005 | AGCCGTTAGTCATGCCGGTG |
| 3710 | SNHG6-005 | CGAAGAGCCGTTAGTCATGC |
| 3711 | SNHG6-005 | CGTTAGTCATGCCGGTGTGG |
| 3712 | SNHG6-005 | CACCACACCGGCATGACTAA |
| 3713 | SNHG6-005 | GCCCGTAGCTGGGCTCTGCG |
| 3714 | SNHG6-005 | CTCCGCCGCCGCCACCACAC |
| 3715 | SNHG6-005 | TAGTCATGCCGGTGTGGTGG |
| 3716 | SNHG9-001 | CGAAAAGCGACTATAAACGC |
| 3717 | SNHG9-001 | AGAGCACTCAGAGCCCGGTG |
| 3718 | SNHG9-001 | TTCGGGTGACGTAGATTCCC |
| 3719 | SNHG9-001 | GAGCACTCAGAGCCCGGTGA |
| 3720 | SNHG9-001 | TGACGTAGATTCCCGGGCCG |
| 3721 | SNHG9-001 | TTTCGGGTGACGTAGATTCC |
| 3722 | SNHG9-001 | TGCCCGTCCGGCCCCAGCCG |
| 3723 | SNHG9-002 | TCGGGTGGATTCCCGAGCCG |
| 3724 | SNHG9-002 | TCGGGAATCCACCCGAAGAG |
| 3725 | SNHG9-002 | GTTTATAGCCACTCTTCGGG |
| 3726 | SNHG9-002 | TGAAGAGGAGAGCGCAATGG |
| 3727 | SNHG9-002 | AGGAGAGCGCAATGGAGGCG |
| 3728 | SNHG9-002 | AAGTGAAGAGGAGAGCGCAA |
| 3729 | SNHG9-002 | TTGCGCTCTCCTCTTCACTT |
| 3730 | TP53 | GGGACACTTTGCGTTCGGGC |
| 3731 | TP53 | TCCGGGGACACTTTGCGTTC |
| 3732 | TP53 | CCAGGGAGCAGGTAGCTGCT |
| 3733 | TP53 | GCCCGAACGCAAAGTGTCCC |
| 3734 | TP53 | CTCCGGGGACACTTTGCGTT |
| 3735 | TP53 | GGACACTTTGCGTTCGGGCT |
| 3736 | TP53 | CAGGTAGCTGCTGGGCTCCG |
| 3737 | TSIX-001 | AAAAATGTAAAAGATCAGCT |
| 3738 | TSIX-001 | GAGAATCACTTGAACCCAGG |
| 3739 | TSIX-001 | GCGCCAGTGCACTCCATCCT |
| 3740 | TSIX-001 | TTGCCCAGGATGGAGTGCAC |

| | | |
|---|---|---|
| 3741 | TSIX-001 | CAGGAGAATCACTTGAACCC |
| 3742 | TSIX-001 | TGTAATCCCAGCTACTCAGC |
| 3743 | TSIX-001 | GAGACAGGGTTTCACCATGT |
| 3744 | XIST-001 | GCGCTTTAAGAACTGAAGGG |
| 3745 | XIST-001 | AGCGCTTTAAGAACTGAAGG |
| 3746 | XIST-001 | GTGCAGAGAGATCTTCAGTC |
| 3747 | XIST-001 | CCATATTTCTTACTCTCTCG |
| 3748 | XIST-001 | CAGCGCTTTAAGAACTGAAG |
| 3749 | XIST-001 | TGCAGCGCTTTAAGAACTGA |
| 3750 | XIST-001 | CTGAAGATCTCTCTGCACTT |

**Supplementary Table 5. MYC Interactors *in vivo* (n=172)**

| | p value |
|---|---|
| AARS2 | 0.000 |
| EDC4 | 0.000 |
| KDM3B | 0.000 |
| DMAP1 | 0.000 |
| IPO4 | 0.000 |
| RUVBL2 | 0.000 |
| NCOR1 | 0.000 |
| WDR82 | 0.001 |
| SAE1 | 0.001 |
| ATP1A1 | 0.001 |
| FIP1L1 | 0.001 |
| TUBB2B | 0.002 |
| PDCD6 | 0.002 |
| GTF2E2 | 0.002 |
| RPL14 | 0.002 |
| SAP30BP | 0.003 |
| SRPRB | 0.003 |
| SAFB | 0.003 |
| RSU1 | 0.003 |
| DHFR | 0.003 |
| SMUG1 | 0.004 |
| TUBB4A | 0.004 |
| RUVBL1 | 0.005 |
| MAP7D1 | 0.005 |
| LSM2 | 0.005 |
| SF1 | 0.005 |
| DKFZp686L22104 | 0.006 |
| NOB1 | 0.006 |
| PRRC2B | 0.006 |
| ECD | 0.006 |
| RPRD2 | 0.006 |
| ATXN10 | 0.006 |
| TOR1AIP1 | 0.006 |
| TAF6L | 0.007 |
| TK1 | 0.007 |
| PTRH2 | 0.007 |
| SCML2 | 0.007 |

| | |
|---|---|
| TUBA1C | 0.007 |
| PSMD11 | 0.008 |
| TRIM4 | 0.008 |
| SF3A1 | 0.008 |
| EP400 | 0.008 |
| NUBP2 | 0.008 |
| GIGYF2 | 0.008 |
| ARID1A | 0.008 |
| DVL2 | 0.009 |
| GTF2F2 | 0.010 |
| TTC37 | 0.010 |
| STAU1 | 0.010 |
| PIGU | 0.010 |
| CLPTM1 | 0.011 |
| EIF4E | 0.011 |
| TRRAP | 0.011 |
| XRCC6 | 0.011 |
| SCAF4 | 0.011 |
| CRTC3 | 0.011 |
| ITPK1 | 0.012 |
| I02 | 0.012 |
| TOR1AIP1 | 0.012 |
| HCFC1 | 0.012 |
| AP3B1 | 0.012 |
| RPRD1B | 0.012 |
| CPSF4 | 0.012 |
| ARF5 | 0.013 |
| CHD1 | 0.014 |
| FAM96B | 0.014 |
| U2SURP | 0.014 |
| GEMIN5 | 0.014 |
| TAF5L | 0.015 |
| TPR | 0.015 |
| ERLIN2 | 0.015 |
| STAT3 | 0.016 |
| ARID3B | 0.016 |
| SAR1B | 0.016 |
| MAD2L1 | 0.017 |
| NCOR2 | 0.017 |
| PBRM1 | 0.017 |
| DIDO1 | 0.017 |

| | |
|---|---|
| IRF2BP2 | 0.017 |
| RACGAP1 | 0.017 |
| ACTL6A | 0.017 |
| EEF1A2 | 0.018 |
| CIZ1 | 0.018 |
| EIF2B2 | 0.018 |
| Z0318 | 0.018 |
| CBWD1 | 0.019 |
| ITPA | 0.019 |
| CSNK2A2 | 0.019 |
| TUBB | 0.020 |
| IQGAP2 | 0.020 |
| TRNT1 | 0.020 |
| NELFA | 0.022 |
| GAPVD1 | 0.022 |
| EIF2S2 | 0.022 |
| RTKN | 0.023 |
| LRRC47 | 0.023 |
| XPO5 | 0.023 |
| PSMB6 | 0.023 |
| TUBB4B | 0.024 |
| CTPS2 | 0.024 |
| EIF5 | 0.024 |
| JUN | 0.024 |
| MGA | 0.025 |
| COG8 | 0.025 |
| HAT1 | 0.026 |
| RPL37A | 0.026 |
| RING1 | 0.026 |
| NUFIP2 | 0.027 |
| DYNC1LI1 | 0.027 |
| TCAF1 | 0.027 |
| WDR5 | 0.028 |
| HAUS8 | 0.028 |
| USP33 | 0.028 |
| GPATCH8 | 0.028 |
| PCGF6 | 0.028 |
| ASB6 | 0.028 |
| SNW1 | 0.029 |
| CDK12 | 0.030 |
| WDR6 | 0.031 |

| | |
|---|---|
| CANX | 0.031 |
| ANKFY1 | 0.032 |
| TOX4 | 0.032 |
| GIPC1 | 0.033 |
| SMAD5 | 0.033 |
| CDCA2 | 0.033 |
| PSME3 | 0.034 |
| SALL2 | 0.034 |
| ATP2B4 | 0.035 |
| MAGED2 | 0.035 |
| USP10 | 0.035 |
| CHD8 | 0.035 |
| ALAS1 | 0.035 |
| API5 | 0.035 |
| R0214 | 0.035 |
| EMD | 0.036 |
| CBLL1 | 0.036 |
| BIN3 | 0.036 |
| MAP4 | 0.036 |
| SETDB1 | 0.036 |
| HBA1 | 0.036 |
| IPO9 | 0.037 |
| SEC22B | 0.037 |
| OSBP | 0.037 |
| MDC1 | 0.037 |
| CEP85 | 0.038 |
| SAR1A | 0.038 |
| TAF6 | 0.038 |
| PPM1G | 0.038 |
| ZBTB33 | 0.039 |
| HSPA14 | 0.039 |
| RIF1 | 0.039 |
| SRP72 | 0.039 |
| SNRNP40 | 0.040 |
| HIST1H1C | 0.041 |
| LIMA1 | 0.041 |
| AP3M1 | 0.041 |
| TTLL12 | 0.042 |
| EMC4 | 0.043 |
| CHORDC1 | 0.044 |
| IPO8 | 0.044 |

| | |
|---|---|
| GATAD2A | 0.045 |
| ZC3H14 | 0.045 |
| AP2B1 | 0.047 |
| C6orf120 | 0.047 |
| FANCI | 0.047 |
| POLR2B | 0.047 |
| KTI12 | 0.047 |
| GON4L | 0.048 |
| EIF3H | 0.048 |
| NUP62 | 0.049 |
| VBP1 | 0.049 |
| RPL30 | 0.049 |

**Supplementary Table 6. RROL-dependent in vivo MYC interactors from BioID assay (n=18)**

| | **p** |
|---|---|
| AARS2 | 0.000 |
| KDM3B | 0.000 |
| PTRH2 | 0.001 |
| SNW1 | 0.001 |
| PRRC2B | 0.001 |
| EDC4 | 0.002 |
| GIGYF2 | 0.003 |
| AP3B1 | 0.004 |
| DMAP1 | 0.005 |
| WDR82 | 0.006 |
| EIF4E | 0.006 |
| RPRD2 | 0.006 |
| CRTC3 | 0.006 |
| EEF1A2 | 0.007 |
| SAE1 | 0.007 |
| RUVBL2 | 0.007 |
| FAM96B | 0.008 |
| ARID3B | 0.009 |

**Supplementary Table 7. MYC interactors in MM cells (>2 cell lines; n=176)**

| Names | total | elements |
|---|---|---|
| AMO1 H929 U266$^{MYC+}$ | 7 | MAX |
| | | CDSN |
| | | NELFCD |
| | | WDR82 |
| | | CAPNS1 |
| | | RPL34 |
| | | RNASEH2A |
| AMO1 H929 | 137 | RPS17 |
| | | NCBP2-AS2 |
| | | HSPB1 |
| | | PMPCA |
| | | AIFM1 |
| | | MRPL4 |
| | | NAXE |
| | | COX6C |
| | | LONP1 |
| | | PTCD3 |
| | | FAHD1 |
| | | PRDX6 |
| | | TUBAL3 |
| | | GRPEL1 |
| | | PDIA3 |
| | | MRPL19 |
| | | PCMT1 |
| | | SRI |
| | | MYO1C |
| | | RPLP1 |
| | | GMPPA |
| | | SF3B6 |
| | | GDI1 |
| | | ARHGEF2 |
| | | EFTUD2 |
| | | COX7A2 |
| | | ACTR1A |
| | | ISOC2 |
| | | CBR4 |
| | | HSPA8 |

| | | |
|---|---|---|
| | | GET4 |
| | | DHX15 |
| | | ARPC3 |
| | | RPS8 |
| | | OSGEP |
| | | RSU1 |
| | | SSR3 |
| | | PPP3CC |
| | | IFI16 |
| | | OXCT1 |
| | | LSM3 |
| | | MOGS |
| | | SNRPE |
| | | NDUFA13 |
| | | FAU |
| | | PAFAH1B3 |
| | | QPCTL |
| | | SELENOF |
| | | HIST1H2BB |
| | | PCNA |
| | | 43710 |
| | | EIF2B2 |
| | | NAP1L4 |
| | | ATP6V1A |
| | | PSIP1 |
| | | MYC |
| | | RFC4 |
| | | GNB2 |
| | | RPS24 |
| | | VCP |
| | | PSMA5 |
| | | ACTN2 |
| | | PSMC1 |
| | | STMN1 |
| | | Ighg1 |
| | | GAR1 |
| | | NDUFB3 |
| | | CNOT9 |
| | | PARP1 |
| | | FMNL1 |
| | | BAG6 |

| | | |
|---|---|---|
| | | PSMA2 |
| | | PSMA4 |
| | | BRI3BP |
| | | WDR61 |
| | | PPIH |
| | | TTLL12 |
| | | LSP1 |
| | | NSF |
| | | NAMPT |
| | | BLMH |
| | | PRSS2 |
| | | RPS10 |
| | | IGKC |
| | | COPE |
| | | UQCC2 |
| | | EIF4A1 |
| | | RPL15 |
| | | MRPL39 |
| | | DDX3Y |
| | | HSPA14 |
| | | PABPC4 |
| | | ARFIP1 |
| | | DKFZp686J1372 |
| | | HNRNPAB |
| | | AIMP1 |
| | | SYNCRIP |
| | | MRPS23 |
| | | SNRPGP15 |
| | | MTAP |
| | | MSI2 |
| | | TMEM109 |
| | | PAFAH1B2 |
| | | SORD |
| | | MCM4 |
| | | RPL9 |
| | | PDHX |
| | | GNG7 |
| | | RPA3 |
| | | ISOC1 |
| | | SEC11A |
| | | CAT |

| | | |
|---|---|---|
| | | CANX |
| | | SRM |
| | | NDUFA9 |
| | | ARF3 |
| | | NUDT5 |
| | | PSMB3 |
| | | HDGF |
| | | SDF2L1 |
| | | MPRIP |
| | | TRMT10C |
| | | PSME3 |
| | | NDUFS3 |
| | | ECI1 |
| | | PAM16 |
| | | USP39 |
| | | RPL38 |
| | | RPL14 |
| | | IDH3B |
| | | EIF3G |
| | | HNRNPA1L2 |
| | | CCT6A |
| | | FKBP2 |
| | | NDUFA5 |
| | | NCLN |
| | | DNAJC19 |
| AMO1 U266$^{MYC+}$ | 9 | LAMTOR3 |
| | | ARPC2 |
| | | FAM207A |
| | | ASL |
| | | SDC1 |
| | | BAG2 |
| | | FLOT1 |
| | | MIF |
| | | MAGED2 |
| H929 U266$^{MYC+}$ | 23 | MRPS35 |
| | | THRAP3 |
| | | YARS2 |
| | | PPIL1 |
| | | C19orf25 |
| | | FAM98B |
| | | ALDH1L2 |

| | | |
|---|---|---|
| | | SEC13 |
| | | PSMC4 |
| | | FLNB |
| | | DKC1 |
| | | AUH |
| | | TPM4 |
| | | GMPPB |
| | | ACTR1B |
| | | RAD50 |
| | | PSMD4 |
| | | HLA-B |
| | | HNRNPA3 |
| | | CDKN2C |
| | | POLR2E |
| | | DDX41 |
| | | TOP1 |

**Supplementary Table 8. RROL-dependent MYC interactors in MM cells (>2 cell lines; n=8)**

| | | |
|---|---|---|
| AMO1 H929 U266$^{MYC+}$ | 1 | WDR82 |
| AMO1 H929 | 5 | RNASEH2A |
| | | COX6C |
| | | ARFIP1 |
| | | IDH3B |
| | | RPA3 |
| H929 U266$^{MYC+}$ | 2 | C19orf25 |
| | | RAD50 |

**Supplementary table 9. t-ASO sequences.**

**Step 1. 16 oligo sequences either in G (2'MOE/DNA gapmeRs, with a 10-mer DNA gap) or SB (fully 2'MOE) configuration.**

Sequence 1:

CCCGGCCGCCTCCCCCCGTCCC

Sequence 2:>>>**selected as G molecule (G2) (20mer)**

CCCTCCCACCCTCCCCCGCC

Sequence 3:

CCCCCCCGACCCCCTGTGTTT

Sequence 4:

TCCTCCAACCTTTTATCGTTT

Sequence 5:

TCGTCCTTTTTTCTCTTGT

Sequence 6:

CTCCCCTTTTTTACCCAGTT

Sequence 7:

CCCTCTCCGGTCGGTAACCTT

Sequence 8:

TCGTCCTTTTTTCTCTTGTAGT

Sequence 9:>>>**selected as SB molecule (SB9) (22mer)**

CCACCCCTAACACTGGTCTTCT

Sequence 10:

CCCCTCCGCCCTCGTCCTC

Sequence 11:

TCTCTCATTTTCATCCAC

Sequence 12:

AGAGTAAAACAAAACAAAA

Sequence 13:

ACAAAATAGAAAAAAAAGGA

Sequence 14:

ATAAAAAGGGATAAGGTCAG

Sequence 15:

TCGTCCTTTTTTCTCTTGTA

Sequence 16:

CCCTATGGACTTATTATCT

**Step 2. Fine tuning of G2 and SB9 as follows.**

**Fine-tuning of Sequence G2 (2'MOE/DNA gapmeRs, with a 10-mer DNA gap):**

**20mers**

1. gCCCTCCCACCCTCCCCCGC
2. cgCCCTCCCACCCTCCCCCG
3. ccgCCCTCCCACCCTCCCCC
4. cccgCCCTCCCACCCTCCCC
5. CCTCCCACCCTCCCCCGCCg
6. CTCCCACCCTCCCCCGCCgc
7. TCCCACCCTCCCCCGCCgca
8. CCCACCCTCCCCCGCCgcac

**18mers**

9. CCCTCCCACCCTCCCCCG
10. CCTCCCACCCTCCCCCGC
11. CTCCCACCCTCCCCCGCC
12. gCCCTCCCACCCTCCCCC

13. cgCCCTCCCACCCTCCCC
14. TCCCACCCTCCCCCGCCg
15. CCCACCCTCCCCCGCCgc >>>selected as G molecule (G2-15)

**Fine-tuning of Sequence SB9 (fully 2'MOE):**

**22mers**

1. aCCACCCCTAACACTGGTCTTC
2. aaCCACCCCTAACACTGGTCTT
3. aaaCCACCCCTAACACTGGTCT
4. caaaCCACCCCTAACACTGGTC
5. CACCCCTAACACTGGTCTTCa
6. ACCCCTAACACTGGTCTTCaa
7. CCCTAACACTGGTCTTCaaa
8. CCTAACACTGGTCTTCaaaa

**20mers**

9. CCACCCCTAACACTGGTCTT
10. CACCCCTAACACTGGTCTTC
11. ACCCCTAACACTGGTCTTCT
12. aCCACCCCTAACACTGGTCT
13. aaCCACCCCTAACACTGGTC
14. CCCCTAACACTGGTCTTCa
15. CCCTAACACTGGTCTTCaa

**18mers**

16. ACCCCTAACACTGGTCTT
17. CACCCCTAACACTGGTCT
18. CCACCCCTAACACTGGTC
19. CCCCTAACACTGGTCTTC >>> selected as SB molecule (SB9-19)

20. CCCTAACACTGGTCTTCT

### Step 3. Lipid conjugation.

G2-15 and SB9-19 have been conjugated with either:

- Palmitic acid (PA)
- Tocopherol (TO)
- Cholesterol (CH)

Lipid were connected to oligos at 5’ end, with a PS (*) or a PO () linkage.

**Step 3b.** The design G2-15-TO has been further tested with a 8-mer DNA gap (G2-15b-TO), which showed increased on target activity.
